# Development and validation of a potent and specific inhibitor for the CLC-2 chloride channel

**DOI:** 10.1101/2020.01.07.897785

**Authors:** Anna K. Koster, Austin L. Reese, Yuri Kuryshev, Xianlan Wen, Keri A. McKiernan, Erin E. Gray, Caiyun Wu, John R. Huguenard, Merritt Maduke, J. Du Bois

## Abstract

CLC-2 is a voltage-gated chloride channel that is widely expressed in many mammalian tissues. In the central nervous system (CNS), CLC-2 is expressed in neurons and glia. Studies to define how this channel contributes to normal and pathophysiological function in the CNS have been controversial, in part due to the absence of precise pharmacological tools for modulating CLC-2 activity. Herein, we describe the development and optimization of AK-42, a specific small-molecule inhibitor of CLC-2 with nanomolar potency (IC_50_ = 17 ± 1 nM). AK-42 displays unprecedented selectivity (>1000-fold) over CLC-1, the closest CLC-2 homolog, and exhibits no off-target engagement against a panel of 58 common channels, receptors, and transporters expressed in brain tissue. Computational docking, validated by mutagenesis and kinetic studies, indicates that AK-42 binds to an extracellular vestibule above the channel pore. In electrophysiological recordings of mouse CA1 hippocampal pyramidal neurons, AK-42 acutely and reversibly inhibits CLC-2 currents; no effect on current is observed on brain slices taken from CLC-2 knockout mice. These results establish AK-42 as a powerful new tool for investigating CLC-2 neurophysiology.

**Significance Statement:** The CLC-2 ion channel facilitates selective passage of Cl^−^ ions across cell membranes. In the central nervous system (CNS), CLC-2 is expressed in both neurons and glia and is proposed to regulate electrical excitability and ion homeostasis. CLC-2 has been implicated in various CNS disorders, including certain types of epilepsy and leukodystrophy. Establishing a causative role for CLC-2 in neuropathologies, however, has been limited by the absence of selective reagents that enable acute and specific channel modulation. Our studies have resulted in the identification of a highly potent, small-molecule inhibitor that enables specific block of CLC-2 Cl^−^ currents in hippocampal brain slices. This precise molecular tool should enable future efforts to identify and treat CLC-2-related disease.

## Introduction

The CLC-2 ion channel is one of nine mammalian CLC homologs, which comprise a family of Cl^−^-selective channels and transporters (1). CLC-2 is expressed in nearly every organ, including brain, heart, intestines, and lungs (2), and is critical for electrogenesis, homeostatic control of cell volume, and maintenance of ion gradients. Malfunction and/or dysregulation of CLC-2 underlie disparate human pathologies (1, 3-9). In epithelia, CLC-2 has been implicated as a clinical target for the treatment of chronic constipation and irritable bowel syndrome (10-12). In renal tissue, CLC-2 mutations are linked to primary aldosteronism, the most common cause of secondary arterial hypertension in humans (13-16). In the central nervous system (CNS), CLC-2 mutations are found in various rare forms of leukodystrophy, characterized by leukoencephalopathy, male infertility, and secondary paroxysmal kinesigenic dyskinesia (an episodic, involuntary movement disorder) (17-23). A phenotype reminiscent of the human disease is observed in CLC-2 genetic knockout mice, which exhibit severe retinal degeneration/blindness, infertility, and vacuolization of myelin in brain tissue (24-26). Additionally, aberrant CLC-2 activity influences neuronal excitability and may be involved in idiopathic generalized epilepsies (27-38). Although a direct causal link between CLC-2 and epilepsy remains controversial (39, 40), *in vitro* electrophysiology recordings of neurons from wild-type and *Clcn2*^*–/–*^ knockout animals strongly suggest a role for CLC-2 in modulating electrical excitability in inhibitory signaling pathways (41-47). These findings motivate subsequent research efforts to gain a deeper understanding of CLC-2 function in the CNS.

Unraveling the details of CLC-2 physiology is challenged, in part, by the absence of selective chemical reagents for modulating channel activity. While mouse knockout models have provided important insights into CLC-2 function, compensatory changes in gene expression and developmental abnormalities that accompany from-birth genetic deletion of CLC-2 complicate phenotypic analysis. A potent, selective chemical reagent targeting CLC-2 could inhibit channel function rapidly and reversibly, a decided advantage over genetic (irreversible) methods for investigating channel physiology. Available Cl^−^-channel inhibitors (48), however, suffer from low potency, with mid-µM to mM concentrations required to inhibit Cl^−^ current, and lack specificity against different Cl^−^ channel subtypes (1, 49-51). As such, the usefulness of these tool compounds for cellular and organismal studies is severely limited. For CLCs, the most potent and selective small-molecule inhibitors (IC_50_ ∼ 0.5 µM) are those targeting CLC-Ka, one of two CLC homologs expressed in the kidney (52-55). For CLC-2, a peptide toxin, GaTx2, has been identified as a selective inhibitor of CLC-2 at concentrations of ∼20 pM (56); however, reduction in peak current saturates at ∼50% with GaTx2 application, thus restricting use of this toxin as a pharmacological probe. All other reported CLC-2 inhibitors are not selective and require application of high concentrations (∼1 mM) (1, 2, 57-60) to inhibit channel function.

The dearth of high-affinity, high-precision tool compounds for investigating CLC-2 physiology inspires the studies described herein. Through a systematic structure-activity analysis, we have developed the most potent small-molecule inhibitor known to-date for any member of the CLC protein family. This compound, AK-42, has an IC_50_ of 17 ± 1 nM against human CLC-2 and displays ∼10,000×greater potency towards CLC-2 compared to the most closely related CLC homolog, CLC-1. In addition, AK-42 is specific for CLC-2 over a diverse panel of 58 CNS receptors, channels, and transporters. We have validated the suitability of AK-42 for physiological studies of the CNS by performing brain slice recordings on both wild-type and CLC-2 knockout mice. Computational and mutagenesis studies identify an AK-42 binding site on the extracellular side of the channel pore, laying the groundwork for understanding the mechanistic underpinnings of channel inhibition and for further development of related tools. The availability of AK-42 as a potent, highly selective reagent for pharmacological knockout of CLC-2 should greatly accelerate physiological studies of this channel.

## Results

### Screening for new CLC-2 inhibitors

Using the IonWorks™ Barracuda (IWB) automated patch-clamp platform, we performed a focused screen of FDA-approved drug molecules against human CLC-2 stably expressed in Chinese hamster ovary (CHO) cells and identified meclofenamate (MCFA) as a ‘hit’ compound (**Figure 1, Figure S1, Table S1, Dataset 1**). MCFA was selected over other ‘hits’ owing to the ease of synthesis and modification of the diarylamine core. Examination of other non-steroidal anti-inflammatory drugs (NSAID) within our initial screen (**Table S1**) revealed that the interaction of MCFA with CLC-2 is complex and does not simply derive from non-specific Coulombic attraction of the anionic carboxylate group to the electropositive ion conduction pore. Most notably, mefenamic acid and diclofenac, which are structurally analogous to MCFA, have negligible inhibitory activity at 30 µM (**Table S1**). A small collection of angiotensin (AT1) receptor antagonists called sartans, which are anionic at physiological pH (due to the presence of a tetrazole, carboxylic acid, or both), also failed to inhibit Cl^−^ currents to any appreciable extent at 30 µM (**Table S1**). Other known small-molecule modulators of CLC homologs (e.g., BIM1, BIM4, DPC, and NFA, **Figure S1**) displayed no efficacy against CLC-2 (**Table S1**). Lubiprostone, a reported CLC-2 activator used to treat chronic constipation (10, 61-67), also had no effect on CLC-2 currents at concentrations up to 120 μM, in agreement with other literature (68-70) (**Table S1**).

**Figure 1.**
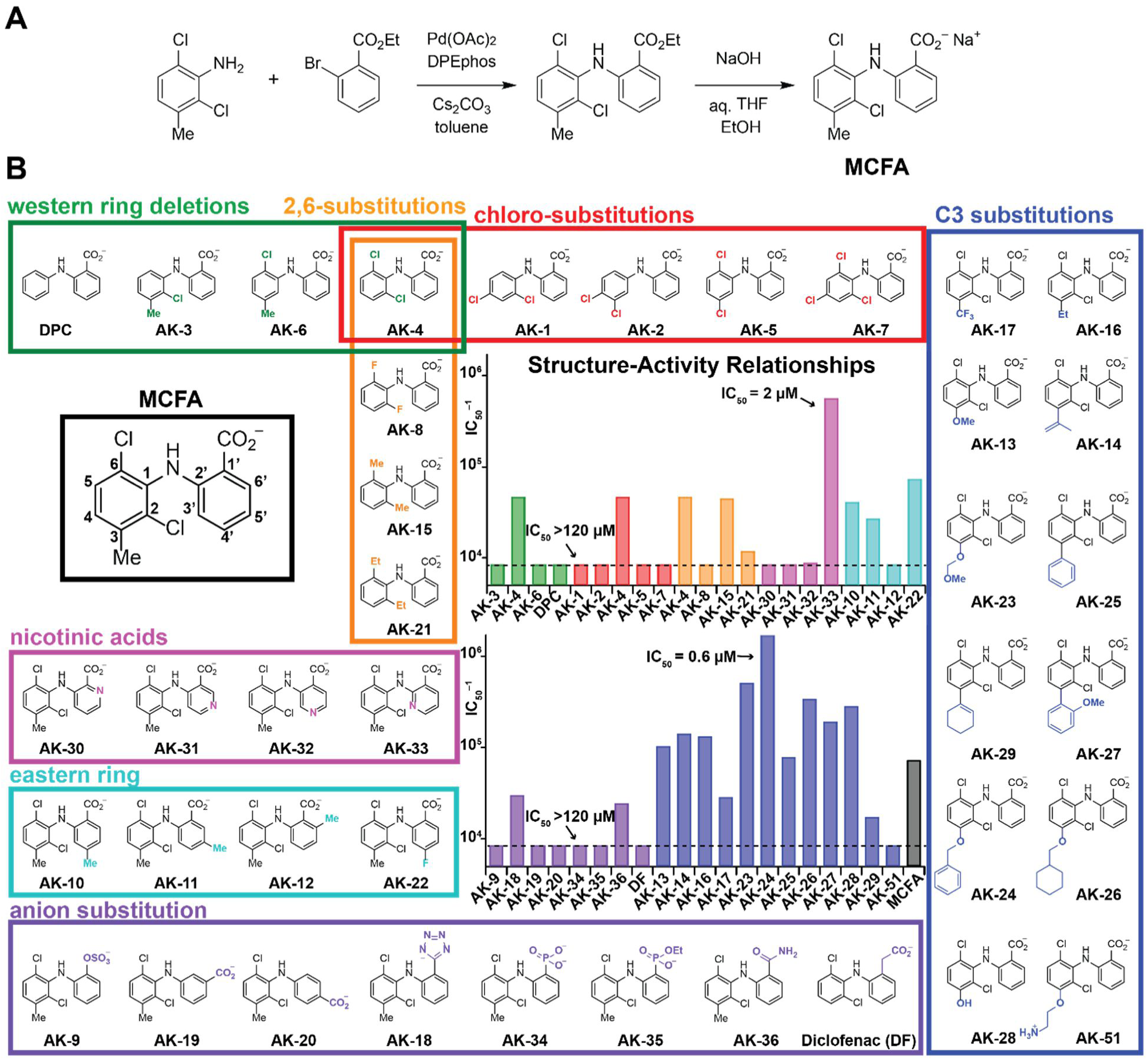
Structure-activity relationship studies to develop a selective CLC-2 inhibitor based on the MCFA ‘hit’ compound. (**A**) General synthetic route for the preparation of MCFA and analogs. (**B**) SAR of MCFA derivatives. The structure of MCFA is shown left (black box) with carbon numbers of the eastern and western rings labeled. The bar graphs show the inverse IC_50_ (IC_50_^−1^) for each compound against CLC-2, estimated from inhibition at four concentrations (n = 3–4 for each concentration). Compounds are grouped by substitution position, as indicated by bar colors that match the box outlines around each group of structures. Compounds that inhibited weakly at the highest concentration tested (usually 120 µM) are indicated with the dotted line (IC_50_ > 120 µM). Results are tabulated in **Table 1**.

### Structure-Activity Relationship (SAR) studies

To enable SAR studies of MCFA, a general synthesis of substituted diarylamines was developed (**Figure 1A**). Using this chemistry, a total of 51 MCFA derivatives were prepared. The results from evaluation of the first 37 derivatives tested (**Figure 1B)** guided the design of an additional 14 compounds (**Figure 2**).

**TABLE 1.** *Top Table:* Approximate IC_50_ values of fenamate derivatives against CLC-2, determined from 4 concentrations using the IWB platform. Compounds are listed according to position of modification, as shown in **Figure 1B**, and groups are color-coded accordingly. *Bottom Table:* IC_50_ data summary for derivatives of AK-24, determined from 4 concentrations using the IWB platform. For both tables, if the IC_50_ was greater than the highest concentration tested, this concentration is listed along with the corresponding % inhibition (in parenthesis).

| Compound | IC<sub>50</sub> (μM) | Compound | IC<sub>50</sub> (μM) |
| --- | --- | --- | --- |
| **Western Ring Deletions** |  | **2,6-Substitutions** |  |
| AK-3 | >120 (43%) | AK-4 | 22 |
| AK-4 | 22 | AK-8 | >120 (41%) |
| AK-6 | >120 (48%) | AK-15 | 23 |
| DPC | >312 (27%) | AK-21 | 86 |
| **Chloro- Substitutions** |  | **Nicotinic Acids** |  |
| AK-1 | >120 (13%) | AK-30 | >222 (12%) |
| AK-2 | >120 (12%) | AK-31 | >42 (16%) |
| AK-4 | 22 | AK-32 | 115 |
| AK-5 | >120 (23%) | AK-33 | 2 |
| AK-7 | >120 (22%) | **Eastern Ring** |  |
| **C3-Substitutions** |  | AK-10 | 25 |
| AK-13 | 10 | AK-11 | 38 |
| AK-14 | 7 | AK-12 | >102 (37%) |
| AK-16 | 8 | AK-22 | 14 |
| AK-17 | 36 | **Anion Substitutions** |  |
| AK-23 | 2 | AK-9 | >120 (28%) |
| AK-24 | 0.6 | AK-18 | 33 |
| AK-25 | 13 | AK-19 | >141 (36%) |
| AK-26 | 3 | AK-20 | >144 (8%) |
| AK-27 | 5 | AK-34 | >195 (6%) |
| AK-28 | 4 | AK-35 | >354 (41%) |
| AK-29 | 49 | AK-36 | 42 |
| AK-51 | >120 (7%) | DF | >363 (5%) |
| MCFA | 14 |  |  |
| Compound | IC<sub>50</sub> (μM) | Compound | IC<sub>50</sub> (μM) |
| AK-24 | 0.6 | AK-44 | 68 |
| AK-37 | >120 (15%) | AK-45 | 1 |
| AK-38 | >120 (4%) | AK-46 | 8 |
| AK-39 | >120 (0%) | AK-47 | 1 |
| AK-40 | >120 (5%) | AK-48 | 16 |
| AK-41 | 82 | AK-49 | 76 |
| AK-42 | <0.12 (90%) | AK-50 | 7 |
| AK-43 | 18 |  |  |

**Figure 2.**
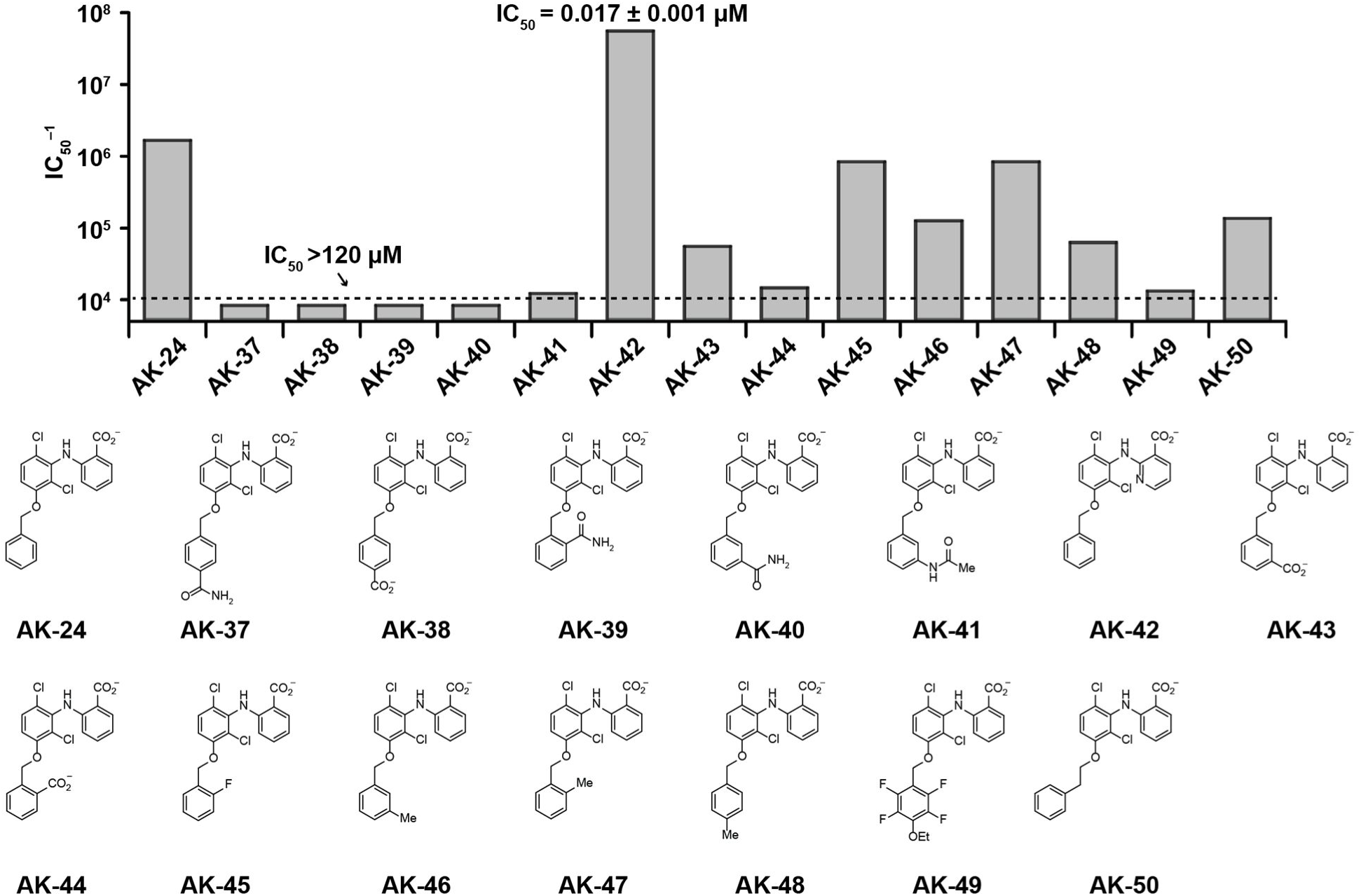
SAR studies around the AK-24 scaffold. The bar graph shows the inverse IC_50_ estimated from inhibition at four concentrations (n = 3–4 each concentration), as in **Figure 1**. Substitutions around the C3-OBn ring are destabilizing. Combining the substitutions of AK-24 and AK-33 has a synergistic effect on inhibitor potency (AK-42). Data are given in **Table 1**.

An initial systematic examination of the western aryl ring of the MCFA scaffold was performed to assess the effects of -Cl and -Me substitution (**Figure 1, Table 1**, green, orange, and red boxes). Removal of either the C2- or C6-Cl group (AK-3, AK-6) resulted in a ∼50% decrease in potency relative to MCFA. By contrast, the potency of AK-4, which lacks the C3-Me group, was relatively unchanged. Other di- and tri-chlorinated analogs, including AK-1, AK-2, AK-5, and AK-7, were ineffective at blocking CLC-2 (IC_50_ ≥ 120 µM). C2,C6-Substitution with Cl- or Me-groups (AK-4 and AK-15, respectively) proved imperative; introduction of sterically smaller (e.g., -F, AK-8) or larger (e.g., -Et, AK-21) substituents significantly reduced inhibitor potency. These results suggest that a twisted geometry of the diarylamine scaffold is necessary for inhibition.

The small reduction in potency of AK-4 versus MCFA (IC_50_ = 22 µM vs. 14 µM) led us to query the effect of replacing the C3-Me group in MCFA with alternative substituents (**Figure 1, Table 1**, blue box). Initially, a series of four compounds was prepared in which a -OMe, isopropenyl, -Et, or -CF_3_, group was introduced at C3 (AK-13, AK-14, AK-16, and AK-17, respectively). With the exception of AK-17, these ligands were nearly 2-fold more potent than MCFA. In a second round of SAR, incorporation of sterically larger groups at C3, which included -OBn and -OCH_2_^c^Hx (AK-24 and AK-26, respectively), resulted in an additional boost in potency. Among these derivatives, AK-24 was identified as the most effective inhibitor with an IC_50_ of 0.6 µM.

We next evaluated the influence of the MCFA carboxylate group on ligand binding (**Figure 1, Table 1**, purple). This study included moving the position of the carboxylate from C1’ to C5’ and C6’ on the eastern ring (AK-20, AK-19) and replacing the carboxylate unit with other charged substituents, including -OSO_3_^−^,-PO_3_^2–^, -PO_2_(OEt)^−^, -tetrazolate, and -CH_2_CO_2_^−^ (AK-9, AK-34, AK-35, AK-18, and diclofenac, respectively). A neutral analog of MCFA, AK-36, in which the carboxylate group was converted to a primary carboxamide, was also prepared. All eight compounds displayed reduced potency relative to MCFA. From these data, the efficacy of carboxamide AK-36 is perhaps most surprising, as this derivative is only ∼3-fold less potent than MCFA despite lacking the anionic charge. This result indicates that ligand-protein binding is not purely dictated by charge-charge interactions.

Additional ligand SAR effects were revealed by varying substituent groups on the eastern ring of MCFA with the western ring held fixed (**Figure 1, Table 1**, magenta and cyan). A Me-scan at positions C4’, C5’, and C6’ (AK-10, AK-11, and AK-12, respectively) showed diminished potency for all three isomers, with the most significant loss resulting from C6’ substitution. A C4’-F derivative (AK-22) also proved less potent than MCFA. Collectively, these findings intimate that the eastern ring of MCFA is sequestered in a restricted binding pocket. Given these data, we prepared four isomeric nicotinic acid derivatives (AK-30, AK-31, AK-32, and AK-33), reasoning that introduction of a N-atom in this region of the inhibitor would alter steric size and potentially enable a stabilizing hydrogen bonding interaction with the protein. While three of these compounds display substantially reduced potency, one derivative, AK-33, is improved 7-fold over MCFA (IC_50_ = 2 µM). Because the four isomers, AK-30, AK-31, AK-32, and AK-33, have similar cLogP values and polar surface areas, this finding suggests that the N-atom at the 3’ position engages in a specific interaction between AK-33 and CLC-2.

In light of our results with AK-24 and AK-33, we synthesized AK-42, a derivative that combines the C3-OBn substituent on the western ring with the C3’-nicotinic acid on the eastern half (**Figure 3A**). AK-42 is almost three orders of magnitude more potent than MCFA (**Figure 3B**). Manual patch-clamp experiments on CHO cells transiently transfected with rat CLC-2 indicate comparable AK-42 potency (IC_50_ = 14 ± 1 nM for rat CLC-2, **Figure S2**) to that determined for human CLC-2 using the IWB (17 ± 1 nM). To our knowledge, AK-42 is the most potent small-molecule inhibitor against any CLC homolog.

**Figure 3.**
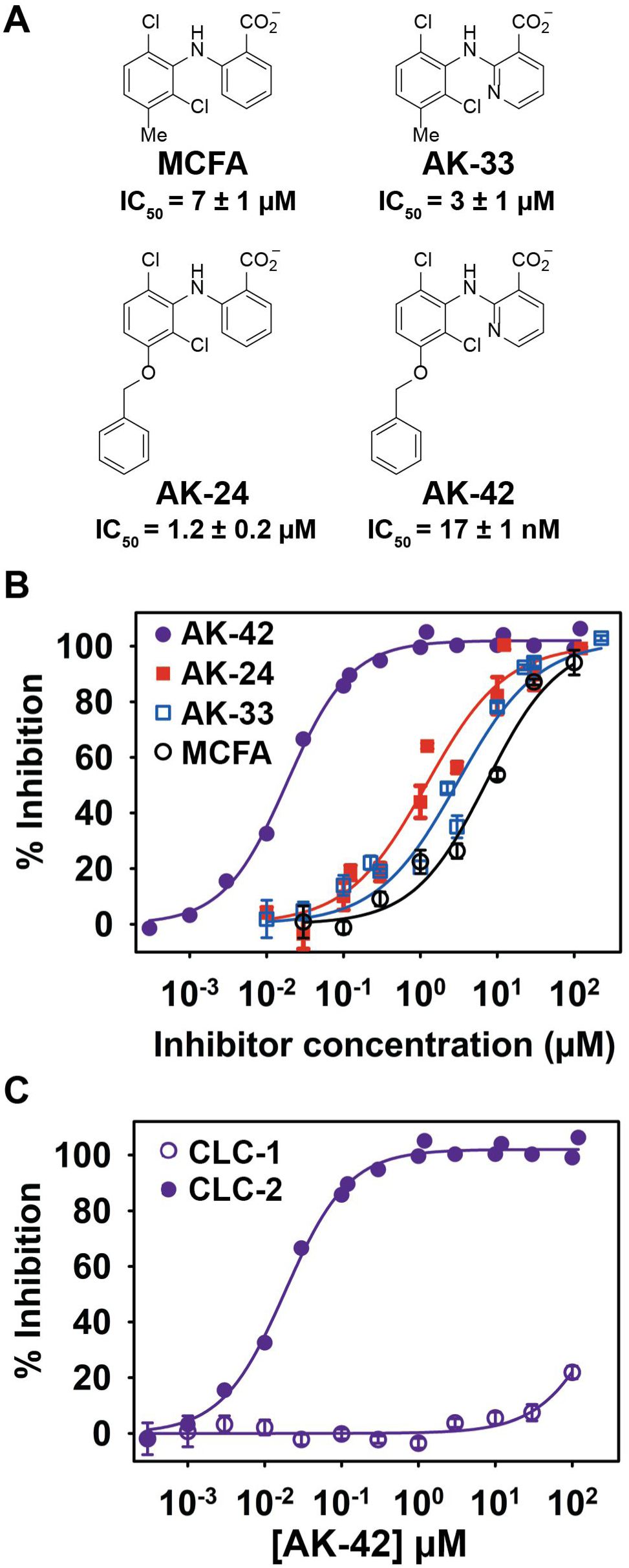
Combining the most potent compounds identified in the SAR studies results in a highly potent and selective CLC-2 inhibitor, AK-42. (**A**) Structures, including the initial ‘hit’ compound MCFA, are shown. IC_50_ values were determined from data in panel (**B**); these IC_50_ values differ slightly from those obtained in the initial screen (**Fig. 1**), which used only four concentrations of each compound. (**B**) Percent inhibition at –120 mV was determined using the IWB, n = 3–4 for each concentration ± SEM. Fits to the Hill equation yield IC_50_ values of 7 ± 1 µM, 1.2 ± 0.2 µM, 3 ± 1 µM, and 0.017 ± 0.001 µM for MCFA, AK-24, AK-33, and AK-42 respectively (**Table S2**). (**C**) Inhibition curves for AK-42 against CLC-1 and CLC-2 determined using the IWB, illustrating that AK-42 is ∼10,000 times more potent against CLC-2 compared to CLC-1, the most closely related CLC homolog (individual data points are shown in **Table S3**).

### AK-42 selectivity, specificity, and binding site

To assess the utility of AK-42 as a tool compound for studies of CLC-2 physiology in the CNS, we evaluated the specificity of this ligand for CLC-2 over other protein targets. Electrophysiological recordings of CLC-1, the most closely related homlog to CLC-2 (∼55% sequence identical), revealed >4 orders of magnitude reduced sensitivity to AK-42 (**Figure 3C**). By comparison, the original ‘hit’ compound, MCFA, exhibits only modest selectivity (<10-fold) between the CLC-1 and CLC-2 (**Table S2**). Additionally, MCFA inhibition of the kidney channel, CLC-Ka, requires mid-µM concentrations (IC_50_ ∼40 µM) (71). Collectively, these results support AK-42 as a uniquely selective agent for targeting CLC-2 over other CLCs.

The value of CLC-2 as a tool compound for studying channel function in the CNS was further assessed by measuring ligand potency against a diverse panel of potential off-target CNS receptors, channels, and transporters. To do so, we employed a comprehensive screen developed and conducted by the National Institutes of Health Psychoactive Drug Screening Program (NIH PDSP) (72). Depending on the identity of the receptor, primary binding assays or functional assays were performed with each target at 10 µM AK-42, a concentration well above the IC_50_ of this compound for CLC-2 and at least 4 times higher than the inhibitor concentrations used in our primary cell experiments (*vide infra*). In these assays, no notable off-target protein binding effects were noted; for the 8 targets exhibiting a response of >50% at 10 µM (**Dataset 2, Table 1**), secondary binding assays confirmed that the effects were weak or negligible (**Dataset 2, Figure 1**).

Computational docking was performed to gain insight into the molecular basis for CLC homolog selectivity and to generate binding site predictions for AK-42. Absent a high-resolution structure of CLC-2, we took advantage of a CLC-2 structural homology model that we developed and refined with 600 µs of molecular dynamics simulations (73). Based on the rapid onset of channel block and the reversibility of inhibition (**Figures S2, S3**), we reasoned that AK-42 is likely binding to an extracellular site on the channel. Docking of AK-42 to the extracellular side of the CLC-2 model preferentially places this ligand near the outer vestibule of the ion permeation pathway, in a similar location to that targeted by anionic inhibitors of CLC-Ka (54, 55). Notably, we found that AK-42 does not dock to the cryo-EM structure of CLC-1 (6COY) (74), in line with our functional studies on selectivity (**Figure 3C**). While the top docking poses in our CLC-2 model vary in the specific interactions predicted, all place AK-42 in the same general binding site. A representative docking pose to the putative CLC-2 binding site, along with a schematic of the 14 residues in the vicinity of AK-42 (compiled from multiple high-scoring docking models), is shown in **Figure 4A, B**. To test the binding site predictions of this model, we first examined mutations of the two charged residues located within this proposed pocket, K210 and K400, reasoning that at least one of these could contribute to electrostatic stabilization of the anionic inhibitor molecule. Mutations at K210, a highly conserved residue, resulted in significantly diminished sensitivity to AK-42 (**Figure 4C, D**). The effectiveness of AK-42 is reduced even against the charge-conserved mutant, K210R. Accordingly, K210 may play a critical role in fixing the inhibitor geometry within the binding pocket, as highlighted in **Figure 4B**. By contrast, no change in inhibitor sensitivity is measured against the K400R mutant as compared to wild-type. These results reveal that K400 does not contribute to inhibitor selectivity for CLC-2 over CLC-1, the latter of which bears an arginine at this position. Finally, introduction of proline at Q399, the residue present at the equivalent position in CLC-1, reduces inhibition by AK-42 (**Figure 4D**). Together, these data support our conclusion that AK-42 binds to the predicted extracellular site on CLC-2.

**Figure 4.**
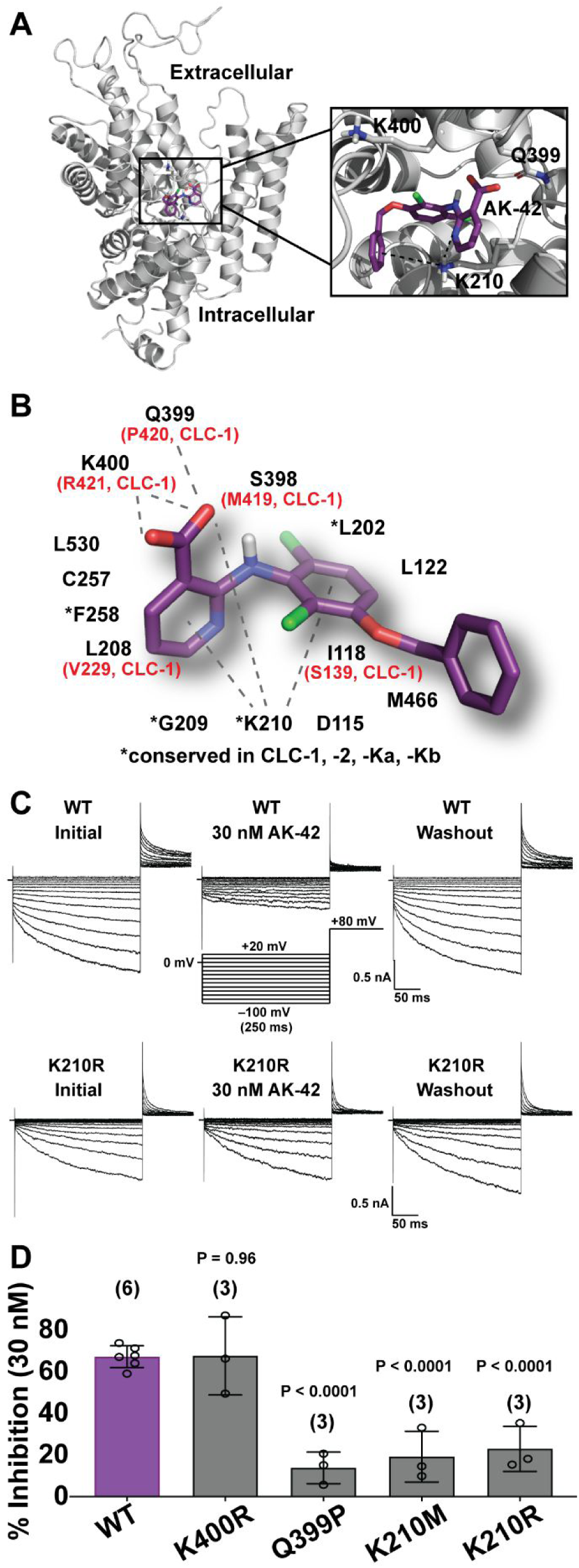
Analysis of AK-42 selectivity and the inhibitory binding site. (**A**) One subunit of the CLC-2 homodimer is shown. AK-42 docks near the extracellular entrance to the chloride permeation pathway, which is tilted towards the viewer. The expanded view highlights three of the residues in the predicted binding pocket. (**B**) Illustration of one of the top poses adopted by AK-42 in molecular docking to CLC-2, together with a schematic of the 14 residues proximal to AK-42 in all docking poses. Residues differing between CLC-1 and CLC-2 are indicated in red. (**C**) Representative CLC-2 current traces from manual patch-clamp experiments. *Top:* WT CLC-2 currents, illustrating reversible CLC-2 inhibition by 30 nM of AK-42, consistent with an extracellular binding site. *Bottom:* K210R CLC-2 currents, illustrating a diminution in AK-42 potency upon mutating K210 to R, supporting the docking model in **B**. (**D**) Testing other residues predicted to interact with AK-42 in the CLC-2 binding site. Mutations of K210 and Q399 produced statistically significant (by unpaired *t*-test) effects on inhibitor binding at 30 nM, whereas the K400R mutation did not. Number of data points per mutant is shown in parenthesis. Summary data are tabulated in **Table S4**. These data support that AK-42 binds in the pocket predicted by docking studies.

### Validation of AK-42 in brain-slice recordings

To validate the effectiveness of AK-42 for neurophysiological studies, we measured CLC-2 currents in CA1 pyramidal cells of acute brain slices using whole-cell patch-clamp electrophysiology. CLC-2 currents were isolated by blocking synaptic transmission, currents mediated by voltage-gated sodium channels, and hyperpolarization-activated cyclic nucleotide–gated (HCN) channels (see **Materials and Methods**). To evoke CLC-2 currents, cells were stepped from –4 mV, a membrane potential at which CLC-2 channels have low open probability, to test potentials ranging from –100 to +30 mV. During the three-second test pulse, CLC-2 current is observed as a slow increase in inward current at the most negative potentials (**Figure 5A**). These data are as expected based on previous studies in hippocampal slice recordings (41, 44, 75-77). Relative to other neuronal ion channels, the CLC-2 channel activates slowly (seconds) following hyperpolarization, thereby facilitating current isolation (75). We measured CLC-2 channel activity in terms of the final steady-state current, as well as the amount of current change over the four-second test step (relaxation current). AK-42 significantly attenuated steady-state currents and eliminated relaxation currents in recorded neurons (**Figure 5A–D**); in contrast, AK-42 had no effect on either measurement in neurons taken from homozygous CLC-2 knockout (*Clcn2*^−/–^) mice, thereby demonstrating target specificity (**Figure 5E–H**). An overlay of recordings from untreated *Clcn*2^−/–^ versus wild-type animals treated with AK-42 shows no discernable difference (**Figure S4, A–B**), confirming that inhibition of CLC-2 current by this compound in wild-type hippocampal CA1 neurons is complete and highly selective.

**Figure 5.**
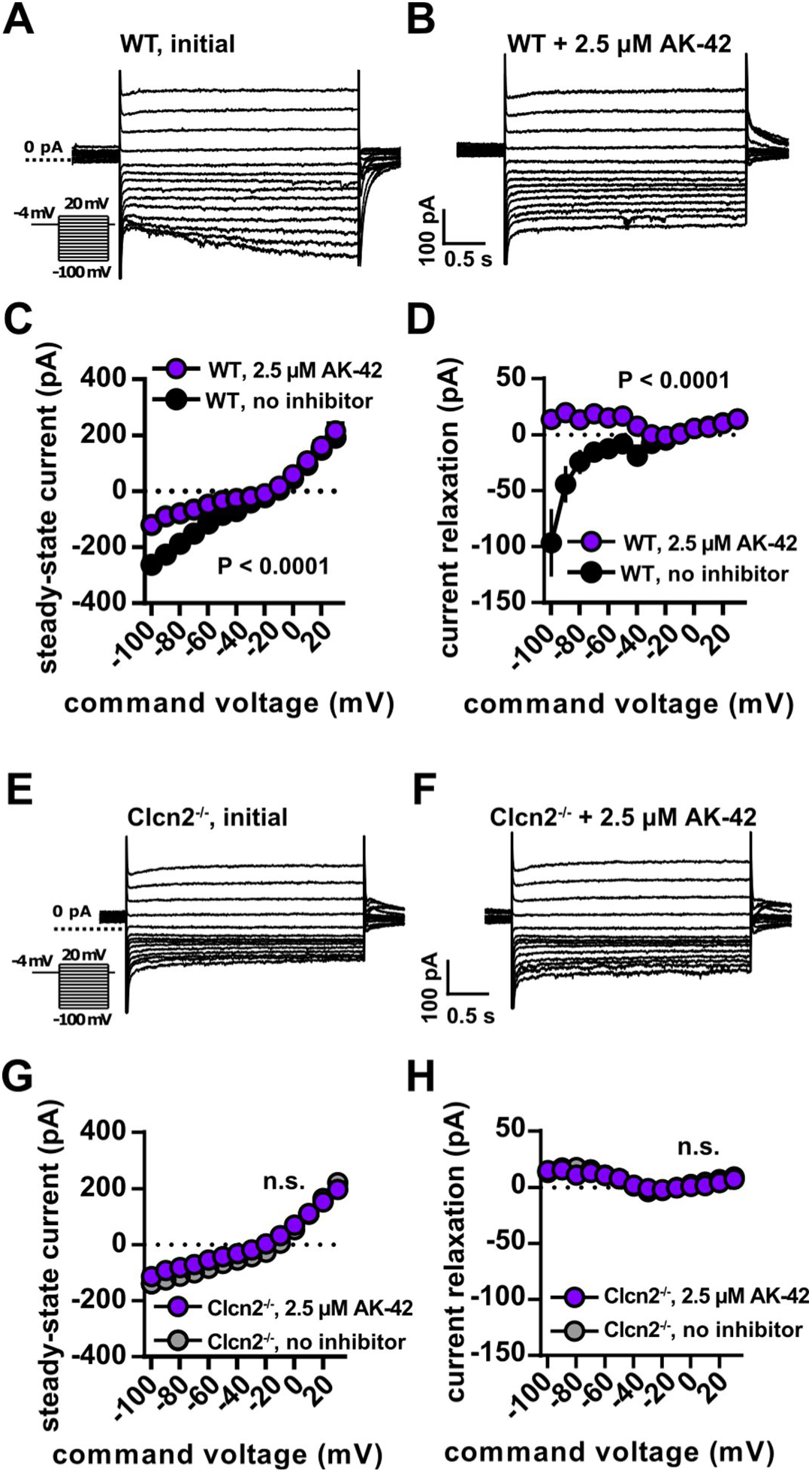
AK-42 inhibits hyperpolarization-activated current in hippocampal cells. (**A–B**) Whole-cell voltage-clamp recordings (protocol in inset) of WT mouse CA1 pyramidal neurons with and without AK-42. Capacitive transients are clipped for display purposes. (**C**) Whole-cell I-V relationship at steady-state both with and without AK-42 (n = 5 cells, 6 slices and 2 animals per group. P < 0.0001 via two-way RM ANOVA). Error bars represent ± SEM throughout. (**D**) Current relaxation measured as the difference between the beginning and end of a 4-s pulse, showing that development of inward current was blocked by AK-42 (n = 6 cells, 6 slices and 2 animals per group. P < 0.0001 via two-way RM ANOVA). (**E–F**) Whole-cell currents from voltage-clamp recordings of *Clcn2*^−/–^ CA1 pyramidal neurons before and after AK-42 treatment. Unlike WT responses, steady-state responses show no differences after AK-42 treatment at negative membrane potentials. Steady-state currents (**G**) and current relaxation (**H**) in *Clcn2*^−/–^ cells show no difference after AK-42 treatment (n = 6 cells, 6 slices and 2 animals per group. P = 0.25 and 0.97 for steady-state and current relaxation measurements, respectively, via two-way RM ANOVA).

In the pilot studies described above, a high concentration of AK-42 (2.5 µM) was used to ensure efficient penetration of the inhibitor into the tissue slice. To confirm potency, we repeated these experiments using a lower, but still saturating, concentration of AK-42 (100 nM). Under these conditions, AK-42 also attenuates relaxation current and peak steady-state current, providing additional validation of our initial results (**Figure S4, C–E**). Current-clamp recordings with 100 nM AK-42 show no differences in firing rate, resting membrane potential, input resistance or membrane time constant, indicating that this compound does not influence neuronal excitability through off-target effects (**Figure S4, F–I**).

Using the same tissue preparation, the time course for inhibition by AK-42 (100 nM) was measured (**Figure S4, C**). Steady-state block of CLC-2 current recorded at –80 mV in hippocampal neurons was achieved within 5–10 minutes of compound application to brain slices. The inhibitor had no effect on currents recorded at –20 mV, a voltage at which little to no CLC-2 current is expected. The 5–10-minute time course for onset of inhibition is slower than the rates derived from our kinetic assays in CHO cells (**Figures S2, S3**), but such differences are expected for a compound that must penetrate tissue to reach the intended target. Following steady-state block, we evaluated the reversibility of CLC-2 inhibition by washing out the compound (perfusion with buffer solution) over several minutes (**Figure S4, C**). The partial recovery of hyperpolarization-induced current (66%) over 25 minutes of washing supports the following conclusions: 1) the AK-42 binding site is extracellular; 2) the compound is not significantly internalized over the course of the experiment; and 3) application of AK-42 can provide acute, selective, and reversible block of CLC-2 in primary neurons.

## Discussion

The numerous outstanding questions surrounding the complex physiology of CLC-2 in the CNS inspired our development of AK-42 as a high-precision molecular tool. Systematic modification of an initial hit compound, MCFA, yielded a small library of 51 derivatives from which the critical molecular features responsible for inhibitor potency were identified. This work culminated in the design and synthesis of AK-42.

The potency and specificity of AK-42 for CLC-2 stands in contrast to all other known Cl^−^ channel small-molecule modulators. Application of this compound to cells expressing CLC-2 enables acute channel inhibition, thus avoiding the interpretive challenges of genetic knockout or gene silencing methods. Electrophysiology slice recordings on hippocampal CA1 neurons provide compelling evidence in support of AK-42 specificity: current vs. voltage plots from data collected on *Clcn2*^−/−^ neurons in the absence and presence of AK-42 mirror those obtained from application of AK-42 to wild-type CA1 neurons. Furthermore, primary binding assays using 10 µM AK-42 (≥100× the concentration required to elicit complete CLC-2 current block in hippocampal slices) show no substantive off-target engagement against a panel of 58 common CNS membrane proteins.

The availability of AK-42 enables subsequent experiments to interrogate CLC-2 structure and channel biophysics. While all CLCs share a homodimeric architecture with two independent Cl^−^ permeation pathways, channel gating properties differ between CLC homologs. The disclosures of two mammalian CLC structures have revealed only subtle structural differences between channels (74, 78), and the detailed biophysical machinery driving functional diversity within the CLC family remains largely enigmatic. Small molecules, such as AK-42, are indispensable tools to advance such investigations.

A desire to understand the basis for AK-42 selectivity and specificity prompted our efforts to identify the inhibitor binding site. Fast and reversible inhibition by AK-42 in electrophysiological recordings of heterologously expressed CLC-2 suggest that AK-42 acts extracellularly. This conclusion is supported by computational docking models of AK-42 to a CLC-2 homology model, as well as by mutagenesis experiments with CLC-2 at positions K210 and Q399 (which result in AK-42 potency reductions at 30 nM). Even conservative mutation of K210 to arginine (K210R) perturbs the ability of AK-42 to block Cl^−^ current.

By contrast, arginine mutation of K400 (K400R) has no influence on AK-42 affinity. From our modeling studies, the putative AK-42 binding pocket occupies an extracellular vestibule, located in the same region as the benzimidazole sulfonate, BIM1, receptor site in CLC-Ka (55). Our results, however, indicate that BIM1 does not inhibit CLC-2 at concentrations exceeding 100 µM, suggesting that subtle differences between channels in this locus can be exploited to target individual CLC homologs. Additional investigations are required to determine exact details of AK-42 binding to CLC-2, as our ligand docking model comprises multiple high-scoring poses within the extracellular pocket. Future studies that leverage available SAR data, along with our ability to generate additional AK-42 derivatives and CLC-2 mutants, should reveal the precise molecular interactions between AK-42 and the channel.

The development of AK-42, together with cell-type specific genetic manipulation of CLC-2, will facilitate future investigations into CLC-2 function in both neurons and glia, including studies to determine the effects of this channel on network excitability, cell volume, and ion homeostasis. The widespread expression of CLC-2 in different cell types found in the CNS (24, 41, 45, 75, 76, 79-81) intimates multiple functional roles for this channel in proper brain function. One hypothesis is that the strong inward rectification of CLC-2 at hyperpolarized potentials maintains low intracellular [Cl^−^] in hippocampal neurons to facilitate GABAergic inhibition (41, 42, 44). Alternatively, it has been argued that CLC-2 regulates neuronal excitability by mediating Cl^−^ *influx* under physiological conditions (43). Other studies have suggested that fast-spiking (PV-BC) basket cells specifically associate with CLC-2-containing GABAergic synapses, where the channel is thought to act as a safety valve for Cl^−^ efflux during intense synchronized network oscillations mediated by PV-BC interneurons (44, 46). In glia, CLC-2 is expressed in oligodendrocytes and astrocytes and is associated with an adhesion molecule known as GlialCAM (79, 81, 82). Glial CLC-2 is postulated to regulate ion and fluid homeostasis, and it may indirectly influence neuronal excitability by facilitating uptake of extracellular K^+^ following intense neuronal firing (20, 24, 83-85). While compelling, these mechanisms have been difficult to parse in the absence of acute and specific methods for modulating CLC-2 function. With the advent of AK-42, we expect that new insights to these questions will be revealed.

## Materials and Methods

### IonWorks™ Barracuda (IWB) electrophysiology recording conditions

Data acquisition and analyses were performed using the IonWorks™ Barracuda systems (Molecular Devices Corporation, Union City, CA, operation software version 2.5.3.359) in the Population Patch Clamp™ (PPC) Mode. Test article (TA) and positive control concentrations were prepared fresh daily by diluting stock solutions into an extracellular buffer solution, and the TA formulations were loaded in a 384-well compound plate. Currents were recorded from groups of 64 cells. HEPES-buffered intracellular solution (in mM: 50 CsCl, 90 CsF, 5 MgCl_2_, 5 EGTA, 10 HEPES, adjusted to pH = 7.4 with CsOH) for whole-cell recordings was loaded into the intracellular compartment of the PPC planar electrode. Extracellular buffer (HB-PS; in mM: 137 NaCl, 4 KCl, 3.8 CaCl_2_, 1 MgCl_2_, 10 HEPES, 10 Glucose, adjusted to pH = 7.3 with NaOH) was loaded into PPC planar electrode plate wells (11 µL per well). The cell suspension was pipetted into the wells of the PPC planar electrode (9 µL per well). After establishment of a whole-cell configuration (the perforated patch; 12 µM escin, 7 min), membrane currents were recorded using a patch-clamp amplifier in the IonWorks™ Barracuda system. The current recordings were performed one time before TA application to the cells (baseline) and one time after application of TA or reference compound. Seal resistance prior to compound application was ≥200 MΩ. Reference compound concentrations were applied to naïve cells (n = 4, where n = replicates/concentration). Each application consisted of a 20 μL addition of a 2 × concentrated test article solution to the total 40-μL final volume of the extracellular well of the PPC electrode. The addition was followed by a single mixing of the PPC well content. Duration of exposure to each compound concentration was 5 minutes.

Prior to initial screening, we validated our assay by using Cd^2+^ and NPPB as positive controls for CLC-2 inhibition. Eight dose-response curves were generated in each experiment on two different days (n = 3–4 per concentration). All Z’ values (a standard metric for evaluating assay quality) exceeded 0.5, indicating acceptable assay variability (86). The Z’ factor for was calculated as: Z’ = 1 – (3 × SD_VC_ + 3 × SD_PC_)/ABS (Mean_VC_ – Mean_PC_), where Mean_VC_ and SD_VC_ are the mean and standard deviation values for a vehicle control, Mean_PC_ and SD_PC_ are the mean and standard deviation values for a positive control (300 µM NPPB or 1 mM Cd^2+^). Initial screening for hit compounds was performed using the ENZO Life Sciences SCREEN-WELL^®^ FDA-approved drug library, v. 2.0 (Product # BML-2843-0100). All compounds in this library are reported in **Dataset 1**. Vehicle controls of 0.3 or 1.0% DMSO (without compound) in recording solution were applied in replicate as additional controls in each experiment.

### Cell line generation

All CLC-2 recordings using the IWB platform were performed with a stably expressing Chinese hamster ovary (CHO) cell line. CHO cells were transfected with human CLC-2 cDNAs (*CLCN2*, GenBank Acc # NM_004366.3). Stable transfectants were selected by expression with the antibiotic-resistance gene (Geneticin/G418) incorporated into the expression plasmid (pcDNA3.1(+)). Selection pressure was maintained by including selection antibiotics in the culture medium. Cells were cultured in Ham’s F-12 media (Life Technologies, Grand Island, NY) supplemented with 10% fetal bovine serum (HyCloneTM Fetal Bovine Serum, U.S.), 100 U/mL penicillin G sodium, 100 µg/mL streptomycin sulfate and 500 µg/mL G418. Before testing, cells in culture dishes were washed twice with Hank’s Balanced Salt Solution (HBSS, Life Technologies, Grand Island, NY), treated with Accutase™ (Innovative Cell Technologies, San Diego, CA) solution for 20 min at room temperature and re-suspended in HBSS (4–6 × 106 cells in 10 mL). Immediately before use, the cells were washed twice in HBSS to remove the Accutase™ and re-suspended in 5 mL of HBSS. CLC-1 recordings were performed using a stably expressing CHO cell line that was generated according to the same methods described for CLC-2.

### IonWorks™ Barracuda (IWB) voltage protocol and data analysis

From a holding potential of –30 mV, CLC-2 currents were elicited by applying 2-second test pulses from –120 to +20 mV in 20-mV increments. Following each test pulse, a 500-ms tail pulse to 0 mV was applied. We required initial currents at the 0-mV tail pulse to be ≥0.2 nA to be included in analysis. All IC_50_ data reported in the manuscript are given according to measurements at the maximum current at –120 mV, the potential at which the maximum number of CLC-2 channels are open. The decrease in current amplitude after compound application (TA) was used to calculate the percent block relative to the positive control, according to the equation: % Inhibition = (1 – I_TA_ / I_Baseline_) x 100%, where I_Baseline_ and I_TA_ are the currents measured before and after addition of compound, respectively. The data were then corrected for run-down according to the equation: %Block’ = 100%– ((%Block–%PC)*(100% / (%VC–%PC))), where %VC and %PC are the mean values of the current inhibition with the vehicle control and the highest concentration of a reference compound (300 µM Cd^2+^). The inhibition curves were fit to the equation: % Inhibition = 100 / [1 + ([IC_50_] / [TA])^n^], where [TA] is the concentration of test article, IC_50_ is the concentration of TA producing half-maximal inhibition, n is the Hill coefficient, and % block is the percentage of current inhibited at each concentration of a TA. Nonlinear least squares fits were solved with the XLfit add-in for Excel (Microsoft, Redmond, WA) (initial screening data) or using the Sigmaplot 13.0 (Systat Software, San Jose, CA) regression analysis (final analyses).

For the kinetics experiments (**Figure S3**), current was elicited with a 2000-ms voltage step to a potential of –120 mV, followed by a 500-ms voltage step to a potential of 0 mV; the holding potential was –30 mV. The test pulses were repeated with a frequency of 0.1 Hz. Six stimulations were applied before application of test articles (1-min baseline) and thirty stimulations were performed after application of test articles (5 min). In the time course for AK-42 application, the channel occupies both the open and closed states; thus, the results reflect a weighted average of AK-42 binding to the different channel conformations. More detailed experiments are necessary to address whether AK-42 inhibition is state-dependent. Findings from our experiments establish AK-42 binding to an extracellular (efficiently reversible) site with a 1:1 AK-42/channel subunit stoichiometry. Current amplitudes were measured as the minimal inward current at test potential –120 mV and the maximal outward current at test potential 0 mV.

The vehicle control data (0.3% DMSO; n = 32) were averaged for each time point. All data were normalized to the time-matching averaged vehicle control. Then data were normalized to the baseline current recorded before application of test articles.

### Cell culture & recording protocols (manual patch-clamp)

CHO K1 cells (ATCC^®^ CCL-61TM) were cultured at 37 °C in F12K media (ATCC, Catalog No. 30-2004) supplemented with 10% fetal bovine serum (GIBCO) and 1% penicillin/streptomycin (GIBCO). At 70–90% confluency, cells were transfected with full-length rat CLC-2 cDNA (449 ng), using Lipofectamine^®^ LTX, opti-MEM^®^, and PLUS™ reagent (Invitrogen). The rat CLC-2 gene was contained in a custom plasmid vector called pFROG (ampicillin resistance), provided by Prof. Michael Pusch (Istituto di Biofisica, CNR, Genova, Italy). Co-transfection with GFP (177 ng, 2.5:1 CLC-2/GFP) was used to estimate transfection efficiency. Cells were transfected 18 hours prior to recording. Borosilicate glass micropipettes (Sutter Instruments, Novato, CA) were pulled and fire-polished to a tip diameter with a resistance of 1.5–3.0 MΩ. For whole-cell patch-clamp recordings, the external solution was composed of 148 mM CsCl, 2 mM CaCl_2_, 10 mM HEPES, and 100 mM D-mannitol (adjusted to pH 7.4 with 2 M aqueous CsOH). The internal solution was composed of 146 mM CsCl, 5 mM EGTA, 10 mM HEPES, 5 mM NaF, and 60 mM D-mannitol (adjusted to pH 7.4 with 2 M aqueous CsOH). If necessary, the osmolarity was adjusted with D-mannitol such that it was slightly higher outside than inside in order to prevent cell swelling, which can make it difficult to establish a high-resistance seal. CLC-2 currents were measured in whole-cell patch-clamp mode, using an Axopatch-200B amplifier (Axon Instruments, Union City, CA), an InstruTECH ITC-16 interface (HEKA Instruments, Holliston, MA) and a Mac Mini computer (Apple, Cupertino, CA). Igor Pro (WaveMetrics, Portland, OR) software was used for stimulation and data collection. The data were sampled at 5 kHz and filtered at 1 kHz. From a holding potential of 0 mV, currents were elicited with 250-ms voltage steps of –100 to +20 mV in 10-mV increments, followed by a 100-ms tail pulse to +80 mV (**Figure 4C**). Note: we used relatively moderate hyperpolarizing potentials (compared to those used to study the full voltage-dependence of CLC-2 (87, 88) to ensure stability of the voltage-clamp seals over the course of AK-42 application and washout. All experiments were carried out at room temperature (22–25 °C). Following initial measurement of currents (“initial”), AK-42 was flowed into the recording chamber, and current was measured again after 5–10 minutes. At least two voltage-step families were measured in order to ensure that steady-state inhibition had been reached (“inhibited”). Currents were measured again following ∼10 minutes of washout (“washout”), again ensuring steady-state had been attained. For fitting dose-response curves (**Figure S2**), we excluded experiments in which there was poor reversibility of inhibition. Poor reversibility was defined as when reversal current differed by >20% from initial current and/or inhibition calculated using washout current differed by >10% from inhibition calculated using initial current. An IC_50_ value and a Hill coefficient (n) were obtained with Sigmaplot 13.0 (Systat Software, San Jose, CA) by fitting dose-response curves to the following equation: I (%) = 1/{1 + (IC_50_/[D])^n^}, in which I (%) is the percentage current inhibition {I (%) = [1 – I_inhibited_ / I_initial_] × 100} at the –100 mV test potential and [D] represents various concentrations of inhibitor. Results were each expressed as a mean ± SEM. Analysis of variance (ANOVA) was used for statistical analysis.

### Mutagenesis strategy

CLC-2 mutants were generated by site-directed mutagenesis using a QuickChange XL Site-Directed Mutagenesis Kit (Agilent Technologies, Cedar Creek, TX). The following table lists primers used to generate expression vectors in this study.

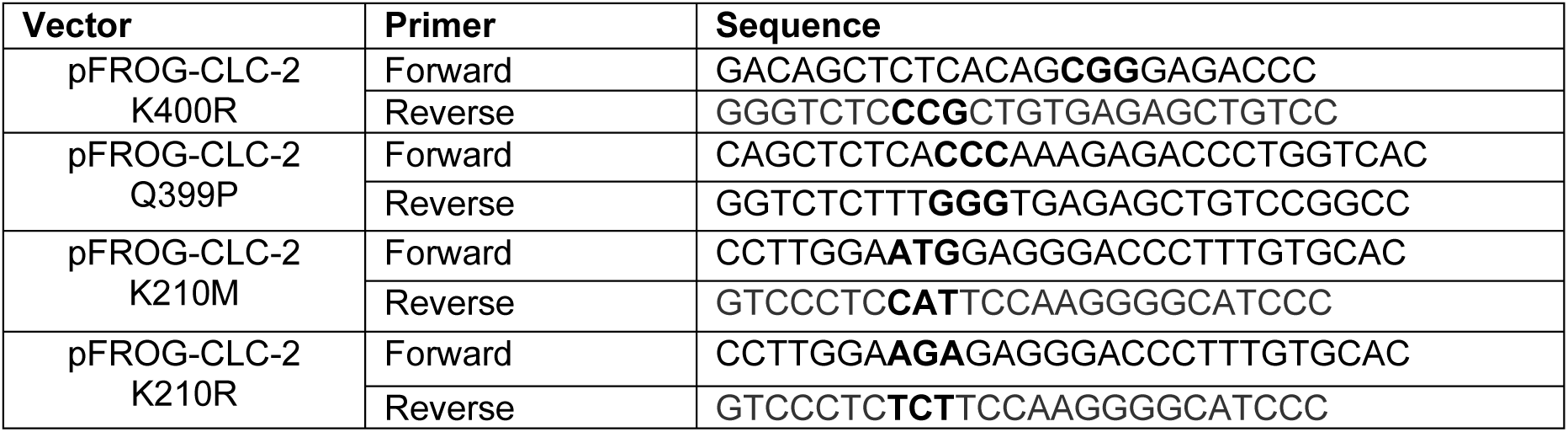

### PDSP screening results

AK-42 was screened against a panel of receptors, transporters, and ion channels to examine specificity for CLC-2 in CNS tissue. A comprehensive primary binding assay screen was performed through the NIH Psychoactive Drug Screening Program (PDSP) at UNC-Chapel Hill. Secondary or functional assays were performed for compounds showing >50% activity in the primary assay. All results are shown in **Dataset 2**.

### CLC-2 KO mice

Heterozygous CLC-2 KO mice (*Clcn*2^+/–^)(26) were bred and litters genotyped to identify littermate *Clcn*2^−/−^ and *Clcn*2^+/+^ mice, which were paired in each experiment. On any experimental day brain slices from a pair of young mice (P30–P45) were prepared, and then slices separated by group with the identity blinded to the investigator.

### Electrophysiological recordings in acute hippocampal slices

250-µM thick coronal slices were prepared from 30–35 day old WT C57/B6J or KO (*Clcn*2^−/−^) mice as described previously (89). After recovery for 1 hour, slices were transferred to a recording chamber at 23.5 °C containing oxygenated ACSF (in mM): 2.5 KCl, 10 glucose, 10 CsCl, 116 NaCl, 1.25 sodium phosphate hydrate, 1.0 MgSO_4_, 2.0 CaCl_2_, 26 NaHCO_3_. Whole-cell voltage-clamp recordings were made under constant perfusion at room temperature from CA1 hippocampal pyramidal cells under conditions that isolate and promote stable recordings of hyperpolarization-activated CLC-2 currents. These include Cs^+^ as the main internal monovalent cation to block K^+^ currents, high internal [Cl^−^] to make E_Cl_ = 0 mV and facilitate recordings at a physiological membrane potential of –80 mV, 10 mM external Cs^+^ to block HCN channels and 50 µM picrotoxin/1.5 mM kynurenic acid to block synaptic responses. Internal solution for voltage-clamp contained the following (in mM): 10 HEPES, 130 CsCl, 10 EGTA, 3 MgATP, 0.3 Na_2_GTP hydrate, 5 QX-314. For long recordings in **Figure S4**, internal solution was modified to include an ATP regenerating system and was comprised of (in mM): 10 HEPES, 125 CsCl, 10 EGTA, 4 MgATP, 0.3 Na_2_GTP, 5 QX-314, 15 phosphocreatine disodium hydrate, 50 U/mL creatine phosphokinase. Internal solution for current-clamp included (in mM): 105 K-gluconate, 10 HEPES, 10 NaCl, 20 KCl, 0.6 EGTA, 5 MgATP, 0.3 Na_2_GTP. ACSF for current-clamp omitted picrotoxin and cesium. Recordings were performed using an Axon Instruments Multiclamp 700A amplifier and Digidata 1550B digitizer (Molecular Devices, San Jose, CA). 10 KHz bessel filtering was applied during the recording.

After achieving whole-cell configuration, cells were allowed to stabilize for 2 minutes before beginning each recording. Recordings in **Figure S4** were obtained (2 per slice), one before AK-42 was added and one after washing the slice for 10 minutes with AK-42. Cell membrane parameters were monitored before and after each voltage-clamp protocol to ensure the stability of each recording. Recordings were excluded if the series resistance was greater than 20 MΩ or changed by more than 25%, or if the holding current at –60 mV exceeded –200 pA. Recordings in **Figure 4** were obtained 1 per slice in ACSF containing either 2.5 µM AK-42 or no inhibitor. Access resistance was measured every 4 minutes. Voltage step protocols were 2 seconds at –4 mV, followed by a 3-second step at the test voltage ranging between –100 mV and +20 mV in 10 mV increments before stepping back to –4 mV for 0.5 seconds. Continuous monitoring of whole-cell currents in **Figure S4, C–E** used a modified voltage protocol utilizing a 4-second conditioning step at –4 mV and a 4-second test step to either –20 or –80 mV. Protocols were run every 30 seconds along with a membrane test.

Data were measured using Clampfit 10.6 (Molecular Devices, San Jose, CA). Steady-state currents were measured as the RMS current in the last 200 ms of the test voltage step. Current relaxation was measured to be the RMS current between 200 and 400 ms into the test voltage step minus the steady state current. Two-way repeated measures ANOVA was run in GraphPad Prism 6.01 (GraphPad Software, San Diego, CA) to compare treatment groups.

### Computational Docking

Initial docking studies involved preparation of the receptor and ligand using the dock prep tool in the UCSF Chimera package (90). The receptor surface and ligand were parameterized using the AMBER parm99 force field (91). A spacing of 0.3 Å was used for the receptor grid. Initial grids included the entire extracellular surface of the protein. Once prepared, docking calculations were performed using Dock6 (92). These calculations employed flexible docking and yielded 1000 ligand orientations. From this data set, the highest scoring conformations were analyzed. The results revealed clustering of ligands in a single extracellular pocket above the ion permeation pathway. Following identification of a general binding site, additional docking poses were generated for the most potent lead compounds, using the Schrödinger Maestro software suite (Version 2019-2, Schrödinger, LLC, New York, NY, 2019). Prior to docking, ligands were optimized using the OPLS3e (93) force field in LigPrep (Schrödinger, LLC, 2019). Possible ligand tautomers and protonation states at pH 7.0 ± 2.0 were sampled using Epik (94) (Schrödinger, LLC, 2019). Ligands were docked to grids centered on the general binding site (residue K210) for each of the conformational states developed from the CLC-2 homology model described in McKiernan, et al. (73), as well as CLC-2 homology models generated from the 5TQQ and 5TR1 cryo-EM structures of bovine CLC-K (74). Alignment of the CLC-2 sequence with these structures for generation of the homology models was accomplished with the UCSF Chimera sequence alignment tool (90). The CLC-2 sequence was then mapped onto the template structure, using the Modeller package (95). Unresolved regions of the template structure were also refined using Modeller. Docking to the CLC-1 cryo-EM structure (6COY) (78) was performed using a grid centered on the equivalent lysine residue for this homolog (K231). Docking grids were generated using the default Maestro settings for receptor rigidity (Van der Waals radius scaling factor of 1.0 with a partial charge cutoff of 0.25). Prepared ligands were docked to these grids using the default settings for Schrödinger Glide (96) (Schrödinger, LLC, 2019, Van der Waals radius scaling factor of 0.80 with a partial charge cutoff of 0.15). Standard precision (SP) docking was performed with flexible ligand sampling (sampling of nitrogen inversion and ring conformations), and post-docking minimization was performed to generate a maximum of 10 poses per ligand. A representative docking pose is illustrated in **Figure 4**.

### Chemical synthesis: General

Reagents were obtained commercially unless otherwise noted. Meclofenamate sodium, *N*-phenylanthranilic acid, and lubiprostone were purchased commercially from Sigma-Aldrich. Diclofenac sodium, indomethacin, and niflumic acid were purchased from Santa Cruz Biotechnology. Salsalate was purchased from ACROS Organics. Aceclofenac was purchased from AK Scientific. GaTx2 was purchased from Tocris Bioscience (Batch No. 1C) and stored at –20 °C, and fresh stock solutions were prepared on the day of testing. Organic solutions were concentrated under reduced pressure (∼20 Torr) by rotary evaporation. Air- and moisture-sensitive liquids and solutions were transferred via syringe or stainless steel cannula. Chromatographic purification of final carboxylate, phosphonate, and sulfate derivatives was accomplished using high performance liquid chromatography on a C18 column (Alltima C18, 10 μM, 22 × 250 mm or SiliaChrom AQ C18, 5 µM, 10 × 250 mm). Thin layer chromatography was performed on EM Science silica gel 60 F254 plates (250 mm). Visualization of the developed chromatogram was accomplished by fluorescence quenching and by staining with aqueous potassium permanganate or aqueous ceric ammonium molybdate (CAM) solution.

Nuclear magnetic resonance (NMR) spectra were acquired on a Varian Inova spectrometer operating at 300, 400, 500 or 600 MHz for ^1^H. Spectra are referenced internally according to residual solvent signals. Data for ^1^H NMR are recorded as follows: chemical shift (δ, ppm), multiplicity (s, singlet; d, doublet; t, triplet; q, quartet; quintet; m, multiplet; br, broad), coupling constant (Hz), integration. Compound concentrations were determined by quantitative NMR, using *N,N*-dimethylformamide as an internal standard in DMSO-*d*_*6*_. Infrared spectra were recorded as thin films using NaCl plates on a Thermo-Nicolet 300 FT-IR spectrometer and are reported in frequency of absorption. Low-resolution mass spectra were obtained from the Vincent Coates Foundation Mass Spectrometry Laboratory and the Stanford ChEM-H facility, using a Shimadzu 20-20 ESI mass spectrometer and a Phenomenex Synergi 4 µm Hydro-RP 80 Å reversed phase column (30 × 2 mm column, gradient flow 0:1→1:0 MeCN/H_2_O with 0.1% formic acid over 4 min). Microwave reactions were performed in a Biotage Initiator microwave reactor.

### Inhibitor stock solution preparation and quantification

Meclofenamic acid (MCFA) derivatives were quantified by ^1^H NMR spectroscopy using distilled *N,N*-dimethylformamide (DMF) as an internal standard. Use of DMF as the internal standard allows for recovery of pure material following lyophilization. Each MCFA derivative was weighed into an Eppendorf tube, using a calibrated analytical balance (Mettler Toledo, Model XS105), and dissolved into DMSO-*d*_*6*_ to a final concentration of 30–100 mM. To ensure complete dissolution, the sealed Eppendorf tube was inverted at least 5 times and then sonicated for ∼60 seconds. Using a calibrated analytical balance, DMF (22 mg) was weighed into a scintillation vial, 3.0 mL of DMSO-*d*_*6*_ (stored in a desiccator jar and preferably newly opened to minimize water contamination in the solvent) was added to the vial via a p1000 micropipette, and the solution was thoroughly mixed by inverting the capped vial at least 10 times (note: to prevent cross-contamination between samples and for convenience, disposable micropipette tips may be used without affecting the accuracy of the measurements). To ensure robustness of the quantification protocol, the 100 mM DMF stock solution was independently quantified in triplicate against stock solutions with known concentrations of fumaronitrile dissolved in DMSO-*d*_*6*_. Fumaronitrile produces a sharp singlet that integrates to 2H at 7.03 ppm in DMSO-*d*_*6*_ (DMSO solvent residual peak referenced to 2.50 ppm) and may be integrated against either of the DMF methyl signals that appear at 2.73 ppm (3H) and 2.89 (3H) ppm. The average of these three NMR measurements was used for calculation of the final DMF internal standard concentration. All measurements were performed at room temperature. Stock solutions were stored frozen and left at ambient temperature for several hours to thaw and vortexed prior to quantification. A relaxation delay time (d1) of 20 s and an acquisition time (at) of 10 s were used during spectral acquisition. The number of scans was typically set to 32 unless a particular compound concentration was low (< 10 mM) such that more scans were required to improve signal/noise. The concentration of each MCFA derivative was determined by comparison with signal integrations for the MCFA derivative and the DMF internal standard.

## Acknowledgements

Receptor binding profiles, and agonist/antagonist functional data were generously provided by the National Institute of Mental Health’s Psychoactive Drug Screening Program, Contract #HHSN-271-2018-00023-C (NIMH PDSP). The NIMH PDSP is directed by Bryan L. Roth MD, PhD at the University of North Carolina at Chapel Hill and Project Officer Jamie Driscoll at NIMH, Bethesda, MD, USA. *Clcn2*^−/−^ mice were generously provided by J. Melvin (University of Rochester). The CLC-2 construct used for manual voltage-clamp recordings was kindly provided by M. Pusch (Istituto di Biofisica, Genova, Italy), with permission from Thomas Jentsch. This work was supported by a Stanford Bio-X Seed grant (to J.D.B and M.M.) and NIH grant # R01NS113611 (to J.D.B., M.M., and J.R.H.). A.K.K. and K.A.M. were supported by the Stanford Center for Molecular Analysis and Design (CMAD). A.K.K. was also supported by a Stanford Interdisciplinary Graduate Fellowship (SIGF) through the Stanford ChEM-H Institute.

## Figures & Tables

**Figure S1.**
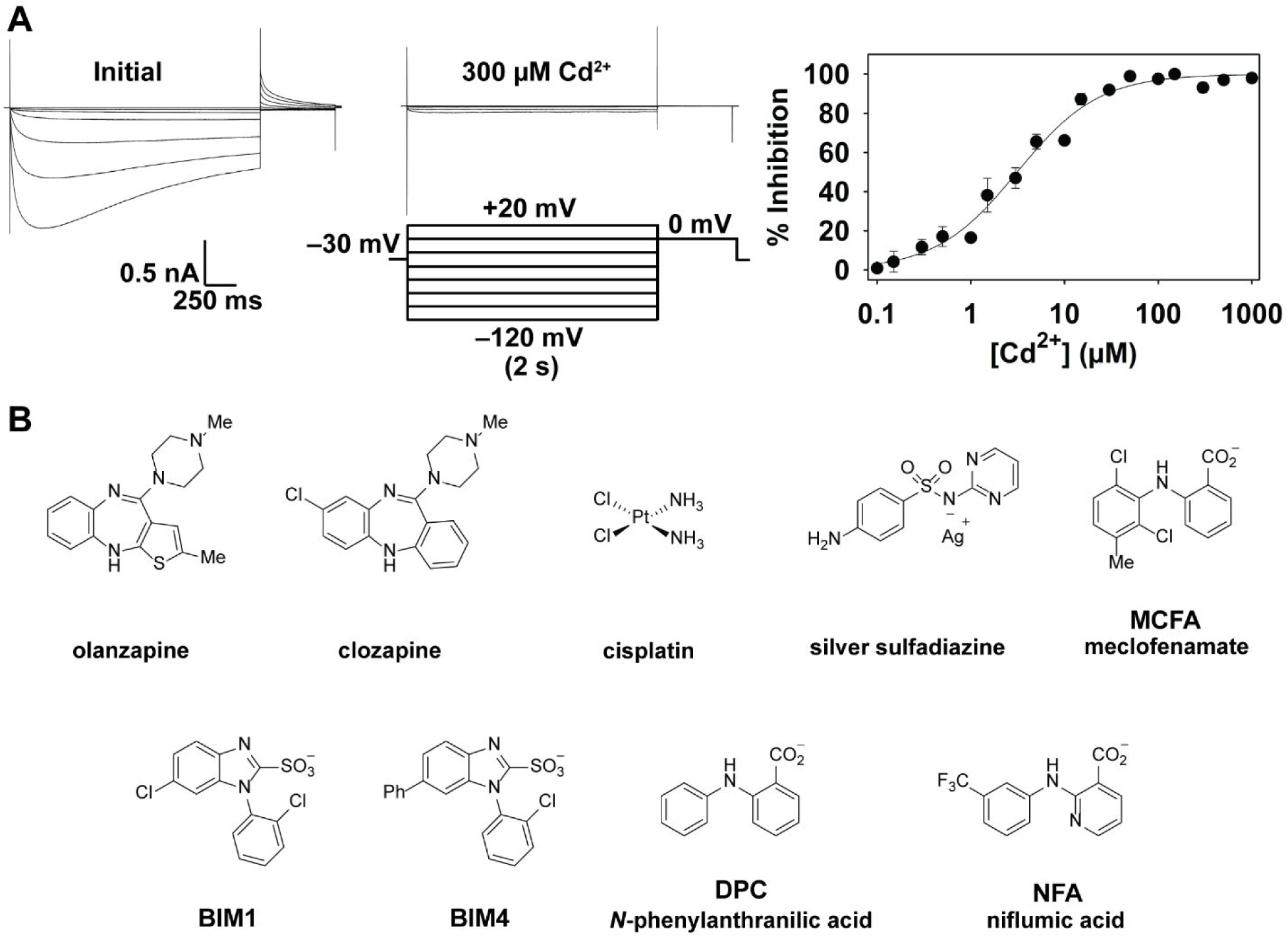
CLC-2 assay on the IWB and identification of ‘hit’ compounds. (**A**) Assay development. *Left:* Representative human CLC-2 currents on IWB measured before (left) or after (right) treatment with positive control Cd^2+^ in response to the voltage protocol shown. The current decay at negative voltages, which is not seen in manual patch-clamp recordings (**Figure 4C** and references (60, 97, 98)) or in a different automated patch-clamp platform (PatchXpress, unpublished data), is likely due to the differences in the intracellular solution, which in this case includes a mixture of Cl^−^ and F^−^. *Right:* Summary data for inhibition of CLC-2 by Cd^2+^ (± SEM, n = 4–32; IC_50_ = 3.1 ± 0.3 µM). Inhibition was calculated using the maximum current at –120 mV in the presence or absence of Cd^2+^. Assay-validation studies showed a Z-factor of 0.83 and 0.73 on separate days. (**B**) Structures of representative compounds. *Top:* Structures of top five ‘hit’ compounds identified in the IWB screen of 772 FDA-approved compounds (ENZO Life Sciences). *Bottom:* Representative structures of compounds known to inhibit other CLC channels (55, 99) but found to be ineffective inhibitors of CLC-2 in our screen. DPC and NFA, like the ‘hit’ compound MCFA, are NSAIDs.

**Figure S2.**
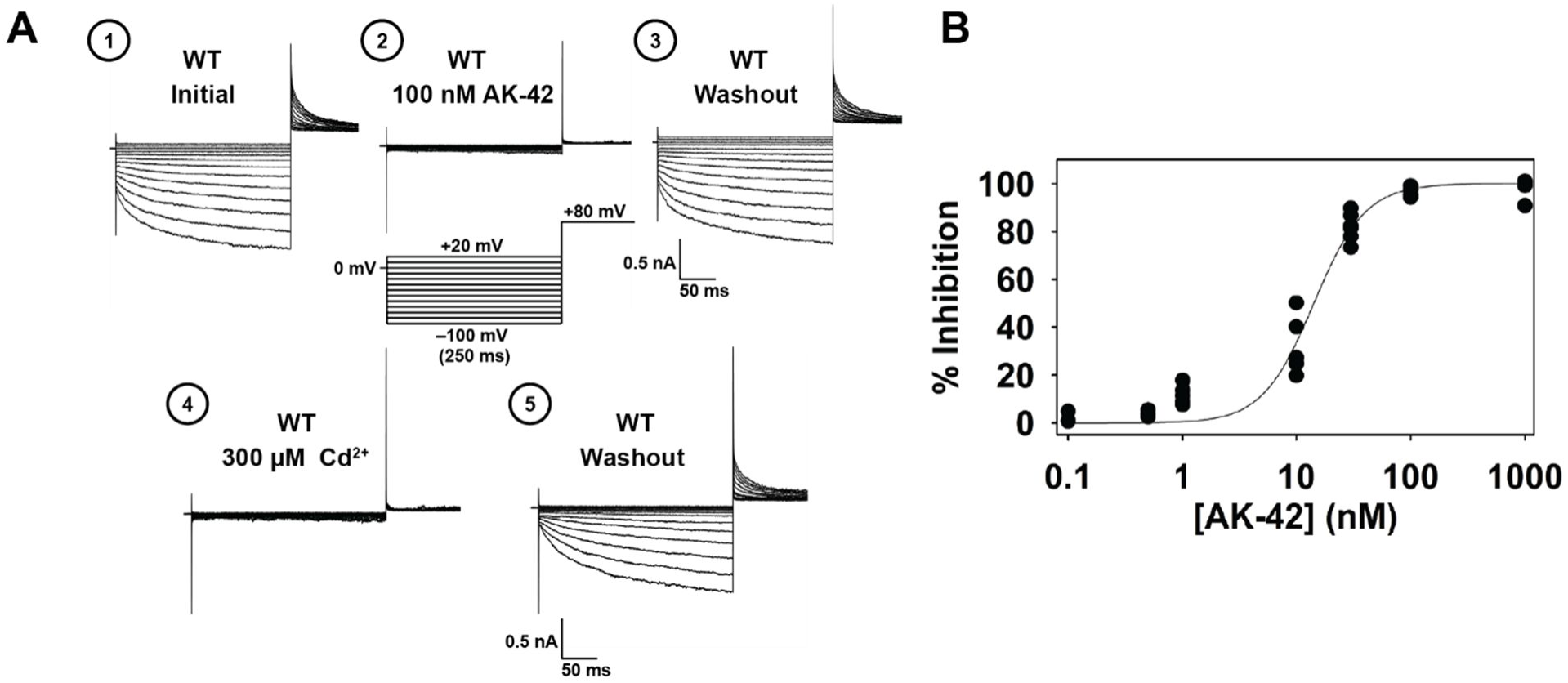
Manual patch-clamp recording of rat CLC-2. (**A**) Representative traces showing rat CLC-2 currents in transiently transfected CHO cells in response to the voltage protocol shown: before, after, and following washout of 100 nM AK-42. A saturating concentration of Cd^2+^ (the low-potency CLC-2 inhibitor used in assay development, **Figure S1**) was added at the end of each experiment (Step 4) to facilitate subtraction of background currents on a given cell; Cd^2+^ was subsequently washed out (Step 5). Steps 1– 5 of a typical experiment are shown. (**B**) Summary inhibition data for AK-42 concentration. Individual data points are shown for inhibition at –100 mV for 0.1 nM (n = 3), 0.5 nM (n = 3), 1 nM (n = 5), 10 nM (n = 5), 30 nM (n = 6), 100 nM (n = 4), 1 µM (n = 4). The fitted IC_50_ value (14 ± 1 nM) is comparable to that obtained for human CLC-2 in the IWB assay (17 ± 1 nM at –120 mV).

**Figure S3.**
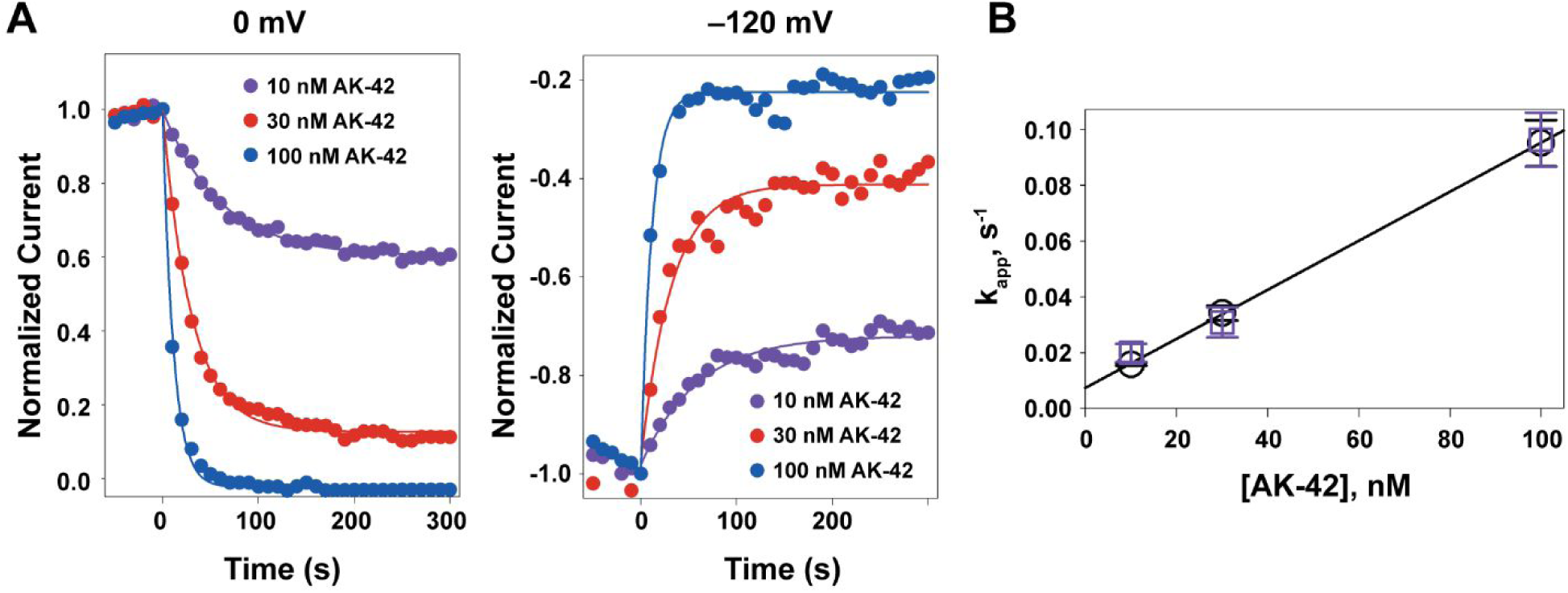
Kinetics of CLC-2 inhibition support that AK-42 acts at a single site from the extracellular side. (**A**) Representative data showing the time course of CLC-2 inhibition by 10, 30, or 100 nM of AK-42. Currents were measured using the IWB; cells were held at the reversal potential (–30 mV), and currents were measured with 2-s test pulses to –120 mV followed by 0.5-s tail pulses to 0 mV, every 10 s. The peak current amplitudes in both the test and tail pulses were measured and plotted as a function of time. Data were fitted with a single exponential function to obtain values for *k*_*app*_ (apparent rate constant). (**B)** Plot of *k*_*app*_ values as a function of AK-42 concentration at –120 mV (purple) or 0 mV (black) for n = 3 (10 nM) or n = 4 (30 nM, 100 nM) cells. The linear relationship between *k*_*app*_ and AK-42 illustrates that inhibition is a first-order process, involving a 1:1 CLC-2 subunit/AK-42 interaction. Regression analysis (fitting simultaneously to both sets of points) yields estimates for on- and off-rates of 9 × 10^5^ M^−1^ s^−1^ (slope) and 8 × 10^−3^ s^−1^ (intercept). While the IWB is not set up to allow measurements of reversal, we confirmed reversibility of inhibition using manual patch-clamp recordings (**Figure S2**), with reversal occurring within ∼10 min, consistent with the off-rate estimated using the IWB.

**Figure S4.**
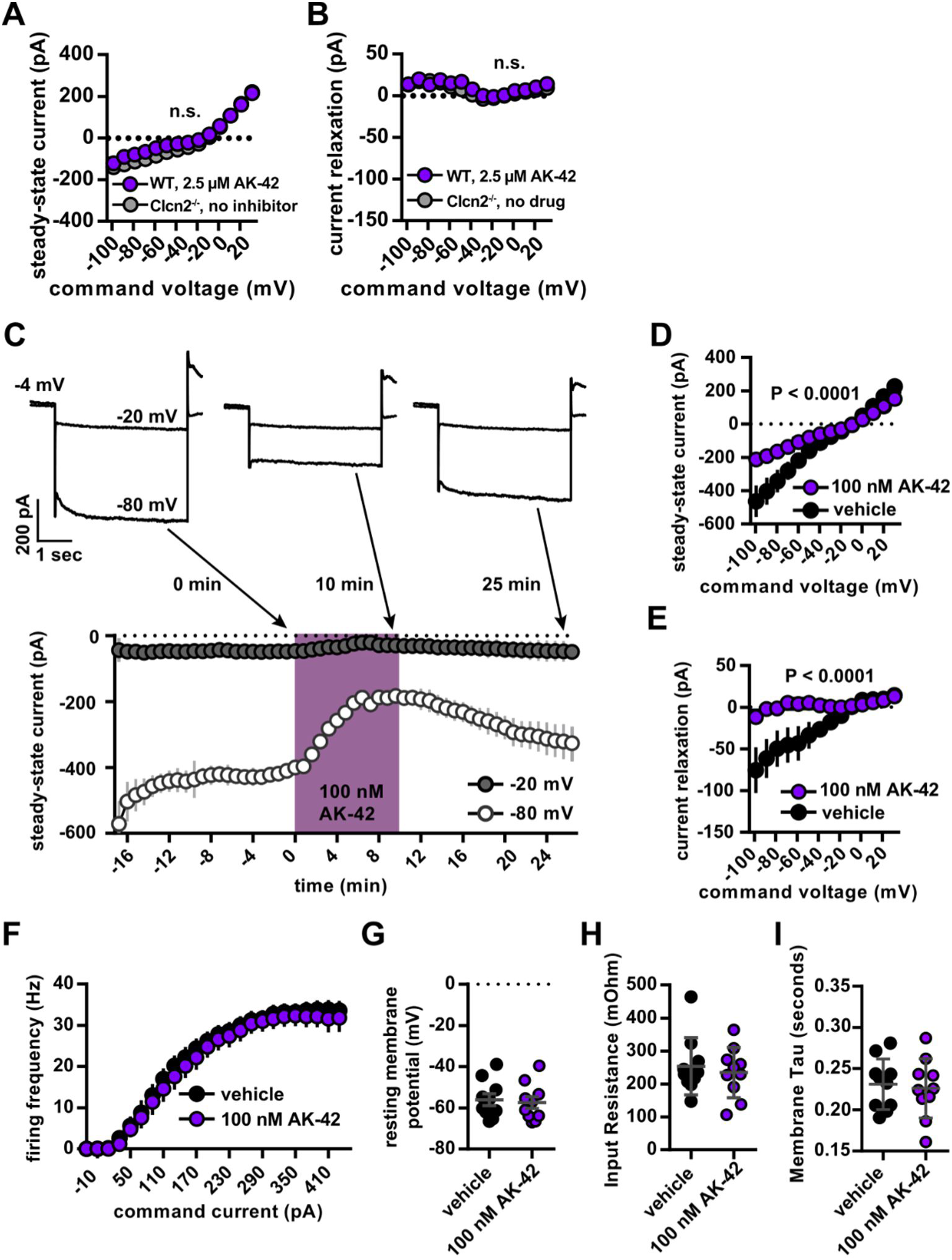
Confirmation of AK-42 efficacy and specificity. (**A–B**): *Clcn2*^−/−^ control. Steady-state currents (**A**) and current relaxation measurements (**B**) are indistinguishable between wild-type cells treated with 2.5 µM AK-42 and untreated *Clcn2*^−/−^ (P = 0.53 and 0.63 for **A** and **B**, respectively via two-way RM ANOVA.). (**C–E**): AK-42 is effective and reversible at 100 nM. (**C**) Time course (bottom) and representative whole-cell current traces (top) show reversible inhibition of CLC-2 by 100 nM AK-42. Note that AK-42 visibly decreases current at hyperpolarized (–80 mV) but not depolarized (–20 mV) potentials. Capacitive transients are clipped for display purposes. (**D–E**) Summary data showing I-V relationship of wild-type whole-cell steady-state currents (**D**) and current relaxation (**E**) before and after 10 minutes of AK-42 treatment at 100 nM (n = 6 cells, 6 slices and 6 animals. P < 0.0001 via two-way RM ANOVA). (**F–I**): Specificity of AK-42 as evidenced by lack of effects on firing frequency and membrane parameters. (**F**) Firing frequency of CA1 pyramidal cells in response to a 500-ms current injection is not changed after the application of 100 nM AK-42 for 10 minutes. Error bars represent ± SEM throughout. (P = 0.66 via two-way RM ANOVA. n = 10 cells, 10 slices from 6 animals). (**G–I**) Membrane parameters remain unchanged after 10 minutes of AK-42 application (100 nM), calculated from current-clamp recordings in panel F. (P = 0.73, 0.61, and 0.71, respectively, via Student’s unpaired *t*-test.)

**TABLE S1.**
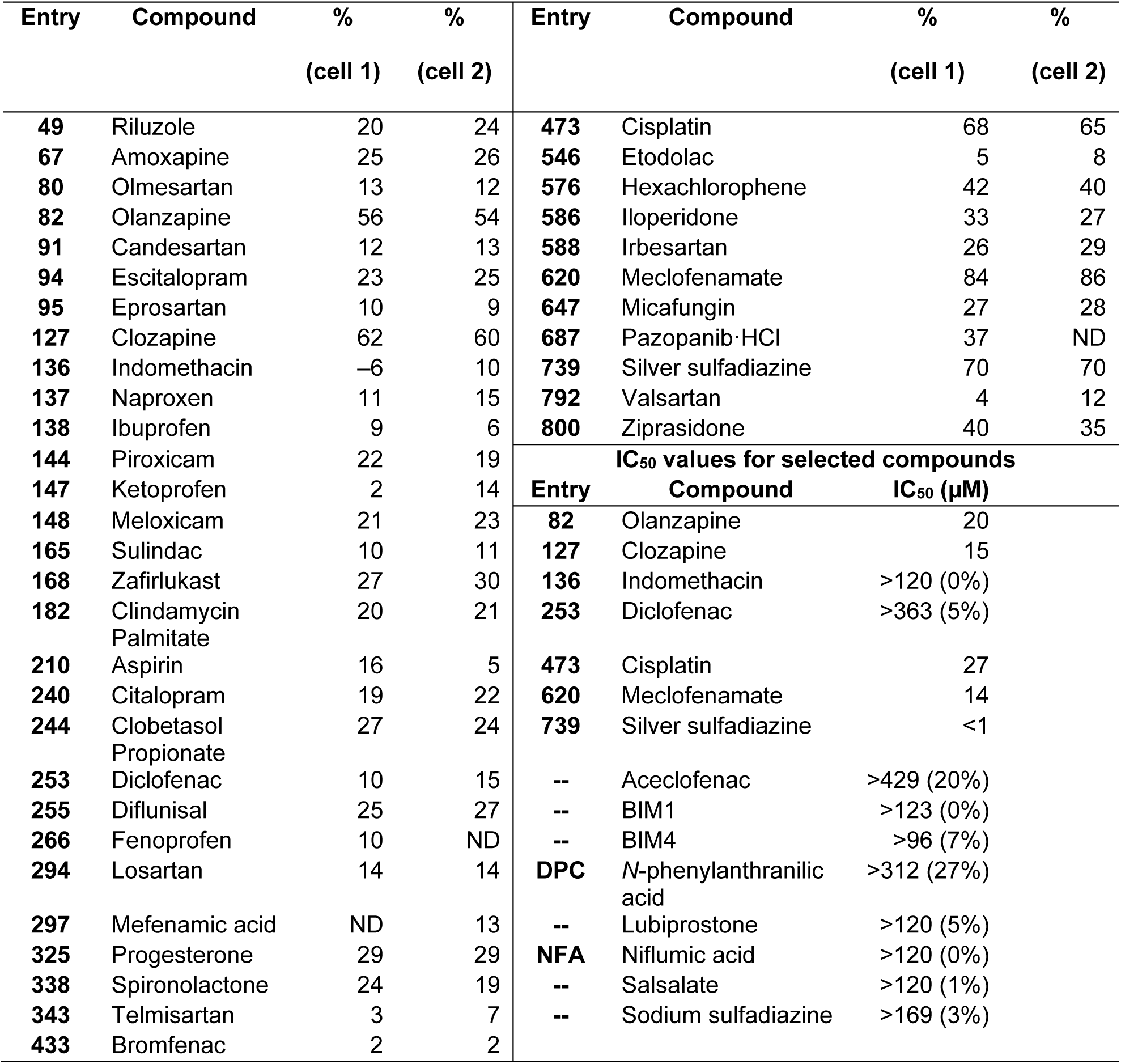
Summary inhibition data for initial ‘hit’ compounds (>20% inhibition of CLC-2 at –120 mV) and for selected NSAIDs and AT_1_ antagonists sampled from the ENZO library. Percent inhibition of CLC-2 current by 30 µM compound at –120 mV is shown for each of two cells. Results from the complete screen (including those shown here) are available in **Dataset 1**. Approximate IC_50_ values of the most potent hit compounds and other selected compounds are shown in the lower right portion of the table. Values were estimated from inhibition measured at 4 concentrations of compound (1, 3, 10, 30 µM, n = 3–4 per concentration) using the IWB platform. For compounds that exhibit little or no inhibition of CLC-2, the IC_50_ is listed as > the highest concentration of compound tested, and the amount of inhibition observed at this concentration is shown in parentheses.

**TABLE S2.** Final IC_50_ values for selected compounds against human CLC-1 and human CLC-2, using the IWB platform. If the IC_50_ was greater than the highest concentration tested, this concentration is listed along with the % inhibition at this concentration (in parenthesis). *For MCFA, the maximum inhibition of CLC-1 at 100 µM was 61%; thus, this IC_50_ value is an approximation.

| Compound | IC<sub>50</sub> (CLC-2) | IC<sub>50</sub> (CLC-1) |
| --- | --- | --- |
| **MCFA** | 7 ± 1 μM | ~50 μM* |
| **AK-24** | 1.2 ± 0.2 μM | >30 μM (25%) |
| **AK-33** | 3 ± 1 μM | >30 μM (5%) |
| **AK-42** | 0.017 ± 0.001 μM | >100 μM (22%) |

**TABLE S3.** % inhibition values for human CLC-1 and human CLC-2 with AK-42, as shown in **Figure 3**. Values of over 100% reflect that some current measurements at –120 mV flipped from negative to slightly positive in the presence of inhibitor.

| Concentration (μM) | % inhibition, CLC-1 | % inhibition, CLC-2 |
| --- | --- | --- |
| **0.0003** | –2 ± 6 (n = 4) | –1 ± 6 (n = 4) |
| **0.001** | 1 ± 6 (n = 4) | 3 ± 3 (n = 4) |
| **0.003** | 3 ± 3 (n = 4) | 15 ± 4 (n = 4) |
| **0.01** | 2 ± 3 (n = 4) | 32 ± 4 (n = 4) |
| **0.03** | –2 ± 2 (n = 8) | 67 ± 2 (n = 8) |
| **0.1** | 0 ± 2 (n = 7) | 86 ± 1 (n = 8) |
| **0.12** | not determined (nd) | 90 ± 5 (n = 4) |
| **0.3** | –2 ± 1 (n = 7) | 95 ± 2 (n = 8) |
| **1** | –4 ± 2 (n = 7) | 100.0 ± 0.3 (n = 8) |
| **1.2** | nd | 105.1 ± 0.7 (n = 4) |
| **3** | 4 ± 2 (n = 4) | 100.3 ± 0.4 (n = 4) |
| **10** | 5 ± 2 (n = 4) | 100.3 ± 0.4 (n = 4) |
| **12** | nd | 104.0 ± 0.8 (n = 3) |
| **30** | 7 ± 3 (n = 4) | 100.3 ± 0.3 (n = 4) |
| **100** | 22 ± 2 (n = 4) | 99.1 ± 0.2 (n = 4) |
| **120** | nd | 106.3 ± 0.5 (n = 4) |

**Table S4.** % inhibition values for WT CLC-2 and four mutants (K400R, Q399P, K210M, K210R) with 30 nM AK-42 at –100 mV, as shown in **Figure 4D**.

| CLC-2 plasmid | % inhibition |
| --- | --- |
| WT | 82 ± 3 (n = 6) |
| K400R | 82 ± 13 (n = 3) |
| Q399P | 17 ± 5 (n = 3) |
| K210M | 23 ± 9 (n = 3) |
| K210R | 27 ± 8 (n = 3) |

## Supplementary Information

All results from the ENZO library screen are available in **Dataset 1**. All specificity data from the PDSP assays are available in **Dataset 2**. Detailed chemical synthesis protocols and characterization data for all inhibitors and intermediates are provided in **Dataset 3**.

