## Supplementary Material for "Development and validation of a potent and specific inhibitor for the CLC-2 chloride channel"

### *Dataset 1: Compound library screening results*

The percent inhibition of current at 30 µM was determined for each compound in the ENZO Life Sciences SCREEN-WELL® FDA-approved drug library, v. 2.0 (Product # BML-2843-0100). Compounds were screened against human CLC-2 stably expressed in CHO cells. Values were acquired on n = 2 cells using the IonWorks™ Barracuda patch clamp electrophysiology system. Both values were reported for block of peak current at –120 mV. If a compound elicited an increase in peak current instead of inhibition, values are report as a negative percentage. Entries labeled as 'VEHICLE' represent controls in which 0.3% DMSO in recording solution was applied between measurements (n = 28, 6 ± 4%). ND = not determined.

**Dataset 1**, **Table 1.** Automated patch-clamp screen of ENZO library at 30 µM against human CLC-2.

| Entry | Compound | % (n = 1) | % (n = 2) | Entry | Compound | % (n = 1) | % (n = 2) |
|---|---|---|---|---|---|---|---|
| **1** | Clindamycin·HCl | 6.4 | 17.2 | **401** | Ambrisentan | 9.1 | 7.2 |
| **2** | Felbamate | 9.4 | 11.7 | **402** | Amcinonide | 3.1 | 5.7 |
| **3** | Cyclosporine A | 10.7 | 16.8 | **403** | Amikacin Disulfate | 11.7 | 10.1 |
| **4** | Donepezil·HCl | 13.5 | –17.1 | **404** | Aminocaproic Acid | 8.3 | 9.8 |
| **5** | Lincomycin·HCl | 9.1 | 14.6 | **405** | Aminohippurate·Na | 4.6 | 5.5 |
| **6** | Mycophenolic Acid | 10.7 | 15.9 | **406** | Aminolevulinic Acid·HCl | 1.3 | 12.3 |
| **7** | Sirolimus (Rapamycin) | 11.1 | 12.7 | **407** | Amlexanox | 6.8 | –11.3 |
| **8** | Spectinomycin·HCl Pentahydrate | 13.0 | 10.1 | **408** | Amphotericin B | 7.5 | 1.7 |
| **9** | Amiodarone·HCl | 7.8 | 11.9 | **409** | Arsenic Trioxide | 3.9 | ND |
| **10** | Nicardipine·HCl | 10.6 | 8.5 | **410** | Artemether | –5.8 | 2.3 |
| **11** | Pimozide | 2.7 | 8.7 | **411** | Articaine·HCl | 13.0 | 3.7 |
| **12** | Loperamide·HCl | 14.4 | –13.6 | **412** | L-Ascorbic Acid | 5.3 | 8.4 |
| **13** | Tolbutamide | 28.8 | 5.0 | **413** | Asenapine Maleate | 15.7 | 1.2 |
| **14** | Glipizide | 14.2 | 11.1 | **414** | Atomoxetine·HCl | 2.3 | 5.5 |
| **15** | Phentolamine·HCl | 4.2 | 8.2 | **415** | Atorvastatin Calcium | 0.4 | 8.1 |
| **16** | Quinine·HCl·$H_2O$ | 15.1 | –6.6 | **416** | Azacitidine | –1.0 | 8.2 |
| **17** | Propafenone·HCl | 7.6 | 9.2 | **417** | Azelaic Acid | 6.7 | 6.9 |
| **18** | Phenytoin | 12.6 | 10.8 | **418** | Azelastine·HCl | –2.1 | –1.7 |
| **19** | Procainamide·HCl | 7.4 | 13.2 | **419** | Bacitracin | 0.1 | –0.8 |
| **20** | Lidocaine·HCl·$H_2O$ | 12.3 | 12.2 | **420** | Baclofen | 0.9 | –0.5 |
| **21** | Flecainide Acetate | 11.8 | 13.7 | **421** | Balsalazide | 4.7 | 10.6 |
| **22** | Rosiglitazone | 12.9 | 14.0 | **422** | Beclomethasone Dipropionate | 1.6 | 4.2 |
| **23** | Amantadine·HCl | 8.5 | 11.6 | **423** | Benazepril·HCl | 6.6 | 1.3 |
| **24** | Prazosin·HCl | 13.3 | 11.7 | **424** | Bendamustine·HCl | 7.0 | 5.0 |
| **25** | Clonidine·HCl | 10.9 | 14.0 | **425** | Bendroflumethiazide | 13.2 | 8.0 |
| **26** | Guanabenz Acetate | 14.9 | 31.3 | **426** | Benztropine Mesylate | –4.6 | –5.8 |
| **27** | Dihydroergotamine Mesylate | 19.8 | 11.2 | **427** | Betaine | 4.6 | 3.5 |
| **28** | Emtricitabine | 9.6 | 12.9 | **428** | Bethanechol Chloride | 4.9 | 7.7 |
| **29** | Betaxolol·HCl | 12.9 | 9.7 | **429** | Bimatoprost | 4.2 | 2.9 |
| **30** | Caffeine | 13.9 | 11.2 | **430** | Biperiden·HCl | –48.8 | –41.9 |

| | | | | | | | |
|---|---|---|---|---|---|---|---|
| **31** | (S)-Timolol Maleate | 6.0 | 9.4 | **431** | Bisoprolol Fumarate | 6.7 | 5.1 |
| **32** | Salbutamol Hemisulfate | 12.4 | 10.8 | **432** | Brimonidine | 4.8 | 4.4 |
| **33** | Pindolol | 11.4 | 14.6 | **433** | Bromfenac | 2.0 | 1.9 |
| **34** | Dobutamine·HCl | 13.3 | 12.5 | **434** | Brompheniramine Maleate | 6.5 | 3.1 |
| **35** | Sotalol·HCl | 10.2 | 11.0 | **435** | Budesonide | 4.8 | 4.2 |
| **36** | Maprotiline·HCl | 11.9 | 13.6 | **436** | Bupropion | 1.8 | 2.0 |
| **37** | Pilocarpine·HCl | 8.2 | 9.2 | **437** | Busulfan | 4.0 | 3.8 |
| **38** | Ipratropium·Br | 9.3 | 13.4 | **438** | Butorphanol-(+)-Tartrate | –33.2 | –23.1 |
| **39** | Tropicamide | 14.0 | 12.6 | **439** | Capreomycin Disulfate | 8.8 | 6.8 |
| **40** | Pancuronium·2Br | 9.9 | 6.9 | **440** | Carbinoxamine Maleate | 4.1 | –0.1 |
| **41** | Ivermectin | 13.5 | 15.3 | **441** | Carglumic Acid | 11.6 | 4.6 |
| **42** | Haloperidol | 14.5 | 12.4 | **442** | VEHICLE | –5.5 | –3.6 |
| **43** | Cimetidine | 10.0 | 15.2 | **443** | Carmustine | 8.4 | –2.3 |
| **44** | Zonisamide | 8.8 | 17.3 | **444** | Cefaclor | 5.2 | 5.1 |
| **45** | Zoledronic Acid Monohydrate | 14.0 | 15.5 | **445** | Cefadroxil | –4.3 | –4.8 |
| **46** | Naltrexone·HCl | 1.0 | 0.5 | **446** | Cefazolin·Na | 8.4 | 6.9 |
| **47** | Zolmitriptan | 10.7 | 21.1 | **447** | Cefdinir | –0.5 | 4.1 |
| **48** | Memantine·HCl | 6.3 | 8.0 | **448** | Cefditoren Pivoxil | 7.1 | 2.6 |
| **49** | Riluzole·HCl | 20.1 | 24.2 | **449** | Cefixime | 4.6 | 3.8 |
| **50** | Propofol | 9.3 | 8.4 | **450** | Cefotetan Disodium | 6.3 | 0.8 |
| **51** | Aminophylline | 14.6 | 11.8 | **451** | Cefoxitin·Na | 10.8 | 5.8 |
| **52** | Nateglinide | 17.0 | 11.6 | **452** | Cefpodoxime Proxetil | 2.0 | 1.3 |
| **53** | (±) Isoproterenol·HCl | 10.3 | 12.9 | **453** | Cefprozil | 9.0 | –2.6 |
| **54** | Acetylcholine Chloride | 1.9 | 15.2 | **454** | Ceftibuten | 1.5 | 7.5 |
| **55** | Atropine Sulfate Monohydrate | 10.6 | 14.0 | **455** | Ceftizoxim·Na | 8.4 | 3.0 |
| **56** | Apomorphine·HCl Hemihydrate | 16.0 | 17.6 | **456** | Ceftriaxone·Na | 2.9 | 5.1 |
| **57** | Chlorpromazine·HCl | –2.6 | 1.7 | **457** | Cefuroxime Axetil | 5.5 | –1.7 |
| **58** | Fluphenazine·HCl | 15.9 | 12.7 | **458** | Cefuroxime·Na | 7.2 | 3.4 |
| **59** | Risperidone | 11.7 | 16.1 | **459** | Cephalexin·$H_2O$ | 3.5 | –3.7 |
| **60** | Diphenhydramine·HCl | 13.7 | 5.7 | **460** | Chenodiol (Chenodeoxycholic Acid) | 9.3 | 8.4 |
| **61** | Promethazine·HCl | 13.3 | 2.1 | **461** | VEHICLE | –7.9 | –3.3 |
| **62** | Ranitidine·HCl | 18.7 | 14.1 | **462** | Chlorhexidine Dihydrochloride | –20.0 | –18.2 |
| **63** | L-(-)-Epinephrine-(+)-Bitartrate | 7.6 | 9.3 | **463** | Chlorothiazide | 5.4 | –1.1 |
| **64** | Norepinephrine Bitartrate Monohydrate | 13.9 | 12.0 | **464** | Chlorpropamide | 1.0 | 9.5 |
| **65** | Quetiapine Fumarate | 7.3 | 13.3 | **465** | Chlorthalidone | 18.8 | 9.0 |
| **66** | Imipramine·HCl | 1.3 | –2.3 | **466** | Chlorzoxazone | 15.8 | 15.6 |
| **67** | Amoxapine | 24.6 | 26.4 | **467** | Ciclesonide | 10.1 | 10.1 |
| **68** | Metoclopramide·HCl | 12.9 | 10.6 | **468** | Ciclopirox | 1.0 | –6.9 |

| | | | |
|---|---|---|---|
| **69** | Nalbuphine·HCl Dihydrate | 12.7 | 10.1 |
| **70** | Carbachol (Carbamylcholine) Chloride | 14.1 | 10.0 |
| **71** | Famotidine | 14.3 | 7.9 |
| **72** | Isoniazid | 14.3 | 12.8 |
| **73** | Ticlopidine·HCl | 12.7 | 11.2 |
| **74** | Clemastine Fumarate | –3.5 | –4.0 |
| **75** | Vardenafil | 9.8 | 13.0 |
| **76** | Linezolid | ND | 6.8 |
| **77** | Docetaxel (Taxotere) | 8.9 | 11.0 |
| **78** | Olopatadine | 14.5 | 14.4 |
| **79** | Tolcapone | 26.7 | 23.3 |
| **80** | Olmesartan | 12.7 | 11.9 |
| **81** | Nisoldipine | 8.9 | 3.9 |
| **82** | Olanzapine | 55.6 | 53.9 |
| **83** | Lovastatin | 7.3 | –1.8 |
| **84** | Lamotrigine | 9.8 | 13.7 |
| **85** | Azathioprine | 4.9 | 11.1 |
| **86** | Sildenafil Citrate | 8.3 | 5.3 |
| **87** | Atovaquone | 9.4 | 5.6 |
| **88** | Sertaconazole | 9.1 | 7.9 |
| **89** | Cefepime·HCl Hydrate | 16.5 | 6.4 |
| **90** | Aripiprazole | 17.7 | 8.3 |
| **91** | Candesartan | 12.1 | 12.6 |
| **92** | Butenafine·HCl | 5.1 | 7.1 |
| **93** | Dorzolamide·HCl | 9.6 | 10.0 |
| **94** | Escitalopram | 22.7 | 24.7 |
| **95** | Eprosartan Mesylate | 10.3 | 8.8 |
| **96** | Entacapone | 4.8 | 8.7 |
| **97** | Bleomycin Sulfate | 3.5 | 10.8 |
| **98** | Guanfacine·HCl | 12.3 | 9.5 |
| **99** | Tizanidine·HCl | 17.5 | 5.8 |
| **100** | Carvedilol | 8.1 | 11.9 |
| **101** | Flumazenil | 14.0 | 9.1 |
| **102** | Gefitinib | 17.0 | 14.5 |
| **103** | Imatinib Mesylate | 6.9 | 9.9 |
| **104** | Idarubicin·HCl | 3.9 | –1.9 |
| **105** | Montelukast·Na | 3.5 | 9.8 |
| **106** | Exemestane | 2.9 | 27.3 |
| **107** | Dinoprostone | 11.1 | 9.2 |
| **108** | Metformin·HCl | 6.5 | 13.7 |
| **109** | Anagrelide | 10.4 | 4.6 |
| **469** | Cidofovir | –10.5 | –7.6 |
| **470** | Cilostazol | 4.0 | 8.0 |
| **471** | Cinacalcet·HCl | 6.1 | 8.7 |
| **472** | Cisatracurium Besylate | 5.0 | 12.2 |
| **473** | Cisplatin | 67.5 | 64.7 |
| **474** | Cladribine | 18.6 | 12.0 |
| **475** | Clavulanate Potassium | 4.6 | 15.6 |
| **476** | Clobazam | 4.3 | 13.7 |
| **477** | Clofazimine | 8.9 | 17.9 |
| **478** | Clomipramine·HCl | –8.3 | –13.2 |
| **479** | Clonazepam | –2.0 | 10.2 |
| **480** | Clotrimazole | –19.9 | –13.8 |
| **481** | Cloxacillin·Na | 6.7 | 5.5 |
| **482** | Colchicine | 9.7 | 6.6 |
| **483** | Colistimethate·Na | 7.7 | 11.3 |
| **484** | Colistin Sulfate | 10.2 | 28.9 |
| **485** | Cortisone Acetate | 6.9 | 9.9 |
| **486** | Cyclobenzaprine·HCl | –5.9 | –3.9 |
| **487** | Cyclopentolate | 4.0 | 4.7 |
| **488** | Cycloserine | 9.3 | 6.6 |
| **489** | Cysteamine·HCl | 10.5 | 9.9 |
| **490** | Dactinomycin (Actinomycin D) | 12.4 | 12.2 |
| **491** | Dalfampridine (4-Aminopyridine) | 4.6 | 8.9 |
| **492** | Dantrolene·Na | 5.5 | 0.9 |
| **493** | Dapsone | –0.2 | 3.1 |
| **494** | Daptomycin | –3.5 | 9.9 |
| **495** | Darifenacin·HBr | 4.3 | 17.9 |
| **496** | Darunavir | 9.8 | 12.8 |
| **497** | Dasatinib | 8.3 | 15.2 |
| **498** | Decitabine | 1.6 | –2.3 |
| **499** | Deferasirox | 5.1 | 0.0 |
| **500** | Deferoxamine Mesylate | 4.6 | 1.9 |
| **501** | Demeclocycline·HCl | 11.4 | 11.5 |
| **502** | Desipramine·HCl | 3.3 | 7.3 |
| **503** | Desogestrel | 6.0 | 5.3 |
| **504** | Desonide | 11.0 | 11.5 |
| **505** | Desoximetasone | 8.3 | 14.8 |
| **506** | Desvenlafaxine Succinate Hydrate | 10.4 | 11.2 |
| **507** | Dexchlorpheniramine Maleate | 3.1 | 9.0 |
| **508** | Dexmedetomidine·HCl | –0.7 | 8.6 |
| **509** | VEHICLE | –3.8 | –0.8 |

| | | | |
|---|---|---|---|
| **110** | Dofetilide | 11.1 | 6.1 |
| **111** | Erlotinib | 14.0 | 10.5 |
| **112** | Tacrine·HCl | 10.7 | 13.5 |
| **113** | Galantamine·HBr | 8.3 | 11.9 |
| **114** | Amiloride·HCl·$2H_2O$ | 11.1 | 11.4 |
| **115** | Amlodipine | 13.7 | 11.6 |
| **116** | Diltiazem·HCl | 4.8 | 13.3 |
| **117** | Nifedipine | 9.1 | 10.0 |
| **118** | Nimodipine | –8.6 | 3.5 |
| **119** | Verapamil·HCl | 7.7 | 12.0 |
| **120** | Gabapentin | 6.5 | 10.4 |
| **121** | Felodipine | 10.6 | 13.2 |
| **122** | Phenoxybenzamine·HCl | 11.8 | 12.9 |
| **123** | Trifluoperazine· HCl | 8.1 | 7.5 |
| **124** | Latanoprost | 9.4 | 5.4 |
| **125** | Alfuzosin | 9.8 | 17.2 |
| **126** | Bromocriptine Mesylate | 10.9 | 14.7 |
| **127** | Clozapine | 62.3 | 60.0 |
| **128** | Acitretin | 4.9 | 2.3 |
| **129** | Calcitriol | 10.2 | 9.9 |
| **130** | Ketoconazole | 0.2 | 0.9 |
| **131** | Cromolyn·Na (Disodium Cromoglycate) | 10.8 | 14.1 |
| **132** | Capsaicin | 12.6 | –4.9 |
| **133** | Dexamethasone | 10.4 | 13.9 |
| **134** | Dipyridamole | 6.0 | 11.2 |
| **135** | Ethacrynic Acid | 13.3 | 16.6 |
| **136** | Indomethacin | –6.1 | 10.2 |
| **137** | Naproxen | 11.1 | 14.6 |
| **138** | Ibuprofen | 9.1 | 6.4 |
| **139** | Bumetanide | 11.0 | 11.6 |
| **140** | Neomycin Sulfate | 5.5 | 10.2 |
| **141** | Auranofin | 8.1 | 7.6 |
| **142** | Captopril | 7.4 | 10.4 |
| **143** | Tranylcypromine Hemisulfate | 13.2 | 10.8 |
| **144** | Piroxicam | 22.2 | 19.4 |
| **145** | Moxifloxacin·HCl | 8.8 | 11.6 |
| **146** | Carbidopa | 12.5 | 13.1 |
| **147** | Ketoprofen | 1.9 | 14.1 |
| **148** | Meloxicam | 21.2 | 23.4 |
| **149** | Terbinafine·HCl | 8.2 | 8.3 |
| **150** | Sodium Phenylbutyrate | 9.6 | 9.2 |
| **151** | Simvastatin | 6.5 | 10.5 |
| **152** | Goserelin Acetate | 0.4 | 4.8 |
| **510** | Dexrazoxane | 2.4 | 11.5 |
| **511** | Diatrizoate Meglumine | 4.2 | 13.0 |
| **512** | Diazepam | 4.8 | 7.4 |
| **513** | Dicloxacillin·Na Monohydrate | 5.1 | 7.1 |
| **514** | Dicyclomine·HCl | –19.1 | –16.5 |
| **515** | Dienogest | 8.7 | 12.4 |
| **516** | VEHICLE | 0.0 | –8.4 |
| **517** | Difluprednate | 3.0 | 1.6 |
| **518** | Digoxin | 10.0 | 8.6 |
| **519** | Dimenhydrinate | 5.6 | 3.9 |
| **520** | Disopyramide | 10.8 | 9.9 |
| **521** | Dopamine·HCl | 15.5 | 8.1 |
| **522** | Doripenem | 7.2 | 14.1 |
| **523** | Doxapram·HCl·$H_2O$ | 6.2 | 6.7 |
| **524** | Doxepin·HCl | 0.8 | –1.1 |
| **525** | Droperidol | 10.0 | 13.6 |
| **526** | Drospirenone | 13.8 | 11.5 |
| **527** | Duloxetine·HCl | 6.3 | 0.5 |
| **528** | Dutasteride | 4.3 | 2.8 |
| **529** | Dyphylline | 8.3 | 7.0 |
| **530** | Econazole Nitrate | –7.7 | –0.6 |
| **531** | VEHICLE | –5.6 | –5.2 |
| **532** | Eflornithine·HCl | 12.0 | 8.5 |
| **533** | Epinastine·HCl | 7.0 | 11.5 |
| **534** | Epirubicin·HCl | –4.2 | –0.8 |
| **535** | Eplerenone | 8.5 | 8.0 |
| **536** | Eptifibatide | 8.4 | 8.5 |
| **537** | Erythromycin | 8.5 | 5.9 |
| **538** | Estramustine Phosphate·Na | 6.4 | 12.9 |
| **539** | Estropipate | 9.4 | 8.8 |
| **540** | Eszopiclone | 6.3 | 4.4 |
| **541** | Ethambutol Dihydrochloride | 5.7 | 6.5 |
| **542** | VEHICLE | –5.3 | –3.9 |
| **543** | Ethinyl Estradiol | 3.6 | –0.5 |
| **544** | Ethionamide | 4.8 | 6.3 |
| **545** | Ethosuximide | 3.6 | 11.2 |
| **546** | Etodolac | 5.2 | 7.8 |
| **547** | Etomidate | 7.9 | 10.7 |
| **548** | Etonogestrel | 0.0 | 5.6 |
| **549** | Everolimus | 6.4 | 8.9 |
| **550** | Ezetimibe | –3.5 | 5.9 |
| **551** | Febuxostat | –1.2 | 5.7 |
| **552** | Fexofenadine·HCl | 3.9 | 5.3 |

| | | | | | | | |
|---|---|---|---|---|---|---|---|
| **153** | Raloxifene·HCl | 14.0 | 12.7 | **553** | Fingolimod | 2.2 | 5.0 |
| **154** | Rifampin (Rifampicin) | 8.9 | 6.6 | **554** | Flavoxate·HCl | 1.3 | 2.0 |
| **155** | Etoposide | 12.8 | 14.3 | **555** | Flucytosine | 4.3 | 8.2 |
| **156** | Mitomycin C | 10.2 | 10.4 | **556** | Fludarabine Phosphate | 8.3 | 10.6 |
| **157** | Delavirdine Mesylate | 7.4 | –2.5 | **557** | Fludrocortisone Acetate | 9.0 | 10.8 |
| **158** | Daunorubicin·HCl | 11.4 | 10.4 | **558** | Flunisolide | 3.9 | 8.3 |
| **159** | Doxorubicin·HCl | 8.4 | 15.3 | **559** | Fluocinonide | 9.9 | 9.6 |
| **160** | Cetirizine HCl | 1.4 | 4.3 | **560** | Fluorometholone | –0.7 | 8.5 |
| **161** | Lapatinib Ditosylate | 19.0 | 12.0 | **561** | Flurandrenolide | 4.0 | 6.9 |
| **162** | Pioglitazone·HCl | 1.6 | 10.8 | **562** | VEHICLE | –10.2 | –6.6 |
| **163** | Rivastigmine Tartrate | –8.5 | 9.2 | **563** | Fluticasone Propionate | 7.5 | 10.4 |
| **164** | Ergotamine Tartrate | 12.7 | 14.7 | **564** | Fluvoxamine Maleate | 3.1 | –0.2 |
| **165** | Sulindac | 10.1 | 11.2 | **565** | Fomepizole | 5.7 | 10.4 |
| **166** | Valproic Acid | 13.9 | 14.6 | **566** | Formoterol | 6.9 | 9.4 |
| **167** | Calcipotriene | –48.1 | 14.1 | **567** | Foscarnet·Na | 6.3 | 11.7 |
| **168** | Zafirlukast | 26.8 | 30.2 | **568** | Fosfomycin Calcium | 13.7 | 5.5 |
| **169** | Zileuton | 6.1 | –15.3 | **569** | Fosphenytoin·Na Pentahydrate | 4.9 | ND |
| **170** | Bortezomib | 10.4 | 7.5 | **570** | Gemifloxacin | 2.3 | 8.9 |
| **171** | Diazoxide | 10.8 | 17.1 | **571** | Glycopyrrolate Iodide | 1.8 | 8.2 |
| **172** | Glyburide | 4.7 | 14.9 | **572** | Griseofulvin | 2.8 | –18.1 |
| **173** | Minoxidil | 34.4 | –0.8 | **573** | Guanidine·HCl | 13.2 | 12.0 |
| **174** | Tolazamide | 10.5 | 8.7 | **574** | Halcinonide | 4.2 | 8.1 |
| **175** | Bexarotene | 7.5 | 12.6 | **575** | Halobetasol Propionate | 11.5 | 9.1 |
| **176** | Tranexamic Acid | 15.4 | 12.0 | **576** | Hexachlorophene | 41.8 | 39.6 |
| **177** | Celecoxib | –4.2 | –4.3 | **577** | Homatropine Methylbromide | 6.3 | 12.8 |
| **178** | Levetiracetam | 12.4 | 8.3 | **578** | Hydralazine·HCl | 5.9 | 6.8 |
| **179** | Letrozole | 9.2 | 12.2 | **579** | Hydrochlorothiazide | 10.3 | 7.7 |
| **180** | Anastrozole | 11.3 | 12.8 | **580** | Hydroflumethiazide | 7.8 | 6.5 |
| **181** | Bicalutamide | 9.5 | 4.3 | **581** | Hydroxocobalamin·HCl | 0.5 | 10.3 |
| **182** | Clindamycin Palmitate·HCl | 19.9 | 21.1 | **582** | Hydroxychloroquine Sulfate | 9.9 | 9.1 |
| **183** | Vorinostat | 9.8 | 10.5 | **583** | Hydroxyurea | 13.2 | 9.4 |
| **184** | Didanosine | 1.5 | 4.9 | **584** | Hydroxyzine Dihydrochloride | –0.1 | –2.1 |
| **185** | Dolasetron | –2.9 | 16.2 | **585** | Ibutilide Fumarate | 25.1 | 10.9 |
| **186** | Enalaprilat Maleate | 7.6 | –2.9 | **586** | Iloperidone | 32.6 | 27.1 |
| **187** | Fluvastatin·Na | 7.5 | 6.0 | **587** | Indinavir | 7.5 | 9.2 |
| **188** | Fosinopril·Na | 5.2 | 11.1 | **588** | Irbesartan | 26.1 | 29.1 |
| **189** | Gemcitabine·HCl | 11.3 | 6.4 | **589** | Irinotecan·HCl | 12.3 | 8.2 |
| **190** | Granisetron·HCl | 12.7 | 8.4 | **590** | Isocarboxazid | 7.6 | 11.5 |
| **191** | Oxaliplatin | 2.3 | 16.0 | **591** | Isosorbide Dinitrate | 7.3 | 12.0 |
| **192** | Atazanavir | 14.1 | 17.2 | **592** | Isotretinoin (13-Cis-Retinoic Acid) | 5.2 | 13.0 |
| **193** | Mycophenolate Mofetil | 8.3 | 7.1 | **593** | Isradipine | 2.7 | 4.1 |
| **194** | Clofarabine | –0.2 | 11.0 | **594** | Kanamycin Sulfate | 5.8 | 9.9 |

| | | | |
|---|---|---|---|
| **195** | Cabergoline | 13.8 | 15.1 |
| **196** | Ibandronate·Na Monohydrate | 7.2 | 11.1 |
| **197** | Imipenem | 12.6 | 10.5 |
| **198** | Lomustine | ND | 5.6 |
| **199** | Adapalene | 13.4 | 10.4 |
| **200** | Meropenem | 7.2 | 12.3 |
| **201** | Oseltamivir Phosphate | 6.0 | 11.7 |
| **202** | Pamidronate Disodium Pentahydrate (Pamidronic Acid) | 13.0 | 9.8 |
| **203** | Pramipexole Dihydrochloride Monohydrate | 12.4 | 3.7 |
| **204** | Triptorelin Acetate | 3.8 | 6.7 |
| **205** | Risredonic Acid | 13.0 | 12.5 |
| **206** | Rocuronium Bromide | 10.5 | 12.6 |
| **207** | Vinorelbine | 15.7 | 12.6 |
| **208** | Salmeterol | 5.9 | 5.1 |
| **209** | Vincristine Sulfate | 12.1 | 8.5 |
| **210** | Aspirin (Acetylsalicylic Acid) | 16.0 | 5.2 |
| **211** | Acyclovir (Acycloguanosine) Zovirax | 10.9 | 14.1 |
| **212** | Zidovudine (3'-Azido-3'-Deoxythymidine) | 10.4 | 13.9 |
| **213** | Allopurinol | 8.6 | 9.5 |
| **214** | Altretamine | 9.9 | 8.2 |
| **215** | Alendronate·Na Trihydrate | 13.3 | 12.1 |
| **216** | Albendazole | ND | 8.2 |
| **217** | Sumatriptan Succinate | 11.5 | 20.0 |
| **218** | Amifostine | 9.4 | 15.9 |
| **219** | 4-Aminosalicylic Acid | 7.3 | 7.2 |
| **220** | Mesalamine (5-Aminosalicylic Acid) | 14.2 | 13.1 |
| **221** | Ampicillin Trihydrate | 11.8 | –8.2 |
| **222** | (±)-Atenolol | 8.2 | 16.6 |
| **223** | Atracurium Besylate | 9.7 | 29.3 |
| **224** | Vinblastine Sulfate | 31.3 | 18.1 |
| **225** | Azithromycin | 15.7 | 15.9 |
| **226** | Aztreonam | 22.8 | 14.0 |
| **227** | Betamethasone | 11.4 | 13.7 |
| **228** | Bisacodyl | 12.2 | 7.6 |
| **229** | Buspirone·HCl | 14.3 | 12.3 |
| **595** | Ketorolac Tromethamine | 9.7 | 11.4 |
| **596** | Labetalol·HCl | 9.5 | 8.0 |
| **597** | Lacosamide | 8.3 | 8.3 |
| **598** | Lactulose | 6.4 | 11.9 |
| **599** | Lamivudine | 10.2 | 7.6 |
| **600** | Lansoprazole | 1.3 | 7.0 |
| **601** | Lenalidomide | 8.7 | 9.0 |
| **602** | Leucovorin Calcium Pentahydrate | 7.2 | 11.3 |
| **603** | Levalbuterol·HCl | 11.2 | 10.9 |
| **604** | Levobunolol·HCl | 8.9 | 10.1 |
| **605** | Levocarnitine | 8.4 | 8.2 |
| **606** | Levocetirizine Dihydrochloride | 5.1 | 1.2 |
| **607** | Levothyroxine·Na | 4.5 | 4.9 |
| **608** | Lindane | 14.0 | 13.5 |
| **609** | Liothyronine·Na | 12.1 | 6.9 |
| **610** | Lopinavir | 4.9 | 0.9 |
| **611** | Lorazepam | 14.0 | 9.3 |
| **612** | Loteprednol Etabonate | 14.4 | 6.9 |
| **613** | Loxapine Succinate | 2.6 | 1.2 |
| **614** | Mafenide·HCl | 7.5 | 7.5 |
| **615** | Malathion | ND | 9.8 |
| **616** | Mannitol | 7.7 | 6.4 |
| **617** | Maraviroc | 8.9 | 6.3 |
| **618** | Mechlorethamine·HCl | 10.9 | 8.6 |
| **619** | Meclizine Dihydrochloride | 3.4 | 4.2 |
| **620** | Meclofenamate·Na | 83.7 | 85.5 |
| **621** | Mefloquine·HCl | –6.6 | –8.7 |
| **622** | Mepenzolate Bromide | 0.8 | 0.7 |
| **623** | Mepivacaine·HCl | 6.0 | 9.2 |
| **624** | Meprobamate | 0.4 | 7.6 |
| **625** | Mequinol | 3.2 | 9.5 |
| **626** | Mercaptopurine Hydrate | 5.4 | 13.2 |
| **627** | Mesna | 14.7 | 10.3 |
| **628** | Mestranol | 11.8 | –1.6 |
| **629** | Metaproterenol Hemisulfate (Orciprenaline) | 7.7 | 6.6 |

| | | | | | | | |
|---|---|---|---|---|---|---|---|
| **230** | Carboplatin | 16.9 | 15.4 | **630** | Metaraminol Bitartrate | 5.7 | 5.9 |
| **231** | Carbamazepine | 14.0 | –17.5 | **631** | Metaxalone | 6.5 | 5.6 |
| **232** | Cefotaxime Acid | –1.8 | 12.8 | **632** | Methacholine Chloride | 6.6 | 12.2 |
| **233** | Ceftazidime | 2.6 | 9.8 | **633** | Methazolamide | 7.1 | 13.0 |
| **234** | Chloramphenicol | 6.1 | 15.1 | **634** | Methenamine Hippurate | 9.7 | 5.8 |
| **235** | Chlorambucil | 7.3 | 12.3 | **635** | Methocarbamol | 6.6 | 8.5 |
| **236** | Chlorpheniramine Maleate | 13.6 | 10.2 | **636** | Methotrexate | 12.0 | 32.6 |
| **237** | Chloroquine Diphosphate | –2.4 | 16.4 | **637** | Methoxsalen (Xanthotoxin) | 15.5 | 9.9 |
| **238** | Thalidomide | 11.1 | 11.0 | **638** | Methscopolamine Bromide ((−)-Scopolamine Methyl Bromide) | 12.5 | 10.4 |
| **239** | Ciprofloxacin | –13.5 | 12.7 | **639** | Methsuximide | 5.8 | 8.4 |
| **240** | Citalopram·HBr | 18.6 | 22.0 | **640** | Methyclothiazide | 16.2 | 16.3 |
| **241** | Clarithromycin | 8.1 | 13.9 | **641** | Methyl Aminolevulinate·HCl | 9.1 | 8.9 |
| **242** | Clomiphene Citrate | 3.8 | 3.2 | **642** | Methylergonovine Maleate | 12.7 | 10.6 |
| **243** | Clopidogrel Hydrogen Sulfate | 9.7 | 14.0 | **643** | Metolazone | 7.5 | 10.2 |
| **244** | Clobetasol Propionate | 27.3 | 23.5 | **644** | Metyrapone | 7.0 | 16.8 |
| **245** | Orphenadrine Citrate | 3.9 | 7.2 | **645** | VEHICLE | –6.3 | –5.4 |
| **246** | Crotamiton | 13.6 | 7.9 | **646** | Mexiletine·HCl | 11.4 | 19.7 |
| **247** | Cyclophosphamide (Free Base) | 11.6 | 0.9 | **647** | Micafungin | 26.7 | 27.9 |
| **248** | Cytarabine | 13.3 | 13.9 | **648** | Miconazole | –2.3 | –0.9 |
| **249** | Dacarbazine | 12.8 | 6.4 | **649** | VEHICLE | –3.8 | –1.1 |
| **250** | Danazol | 0.7 | 8.0 | **650** | Midodrine·HCl | 8.4 | 10.0 |
| **251** | Desloratadine | 9.2 | 8.4 | **651** | Miglitol | 14.7 | 9.8 |
| **252** | Dextromethorphan | 11.9 | 9.5 | **652** | Milnacipran·HCl | 10.2 | –4.1 |
| **253** | Diclofenac·Na Salt | 9.5 | 14.9 | **653** | Mirtazapine | 5.2 | 6.5 |
| **254** | Zalcitabine (2',3'-Dideoxycytidine) | 10.5 | 17.3 | **654** | Mitotane | 7.5 | 12.2 |
| **255** | Diflunisal | 24.7 | 27.0 | **655** | Modafinil | 9.1 | 11.5 |
| **256** | Disulfiram | 15.3 | 4.5 | **656** | Moexipril·HCl | 8.1 | 9.4 |
| **257** | Doxazosin Mesylate | 14.6 | 14.7 | **657** | Mometasone Furoate | 17.5 | 12.0 |
| **258** | Doxycycline Monohydrate | 7.0 | 7.4 | **658** | Mupirocin | 8.3 | 8.4 |
| **259** | Enalapril | 9.8 | 10.3 | **659** | Nadolol | 9.3 | 6.6 |
| **260** | Esomeprazole Potassium | 9.6 | 10.6 | **660** | Nafcillin·Na | 9.2 | 17.5 |
| **261** | Estradiol | 14.3 | 12.1 | **661** | Naftifine·HCl | 5.4 | 12.5 |
| **262** | Estrone | 13.9 | 11.7 | **662** | Naratriptan·HCl | 14.2 | 11.2 |
| **263** | Etidronate Disodium | 6.6 | 14.1 | **663** | Natamycin | 9.0 | 10.7 |
| **264** | Famciclovir | 11.8 | 12.9 | **664** | Nebivolol·HCl | –22.3 | 4.5 |
| **265** | Fenoldopam Mesylate | 8.9 | 13.6 | **665** | Nelarabine | 30.3 | 15.2 |
| **266** | Fenoprofen Calcium | 10.2 | ND | **666** | Nepafenac | 10.2 | 7.9 |
| **267** | Fenofibrate | 6.8 | 13.5 | **667** | Nevirapine | 9.9 | 11.5 |

| | | | |
|---|---|---|---|
| **268** | Finasteride | 1.2 | 5.1 |
| **269** | Fluorouracil (5-Fluorouracil) | 13.7 | 10.1 |
| **270** | Flurbiprofen | 9.3 | 10.7 |
| **271** | Amitriptyline·HCl | –27.8 | –5.2 |
| **272** | Floxuridine | 12.7 | 13.8 |
| **273** | Fluocinolone Acetonide | 12.1 | 3.6 |
| **274** | Flutamide | 1.7 | 15.3 |
| **275** | Fluconazole | 12.4 | 11.1 |
| **276** | Furosemide | 13.9 | 2.6 |
| **277** | Ganciclovir | 15.0 | 7.4 |
| **278** | Gatifloxacin | 6.6 | 8.5 |
| **279** | Gentamycin Sulfate | 12.9 | 10.2 |
| **280** | Gemfibrozil | 6.3 | 8.8 |
| **281** | Glimepiride | 11.3 | 6.6 |
| **282** | Hydrocortisone | 12.6 | 9.2 |
| **283** | Hydrocortisone Acetate | 9.1 | 11.7 |
| **284** | Idoxuridine | 10.8 | 9.6 |
| **285** | Ifosfamide | 6.3 | 9.4 |
| **286** | Imiquimod | 9.1 | 10.0 |
| **287** | Indapamide | 11.6 | 8.4 |
| **288** | Itraconazole | 10.4 | –0.2 |
| **289** | Levonorgestrel | 13.2 | 6.2 |
| **290** | Levofloxacin·HCl | 5.6 | 6.7 |
| **291** | Leflunomide | 3.8 | 4.2 |
| **292** | Lisinopril·2$H_2O$ | 11.0 | 11.5 |
| **293** | Loratadine | 6.4 | 3.7 |
| **294** | Losartan Potassium | 14.3 | 13.9 |
| **295** | Mebendazole | 9.6 | 6.4 |
| **296** | Medroxyprogester-one Acetate | 18.8 | 14.9 |
| **297** | Mefenamic Acid | ND | 12.9 |
| **298** | Melphalan | 9.3 | 4.9 |
| **299** | Methyldopa Sesquihydrate (L-A-Methyl-Dopa Sesquihyrate) | 10.8 | 9.3 |
| **300** | Methylprednisolone | 10.1 | 9.7 |
| **301** | Metoprolol Tartrate | 14.1 | 10.8 |
| **302** | Methimazole | 11.7 | 12.1 |
| **303** | Metronidazole | 15.7 | 9.8 |
| **304** | Minocycline | 10.3 | 11.2 |
| **305** | Mitoxantrone·HCl | 9.1 | 2.7 |
| **306** | Paclitaxel (Taxol) | 11.8 | 13.3 |
| **307** | Nabumetone | 11.0 | 3.0 |
| **668** | Niacin (Vitamin B3, Nicotinic Acid And Vitamin Pp) | 6.0 | 9.4 |
| **669** | Nicotine | 10.8 | 9.8 |
| **670** | Nilotinib | 6.4 | 10.7 |
| **671** | Nilutamide | 3.7 | 5.2 |
| **672** | Nitazoxanide | 23.3 | 9.2 |
| **673** | Nitisinone | 17.7 | 15.4 |
| **674** | Nitrofurantoin | 6.0 | 4.9 |
| **675** | Nizatidine | 11.9 | 15.1 |
| **676** | Nortriptyline·HCl | –3.7 | –18.1 |
| **677** | Olsalazine·Na | 10.3 | –0.9 |
| **678** | Orlistat (Tetrahydrolipstatin) | 10.0 | 7.3 |
| **679** | Oxaprozin | 5.4 | 12.3 |
| **680** | Oxazepam | 1.6 | 8.1 |
| **681** | Oxtriphylline | 14.6 | 16.1 |
| **682** | Oxybutynin Chloride | 8.3 | 9.0 |
| **683** | Oxytetracycline·HCl | 3.9 | 8.5 |
| **684** | Paliperidone | 9.5 | 10.7 |
| **685** | Palonosetron·HCl | 16.9 | 9.0 |
| **686** | Paromomycin Sulfate | 9.9 | 3.1 |
| **687** | Pazopanib·HCl | 37.4 | ND |
| **688** | Pemetrexed Disodium | 7.8 | 7.7 |
| **689** | Pemirolast Potassium | 8.8 | 12.1 |
| **690** | Penicillamine (D-Penicillamine) | –2.0 | 7.3 |
| **691** | Penicillin G Potassium (Benzylpenicillin) | 7.6 | 8.3 |
| **692** | Pentamidine Isethionate | –3.3 | –2.4 |
| **693** | Pentostatin | 9.8 | 10.5 |
| **694** | Perindopril Erbumine | 8.2 | –12.6 |
| **695** | Permethrin | 11.5 | 12.7 |
| **696** | Perphenazine | –9.3 | –8.7 |
| **697** | Phenelzine Sulfate | 6.4 | 14.4 |
| **698** | Phenylephrine | 13.1 | 11.1 |
| **699** | Phytonadione | 12.0 | 15.5 |
| **700** | Pimecrolimus | 3.6 | 13.6 |
| **701** | Pitavastatin Calcium | 14.0 | 10.8 |
| **702** | VEHICLE | –6.8 | –4.0 |
| **703** | Podofilox | 6.1 | 11.6 |
| **704** | Posaconazole | 8.4 | 10.5 |
| **705** | Pralidoxime Chloride | 14.6 | 14.1 |
| **706** | Prasugrel | 7.2 | 7.5 |
| **707** | Pravastatin·Na | 6.6 | 3.0 |

| | | | |
|---|---|---|---|
| **308** | Naphazoline·HCl | 4.8 | 7.1 |
| **309** | Nefazodone·HCl | 11.3 | –7.5 |
| **310** | Norethindrone | 11.2 | 7.4 |
| **311** | Norfloxacin | 6.8 | 12.0 |
| **312** | Nystatin | 18.3 | 18.6 |
| **313** | Ofloxacin | 5.0 | 12.2 |
| **314** | Omeprazole | 9.9 | –2.5 |
| **315** | Oxcarbazepine | 12.6 | 7.4 |
| **316** | Oxiconazole Nitrate | 4.8 | 0.4 |
| **317** | Oxacillin sodium salt monohydrate | 13.0 | 17.5 |
| **318** | Pantoprazole | 8.4 | 6.6 |
| **319** | Paroxetine·HCl | 6.5 | –20.9 |
| **320** | Penciclovir | 10.6 | 11.7 |
| **321** | Pentoxifylline | 5.7 | 4.9 |
| **322** | Penicillin V Potassium | –1.8 | 6.1 |
| **323** | Piperacillin | –1.1 | 5.5 |
| **324** | Prednisolone | 5.0 | 1.5 |
| **325** | Progesterone | 28.5 | 28.6 |
| **326** | Procarbazine·HCl | 8.4 | 6.3 |
| **327** | Prednisone | 6.6 | 8.3 |
| **328** | Primaquine Phosphate | ND | 2.3 |
| **329** | Praziquantel | 2.9 | 9.6 |
| **330** | Quinapril·HCl | 8.5 | 1.9 |
| **331** | Ranolazine·2HCl | 4.8 | 8.1 |
| **332** | Ramipril | 6.3 | –1.1 |
| **333** | Ribavirin | 5.1 | 9.3 |
| **334** | Nelfinavir Mesylate | 3.0 | 2.1 |
| **335** | Rimantadine·HCl | 3.4 | 3.3 |
| **336** | Propranolol·HCl | –0.9 | 2.3 |
| **337** | Scopolamine·HBr | –0.4 | 4.7 |
| **338** | Spironolactone | 24.3 | 19.2 |
| **339** | Streptomycin Sulfate | 6.9 | 7.4 |
| **340** | Sulfadiazine | 6.4 | 30.0 |
| **341** | Sulfasalazine | ND | 8.2 |
| **342** | Tamsulosin·HCl | ND | 9.5 |
| **343** | Telmisartan | 2.6 | 7.1 |
| **344** | Terazosin·HCl | 6.6 | –8.4 |
| **345** | Tetracycline | 5.7 | 6.1 |
| **346** | Temozolomide | 4.3 | 4.3 |
| **347** | Tinidazole | 9.2 | 7.2 |
| **348** | Tobramycin | 1.1 | 8.3 |
| **349** | Topotecan·HCl | 5.3 | 1.0 |
| **350** | Toremifene Base | –2.8 | –7.4 |
| **351** | Tolmetin sodium dihydrate | 15.0 | 12.1 |
| **352** | Amoxicillin | ND | 9.8 |
| **353** | Tramadol·HCl | –2.6 | –1.8 |
| **354** | Trimethoprim | 5.5 | 12.6 |
| **708** | Pregabalin | –5.5 | –6.9 |
| **709** | Prilocaine·HCl | 8.1 | 11.2 |
| **710** | Primidone | 8.8 | 14.6 |
| **711** | Probenecid | 11.5 | 8.2 |
| **712** | VEHICLE | –5.5 | –19.1 |
| **713** | Proparacaine·HCl | 7.5 | 11.6 |
| **714** | Propylthiouracil | 8.1 | 10.8 |
| **715** | Protriptyline·HCl | 7.6 | 15.9 |
| **716** | Pyrazinamide | 6.8 | 12.6 |
| **717** | Pyridostigmine Bromide | –0.3 | 7.7 |
| **718** | Pyrimethamine | –2.2 | 0.5 |
| **719** | Quinidine·HCl·$H_2O$ | –3.0 | 11.8 |
| **720** | Rabeprazole·Na | 12.1 | 5.3 |
| **721** | Raltegravir | 6.0 | 11.0 |
| **722** | Ramelteon | 7.7 | 10.4 |
| **723** | Rasagiline Mesylate | 8.6 | 8.5 |
| **724** | Regadenoson | 4.2 | 5.0 |
| **725** | Repaglinide | 5.9 | 14.8 |
| **726** | Reserpine | 8.3 | 10.6 |
| **727** | Rifabutin | 11.0 | –5.5 |
| **728** | Rifapentine | 0.4 | 7.4 |
| **729** | Rifaximin | 8.2 | ND |
| **730** | Ritonavir | –4.3 | 13.4 |
| **731** | Rizatriptan Benzoate | 10.7 | 12.5 |
| **732** | Ropinirole·HCl | 9.1 | 11.0 |
| **733** | Ropivacaine·HCl Monohydrate | 14.5 | 8.3 |
| **734** | Rosuvastatin Calcium | 9.4 | 7.2 |
| **735** | Rufinamide | 13.3 | 9.3 |
| **736** | Saquinavir Mesylate | 4.2 | 6.1 |
| **737** | Selegiline·HCl | 7.4 | 4.4 |
| **738** | Sertraline·HCl | –16.1 | –6.9 |
| **739** | Silver Sulfadiazine | 69.7 | 69.9 |
| **740** | Sitagliptin Phosphate | 8.7 | 7.6 |
| **741** | Sorafenib Tosylate | 11.6 | 20.7 |
| **742** | Stavudine | 11.2 | 10.0 |
| **743** | Streptozocin | 10.0 | 13.7 |
| **744** | Sulconazole Nitrate | –1.4 | –4.0 |
| **745** | Sulfacetamide·Na | 11.7 | 8.3 |
| **746** | Sulfamethoxazole | 9.5 | 4.5 |
| **747** | Sulfanilamide | 8.3 | 8.4 |
| **748** | Sunitinib Malate | 4.4 | 8.2 |
| **749** | Tacrolimus (Fk506) | 10.2 | 5.0 |
| **750** | Tadalafil | 8.2 | –4.5 |
| **751** | Tazarotene | 12.8 | 14.1 |
| **752** | Telbivudine | 11.1 | 2.4 |
| **753** | Telithromycin | 12.9 | 11.0 |
| **754** | Temazepam | 6.4 | 11.6 |

| | | | |
|---|---|---|---|
| **355** | Valacyclovir·HCl | 1.0 | 21.9 |
| **356** | Vecuronium Bromide | 4.0 | 13.4 |
| **357** | Venlafaxine·HCl | 2.6 | 8.0 |
| **358** | Bupivacaine·HCl | 9.4 | 2.1 |
| **359** | Ketotifen Fumarate | 2.9 | 2.6 |
| **360** | Naloxone·HCl | –12.5 | –12.7 |
| **361** | Fluoxetine·HCl | 1.7 | 4.1 |
| **362** | Ondansetron | 5.5 | 11.1 |
| **363** | Tiotropium Bromide | 16.2 | 6.6 |
| **364** | Thioridazine·HCl | –6.5 | –7.3 |
| **365** | Amrinone | 7.0 | 6.0 |
| **366** | Milrinone | 5.0 | 9.8 |
| **367** | Alprostadil | 5.9 | 5.5 |
| **368** | Misoprostol | 0.9 | 0.7 |
| **369** | Argatroban | 6.9 | 9.9 |
| **370** | Cilastatin·Na | 5.5 | 2.5 |
| **371** | Butoconazole Nitrate | 1.7 | –0.2 |
| **372** | Mifepristone | 3.8 | 7.0 |
| **373** | Megestrol Acetate | 15.8 | 16.7 |
| **374** | Tamoxifen Citrate | –11.2 | –11.3 |
| **375** | Aprepitant | –24.2 | –17.1 |
| **376** | Bosentan | 3.5 | 3.1 |
| **377** | Efavirenz | –7.5 | –2.6 |
| **378** | Miglustat (*N*-Butyldeoxynojirimycin·HCl) | 6.3 | 7.0 |
| **379** | Fulvestrant | 6.2 | 5.4 |
| **380** | Esmolol | 11.0 | 6.7 |
| **381** | Capecitabine | 8.2 | 3.6 |
| **382** | Succinylcholine Chloride·$2H_2O$ | 8.9 | 4.2 |
| **383** | Cyproheptadine·HCl Sesquihydrate | –6.2 | –7.2 |
| **384** | Abacavir Sulfate | 3.7 | 4.3 |
| **385** | Acamprosate | 8.1 | 4.2 |
| **386** | Acarbose | 4.0 | 10.1 |
| **387** | Acebutolol·HCl | 7.6 | 7.4 |
| **388** | Acetaminophen | 2.7 | 2.4 |
| **389** | Acetazolamide | 8.0 | 6.6 |
| **390** | Acetohexamide | 9.0 | 11.4 |
| **391** | Acetohydroxamic Acid | –0.3 | 5.3 |
| **392** | Acetylcysteine | 9.3 | 8.2 |
| **393** | Acrivastine | 8.1 | 2.9 |
| **394** | Adefovir Dipivoxil | 6.0 | 2.7 |
| **395** | Adenosine | 5.7 | 5.9 |
| **396** | VEHICLE | –7.7 | –9.7 |
| **397** | Alitretinoin | –4.5 | 8.7 |

| | | | |
|---|---|---|---|
| **755** | Temsirolimus | 8.7 | –0.3 |
| **756** | Teniposide | 1.0 | 8.9 |
| **757** | Tenofovir | 10.1 | 8.9 |
| **758** | Terbutaline Hemisulfate | 11.0 | 11.5 |
| **759** | Terconazole | 6.4 | 9.8 |
| **760** | Testosterone Enanthate | 15.0 | 5.5 |
| **761** | Tetrabenazine | 12.4 | 11.9 |
| **762** | Tetrahydrozoline·HCl | 1.7 | 10.9 |
| **763** | Theophylline | 8.5 | 12.3 |
| **764** | Thioguanine(6-Thioguanine) | 10.1 | 9.9 |
| **765** | Thiotepa | –14.8 | 4.8 |
| **766** | VEHICLE | –7.0 | –4.4 |
| **767** | Tiagabine·HCl | 6.7 | 5.8 |
| **768** | Tigecycline | 8.9 | 16.2 |
| **769** | Tiludronate Disodium | 5.0 | 7.3 |
| **770** | Tiopronin | –1.3 | 5.1 |
| **771** | Tirofiban·HCl | 12.0 | 1.0 |
| **772** | Tolterodine Tartrate | 3.3 | 5.6 |
| **773** | Tolvaptan | 2.4 | 7.7 |
| **774** | Topiramate | 10.7 | 12.4 |
| **775** | Torsemide | 5.4 | 1.8 |
| **776** | Trandolapril | 2.9 | 12.8 |
| **777** | Travoprost | 7.8 | 7.9 |
| **778** | Trazodone·HCl | 9.0 | 3.7 |
| **779** | Tretinoin | 11.5 | 7.3 |
| **780** | Triamcinolone Acetonide | 10.4 | 10.6 |
| **781** | Triamterene | 11.8 | 25.6 |
| **782** | Triazolam | 12.7 | 11.6 |
| **783** | Trientine Dihydrochloride | 9.4 | 2.8 |
| **784** | Trihexyphenidyl·HCl | 5.0 | 9.0 |
| **785** | Trimethadione | 9.1 | 10.3 |
| **786** | Trimethobenzamide·HCl | 7.2 | 5.7 |
| **787** | Trimipramine Maleate | 9.0 | 24.3 |
| **788** | Trospium Chloride | 6.4 | 11.7 |
| **789** | Ursodiol | 5.4 | 8.8 |
| **790** | Valganciclovir·HCl | –3.8 | 0.9 |
| **791** | Valproate·Na | –12.2 | –10.3 |
| **792** | Valsartan | 4.0 | 11.8 |
| **793** | Vancomycin·HCl | 10.3 | 9.2 |
| **794** | Varenicline Tartrate | –10.8 | –10.9 |
| **795** | Vigabatrin | 10.9 | 7.0 |
| **796** | Voriconazole | 11.6 | 10.3 |
| **797** | Warfarin·Na | 6.8 | 8.3 |

| | | | | | | | |
|---|---|---|---|---|---|---|---|
| **398** | Almotriptan | 6.1 | 1.1 | **798** | Zaleplon | 4.7 | 1.8 |
| **399** | Alosetron·HCl | 4.0 | 5.0 | **799** | Zanamivir | 7.4 | 5.2 |
| **400** | VEHICLE | –7.3 | –2.2 | **800** | Ziprasidone | 39.7 | 34.7 |

### *Dataset 2: PDSP screening results*

To examine the specificity of AK-42 for CLC-2 in the brain, this compound was screened against a panel of CNS receptors, transporters, and ion channels. The primary screen involved either a comprehensive binding assay (55 of 58 targets) or a functional assay (3 targets for which no binding assays were available). Results are summarized in **Dataset 2**, **Table 1**. Data for functional assays and for secondary binding assays are shown in **Dataset 2**, **Figure 1**.

**Dataset 2**, **Table 1.** Primary specificity-screening assay data. Compounds eliciting >50% mean effect at 10 μM (n = 4, highlighted in gray) were subjected to additional assays (raw data shown in **Dataset 2**, **Figure 1**). Entries marked with * denote targets that were tested via functional assays instead of binding assays.

| Entry | Receptor | Mean % inhibition (10 μM AK-42) |
|---|---|---|
| **1** | 5-HT1A | 53.99 |
| **2** | 5-HT1B | 49.89 |
| **3** | 5-HT1D | 44.06 |
| **4** | 5-HT1E | 10.43 |
| **5** | 5-HT2A | 18.91 |
| **6** | 5-HT2B | 45.16 |
| **7** | 5-HT2C | –13.39 |
| **8** | 5-HT3 | 28.75 |
| **9** | 5-HT5A | 28.19 |
| **10** | 5-HT6 | 18.68 |
| **11** | 5-HT7 | 17.51 |
| **12** | A2A | –8.70 |
| **13** | Alpha1A | 12.29 |
| **14** | Alpha1B | 4.90 |
| **15** | Alpha1D | –10.12 |
| **16** | Alpha2A | –2.55 |
| **17** | Alpha2B | –28.62 |
| **18** | Alpha2C | 3.41 |
| **19** | AMPA | 28.20 |
| **20** | Beta1 | 22.04 |
| **21** | Beta2 | 4.77 |
| **22** | Beta3 | 24.76 |
| **23** | BZP rat brain site | 29.32 |
| **24** | Calcium channel | 23.69 |
| **25** | D1 | –9.96 |
| **26** | D2 | 7.69 |
| **27** | D3 | –4.66 |
| **28** | D4 | –0.65 |
| **29** | D5 | 31.58 |

| | | |
|---|---|---|
| **30** | DAT | 9.23 |
| **31** | DOR | 16.59 |
| **32** | $GABA_A$ | –4.29 |
| **33** | H1 | 15.48 |
| **34** | H2 | 5.35 |
| **35** | H3 | –0.92 |
| **36** | H4 | 2.23 |
| **37** | HERG | –43.87 |
| **38** | KA | 36.60 |
| **39** | KOR | 4.11 |
| **40** | M1 | 11.29 |
| **41** | M2 | –8.25 |
| **42** | M3 | 54.64 |
| **43** | M4 | 56.16 |
| **44** | M5 | 136.55 |
| **45** | mGluR1* | -- |
| **46** | mGluR5* | -- |
| **47** | MOR | –0.18 |
| **48** | NET | 17.36 |
| **49** | NMDA | 24.18 |
| **50** | NOP | 21.14 |
| **51** | Oxytocin | –1.41 |
| **52** | PBR | 8.22 |
| **53** | SERT | 30.25 |
| **54** | Sigma 1 | 38.52 |
| **55** | Sigma 2 | –8.14 |
| **56** | V1A | 23.96 |
| **57** | V1B | 40.53 |
| **58** | Y2* | -- |

**Dataset 2, Figure 1**. Screen results from functional secondary binding assays (5-HT1A, HERG, M3, M4, and M5) and from functional assays (mGluR1, mGluR5, and Y2). Secondary binding assays were performed in a 96-well plate format in which 12 concentrations of the test compound, ranging from 0.1 nM to 10 μM (n = 3), were evaluated relative to the same concentrations of an appropriate known receptor agonist or antagonist as a positive control (n = 3). Competitive binding of each applied compound was measured as the remaining binding of a radiolabeled ligand appropriate to the receptor of interest in counts per million (CPM). Points represent the average CPM for each compound concentration ± SEM. The radioligands and control compounds for each assay are indicated in each plot. Each plot represents data from a given day. Assays that were repeated on subsequent days are shown as replicates in individual plots (HERG, M3, and M5) as a measure of reproducibility. Functional assays were performed by measuring changes in fluorescence upon $Ca^{2+}$ flux (mGluR1 and mGluR5) or transcriptional activation of a β-lactamase reporter construct (Y2, Tango GPCR assay) relative to a known chemical modulator. (See https://pdspdb.unc.edu/pdspWeb/ for detailed protocols.)

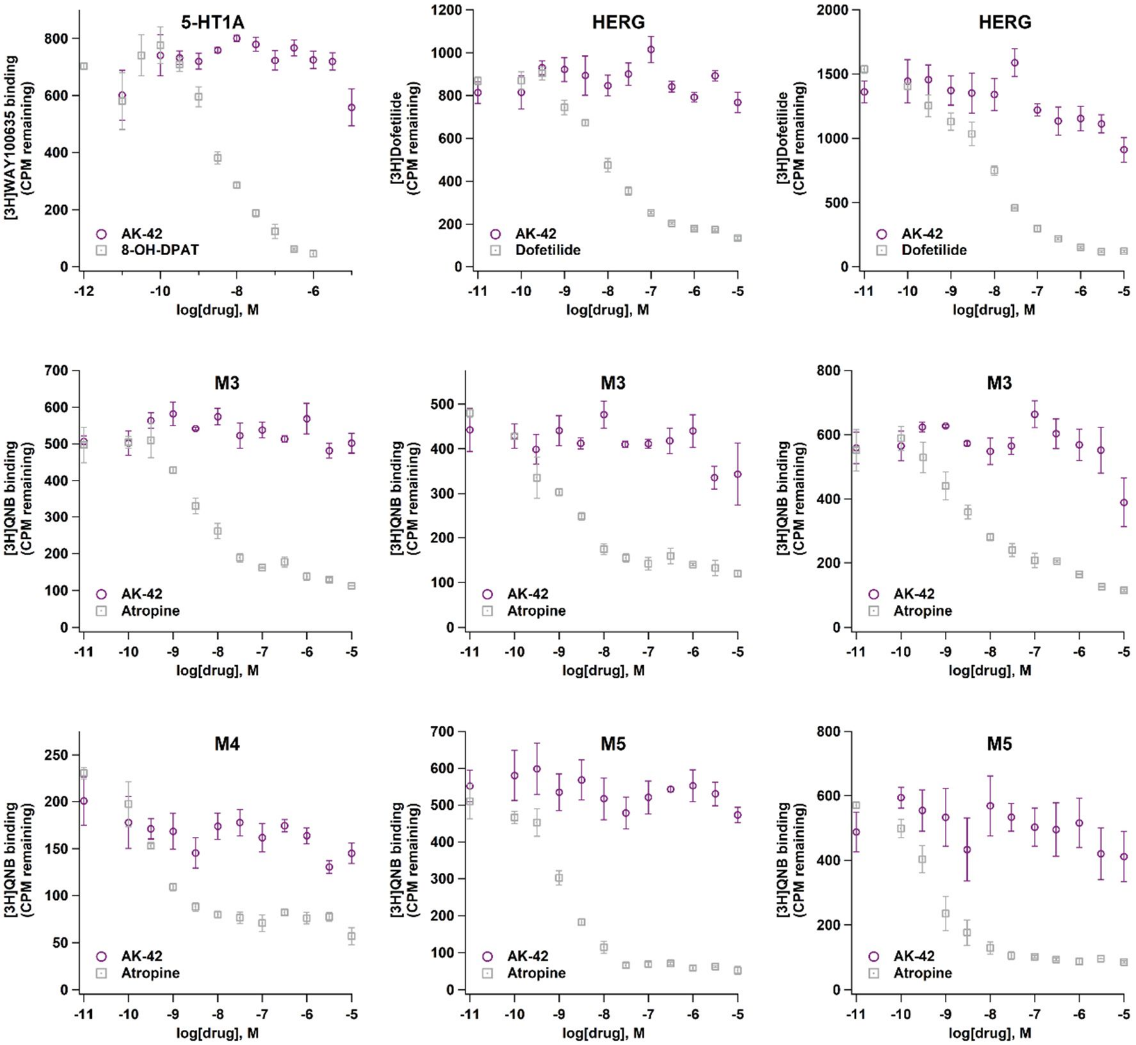

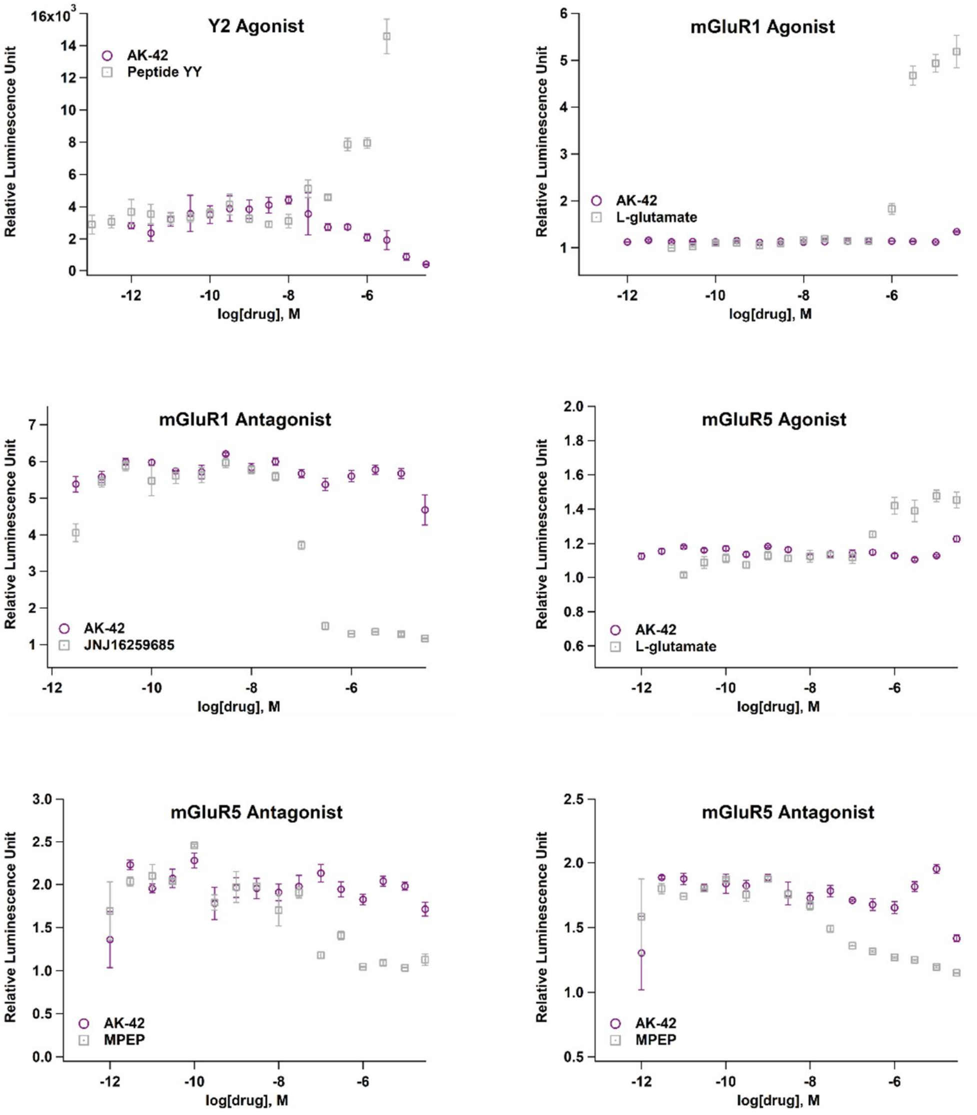
Y2 Agonist
AK-42
Peptide YY
Relative Luminescence Unit
log[drug], M
16x10^3
mGluR1 Agonist
AK-42
L-glutamate
Relative Luminescence Unit
log[drug], M
mGluR1 Antagonist
AK-42
JNJ16259685
Relative Luminescence Unit
log[drug], M
mGluR5 Agonist
AK-42
L-glutamate
Relative Luminescence Unit
log[drug], M
mGluR5 Antagonist
AK-42
MPEP
Relative Luminescence Unit
log[drug], M
mGluR5 Antagonist
AK-42
MPEP
Relative Luminescence Unit
log[drug], M

### *Dataset 3: Chemical Synthesis, General*

All reagents were obtained commercially unless otherwise noted. Meclofenamate sodium, *N*-phenylanthranilic acid, and lubiprostone were purchased commercially from Sigma-Aldrich. Diclofenac sodium, indomethacin, and niflumic acid were purchased from Santa Cruz Biotechnology. Salsalate was purchased from ACROS Organics. Aceclofenac was purchased from AK Scientific. Organic solutions were concentrated under reduced pressure (~20 Torr) by rotary evaporation. Air- and moisture-sensitive liquids and solutions were transferred via syringe or stainless steel cannula. Chromatographic purification of the desired carboxylate, phosphonate, and sulfate inhibitors was accomplished using high performance liquid chromatography on a C18 column (Alltima C18, 10 μM, 22 × 250 mm or SiliaChrom AQ C18, 5 μM, 10 × 250 mm). Thin layer chromatography was performed on EM Science silica gel 60 F254 plates (250 mm). Visualization of the developed chromatogram was accomplished by fluorescence quenching and by staining with aqueous potassium permanganate or aqueous ceric ammonium molybdate (CAM) solution.

Nuclear magnetic resonance (NMR) spectra were acquired on a Varian Inova spectrometer operating at 300, 400, 500, or 600 MHz for $^1$H spectra are referenced internally according to residual solvent signals. Data for $^1$H NMR are recorded as follows: chemical shift (δ, ppm), multiplicity (s, singlet; d, doublet; t, triplet; q, quartet; quintet; m, multiplet; br, broad), coupling constant (Hz), integration. Compound concentrations were determined by quantitative NMR in DMSO-$d_6$ using *N,N*-dimethylformamide as an internal standard. Infrared spectra were recorded as thin films using NaCl plates on a Thermo-Nicolet 300 FT-IR spectrometer and are reported in frequency of absorption. Low-resolution mass spectra were obtained from the Vincent Coates Foundation Mass Spectrometry Laboratory and the Stanford ChEM-H facility, using a Shimadzu 20-20 ESI mass spectrometer and a Phenomenex Synergi 4 μm Hydro-RP 80 Å reversed phase column (30 × 2 mm column, gradient flow 0:1→1:0 MeCN/$H_2O$ with 0.1% formic acid over 4 min). Microwave reactions were performed in a Biotage Initiator microwave reactor.

### *Dataset 3, Inhibitor Stock Solution Preparation and Quantification*

Meclofenamic acid (MCFA) derivatives were quantified by $^1$H NMR spectroscopy using distilled *N,N*-dimethylformamide (DMF) as an internal standard. Use of DMF as the internal standard allows for recovery of pure material following lyophilization. Each MCFA derivative was weighed into an Eppendorf tube, using a calibrated analytical balance (Mettler Toledo, Model XS105), and dissolved into DMSO-$d_6$ to a final concentration of 30–100 mM. To ensure complete dissolution, the sealed Eppendorf tube was inverted at least 5 times and then sonicated for ~60 seconds. Using a calibrated analytical balance, DMF (22 mg) was weighed into a scintillation vial, 3.0 mL DMSO-$d_6$ (stored in a desiccator jar and preferably newly opened to minimize water contamination in the solvent) was added to the vial via a p1000 micropipette, and the solution thoroughly mixed by inverting the capped vial at least 10 times (note: to prevent cross-contamination between samples and for convenience, disposable micropipette tips may be used without affecting the accuracy of the measurements). To ensure robustness of the quantification protocol, the 100 mM DMF stock solution was independently quantified in triplicate against stock solutions with known concentrations of fumaronitrile dissolved in DMSO-$d_6$. Fumaronitrile produces a sharp singlet that integrates to 2H at 7.03 ppm in DMSO-$d_6$ (DMSO solvent residual peak referenced to 2.50 ppm) and may be integrated against either of the DMF methyl signals that appear at 2.73 ppm (3H) and 2.89 (3H) ppm. The average of these three NMR measurements was used for calculation of the final DMF internal standard concentration. All measurements were performed at room temperature. Stock solutions were stored frozen, and left at ambient temperature for several hours to thaw and vortexed prior to quantification. A relaxation delay time (d1) of 20 s and an acquisition time (at) of 10 s were used during spectral acquisition. The number of scans (nt) was typically set to 32 unless a particular compound concentration was low (< 10 mM), such that more scans were required to improve signal/noise. The concentration of each MCFA derivative was determined by comparison with signal integrations for the MCFA derivative and the DMF internal standard.

### *Dataset 3: Chemical synthesis, Experimental protocols and characterization data*

*Meclofenamate derivatives.* A general two-step protocol (Buchwald-Hartwig cross-coupling,[1] followed by ester hydrolysis) was used to transform commercially available anilines into meclofenamate derivatives. Additional transformations were required to obtain several derivatives for which the requisite starting materials are not commercially available. Experimental details are provided for these compounds.

#### *Synthesis of C3-substituted 2,6-dichloronitrobenzenes*

*Methylation of 2,6-dichloro-3-nitrophenol*

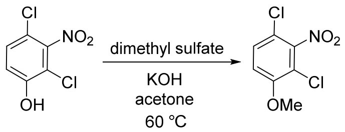

2,4-Dichloro-3-nitrophenol (0.5 g, 2.4 mmol) was dissolved in 5 mL of acetone, and the solution was cooled to 0 °C. Potassium hydroxide (146 mg, 2.6 mmol, 1.1 equiv) was added, followed by dropwise addition of dimethyl sulfate (656 mg, 5.2 mmol, 2.2 equiv). The mixture was warmed to room temperature and stirred for 3 h. The reaction was stirred for an additional 30 min at 60 °C until disappearance of the yellow color was observed. The reaction mixture was cooled to room temperature and diluted with 10 mL of EtOAc. The contents were transferred to a separatory funnel with an additional ~10 mL of EtOAc, and the combined filtrates were washed with 2 x 20 mL of $H_2O$ and 1 x 20 mL of saturated aqueous NaCl. The organic fraction was dried over $MgSO_4$, filtered and concentrated under reduced pressure. The oily residue was re-dissolved in $CH_2Cl_2$ to which ~1 g of silica gel was then added. The suspension was concentrated under reduced pressure and the solid material dry loaded onto a silica gel column pre-packed in pentane. Purification of this material by chromatography on silica gel (gradient elution: 0:1→1:19 $Et_2O$/pentane) furnished the desired product as a white solid (112 mg, 21%). TLC $R_f$ = 0.3 (9:1 hexanes/acetone); $^1$H NMR

1. Sadighi, J. P.; Harris, M. C.; Buchwald, S. L. A Highly Active Palladium Catalyst System for the Arylation of Anilines. *Tetrahedron Lett.* **1998**, *39* (30), 5327–5330.

($CDCl_3$, 300 MHz) δ 7.38 (d, $J$ = 9.1 Hz, 1H), 6.98 (d, $J$ = 9.1 Hz, 1H), 3.95 (s, 3H) ppm; IR (thin film) ν 1539, 1480, 1454, 1438, 1362, 1300, 1291 $cm^{-1}$.

#### *Triflation of 2,6-dichloro-3-nitrophenol*

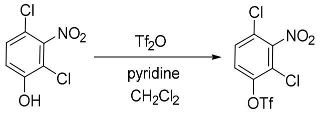

To a 250-mL flame-dried flask under $N_2$ was added 2,4-dichloro-3-nitrophenol (1.0 g, 4.8 mmol), 48 mL of $CH_2Cl_2$, and freshly distilled pyridine (0.46 g, 5.8 mmol, 1.2 equiv). The flask was placed in an ice bath, and triflic anhydride (1.36 g, 4.8 mmol) was add dropwise via syringe. The solution was stirred at 0 °C for 15 min, then warmed to room temperature and stirred for an additional 40 min. Following this time, the reaction mixture was poured into a separatory funnel and the organic layer was washed successively with 1 x 50 mL of saturated aqueous $NaHCO_3$, 1 x 50 mL of $H_2O$, and 1 x 50 mL of saturated aqueous NaCl. The organic fraction was dried over $Na_2SO_4$, filtered and concentrated under reduced pressure. The oily residue was re-dissolved in $CH_2Cl_2$ to which ~1 g of silica gel was then added. The suspension was concentrated under reduced pressure and the solid material dry loaded onto a silica gel column pre-packed in hexanes. Purification of this material by chromatography on silica gel (gradient elution: 0:1→1:3 acetone/hexanes) furnished the desired product as a white solid (1.43 g, 88%). TLC $R_f$ = 0.5 (9:1 hexanes/acetone); $^1H$ NMR ($CDCl_3$, 300 MHz) δ 7.57 (d, $J$ = 9.0 Hz, 1H), 7.49 (d, $J$ = 9.1 Hz, 1H) ppm; IR (thin film) ν 1556, 1456, 1436, 1359, 1221, 1135 $cm^{-1}$.

#### *Cross-coupling of aryl triflates and trifluoroborate salts*

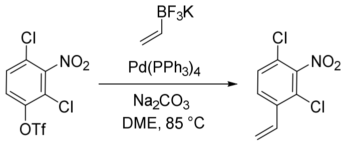

To a two-necked 25-mL flask was added 2,4-dichloro-3-nitrophenyl trifluoromethanesulfonate (300 mg, 0.88 mmol), potassium vinyltrifluoroborate (118 mg, 0.88 mmol), and $Na_2CO_3$ (281 mg, 2.6 mmol, 3.0 equiv). The flask was stoppered with a rubber septum and a reflux condenser, and flushed for ~5 min with argon. Dimethoxyethane (6.0 mL) was sparged with argon for 10 min, and then added via cannula to the flask containing the solid materials. To this mixture was added a solution of $Pd(PPh_3)_4$ (51 mg, 0.06 mmol, 0.05 equiv) in 2.5 mL of benzene under argon. The contents were stirred at 85 °C under argon for 6 h. Following this time, the reaction mixture was cooled to room temperature, diluted with 100 mL of $Et_2O$, and transferred to a separatory funnel. The organic phase was washed with 2 x 100 mL of $H_2O$ and 1 x 100 mL of saturated aqueous NaCl, dried over $Na_2SO_4$, filtered and concentrated under reduced pressure. The oily residue was re-dissolved in $CH_2Cl_2$ to which ~1 g of silica gel was then added. The suspension was concentrated under reduced pressure and the solid material dry loaded onto a silica gel column pre-packed in hexanes. Purification of this material by chromatography on silica gel (gradient elution: 0:1→1:19

acetone/hexanes) furnished the desired product as a white solid (115 mg, 60%). TLC $R_f$ = 0.33 (19:1 hexanes/acetone); $^1$H NMR ($CDCl_3$, 300 MHz) δ 7.62 (dd, *J* = 8.6, 0.6 Hz, 1H), 7.40 (dd, *J* = 8.6, 0.6 Hz, 1H), 7.01 (ddt, *J* = 17.5, 11.0, 0.6 Hz, 1H), 5.83 (dd, *J* = 17.4, 0.6 Hz, 1H), 5.57 (dd, *J* = 11.0, 0.6 Hz, 1H) ppm; IR (thin film) ν 1536, 1456, 1361 cm$^{-1}$.

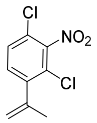

Prepared according to the above procedure using the appropriate tetrafluoroborate salt; colorless oil (30 mg, 44%). TLC $R_f$ = 0.40 (19:1 hexanes/acetone); $^1$H NMR ($CDCl_3$, 300 MHz) δ 7.39 (d, *J* = 8.4 Hz, 1H), 7.28 (d, *J* = 8.4 Hz, 1H), 5.34 (q, *J* = 1.5 Hz, 1H), 5.03 (t, *J* = 1.1 Hz, 1H), 2.08 (s, 3H) ppm; IR (thin film) ν 1548, 1460, 1362 cm$^{-1}$.

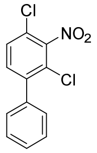

Prepared according to the above procedure using the appropriate tetrafluoroborate salt; white solid (78 mg, 33%). TLC $R_f$ = 0.36 (19:1 hexanes/EtOAc); $^1$H NMR ($CDCl_3$, 500 MHz) δ 7.51–7.45 (m, 4H), 7.43 (d, *J* = 8.4 Hz, 1H), 7.41–7.38 (m, 2H) ppm; IR (thin film) ν 3089, 3063, 3031, 2892, 1539, 1447, 1456, 1359, 1253 cm$^{-1}$.

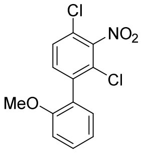

Prepared according to the above procedure using the appropriate tetrafluoroborate salt; colorless film (99 mg, 20%). TLC $R_f$ = 0.24 (19:1 hexanes/EtOAc); $^1$H NMR ($CDCl_3$, 500 MHz) δ 7.45 (d, *J* = 8.4 Hz, 1H), 7.44 (td, *J* = 7.5, 1.8 Hz, 1H), 7.37 (d, *J* = 8.4 Hz, 1H), 7.16 (dd, *J* = 7.5, 1.8 Hz, 1H), 7.05 (td, *J* = 7.5, 1.0 Hz, 1H), 7.00 (d, *J* = 8.4 Hz, 1H), 3.79 (s, 3H) ppm; IR (thin film) ν 2934, 2838, 1583, 1548, 1498, 1464, 1436, 1360, 1275, 1241 cm$^{-1}$.

*Cross-coupling of aryl triflates and vinyl boronic acids*

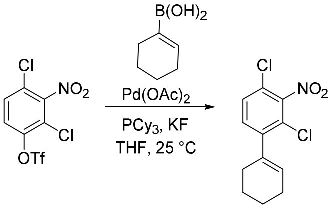

The reaction was performed following a general protocol described by Littke, et al.[2] To a 5-mL oven-dried round-bottom flask was added 2,4-dichloro-3-nitrophenyl trifluoromethanesulfonate (300 mg, 0.88 mmol), cyclohex-1-en-1-yl boronic acid (122 mg, 0.97 mmol, 1.1 equiv), anhydrous potassium fluoride (169 mg, 2.9 mmol, 3.3 equiv), and $Pd(OAc)_2$ (9.9 mg, 0.044 mmol, 0.05 equiv). The flask was stoppered with a rubber septum and flushed with argon for 5 min. In a glovebox, tricyclohexylphosphine (15 mg, 0.053 mmol, 0.06 equiv) was transferred to a round-bottom flask and sealed with a new rubber septum. The flask was removed from the glovebox, and 2.0 mL of dry, argon-purged THF was added. The phosphine solution was sparged with argon for 5 min and then added via cannula to the flask containing the starting materials and Pd catalyst. The reaction was stirred under argon for 8 h. Following this time, the reaction mixture was diluted with 5 mL of $Et_2O$ and filtered through a plug of silica gel. The flask and filter cake were rinsed with 3 x 5 mL of $Et_2O$. The combined filtrates were concentrated under reduced pressure to an oily residue. Purification of this material by chromatography on silica gel (gradient elution: 0:1→1:19 EtOAc/hexanes) furnished the desired product as a white solid (136 mg, 57%). TLC $R_f$ = 0.53 (19:1 hexanes/acetone); $^1H$ NMR ($CDCl_3$, 500 MHz) δ 7.35 (d, $J$ = 8.3 Hz, 1H), 7.23 (d, $J$ = 8.3 Hz, 1H), 5.73 (tt, $J$ = 3.7, 1.8 Hz, 1H), 2.25–2.22 (m, 2H), 2.21–2.16 (m, 2H), 1.79–1.72 (m, 2H), 1.71–1.66 (m, 2H) ppm; IR (thin film) ν 2933, 2859, 2836, 1549, 1456, 1362, 1256, 1188, 1137 $cm^{-1}$.

*Alkylation of 2,6-dichloro-3-nitrophenol*

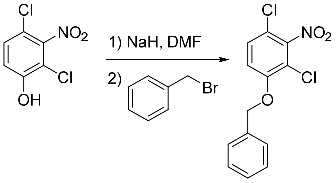

To an ice-cold suspension of sodium hydride (35 mg, 1.44 mmol, 1.2 equiv) in 1.0 mL of anhydrous DMF was added via syringe a solution of 2,4-dichloro-3-nitrophenol (250 mg, 1.20 mmol) in 1.0 mL DMF. After evolution of $H_2$ ceased (~30 min), neat benzyl bromide (206 mg, 1.20 mmol) was added via syringe. The reaction mixture was warmed to room temperature and stirred for 1.5 h. During this time, the color of the reaction changed from bright red-orange to pale yellow (Note: reaction times vary according to the reactivity of the electrophile and require up to 24 h at ambient temperature until the yellow color is largely

2. Littke, A. F.; Dai, C.; Fu, G. C. Versatile Catalysts for the Suzuki Cross-Coupling of Arylboronic Acids with Aryl and Vinyl Halides and Triflates under Mild Conditions. *J. Am. Chem. Soc.* **2000**, *122*, 4020–4028.

extinguished. Certain reactions, particularly those using mesylate-base electrophiles, require elevated temperatures, as indicated below). Following completion, the contents were transferred to a separatory funnel with 50 mL of $Et_2O$. The organic layer was washed with 3 x 50 mL of $H_2O$ and 2 x 50 mL of 1.0 M aqueous NaOH, dried over $Na_2SO_4$, filtered and concentrated under reduced pressure to a white solid. Purification of this material by recrystallization from ~25 mL of hot/cold hexanes afforded the desired product as white crystals (146 mg, 41%). TLC $R_f$ = 0.40 (3:1 hexanes/EtOAc); $^1$H NMR ($CDCl_3$, 500 MHz) δ 7.46–7.34 (m, 5H), 7.32 (d, *J* = 9.0 Hz, 1H), 7.00 (d, *J* = 9.0 Hz, 1H), 5.21 (s, 2H) ppm; IR (thin film) ν 1541, 1479, 1449, 1364, 1298 $cm^{-1}$.

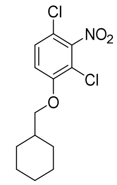

Prepared according to the above procedure with 2,4-dichloro-3-nitrophenol (250 mg, 1.20 mmol) and freshly prepared cyclohexylmethyl methanesulfonate (231 mg, 1.20 mmol). After stirring for 1.5 h at room temperature, the reaction flask was heated to 120 °C in an oil bath and stirred for an additional 4 h. Product obtained as an off-white solid (141 mg, 39%). TLC $R_f$ = 0.56 (3:1 hexanes/EtOAc); $^1$H NMR ($CDCl_3$, 500 MHz) δ 7.33 (d, *J* = 9.0 Hz, 1H), 6.94 (d, *J* = 9.1 Hz, 1H), 3.84 (d, *J* = 5.9 Hz, 2H), 1.91–1.81 (m, 3H), 1.81–1.75 (m, 2H), 1.75–1.68 (m, 1H), 1.36–1.26 (m, 2H), 1.21 (tt, *J* = 12.7, 3.2 Hz, 1H), 1.13–1.03 (m, 2H) ppm; IR (thin film) ν 2925, 2852, 1544, 1478, 1459, 1360, 1293, 1267 $cm^{-1}$.

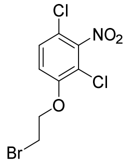

Prepared according to the above procedure with 2,4-dichloro-3-nitrophenol (1.5 g, 7.21 mmol) and 1,2-dibromoethane (208 mg, 8.65 mmol, 1.2 equiv). Following the addition of 1,2-dibromoethane, the mixture was stirred at 110 °C for 6 h. Product was obtained as a white solid (662 mg, 32%). TLC $R_f$ = 0.20 (3:1 hexanes/EtOAc); $^1$H NMR ($CDCl_3$, 400 MHz) δ 7.38 (d, *J* = 9.0 Hz, 1H), 6.99 (d, *J* = 9.0 Hz, 1H), 4.39 (t, *J* = 6.2 Hz, 2H), 3.69 (t, *J* = 6.2 Hz, 2H) ppm; IR (thin film) ν 1537, 1364, 1296 $cm^{-1}$.

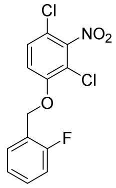

Prepared according to the above procedure with 2,4-dichloro-3-nitrophenol (250 mg, 1.2 mmol) and 2-fluorobenzyl bromide (227 mg, 1.2 mmol). The product was isolated as a white solid (294 mg, 77%). TLC $R_f$ = 0.22 (19:1 hexanes/EtOAc); $^1$H NMR ($CDCl_3$, 500 MHz) δ 7.51 (td, *J* = 7.5, 1.7 Hz, 1H), 7.39–7.34 (m,

2H), 7.20 (td, $J$ = 7.6, 1.1 Hz, 1H), 7.11 (ddd, $J$ = 9.7, 8.2, 1.1 Hz, 1H), 7.06 (d, $J$ = 9.1 Hz, 1H), 5.26 (s, 2H) ppm; IR (thin film) ν 3093, 2911, 1585, 1542, 1492, 1478, 1455, 1364, 1303, 1274, 1234 cm$^{-1}$.

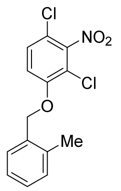

Prepared according to the above procedure with 2,4-dichloro-3-nitrophenol (250 mg, 1.2 mmol) and 2-methylbenzyl bromide (266 mg, 1.44 mmol, 1.2 equiv). The product was isolated as an off-white solid (293 mg, 78%). TLC $R_f$ = 0.18 (9:1 hexanes/EtOAc); $^1$H NMR (CDCl$_3$, 400 MHz) δ 7.38 (d, $J$ = 7.2 Hz, 1H), 7.35 (dd, $J$ = 9.0, 1.1 Hz, 1H), 7.32–7.27 (m, 1H), 7.25–7.21 (m, 2H), 7.05 (d, $J$ = 9.0 Hz, 1H), 5.16 (s, 2H), 2.39 (s, 3H) ppm; IR (thin film) ν 2919, 1592, 1542, 1495, 1479, 1451, 1364, 1300, 1285, 1272, 1194, 1100 cm$^{-1}$.

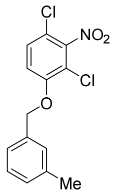

Prepared according to the above procedure with 2,4-dichloro-3-nitrophenol (250 mg, 1.2 mmol) and 3-methylbenzyl bromide (222 mg, 1.2 mmol). The product was isolated as a white solid (239 mg, 64%). TLC $R_f$ = 0.26 (19:1 hexanes/EtOAc); $^1$H NMR (CDCl$_3$, 500 MHz) δ 7.31 (d, $J$ = 9.1, 1H), 7.29 (t, $J$ = 7.4, 1H), 7.23–7.16 (m, 3H), 7.00 (d, $J$ = 9.1 Hz, 1H), 5.17 (s, 2H), 2.38 (s, 3H) ppm; IR (thin film) ν 3101, 3043, 2916, 1592, 1539, 1479, 1450, 1363, 1302, 1273, 1100 cm$^{-1}$.

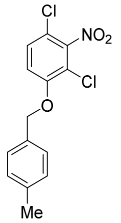

Prepared according to the above procedure with 2,4-dichloro-3-nitrophenol (312 mg, 1.5 mmol) and 4-methylbenzyl bromide (333 mg, 1.8 mmol, 1.2 equiv). The product was isolated as a white solid (342 mg, 73%). TLC $R_f$ = 0.24 (9:1 hexanes/EtOAc); $^1$H NMR (CDCl$_3$, 400 MHz) δ 7.34–7.28 (m, 3H), 7.21 (d, $J$ = 7.8 Hz, 2H), 7.00 (d, $J$ = 9.2 Hz, 1H), 5.16 (s, 2H), 2.37 (s, 3H) ppm; IR (thin film) ν 3094, 2908, 2867, 1866, 1594, 1537, 1482, 1455, 1363, 1312, 1299, 1274 cm$^{-1}$.

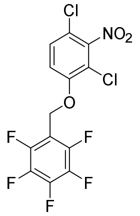

Prepared according to the above procedure with 2,4-dichloro-3-nitrophenol (312 mg, 1.5 mmol) and freshly prepared (perfluorophenyl)methyl methanesulfonate (425 mg, 1.54 mmol, 1.03 equiv). The product was isolated as a pale yellow solid (425 mg, 73%). TLC $R_f$ = 0.30 (4:1 hexanes/EtOAc); $^1$H NMR ($CDCl_3$, 400 MHz) δ 7.43 (d, *J* = 9.0 Hz, 1H), 7.14 (d, *J* = 9.0 Hz, 1H), 5.22 (s, 2H) ppm; $^{19}$F NMR ($CDCl_3$, 376 MHz) δ –142.18 (dd, *J* = 22.1, 8.2 Hz, 2F), –151.06 (t, *J* = 20.8 Hz, 1F), –160.92 (td, *J* = 20.6, 7.0 Hz, 2F) ppm; IR (thin film) ν 1658, 1590, 1549, 1525, 1508, 1465, 1435, 1389, 1362, 1313, 1288, 1264, 1195, 1135, 1104 $cm^{-1}$.

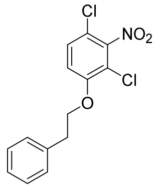

Prepared according to the above procedure with 2,4-dichloro-3-nitrophenol (250 mg, 1.2 mmol) and (2-bromoethyl)benzene (266 mg, 1.44 mmol, 1.2 equiv). The product was isolated as a white solid (169 mg, 45%). TLC $R_f$ = 0.16 (9:1 hexanes/EtOAc); $^1$H NMR ($CDCl_3$, 500 MHz) δ 7.37–7.25 (m, 6H), 6.92 (d, *J* = 9.1 Hz, 1H), 4.24 (t, *J* = 6.8 Hz, 2H), 3.17 (t, *J* = 6.7 Hz, 2H) ppm; IR (thin film) ν 3084, 3031, 2955, 2884, 1889, 1603, 1584, 1541, 1496, 1478, 1457, 1431, 1389, 1367, 1293, 1269, 1193, 1157 $cm^{-1}$.

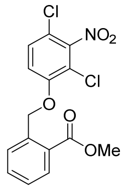

Prepared according to the above procedure with 2,4-dichloro-3-nitrophenol (1.24 g, 6.0 mmol) and methyl 2-(bromomethyl)benzoate (1.50 g, 6.6 mmol, 1.1 equiv). The product was isolated as a white solid (1.71 g, 81%). TLC $R_f$ = 0.5 (3:1 hexanes/EtOAc); $^1$H NMR ($CDCl_3$, 500 MHz) δ 8.08 (dd, *J* = 7.8, 1.3 Hz, 1H), 7.80 (d, *J* = 7.9 Hz, 1H), 7.62 (td, *J* = 7.8, 1.1 Hz 1H), 7.44 (t, *J* = 7.8 Hz, 1H), 7.35 (dd, *J* = 9.0, 0.7 Hz, 1H), 7.09 (d, *J* = 9.0, 1H), 5.63 (s, 2H), 3.92 (s, 3H) ppm; IR (thin film) ν 1722, 1543, 1475, 1432, 1365, 1302, 1267, 1143 $cm^{-1}$.

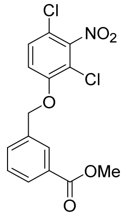

Prepared according to the above procedure with 2,4-dichloro-3-nitrophenol (1.24 g, 6.0 mmol) and methyl 3-(bromomethyl)benzoate (1.50 g, 6.6 mmol, 1.1 equiv). The product was isolated as an off-white solid (1.68 g, 79%). TLC $R_f$ = 0.39 (3:1 hexanes/EtOAc); $^1$H NMR ($CDCl_3$, 500 MHz) δ 8.09 (s, 1H), 8.04 (d, *J* = 7.8 Hz, 1H), 7.65 (d, *J* = 7.7 Hz, 1H), 7.51 (t, *J* = 7.7 Hz, 1H), 7.34 (d, *J* = 9.0 Hz, 1H), 7.01 (d, *J* = 9.0 Hz, 1H), 5.23 (s, 2H), 3.94 (s, 3H) ppm; IR (thin film) ν 1717, 1544, 1476, 1362, 1306, 1102 cm$^{-1}$.

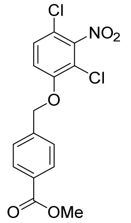

Prepared according to the above procedure with 2,4-dichloro-3-nitrophenol (1.24 g, 6.0 mmol) and methyl 4-(bromomethyl)benzoate (1.50 g, 6.6 mmol, 1.1 equiv). The product was isolated as a white solid (1.75 g, 83%). TLC $R_f$ = 0.32 (3:1 hexanes/EtOAc); $^1$H NMR ($CDCl_3$, 500 MHz) δ 8.08 (d, *J* = 8.0 Hz, 2H), 7.50 (d, *J* = 8.0 Hz, 2H), 7.33 (d, *J* = 9.0 Hz, 1H), 6.98 (d, *J* = 9.0 Hz, 1H), 5.25 (s, 2H), 3.93 (s, 3H) ppm; IR (thin film) ν 1705, 1551, 1480, 1449, 1309, 1287, 1120 cm$^{-1}$.

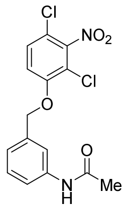

Prepared according to the above procedure with 2,4-dichloro-3-nitrophenol (194 mg, 0.93 mmol) and freshly prepared 3-acetamidobenzyl methanesulfonate (250 mg, 1.20 mmol, 1.3 equiv). The product was isolated as a tan solid (270 mg, 82%). TLC $R_f$ = 0.43 (19:1 $CH_2Cl_2$/MeOH); $^1$H NMR (acetone-$d_6$, 500 MHz) δ 9.27 (s, 1H), 7.84 (d, *J* = 2.8 Hz, 1H), 7.66–7.56 (m, 2H), 7.47 (d, *J* = 9.0 Hz, 1H), 7.35–7.27 (m, 1H), 7.20 (d, *J* = 7.6 Hz, 1H), 5.33 (s, 2H), 2.08 (s, 3H) ppm; IR (thin film) ν 3303, 3088, 1669, 1615, 1596, 1545, 1476, 1364, 1295, 1267, 1196 cm$^{-1}$.

*Phthalimide synthesis*

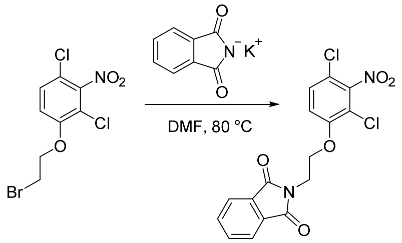

To a solution of 1-(2-bromoethoxy)-2,4-dichloro-3-nitrobenzene (250 mg, 0.79 mmol) in 1.6 mL of DMF was added potassium phthalimide (176 mg, 0.95 mmol, 1.2 equiv). The reaction was stirred at 80 °C for 7.5 h. Following this time, the reaction mixture was transferred to a separatory funnel with 100 mL of EtOAc. The organic layer was washed with 1 x 100 mL of $H_2O$, 2 x 100 mL of saturated aqueous $Na_2CO_3$, and 2 x 100 mL of saturated aqueous NaCl (2 x 100 mL), dried over $Na_2SO_4$, filtered and concentrated under reduced pressure to a white solid. Purification of this material by recrystallization from ~10 mL of hot/cold acetone afforded the desired product as a white solid (173 mg, 57%). TLC $R_f$ = 0.44 (3:2 hexanes/EtOAc); $^1$H NMR ($CDCl_3$, 500 MHz) δ 7.88 (dd, *J* = 5.4, 3.0 Hz, 2H), 7.75 (dd, *J* = 5.5, 3.0 Hz, 2H), 7.34 (d, *J* = 9.1 Hz, 1H), 6.98 (d, *J* = 9.1 Hz, 1H), 4.33 (t, *J* = 5.7 Hz, 2H), 4.18 (t, *J* = 5.7 Hz, 2H) ppm; IR (thin film) ν 1714, 1544, 1467, 1394, 1365, 1295, 1268 cm$^{-1}$.

*Ester hydrolysis*

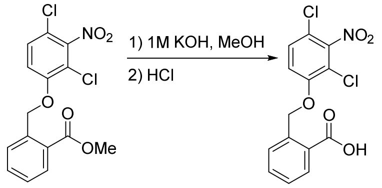

A screw-capped vial was charged with methyl 2-((2,4-dichloro-3-nitrophenoxy)methyl)benzoate (500 mg, 1.40 mmol) and 20 mL of a 1.0 M methanolic solution of KOH. The mixture was stirred for 24 h then warmed to 50 °C and stirred for an additional 2 h. During this time the opaque mixture transitioned to a clear yellow solution. The solution was cooled to ambient temperature and acidified to pH 1 by dropwise addition of 5 mL of concentrated aqueous HCl. Acidification of the reaction mixture resulted in precipitation of a white solid. Following the addition of 50 mL of $H_2O$, the desired product was isolated by vacuum filtration through a Büchner funnel as a white solid (405 mg, 84%). TLC $R_f$ = 0.28 (19:1 $CH_2Cl_2$/MeOH); $^1$H NMR (acetone-$d_6$, 400 MHz) δ 8.12 (dd, *J* = 7.8, 1.5 Hz, 1H), 7.81 (d, *J* = 7.8, 1H), 7.73–7.63 (m, 2H), 7.54 (t, *J* = 7.5, 1H), 7.47 (d, *J* = 9.0, 1H), 5.74 (s, 2H) ppm; IR (thin film) ν 2851, 1686, 1541, 1475, 1415, 1363, 1298, 1271, 1198 cm$^{-1}$.

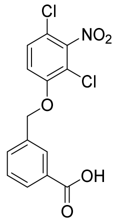

Prepared according to the above procedure; white solid (436 mg, 91%). TLC $R_f$ = 0.28 (19:1 $CH_2Cl_2$/MeOH); $^1$H NMR (acetone-$d_6$, 400 MHz) δ 8.20 (s, 1H), 8.04 (d, *J* = 7.8 Hz, 1H), 7.82 (d, *J* = 7.8 Hz, 1H), 7.69 (d, *J* = 9.2 Hz, 1H), 7.62–7.49 (m, 2H), 5.48 (s, 2H) ppm; IR (thin film) ν 2824, 1567, 1680, 1544, 1480, 1365, 1297, 1210 cm$^{-1}$.

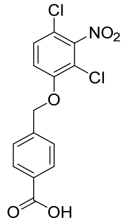

Prepared according to the above procedure; white solid (410 mg, 86%). TLC $R_f$ = 0.28 (19:1 $CH_2Cl_2$/MeOH); $^1$H NMR (acetone-$d_6$, 400 MHz) δ 8.10 (d, *J* = 8.4 Hz, 2H), 7.68 (d, *J* = 9.2 Hz, 3H), 7.53 (d, *J* = 9.2 Hz, 1H), 5.50 (s, 2H) ppm; IR (thin film) ν 2900, 2551, 1687, 1542, 1478, 1447, 1426, 1363, 1297, 1263 cm$^{-1}$.

#### *MOM protection of 2,6-dichloro-3-nitrophenol*

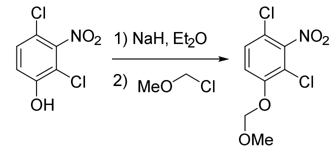

To an ice-cold suspension of NaH (70 mg, 2.92 mmol, 1.2 equiv) in 3.2 mL of anhydrous $Et_2O$ was added 2,4-dichloro-3-nitrophenol (505 mg, 2.43 mmol) in 1.0 mL of DMF under $N_2$. After evolution of $H_2$ ceased (approximately 30 min), chloromethyl methyl ether (196 mg, 1.0 mmol) was added dropwise via syringe. The reaction was warmed to ambient temperature and stirred for 1 h. Following this time, the reaction mixture was quenched by the addition of 1 mL of $H_2O$ and transferred to a separatory funnel with 50 mL of $Et_2O$. The organic layer was washed with 2 x 50 mL of $H_2O$ and 1 x 50 mL of saturated aqueous NaCl, dried over $MgSO_4$, filtered and concentrated under reduce pressure to a white solid (588 mg, 95%). This material was judged to be sufficiently pure by $^1$H NMR for use in the subsequent step without further purification. TLC $R_f$ = 0.25 (19:1 hexanes/EtOAc); $^1$H NMR ($CDCl_3$, 300 MHz) 7.35 (d, *J* = 9.1 Hz, 1H) 7.26 (d, *J* = 9.1 Hz, 1H) 5.28 (s, 2H) 3.51 (s, 3H) ppm; IR (thin film) ν 3092, 2971, 2948, 2909, 1589, 1545, 1465, 1367, 1265, 1152 cm$^{-1}$.

*Synthesis of primary and secondary amides*

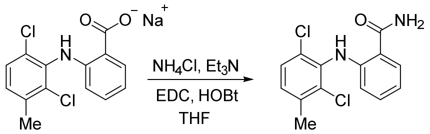

**AK-36**. The reaction was performed following a general protocol described by Suzuki, et al.[3] To a suspension of meclofenamic sodium acetate (75 mg, 0.24 mmol), $Et_3N$ (0.33 mL, 2.4 mmol, 10 equiv), and $NH_4Cl$ (126 mg, 2.4 mmol, 10 equiv) in 2.0 mL of THF were added successively 1-ethyl-3(3-dimethylamino)carbodiimide (131 mg, 0.68 mmol, 2.9 equiv) and 1-hydroxybenzotriazole monohydrate (112 mg, 0.73 mmol, 3.1 equiv). The reaction mixture was stirred for 23 h, then poured into a separatory funnel containing 50 mL of $H_2O$. The aqueous layer was extracted with 1 x 50 mL of EtOAc. The organic fraction was washed with saturated aqueous 1 x 50 mL of $NaHCO_3$ and 1 x 50 mL saturated aqueous NaCl, dried over $Na_2SO_4$, filtered and concentrated under reduced pressure to a white solid. Purification of this material by chromatography on silica gel (1:1 hexanes/EtOAc) furnished the desired product as a white solid (53 mg, 76%). TLC $R_f$ = 0.50 (1:1 hexanes/EtOAc); $^1$H NMR (DMSO-$d_6$, 600 MHz) δ 10.09 (s, 1H), 8.09 (s, 1H), 7.72 (dd, *J* = 7.9, 1.5 Hz, 1H), 7.48 (d, *J* = 8.3 Hz, 2H), 7.30 (d, *J* = 8.5 Hz, 1H), 7.22 (td, *J* = 8.5, 1.5 Hz, 1H), 6.75 (td, *J* = 7.9, 1.1 Hz, 1H), 6.18 (dd, *J* = 8.4, 1.0 Hz, 1H), 2.37 (s, 3H) ppm; IR (thin film) ν 3482, 3191, 1660, 1616, 1581, 1506, 1449, 1392, 1287 cm$^{-1}$; LRMS (ES$^+$) calcd 295.03 for $C_{14}H_{12}Cl_2N_2O$ $[M+H]^+$ found 294.7 ($M^+$).

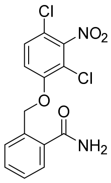

Prepared according to the above procedure from the corresponding carboxylic acid (See *Synthesis of C3-substituted 2,6-dichloronitrobenzenes* for carboxylic acid synthesis); off-white solid (187 mg, 63%). TLC $R_f$ = 0.37 (19:1 $CH_2Cl_2$/MeOH); $^1$H NMR (acetone-$d_6$, 400 MHz) δ 7.73–7.68 (m, 2H), 7.66 (d, *J* = 9.2 Hz, 1H), 7.57–7.49 (m, 2H), 7.46–7.42 (m, 2H), 6.84 (s, 1H), 5.62 (s, 2H) ppm; IR (thin film) ν 3467, 1679, 1540, 1470, 1361, 1278 cm$^{-1}$.

---

3. Suzuki, T.; Imai, K.; Nakagawa, H.; Miyata, N. 2-Anilinobenzamides as SIRT Inhibitors. *ChemMedChem* **2006**, *1*, 1059–1062.

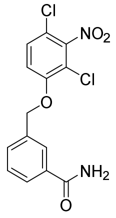

Prepared according to the above procedure from the corresponding carboxylic acid (See *Synthesis of C3-substituted 2,6-dichloronitrobenzenes* for carboxylic acid synthesis); off-white solid (223 mg, 74%). TLC $R_f$ = 0.33 (19:1 $CH_2Cl_2$/MeOH); $^1$H NMR (acetone-d$_6$, 400 MHz) δ 8.10 (s, 1H), 7.94 (d, *J* = 7.9 Hz, 1H), 7.75–7.64 (m, 2H), 7.55–7.51 (m, 3H), 6.70 (s, 1H), 5.44 (s, 2H) ppm; IR (thin film) ν 3357, 3182, 1654, 1539, 1500, 1476, 1407, 1364, 1301, 1199 cm$^{-1}$.

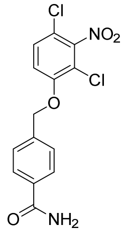

Prepared according to the above procedure from the corresponding carboxylic acid (See *Synthesis of C3-substituted 2,6-dichloronitrobenzenes* for carboxylic acid synthesis); off-white solid (222 mg, 74%). TLC $R_f$ = 0.33 (19:1 $CH_2Cl_2$/MeOH); $^1$H NMR (acetone-d$_6$, 400 MHz) δ 7.99 (d, *J* = 8.2 Hz, 2H), 7.67 (d, *J* = 8.9 Hz, 1H), 7.63 (d, *J* = 8.2 Hz, 2H), 7.50 (d, *J* = 9.2 Hz, 2H), 6.68 (s, 1H), 5.46 (s, 2H) ppm; IR (thin film) ν 3405, 1672, 1615, 1544, 1476, 1448, 1363, 1291, 1263, 1197 cm$^{-1}$.

### *Synthesis of substituted anilines*

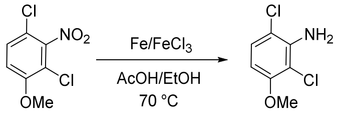

To a solution of 1,3-dichloro-4-methoxy-2-nitrobenzene (60 mg, 0.27 mmol) in 2.7 mL of EtOH was added Fe powder (106 mg, 1.89 mmol, 7.0 equiv), anhydrous $FeCl_3$ (6.7 mg, 0.04 mmol, 0.15 equiv), and glacial acetic acid (0.1 mL, 1.70 mmol, 6.3 equiv). The reaction flask was sealed and the contents stirred at 70 °C for 2 h. Following this time, the solution was cooled to room temperature, diluted with 10 mL of EtOAc, and filtered through a pad of Celite. The flask and filter cake were washed with EtOAc and the combined filtrates were concentrated under reduced pressure. The oily residue was re-dissolved in $CH_2Cl_2$ to which ~0.5 g of silica gel was then added. The suspension was concentrated under reduced pressure and the solid material dry loaded onto a silica gel column pre-packed in hexanes. Purification of this material by chromatography on silica gel (gradient elution: 0:1→1:3 EtOAc/hexanes) furnished the desired product as a white solid (36 mg, 70%). TLC $R_f$ = 0.27 (19:1 hexanes/acetone); $^1$H NMR ($CDCl_3$, 300 MHz) δ 7.12 (d, *J* = 8.9 Hz, 1H), 6.30 (d, *J* = 8.9 Hz, 1H), 3.86 (s, 3H) ppm; IR (thin film) ν 3485, 3387, 1610, 1475, 1306, 1247, 1121 cm$^{-1}$.

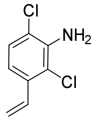

Prepared according to the above procedure; white solid (475 mg, 92%). TLC $R_f$ = 0.54 (9:1 hexanes/EtOAc); $^1$H NMR ($CDCl_3$, 300 MHz) δ 7.15 (d, *J* = 8.5 Hz, 1H), 7.02 (dd, *J* = 17.4, 10.7 Hz, 1H), 6.91 (d, *J* = 8.4 Hz, 1H), 5.71 (d, *J* = 17.4 Hz, 1H), 5.37 (d, *J* = 10.7 Hz, 1H), 4.49 (br s, 2H) ppm; IR (thin film) ν 3488, 3391, 1603, 1471, 1401 $cm^{-1}$.

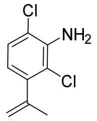

Prepared according to the above procedure; colorless oil (20 mg, 65%). TLC $R_f$ = 0.59 (19:1 hexanes/acetone); $^1$H NMR ($CDCl_3$, 500 MHz) δ 7.13 (d, *J* = 8.2 Hz, 1H), 6.54 (d, *J* = 8.2 Hz, 1H), 5.20 (quintet, *J* = 1.6 Hz, 1H), 4.94 (dq, *J* = 1.9, 1.0 Hz, 1H), 4.52 (s, 2H), 2.06 (s, 3H) ppm; IR (thin film) ν 3490, 3393, 2920, 1604, 1468, 1422 $cm^{-1}$.

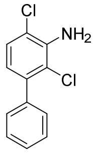

Prepared according to the above procedure; colorless oil (65 mg, 94%). TLC $R_f$ = 0.50 (19:1 hexanes/EtOAc); $^1$H NMR ($CDCl_3$, 500 MHz) δ 7.46–7.36 (m, 5H), 7.23 (d, *J* = 8.2 Hz, 1H), 6.68 (d, *J* = 8.3 Hz, 1H), 4.60 (s, 2H) ppm; IR (thin film) ν 3488, 3390, 3060, 3031, 1603, 1579, 1548, 1464, 1419 $cm^{-1}$.

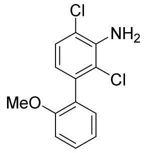

Prepared according to the above procedure; colorless oil (74 mg, 83%). TLC $R_f$ = 0.37 (19:1 hexanes/EtOAc); $^1$H NMR ($CDCl_3$, 500 MHz) δ 7.38 (td, *J* = 7.5, 1.8 Hz, 1H), 7.22 (d, *J* = 8.2 Hz, 1H), 7.16 (dd, *J* = 7.5, 1.8 Hz, 1H), 7.02 (td, *J* = 7.4, 1.1 Hz, 1H), 6.98 (dd, *J* = 8.3, 1.0 Hz, 1H), 6.64 (d, *J* = 8.2 Hz, 1H), 4.54 (s, 2H), 3.79 (s, 3H) ppm; IR (thin film) ν 3486, 3387, 2937, 2835, 1603, 1582, 1548, 1497, 1467, 1435, 1420, 1280, 1254, 1239 $cm^{-1}$.

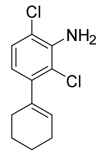

Prepared according to the above procedure; pale yellow film (38 mg, 85%). TLC $R_f$ = 0.69 (19:1 hexanes/EtOAc); $^1$H NMR ($CDCl_3$, 500 MHz) δ 7.10 (d, *J* = 8.2 Hz, 1H), 6.50 (d, *J* = 8.2 Hz, 1H), 5.64–5.62 (m, 1H), 4.48 (s, 2H), 2.25–2.21 (m, 2H), 2.19–2.14 (m, 2H), 1.79–1.71 (m, 2H), 1.69–1.65 (m, 2H) ppm; IR (thin film) ν 3490, 3392, 2931, 2857, 2835, 1603, 1547, 1467, 1421 cm$^{-1}$.

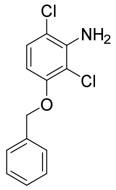

Prepared according to the above procedure; white solid (111 mg, 85%). TLC $R_f$ = 0.39 (19:1 hexanes/EtOAc); $^1$H NMR ($CDCl_3$, 400 MHz) δ 7.44 (d, *J* = 7.5 Hz, 2H), 7.39 (t, *J* = 6.8 Hz, 2H), 7.32 (t, *J* = 7.1 Hz, 1H), 7.08 (d, *J* = 8.9 Hz, 1H), 6.34 (d, *J* = 8.9 Hz, 1H), 5.12 (s, 2H), 4.49 (s, 2H) ppm; IR (thin film) ν 3302, 1615, 1476, 1452, 1303, 1248 cm$^{-1}$.

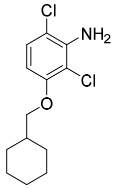

Prepared according to the above procedure; off-white solid (104 mg, 83%). TLC $R_f$ = 0.45 (19:1 hexanes/EtOAc); $^1$H NMR ($CDCl_3$, 400 MHz) δ 7.08 (d, *J* = 8.8 Hz, 1H), 6.27 (d, *J* = 8.9 Hz, 1H), 4.45 (s, 2H), 3.77 (d, *J* = 6.1 Hz, 2H), 1.92–1.82 (m, 3H), 1.81–1.66 (m, 4H), 1.36–1.17 (m, 2H), 1.08 (m, 2H) ppm; IR (thin film) ν 3332, 2921, 2857, 1614, 1478, 1449, 1309, 1118 cm$^{-1}$.

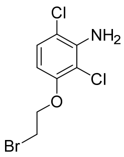

Prepared according to the above procedure; white solid (107 mg, 79%). TLC $R_f$ = 0.54 (9:1 hexanes/EtOAc); $^1$H NMR ($CDCl_3$, 500 MHz) δ 7.11 (d, *J* = 8.9 Hz, 1H), 6.29 (d, *J* = 8.9 Hz, 1H), 4.40 (s, 2H), 4.30 (t, *J* = 6.5 Hz, 2H), 3.66 (t, *J* = 6.5 Hz, 2H) ppm; IR (thin film) ν 3438, 3330, 2942, 2882, 1615, 1476, 1462, 1306, 1246 cm$^{-1}$.

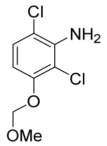

Prepared according to the above procedure; white solid (301 mg, 85%). TLC $R_f$ = 0.39 (19:1 hexanes/EtOAc); $^1$H NMR ($CDCl_3$, 400 MHz) δ 7.10 (d, *J* = 8.9 Hz, 1H), 6.54 (d, *J* = 8.9 Hz, 1H), 5.21 (s, 2H), 4.48 (s, 2H), 3.50 (s, 3H) ppm; IR (thin film) ν 3438, 3391, 3317, 2961, 2833, 1615, 1476, 1446, 1160 $cm^{-1}$.

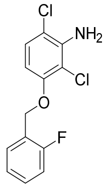

Prepared according to the above procedure; white solid (225 mg, 98%). TLC $R_f$ = 0.30 (19:1 hexanes/$Et_2O$); $^1$H NMR ($CDCl_3$, 400 MHz) δ 7.57 (td, *J* = 7.5, 0.9 Hz, 1H), 7.35–7.28 (m, 1H), 7.17 (t, *J* = 7.5 Hz, 1H), 7.13–7.04 (m, 2H), 6.38 (d, *J* = 8.9 Hz, 1H), 5.18 (s, 2H), 4.51 (s, 2H) ppm; $^{19}$F NMR ($CDCl_3$, 376 MHz) δ –119.19 to –119.31 (m, 1F) ppm; IR (thin film) ν 3293, 1616, 1587, 1493, 1477, 1451, 1388, 1313, 1302, 1230, 1114 $cm^{-1}$.

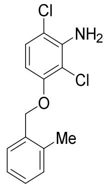

Prepared according to the above procedure; white solid (192 mg, 97%). TLC $R_f$ = 0.30 (19:1 hexanes/ $Et_2O$); $^1$H NMR ($CDCl_3$, 400 MHz) δ 7.45 (d, *J* = 7.3 Hz, 1H), 7.29–7.17 (m, 3H), 7.11 (d, *J* = 8.9 Hz, 1H), 6.38 (d, *J* = 8.9 Hz, 1H), 5.07 (s, 2H), 2.38 (s, 3H) ppm; IR (thin film) ν 3482, 3384, 2926, 1608, 1450, 1376, 1304, 1247, 1109 $cm^{-1}$.

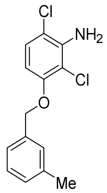

Prepared according to the above procedure; white solid (186 mg, 94%). TLC $R_f$ = 0.32 (19:1 hexanes/ $Et_2O$); $^1$H NMR ($CDCl_3$, 300 MHz) δ 7.29–7.20 (m, 3H), 7.13 (d, *J* = 7.2 Hz, 1H), 7.07 (d, *J* = 8.9 Hz, 1H), 6.34 (d, *J* = 8.9 Hz, 1H), 5.08 (s, 2H), 2.37 (s, 3H) ppm; IR (thin film) ν 3488, 3389, 3027, 2920, 1610, 1567, 1475, 1377, 1307, 1244, 1168, 1117 $cm^{-1}$.

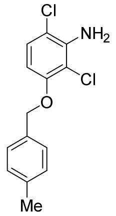

Prepared according to the above procedure; white solid (192 mg, 97%). TLC $R_f$ = 0.30 (19:1 hexanes/$Et_2O$); $^1H$ NMR ($CDCl_3$, 400 MHz) δ 7.33 (d, *J* = 8.0 Hz, 2H), 7.19 (d, *J* = 8.3 Hz, 2H), 7.07 (d, *J* = 8.9 Hz, 1H), 6.34 (d, *J* = 8.9 Hz, 1H), 5.08 (s, 2H), 2.36 (s, 3H) ppm; IR (thin film) ν 3441, 3319, 2920, 1614, 1586, 1519, 1477, 1451, 1380, 1302, 1247, 1166, 1115 $cm^{-1}$.

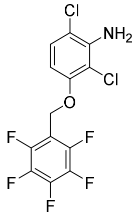

Prepared according to the above procedure; white solid (180 mg, 72%). TLC $R_f$ = 0.27 (19:1 hexanes/$Et_2O$); $^1H$ NMR ($CDCl_3$, 400 MHz) δ 7.14 (d, *J* = 8.8 Hz, 1H), 6.43 (d, *J* = 8.9 Hz, 1H), 5.14 (s, 2H) ppm; $^{19}F$ NMR ($CDCl_3$, 376 MHz) δ –142.32 (dd, *J* = 22.2, 8.6 Hz, 2F), –152.59 (t, *J* = 20.8 Hz, 1F), –161.74 (td, *J* = 21.1, 7.5 Hz, 2F) ppm. IR (thin film) ν 3442, 3338, 1659, 1613, 1525, 1510, 1479, 1468, 1452, 1389, 1300, 1248, 1135 $cm^{-1}$.

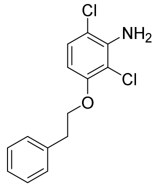

Prepared according to the above procedure; colorless residue that slowly solidified (146 mg, 99%). TLC $R_f$ = 0.30 (19:1 hexanes/$Et_2O$); $^1H$ NMR ($CDCl_3$, 400 MHz) δ 7.36–7.30 (m, 4H), 7.29–7.23 (m, 1H), 7.08 (d, *J* = 8.9 Hz, 1H), 6.27 (d, *J* = 8.9 Hz, 1H), 4.49 (s, 2H), 4.18 (t, *J* = 7.0 Hz, 2H), 3.15 (t, *J* = 7.0 Hz, 2H) ppm; IR (thin film) ν 3487, 3388, 2928, 1610, 1497, 1476, 1465, 1386, 1307, 1247, 1168, 1117 $cm^{-1}$.

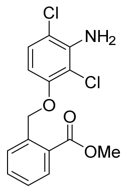

Prepared according to the above procedure; off-white solid (212 mg, 77%). TLC $R_f$ = 0.37 (9:1 hexanes/EtOAc); $^1H$ NMR ($CDCl_3$, 400 MHz) δ 8.04 (dd, *J* = 7.8, 1.4 Hz, 1H), 7.87 (dd, *J* = 8.0, 1.1 Hz, 1H), 7.59 (td, *J* = 7.7, 1.4 Hz, 1H), 7.39 (t, *J* = 7.4 Hz, 1H), 7.09 (d, *J* = 8.9 Hz, 1H), 6.39 (d, *J* = 8.9 Hz, 1H),

5.54 (s, 2H), 4.52 (s, 2H), 3.91 (s, 3H) ppm; IR (thin film) ν 3432, 3320, 2942, 1719, 1616, 1474, 1313, 1262, 1140 $cm^{-1}$.

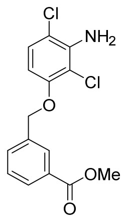

Prepared according to the above procedure; off-white solid (243 mg, 89%). TLC $R_f$ = 0.20 (9:1 hexanes/EtOAc); [1]H NMR ($CDCl_3$, 400 MHz) δ 8.10 (s, 1H), 8.00 (d, *J* = 8.0 Hz, 1H), 7.68 (d, *J* = 7.6 Hz, 1H), 7.47 (t, *J* = 7.6 Hz, 1H), 7.08 (d, *J* = 8.8 Hz, 1H), 6.32 (d, *J* = 8.8 Hz, 1H), 5.15 (s, 2H), 4.52 (s, 2H), 3.93 (s, 3H) ppm; IR (thin film) ν 3484, 3381, 2951, 1720, 1611, 1475, 1308, 1289, 1248, 1205, 1110 $cm^{-1}$.

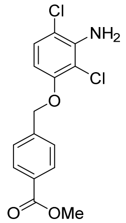

Prepared according to the above procedure; off-white solid (228 mg, 83%). TLC $R_f$ = 0.27 (9:1 hexanes/EtOAc); [1]H NMR ($CDCl_3$, 400 MHz) δ 8.06 (d, *J* = 7.8 Hz, 2H), 7.52 (d, *J* = 7.8 Hz, 2H), 7.08 (d, *J* = 8.9 Hz, 1H), 6.30 (d, *J* = 8.9 Hz, 1H), 5.17 (s, 2H), 4.53 (s, 2H), 3.92 (s, 3H) ppm; IR (thin film) ν 3471, 3381, 2952, 1716, 1614, 1475, 1435, 1310, 1283, 1111 $cm^{-1}$.

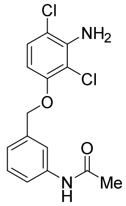

Prepared according to the above procedure; tan solid (155 mg, 91%). TLC $R_f$ = 0.43 (19:1 $CH_2Cl_2$/MeOH); [1]H NMR (DMSO-$d_6$, 500 MHz) δ 9.98 (s, 1H), 7.65 (s, 1H), 7.53 (d, *J* = 7.9 Hz, 1H), 7.30 (t, *J* = 7.8 Hz, 1H), 7.15 (d, *J* = 8.9 Hz, 1H), 7.09 (d, *J* = 7.6 Hz, 1H), 6.44 (d, *J* = 9.0 Hz, 1H), 5.48 (s, 2H), 5.11 (s, 2H), 2.03 (s, 3H) ppm; IR (thin film) ν 3311, 1666, 1612, 1552, 1473, 1370, 1306, 1246, 1199 $cm^{-1}$.

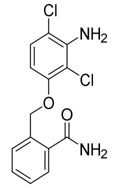

Prepared according to the above procedure; off-white solid (133 mg, 97%). TLC $R_f$ = 0.37 (19:1 $CH_2Cl_2$/MeOH); $^1$H NMR (DMSO-$d_6$, 500 MHz) δ 7.91 (s, 1H), 7.60 (d, *J* = 7.6 Hz, 1H), 7.56 (d, *J* = 7.6 Hz, 1H), 7.50 (t, *J* = 7.6 Hz, 1H), 7.47 (s, 1H), 7.40 (t, *J* = 7.4 Hz, 1H), 7.16 (d, *J* = 8.8 Hz, 1H), 6.38 (d, *J* = 8.9 Hz, 1H), 5.50 (s, 2H), 5.34 (s, 2H) ppm; IR (thin film) ν 3384, 3188, 1645, 1607, 1594, 1473, 1310, 1246, 1200, 1117 $cm^{-1}$.

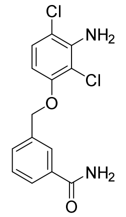

Prepared according to the above procedure; off-white solid (190 mg, 99%). TLC $R_f$ = 0.33 (19:1 $CH_2Cl_2$/MeOH); $^1$H NMR (DMSO-$d_6$, 500 MHz) δ 8.01 (s, 1H), 7.95 (s, 1H), 7.83 (d, *J* = 7.6 Hz, 1H), 7.60 (d, *J* = 7.6 Hz, 1H), 7.49 (t, *J* = 7.6 Hz, 1H), 7.41 (s, 1H), 7.17 (d, *J* = 8.9 Hz, 1H), 6.49 (d, *J* = 8.9 Hz, 1H), 5.50 (s, 2H), 5.19 (s, 2H) ppm; IR (thin film) ν 3345, 3182, 1651, 1606, 1474, 1452, 1406, 1375, 1305, 1244, 1200, 1116 $cm^{-1}$.

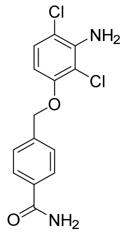

Prepared according to the above procedure; off-white solid (174 mg, 95%). TLC $R_f$ = 0.33 (19:1 $CH_2Cl_2$/MeOH); $^1$H NMR (DMSO-$d_6$, 500 MHz) δ 7.98 (s, 1H), 7.89 (d, *J* = 8.1 Hz, 2H), 7.51 (d, *J* = 8.1 Hz, 2H), 7.39 (s, 1H), 7.17 (d, *J* = 8.9 Hz, 1H), 6.47 (d, *J* = 9.0 Hz, 1H), 5.51 (s, 2H), 5.22 (s, 2H) ppm; IR (thin film) ν 3338, 3178, 1641, 1614, 1572, 1477, 1452, 1304, 1120 $cm^{-1}$.

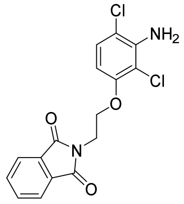

Prepared according to the above procedure; pale yellow solid (79 mg, 54%). TLC $R_f$ = 0.44 (3:2 hexanes/EtOAc); $^1$H NMR ($CDCl_3$, 500 MHz) δ 7.86 (dd, *J* = 5.4, 3.0 Hz, 2H), 7.72 (dd, *J* = 5.4, 3.0 Hz, 2H), 7.06 (d, *J* = 8.9 Hz, 1H), 6.28 (d, *J* = 8.9 Hz, 1H), 4.44 (s, 2H), 4.23 (t, *J* = 5.7 Hz, 2H), 4.15 (t, *J* = 5.7 Hz, 2H) ppm; IR (thin film) ν 3480, 3374, 2918, 2849, 1712, 1611, 1464, 1393, 1306 $cm^{-1}$.

### *Hydrogenation of styrenyl olefins*

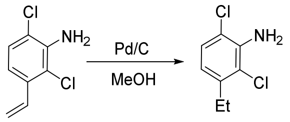

Pd/C (5 wt %, 1.0 mg) was added to a solution of 2,6-dichloro-3-vinylaniline (20 mg, 0.11 mmol) in 2.5 mL of MeOH. The reaction flask was sealed with a septum and fitted with a $N_2$ gas inlet and an 18-gauge needle. The flask was flushed with $N_2$ for 5 min. The $N_2$ line was then replaced with a balloon of $H_2$ and the headspace of the flask was swept with $H_2$. The outlet needle was removed, the flask equipped with a fresh $H_2$ balloon, and the black suspension was stirred for exactly 1 h (Note: reaction times beyond 1 h resulted in inseparable mixtures of proto-dehalogenation products). Following this time, the mixture was sparged for 5 min with a gentle-stream of $N_2$ gas, diluted with 10 mL of EtOAc, and filtered through a pad of Celite. The flask and filter cake were washed with EtOAc and the combined filtrates were concentrated under reduced pressure. The oily residue was re-dissolved in $CH_2Cl_2$ to which ~0.5 g of silica gel was then added. The suspension was concentrated under reduced pressure and the solid material dry loaded onto a silica gel column pre-packed in hexanes. Purification of this material by chromatography on silica gel (gradient elution, 0:1→1:3 EtOAc/hexanes) furnished the desired product as a clear oil (10.3 mg, 51%). TLC $R_f$ = 0.38 (19:1 hexanes/EtOAc); $^1$H NMR ($CDCl_3$, 300 MHz) δ 7.13 (d, *J* = 8.3 Hz, 1H), 6.61 (d, *J* = 8.4 Hz, 1H), 4.49 (s, 2H), 2.72 (q, *J* = 7.5 Hz, 2H), 1.23 (t, *J* = 7.5 Hz, 3H) ppm; IR (thin film) ν 3489, 3392, 2970, 2935, 2873, 1607, 1471, 1432 $cm^{-1}$.

### *Synthesis of substituted 2-bromo esters*

#### *Fischer esterification*

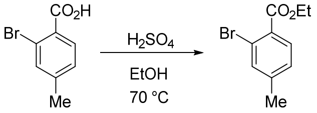

2-Bromo-4-methylbenzoic acid (500 mg, 2.3 mmol) was dissolved in 6.0 mL of EtOH to which activated 3 Å molecular sieves (~0.5 g) were then added. Concentrated $H_2SO_4$ (1.0 mL) was added dropwise, the flask was fitted with a refluxed condenser, and the suspension was stirred at 70 °C for 3 h. Following this time, the mixture was cooled to room temperature and filtered through a pad of Celite. The flask and filter cake were rinsed with ~20 mL of EtOAc. The combined filtrates were transferred to a separatory funnel containing 70 mL of EtOAc. The organic fraction was washed with 1 x 70 mL of $H_2O$, 1 x 70 mL of saturated aqueous $NaHCO_3$, and 1 x 70 mL of saturated aqueous NaCl. The organic layer was collected, dried over $MgSO_4$, filtered and concentrated under reduced pressure. Purification of this material by chromatography on silica gel (gradient elution, 0:1→1:19 acetone/hexanes) furnished the desired product as a clear, colorless oil (395 mg, 71%). TLC $R_f$ = 0.57 (19:1 hexanes/acetone); $^1$H NMR ($CDCl_3$, 300 MHz) δ 7.71 (d, *J* = 7.9 Hz, 1H), 7.48 (s, 1H), 7.15 (d, *J* = 7.9 Hz, 1H), 4.38 (q, *J* = 7.1 Hz, 2H), 2.36 (s, 3H), 1.39 (t, *J* = 7.1 Hz, 3H) ppm; IR (thin film) ν 2982, 1732, 1447, 1408, 1366, 1291, 1245, 1182, 1146 cm$^{-1}$.

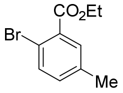

Prepared according to the above procedure; colorless oil (461 mg, 82%). TLC $R_f$ = 0.57 (19:1 hexanes/acetone); $^1$H NMR ($CDCl_3$, 300 MHz) δ 7.57 (dd, *J* = 2.3, 0.8 Hz, 1H), 7.51 (d, *J* = 8.2 Hz, 1H), 7.12 (ddq, *J* = 8.2, 2.3, 0.7 Hz, 1H), 4.39 (q, *J* = 7.1 Hz, 2H), 2.33 (s, 3H), 1.40 (t, *J* = 7.1 Hz, 3H) ppm; IR (thin film) ν 2982, 1732, 1471, 1297, 1251, 1203, 1110 cm$^{-1}$.

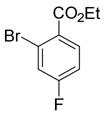

Prepared according to the above procedure; colorless oil (374 mg, 66%). TLC $R_f$ = 0.41 (19:1 hexanes/acetone); $^1$H NMR ($CDCl_3$, 300 MHz) δ 7.86 (dd, *J* = 8.7, 6.0 Hz, 1H), 7.40 (dd, *J* = 8.3, 2.5 Hz, 1H), 7.07 (ddd, *J* = 8.8, 7.7, 2.5 Hz, 1H), 4.39 (q, *J* = 7.1 Hz, 2H), 1.40 (t, *J* = 7.1 Hz, 3H) ppm; IR (thin film) ν 3081, 2983, 1732, 1596, 1487, 1385, 1366, 1286, 1253, 1208, 1109 cm$^{-1}$.

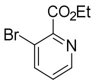

Prepared according to the above procedure; colorless oil (364 mg, 64%). TLC $R_f$ = 0.13 (19:1 hexanes/acetone); $^1$H NMR ($CDCl_3$, 300 MHz) δ 8.61 (dd, *J* = 4.6, 1.4 Hz, 1H), 7.99 (dd, *J* = 8.2, 1.4 Hz, 1H), 7.29 (dd, *J* = 8.2, 4.6 Hz, 1H), 4.49 (q, *J* = 7.1 Hz, 2H), 1.44 (t, *J* = 7.1 Hz, 3H) ppm; IR (thin film) ν 3056, 2983, 1732, 1570, 1424, 1367, 1301, 1205, 1174, 1141 cm$^{-1}$.

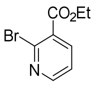

Prepared according to the above procedure (the corresponding methyl ester may be prepared in >90%

yield via the protocol described in *Esterification through EDC coupling* and alternatively used for the synthesis of AK-44); colorless oil (140 mg, 31%). TLC $R_f$ = 0.37 (3:1 hexanes/acetone); $^1$H NMR ($CDCl_3$, 300 MHz, appears as a mixture of protonated and free base) δ 8.48 (dd, *J* = 4.7, 2.1 Hz, 1H), 8.07 (dd, *J* = 7.7, 2.1 Hz, 1H), 7.34 (dd, *J* = 7.7, 4.8 Hz, 1H), 4.43 (q, *J* = 7.1 Hz, 2H), 1.42 (t, *J* = 7.1, 3H) ppm; IR (thin film) ν 2983, 1732, 1578, 1557, 1402, 1367, 1301, 1275, 1245, 1142 $cm^{-1}$.

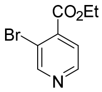

Prepared according to the above procedure; colorless oil (130 mg, 29%). TLC $R_f$ = 0.40 (3:1 hexanes/acetone); $^1$H NMR ($CDCl_3$, 300 MHz) δ 8.85 (s, 1H), 8.62 (d, *J* = 4.9 Hz, 1H), 7.62 (dd, *J* = 4.9, 0.7 Hz, 1H), 4.44 (q, *J* = 7.1 Hz, 2H), 1.42 (t, *J* = 7.1 Hz, 3H) ppm; IR (thin film) ν 2983, 1737, 1470, 1398, 1367, 1303, 1270, 1214, 1181, 1129 $cm^{-1}$.

*Esterification with MeI on sterically hindered 2,6-disubstituted substrates*

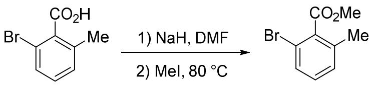

To an ice-cold suspension of NaH (59 mg, 2.45 mmol, 1.2 equiv) in 3.0 mL of anhydrous DMF was added dropwise via cannula a solution of 2-bromo-6-methylbenzoic acid (439 mg, 2.04 mmol) in 9 mL of DMF. Gas evolution ensued immediately and stirring at 0 °C continued until bubbling ceased (~30 min). Iodomethane (0.25 mL, 4.08 mmol, 2.0 equiv) was added dropwise via syringe, the solution was warmed to ambient temperature and stirred for 2 h. The flask was then fitted with a reflux condenser and the contents stirred at 80 °C for 1 h. Upon heating, the opaque reaction mixture slowly transitioned to clear yellow. The reaction was cooled to room temperature and transferred to a separatory funnel with 200 mL of $Et_2O$. The ethereal layer was washed with 4 x 200 mL $H_2O$, dried over $MgSO_4$, filtered and concentrated under reduced pressure to an oily residue. Purification of this material by chromatography on silica gel (gradient elution, 0:1→1:19 EtOAc/hexanes) furnished the desired product as a colorless oil (363 mg, 78%). TLC $R_f$ = 0.30 (19:1 hexanes/acetone); $^1$H NMR ($CDCl_3$, 300 MHz) δ 7.44–7.36 (m, 1H), 7.21–7.12 (m, 2H), 3.95 (s, 3H), 2.33 (s, 3H) ppm; IR (thin film) ν 2952, 1738, 1594, 1565, 1451, 1278, 1245, 1178, 1150, 1103 $cm^{-1}$.

*Esterification through EDC coupling*

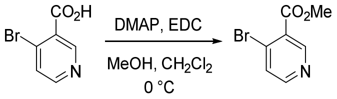

The following procedure is adapted from Stangeland, et al.[4] To an ice-cold suspension of 4-bromonicotinic acid (102 mg, 0.50 mmol) in 2.0 mL of $CH_2Cl_2$ was added successively 4-dimethylaminopyridine (6.2 mg, 0.05 mmol, 0.1 equiv), freshly distilled MeOH (80 μL, 2.0 mmol, 4.0 equiv), and 1-ethyl-3-(3-dimethylaminopropyl)carbodiimide (107 mg, 0.55 mmol, 1.1 equiv). Upon addition of EDC, the solution changed from opaque to clear. The reaction was stirred for 3 h at 0 °C and then quenched by the addition of 3 mL of $H_2O$. The reaction was transferred to a separatory funnel with 30 mL of $H_2O$ and extracted with 1 x 30 mL of $CH_2Cl_2$. The organic layer was washed with 2 x 30 mL of saturated aqueous NaCl, dried over $Na_2SO_4$, filtered and concentrated under reduced pressure to a yellow oil (97 mg, 89%). This material was judged to be sufficiently pure by $^1$H NMR for use in the subsequent step without further purification. TLC $R_f$ = 0.32 (3:1 hexanes/EtOAc); $^1$H NMR ($CDCl_3$, 500 MHz) δ 8.97 (s, 1H), 8.45 (d, $J$ = 5.3 Hz, 1H), 7.61 (d, $J$ = 5.3 Hz, 1H), 3.96 (s, 3H) ppm; IR (thin film) ν 2953, 1738, 1569, 1548, 1466, 1435, 1396, 1291, 1272, 1217, 1127 cm$^{-1}$.

#### *Buchwald-Hartwig amination*

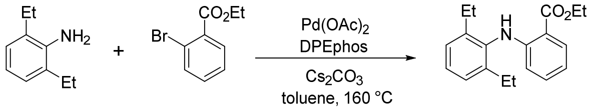

The following procedure is adapted from Sadighi, et al.**Error! Bookmark not defined.** An oven-dried 5-mL microwave vial was charged with 2,6-diethylaniline (200 mg, 1.34 mmol), ethyl 2-bromobenzoate (307 mg, 1.34 mmol), and DPEphos (54 mg, 0.10 mmol, 0.075 equiv). The vial was sealed with a septum and purged with argon. Anhydrous toluene (3.0 mL) was added and the solution was sparged with a gentle stream of argon for ~5 min. The septum was quickly removed and $Pd(OAc)_2$ (15 mg, 0.067 mmol, 0.05 equiv) was added in a single portion. The vial was resealed and the yellow solution was stirred under argon for 10 min. Following this time, the septum was quickly removed and $Cs_2CO_3$ (611 mg, 1.88 mmol, 1.4 equiv) was added. The headspace of the vial was flushed with argon and the vessel was quickly resealed with a crimped microwave vial cap. The reaction was stirred at 160 °C in a microwave reactor for 2 h (Note: the reaction may also be performed with conventional heating at 115 °C for 20 h; reaction yields are comparable). The reaction mixture was cooled to room temperature, diluted with 10 mL of EtOAc, and filtered through a plug of Celite. The flask and filter cake were rinsed with ~30 mL of EtOAc, and the combined filtrates were concentrated under reduced pressure. The brown residue was re-dissolved in $CH_2Cl_2$ to which ~1 g of silica gel was then added. The suspension was concentrated under reduced pressure and the solid material dry loaded onto a silica gel column pre-packed in pentane. Purification by chromatography on silica gel (gradient elution, 0:1→1:19 $Et_2O$/pentane) furnished the desired product as a white solid (180 mg, 45%). TLC $R_f$ = 0.40 (19:1 hexanes/acetone); $^1$H NMR ($CDCl_3$, 300 MHz) δ 9.05 (s, 1H), 7.97 (dd, $J$ = 8.0, 1.7 Hz, 1H), 7.26–7.13 (m,

---

4. Stangeland, E. L.; Patterson, L. J.; Zipfel, S. 1-(2-phenoxymethylheteroaryl)piperidine and piperazine compounds. US20110230495 A1, September 22, 2011.

4H), 6.61 (ddd, $J$ = 8.1, 7.1, 1.2 Hz, 1H), 6.19 (dd, $J$ = 8.5, 1.2 Hz, 1H), 4.39 (q, $J$ = 7.1 Hz, 2H), 2.55 (dq, $J$ = 14.6, 7.3 Hz, 4H), 1.43 (t, $J$ = 7.1 Hz, 3H), 1.13 (t, $J$ = 7.6 Hz, 6H) ppm; IR (thin film) ν 3320, 2967, 2934, 1680, 1578, 1506, 1453, 1253, 1231 cm$^{-1}$.

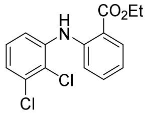

Prepared according to the above procedure; pale yellow residue (207 mg, 67%). TLC $R_f$ = 0.6 (9:1 hexanes/acetone); $^1$H NMR ($CDCl_3$, 400 MHz) δ 9.70 (s, 1H), 8.03 (d, $J$ = 8.0, 1H), 7.45–7.34 (m, 2H), 7.30 (d, $J$ = 8.4 Hz, 1H), 7.15–7.12 (m, 2H), 6.87 (t, $J$ = 7.6 Hz, 1H), 4.39 (q, $J$ = 7.1 Hz, 2H), 1.42 (t, $J$ = 7.1, 3H) ppm; IR (thin film) ν 3299, 3261, 2982, 1687, 1581, 1522, 1448, 1402, 1368, 1320, 1249, 1185, 1164, 1144, 1084 cm$^{-1}$.

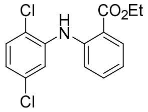

Prepared according to the above procedure; white crystalline solid (174 mg, 56%). TLC $R_f$ = 0.72 (9:1 hexanes/acetone); $^1$H NMR ($CDCl_3$, 400 MHz) δ 9.68 (s, 1H), 8.03 (dd, $J$ = 8.0, 1.7 Hz, 1H), 7.49 (d, $J$ = 2.4 Hz, 1H), 7.45–7.40 (m, 1H), 7.38–7.32 (m, 2H), 6.94–6.86 (m, 2H), 4.39 (q, $J$ = 7.1 Hz, 2H), 1.42 (t, $J$ = 7.2 Hz, 3H) ppm; IR (thin film) ν 3293, 2986, 1689, 1583, 1520, 1456, 1410, 1256, 1224, 1085 cm$^{-1}$.

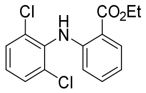

Prepared according to the above procedure (Note: the reaction was stirred for 1 h at 160 °C; these conditions were employed prior to reaction optimization); off-white solid (62 mg, 17%). TLC $R_f$ = 0.58 (9:1 hexanes/acetone); $^1$H NMR ($CDCl_3$, 400 MHz) δ 9.41 (s, 1H), 8.02 (d, $J$ = 8.0 Hz, 1H), 7.42 (d, $J$ = 8.1 Hz, 2H), 7.30–7.24 (m, 1H), 7.15 (t, $J$ = 8.1 Hz, 1H), 6.78 (t, $J$ = 7.5 Hz, 1H), 6.35 (d, $J$ = 8.5 Hz, 1H), 4.40 (q, $J$ = 7.1 Hz, 2H), 1.43 (t, $J$ = 7.1 Hz, 3H) ppm; IR (thin film) ν 3297, 3081, 2981, 1683, 1586, 1506, 1452, 1314, 1256, 1162, 1144, 1084 cm$^{-1}$.

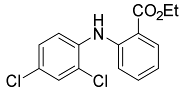

Prepared according to the above procedure with conventional heating at 100 °C for 16 h (unoptimized reaction conditions); off-white solid (23 mg, 7%). TLC 0.60 $R_f$ = (9:1 hexanes/acetone); $^1$H NMR ($CDCl_3$, 500 MHz) δ 9.61 (s, 1H), 8.02 (dd, $J$ = 8.0, 1.7 Hz, 1H), 7.46–7.40 (m, 2H), 7.37 (ddd, $J$ = 8.7, 7.1, 1.7 Hz, 1H), 7.22 (d, $J$ = 8.5, 1H), 7.18 (dd, $J$ = 8.7, 2.4 Hz, 1H), 6.85 (t, $J$ = 8.1 Hz, 1H), 4.39 (q, $J$ = 7.1 Hz, 2H), 1.42 (t, $J$ = 7.1 Hz, 3H) ppm; IR (thin film) ν 3294, 2925, 1690, 1592, 1521, 1479, 1455, 1321, 1255, 1225 cm$^{-1}$.

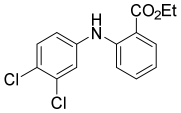

Prepared according to the above procedure with conventional heating at 100 °C for 16 h. rac-BINAP (0.15 equiv) was used in place of DPEphos (unoptimized reaction conditions); white solid (42 mg, 14%). TLC $R_f$ = 0.60 (9:1 hexanes/acetone); $^1$H NMR ($CDCl_3$, 500 MHz) δ 9.56 (s, 1H), 8.02 (dd, *J* = 8.1, 1.7 Hz, 1H), 7.41–7.35 (m, 3H), 7.27 (d, *J* = 7.9 Hz, 1H), 7.08 (dd, *J* = 8.7, 2.6 Hz, 1H), 6.84 (t, *J* = 8.1 Hz, 1H), 4.38 (q, *J* = 7.1 Hz, 2H), 1.43 (t, *J* = 7.1 Hz, 3H) ppm; IR (thin film) v 3307, 2985, 2927, 1683, 1587, 1562, 1517, 1476, 1454, 1368, 1323, 1257, 1230, 1149 cm$^{-1}$.

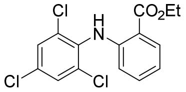

Prepared according to the above procedure; off-white solid (188 mg, 55%). TLC $R_f$ = 0.68 (24:1 pentane/$Et_2O$); $^1$H NMR ($CDCl_3$, 500 MHz) δ 9.37 (s, 1H), 8.01 (dd, *J* = 8.1, 1.7 Hz, 1H), 7.44 (s, 2H), 7.29 (ddd, *J* = 8.7, 7.2, 1.7 Hz, 1H), 6.80 (ddd, *J* = 8.1, 7.1, 1.1 Hz, 1H), 6.32 (dd, *J* = 8.4, 1.1 Hz, 1H), 4.40 (q, *J* = 7.1 Hz, 2H), 1.43 (t, *J* = 7.3 Hz, 3H) ppm; IR (thin film) v 3307, 3074, 2984, 1682, 1583, 1505, 1450, 1369, 1312, 1253, 1145, 1091 cm$^{-1}$.

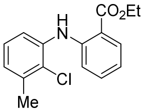

Prepared according to the above procedure; 1 h reaction time (unoptimized reaction conditions); white solid (20 mg, 7%). TLC $R_f$ = 0.60 (9:1 hexanes/acetone); $^1$H NMR ($CDCl_3$, 500 MHz) δ 9.65 (s, 1H), 8.04 (dd, *J* = 8.0, 1.7 Hz, 1H), 7.39–7.35 (m, 2H), 7.31–7.29 (m, 1H), 7.13 (t, *J* = 7.8 Hz, 1H), 6.96 (d, *J* = 7.6 Hz, 1H), 6.83 (t, *J* = 7.7 Hz, 1H), 4.42 (q, *J* = 7.1 Hz, 2H), 2.45 (s, 3H), 1.45 (t, *J* = 7.1 Hz, 3H) ppm; IR (thin film) v 3307, 2981, 1687, 1594, 1577, 1520, 1472, 1454, 1323, 1253, 1224 cm$^{-1}$.

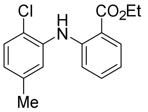

Prepared according to the above procedure; yellow oil (80 mg, 28%). TLC $R_f$ = 0.68 (24:1 pentane/$Et_2O$); $^1$H NMR ($CDCl_3$, 400 MHz) δ 9.54 (s, 1H), 8.00 (dd, *J* = 8.0, 1.7 Hz, 1H), 7.38–7.28 (m, 3H), 7.27–7.21 (m, 1H), 6.83–6.77 (m, 2H), 4.38 (q, *J* = 7.5 Hz, 2H), 2.30 (s, 3H), 1.41 (t, *J* = 7.1 Hz, 3H) ppm; IR (thin film) v 3306, 2980, 2927, 1687, 1585, 1522, 1455, 1368, 1319, 1255, 1228, 1163, 1145, 1083 cm$^{-1}$.

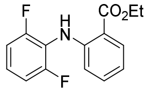

Prepared according to the above procedure; colorless oil (140 mg, 50%). TLC $R_f$ = 0.61 (24:1 pentane/$Et_2O$); $^1H$ NMR ($CDCl_3$, 400 MHz) δ 9.21 (s, 1H), 8.01 (dd, *J* = 8.0, 1.7 Hz, 1H), 7.33 (t, *J* = 7.9 Hz, 1H), 7.20–7.11 (m, 1H), 6.99 (t, *J* = 8.0 Hz, 2H), 6.79 (t, *J* = 7.6 Hz, 1H), 6.60 (d, *J* = 8.0 Hz, 1H), 4.39 (q, *J* = 7.2 Hz, 2H), 1.42 (t, *J* = 7.2 Hz, 3H) ppm; IR (thin film) ν 3308, 2983, 1683, 1584, 1519, 1474, 1455, 1320, 1280, 1245, 1164, 1145, 1085 $cm^{-1}$.

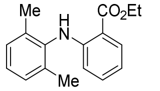

Prepared according to the above procedure; 1.5 h reaction time (unoptimized reaction conditions); yellow oil (190 mg, 40%). TLC $R_f$ = 0.58 (19:1 hexanes/acetone); $^1H$ NMR ($CDCl_3$, 500 MHz) δ 9.04 (s, 1H), 7.99 (dd, *J* = 8.0, 1.4 Hz, 1H), 7.20 (m, 1H), 7.14 (m, 3H), 6.63 (ddd, *J* = 8.1, 7.0, 1.1 Hz, 1H), 6.20 (dd, *J* = 8.6, 1.2 Hz, 1H), 4.39 (q, *J* = 7.1 Hz, 2H), 2.21 (s, 6H), 1.43 (t, *J* = 7.1 Hz, 3H) ppm; IR (thin film) ν 3320, 2980, 1682, 1579, 1507, 1454, 1251, 1160, 1142 $cm^{-1}$.

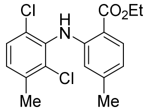

Prepared according to the above procedure; white solid (115 mg, 40%). TLC $R_f$ = 0.52 (19:1 hexanes/acetone); $^1H$ NMR ($CDCl_3$, 300 MHz) δ 9.37 (s, 1H), 7.90 (d, *J* = 8.1 Hz, 1H), 7.32 (d, *J* = 8.1 Hz, 1H), 7.11 (d, *J* = 8.3 Hz, 1H), 6.58 (d, *J* = 8.2 Hz, 1H), 6.10 (s, 1H), 4.38 (q, *J* = 7.1 Hz, 2H), 2.42 (s, 3H), 2.22 (s, 3H), 1.42 (t, *J* = 7.1 Hz, 3H) ppm; IR (thin film) ν 3287, 2978, 2926, 1677, 1615, 1572, 1510, 1460, 1241, 1156 $cm^{-1}$.

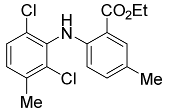

Prepared according to the above procedure; white solid (32 mg, 10%). TLC $R_f$ = 0.52 (19:1 hexanes/acetone); $^1H$ NMR ($CDCl_3$, 300 MHz) δ 9.22 (s, 1H), 7.83–7.78 (m, 1H), 7.29 (d, *J* = 8.2 Hz, 1H), 7.11–7.07 (m, 2H), 6.25 (d, *J* = 8.5 Hz, 1H), 4.40 (q, *J* = 7.1 Hz, 2H), 2.40 (s, 3H), 2.27 (s, 3H), 1.43 (t, *J* = 7.1 Hz, 3H) ppm; IR (thin film) ν 3306, 2981, 2924, 1685, 1584, 1511, 1461, 1256, 1234, 1207 $cm^{-1}$.

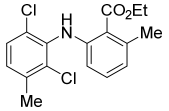

Prepared according to the above procedure; white solid (120 mg, 50%). TLC $R_f$ = 0.42 (19:1 hexanes/acetone); $^1H$ NMR ($CDCl_3$, 300 MHz) δ 8.24 (s, 1H), 7.27 (d, *J* = 8.0 Hz, 1H), 7.13–7.02 (m, 2H),

6.69 (d, *J* = 7.6 Hz, 1H), 6.23 (d, *J* = 8.4 Hz, 1H), 3.96 (s, 3H), 2.48 (s, 3H), 2.39 (s, 3H) ppm; IR (thin film) ν 3289, 2947, 1677, 1583, 1492, 1460, 1303, 1253 cm$^{-1}$.

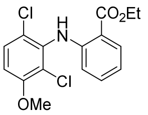

Prepared according to the above procedure; white solid (33 mg, 52%). TLC $R_f$ = 0.32 (19:1 hexanes/acetone); $^1$H NMR ($CDCl_3$, 300 MHz) δ 9.42 (s, 1H), 8.01 (dd, *J* = 8.0, 1.7 Hz, 1H), 7.36 (d, *J* = 9.0 Hz, 1H), 7.27 (ddd, *J* = 8.6, 7.1, 1.7 Hz, 1H), 6.82 (d, *J* = 9.0 Hz, 1H), 6.77 (ddd, *J* = 8.1, 7.1, 1.1 Hz, 1H), 6.37 (dd, *J* = 8.4, 1.1 Hz, 1H), 4.40 (q, *J* = 7.1 Hz, 2H), 3.93 (s, 3H), 1.43 (t, *J* = 7.1 Hz, 3H) ppm; IR (thin film) ν 3300, 2980, 1683, 1578, 1508, 1466, 1455, 1252 cm$^{-1}$.

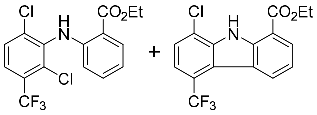

Prepared according to the above procedure; isolated as an inseparable mixture of products and resolved by reversed-phase HPLC following ester hydrolysis (see below for details); white solid (137 mg); TLC $R_f$ = 0.33 (19:1 hexanes/EtOAc, single spot for both products); $^1$H NMR ($CDCl_3$, 400 MHz), see spectrum below; IR (thin film) ν 3435, 1688, 1586, 1501, 1322, 1257, 1134 cm$^{-1}$.

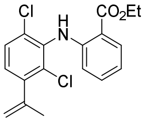

Prepared according to the above procedure; isolated as an inseparable mixture of products and resolved by HPLC following ester hydrolysis (see below for details); white solid (93 mg); TLC $R_f$ = 0.9 (19:1 hexanes/acetone); IR (thin film) ν 3492, 3392, 3299, 2976, 1683, 1604, 1588, 1506, 1454, 1369, 1318, 1251 cm$^{-1}$.

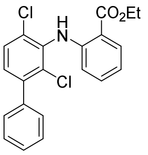

Prepared according to the above procedure; white solid (6 mg, 6%). TLC $R_f$ = 0.43 (19:1 hexanes/EtOAc); $^1$H NMR ($CDCl_3$, 500 MHz) δ 9.49 (s, 1H), 8.02 (dd, *J* = 8.0, 1.6 Hz, 1H), 7.48–7.38 (m, 6H), 7.31 (ddd, *J* = 8.6, 7.1, 1.7 Hz, 1H), 7.21 (d, *J* = 8.3 Hz, 1H), 6.78 (ddd, *J* = 8.1, 7.1, 1.1 Hz, 1H), 6.44 (dd, *J* = 8.4, 1.1 Hz, 1H), 4.40 (q, *J* = 7.1 Hz, 2H), 1.43 (t, *J* = 7.1 Hz, 3H) ppm; IR (thin film) ν 3298, 1684, 1535, 1508, 1452, 1368, 1316, 1251, 1162, 1144 cm$^{-1}$.

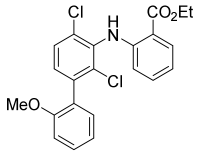

Prepared according to the above procedure; colorless film (26 mg, 22%). TLC $R_f$ = 0.23 (19:1 hexanes/EtOAc); $^1$H NMR ($CDCl_3$, 500 MHz) δ 9.49 (s, 1H), 8.02 (dd, *J* = 8.0, 1.6 Hz, 1H), 7.45 (d, *J* = 8.2 Hz, 1H), 7.40 (ddd, *J* = 8.3, 7.4, 1.7 Hz, 1H), 7.31 (t, *J* = 7.3 Hz, 1H), 7.21 (dd, *J* = 7.5, 1.7 Hz, 1H), 7.17 (d, *J* = 8.3 Hz, 1H), 7.04 (td, *J* = 7.4, 1.1 Hz, 1H), 6.99 (d, *J* = 8.3 Hz, 1H), 6.78 (t, *J* = 7.6 Hz, 1H), 6.48 (d, *J* = 52.5 Hz, 1H), 4.40 (q, *J* = 7.1 Hz, 2H), 3.80 (s, 3H), 1.43 (t, *J* = 7.1, 1.0 Hz, 3H) ppm; IR (thin film) ν 3299, 2980, 1683, 1584, 1508, 1463, 1451, 1251 $cm^{-1}$.

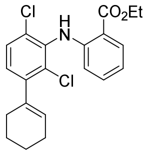

Prepared according to the above procedure; colorless film (21 mg, 12%). TLC $R_f$ = 0.40 (19:1 hexanes/EtOAc); $^1$H NMR ($CDCl_3$, 500 MHz) δ 9.40 (s, 1H), 8.01 (dd, *J* = 8.0, 1.7 Hz, 1H), 7.34 (d, *J* = 8.3 Hz, 1H), 7.38 (td, *J* = 8.4, 1.6 Hz, 1H), 7.03 (d, *J* = 8.3 Hz, 1H), 6.76 (td, *J* = 7.1, 1.1 Hz, 1H), 6.36 (dd, *J* = 8.4, 1.0 Hz, 1H), 5.69 (tt, *J* = 3.7, 1.8 Hz, 1H), 4.40 (q, *J* = 7.1 Hz, 2H), 2.32–2.24 (m, 2H), 2.22–2.13 (m, 2H), 1.80–1.72 (m, 2H), 1.72–1.64 (m, 2H), 1.42 (t, *J* = 7.1 Hz, 3H) ppm; IR (thin film) 3299, 2932, 1685, 1585, 1508, 1453, 1368, 1251, 1162, 1144 $cm^{-1}$.

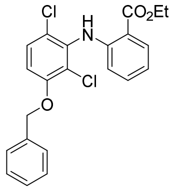

Prepared according to the above procedure; white solid (42 mg, 24%). TLC $R_f$ = 0.31 (19:1 hexanes/EtOAc); $^1$H NMR ($CDCl_3$, 500 MHz) δ 9.42 (s, 1H), 8.01 (ddd, *J* = 8.0, 1.7, 0.4 Hz, 1H), 7.49–7.27 (m, 7H), 6.85 (d, *J* = 9.0 Hz, 1H), 6.78 (ddd, *J* = 8.2, 7.1, 1.1 Hz, 1H), 6.38 (dd, *J* = 8.4, 1.1 Hz, 1H), 5.19 (s, 2H), 4.40 (q, *J* = 7.1 Hz, 2H), 1.43 (t, *J* = 7.1 Hz, 3H) ppm; IR (thin film) ν 3299, 1683, 1578, 1507, 1451, 1369, 1296, 1251, 1162 $cm^{-1}$.

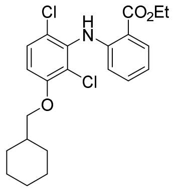

Prepared according to the above procedure; colorless film (16 mg, 10%). TLC $R_f$ = 0.43 (19:1 hexanes/EtOAc); $^1$H NMR ($CDCl_3$, 500 MHz) δ 9.38 (s, 1H), 8.00 (dd, *J* = 8.0, 1.6 Hz, 1H), 7.32 (d, *J* = 8.9 Hz, 1H), 7.27 (td, *J* = 8.5, 7.8, 1.6 Hz, 1H), 6.81–6.74 (m, 2H), 6.37 (d, *J* = 8.4 Hz, 1H), 4.40 (q, *J* = 7.1 Hz, 2H), 3.83 (d, *J* = 6.1 Hz, 2H), 1.92–1.84 (m, 3H), 1.81–1.68 (m, 4H), 1.42 (t, *J* = 7.1 Hz, 3H), 1.36–1.18 (m, 2H), 1.15–1.05 (m, 2H) ppm; IR (thin film) ν 3300, 2925, 2853, 1684, 1577, 1509, 1452, 1298, 1250, 1162 cm$^{-1}$.

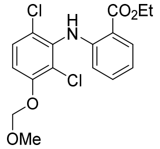

Prepared according to the above procedure; white solid (135 mg, 37%). TLC $R_f$ = 0.31 (19:1 hexanes/EtOAc); $^1$H NMR ($CDCl_3$, 400 MHz) δ 9.42 (s, 1H), 8.01 (dd, *J* = 8.0, 1.7 Hz, 1H), 7.33 (d, *J* = 9.0 Hz, 1H), 7.29 (ddd, *J* = 8.6, 7.1, 1.7 Hz, 1H), 7.09 (d, *J* = 9.0 Hz, 1H), 6.78 (ddd, *J* = 8.2, 7.1, 1.1 Hz, 1H), 6.37 (dd, *J* = 8.5, 1.0 Hz, 1H), 5.27 (s, 2H), 4.40 (q, *J* = 7.1 Hz, 2H), 3.54 (s, 3H), 1.43 (t, *J* = 7.1 Hz, 3H) ppm; IR (thin film) ν 3299, 2980, 1682, 1580, 1505, 1454, 1251, 1162 cm$^{-1}$.

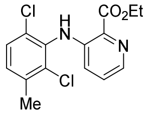

Prepared according to the above procedure. Reaction mixture was diluted with 1:9 MeOH/$CH_2Cl_2$ instead of EtOAc before filtering through Celite due to limited compound solubility in less polar solvents; white solid (45 mg, 14%). TLC $R_f$ = 0.10 (9:1 hexanes/EtOAc); $^1$H NMR ($CDCl_3$, 400 MHz) δ 9.40 (s, 1H), 8.17 (d, *J* = 4.3 Hz, 1H), 7.32 (d, *J* = 8.2 Hz, 1H), 7.21 (dd, *J* = 8.6, 4.3 Hz, 1H), 7.14 (d, *J* = 8.3 Hz, 1H), 6.71 (d, *J* = 8.6 Hz, 1H), 4.53 (q, *J* = 7.1 Hz, 2H), 2.41 (s, 3H), 1.49 (t, *J* = 7.1 Hz, 3H) ppm; IR (thin film) ν 3301, 2982, 1683, 1591, 1489, 1453, 1402, 1373, 1321, 1249, 1199 cm$^{-1}$.

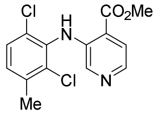

Prepared according to the above procedure. Reaction mixture was diluted with 1:9 MeOH/$CH_2Cl_2$ instead of EtOAc before filtering through Celite due to limited compound solubility in less polar solvents. In the filtration step, the ethyl ester transesterified to the methyl ester; pale yellow solid (48 mg, 26%). TLC $R_f$ = 0.42 (3:1 hexanes/EtOAc); $^1$H NMR ($CDCl_3$, 400 MHz) δ 9.00 (s, 1H), 8.07 (d, *J* = 5.2 Hz, 1H), 7.82 (s, 1H),

7.72 (d, $J$ = 5.1 Hz, 1H), 7.33 (d, $J$ = 8.2 Hz, 1H), 7.15 (d, $J$ = 8.2 Hz, 1H), 3.98 (s, 3H), 2.42 (s, 3H) ppm; IR (thin film) ν 3320, 2951, 1694, 1562, 1494, 1459, 1441, 1424, 1301, 1226, 1202, 1179 cm$^{-1}$.

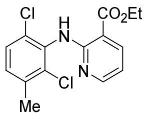

Prepared according to the above procedure. Reaction mixture was diluted with 1:9 MeOH/$CH_2Cl_2$ instead of EtOAc before filtering through Celite due to limited compound solubility in less polar solvents; pale yellow solid (72 mg, 36%). TLC $R_f$ = 0.65 (3:1 hexanes/EtOAc); $^1$H NMR ($CDCl_3$, 500 MHz) δ 9.56 (s, 1H), 8.30–8.20 (m, 2H), 7.30 (d, $J$ = 8.3 Hz, 1H), 7.11 (d, $J$ = 8.2 Hz, 1H), 6.71 (dd, $J$ = 7.7, 4.9 Hz, 1H), 4.42 (q, $J$ = 7.1 Hz, 2H), 2.40 (s, 3H), 1.43 (t, $J$ = 7.1 Hz, 3H) ppm; IR (thin film) ν 3313, 2984, 1679, 1584, 1487, 1458, 1382, 1295, 1244, 1141 cm$^{-1}$.

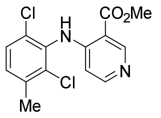

Prepared according to the above procedure. Reaction mixture was diluted with 1:9 MeOH/$CH_2Cl_2$ instead of EtOAc before filtering through Celite due to limited compound solubility in less polar solvents; off-white solid (7 mg, 5%). TLC $R_f$ = 0.20 (3:1 hexanes/EtOAc); $^1$H NMR ($CDCl_3$, 500 MHz) δ 9.60 (s, 1H), 9.03 (s, 1H), 8.23 (d, $J$ = 6.1 Hz, 1H), 7.34 (d, $J$ = 8.3 Hz, 1H), 7.20 (d, $J$ = 8.3 Hz, 1H), 6.16 (d, $J$ = 6.0 Hz, 1H), 3.98 (s, 3H), 2.42 (s, 3H) ppm; IR (thin film) ν 3288, 1692, 1596, 1572, 1502, 1459, 1316, 1226, 1113 cm$^{-1}$.

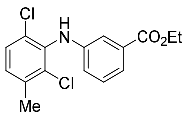

Prepared according to the above procedure; colorless oil (110 mg, 34%). TLC $R_f$ = 0.26 (19:1 hexanes/EtOAc); $^1$H NMR ($CDCl_3$, 500 MHz) δ 7.58 (ddd, $J$ = 7.7, 1.6, 1.0 Hz, 1H), 7.41 (dd, $J$ = 2.6, 1.6 Hz, 1H), 7.27 (td, $J$ = 8.0, 2.0 Hz, 2H), 7.05 (d, $J$ = 8.3 Hz, 1H), 6.81 (ddd, $J$ = 8.1, 2.5, 1.0 Hz, 1H), 5.88 (s, 1H), 4.35 (q, $J$ = 7.1 Hz, 2H), 2.40 (s, 3H), 1.37 (t, $J$ = 7.1 Hz, 3H) ppm; IR (thin film) ν 3353, 2981, 1715, 1608, 1590, 1510, 1484, 1462, 1392, 1368, 1290, 1231, 1106 cm$^{-1}$.

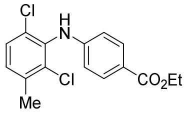

Prepared according to the above procedure; white solid (145 mg, 45%). TLC $R_f$ = 0.19 (19:1 hexanes/EtOAc); $^1$H NMR ($CDCl_3$, 500 MHz) δ 7.92 (d, $J$ = 8.6 Hz, 2H), 7.30 (d, $J$ = 8.3 Hz, 1H), 7.10 (d, $J$ = 8.3 Hz, 1H), 6.63 (d, $J$ = 8.6 Hz, 2H), 5.95 (s, 1H), 4.33 (q, $J$ = 7.1 Hz, 2H), 2.41 (s, 3H), 1.36 (t, $J$ = 7.1 Hz, 3H) ppm; IR (thin film) ν 3390, 3329, 2981, 1694, 1607, 1519, 1462, 1367, 1311, 1278, 1174, 1108 cm$^{-1}$.

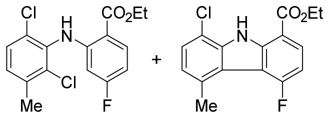

Prepared according to the above procedure; isolated as an inseparable mixture of products (~2.5:1 desired product/carbazole) and resolved by reversed-phase HPLC following ester hydrolysis; pale pink solid (184 mg); TLC $R_f$ = 0.64 (9:1 hexanes/EtOAc, single spot for both products); $^1$H NMR ($CDCl_3$, 400 MHz), see spectrum below; IR (thin film) ν 3427, 3293, 2982, 2930, 1687, 1617, 1591, 1513, 1461, 1258, 1128 cm$^{-1}$.

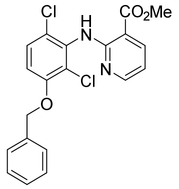

Prepared according to the above procedure; colorless residue (205 mg, 55%). TLC $R_f$ = 0.43 (4:1 hexanes/EtOAc); $^1$H NMR ($CDCl_3$, 500 MHz) δ 9.56 (s, 1H), 8.29–8.20 (m, 2H), 7.46 (d, *J* = 7.6 Hz, 2H), 7.39 (t, *J* = 7.4 Hz, 2H), 7.35–7.30 (m, 2H), 6.87 (d, *J* = 8.9 Hz, 1H), 6.75–6.69 (m, 1H), 5.17 (s, 2H), 3.96 (s, 3H) ppm; IR (thin film) ν 1737, 1693, 1577, 1495, 1452, 1396, 1292, 1245, 1137 cm$^{-1}$.

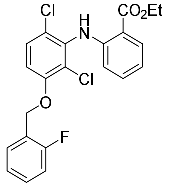

Prepared according to the above procedure; colorless residue (97 mg, 80%). TLC $R_f$ = 0.23 (19:1 hexanes/EtOAc); $^1$H NMR ($CDCl_3$, 500 MHz) δ 9.43 (s, 1H), 8.02 (dd, *J* = 8.0, 1.6 Hz, 1H), 7.59 (t, *J* = 7.6 Hz, 1H), 7.36–7.26 (m, 3H), 7.19 (t, *J* = 7.6 Hz, 1H), 7.12–7.07 (m, 1H), 6.90 (d, *J* = 9.0 Hz, 1H), 6.78 (t, *J* = 7.8 Hz, 1H), 6.38 (d, *J* = 8.3 Hz, 1H), 5.25 (s, 2H), 4.41 (q, *J* = 7.1 Hz, 2H), 1.43 (t, *J* = 7.1 Hz, 3H) ppm; $^{19}$F NMR ($CDCl_3$, 376 MHz) δ –119.10 (dt, *J* = 10.2, 6.6 Hz, 1F) ppm; IR (thin film) ν 3298, 2981, 1683, 1608, 1578, 1507, 1453, 1382, 1297, 1252, 1163, 1146 cm$^{-1}$.

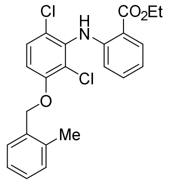

Prepared according to the above procedure; viscous, colorless oil that slowly solidified (82 mg, 48%). TLC $R_f$ = 0.19 (19:1 hexanes/$Et_2O$); $^1$H NMR ($CDCl_3$, 400 MHz) δ 9.43 (s, 1H), 8.00 (dd, *J* = 8.0, 1.7 Hz, 1H), 7.44 (d, *J* = 6.9 Hz, 1H), 7.31 (dd, *J* = 8.9, 0.5 Hz, 1H), 7.29–7.17 (m, 4H), 6.87 (d, *J* = 8.9 Hz, 1H), 6.80–6.72 (m, 1H), 6.37 (d, *J* = 8.4 Hz, 1H), 5.11 (s, 2H), 4.38 (q, *J* = 7.1 Hz, 2H), 2.38 (s, 3H), 1.41 (t, *J* = 7.1

Hz, 3H) ppm; IR (thin film) ν 3298, 2980, 1683, 1606, 1577, 1506, 1453, 1369, 1295, 1251, 1163, 1145 $cm^{-1}$.

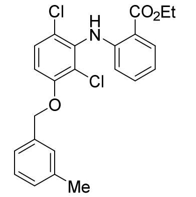

Prepared according to the above procedure; viscous, colorless oil (87 mg, 50%). TLC $R_f$ = 0.20 (19:1 hexanes/ $Et_2O$); $^1H$ NMR ($CDCl_3$, 400 MHz) δ 9.43 (s, 1H), 8.01 (dd, *J* = 8.0, 1.7 Hz, 1H), 7.33–7.22 (m, 5H), 7.14 (d, *J* = 7.0 Hz, 1H), 6.84 (d, *J* = 9.0 Hz, 1H), 6.77 (ddd, *J* = 8.0, 7.2, 0.9 Hz, 1H), 6.38 (d, *J* = 8.4 Hz, 1H), 5.13 (s, 2H), 4.40 (q, *J* = 7.1 Hz, 2H), 2.37 (s, 3H), 1.42 (t, *J* = 7.1 Hz, 3H) ppm; IR (thin film) ν 3298, 2980, 1684, 1606, 1578, 1508, 1452, 1369, 1298, 1250, 1162, 1145 $cm^{-1}$.

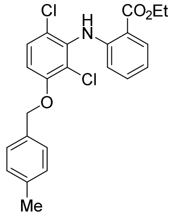

Prepared according to the above procedure; viscous, colorless oil that slowly solidified (85 mg, 49%). TLC $R_f$ = 0.19 (19:1 hexanes/ $Et_2O$); $^1H$ NMR ($CDCl_3$, 400 MHz) δ 9.42 (s, 1H), 8.01 (dd, *J* = 8.0, 1.7 Hz, 1H), 7.35 (d, *J* = 8.1 Hz, 2H), 7.32–7.25 (m, 2H), 7.20 (d, *J* = 8.2 Hz, 2H), 6.85 (d, *J* = 9.0 Hz, 1H), 6.77 (ddd, *J* = 8.2, 7.2, 1.1 Hz, 1H), 6.37 (d, *J* = 8.4 Hz, 1H), 5.15 (s, 2H), 4.40 (q, *J* = 7.1 Hz, 2H), 2.37 (s, 3H), 1.43 (t, *J* = 7.1 Hz, 3H) ppm; IR (thin film) ν 3298, 2981, 2926, 1683, 1605, 1578, 1507, 1452, 1369, 1314, 1296, 1250, 1163, 1145 $cm^{-1}$.

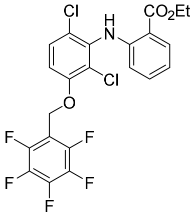

Prepared according to the above procedure; white solid (57 mg, 37%). TLC $R_f$ = 0.25 (9:1 hexanes/ $Et_2O$); $^1H$ NMR ($CDCl_3$, 400 MHz) δ 9.43 (s, 1H), 8.00 (dd, *J* = 8.0, 1.7 Hz, 1H), 7.39 (d, *J* = 8.9 Hz, 1H), 7.31–7.25 (m, 1H), 6.95 (d, *J* = 9.0 Hz, 1H), 6.78 (ddd, *J* = 8.0, 7.2, 1.1 Hz, 1H), 6.34 (d, *J* = 8.5 Hz, 1H), 5.20 (s, 2H), 4.40 (q, *J* = 7.1 Hz, 2H), 1.42 (t, *J* = 7.1 Hz, 3H) ppm; $^{19}F$ NMR ($CDCl_3$, 376 MHz) δ –142.23 (dd, *J* = 22.3, 8.6 Hz, 2F), –152.13 (t, *J* = 20.8 Hz, 1F), –161.38 to –161.60 (m, 2F) ppm; IR (thin film) ν 3299, 2984, 1685, 1658, 1581, 1508, 1453, 1383, 1312, 1292, 1251, 1163, 1134 $cm^{-1}$.

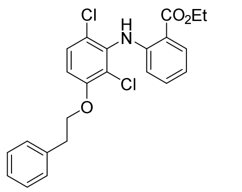

Prepared according to the above procedure; viscous, colorless oil (67 mg, 39%). TLC $R_f$ = 0.29 (9:1 hexanes/$Et_2O$); $^1H$ NMR ($CDCl_3$, 400 MHz) δ 9.39 (s, 1H), 8.00 (dd, *J* = 8.0, 1.7 Hz, 1H), 7.35–7.20 (m, 7H), 6.79–6.72 (m, 2H), 6.34 (d, *J* = 8.4 Hz, 1H), 4.38 (q, *J* = 7.1 Hz, 2H), 4.21 (t, *J* = 6.9 Hz, 2H), 3.15 (t, *J* = 6.9 Hz, 2H), 1.41 (t, *J* = 7.1 Hz, 3H) ppm; IR (thin film) ν 3299, 3028, 2980, 1683, 1605, 1578, 1508, 1453, 1388, 1368, 1296, 1251, 1163, 1145 $cm^{-1}$.

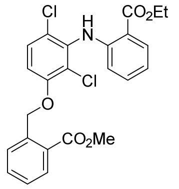

Prepared according to the above procedure; off-white solid (22 mg, 30%). TLC $R_f$ = 0.17 (19:1 hexanes/EtOAc); $^1H$ NMR ($CDCl_3$, 500 MHz) δ 9.44 (s, 1H), 8.04 (dd, *J* = 17.5, 7.9 Hz, 2H), 7.90 (d, *J* = 7.9 Hz, 1H), 7.61 (t, *J* = 7.7 Hz, 1H), 7.40 (t, *J* = 7.7 Hz, 1H), 7.35–7.28 (m, 2H), 6.92 (d, *J* = 8.9 Hz, 1H), 6.78 (t, *J* = 7.6 Hz, 1H), 6.40 (d, *J* = 8.4 Hz, 1H), 5.61 (s, 2H), 4.41 (q, *J* = 7.2 Hz, 2H), 3.93 (s, 3H), 1.43 (t, *J* = 7.0 Hz, 3H) ppm; IR (thin film) ν 3299, 2981, 2951, 1718, 1684, 1579, 1509, 1452, 1300, 1251, 1139 $cm^{-1}$.

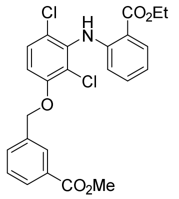

Prepared according to the above procedure; white solid (16 mg, 27%). TLC $R_f$ = 0.59 (3:1 hexanes/EtOAc); $^1H$ NMR ($CDCl_3$, 400 MHz) δ 9.43 (s, 1H), 8.13 (s, 1H), 8.07–7.98 (m, 2H), 7.70 (d, *J* = 7.7 Hz, 1H), 7.49 (t, *J* = 7.7 Hz, 1H), 7.34–7.26 (m, 2H), 6.85 (d, *J* = 8.9 Hz, 1H), 6.78 (t, *J* = 7.5 Hz, 1H), 6.37 (d, *J* = 8.4 Hz, 1H), 5.22 (s, 2H), 4.41 (q, *J* = 7.2 Hz, 2H), 3.93 (s, 3H), 1.43 (t, *J* = 7.2 Hz, 3H) ppm; IR (thin film) ν 3446, 1727, 1693, 1606, 1505, 1431, 1273, 1208, 1181, 1146 $cm^{-1}$.

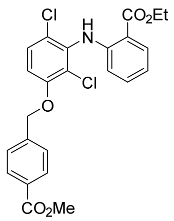

Prepared according to the above procedure; white solid (111 mg, 75%). TLC $R_f$ = 0.48 (3:1 hexanes/EtOAc); $^1$H NMR ($CDCl_3$, 500 MHz) δ 9.44 (s, 1H), 8.07 (d, *J* = 8.2 Hz, 2H), 8.01 (dd, *J* = 8.2, 1.7 Hz, 1H), 7.55 (d, *J* = 8.2 Hz, 2H), 7.32 (d, *J* = 9.0 Hz, 1H), 7.31–7.28 (m, 1H), 6.82 (d, *J* = 9.0 Hz, 1H), 6.78 (ddd, *J* = 8.2, 7.2, 1.1 Hz, 1H), 6.37 (d, *J* = 8.3 Hz, 1H), 5.24 (s, 2H), 4.41 (q, *J* = 7.1 Hz, 2H), 3.93 (s, 3H), 1.43 (t, *J* = 7.1 Hz, 3H) ppm; IR (thin film) ν 3299, 2951, 1722, 1683, 1578, 1508, 1451, 1280, 1251, 1109 cm$^{-1}$.

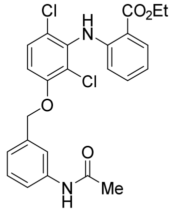

Prepared according to the above procedure. Reaction mixture was diluted with 1:4 MeOH/$CH_2Cl_2$ instead of EtOAc before filtering through Celite due to limited compound solubility in less polar solvents; white solid (33 mg, 24%). TLC $R_f$ = 0.17 (49:1 $CH_2Cl_2$/MeOH); $^1$H NMR ($CDCl_3$, 500 MHz) δ 9.42 (s, 1H), 8.01 (dd, *J* = 8.0, 1.7 Hz, 1H), 7.61 (s, 1H), 7.49 (d, *J* = 8.0 Hz, 1H), 7.38–7.27 (m, 3H), 7.21 (d, *J* = 6.9 Hz, 2H), 6.83 (d, *J* = 9.0 Hz, 1H), 6.78 (ddd, *J* = 8.1, 7.1, 1.1 Hz, 1H), 6.38 (dd, *J* = 8.5, 1.0 Hz, 1H), 5.17 (s, 2H), 4.40 (q, *J* = 7.1 Hz, 2H), 2.19 (s, 3H), 1.43 (t, *J* = 7.1 Hz, 3H) ppm; IR (thin film) ν 3302, 1678, 1561, 1487, 1451, 1298, 1249 cm$^{-1}$.

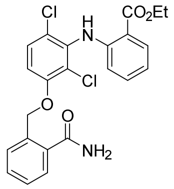

Prepared according to the above procedure with 15 mol % catalyst. Reaction mixture was diluted with 1:4 MeOH/$CH_2Cl_2$ instead of EtOAc before filtering through Celite due to limited compound solubility in less polar solvents; white solid (59 mg, 50%). TLC $R_f$ = 0.22 (49:1 hexanes/EtOAc); $^1$H NMR ($CDCl_3$, 500 MHz) δ 11.74 (s, 1H), 8.83 (d, *J* = 8.4 Hz, 1H), 8.10 (d, *J* = 8.0 Hz, 1H), 7.85 (d, *J* = 7.8 Hz, 1H), 7.78 (d, *J* = 7.6 Hz, 1H), 7.60 (t, *J* = 7.9 Hz, 1H), 7.55 (t, *J* = 7.7 Hz, 1H), 7.46 (t, *J* = 7.5 Hz, 1H), 7.15 (t, *J* = 7.8 Hz, 1H), 7.05 (d, *J* = 8.9 Hz, 1H), 6.41 (d, *J* = 8.9 Hz, 1H), 5.52 (s, 2H), 4.46 (s, 2H), 4.38 (q, *J* = 7.1 Hz, 2H), 1.41 (t, *J* = 7.1 Hz, 3H) ppm; IR (thin film) ν 3299, 1676, 1607, 1531, 1509, 1474, 1448, 1313, 1251 cm$^{-1}$.

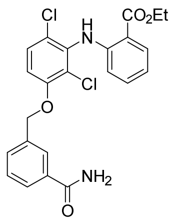

Prepared according to the above procedure with 15 mol % catalyst. Reaction mixture was diluted with 1:4 MeOH/$CH_2Cl_2$ instead of EtOAc before filtering through Celite due to limited compound solubility in less polar solvents; white solid (58 mg, 49%). TLC $R_f$ = 0.16 (9:1 hexanes/EtOAc); $^1$H NMR ($CDCl_3$, 500 MHz) δ 12.13 (s, 1H), 8.93 (d, *J* = 8.5 Hz, 1H), 8.14–8.07 (m, 2H), 7.99 (d, *J* = 7.9 Hz, 1H), 7.67 (d, *J* = 7.8 Hz, 1H), 7.61 (t, *J* = 7.8 Hz, 1H), 7.55 (t, *J* = 7.7 Hz, 1H), 7.13 (t, *J* = 7.6 Hz, 1H), 7.08 (d, *J* = 8.9 Hz, 1H), 6.36 (d, *J* = 8.9 Hz, 1H), 5.21 (s, 2H), 4.52 (s, 2H), 4.42 (q, *J* = 7.1 Hz, 2H), 1.43 (t, *J* = 7.1 Hz, 3H) ppm; IR (thin film) ν 3377, 1675, 1608, 1589, 1531, 1475, 1449, 1308, 1254 $cm^{-1}$.

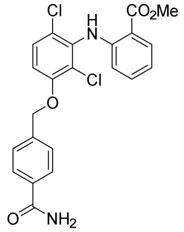

Prepared according to the above procedure. Reaction mixture was diluted with 1:4 MeOH/$CH_2Cl_2$ instead of EtOAc before filtering through Celite due to limited compound solubility in less polar solvents. In the filtration step, the ethyl ester transesterified to the methyl ester; white solid (8 mg, 7%). TLC $R_f$ = 0.52 (49:1 $CH_2Cl_2$/MeOH); $^1$H NMR (DMSO-$d_6$, 500 MHz) δ 11.60 (s, 1H), 8.55 (d, *J* = 8.2 Hz, 1H), 8.02 (dd, *J* = 7.9, 1.7 Hz, 1H), 7.99 (d, *J* = 8.1 Hz, 2H), 7.72–7.64 (m, 3H), 7.26 (t, *J* = 7.7 Hz, 1H), 7.18 (d, *J* = 8.9 Hz, 1H), 6.49 (d, *J* = 8.9 Hz, 1H), 5.53 (s, 2H), 5.27 (s, 2H), 3.90 (s, 3H) ppm; IR (thin film) ν 3309, 2920, 1679, 1609, 1590, 1535, 1477, 1450, 1386, 1312, 1251 $cm^{-1}$.

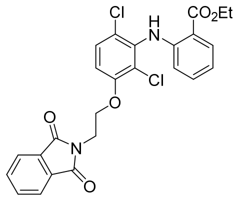

Prepared according to the above procedure; colorless oil (109 mg, 97%). TLC $R_f$ = 0.58 (3:2 hexanes/EtOAc); $^1$H NMR ($CDCl_3$, 400 MHz) δ 9.34 (s, 1H), 7.98 (dd, *J* = 8.0, 1.7 Hz, 1H), 7.85 (dd, *J* = 5.5, 3.0 Hz, 2H), 7.71 (dd, *J* = 5.5, 3.0 Hz, 2H), 7.31 (d, *J* = 8.9 Hz, 1H), 7.26–7.20 (m, 1H), 6.82 (d, *J* = 9.0 Hz, 1H), 6.75 (ddd, *J* = 8.1, 7.1, 1.1 Hz, 1H), 6.32 (d, *J* = 8.4 Hz, 1H), 4.37 (q, *J* = 7.1 Hz, 2H), 4.31 (t, *J* = 5.6 Hz, 2H), 4.18–4.17 (m, 2H), 1.41 (t, *J* = 7.1 Hz, 3H) ppm; IR (thin film) ν 3299, 2982, 1775, 1715, 1683, 1579, 1507, 1453, 1394, 1297, 1251 $cm^{-1}$.

*MOM group deprotection*

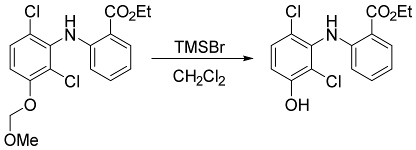

A flame-dried 5-mL round-bottom flask was charged with ethyl 2-((2,6-dichloro-3-(methoxymethoxy)phenyl)amino)benzoate (25.5 mg, 0.069 mmol). The flask was sealed with a septum, and the headspace flushed with $N_2$, and 2.0 mL of anhydrous $CH_2Cl_2$ was added. Following dissolution of the starting material, the flask was placed in an ice bath, and bromotrimethylsilane (0.05 mL, 0.379 mmol, 5.5 equiv) was added dropwise via syringe. The mixture was stirred at 0 °C for 1 h, following which time the reaction was warmed to ambient temperature and stirred for an additional 3 h. The reaction was quenched by the addition of 1 mL of $H_2O$ and transferred to a separatory funnel with 20 mL of EtOAc. The organic fraction was washed with 2 x 20 mL of $H_2O$ and 1 x 20 mL of saturated aqueous NaCl, dried over $Na_2SO_4$, filtered, and concentrated under reduced pressure to give a white solid (19 mg, 85%). This material was judged to be sufficiently pure by $^1$H NMR for use in the subsequent step without further purification. TLC $R_f$ = 0.16 (9:1 hexanes/acetone); $^1$H NMR ($CDCl_3$, 400 MHz) δ 9.40 (s, 1H), 8.01 (d, *J* = 8.1 Hz, 1H), 7.36–7.22 (m, 2H), 6.94 (d, *J* = 8.9 Hz, 1H), 6.79 (t, *J* = 7.5 Hz, 1H), 6.37 (d, *J* = 8.5 Hz, 1H), 5.61 (s, 1H), 4.40 (q, *J* = 7.1 Hz, 2H), 1.43 (t, *J* = 7.1 Hz, 3H) ppm; IR (thin film) v 3308, 2926, 1683, 1580, 1506, 1452, 1253 $cm^{-1}$.

*Ester hydrolysis*

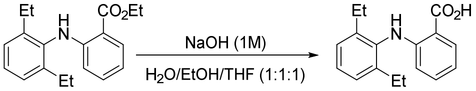

**AK-21**. A screw-capped vial was charged with ethyl 2-((2,6-diethylphenyl)amino)benzoate (180 mg, 0.6 mmol) and 3.0 mL of a 1.0 M solution of NaOH in 1:1:1 mixture of $H_2O$/EtOH/THF. The reaction was stirred until thin-layer chromatography indicated the complete consumption of starting material (~24 h). Following this time, the solution was acidified to pH 2 with 2–3 drops of concentrated aqueous HCl, which resulted in the immediate precipitation of a white solid. The solid product was isolated by vacuum filtration through a small Büchner funnel and purified by reversed-phase HPLC (Alltima C18, 10 μM, 22 x 250 mm column, eluting with a gradient flow of 0:100→70:30 MeCN/0.1% TFA in $H_2O$ over 10 min, then 70:30–100:0 MeCN/0.1% TFA over 25 min, 254 nm UV detection, flow rate = 12.0 mL/min, $R_T$ = 23.3–25.2 min = 86–88% MeCN); white solid (32 mg, 20%); $^1$H NMR (acetone-$d_6$, 500 MHz) δ 9.32 (s, 1H), 7.99 (dd, *J* = 8.1, 1.7 Hz, 1H), 7.29–7.17 (m, 4H), 6.65 (t, *J* = 7.5 Hz, 1H), 6.16 (d, *J* = 8.5 Hz, 1H), 2.54 (dq, *J* = 14.6, 7.2 Hz, 4H), 1.10 (t, *J* = 7.6 Hz, 6H) ppm. LRMS ($ES^-$) calcd 268.13 for $C_{17}H_{18}NO_2^-$ found 267.7 ($M^-$).

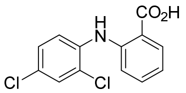

**AK-1**. Prepared according to the above procedure. Purification by reversed-phase HPLC (Alltima C18, 10 μM, 22 x 250 mm column, eluting with a gradient flow over 30 min of 0:1→1:0 MeCN/0.1% TFA in $H_2O$, 254 nm UV detection, flow rate = 12.0 mL/min, $R_T$ = 30.6–31.6 min); white solid (5 mg, 26%); $^1$H NMR (DMSO-d$_6$, 600 MHz) δ 9.87 (s, 1H), 7.96 (d, *J* = 8.0 Hz, 1H), 7.70 (d, *J* = 2.5 Hz, 1H), 7.57 (d, *J* = 8.7 Hz, 1H), 7.49–7.44 (m, 1H), 7.39 (dd, *J* = 8.8, 2.4 Hz, 1H), 7.23 (d, *J* = 8.4 Hz, 1H), 6.92 (t, *J* = 7.4 Hz, 1H) ppm; LRMS (ES$^-$) calcd 279.99 for $C_{13}H_8Cl_2NO_2^-$ found 280.0 (M$^-$).

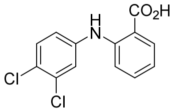

**AK-2**. Prepared according to the above procedure. Purification by reversed-phase HPLC (Alltima C18, 10 μM, 22 x 250 mm column, eluting with a gradient flow over 30 min of 0:1→1:0 MeCN/0.1% TFA in $H_2O$, 254 nm UV detection, flow rate = 12.0 mL/min, $R_T$ = 30.8–31.9 min); white solid (24 mg, 63%); $^1$H NMR (DMSO-d$_6$, 600 MHz) δ 9.64 (s, 1H), 7.93 (d, *J* = 7.8 Hz, 1H), 7.53 (d, *J* = 8.7 Hz, 1H), 7.48–7.44 (m, 2H), 7.30 (d, *J* = 8.4 Hz, 1H), 7.23 (dd, *J* = 9.0, 2.4 Hz, 1H), 6.90 (t, *J* = 7.6 Hz, 1H) ppm; LRMS (ES$^-$) calcd 279.99 for $C_{13}H_8Cl_2NO_2^-$ found 280.0 (M$^-$).

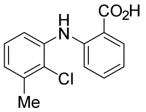

**AK-3**. Prepared according to the above procedure. Purification by reversed-phase HPLC (Alltima C18, 10 μM, 22 x 250 mm column, eluting with a gradient flow over 30 min of 0:1→1:0 MeCN/0.1% TFA in $H_2O$, 254 nm UV detection, flow rate = 12.0 mL/min, $R_T$ = 29.5–30.7 min); white solid (4 mg, yield not determined); $^1$H NMR (DMSO-d$_6$, 600 MHz) δ 9.83 (s, 1H), 7.93 (d, *J* = 7.9 Hz, 1H), 7.43 (t, *J* = 7.8 Hz, 1H), 7.38 (d, *J* = 8.0 Hz, 1H), 7.22 (t, *J* = 7.8 Hz, 1H), 7.19 (d, *J* = 8.5 Hz, 1H), 7.06 (d, *J* = 7.4 Hz, 1H), 6.86 (t, *J* = 7.5 Hz, 1H), 2.37 (s, 3H) ppm; LRMS (ES$^-$) calcd 260.05 for $C_{14}H_{11}ClNO_2^-$ found 260.0 (M$^-$).

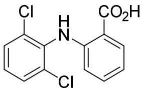

**AK-4**. Prepared according to the above procedure. Purification by reversed-phase HPLC (Alltima C18, 10 μM, 22 x 250 mm column, eluting with a gradient flow over 30 min of 0:1→1:0 MeCN/0.1% TFA in $H_2O$, 254 nm UV detection, flow rate = 12.0 mL/min, $R_T$ = 27.8–28.9 min); white solid (12 mg, 49%); $^1$H NMR (DMSO-d$_6$, 600 MHz) δ 9.51 (s, 1H), 7.90 (d, *J* = 7.6 Hz, 1H), 7.63 (d, *J* = 8.1 Hz, 2H), 7.37 (t, *J* = 8.1 Hz, 1H), 7.33 (t, *J* = 7.8 Hz, 1H), 6.79 (t, *J* = 7.4 Hz, 1H), 6.22 (d, *J* = 8.1 Hz, 1H) ppm; LRMS (ES$^-$) calcd 279.99 for $C_{13}H_8Cl_2NO_2^-$ found 280.0 (M$^-$).

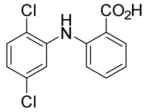

**AK-5**. Prepared according to the above procedure. Purification by reversed-phase HPLC (Alltima C18, 10 µM, 22 x 250 mm column, eluting with a gradient flow over 30 min of 0:1→1:0 MeCN/0.1% TFA in $H_2O$, 254 nm UV detection, flow rate = 12.0 mL/min, $R_T$ = 30.3–32.0 min); white solid (21 mg, 38%); $^1$H NMR (DMSO-$d_6$, 600 MHz) δ 9.90 (s, 1H), 7.96 (d, *J* = 8.0 Hz, 1H), 7.58–7.48 (m, 3H), 7.32 (d, *J* = 8.4 Hz, 1H), 7.10 (dd, *J* = 8.7, 2.5 Hz, 1H), 6.96 (t, *J* = 7.2 Hz, 1H) ppm; LRMS ($ES^-$) calcd 279.99 for $C_{13}H_8Cl_2NO_2^-$ found 280.0 ($M^-$).

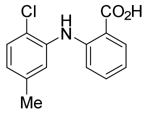

**AK-6**. Prepared according to the above procedure. Purification by reversed-phase HPLC (Alltima C18, 10 µM, 22 x 250 mm column, eluting with a gradient flow over 30 min of 0:1→1:0 MeCN/0.1% TFA in $H_2O$, 254 nm UV detection, flow rate = 12.0 mL/min, $R_T$ = 29.4–30.6 min); white solid (23 mg, 48%); $^1$H NMR (DMSO-$d_6$, 600 MHz) δ 9.77 (s, 1H), 7.93 (dd, *J* = 8.0, 1.7 Hz, 1H), 7.45 (ddd, *J* = 8.7, 7.1, 1.7 Hz, 1H), 7.41 (d, *J* = 8.1 Hz, 1H), 7.37 (d, *J* = 2.1 Hz, 1H), 7.21 (dd, *J* = 8.5, 1.0 Hz, 1H), 6.91 (dd, *J* = 8.2, 2.1 Hz, 1H), 6.87 (ddd, *J* = 8.1, 7.1, 1.1 Hz, 1H), 2.29 (s, 3H) ppm; LRMS ($ES^-$) calcd 260.05 for $C_{14}H_{11}ClNO_2^-$ found 260.0 ($M^-$).

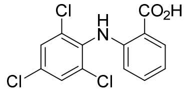

**AK-7**. Prepared according to the above procedure. Purification by reversed-phase HPLC (Alltima C18, 10 µM, 22 x 250 mm column, eluting with a gradient flow over 30 min of 0:1→1:0 MeCN/0.1% TFA in $H_2O$, 254 nm UV detection, flow rate = 12.0 mL/min, $R_T$ = 30.7–31.6 min); white solid (9 mg, 17%); $^1$H NMR (DMSO-$d_6$, 600 MHz) δ 9.48 (s, 1H), 7.89 (dd, *J* = 7.9, 1.7 Hz, 1H), 7.82 (s, 2H), 7.32 (ddd, *J* = 8.6, 7.1, 1.7 Hz, 1H), 6.79 (t, *J* = 7.7 Hz, 1H), 6.24 (d, *J* = 8.4 Hz, 1H) ppm; LRMS ($ES^-$) calcd 313.95 for $C_{13}H_7Cl_3NO_2^-$ found 315.9 ($M^-$).

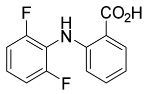

**AK-8**. Prepared according to the above procedure. Purification by reversed-phase HPLC (Alltima C18, 10 µM, 22 x 250 mm column, eluting with a gradient flow over 30 min of 0:1→1:0 MeCN/0.1% TFA in $H_2O$, 254 nm UV detection, flow rate = 12.0 mL/min, $R_T$ = 26.0–27.6 min); white solid (32 mg, 31%); $^1$H NMR (DMSO-$d_6$, 600 MHz) δ 9.31 (s, 1H), 7.90 (dd, *J* = 8.0, 1.7 Hz, 1H), 7.41–7.31 (m, 2H), 7.25 (t, *J* = 8.3 Hz,

2H), 6.81 (t, *J* = 7.6 Hz, 1H), 6.49 (d, *J* = 8.2 Hz, 1H) ppm; LRMS (ES$^-$) calcd 248.05 for $C_{13}H_8F_2NO_2^-$ found 248.0 (M$^-$).

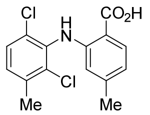

**AK-10**. Prepared according to the above procedure. Purification by reversed-phase HPLC (Alltima C18, 10 μM, 22 x 250 mm column, eluting with a gradient flow over 30 min of 0:1→1:0 MeCN/0.1% TFA in $H_2O$, 254 nm UV detection, flow rate = 12.0 mL/min, $R_T$ = 30.4–30.6 min); white solid (10 mg, 37%); $^1$H NMR (DMSO-d$_6$, 600 MHz) δ 9.51 (s, 1H), 7.78 (d, *J* = 8.1 Hz, 1H), 7.51 (d, *J* = 8.3 Hz, 1H), 7.35 (d, *J* = 8.3 Hz, 1H), 6.60 (dd, *J* = 8.0, 1.1 Hz, 1H), 5.99 (s, 1H), 2.38 (s, 3H), 2.15 (s, 3H) ppm; LRMS (ES$^-$) calcd 308.03 for $C_{15}H_{12}Cl_2NO_2^-$ found 308.1 (M$^-$).

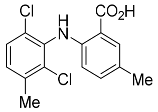

**AK-11**. Prepared according to the above procedure. Purification by reversed-phase HPLC (Alltima C18, 10 μM, 22 x 250 mm column, eluting with a gradient flow over 30 min of 0:1→1:0 MeCN/0.1% TFA in $H_2O$, 254 nm UV detection, flow rate = 12.0 mL/min, $R_T$ = 30.8–32.0 min); white solid (7 mg, 24%); $^1$H NMR (DMSO-d$_6$, 600 MHz) δ 9.37 (s, 1H), 7.70 (d, *J* = 2.1 Hz, 1H), 7.50 (d, *J* = 8.3 Hz, 1H), 7.33 (d, *J* = 8.4 Hz, 1H), 7.13 (dd, *J* = 8.2, 2.1 Hz, 1H), 6.54 (s, 1H), 6.12 (d, *J* = 8.5 Hz, 1H), 2.37 (s, 3H), 2.21 (s, 3H) ppm; LRMS (ES$^-$) calcd 308.03 for $C_{15}H_{12}Cl_2NO_2^-$ found 308.0 (M$^-$).

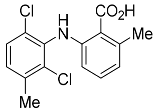

**AK-12**. Prepared according to the above procedure. Purification by reversed-phase HPLC (Alltima C18, 10 μM, 22 x 250 mm column, eluting with a gradient flow over 30 min of 0:1→1:0 MeCN/0.1% TFA in $H_2O$, 254 nm UV detection, flow rate = 12.0 mL/min, $R_T$ = 29.9–30.6 min); white solid (2 mg, 13%); $^1$H NMR (DMSO-d$_6$, 600 MHz) δ 8.33 (s, 1H), 7.47 (d, *J* = 8.3 Hz, 1H), 7.29 (d, *J* = 8.3 Hz, 1H), 7.09 (t, *J* = 7.9 Hz, 1H), 6.67 (d, *J* = 7.5 Hz, 1H), 6.53 (s, 1H), 6.05 (d, *J* = 8.3 Hz, 1H), 2.42 (s, 3H), 2.37 (s, 3H) ppm; LRMS (ES$^-$) calcd 308.03 for $C_{15}H_{12}Cl_2NO_2^-$ found 308.1 (M$^-$).

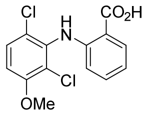

**AK-13**. Prepared according to the above procedure. Purification by reversed-phase HPLC (Alltima C18, 10 μM, 22 x 250 mm column, eluting with a gradient flow over 30 min of 0:1→1:0 MeCN/0.1% TFA in $H_2O$, 254 nm UV detection, flow rate = 12.0 mL/min, $R_T$ = 27.6–28.2 min); white solid (2 mg, 24%); $^1$H NMR

(DMSO-d6, 600 MHz) δ 9.50 (s, 1H), 7.89 (dd, *J* = 7.8, 1.6 Hz, 1H), 7.57 (d, *J* = 9.1 Hz, 1H), 7.35–7.30 (m, 1H), 7.17 (d, *J* = 9.1 Hz, 1H), 6.78 (t, *J* = 7.7 Hz, 1H), 6.23 (d, *J* = 8.5 Hz, 1H), 3.91 (s, 3H) ppm; LRMS (ES⁻) calcd 310.00 for $C_{14}H_{10}Cl_2NO_3^-$ found 310.1 ($M^-$).

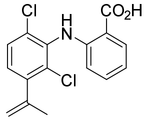

**AK-14**. Prepared according to the above procedure. Purification by reversed-phase HPLC (Alltima C18, 10 μM, 22 x 250 mm column, eluting with a gradient flow over 30 min of 0:1→1:0 MeCN/0.1% TFA in $H_2O$, 254 nm UV detection, flow rate = 12.0 mL/min, $R_T$ = 31.0–31.5 min); white solid (1 mg, 7%); $^1H$ NMR (DMSO-d6, 600 MHz) δ 9.56 (s, 1H), 7.90 (dd, *J* = 7.9, 1.6 Hz, 1H), 7.59 (dd, *J* = 8.4, 0.9 Hz, 1H), 7.34 (t, *J* = 7.6 Hz, 1H), 7.26 (d, *J* = 8.3 Hz, 1H), 6.78 (t, *J* = 7.5 Hz, 1H), 6.24 (d, *J* = 8.4 Hz, 1H), 5.31 (t, *J* = 1.7 Hz, 1H), 5.01 (s, 1H), 2.07 (s, 3H) ppm. LRMS ($ES^-$) calcd 320.03 for $C_{16}H_{12}Cl_2NO_2^-$ found 320.0 ($M^-$).

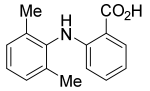

**AK-15**. Prepared according to the above procedure. Purification by reversed-phase HPLC (Alltima C18, 10 μM, 22 x 250 mm column, eluting with a gradient flow over 30 min of 0:1→1:0 MeCN/0.1% TFA in $H_2O$, 254 nm UV detection, flow rate = 12.0 mL/min, $R_T$ = 29.5–30.2 min); white solid (1 mg, 2%); $^1H$ NMR (DMSO-d6, 600 MHz) δ 9.16 (s, 1H), 7.86 (dd, *J* = 8.0, 1.7 Hz, 1H), 7.25 (t, *J* = 7.5 Hz, 1H), 7.20–7.12 (m, 3H), 6.64 (t, *J* = 7.5 Hz, 1H), 6.53 (s, 1H), 6.07 (d, *J* = 8.5 Hz, 1H), 2.12 (s, 6H) ppm; LRMS ($ES^-$) calcd 240.11 for $C_{15}H_{14}NO_2^-$ found 240.2 ($M^-$).

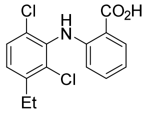

**AK-16**. Prepared according to the above procedure. Purification by reversed-phase HPLC (Alltima C18, 10 μM, 22 x 250 mm column, eluting with a gradient flow over 30 min of 0:1→1:0 MeCN/0.1% TFA in $H_2O$, 254 nm UV detection, flow rate = 12.0 mL/min, $R_T$ = 30.8–32.2 min); white solid (32 mg, 39%); $^1H$ NMR (DMSO-d6, 600 MHz) δ 9.53 (s, 1H), 7.90 (dd, *J* = 8.0, 1.6 Hz, 1H), 7.55 (d, *J* = 8.4 Hz, 1H), 7.35 (d, *J* = 8.4 Hz, 1H), 7.34–7.30 (m, 1H), 6.77 (ddd, *J* = 8.1, 7.1, 1.1 Hz, 1H), 6.19 (dd, *J* = 8.4, 1.1 Hz, 1H), 2.75 (q, *J* = 7.4 Hz, 2H), 1.20 (t, *J* = 7.5 Hz, 3H) ppm; LRMS ($ES^-$) calcd 308.03 for $C_{15}H_{12}Cl_2NO_2^-$ found 308.0 ($M^-$).

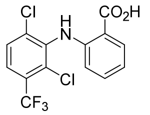

**AK-17**. Prepared according to the above procedure. Purification by reversed-phase HPLC (Alltima C18, 10 μM, 22 x 250 mm column, eluting with a gradient flow over 30 min of 0:1→1:0 MeCN/0.1% TFA in $H_2O$, 254 nm UV detection, flow rate = 12.0 mL/min, $R_T$ = 33.2–34.2 min); white solid (2 mg, 9%); $^1H$ NMR (acetonitrile-$d_3$, 500 MHz) δ 9.50 (s, 1H), 7.99 (dd, *J* = 7.9, 1.6 Hz, 1H), 7.77–7.63 (m, 2H), 7.34 (ddd, *J* = 8.6, 7.2, 1.7 Hz, 1H), 6.84 (ddd, *J* = 8.1, 7.1, 1.1 Hz, 1H), 6.30 (dd, *J* = 8.5, 1.1 Hz, 1H) ppm; LRMS (ES$^-$) calcd 347.98 for $C_{14}H_7Cl_2F_3NO_2^-$ found 348.0 (M$^-$).

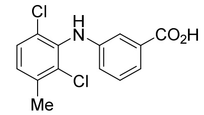

**AK-19**. Prepared according to the above procedure. Purification by reversed-phase HPLC (Alltima C18, 10 μM, 22 x 250 mm column, eluting with a gradient flow of 0:100→70:30 MeCN/0.1% TFA in $H_2O$ over 10 min, then 70:30→100:0 MeCN/0.1% TFA over 25 min, 254 nm UV detection, flow rate = 12.0 mL/min, $R_T$ = 17.1–17.6 min = 79% MeCN); white solid (32 mg, 29%); $^1H$ NMR (DMSO-$d_6$, 500 MHz) δ 8.11 (s, 1H), 7.47 (d, *J* = 8.3 Hz, 1H), 7.31–7.27 (m, 2H), 7.24 (t, *J* = 7.8 Hz, 1H), 7.08 (s, 1H), 6.74 (dt, *J* = 8.0, 1.2 Hz, 1H), 2.37 (s, 3H) ppm. LRMS (ES$^-$) calcd 294.01 for $C_{14}H_{10}Cl_2NO_2^-$ found 293.6 (M$^-$).

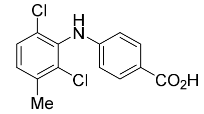

**AK-20**. Prepared according to the above procedure. Purification by reversed-phase HPLC (Alltima C18, 10 μM, 22 x 250 mm column, eluting with a gradient flow of 0:100→70:30 MeCN/0.1% TFA in $H_2O$ over 10 min, then 70:30→100:0 MeCN/0.1% TFA over 25 min, 254 nm UV detection, flow rate = 12.0 mL/min, $R_T$ = 16.7–17.8 min = 78–79% MeCN); white solid (12 mg, 9%); $^1H$ NMR (acetone-$d_6$, 500 MHz) δ 7.86 (d, *J* = 8.7 Hz, 2H), 7.69 (s, 1H), 7.45 (d, *J* = 8.3 Hz, 1H), 7.32 (d, *J* = 8.3 Hz, 1H), 6.66 (d, *J* = 8.7 Hz, 2H), 2.41 (s, 3H) ppm. LRMS (ES$^-$) calcd 294.01 for $C_{14}H_{10}Cl_2NO_2^-$ found 293.5 (M$^-$).

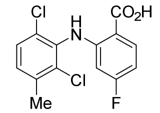

**AK-22**. Prepared according to the above procedure. Purification by reversed-phase HPLC (Alltima C18, 10 μM, 22 x 250 mm column, eluting with a gradient flow of 0:100→70:30 MeCN/0.1% TFA in $H_2O$ over 10 min, then 70:30→100:0 MeCN/0.1% TFA over 25 min, 254 nm UV detection, flow rate = 12.0 mL/min, $R_T$ = 21.6–23.2 min = 84–86% MeCN); white solid (55 mg, 45%); $^1H$ NMR (DMSO-$d_6$, 500 MHz) δ 9.71 (s, 1H), 7.97 (t, *J* = 7.8 Hz, 1H), 7.53 (d, *J* = 8.3 Hz, 1H), 7.38 (d, *J* = 8.3 Hz, 1H), 6.59 (td, *J* = 8.5, 2.3 Hz, 1H), 5.87 (dd, *J* = 11.7, 2.1 Hz, 1H), 2.38 (s, 3H) ppm. LRMS (ES$^-$) calcd 312.00 for $C_{14}H_9Cl_2FNO_2^-$ found 311.6 (M$^-$).

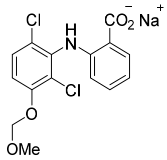

**AK-23**. Prepared according to the above procedure. The solution containing the sodium carboxylate salt was not acidified prior to isolation of the impure material to avoid MOM group deprotection. Purification by reversed-phase HPLC (Alltima C18, 10 μM, 22 x 250 mm column, eluting with a gradient flow of 0:1 MeCN/$H_2O$ over 5 min, then 0:1→1:0 MeCN/$H_2O$ over 30 min, 254 nm UV detection, flow rate = 12.0 mL/min, $R_T$ = 19.8–20.1 min = 49–50% MeCN); white solid (10 mg, 9%); $^1$H NMR (DMSO-$d_6$, 500 MHz) δ 7.86 (dd, *J* = 7.6, 1.8 Hz, 1H), 7.46 (d, *J* = 9.0 Hz, 1H), 7.10 (d, *J* = 9.0 Hz, 1H), 7.02 (td, *J* = 7.5, 1.9 Hz, 1H), 6.60 (td, *J* = 7.4, 1.1 Hz, 1H), 6.09 (dd, *J* = 8.2, 1.1 Hz, 1H), 5.31 (s, 2H), 3.42 (s, 3H) ppm. LRMS (ES$^-$) calcd 340.01 for $C_{15}H_{12}Cl_2NO_4^-$ found 339.6 ($M^-$).

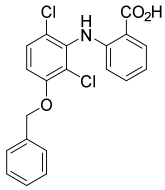

**AK-24**. Prepared according to the above procedure. Purification by reversed-phase HPLC (Alltima C18, 10 μM, 22 x 250 mm column, eluting with a gradient flow of 0:100→70:30 MeCN/0.1% TFA in $H_2O$ over 10 min, then 70:30→100:0 MeCN/0.1% TFA over 25 min, 254 nm UV detection, flow rate = 12.0 mL/min, $R_T$ = 22.2–24.0 min = 85–87% MeCN); white solid (13 mg, 32%); $^1$H NMR (DMSO-$d_6$, 500 MHz) δ 9.52 (s, 1H), 7.90 (dd, *J* = 8.0, 1.6 Hz, 1H), 7.56 (d, *J* = 9.0 Hz, 1H), 7.50–7.47 (m, 2H), 7.42 (t, *J* = 7.4 Hz, 2H), 7.37–7.30 (m, 2H), 7.26 (d, *J* = 9.1 Hz, 1H), 6.78 (ddd, *J* = 8.0, 7.1, 1.1 Hz, 1H), 6.23 (dd, *J* = 8.4, 1.0 Hz, 1H), 5.26 (s, 2H) ppm. LRMS (ES$^-$) calcd 386.04 for $C_{20}H_{14}Cl_2NO_3^-$ found 385.7 ($M^-$).

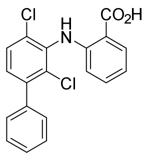

**AK-25**. Prepared according to the above procedure. Purification by reversed-phase HPLC (Alltima C18, 10 μM, 22 x 250 mm column, eluting with a gradient flow of 0:100→70:30 MeCN/0.1% TFA in $H_2O$ over 10 min, then 70:30→100:0 MeCN/0.1% TFA over 25 min, 254 nm UV detection, flow rate = 12.0 mL/min, $R_T$ = 23.3–24.5 min = 86–87% MeCN); white solid (1 mg, 18%); $^1$H NMR (acetone-$d_6$, 500 MHz) δ 9.69 (s, 1H), 8.05 (dd, *J* = 8.0, 1.7 Hz, 1H), 7.64 (d, *J* = 8.3 Hz, 1H), 7.51–7.36 (m, 7H), 6.83 (ddd, *J* = 8.1, 7.1, 1.1 Hz, 1H), 6.45 (dd, *J* = 8.4, 1.0 Hz, 1H) ppm. LRMS (ES$^-$) calcd 356.03 for $C_{19}H_{12}Cl_2NO_2^-$ found 355.6 ($M^-$).

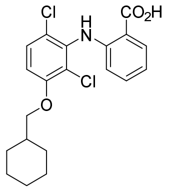

**AK-26**. Prepared according to the above procedure. Purification by reversed-phase HPLC (Alltima C18, 10 μM, 22 x 250 mm column, eluting with a gradient flow of 0:100→70:30 MeCN/0.1% TFA in $H_2O$ over 10 min, then 70:30→100:0 MeCN/0.1% TFA over 25 min, 254 nm UV detection, flow rate = 12.0 mL/min, $R_T$ = 30.2–31.6 min = 94–96% MeCN); white solid (1 mg, 9%); $^1$H NMR (acetone-$d_6$, 500 MHz) δ 9.57 (s, 1H), 8.03 (dd, *J* = 8.0, 1.7 Hz, 1H), 7.48 (d, *J* = 9.0 Hz, 1H), 7.35 (td, *J* = 8.6, 7.1, 1.5 Hz, 1H), 7.12 (d, *J* = 9.0 Hz, 1H), 6.81 (td, *J* = 8.1, 7.1, 1.1 Hz, 1H), 6.35 (dd, *J* = 8.6, 1.1 Hz, 1H), 3.95 (d, *J* = 6.2 Hz, 2H), 1.95–1.82 (m, 3H), 1.80–1.66 (m, 3H), 1.32 (tt, *J* = 12.4, 3.2 Hz, 2H), 1.23 (tt, *J* = 12.4, 3.1 Hz, 1H), 1.15 (qd, *J* = 12.3, 3.4 Hz, 2H) ppm. LRMS ($ES^-$) calcd 392.08 for $C_{20}H_{20}Cl_2NO_3^-$ found 391.8 ($M^-$).

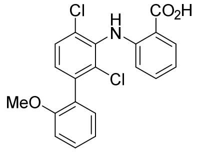

**AK-27**. Prepared according to the above procedure. Purification by reversed-phase HPLC (Alltima C18, 10 μM, 22 x 250 mm column, eluting with a gradient flow of 0:100→70:30 MeCN/0.1% TFA in $H_2O$ over 10 min, then 70:30→100:0 MeCN/0.1% TFA over 25 min, 254 nm UV detection, flow rate = 12.0 mL/min, $R_T$ = 22.2–23.2 min = 85–86% MeCN); white solid (3 mg, 12%); $^1$H NMR (acetone-$d_6$, 500 MHz) δ 9.70 (s, 1H), 8.04 (dd, *J* = 8.0, 1.6 Hz, 1H), 7.60 (d, *J* = 8.4 Hz, 1H), 7.42 (ddd, *J* = 8.3, 7.4, 1.7 Hz, 1H), 7.39 (s, 1H), 7.28 (d, *J* = 8.3 Hz, 1H), 7.23 (dd, *J* = 7.5, 1.8 Hz, 1H), 7.12 (dd, *J* = 8.3, 1.0 Hz, 1H), 7.04 (td, *J* = 7.5, 1.0 Hz, 1H), 6.82 (t, *J* = 7.6 Hz, 1H), 6.47–6.41 (m, 1H), 3.80 (s, 3H) ppm. LRMS ($ES^-$) calcd 386.04 for $C_{20}H_{14}Cl_2NO_3^-$ found 385.7 ($M^-$).

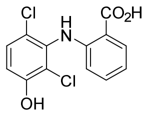

**AK-28**. Prepared according to the above procedure. Purification by reversed-phase HPLC (Alltima C18, 10 μM, 22 x 250 mm column, eluting with a gradient flow of 0:100→70:30 MeCN/0.1% TFA in $H_2O$ over 10 min, then 70:30→100:0 MeCN/0.1% TFA over 25 min, 254 nm UV detection, flow rate = 12.0 mL/min, $R_T$ = 15.4–16.0 min = 76–77% MeCN); white solid (4 mg, 23%); $^1$H NMR (acetone-$d_6$, 600 MHz) δ 9.55 (s, 1H), 8.03 (dd, *J* = 8.0, 1.7 Hz, 1H), 7.41–7.33 (m, 2H), 7.05 (d, *J* = 8.9 Hz, 1H), 6.80 (ddd, *J* = 8.2, 7.1, 1.1 Hz, 1H), 6.37 (ddd, *J* = 8.5, 3.3, 1.0 Hz, 1H) ppm. LRMS ($ES^-$) calcd 295.99 for $C_{13}H_8Cl_2NO_3^-$ found 295.6 ($M^-$).

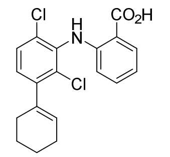

**AK-29**. Prepared according to the above procedure. Purification by reversed-phase HPLC (Alltima C18, 10 μM, 22 x 250 mm column, eluting with a gradient flow of 0:100→70:30 MeCN/0.1% TFA in $H_2O$ over 10 min, then 70:30→100:0 MeCN/0.1% TFA over 25 min, 254 nm UV detection, flow rate = 12.0 mL/min, $R_T$ = 29.0–30.0 min = 93–94% MeCN); white solid (3 mg, 28%); $^1$H NMR (acetone-d$_6$, 600 MHz) δ 9.60 (s, 1H), 8.04 (dd, *J* = 8.1, 1.7 Hz, 1H), 7.50 (d, *J* = 8.3 Hz, 1H), 7.35 (ddd, *J* = 8.6, 7.1, 1.7 Hz, 1H), 7.19 (d, *J* = 8.3 Hz, 1H), 6.81 (ddd, *J* = 8.2, 7.1, 1.2 Hz, 1H), 6.37–6.30 (m, 1H), 5.71 (tt, *J* = 3.7, 1.8 Hz, 1H), 2.32–2.29 (m, 2H), 2.20–2.16 (m, 2H), 1.80–1.73 (m, 2H), 1.72–1.64 (m, 2H) ppm. LRMS (ES$^-$) calcd 360.06 for $C_{19}H_{16}Cl_2NO_2^-$ found 359.7 (M$^-$).

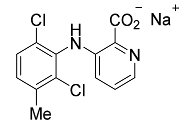

**AK-30**. Prepared according to the above procedure. The solution containing the sodium carboxylate salt was not acidified prior to isolation of the impure material. Purification by reversed-phase HPLC (Alltima C18, 10 μM, 22 x 250 mm column, eluting with a gradient flow of 0:1 MeCN/$H_2O$ over 5 min, then 0:1→1:0 MeCN/$H_2O$ over 30 min, 254 nm UV detection, flow rate = 12.0 mL/min, $R_T$ = 19.8–22.0 min = 49–56% MeCN); white solid (35 mg, 79%); $^1$H NMR (methanol-d$_4$, 500 MHz) δ 7.96 (dd, *J* = 4.6, 1.3 Hz, 1H), 7.39 (d, *J* = 8.3 Hz, 1H), 7.27 (dd, *J* = 8.5, 4.6 Hz, 1H), 7.24 (dd, *J* = 8.3, 0.7 Hz, 1H), 6.77 (dd, *J* = 8.5, 1.3 Hz, 1H), 2.41 (s, 3H) ppm. LRMS (ES$^-$) calcd 295.00 for $C_{13}H_9Cl_2N_2O_2^-$ found 294.6 (M$^-$).

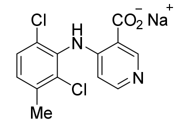

**AK-31**. Prepared according to the above procedure. The solution containing the sodium carboxylate salt was not acidified prior to isolation of the impure material. Purification by reversed-phase HPLC (Alltima C18, 10 μM, 22 x 250 mm column, eluting with a gradient flow of 0:1 MeCN/$H_2O$ over 5 min, then 0:1→1:0 MeCN/$H_2O$ over 30 min, 254 nm UV detection, flow rate = 12.0 mL/min, $R_T$ = 19.6–20.8 min = 49–56% MeCN); white solid (7 mg, 90%); $^1$H NMR (DMSO-d$_6$, 500 MHz) δ 8.74 (s, 1H), 8.13 (d, *J* = 6.6 Hz, 1H), 7.56 (d, *J* = 8.4 Hz, 1H), 7.44 (d, *J* = 8.4 Hz, 1H), 6.27 (d, *J* = 6.7 Hz, 1H), 2.38 (s, 3H) ppm. LRMS (ES$^-$) calcd 295.00 for $C_{13}H_9Cl_2N_2O_2^-$ found 294.6 (M$^-$).

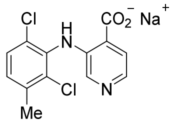

**AK-32**. Prepared according to the above procedure. The solution containing the sodium carboxylate salt was not acidified prior to isolation of the impure material. Purification by reversed-phase HPLC (Alltima C18, 10 μM, 22 x 250 mm column, eluting with a gradient flow of 0:1 MeCN/$H_2O$ over 5 min, then 0:1→1:0 MeCN/$H_2O$ over 30 min, 254 nm UV detection, flow rate = 12.0 mL/min, $R_T$ = 16.0–19.2 min = 36–47% MeCN); white solid (40 mg, 81%); $^1$H NMR (methanol-$d_4$, 500 MHz) δ 7.89 (d, *J* = 5.0 Hz, 1H), 7.81 (d, *J* = 5.1 Hz, 1H), 7.48 (s, 1H), 7.39 (d, *J* = 8.3 Hz, 1H), 7.22 (d, *J* = 8.3 Hz, 1H), 2.42 (s, 3H) ppm. LRMS (ES$^-$) calcd 295.00 for $C_{13}H_9Cl_2N_2O_2^-$ found 294.6 (M$^-$).

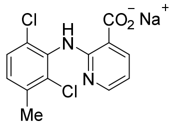

**AK-33**. Prepared according to the above procedure. The solution containing the sodium carboxylate salt was not acidified prior to isolation of the impure material. Purification by reversed-phase HPLC (Alltima C18, 10 μM, 22 x 250 mm column, eluting with a gradient flow of 0:1 MeCN/$H_2O$ over 5 min, then 0:1→1:0 MeCN/$H_2O$ over 30 min, 254 nm UV detection, flow rate = 12.0 mL/min, $R_T$ = 14.4–18.2 min = 31–44% MeCN); white solid (70 mg, 99%); $^1$H NMR (methanol-$d_4$, 500 MHz) δ 8.27 (dd, *J* = 7.5, 1.9 Hz, 1H), 7.89 (dd, *J* = 5.0, 2.0 Hz, 1H), 7.33 (d, *J* = 8.3 Hz, 1H), 7.18 (d, *J* = 8.3 Hz, 1H), 6.70 (dd, *J* = 7.5, 5.0 Hz, 1H), 2.40 (s, 3H) ppm. LRMS (ES$^-$) calcd 295.00 for $C_{13}H_9Cl_2N_2O_2^-$ found 294.6 (M$^-$).

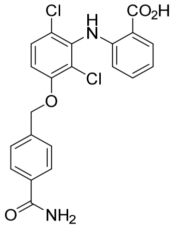

**AK-37**. Prepared according to the above procedure. Purification by reversed-phase HPLC (Alltima C18, 10 μM, 22 x 250 mm column, eluting with a gradient flow of 0:100→70:30 MeCN/0.1% TFA in $H_2O$ over 10 min, then 70:30→100:0 MeCN/0.1% TFA over 25 min, 254 nm UV detection, flow rate = 12.0 mL/min, $R_T$ = 21.2–22.2 min = 71–73% MeCN); white solid (22 mg, yield not determined); $^1$H NMR (DMSO-$d_6$, 600 MHz) δ 8.70 (d, *J* = 8.4 Hz, 1H), 8.06 (dd, *J* = 8.0, 1.6 Hz, 1H), 7.99 (d, *J* = 8.2 Hz, 2H), 7.65 (d, *J* = 7.9 Hz, 3H), 7.20 (t, *J* = 7.5 Hz, 1H), 7.18 (d, *J* = 8.9 Hz, 1H), 6.49 (d, *J* = 8.9 Hz, 1H), 5.50 (s, 2H), 5.27 (s, 2H) ppm; LRMS (ES$^-$) calcd 429.04 for $C_{21}H_{15}Cl_2N_2O_4^-$ found 429.1 (M$^-$).

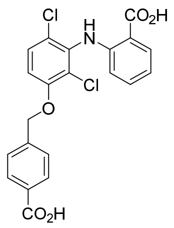

**AK-38**. Prepared according to the above procedure. Purification by reversed-phase HPLC (Alltima C18, 10 μM, 22 x 250 mm column, eluting with a gradient flow of 0:100→70:30 MeCN/0.1% TFA in $H_2O$ over 10 min, then 70:30→100:0 MeCN/0.1% TFA over 25 min, 254 nm UV detection, flow rate = 12.0 mL/min, $R_T$ = 12.0–15.2 min = 72–76% MeCN); white solid (22 mg, 65%); $^1$H NMR (DMSO-$d_6$, 500 MHz) δ 9.53 (s, 1H), 7.98 (d, *J* = 8.2 Hz, 2H), 7.90 (dd, *J* = 8.0, 1.8 Hz, 1H), 7.60 (d, *J* = 8.3 Hz, 2H), 7.57 (d, *J* = 9.0 Hz, 1H), 7.33 (t, *J* = 7.4 Hz, 1H), 7.24 (d, *J* = 9.2 Hz, 1H), 6.79 (t, *J* = 7.4 Hz, 1H), 6.24 (d, *J* = 8.4 Hz, 1H), 5.36 (s, 2H) ppm; LRMS (ES$^-$) calcd 430.03 for $C_{21}H_{14}Cl_2NO_5^-$ found 430.0 (M$^-$).

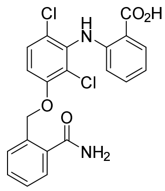

**AK-39**. Prepared according to the above procedure. Purification by reversed-phase HPLC (Alltima C18, 10 μM, 22 x 250 mm column, eluting with a gradient flow of 0:100→70:30 MeCN/0.1% TFA in $H_2O$ over 10 min, then 70:30→100:0 MeCN/0.1% TFA over 25 min, 254 nm UV detection, flow rate = 12.0 mL/min, $R_T$ = 19.4–20.2 min = 81–82% MeCN); white solid (21 mg, 38%); $^1$H NMR (DMSO-$d_6$, 600 MHz) δ 11.77 (s, 1H), 8.57 (d, *J* = 8.4 Hz, 1H), 8.02 (dd, *J* = 7.9, 1.7 Hz, 1H), 7.76 (d, *J* = 7.7 Hz, 1H), 7.70 (d, *J* = 7.7 Hz, 1H), 7.64–7.59 (m, 2H), 7.53 (t, *J* = 7.5 Hz, 1H), 7.20 (t, *J* = 7.7 Hz, 1H), 7.13 (d, *J* = 8.9 Hz, 1H), 6.43 (d, *J* = 8.9 Hz, 1H), 5.40 (s, 4H) ppm; LRMS (ES$^-$) calcd 429.04 for $C_{21}H_{15}Cl_2N_2O_4^-$ found 429.1 (M$^-$).

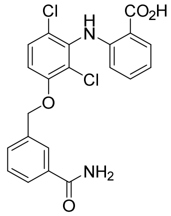

**AK-40**. Prepared according to the above procedure. Purification by reversed-phase HPLC (Alltima C18, 10 μM, 22 x 250 mm column, eluting with a gradient flow of 0:100→70:30 MeCN/0.1% TFA in $H_2O$ over 10 min, then 70:30→100:0 MeCN/0.1% TFA over 25 min, 254 nm UV detection, flow rate = 12.0 mL/min, $R_T$ = 20.0–20.8 min = 82–83% MeCN); white solid (41 mg, 75%); $^1$H NMR (DMSO-$d_6$, 600 MHz) δ 12.24 (s, 1H), 8.70 (d, *J* = 8.3 Hz, 1H), 8.07–8.05 (m, 2H), 7.91 (d, *J* = 7.8 Hz, 1H), 7.71 (d, *J* = 7.7 Hz, 1H), 7.67 (t, *J* = 7.7 Hz, 1H), 7.63 (t, *J* = 7.7 Hz, 1H), 7.22 (t, *J* = 7.6 Hz, 1H), 7.18 (d, *J* = 8.9 Hz, 1H), 6.51 (d, *J* = 8.9 Hz, 1H), 5.49 (s, 2H), 5.26 (s, 2H) ppm; LRMS (ES$^-$) calcd 429.04 for $C_{21}H_{15}Cl_2N_2O_4^-$ found 429.1 (M$^-$).

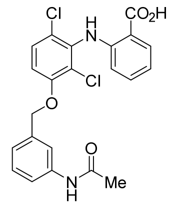

**AK-41**. Prepared according to the above procedure. Purification by reversed-phase HPLC (Alltima C18, 10 μM, 22 x 250 mm column, eluting with a gradient flow of 0:100→70:30 MeCN/0.1% TFA in $H_2O$ over 10 min, then 70:30→100:0 MeCN/0.1% TFA over 25 min, 254 nm UV detection, flow rate = 12.0 mL/min, $R_T$ = 14.2–15.0 min = 75–76% MeCN); white solid (10 mg, 33%); $^1$H NMR (DMSO-$d_6$, 600 MHz) δ 9.99 (s, 1H), 9.54 (s, 1H), 7.90 (d, *J* = 8.0 Hz, 1H), 7.70 (s, 1H), 7.56–7.54 (m, 2H), 7.32 (t, *J* = 7.7 Hz, 2H), 7.23 (d, *J* = 9.1 Hz, 1H), 7.14 (d, *J* = 7.5 Hz, 1H), 6.78 (t, *J* = 7.5 Hz, 1H), 6.24 (d, *J* = 8.4 Hz, 1H), 5.23 (s, 2H), 2.04 (s, 3H) ppm; LRMS (ES$^-$) calcd 443.06 for $C_{22}H_{17}Cl_2N_2O_4^-$ found 443.1 (M$^-$).

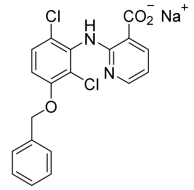

**AK-42**. Prepared according to the above procedure. The solution containing the sodium carboxylate salt was not acidified prior to isolation of the impure material. Purification by reversed-phase HPLC (Alltima C18, 10 μM, 22 x 250 mm column, eluting with a gradient flow of 0:100→70:30 MeCN/ $H_2O$ over 10 min, then 70:30→100:0 MeCN/$H_2O$ over 25 min, 254 nm UV detection, flow rate = 12.0 mL/min, $R_T$ = 12.2–13.7 min = 73–75% MeCN); white solid (11 mg, 78%); $^1$H NMR (DMSO-$d_6$, 600 MHz) δ 8.07 (dd, *J* = 7.3, 2.1 Hz, 1H), 7.88 (d, *J* = 3.4 Hz, 1H), 7.49 (d, *J* = 7.5 Hz, 2H), 7.43–7.40 (m, 3H), 7.35 (t, *J* = 7.3 Hz, 1H), 7.11 (d, *J* = 9.0 Hz, 1H), 6.62 (dd, *J* = 7.4, 4.8 Hz, 1H), 5.23 (s, 2H) ppm; LRMS (ES$^-$) calcd 387.03 for $C_{19}H_{13}Cl_2N_2O_3^-$ found 387.0 (M$^-$).

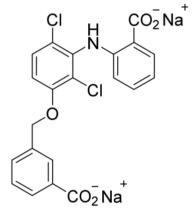

**AK-43**. Prepared according to the above procedure. The solution containing the sodium carboxylate salt was not acidified prior to isolation of the impure material. Purification by reversed-phase HPLC (Alltima C18, 10 μM, 22 x 250 mm column, eluting with a gradient flow of 0:100→70:30 MeCN/ $H_2O$ over 10 min, then 70:30→100:0 MeCN/$H_2O$ over 25 min, 254 nm UV detection, flow rate = 12.0 mL/min, $R_T$ = 11.2–14.2 min = 71–75% MeCN); white solid (6 mg, 37%); $^1$H NMR (DMSO-$d_6$, 600 MHz) δ 8.04 (s, 1H), 7.89 (d, *J* =

7.8 Hz, 1H), 7.84 (d, *J* = 7.7 Hz, 1H), 7.64 (d, *J* = 7.3 Hz, 1H), 7.52–7.43 (m, 2H), 7.11 (d, *J* = 8.7 Hz, 1H), 7.02 (t, *J* = 6.6 Hz, 1H), 6.59 (t, *J* = 7.4 Hz, 1H), 6.09 (d, *J* = 8.1 Hz, 1H), 5.30 (s, 2H) ppm; LRMS ($ES^-$) calcd 430.03 for $C_{21}H_{14}Cl_2NO_5^-$ found 430.0 ($M^-$).

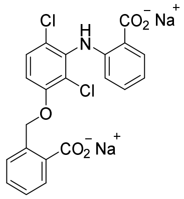

**AK-44**. Prepared according to the above procedure. The solution containing the sodium carboxylate salt was not acidified prior to isolation of the impure material. Purification by reversed-phase HPLC (Alltima C18, 10 μM, 22 x 250 mm column, eluting with a gradient flow of 0:100→70:30 MeCN/$H_2O$ over 10 min, then 70:30→100:0 MeCN/ $H_2O$ over 25 min, 254 nm UV detection, flow rate = 12.0 mL/min, $R_T$ = 10.8–12.2 min = 71–73% MeCN); white solid (9 mg, 40%); $^1$H NMR (DMSO-d$_6$, 600 MHz) δ 7.86 (dd, *J* = 7.7, 1.8 Hz, 1H), 7.83 (d, *J* = 7.6 Hz, 1H), 7.58 (d, *J* = 7.7 Hz, 1H), 7.44 (d, *J* = 9.0 Hz, 1H), 7.41 (t, *J* = 7.8 Hz, 1H), 7.30 (t, *J* = 7.4 Hz, 1H), 7.03 (t, *J* = 7.6 Hz, 1H), 7.00 (d, *J* = 9.1 Hz, 1H), 6.60 (t, *J* = 7.5 Hz, 1H), 6.12 (d, *J* = 8.1 Hz, 1H), 5.65 (d, *J* = 8.1 Hz, 2H) ppm; LRMS ($ES^-$) calcd 430.03 for $C_{21}H_{14}Cl_2NO_5^-$ found 430.1 ($M^-$).

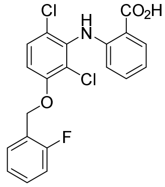

**AK-45**. Prepared according to the above procedure. Purification by reversed-phase HPLC (Alltima C18, 10 μM, 22 x 250 mm column, eluting with a gradient flow of 0:100→70:30 MeCN/0.1% TFA in $H_2O$ over 10 min, then 70:30→100:0 MeCN/0.1% TFA over 25 min, 254 nm UV detection, flow rate = 12.0 mL/min, $R_T$ = 21.3–22.5 min = 84–85% MeCN); white solid (5 mg, 6%); $^1$H NMR (DMSO-d$_6$, 600 MHz) δ 9.48 (s, 1H), 7.85 (dd, *J* = 7.9, 1.7 Hz, 1H), 7.58–7.52 (m, 2H), 7.43–7.39 (m, 1H), 7.30–7.21 (m, 4H), 6.74 (t, *J* = 7.8, 1H), 6.19 (d, *J* = 8.4 Hz, 1H), 5.26 (s, 2H) ppm; $^{19}$F NMR (acetone-d$_6$, 376 MHz) δ –120.18 (dt, *J* = 10.8, 7.0 Hz, 1F) ppm; LRMS ($ES^-$) calcd 404.03 for $C_{20}H_{13}Cl_2FNO_3^-$ found 404.1 ($M^-$).

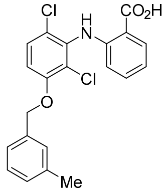

**AK-46**. Prepared according to the above procedure. Purification by reversed-phase HPLC (Alltima C18, 10 μM, 22 x 250 mm column, eluting with a gradient flow of 0:100→70:30 MeCN/0.1% TFA in $H_2O$ over 10 min, then 70:30→100:0 MeCN/0.1% TFA over 25 min, 254 nm UV detection, flow rate = 12.0 mL/min, $R_T$

= 23.7–28.9 min = 86–93% MeCN); white solid (47 mg, 58%); $^{1}$H NMR (DMSO-d$_6$, 600 MHz) δ 9.51 (s, 1H), 7.89 (d, *J* = 8.1 Hz, 1H), 7.56 (d, *J* = 9.1 Hz, 1H), 7.35–7.22 (m, 5H), 7.17 (d, *J* = 7.4 Hz, 1H), 6.78 (t, *J* = 7.6 Hz, 1H), 6.23 (d, *J* = 8.3 Hz, 1H), 5.22 (s, 2H), 2.33 (s, 3H) ppm; LRMS (ES$^-$) calcd 400.05 for $C_{21}H_{16}Cl_2NO_3^-$ found 400.0 (M$^-$).

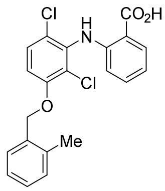

**AK-47**. Prepared according to the above procedure. Purification by reversed-phase HPLC (Alltima C18, 10 μM, 22 x 250 mm column, eluting with a gradient flow of 0:100→70:30 MeCN/0.1% TFA in $H_2O$ over 10 min, then 70:30→100:0 MeCN/0.1% TFA over 25 min, 254 nm UV detection, flow rate = 12.0 mL/min, $R_T$ = 23.2–29.2 min = 86–93% MeCN); white solid (53 mg, 69%); $^{1}$H NMR (DMSO-d$_6$, 600 MHz) δ 9.52 (s, 1H), 7.89 (d, *J* = 8.0 Hz, 1H), 7.58 (d, *J* = 9.0 Hz, 1H), 7.47 (d, *J* = 7.6 Hz, 1H), 7.32 (d, *J* = 9.1 Hz, 2H), 7.28–7.21 (m, 3H), 6.78 (t, *J* = 7.6 Hz, 1H), 6.24 (d, *J* = 8.5 Hz, 1H), 5.24 (s, 2H), 2.36 (s, 3H) ppm; LRMS (ES$^-$) calcd 400.05 for $C_{21}H_{16}Cl_2NO_3^-$ found 400.0 (M$^-$).

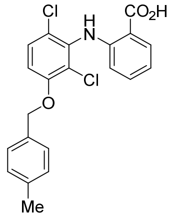

**AK-48**. Prepared according to the above procedure. Purification by reversed-phase HPLC (Alltima C18, 10 μM, 22 x 250 mm column, eluting with a gradient flow of 0:100→70:30 MeCN/0.1% TFA in $H_2O$ over 10 min, then 70:30→100:0 MeCN/0.1% TFA over 25 min, 254 nm UV detection, flow rate = 12.0 mL/min, $R_T$ = 23.8–29.9 min = 86–94% MeCN); white solid (49 mg, 61%); $^{1}$H NMR (DMSO-d$_6$, 600 MHz) δ 9.51 (s, 1H), 7.89 (d, *J* = 8.0 Hz, 1H), 7.55 (d, *J* = 9.7 Hz, 1H), 7.36 (d, *J* = 7.5 Hz, 2H), 7.32 (t, *J* = 7.9 Hz, 1H), 7.24 (d, *J* = 9.4 Hz, 1H), 7.22 (d, *J* = 7.7 Hz, 2H), 6.78 (t, *J* = 7.6 Hz, 1H), 6.23 (d, *J* = 8.2 Hz, 1H), 5.21 (s, 2H), 2.31 (s, 3H) ppm; LRMS (ES$^-$) calcd 400.05 for $C_{21}H_{16}Cl_2NO_3^-$ found 400.0 (M$^-$).

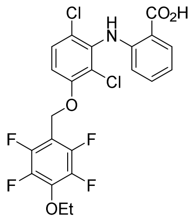

**AK-49**. Prepared according to the above procedure. Nucleophilic aromatic *para*-substitution of the Buchwald-Hartwig coupling product occurred with the ethanolic solvent upon stirring at ambient temperature for 48 h. Purification by reversed-phase HPLC (Alltima C18, 10 μM, 22 x 250 mm column, eluting with a gradient flow of 0:100→70:30 MeCN/0.1% TFA in $H_2O$ over 10 min, then 70:30→100:0 MeCN/0.1% TFA over 25 min, 254 nm UV detection, flow rate = 12.0 mL/min, $R_T$ = 24.2–25.8 min = 87–89% MeCN); off-white solid (44 mg, 80%); $^1H$ NMR (DMSO-d$_6$, 600 MHz) δ 9.50 (s, 1H), 7.89 (dd, *J* = 7.9, 1.7 Hz, 1H), 7.63 (d, *J* = 9.0 Hz, 1H), 7.39 (d, *J* = 9.1 Hz, 1H), 7.32 (ddd, *J* = 8.6, 7.3, 1.7 Hz, 1H), 6.78 (t, *J* = 7.6 Hz, 1H), 6.20 (d, *J* = 8.3 Hz, 1H), 5.31 (s, 2H), 4.35 (q, *J* = 7.0 Hz, 2H), 1.34 (t, *J* = 7.0 Hz, 3H) ppm; $^{19}F$ NMR (acetone-d$_6$, 376 MHz) δ –146.36 (dd, *J* = 20.7, 8.5 Hz, 2F), –159.17 (dd, *J* = 20.7, 8.5 Hz, 2F) ppm; LRMS (ES$^-$) calcd 502.02 for $C_{22}H_{14}Cl_2F_4NO_4^-$ found 502.05 (M$^-$).

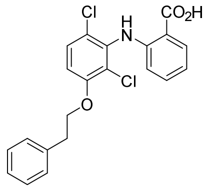

**AK-50**. Prepared according to the above procedure. Purification by reversed-phase HPLC (Alltima C18, 10 μM, 22 x 250 mm column, eluting with a gradient flow of 0:100→70:30 MeCN/0.1%TFA in $H_2O$ over 10 min, then 70:30→100:0 MeCN/0.1%TFA over 25 min, 254 nm UV detection, flow rate = 12.0 mL/min, $R_T$ = 23.2–26.0 min = 86–89% MeCN); white solid (53 mg, 88%); $^1H$ NMR (DMSO-d$_6$, 600 MHz) δ 9.49 (s, 1H), 7.89 (d, *J* = 7.6 Hz, 1H), 7.53 (dd, *J* = 9.1, 1.4 Hz, 1H), 7.37–7.29 (m, 5H), 7.24–7.17 (m, 2H), 6.78 (t, *J* = 7.5 Hz, 1H), 6.21 (d, *J* = 8.3 Hz, 1H), 4.31 (t, *J* = 6.5 Hz, 2H), 3.08 (t, *J* = 6.5 Hz, 2H) ppm; LRMS (ES$^-$) calcd 400.05 for $C_{21}H_{16}Cl_2NO_3^-$ found 400.0 (M$^-$).

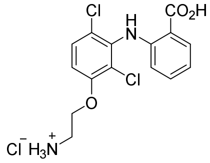

**AK-51**. Prepared according to the above procedure from the corresponding phthalimide-protected ethyl ester. Hydrolysis of the phthalimide group was incomplete, even after heating at 70 °C for 3 h at pH 12. The mixture was acidified to pH 2 with concentrated HCl and purified by reversed-phase HPLC (Alltima C18, 10 μM, 22 x 250 mm column, eluting with a gradient flow of 0:100→70:30 MeCN/0.1% TFA in $H_2O$ over 10 min, then 70:30→100:0 MeCN/0.1% TFA over 25 min, 254 nm UV detection, flow rate = 12.0 mL/min, $R_T$ = 13.0–16.0 min = 74–77% MeCN); To liberate the free amine, the white solid obtained following lyophilization was dissolved in 0.8 mL of EtOH and stirred with an excess of $MeNH_2$ (0.8 mL of a 2.0 M solution in THF, 1.6 mmol, 33.0 equiv) at 70 °C for 5 h. After cooling to ambient temperature, the reaction pH was adjusted to pH 2 with 6.0 M aqueous HCl. Purification by reversed-phase HPLC (Alltima C18, 10 μM, 22 x 250 mm column, eluting with a gradient flow of 0:100→74:26 MeCN/10 mM heptafluoro-butryic acid (HFBA) in $H_2O$ over 5 min, then 74:26→78:22 MeCN/10 mM HFBA in $H_2O$ over 25 min, 254 nm UV detection, flow rate = 12.0 mL/min, $R_T$ = 9.2–9.8 min = 74.7–74.8% MeCN); white solid (4 mg, 21% over two steps); $^1$H NMR (DMSO-d$_6$, 600 MHz) δ 7.90 (dd, *J* = 8.0, 1.7 Hz, 1H), 7.58 (d, *J* = 9.1 Hz, 1H), 7.33 (ddd, *J* = 8.6, 7.1, 1.7 Hz, 1H), 7.19 (d, *J* = 9.1 Hz, 1H), 6.80 (ddd, *J* = 8.1, 7.1, 1.1 Hz, 1H), 6.22 (dd, *J* = 8.5, 1.1 Hz, 1H), 4.29 (t, *J* = 5.2 Hz, 2H), 3.27 (t, *J* = 5.1 Hz, 2H) ppm; LRMS (ES$^-$) calcd 339.03 for $C_{15}H_{13}Cl_2N_2O_3^-$ found 339.0 (M$^-$).

#### *Synthesis of sulfate, phosphonate, and tetrazole derivatives*

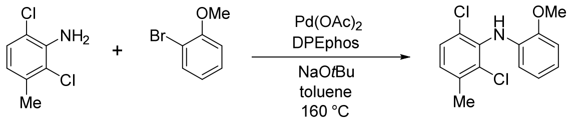

The following procedure is adapted from Sadighi, et al.**Error! Bookmark not defined.** An oven-dried 5-mL microwave vial was charged with 2,6-dichloro-3-methylaniline (176 mg, 1.0 mmol), 2-bromoanisole (187 mg, 1.0 mmol), and DPEphos (40 mg, 0.075 mmol, 0.075 equiv). The vial was sealed with a septum and purged with argon prior to the addition of 3.0 mL of toluene. The solution was sparged with argon for ~5 min. Under a blanket of argon, the septum was briefly removed and Pd(OAc)$_2$ (11 mg, 0.05 mmol, 0.05 equiv) was added. The vial was resealed, and the orange solution was stirred at room temperature for 10 min. Following this time, the septum was quickly removed and sodium *t*-butoxide (135 mg, 1.4 mmol, 1.4 equiv) was added. The headspace of the vial was flushed with argon and quickly resealed with a crimped microwave vial cap. The reaction was stirred at 160 °C in a microwave reactor for 2 h. The reaction mixture was cooled to room temperature, diluted with 10 mL of EtOAc, and filtered through a plug of Celite. The flask and filter cake were rinsed with ~30 mL of EtOAc, and the combined filtrates were concentrated under reduced pressure. The brown residue was re-dissolved in $CH_2Cl_2$ to which ~1 g of silica gel was then added. The suspension was concentrated under reduced pressure and the solid material dry loaded onto a silica gel column pre-packed in pentane. Purification of this material was accomplished by chromatography on silica gel (gradient elution: 0:1→1:19 $Et_2O$/pentane) furnished the desired product as a white solid (142 mg, 50%). TLC $R_f$ = 0.6 (19:1 hexanes/acetone); $^1$H NMR (CDCl$_3$, 400 MHz) δ 7.29 (d, *J* = 8.3 Hz, 1H), 7.04 (dd, *J* = 8.2, 0.7 Hz, 1H), 6.91 (dd, *J* = 7.9, 1.6 Hz, 1H), 6.86 (td, *J* = 7.9, 1.8 Hz, 1H), 6.80 (td, *J* = 7.6, 1.6 Hz, 1H), 6.36 (dd, *J* = 7.6, 1.7 Hz, 1H), 3.97 (s, 3H), 2.41 (s, 3H) ppm; IR (thin film) ν 3372, 1598, 1506, 1452, 1428, 1247, 1225, 1117 cm$^{-1}$.

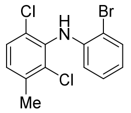

Prepared according to the above procedure substituting 1,2-dibromobenzene for 2-bromoanisole; white solid (453 mg, 55%). TLC $R_f$ = 0.71 (19:1 hexanes/EtOAc); $^1$H NMR (CDCl$_3$, 300 MHz) δ 7.54 (dd, *J* = 7.9, 1.5 Hz, 1H), 7.30 (d, *J* = 8.3 Hz, 1H), 7.20–7.12 (m, 2H), 6.74 (ddd, *J* = 7.9, 7.3, 1.5 Hz, 1H), 6.35 (dd, *J* = 8.1, 1.5 Hz, 1H), 6.12 (s, 1H), 2.41 (s, 3H) ppm; IR (thin film) ν 3376, 3062, 2954, 2922, 2851, 1595, 1500, 1464, 1448, 1400, 1305 cm$^{-1}$.

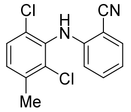

Prepared according to the above procedure substituting 2-bromobenzonitrile for 2-bromoanisole; white solid (128 mg, 46%). TLC $R_f$ = 0.42 (19:1 hexanes/EtOAc); $^1$H NMR ($CDCl_3$, 500 MHz) δ 7.54 (d, *J* = 7.6 Hz, 1H), 7.37–7.29 (m, 2H), 7.15 (d, *J* = 8.3 Hz, 1H), 6.88 (t, *J* = 7.6 Hz, 1H), 6.37 (d, *J* = 8.4 Hz, 1H), 6.23 (s, 1H), 2.41 (s, 3H), ppm; IR (thin film) ν 3306, 2222, 1605, 1578, 1512, 1452, 1399, 1317, 1296, 1163 $cm^{-1}$.

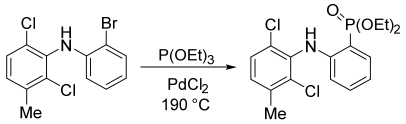

The following procedure is adapted from Ghalib, et al.[5] To a 2 mL oven-dried conical reaction vial was added successively *N*-(2-bromophenyl)-2,6-dichloro-3-methylaniline (263 mg, 0.79 mmol), neat triethylphosphite (0.3 mL, 1.75 mmol, 2.2 equiv), and $PdCl_2$ (28 mg, 0.16 mmol, 0.2 equiv). The vial was purged with argon for 2 min then capped with a sturdy screw-cap and placed in a pre-heated sand bath at 180–190 °C. The reaction was stirred at this temperature for 3 h, over which time the mixture changed in color from pale yellow to dark brownish-black. The reaction mixture was cooled to ambient temperature, diluted with 10 mL of $CH_2Cl_2$, and filtered through a plug of Celite. The vial and filter cake were rinsed with 3 x 30 mL of $CH_2Cl_2$. The combined filtrates were concentrated under reduced pressure. The oily brown residue was re-dissolved in $CH_2Cl_2$ to which ~1 g of silica gel was then added. The suspension was concentrated under reduced pressure and the solid material dry loaded onto a silica gel column pre-packed in hexanes. Purification by chromatography on silica gel (gradient elution: 0:1→1:3 EtOAc/hexanes) provided the desired product as yellow-brown oil (265 mg, 86%). TLC $R_f$ = 0.30 (3:1 hexanes/EtOAc); $^1$H NMR ($CDCl_3$, 300 MHz) δ 8.27 (s, 1H), 7.62 (ddd, *J* = 14.5, 7.7, 1.6 Hz, 1H), 7.37–7.29 (m, 2H), 7.06 (d, *J* = 8.3 Hz, 1H), 6.84 (td, *J* = 7.4, 3.2 Hz, 1H), 6.33 (t, *J* = 7.5 Hz, 1H), 4.31–4.05 (m, 4H), 2.39 (s, 3H), 1.36 (t, *J* = 7.1, 1.8 Hz, 6H) ppm; $^{31}$P NMR ($CDCl_3$, 400 MHz) δ 21.73 ppm; IR (thin film) ν 3259, 2982, 1600, 1512, 1449, 1391, 1218, 1141 $cm^{-1}$.

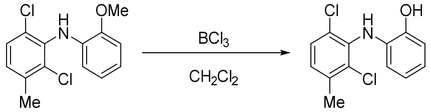

To an ice-cold solution of 2,6-dichloro-N-(2-methoxyphenyl)-3-methylaniline (51 mg, 0.18 mmol) in 2.2 mL of anhydrous $CH_2Cl_2$ was added boron trichloride (0.6 mL of 1.0 M in hexanes, 0.6 mmol, 3.3 equiv) was added dropwise. The reaction was warmed to room temperature over 1 h and stirred for an additional 4 h.

5. Ghalib, M.; Jones, P. G.; Lysenko, S.; Heinicke, J. W. Enantiomerically Pure N Chirally Substituted 1,3-Benzazaphospholes: Synthesis, Reactivity toward *t*-BuLi, and Conversion to Functionalized Benzazaphospholes and Catalytically Useful Dihydrobenzazaphospholes. *Organometallics* **2014**, *33*, 804–816.

Following this time, the reaction was quenched with 2 mL of saturated aqueous $NH_4Cl$ and transferred to a separatory funnel with 60 mL of EtOAc. The organic fraction was washed with 1 x 60 mL of saturated aqueous $NH_4Cl$ and 2 x 60 mL of saturated aqueous NaCl, dried over $MgSO_4$, filtered and concentrated under reduced pressure. Purification of this material was accomplished by chromatography on silica gel (gradient elution, 0:1→1:3 EtOAc/hexanes) to furnish the desired product as a clear oil (13 mg, 27%, note: the product is extremely prone to oxidation and turns turquoise upon prolonged exposure to air). TLC $R_f$ = 0.15 (19:1 hexanes/acetone); $^1$H NMR ($CDCl_3$, 300 MHz) δ 7.22 (d, *J* = 8.3 Hz, 1H), 7.05–6.92 (m, 3H), 6.78 (td, *J* = 7.4, 1.9 Hz, 1H), 6.62 (dd, *J* = 7.9, 1.5 Hz, 1H), 5.93 (s, 1H), 2.37 (s, 3H) ppm; IR (thin film) ν 3388, 1595, 1511, 1494, 1451, 1222 $cm^{-1}$.

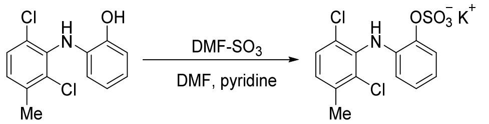

**AK-9**. A flask containing anhydrous sulfur trioxide *N,N*-dimethylformamide complex (50 mg, 0.33 mmol, 6.8 equiv) was placed in an ice bath to which a solution of 2-((2,6-dichloro-3-methyl-phenyl)amino)phenol (9.8 mg, 0.048 mmol) in 750 • L of a 1:4 mixture of freshly distilled pyridine/DMF was added dropwise via syringe. Following the addition, the flask was removed from the ice bath and the mixture was stirred at ambient temperature for 1 h, 30 °C for 2 h, and 45 °C for 2 h. The reaction was quenched by the addition of 0.1 mL of saturated aqueous KOH. All volatiles were removed in vacuo to a yellow residue, which was triturated successively with 1 mL of hexanes, 1 mL of acetone, and 1 mL of EtOAc. The impure material was dissolved in 2 mL of DMSO and filtered through a 13 mm syringe filter with a 0.45 μm PTFE membrane to remove any particulate matter. The desired product was obtained following reversed-phase HPLC (Alltima C18, 10 μM, 22 x 250 mm column, eluting with gradient flow over 30 min of 0:1→1:0 $MeCN/H_2O$, 254 nm UV detection, flow rate = 12.0 mL/min, $R_T$ = 17.4–18.4 min = 58–61% MeCN); white powder (6 mg, 43%). $^1$H NMR (DMSO-$d_6$, 600 MHz) δ 7.44 (s, 1H), 7.39 (d, *J* = 8.3 Hz, 1H), 7.15 (dd, *J* = 8.2, 0.8 Hz, 1H), 7.12 (dd, *J* = 7.9, 1.6 Hz, 1H), 6.88 (td, *J* = 7.7, 1.6 Hz, 1H), 6.77 (td, *J* = 7.6, 1.6 Hz, 1H), 6.23 (dd, *J* = 7.9, 1.6 Hz, 1H), 2.35 (s, 3H) ppm; LRMS ($ES^-$) calcd 345.97 for $C_{13}H_{10}Cl_2NO_4S^-$ found 345.9 ($M^-$).

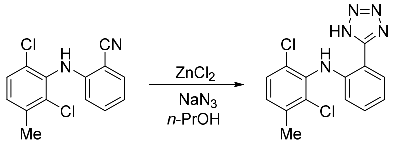

**AK-18**. The following procedure is adapted from Vorona, et al.[6] A 1 mL microscale reaction vial was charged with 2-((2,6-dichloro-3-methylphenyl)amino)benzonitrile (21 mg, 0.076 mmol), 0.2 mL of *n*-propanol, zinc chloride (10 mg, 0.076 mmol), and sodium azide (6 mg, 0.091 mmol, 1.2 equiv). The vial was capped and sealed with Teflon tape, and the mixture was stirred at 95 °C for 3 h. Upon cooling, 0.1 mL of 1.0 M aqueous

6. Vorona, S.; Artamonova, T.; Zevatskii, Y.; Myznikov, L. An Improved Protocol for the Preparation of 5-Substituted Tetrazoles from Organic Thiocyanates and Nitriles. *Synthesis* **2014**, *46*, 781–786.

NaOH was added, resulting in the immediate formation of a white precipitate. The reaction mixture was filtered through a cotton plug. The flask and cotton filter were rinsed with ~3 mL of *n*-propanol. The pH of the filtrate was adjusted to pH 1 with ~1–3 drops of concentrated aqueous HCl. The solution was transferred to a separatory funnel with 5 mL of $H_2O$ and extracted with 5 mL of EtOAc. The organic fraction was washed with 2 x 5 mL of 1.0 M aqueous HCl, dried over $Na_2SO_4$, filtered and concentrated under reduced pressure. Purification of this material by chromatography on silica gel (2:3 hexanes/EtOAc) provided the tetrazole product as white solid (3 mg, 13%). TLC $R_f$ = 0.13 (2:3 hexanes/EtOAc); $^1$H NMR ($CDCl_3$, 500 MHz) δ 8.74 (s, 1H), 7.85 (d, *J* = 7.8 Hz, 1H), 7.34–7.28 (m, 2H), 7.13 (d, *J* = 8.3 Hz, 1H), 6.96 (t, *J* = 7.5 Hz, 1H), 6.51 (d, *J* = 8.3 Hz, 1H), 2.42 (s, 3H) ppm; IR (thin film) ν 2924, 1702, 1616, 1590, 1552, 1480, 1454, 1312, 1286, cm$^{-1}$; LRMS (ES$^-$) calcd 318.03 for $C_{14}H_{10}Cl_2N_5^-$ found 317.6 (M$^-$).

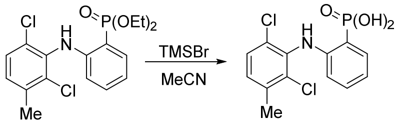

**AK-34**. Diethyl (2-((2,6-dichloro-3-methylphenyl)amino)-phenyl)phosphonate (71 mg, 0.18 mmol) was dissolved in 2.0 mL of MeCN, the solution cooled to 0 °C, and bromotrimethylsilane (0.2 mL, 1.5 mmol, 8.3 equiv) added dropwise via syringe. The flask was removed from the ice bath, equipped with a reflux condenser, and placed in an oil bath at 80 °C. The mixture was stirred at reflux under $N_2$ for 8 h. Following this time, the reaction was cooled to ambient temperature and stirred for an additional 18 h. The reaction was quenched by the addition of 1 mL of $H_2O$ and stirred vigorously for 15 min prior to concentrating the solution under reduced pressure. The yellow-orange residue was transferred to a separatory funnel with 100 mL of EtOAc. The organic layer was washed with 1 x 100 mL of 1.0 M aqueous HCl and 2 x 100 mL of $H_2O$, dried over $Na_2SO_4$, filtered and concentrated under reduced pressure. The desired product was obtained following reversed-phase HPLC (Alltima C18, 10 μM, 22 x 250 mm column, eluting with a gradient flow of 0:100→70:30 MeCN/0.1% TFA in $H_2O$ over 10 min, then 70:30→100:0 MeCN/0.1% TFA over 25 min, 254 nm UV detection, flow rate = 12.0 mL/min, $R_T$ = 13.8–15.8 min = 75–77% MeCN); white powder (49 mg, 81%). $^1$H NMR (acetone-$d_6$, 500 MHz) δ 7.70 (ddd, *J* = 15.1, 7.6, 1.7 Hz, 1H), 7.37 (d, *J* = 8.3 Hz, 1H), 7.28–7.19 (m, 2H), 6.81 (td, *J* = 7.3, 2.9 Hz, 1H), 6.28 (t, *J* = 7.4 Hz, 1H), 2.37 (s, 3H) ppm; LRMS (ES$^-$) calcd 328.98 for $C_{13}H_{10}Cl_2NO_3P_2^-$ found 329.6 (M$^-$).

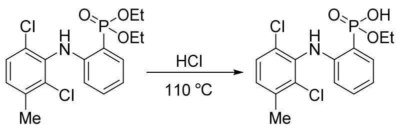

**AK-35**. Diethyl (2-((2,6-dichloro-3-methylphenyl)amino)phenyl)phosphonate (35 mg, 0.09 mmol) was suspended in 2.0 mL of concentrated aqueous HCl. The flask was equipped with a reflux condenser and the contents were stirred at 110 °C for 5 h. The mixture was cooled to room temperature and stirred for an additional 12 h. Following this time, the aqueous yellow solution was decanted, leaving a bright violet residue. This purple film was dissolved in ~5 mL of EtOAc and concentrated under reduced pressure. The desired product was obtained following reversed-phase HPLC (Alltima C18, 10 μM, 22 x 250 mm column, eluting with gradient flow of 0:100→70:30 MeCN/0.1% TFA in $H_2O$ over 10 min, then 70:30→100:0 MeCN/0.1% TFA over 25 min, 254 nm UV detection, flow rate = 12.0 mL/min, $R_T$ = 16.8–19.0 min = 78–81% MeCN); white powder (23 mg, 72%). $^1$H NMR (acetone-$d_6$, 500 MHz) δ 7.67 (dd, *J* = 14.4, 7.6 Hz, 1H), 7.41 (d, *J* = 8.2 Hz, 1H), 7.31–7.22 (m, 2H), 6.84 (t, *J* = 6.4 Hz, 1H), 6.27 (t, *J* = 7.4 Hz, 1H), 4.03 (quintet, *J* = 7.3 Hz, 2H), 2.39 (s, 3H), 1.22 (t, *J* = 7.0 Hz, 3H) ppm; LRMS (ES$^-$) calcd 358.02 for $C_{15}H_{15}Cl_2NO_3P^-$ found 357.6 ($M^-$).

*Dataset 3: NMR and IR characterization spectra*

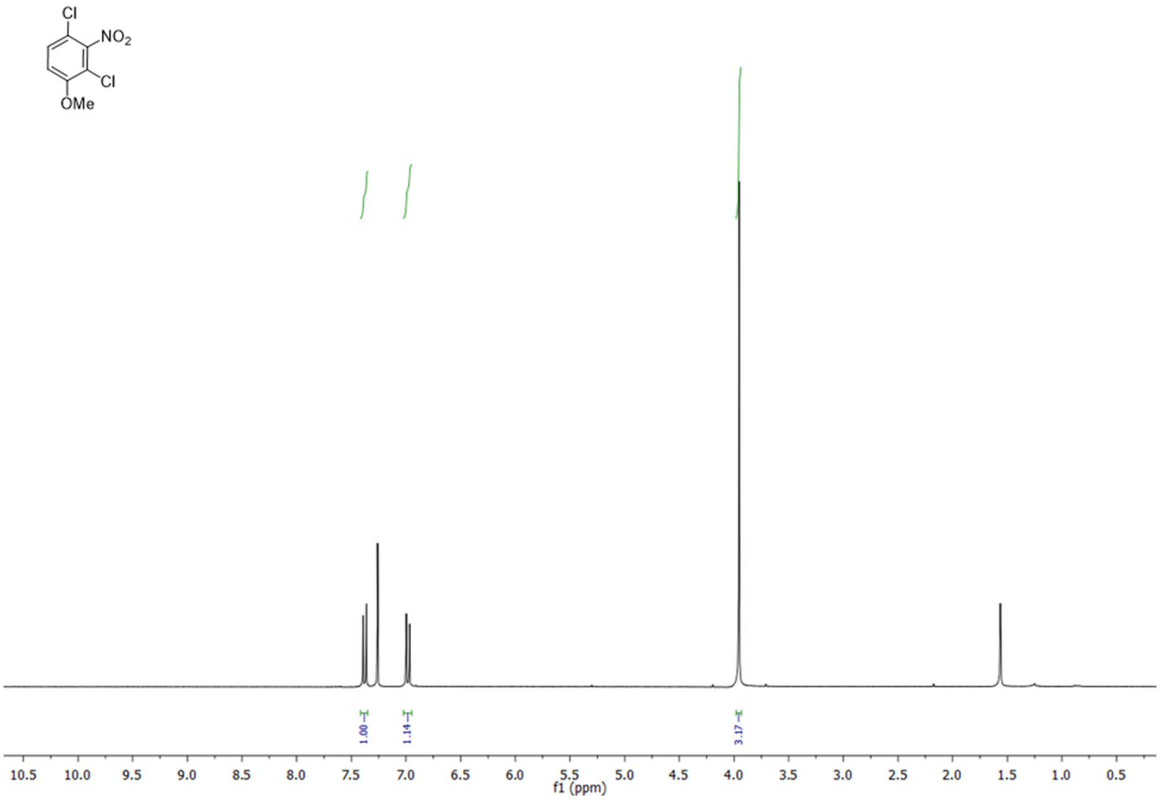

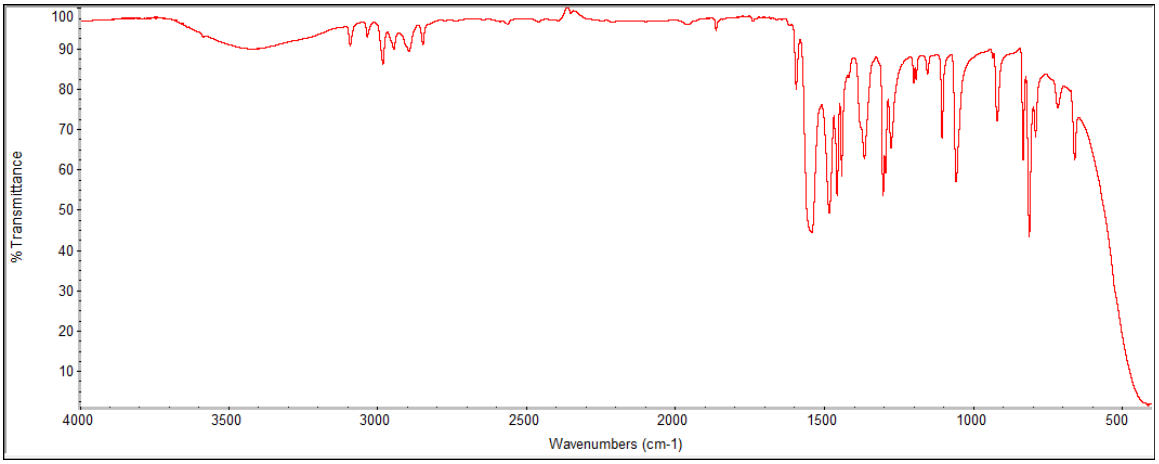

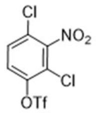

Cl
NO2
Cl
OTf

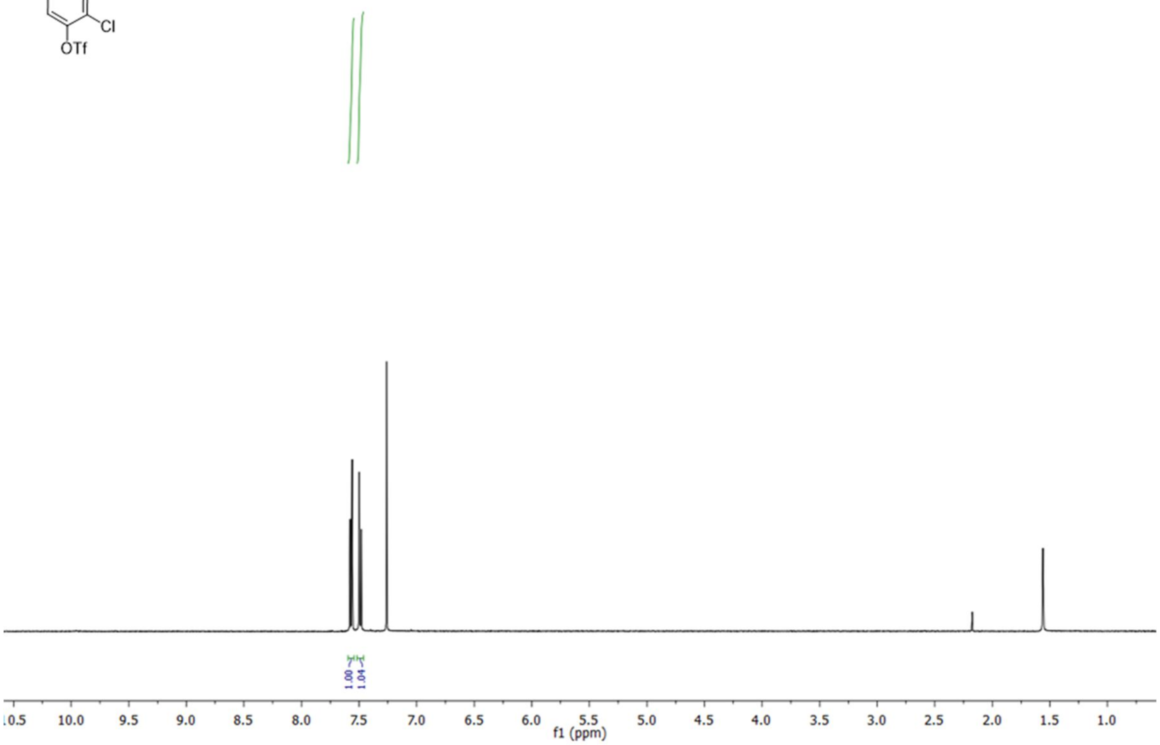

1.00
1.04
f1 (ppm)

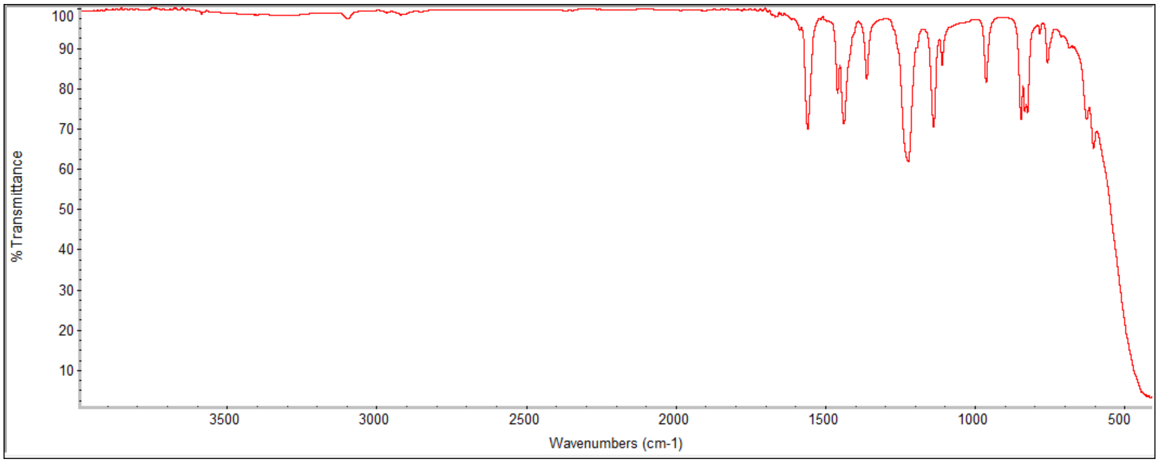

% Transmittance
Wavenumbers (cm-1)

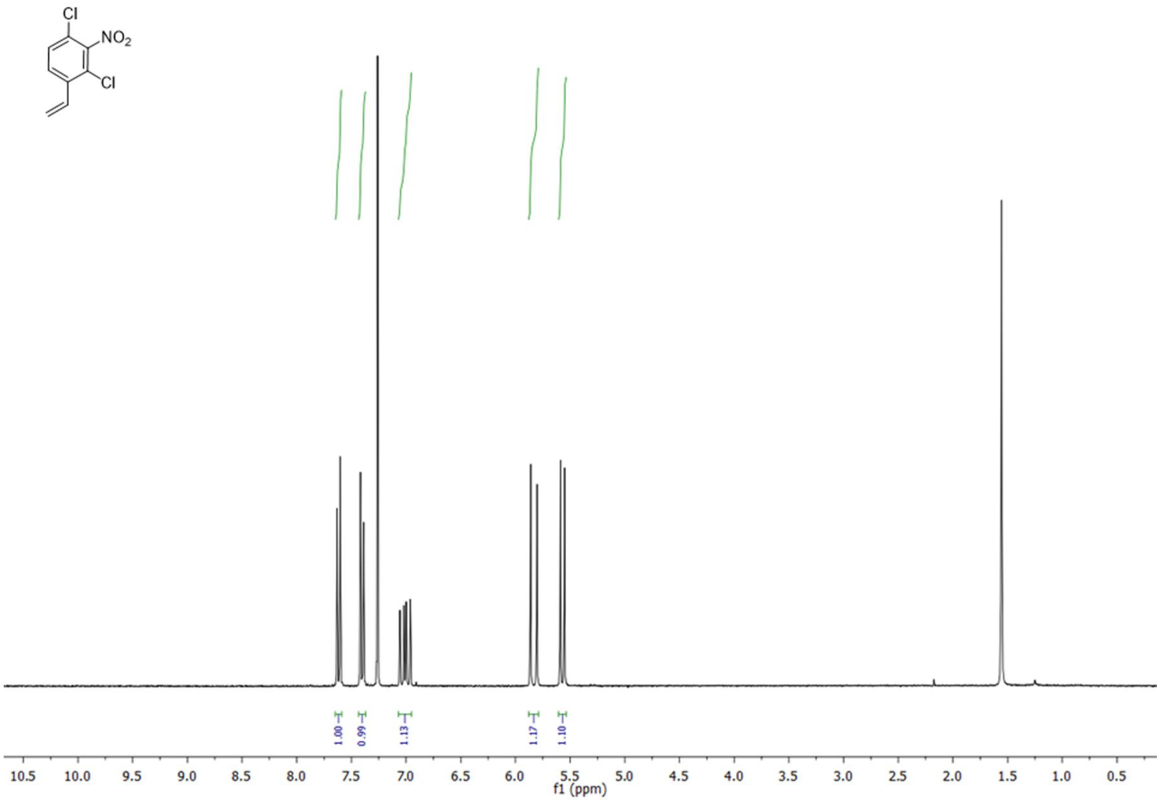
Cl
NO2
Cl
1.00
0.99
1.13
1.17
1.10
10.5 10.0 9.5 9.0 8.5 8.0 7.5 7.0 6.5 6.0 5.5 5.0 4.5 4.0 3.5 3.0 2.5 2.0 1.5 1.0 0.5
f1 (ppm)

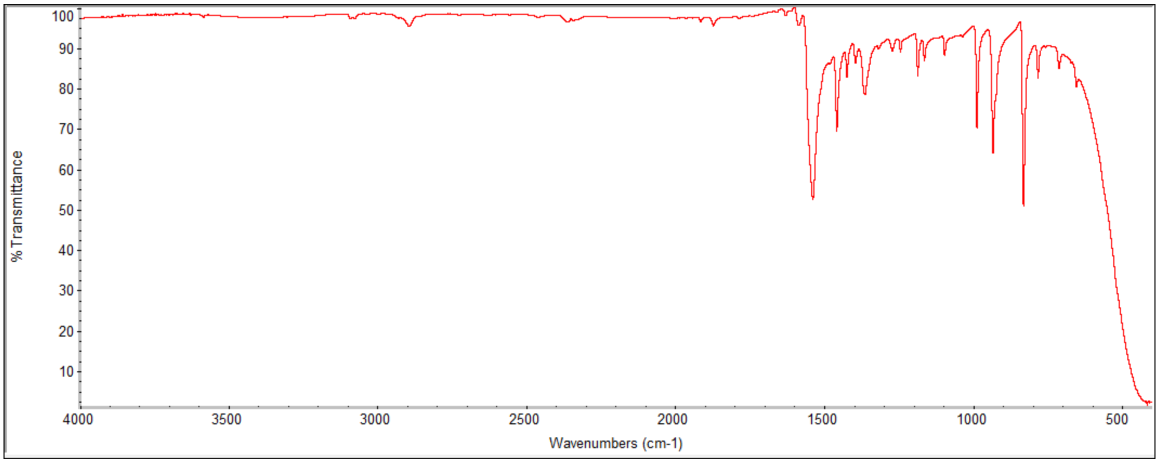
% Transmittance
100 90 80 70 60 50 40 30 20 10
4000 3500 3000 2500 2000 1500 1000 500
Wavenumbers (cm-1)

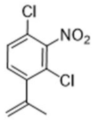
Cl
NO2
Cl

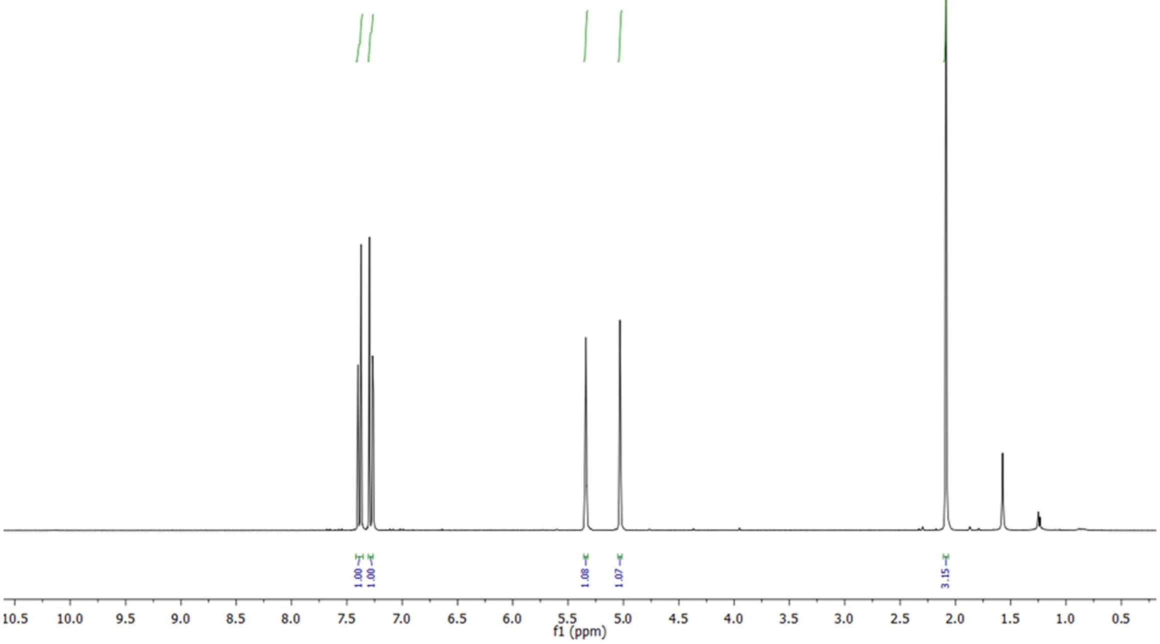
f1 (ppm)

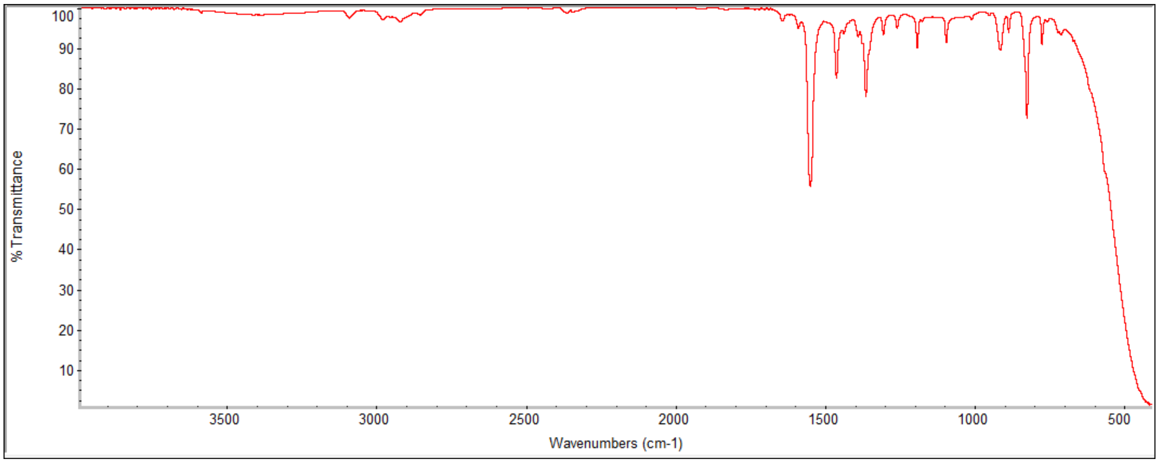
% Transmittance
Wavenumbers (cm-1)

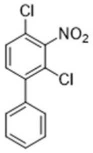
Cl
NO2
Cl

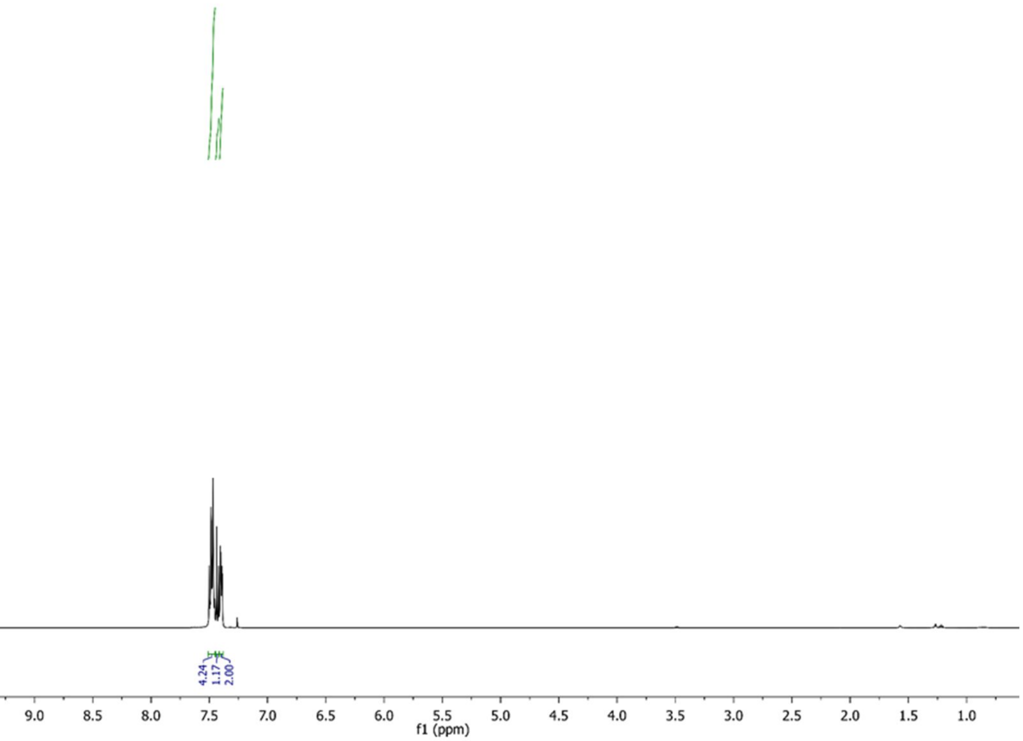
4.24
1.17
2.00
10.0 9.5 9.0 8.5 8.0 7.5 7.0 6.5 6.0 5.5 5.0 4.5 4.0 3.5 3.0 2.5 2.0 1.5 1.0
f1 (ppm)

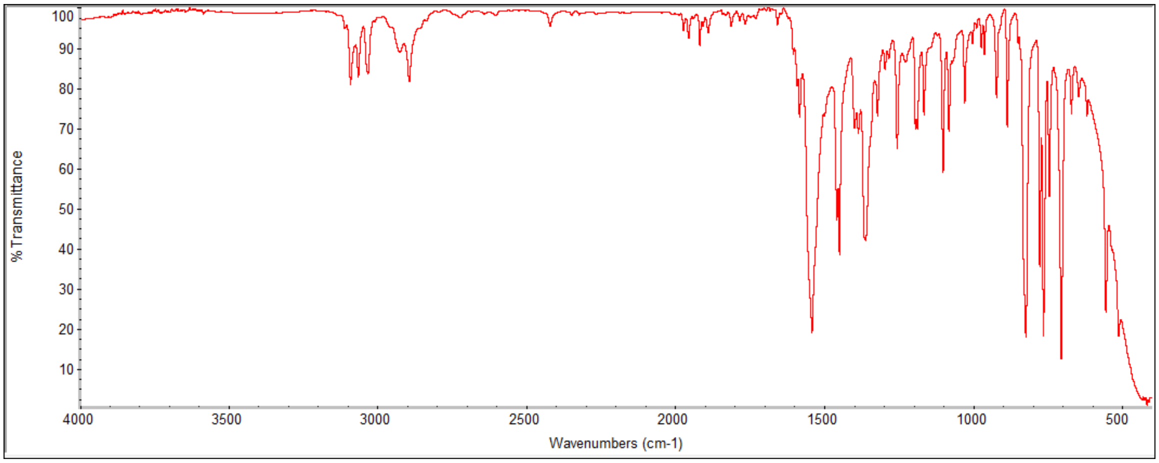
% Transmittance
100 90 80 70 60 50 40 30 20 10
4000 3500 3000 2500 2000 1500 1000 500
Wavenumbers (cm-1)

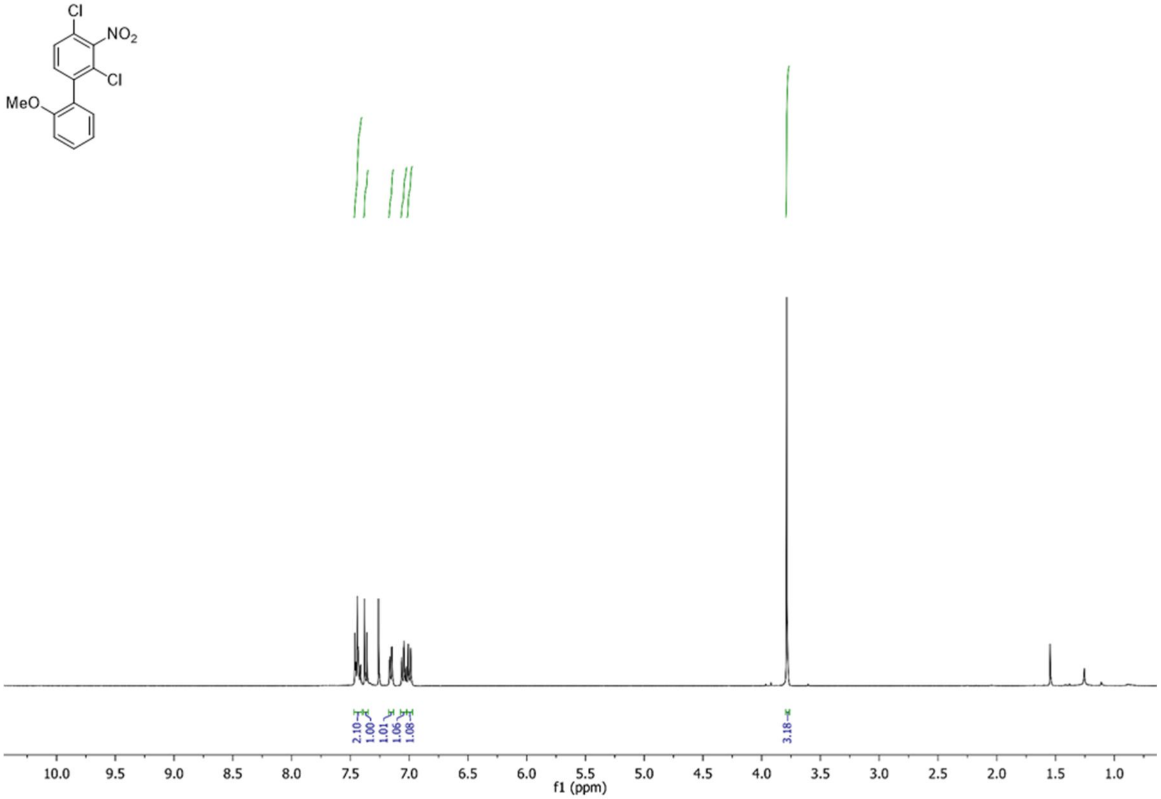
Cl
NO2
Cl
MeO
2.10
1.00
1.01
1.06
1.08
3.18
10.0 9.5 9.0 8.5 8.0 7.5 7.0 6.5 6.0 5.5 5.0 4.5 4.0 3.5 3.0 2.5 2.0 1.5 1.0
f1 (ppm)

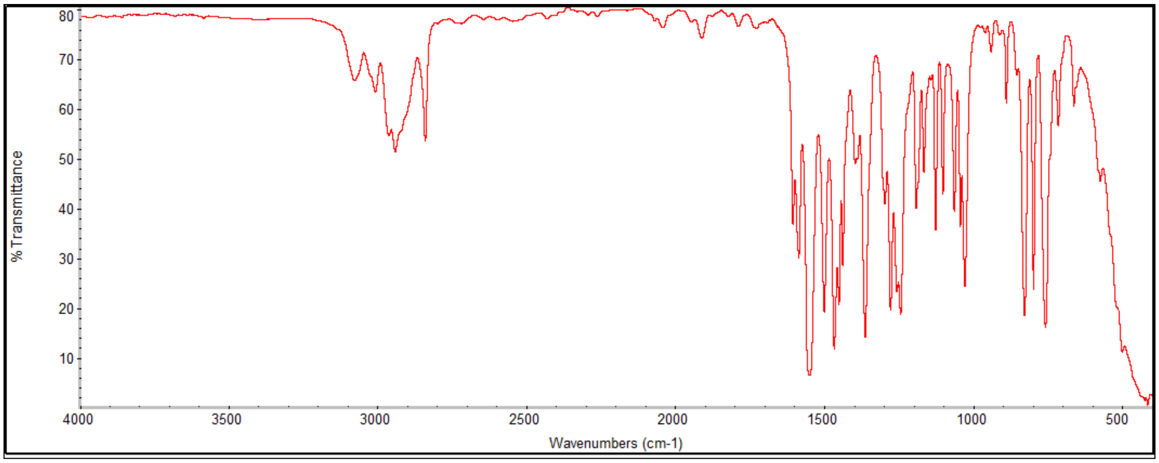
% Transmittance
80 70 60 50 40 30 20 10
4000 3500 3000 2500 2000 1500 1000 500
Wavenumbers (cm-1)

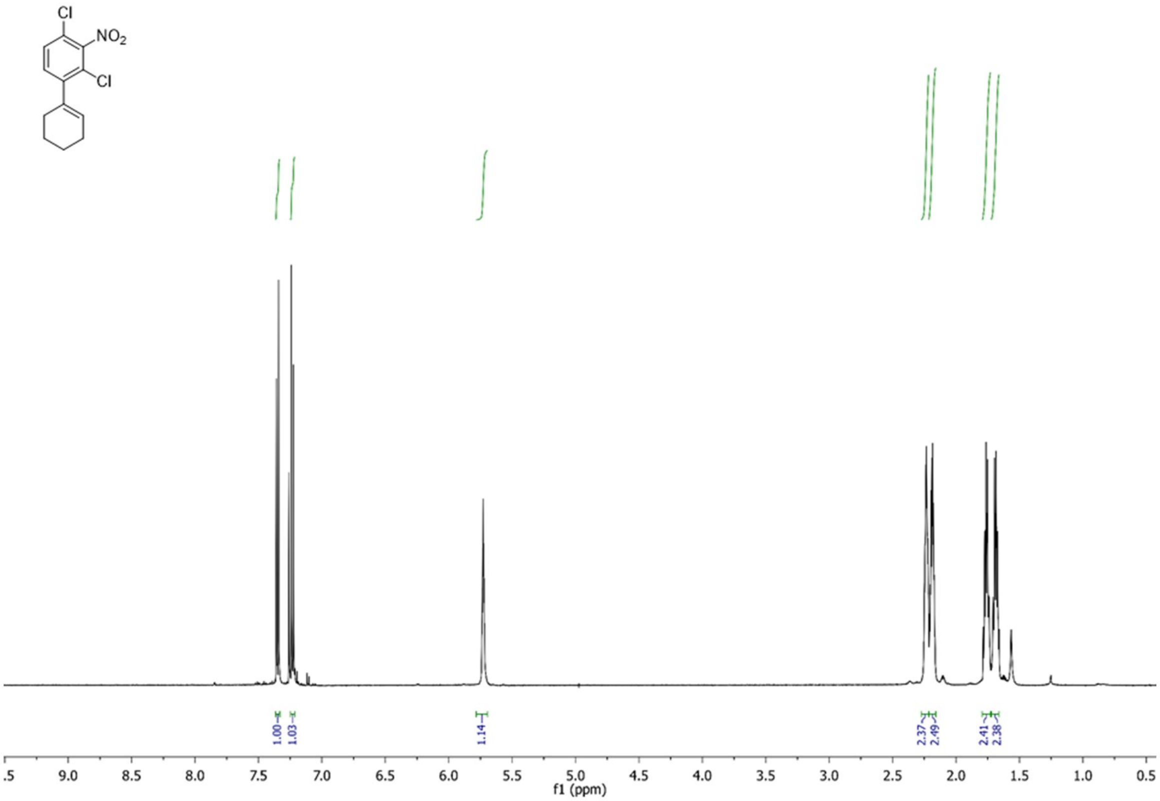
Cl
NO2
Cl
1.00
1.03
1.14
2.37
2.49
2.41
2.38
f1 (ppm)

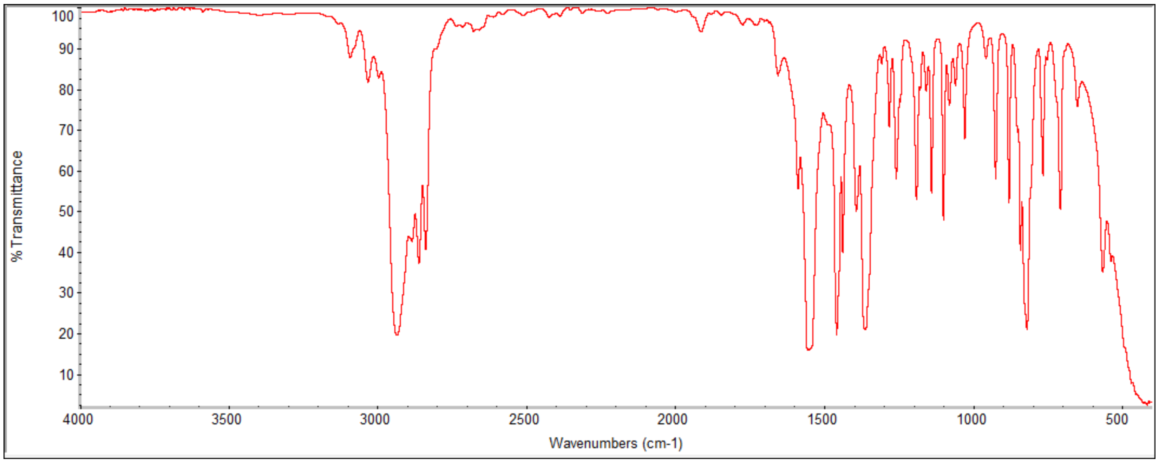
% Transmittance
Wavenumbers (cm-1)

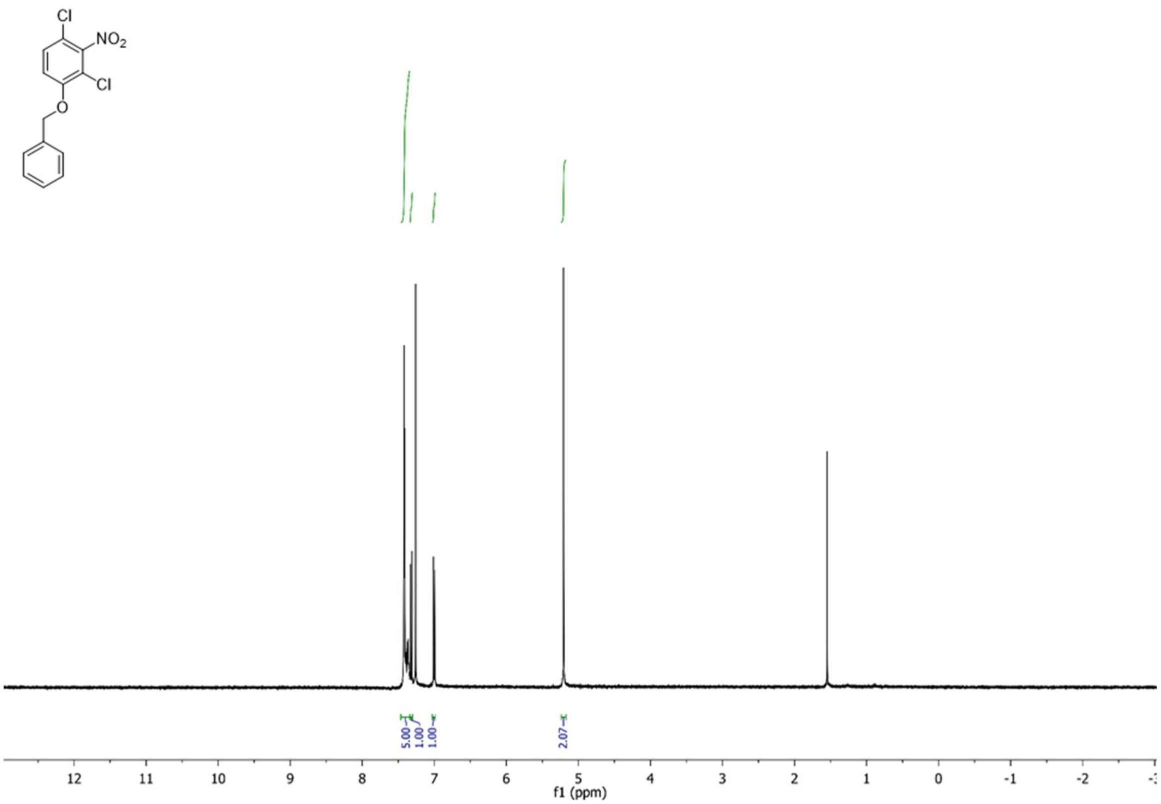
Cl
NO2
Cl
O
5.00
1.00
1.00
2.07
12 11 10 9 8 7 6 5 4 3 2 1 0 -1 -2
f1 (ppm)

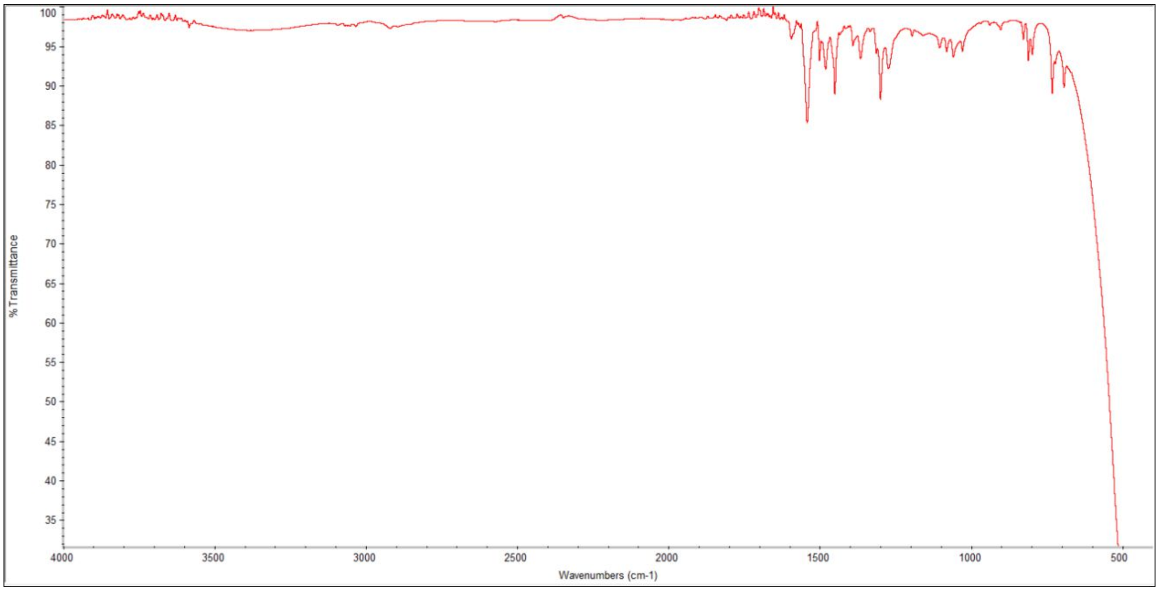
100 95 90 85 80 75 70 65 60 55 50 45 40 35
% Transmittance
4000 3500 3000 2500 2000 1500 1000 500
Wavenumbers (cm-1)

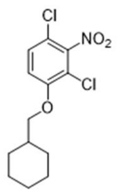
Cl
NO2
Cl
O

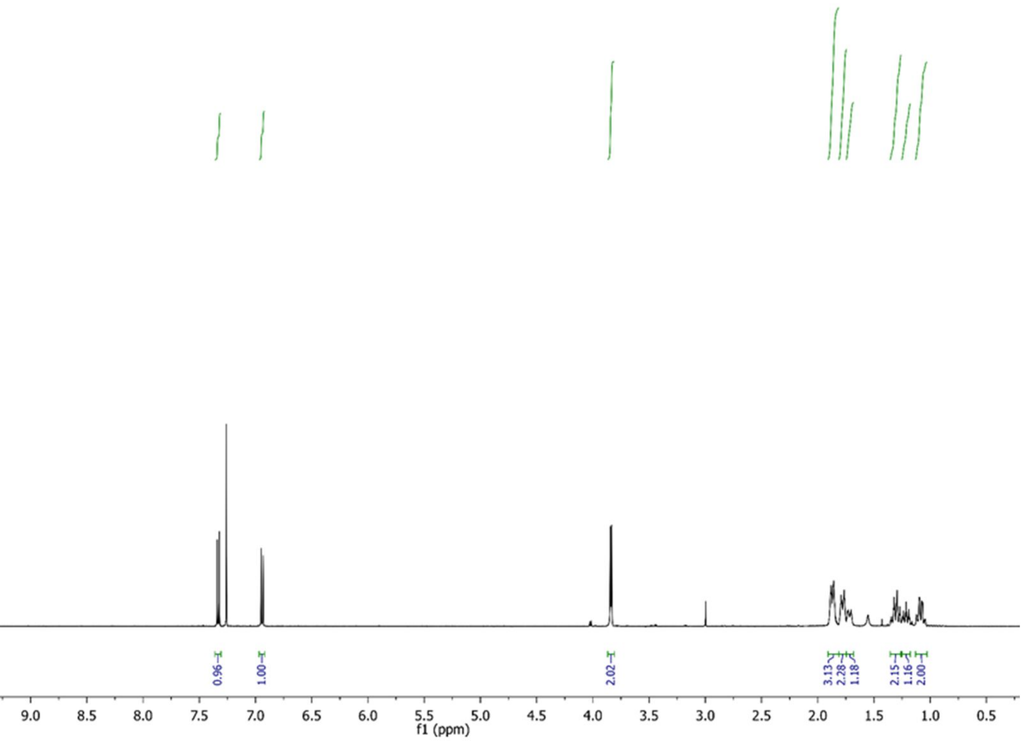
0.96
1.00
2.02
3.13
2.28
1.18
2.15
1.16
2.00
10.0 9.5 9.0 8.5 8.0 7.5 7.0 6.5 6.0 5.5 5.0 4.5 4.0 3.5 3.0 2.5 2.0 1.5 1.0 0.5
f1 (ppm)

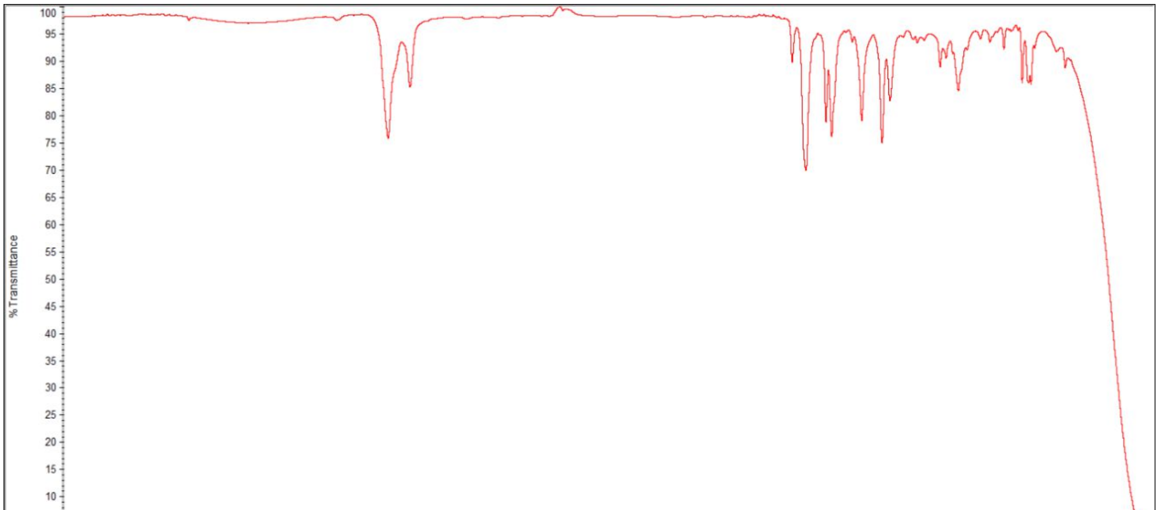
% Transmittance
Wavenumbers (cm-1)
4000 3500 3000 2500 2000 1500 1000 500

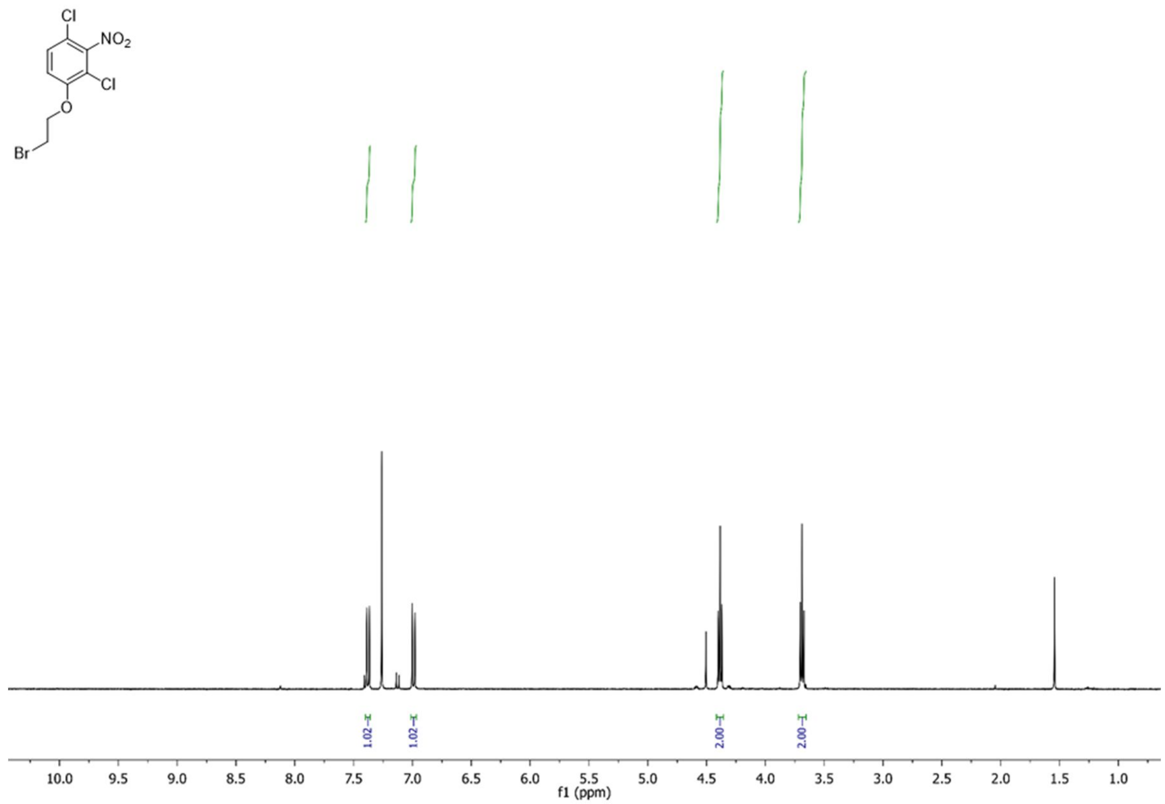
Cl
NO2
Cl
O
Br
1.02
1.02
2.00
2.00
10.0 9.5 9.0 8.5 8.0 7.5 7.0 6.5 6.0 5.5 5.0 4.5 4.0 3.5 3.0 2.5 2.0 1.5 1.0
f1 (ppm)

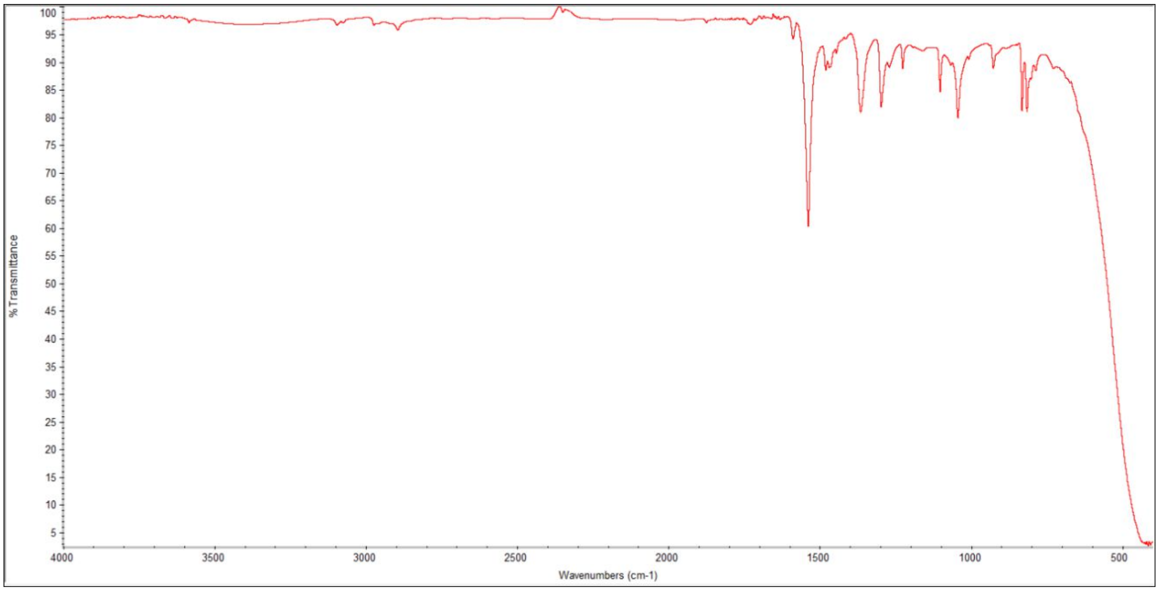
% Transmittance
Wavenumbers (cm-1)
4000 3500 3000 2500 2000 1500 1000 500

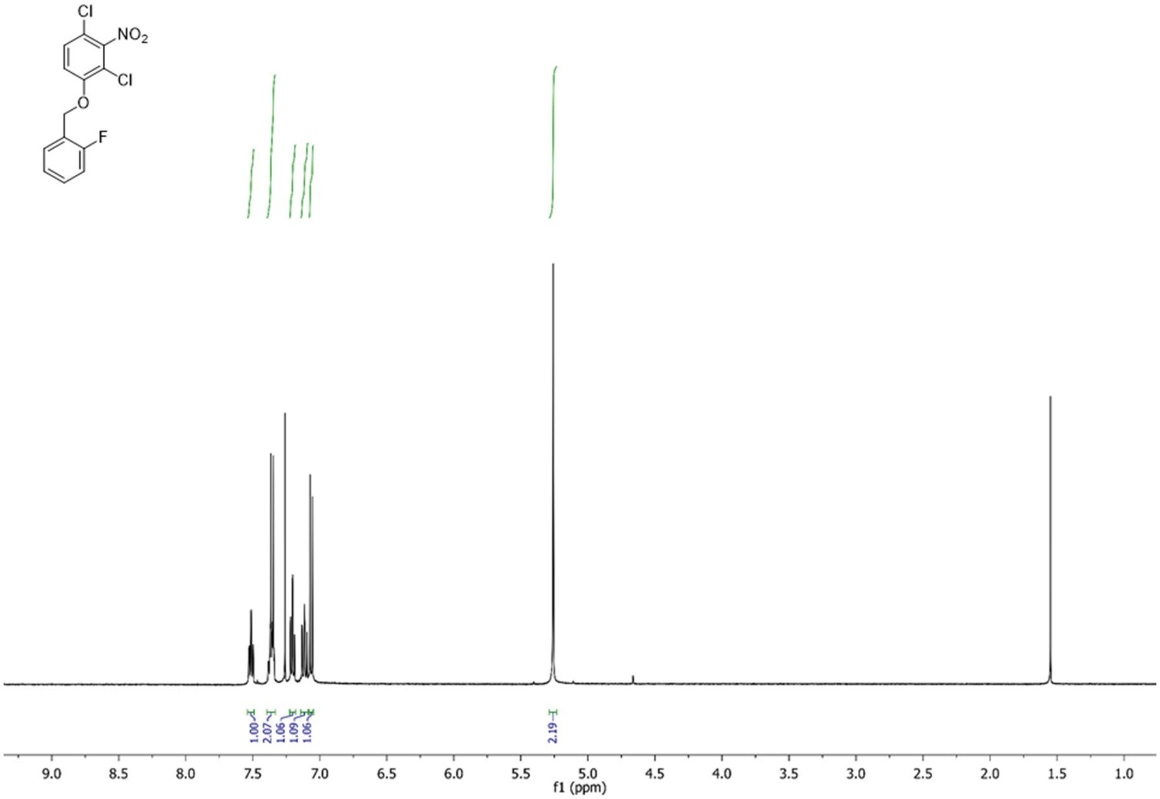
Cl
NO2
Cl
O
F
1.00
2.07
1.06
1.09
1.06
2.19
9.0 8.5 8.0 7.5 7.0 6.5 6.0 5.5 5.0 4.5 4.0 3.5 3.0 2.5 2.0 1.5 1.0
f1 (ppm)

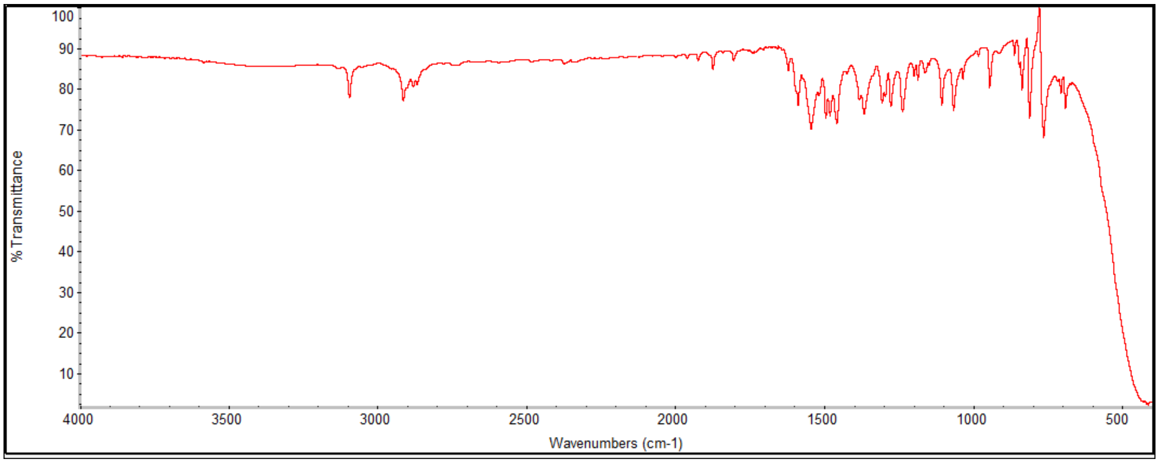
% Transmittance
100 90 80 70 60 50 40 30 20 10
4000 3500 3000 2500 2000 1500 1000 500
Wavenumbers (cm-1)

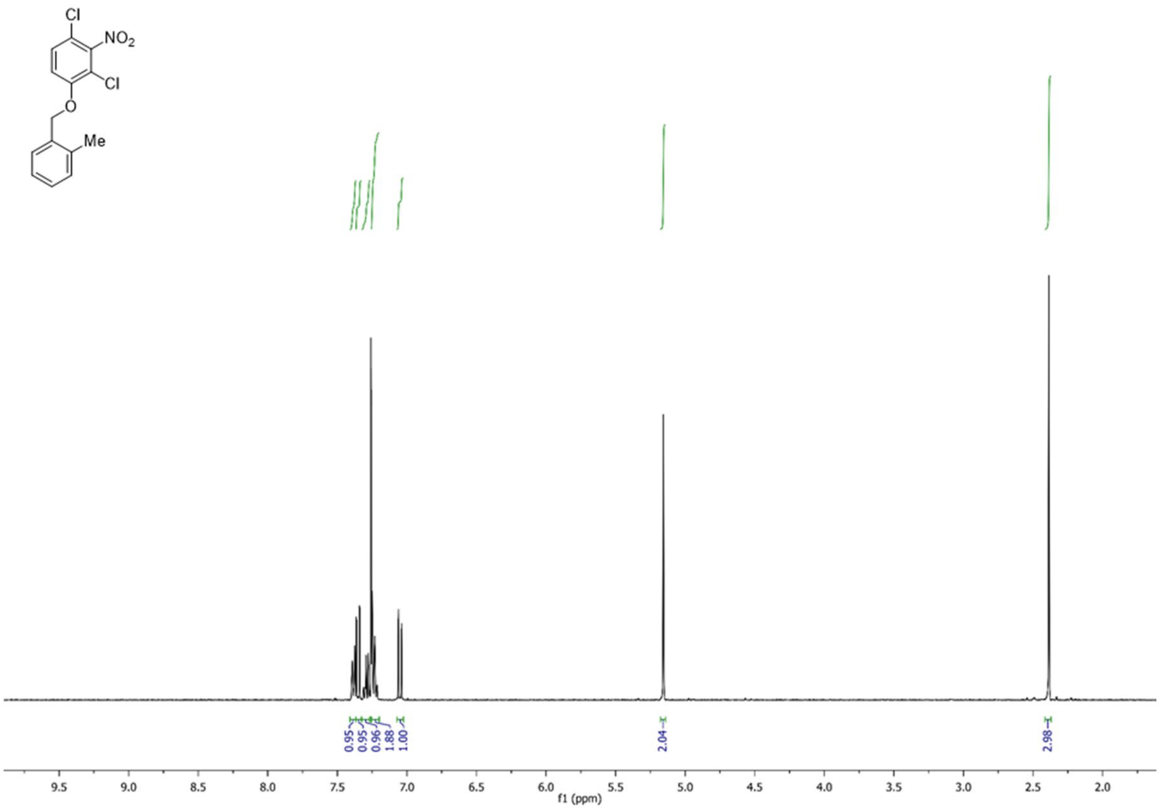
Cl
NO2
Cl
O
Me
0.95
0.95
0.96
1.88
1.00
2.04
2.98
9.5
9.0
8.5
8.0
7.5
7.0
6.5
6.0
5.5
5.0
4.5
4.0
3.5
3.0
2.5
2.0
f1 (ppm)

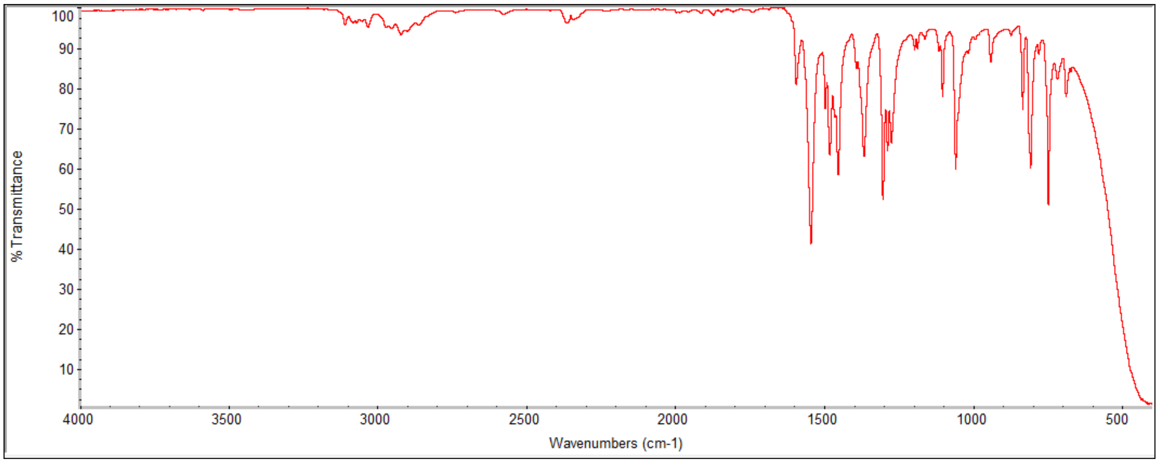
% Transmittance
100
90
80
70
60
50
40
30
20
10
4000
3500
3000
2500
2000
1500
1000
500
Wavenumbers (cm-1)

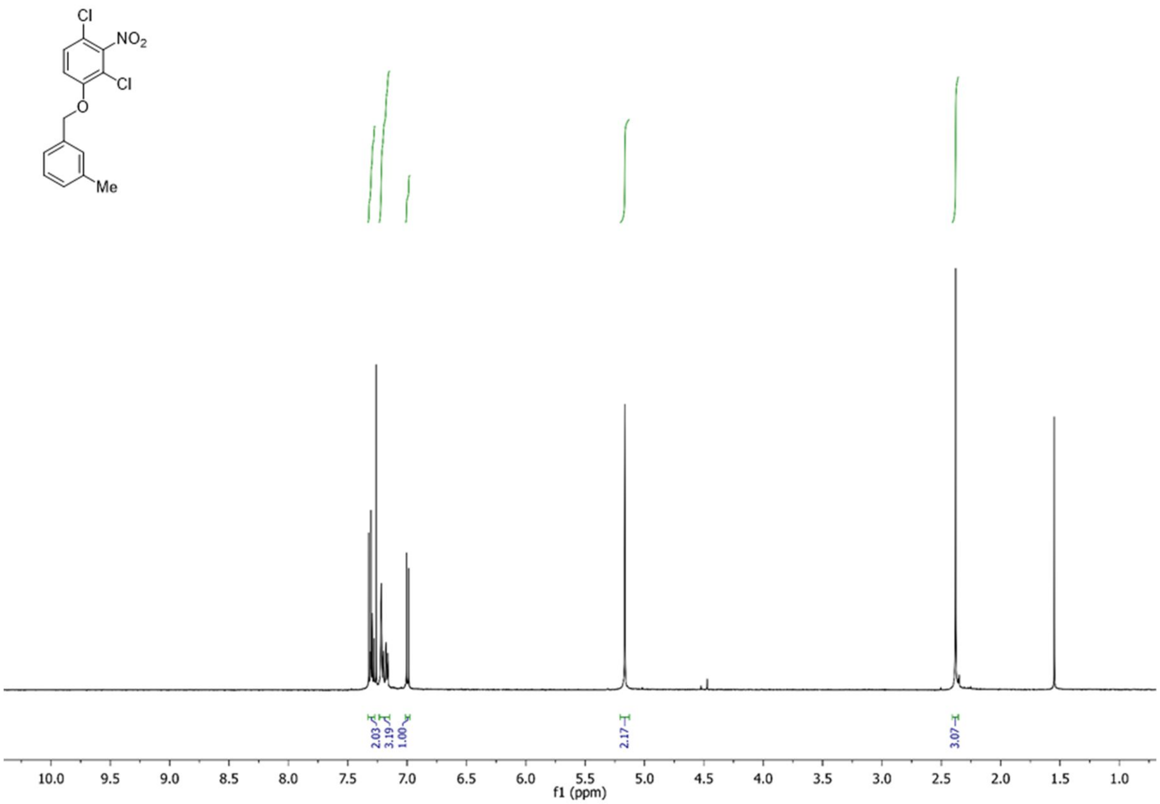
2.03
3.19
1.00
2.17
3.07
10.0
9.5
9.0
8.5
8.0
7.5
7.0
6.5
6.0
5.5
5.0
4.5
4.0
3.5
3.0
2.5
2.0
1.5
1.0
f1 (ppm)

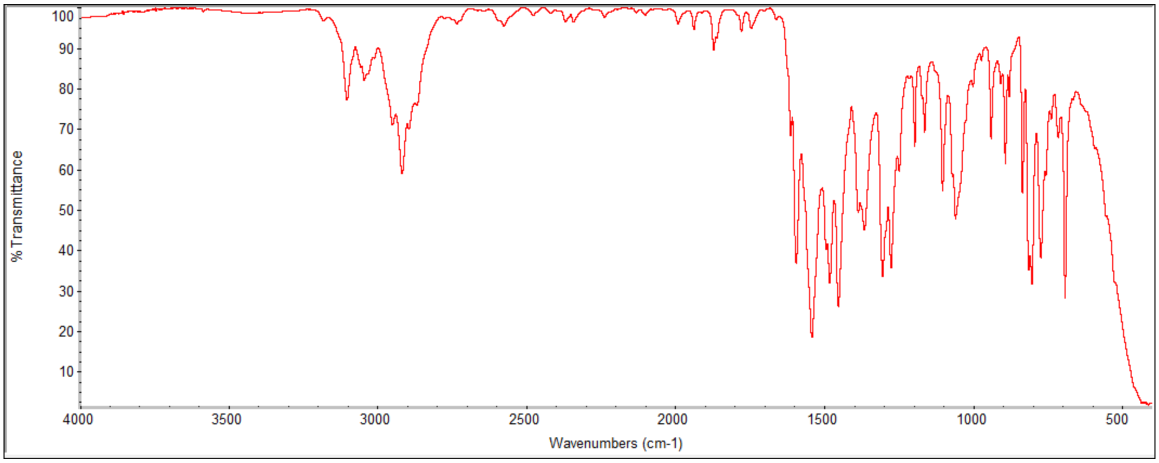
% Transmittance
Wavenumbers (cm-1)
100
90
80
70
60
50
40
30
20
10
4000
3500
3000
2500
2000
1500
1000
500

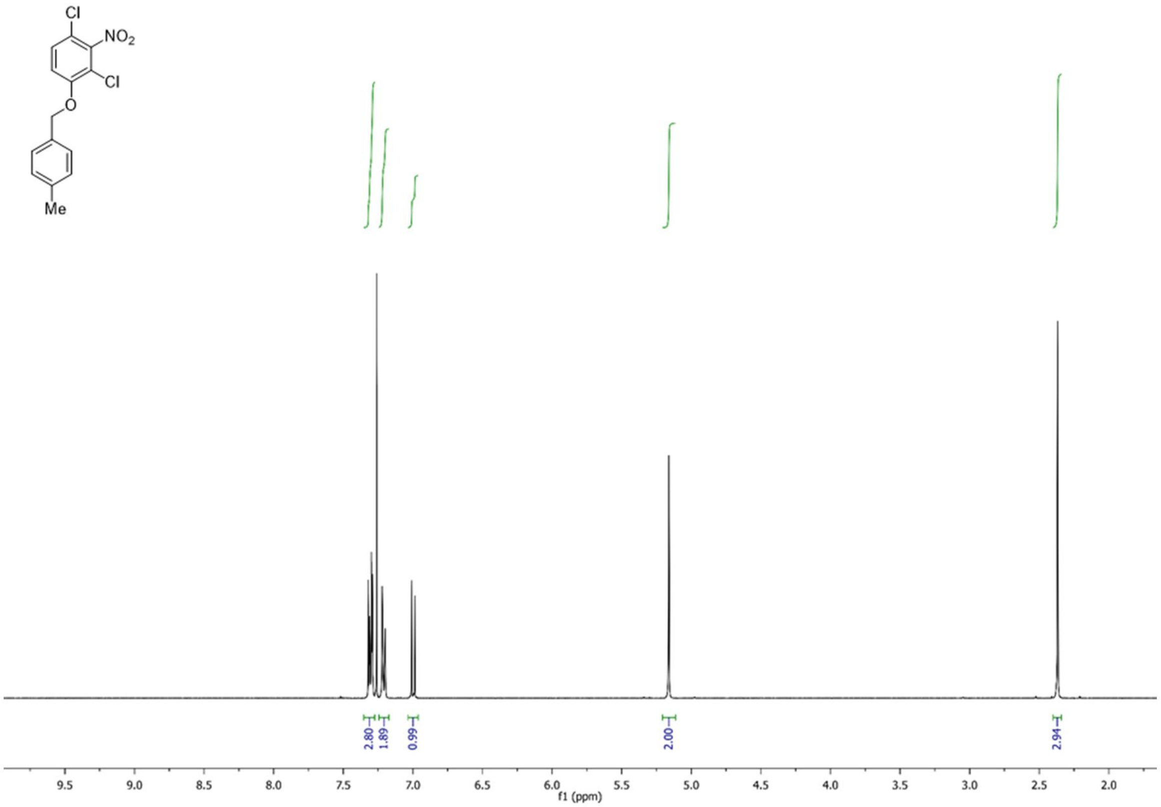
Cl
NO2
Cl
O
Me
2.80
1.89
0.99
2.00
2.94
9.5
9.0
8.5
8.0
7.5
7.0
6.5
6.0
5.5
5.0
4.5
4.0
3.5
3.0
2.5
2.0
f1 (ppm)

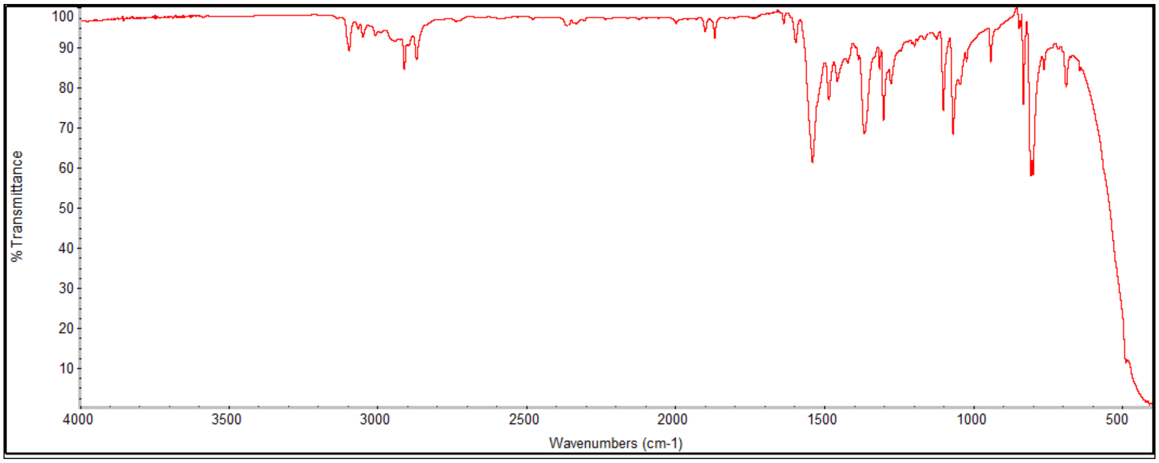
100
90
80
70
60
50
40
30
20
10
% Transmittance
4000
3500
3000
2500
2000
1500
1000
500
Wavenumbers (cm-1)

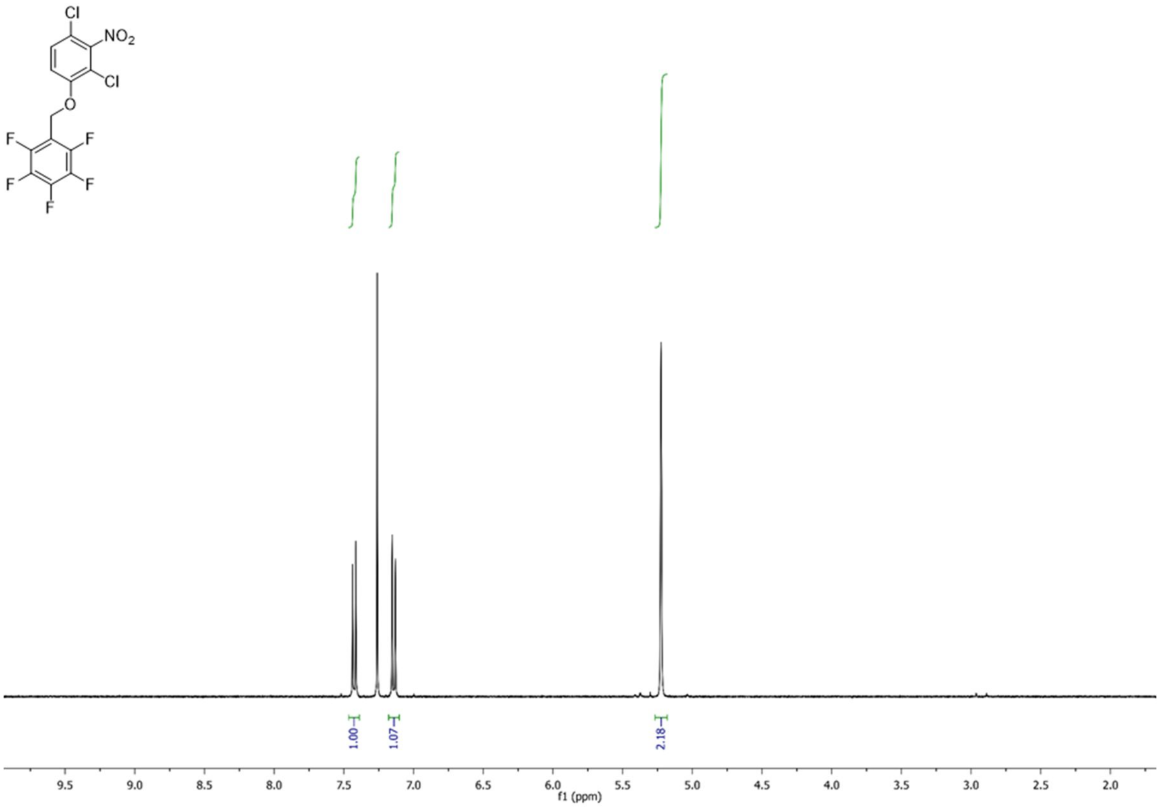
Cl
NO2
Cl
O
F
F
F
F
F
1.00
1.07
2.18
9.5
9.0
8.5
8.0
7.5
7.0
6.5
6.0
5.5
5.0
4.5
4.0
3.5
3.0
2.5
2.0
f1 (ppm)

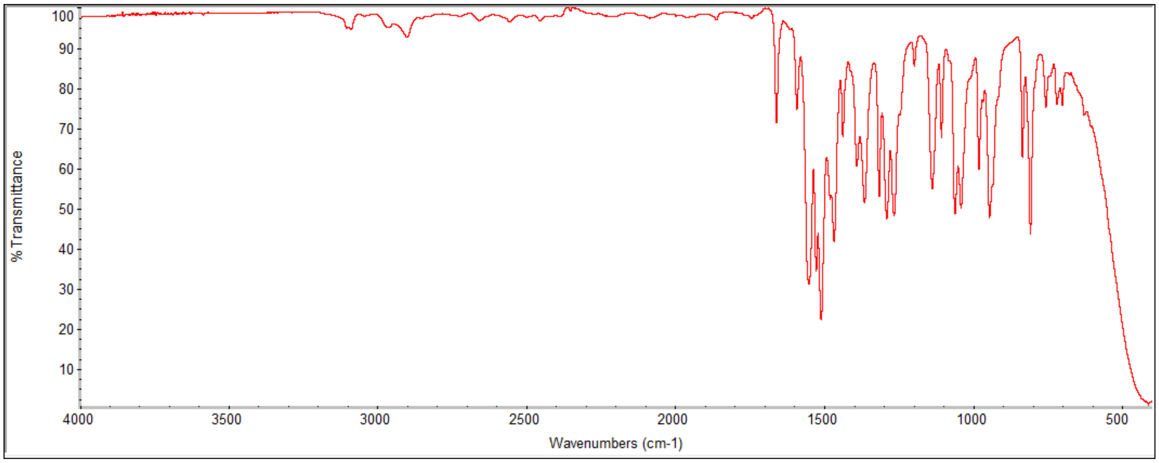
% Transmittance
100
90
80
70
60
50
40
30
20
10
4000
3500
3000
2500
2000
1500
1000
500
Wavenumbers (cm-1)

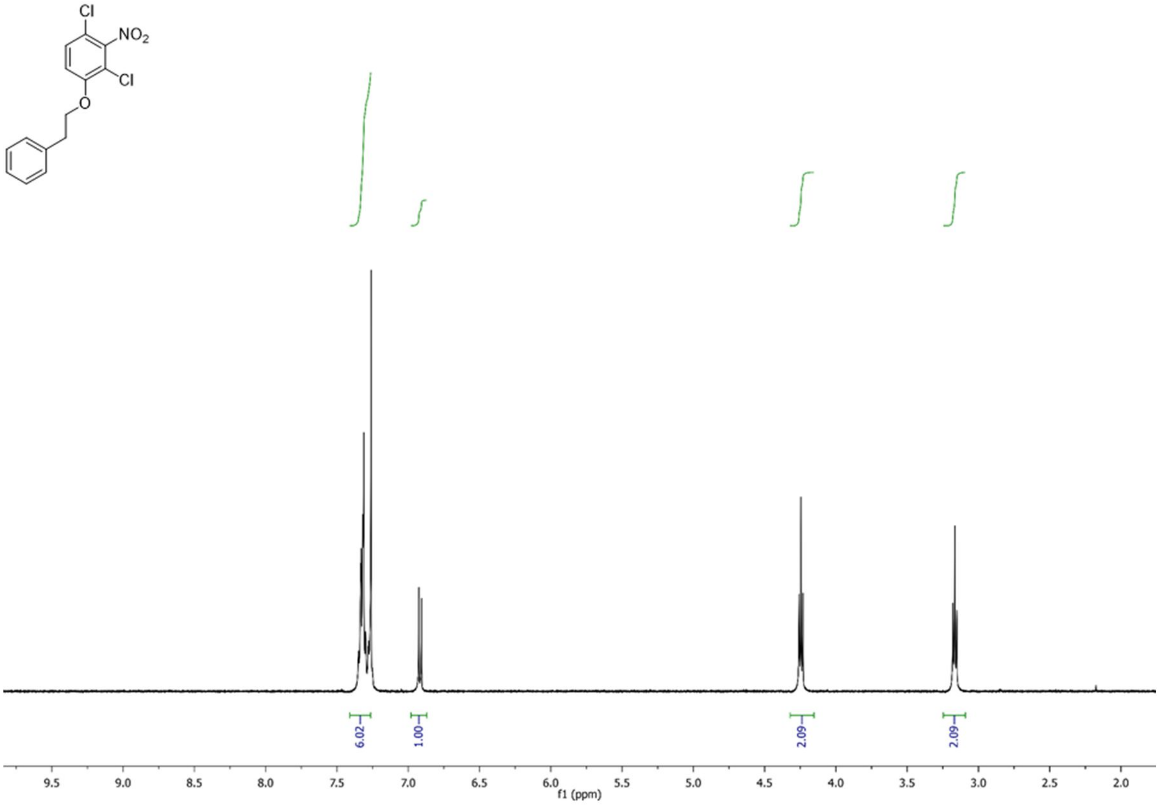

Cl
$NO_2$
Cl
O
6.02
1.00
2.09
2.09
9.5
9.0
8.5
8.0
7.5
7.0
6.5
6.0
5.5
5.0
4.5
4.0
3.5
3.0
2.5
2.0
f1 (ppm)

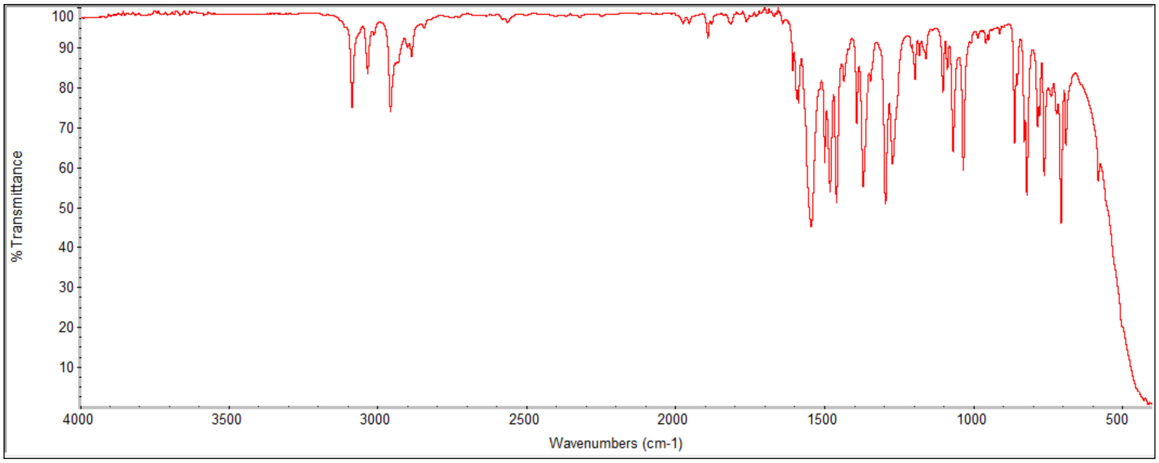

% Transmittance
100
90
80
70
60
50
40
30
20
10
4000
3500
3000
2500
2000
1500
1000
500
Wavenumbers (cm-1)

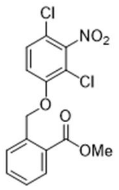
Cl
NO2
Cl
O
O
OMe

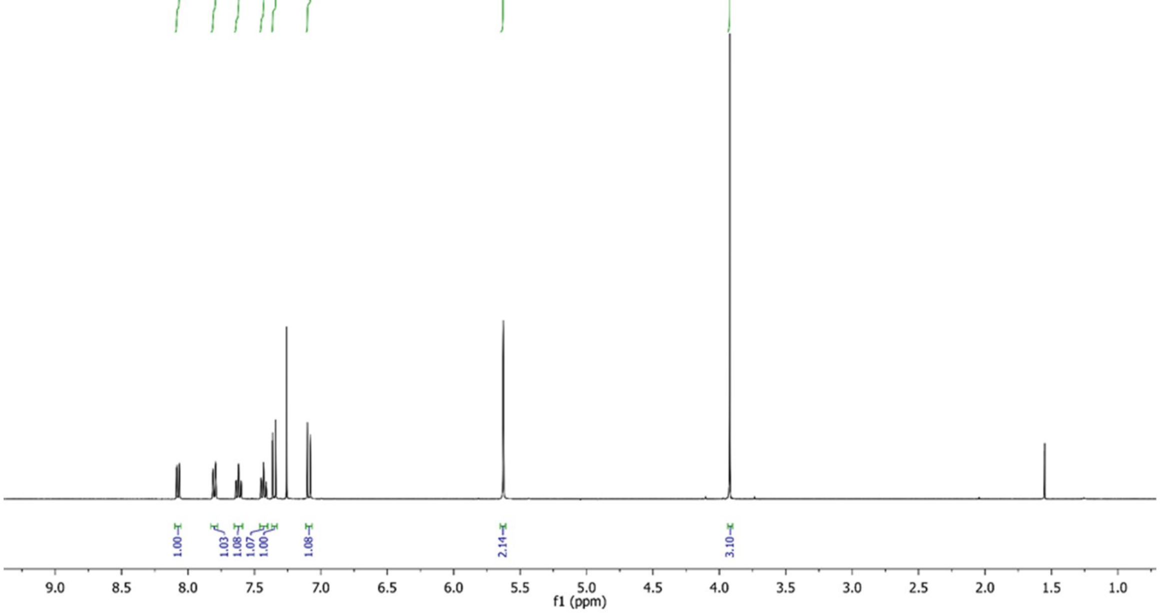
1.00
1.03
1.08
1.07
1.00
1.08
2.14
3.10
9.0
8.5
8.0
7.5
7.0
6.5
6.0
5.5
5.0
4.5
4.0
3.5
3.0
2.5
2.0
1.5
1.0
f1 (ppm)

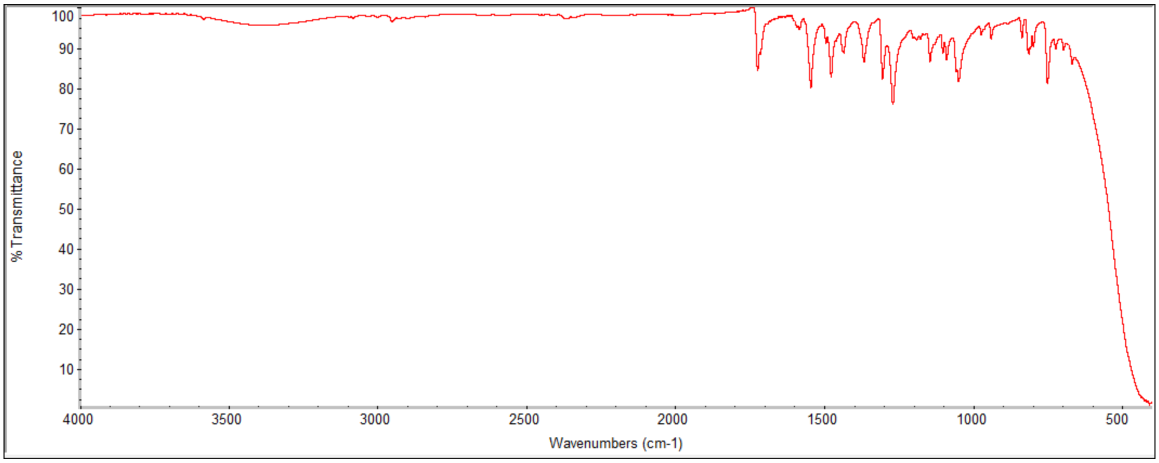
100
90
80
70
60
50
40
30
20
10
% Transmittance
4000
3500
3000
2500
2000
1500
1000
500
Wavenumbers (cm-1)

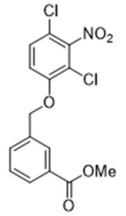
Cl
NO2
Cl
O
OMe
O

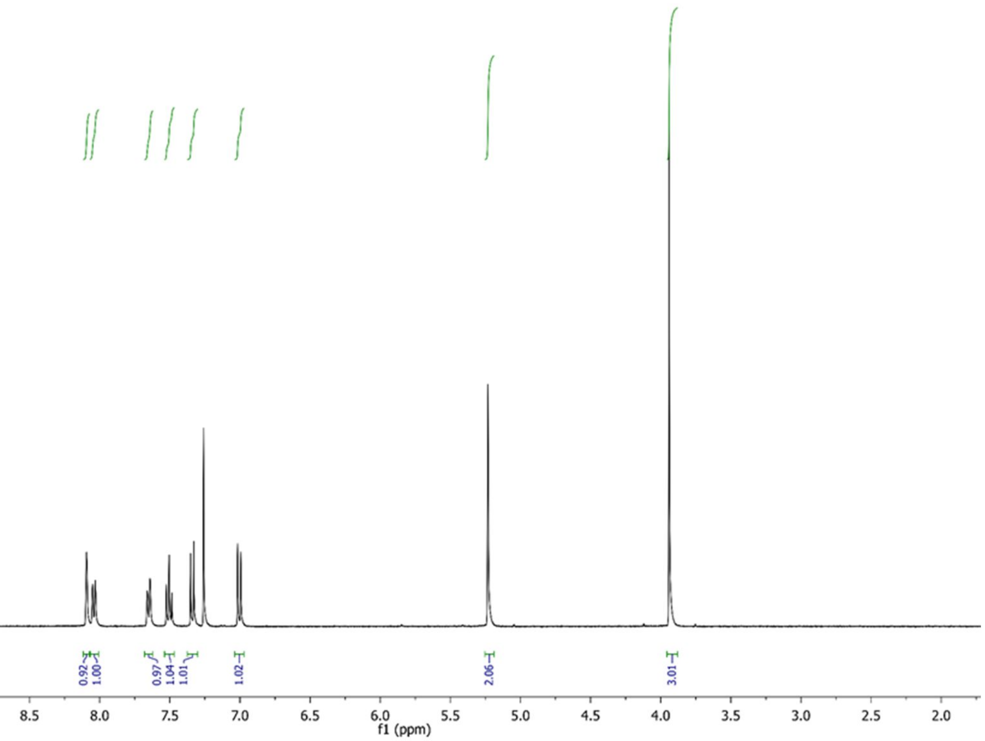
0.92
1.00
0.97
1.04
1.01
1.02
2.06
3.01
9.5
9.0
8.5
8.0
7.5
7.0
6.5
6.0
5.5
5.0
4.5
4.0
3.5
3.0
2.5
2.0
f1 (ppm)

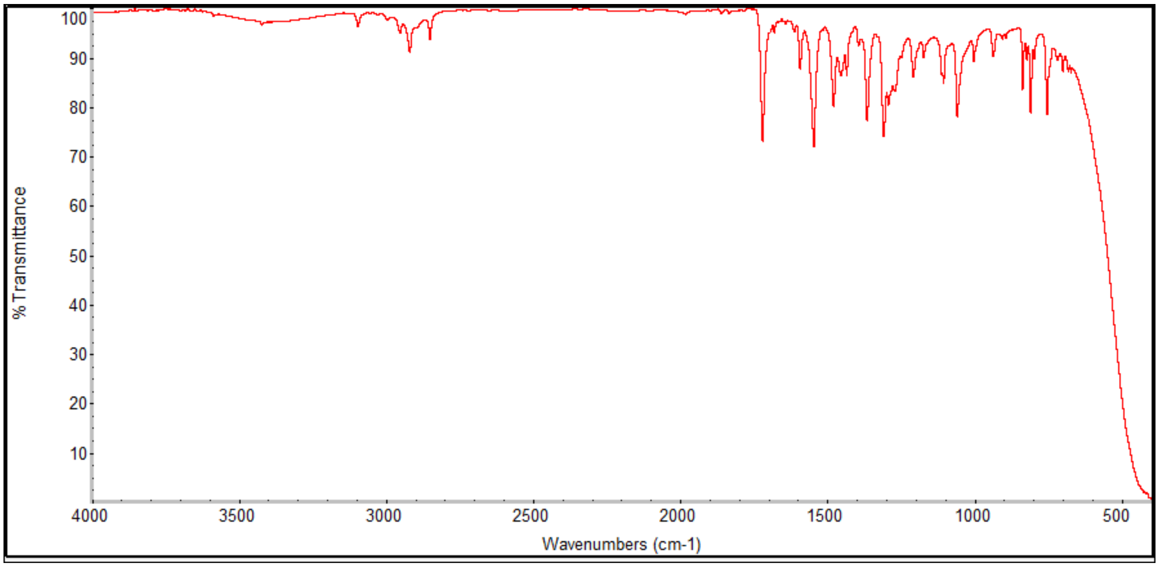
% Transmittance
100
90
80
70
60
50
40
30
20
10
4000
3500
3000
2500
2000
1500
1000
500
Wavenumbers (cm-1)

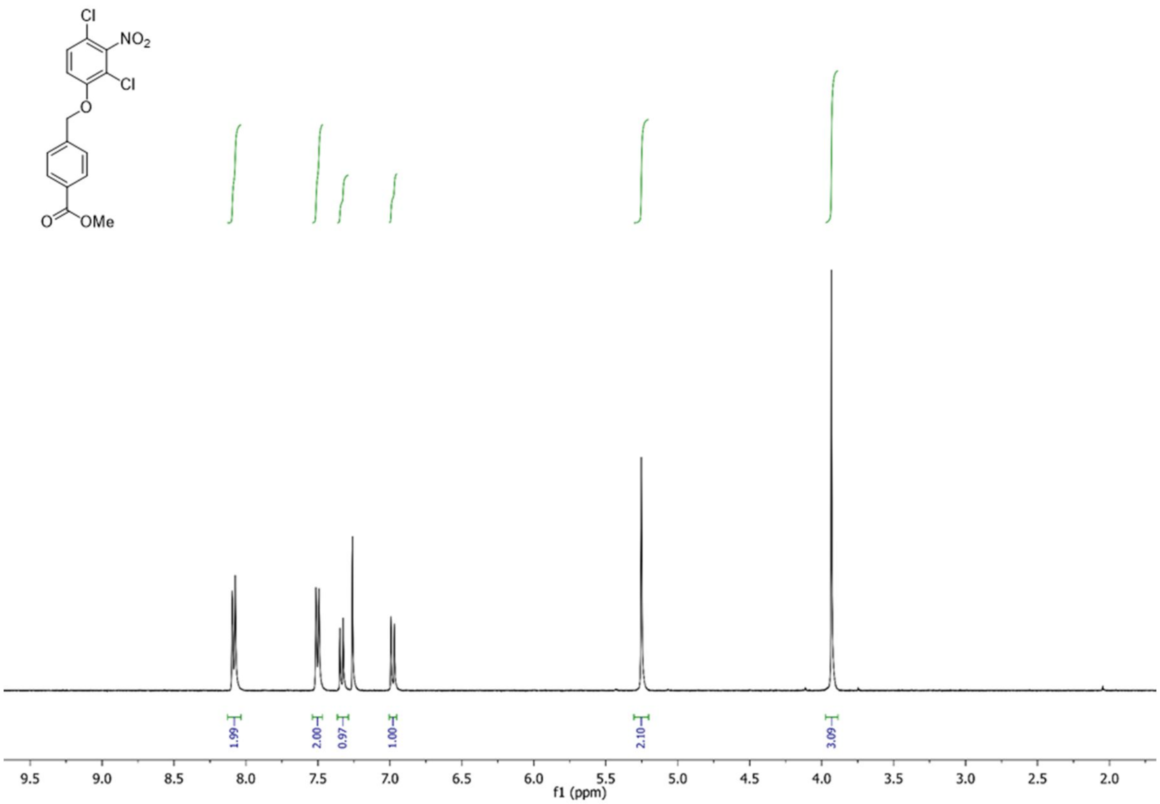
Cl
NO2
Cl
O
O
OMe
1.99
2.00
0.97
1.00
2.10
3.09
9.5
9.0
8.5
8.0
7.5
7.0
6.5
6.0
5.5
5.0
4.5
4.0
3.5
3.0
2.5
2.0
f1 (ppm)

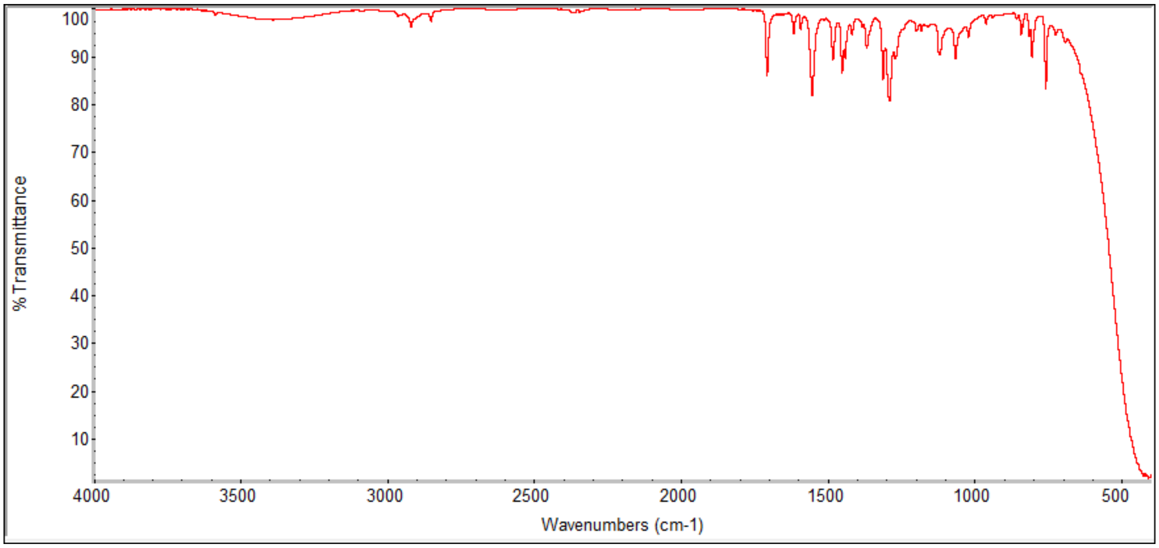
%Transmittance
Wavenumbers (cm-1)
100
90
80
70
60
50
40
30
20
10
4000
3500
3000
2500
2000
1500
1000
500

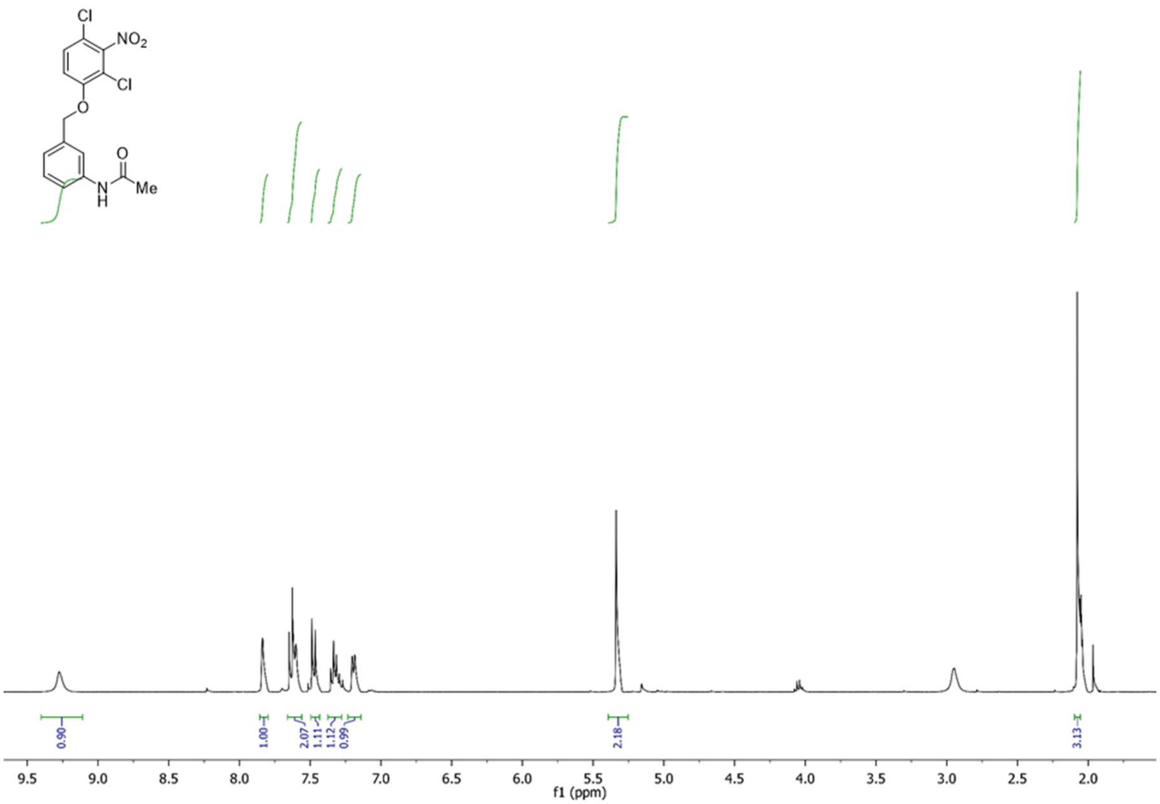
Cl
NO2
Cl
O
O
N
H
Me
0.90
1.00
2.07
1.11
1.12
0.99
2.18
3.13
9.5
9.0
8.5
8.0
7.5
7.0
6.5
6.0
5.5
5.0
4.5
4.0
3.5
3.0
2.5
2.0
f1 (ppm)

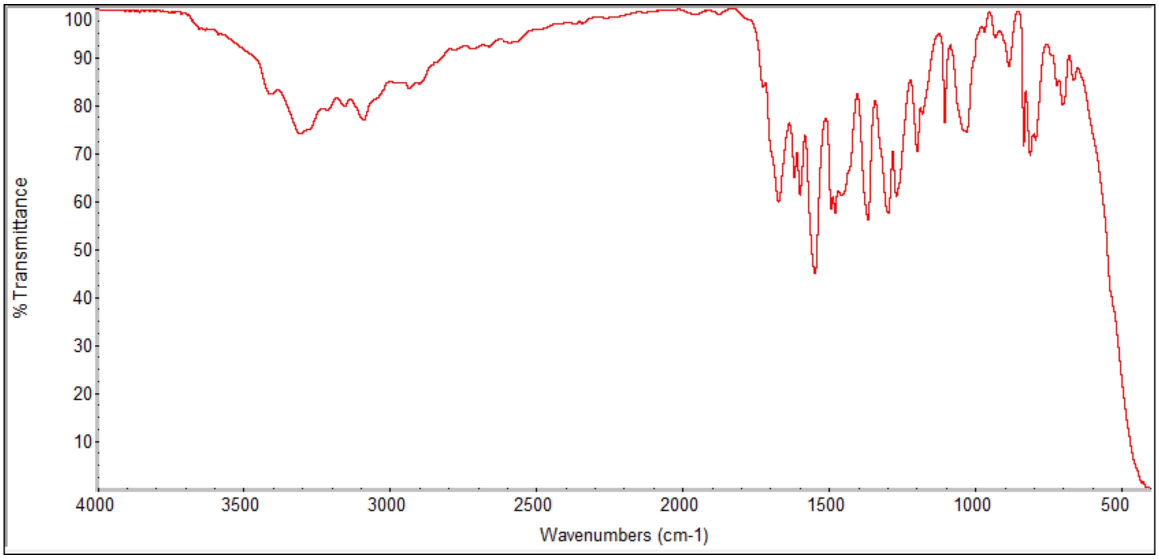
% Transmittance
Wavenumbers (cm-1)

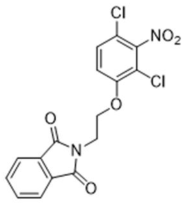
Cl
NO2
Cl
O
O
N
O

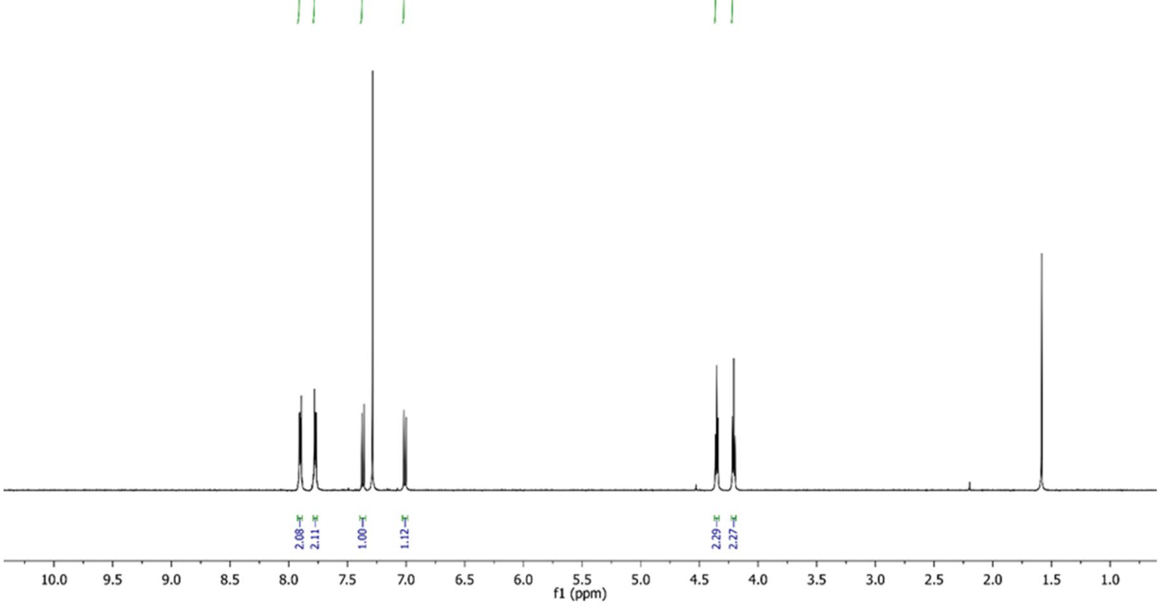
2.08
2.11
1.00
1.12
2.29
2.27
10.0
9.5
9.0
8.5
8.0
7.5
7.0
6.5
6.0
5.5
5.0
4.5
4.0
3.5
3.0
2.5
2.0
1.5
1.0
f1 (ppm)

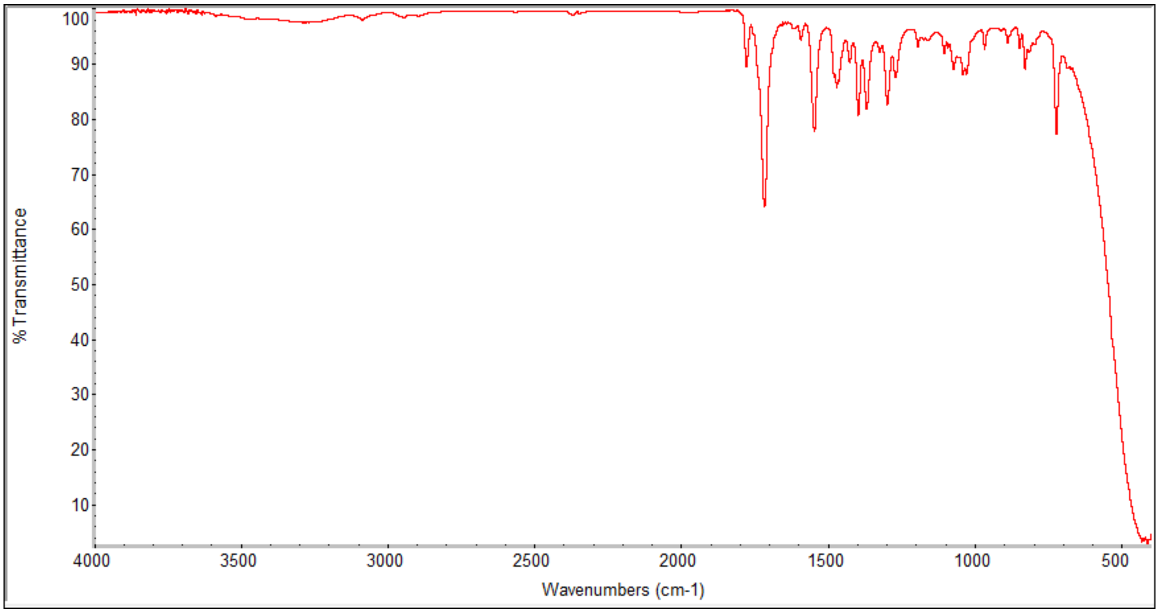
100
90
80
70
60
50
40
30
20
10
% Transmittance
4000
3500
3000
2500
2000
1500
1000
500
Wavenumbers (cm-1)

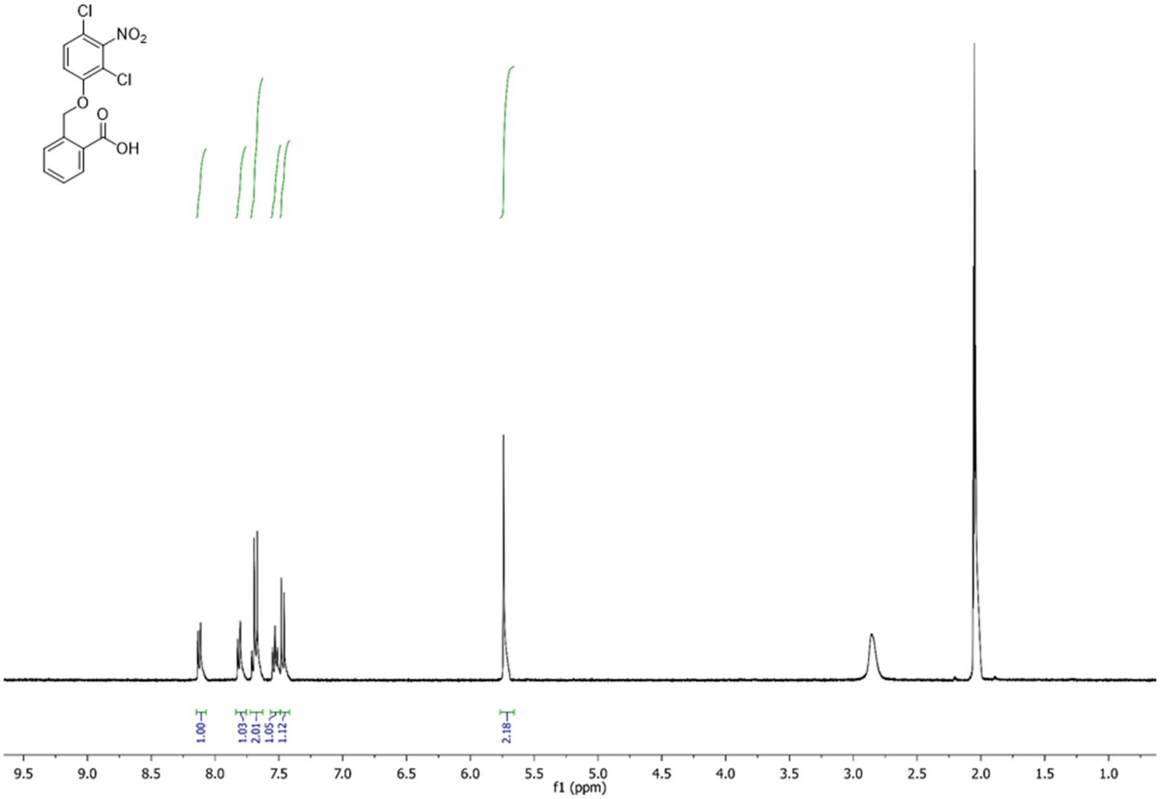
Cl
NO2
Cl
O
O
OH
1.00
1.03
2.01
1.05
1.12
2.18
9.5 9.0 8.5 8.0 7.5 7.0 6.5 6.0 5.5 5.0 4.5 4.0 3.5 3.0 2.5 2.0 1.5 1.0
f1 (ppm)

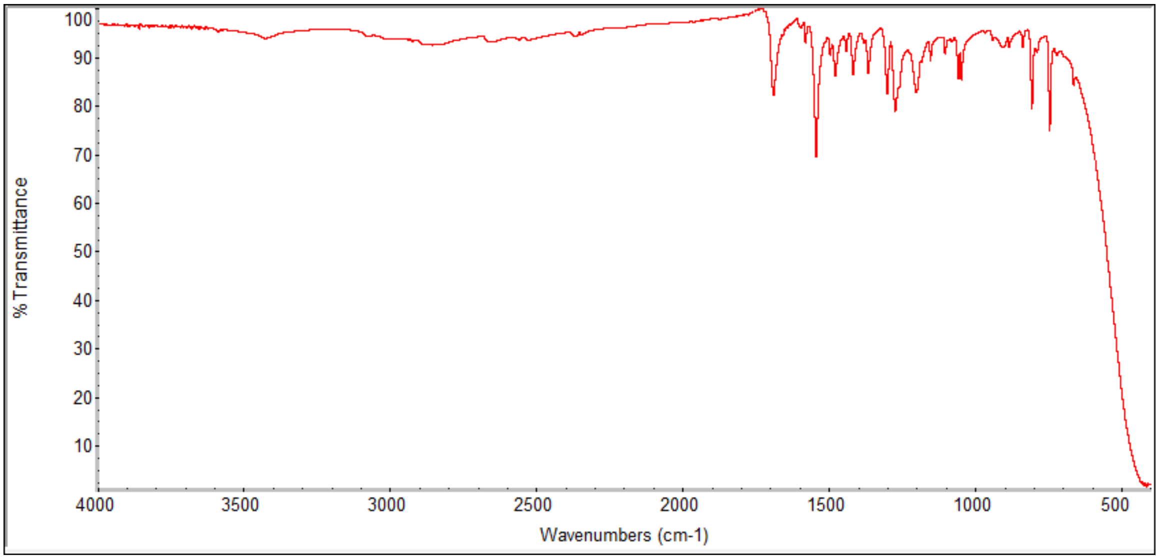
% Transmittance
100 90 80 70 60 50 40 30 20 10
4000 3500 3000 2500 2000 1500 1000 500
Wavenumbers (cm-1)

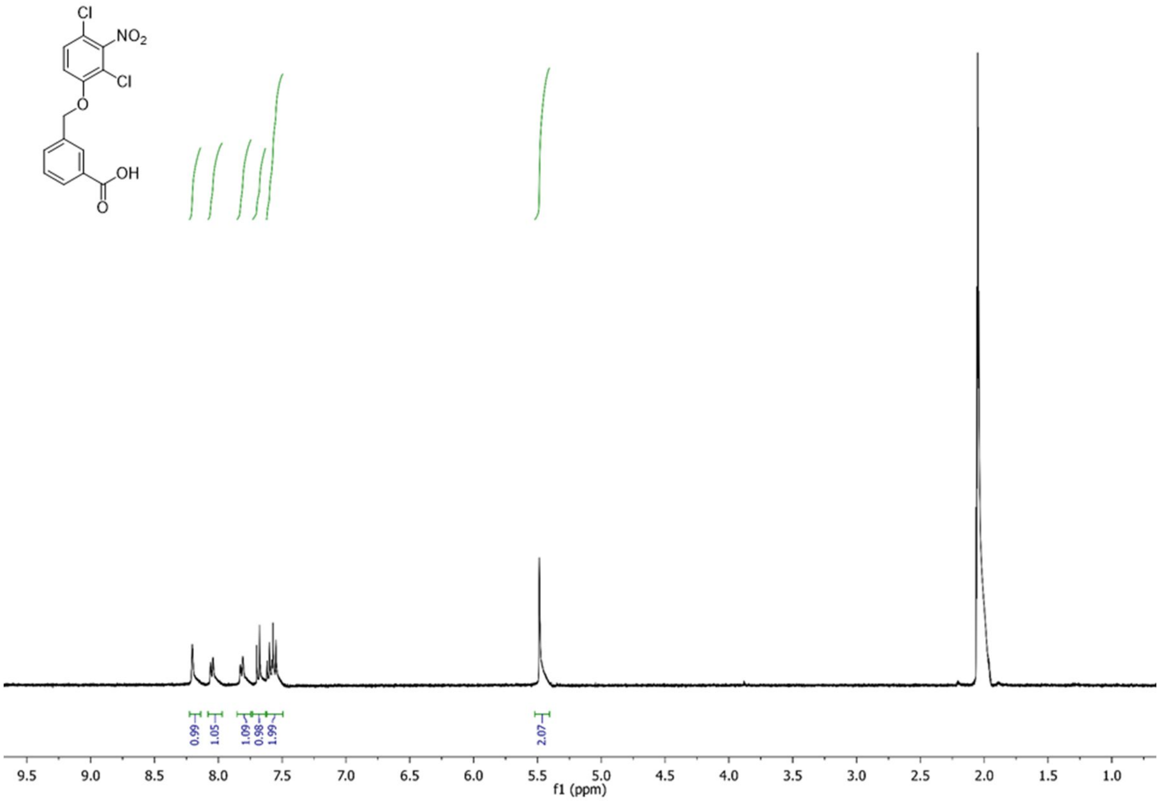
Cl
NO2
Cl
O
OH
O
0.99
1.05
1.09
0.98
1.99
2.07
9.5
9.0
8.5
8.0
7.5
7.0
6.5
6.0
5.5
5.0
4.5
4.0
3.5
3.0
2.5
2.0
1.5
1.0
f1 (ppm)

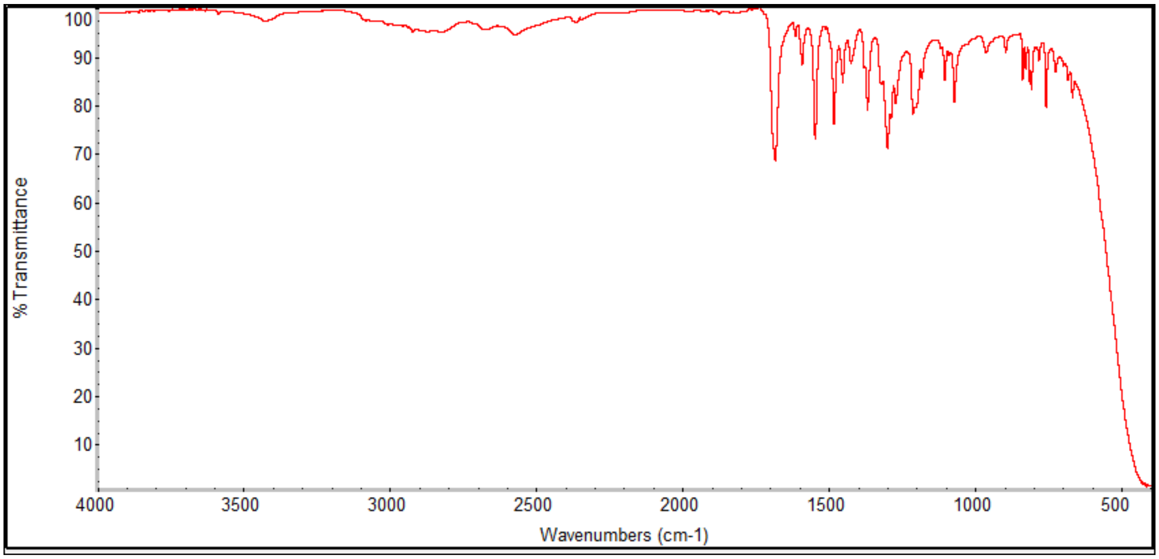
% Transmittance
100
90
80
70
60
50
40
30
20
10
4000
3500
3000
2500
2000
1500
1000
500
Wavenumbers (cm-1)

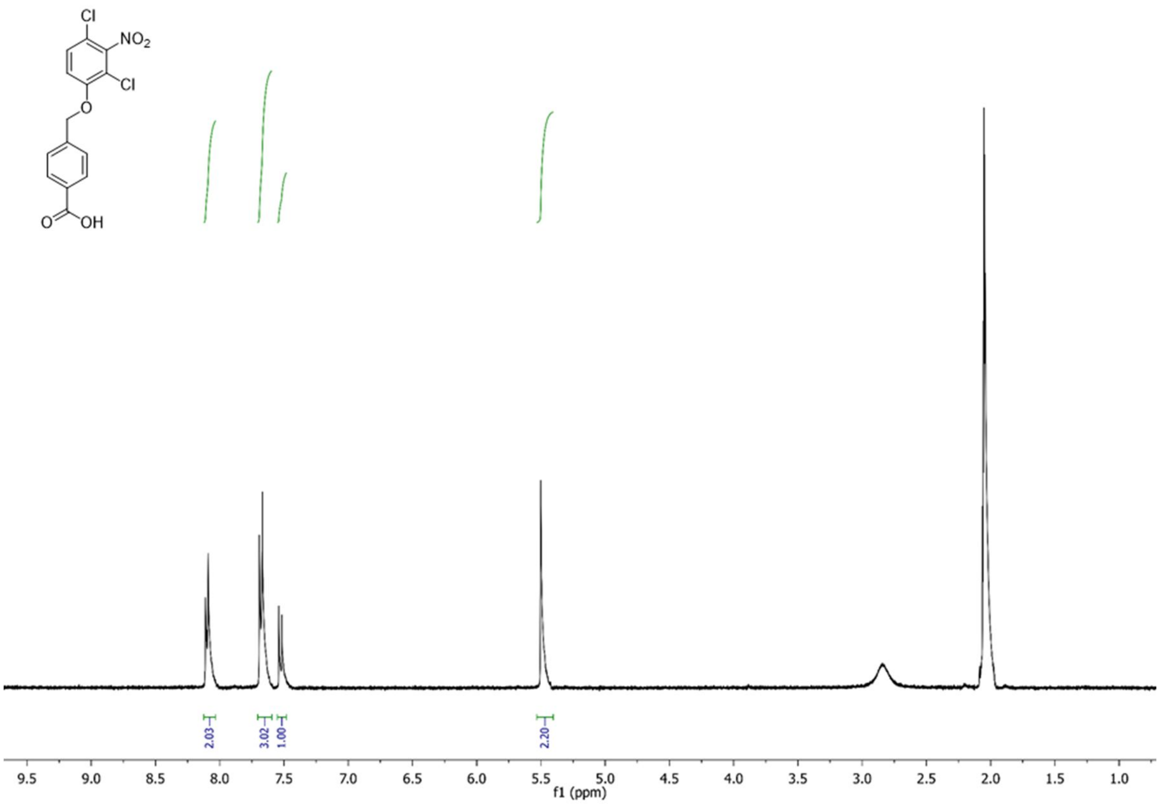

Cl
NO2
Cl
O
O
OH
2.03
3.02
1.00
2.20
9.5
9.0
8.5
8.0
7.5
7.0
6.5
6.0
5.5
5.0
4.5
4.0
3.5
3.0
2.5
2.0
1.5
1.0
f1 (ppm)

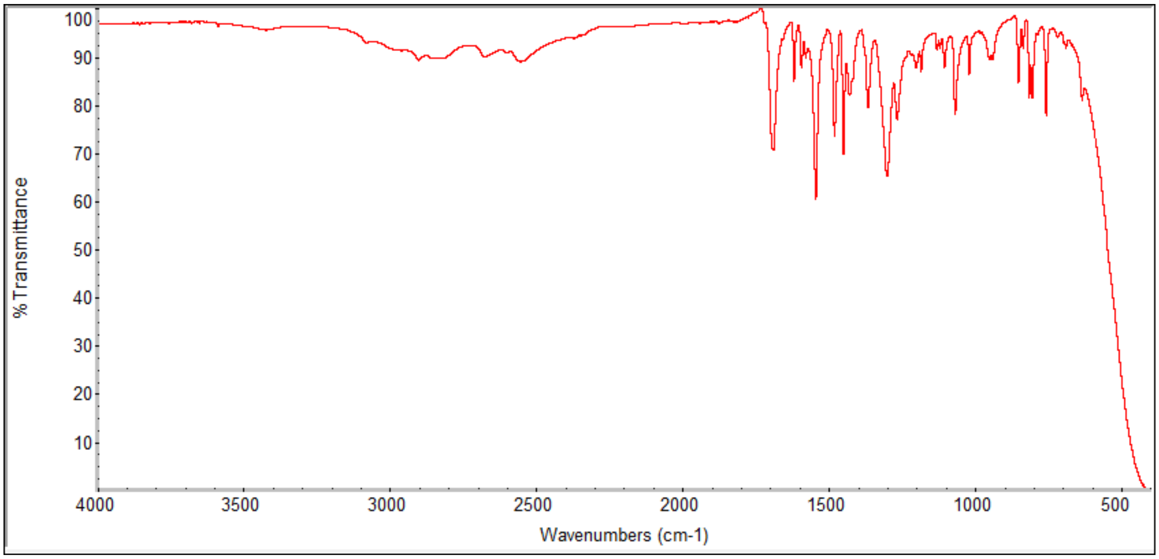

%Transmittance
100
90
80
70
60
50
40
30
20
10
4000
3500
3000
2500
2000
1500
1000
500
Wavenumbers (cm-1)

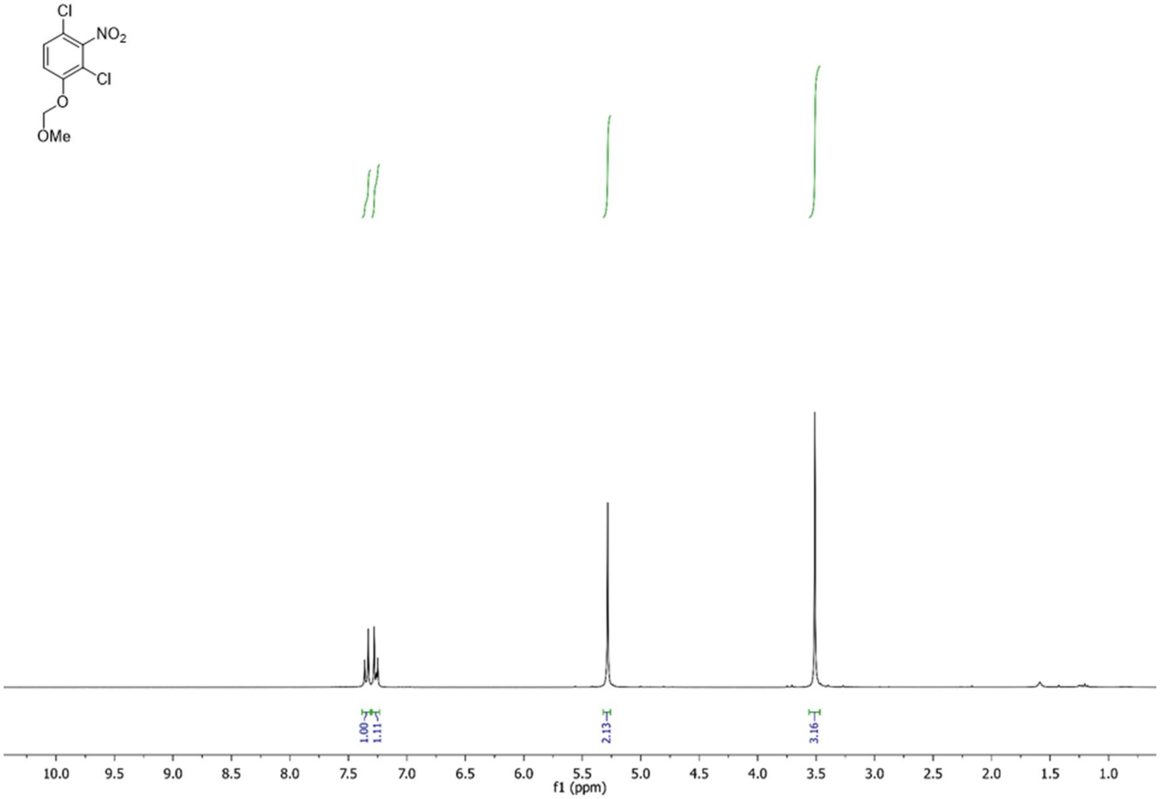
Cl
NO2
Cl
O
OMe
1.00
1.11
2.13
3.16
10.0 9.5 9.0 8.5 8.0 7.5 7.0 6.5 6.0 5.5 5.0 4.5 4.0 3.5 3.0 2.5 2.0 1.5 1.0
f1 (ppm)

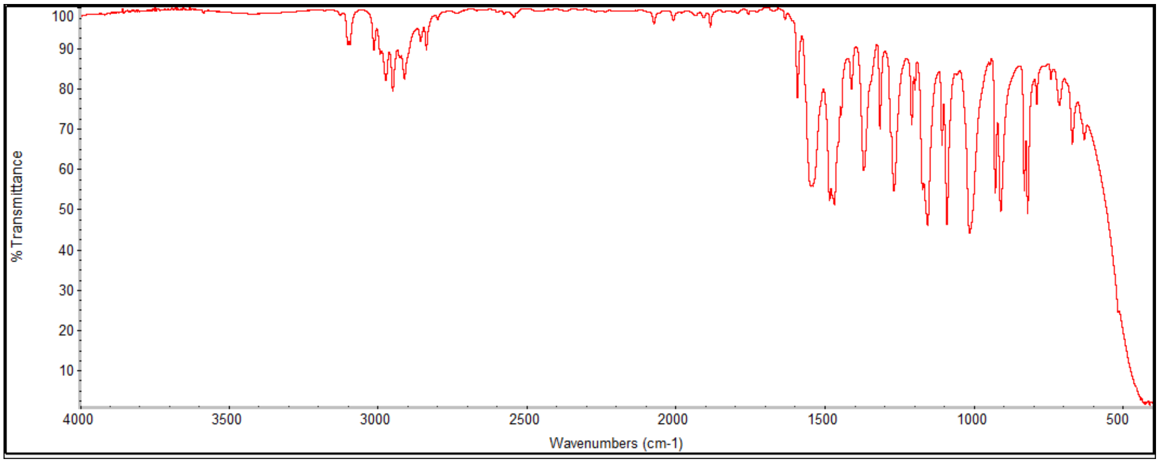
% Transmittance
100 90 80 70 60 50 40 30 20 10
4000 3500 3000 2500 2000 1500 1000 500
Wavenumbers (cm-1)

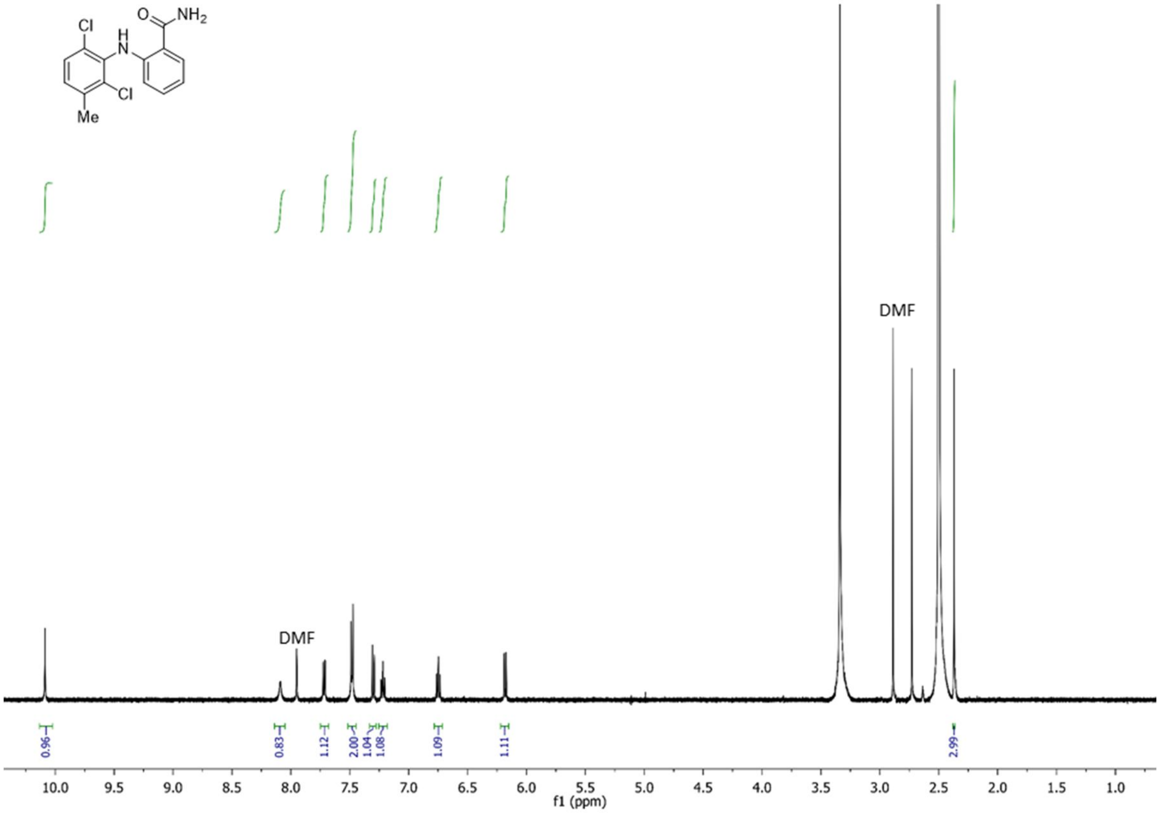

Cl
H
N
O
NH2
Cl
Me
DMF
DMF
0.96
0.83
1.12
2.00
1.04
1.08
1.09
1.11
2.99
10.0
9.5
9.0
8.5
8.0
7.5
7.0
6.5
6.0
5.5
5.0
4.5
4.0
3.5
3.0
2.5
2.0
1.5
1.0
f1 (ppm)

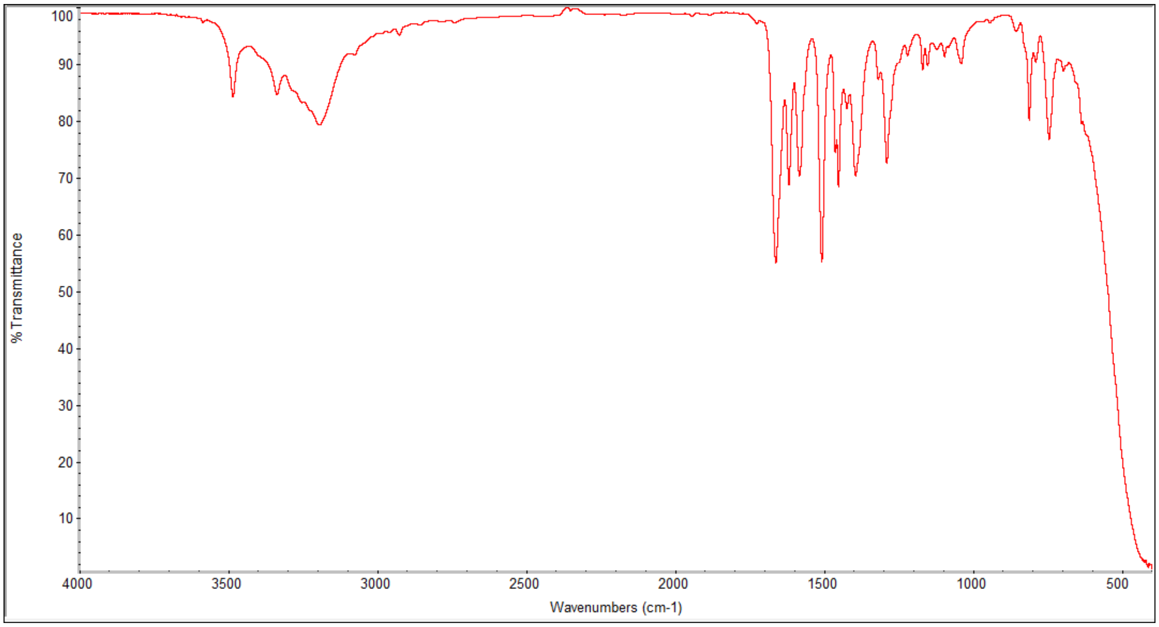

% Transmittance
Wavenumbers (cm-1)
100
90
80
70
60
50
40
30
20
10
4000
3500
3000
2500
2000
1500
1000
500

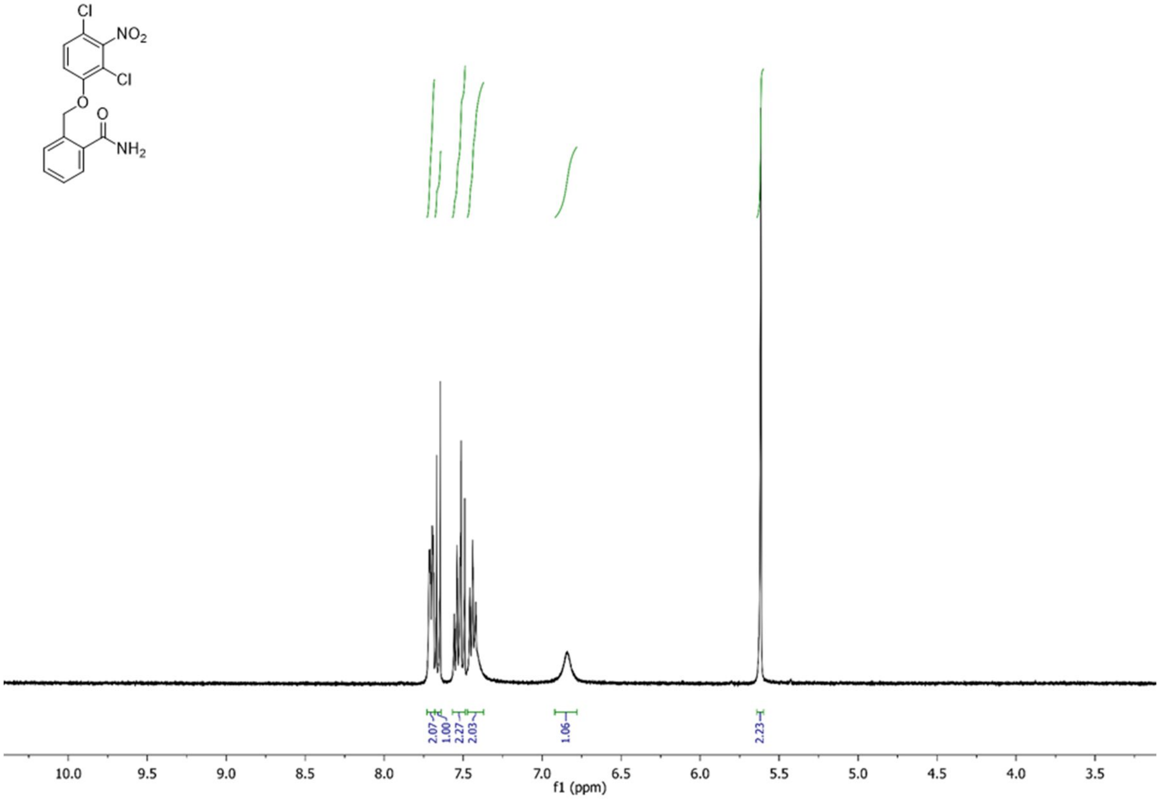
Cl
NO2
Cl
O
O
NH2
2.07
1.00
2.27
2.03
1.06
2.23
10.0
9.5
9.0
8.5
8.0
7.5
7.0
6.5
6.0
5.5
5.0
4.5
4.0
3.5
f1 (ppm)

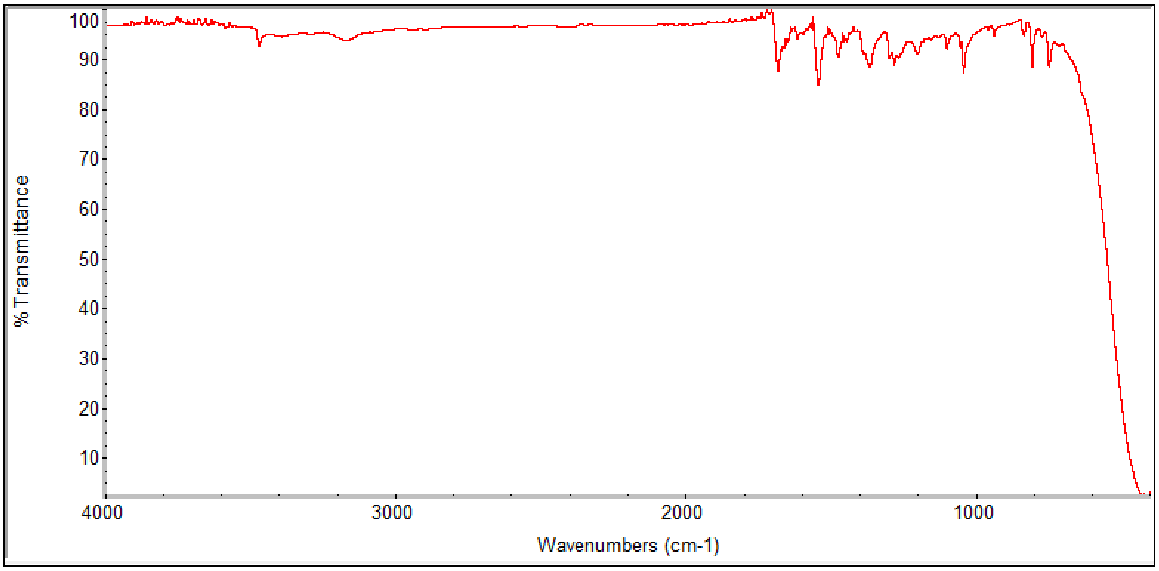
100
90
80
70
60
50
40
30
20
10
% Transmittance
4000
3000
2000
1000
Wavenumbers (cm-1)

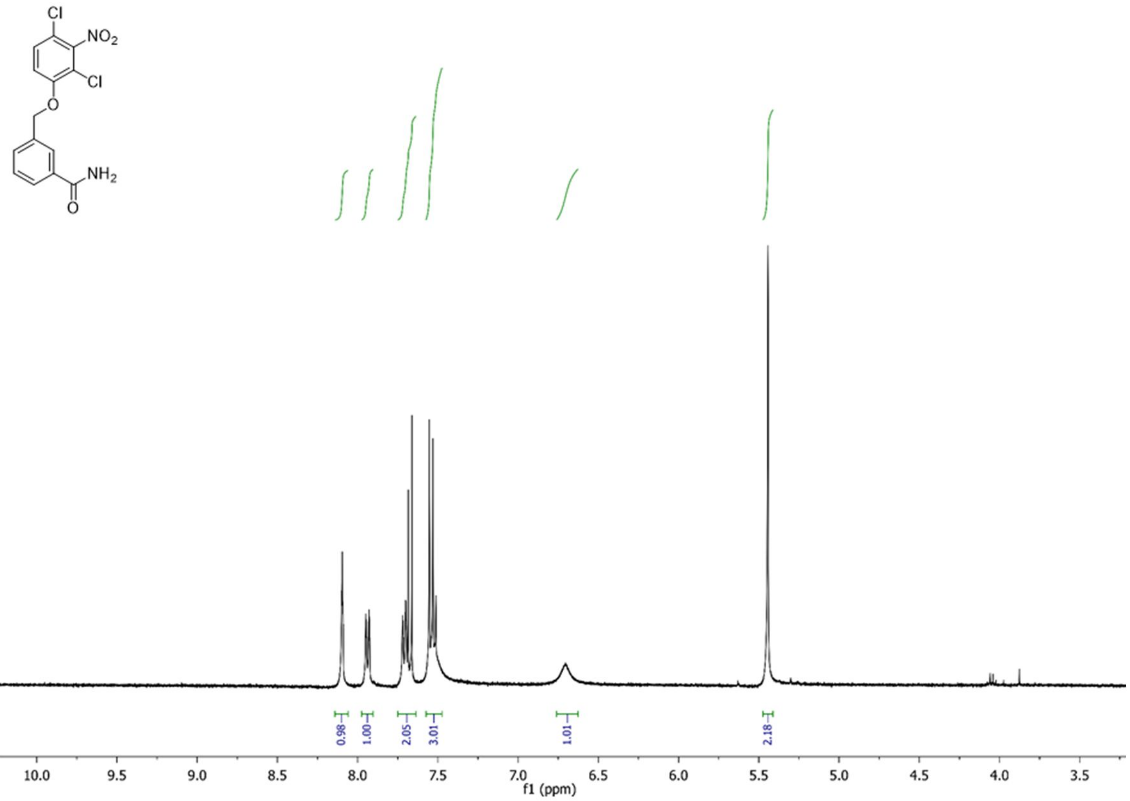
Cl
NO2
Cl
O
NH2
O
0.98
1.00
2.05
3.01
1.01
2.18
10.0 9.5 9.0 8.5 8.0 7.5 7.0 6.5 6.0 5.5 5.0 4.5 4.0 3.5
f1 (ppm)

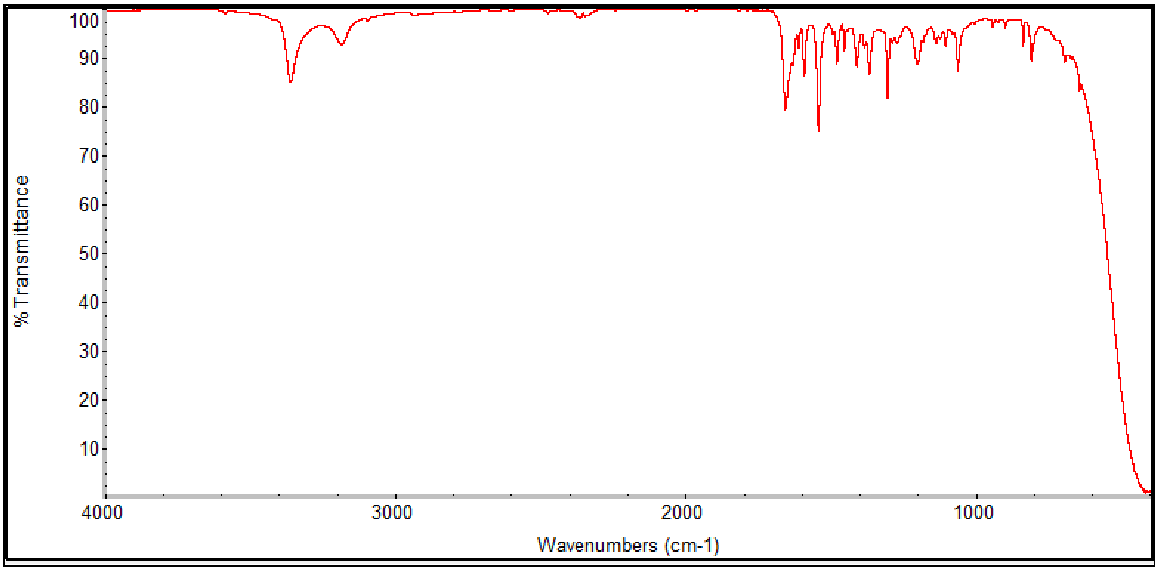
% Transmittance
100 90 80 70 60 50 40 30 20 10
4000 3000 2000 1000
Wavenumbers (cm-1)

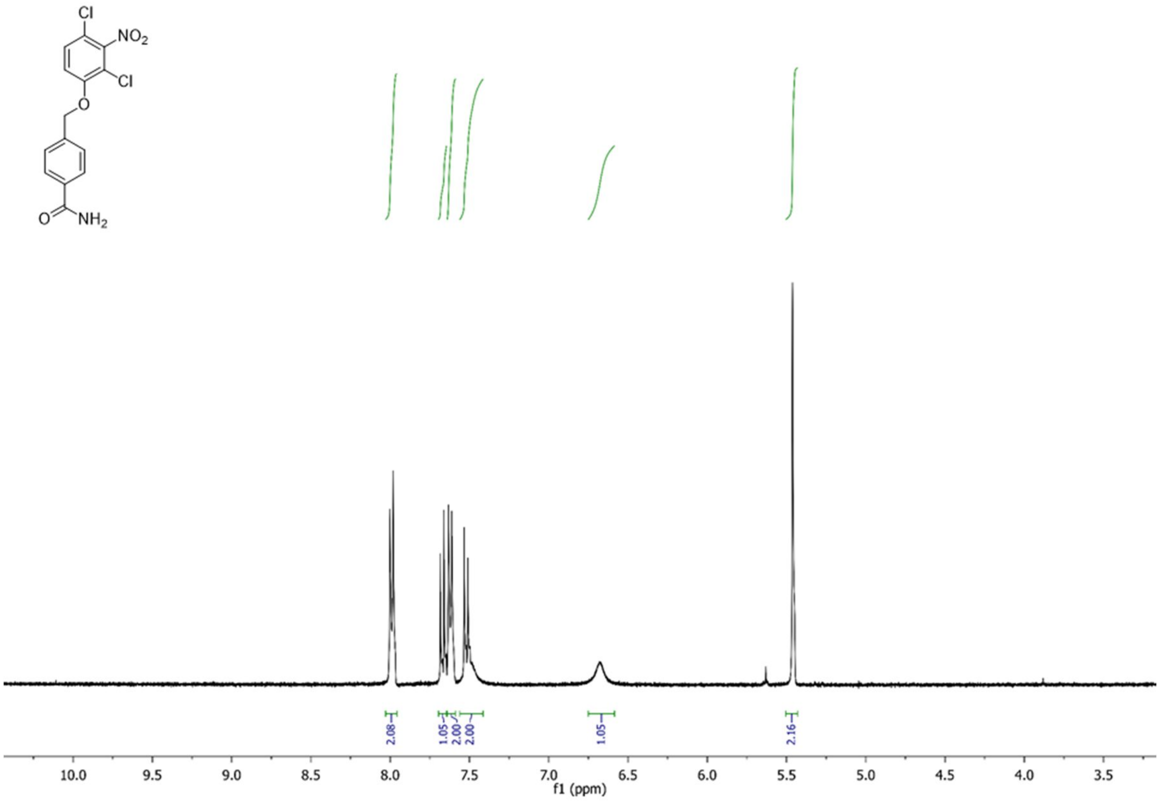
Cl
NO2
Cl
O
O
NH2
2.08
1.05
2.00
2.00
1.05
2.16
10.0
9.5
9.0
8.5
8.0
7.5
7.0
6.5
6.0
5.5
5.0
4.5
4.0
3.5
f1 (ppm)

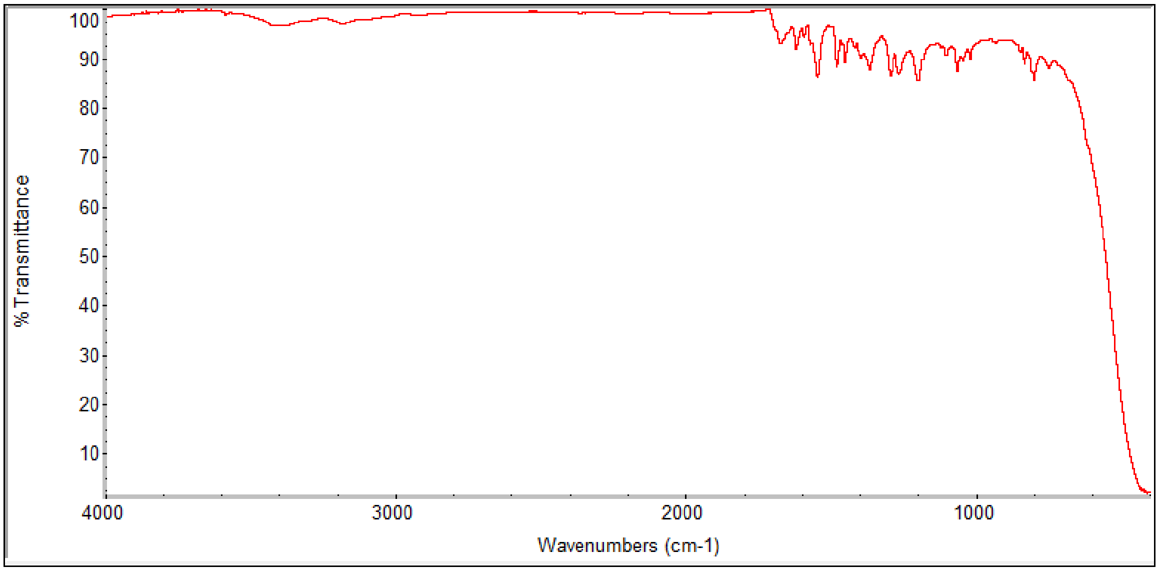
% Transmittance
100
90
80
70
60
50
40
30
20
10
4000
3000
2000
1000
Wavenumbers (cm-1)

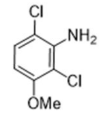
Cl
NH$_2$
Cl
OMe

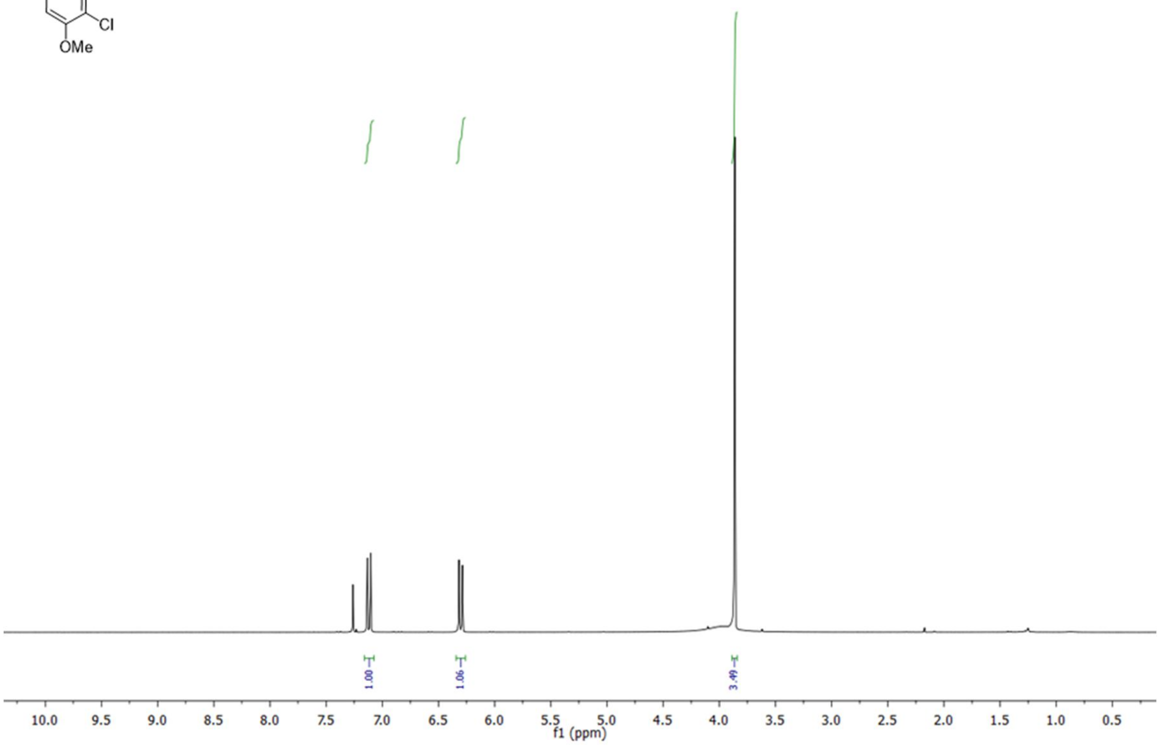
1.00
1.06
3.49
10.0
9.5
9.0
8.5
8.0
7.5
7.0
6.5
6.0
5.5
5.0
4.5
4.0
3.5
3.0
2.5
2.0
1.5
1.0
0.5
f1 (ppm)

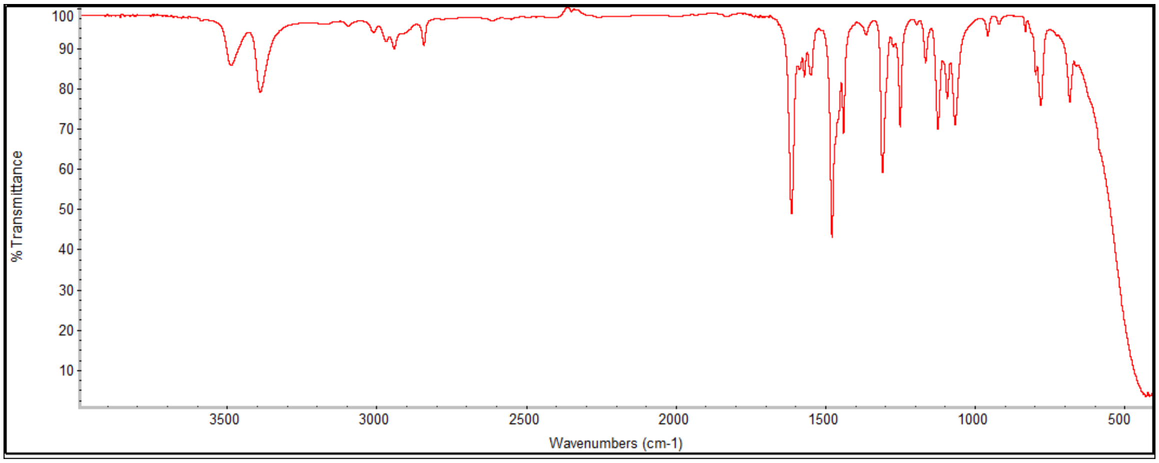
% Transmittance
100
90
80
70
60
50
40
30
20
10
3500
3000
2500
2000
1500
1000
500
Wavenumbers (cm-1)

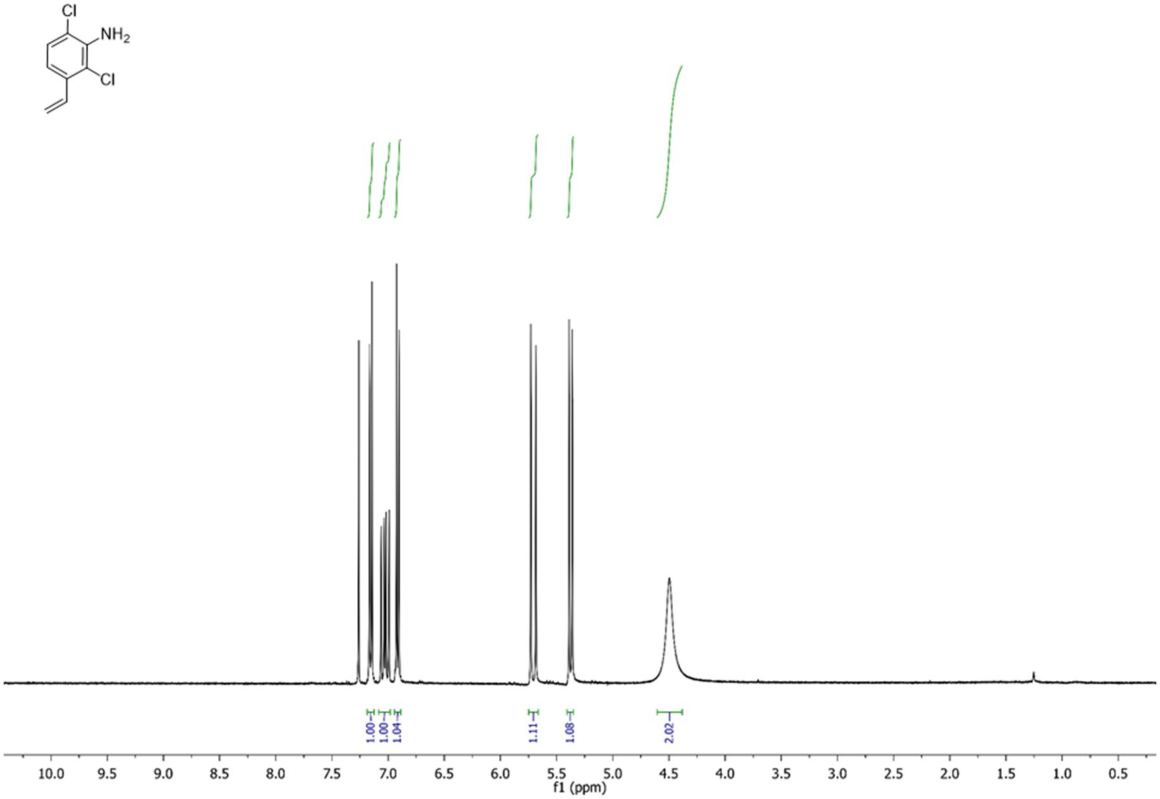
Cl
NH2
Cl
1.00
1.00
1.04
1.11
1.08
2.02
10.0 9.5 9.0 8.5 8.0 7.5 7.0 6.5 6.0 5.5 5.0 4.5 4.0 3.5 3.0 2.5 2.0 1.5 1.0 0.5
f1 (ppm)

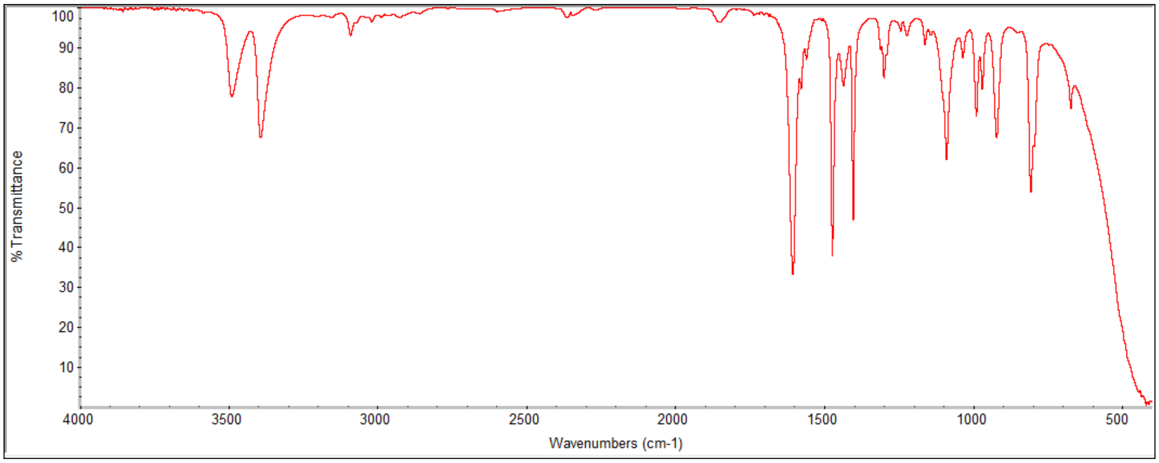
% Transmittance
100
90
80
70
60
50
40
30
20
10
4000 3500 3000 2500 2000 1500 1000 500
Wavenumbers (cm-1)

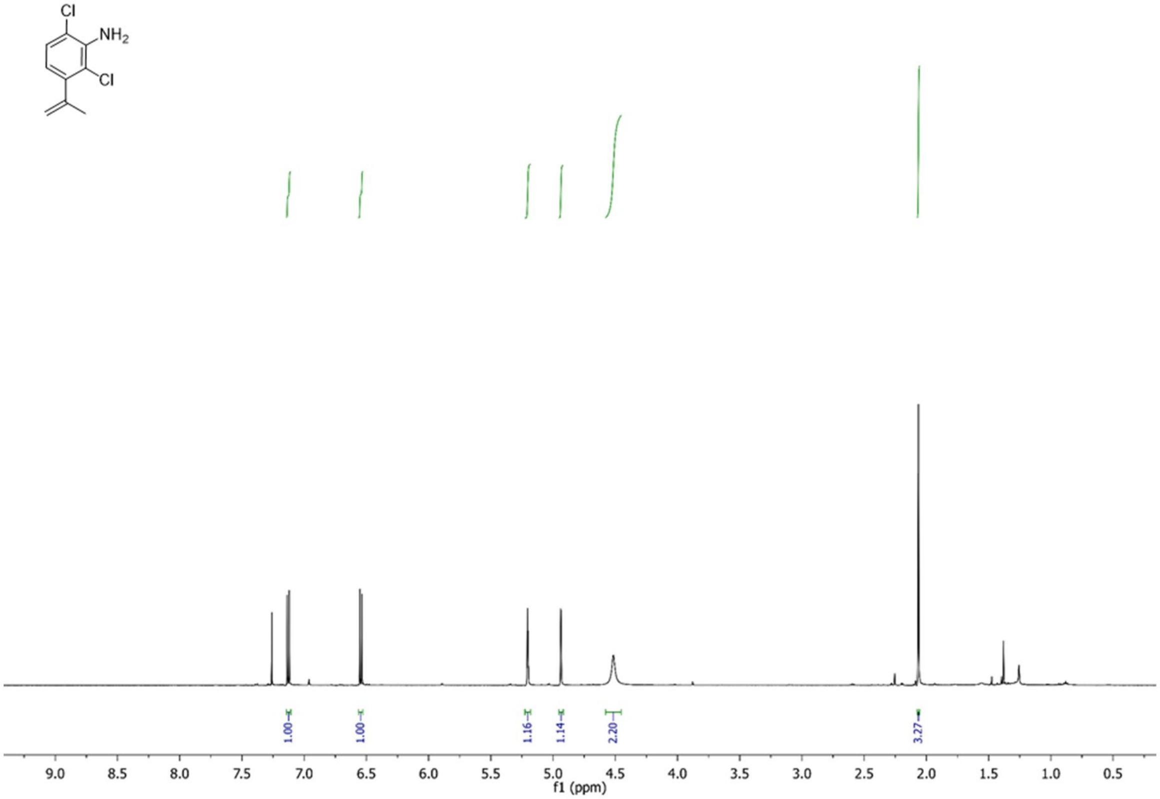
Cl
NH2
Cl
1.00
1.00
1.16
1.14
2.20
3.27
9.0 8.5 8.0 7.5 7.0 6.5 6.0 5.5 5.0 4.5 4.0 3.5 3.0 2.5 2.0 1.5 1.0 0.5
f1 (ppm)

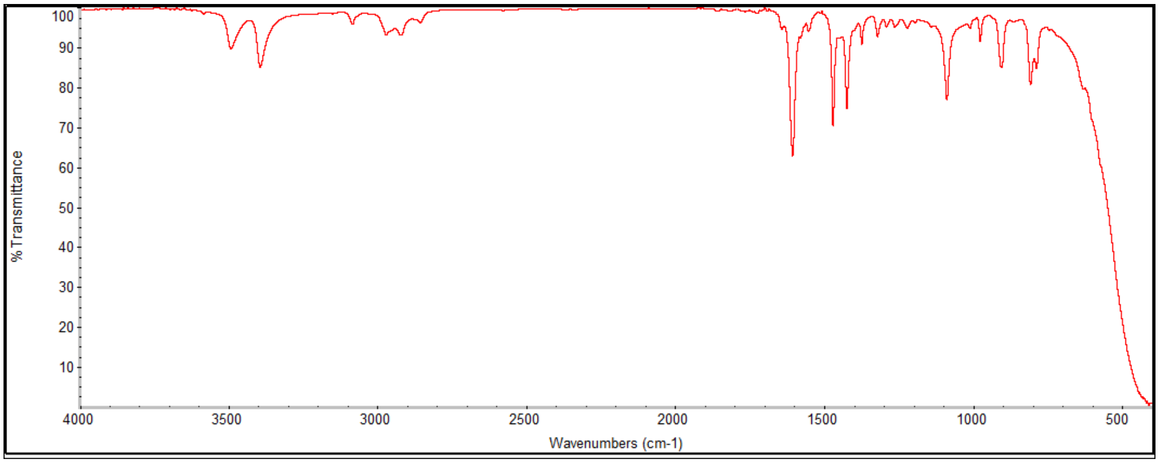
% Transmittance
100 90 80 70 60 50 40 30 20 10
4000 3500 3000 2500 2000 1500 1000 500
Wavenumbers (cm-1)

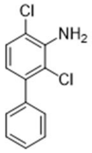
Cl
NH2
Cl

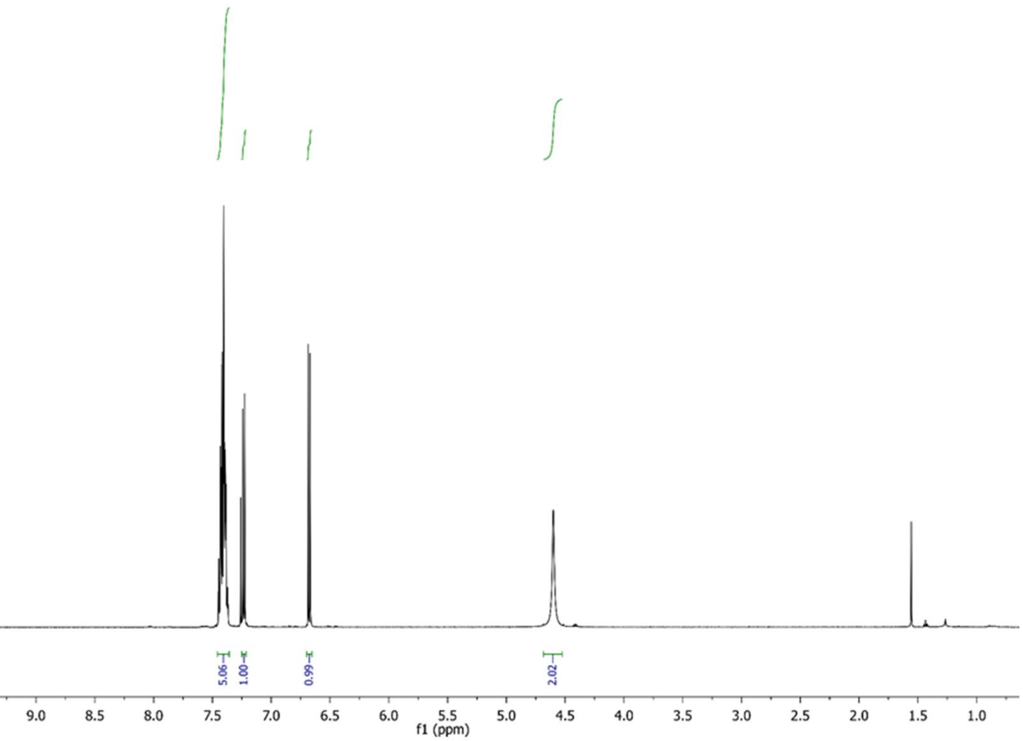
5.06
1.00
0.99
2.02
10.0 9.5 9.0 8.5 8.0 7.5 7.0 6.5 6.0 5.5 5.0 4.5 4.0 3.5 3.0 2.5 2.0 1.5 1.0
f1 (ppm)

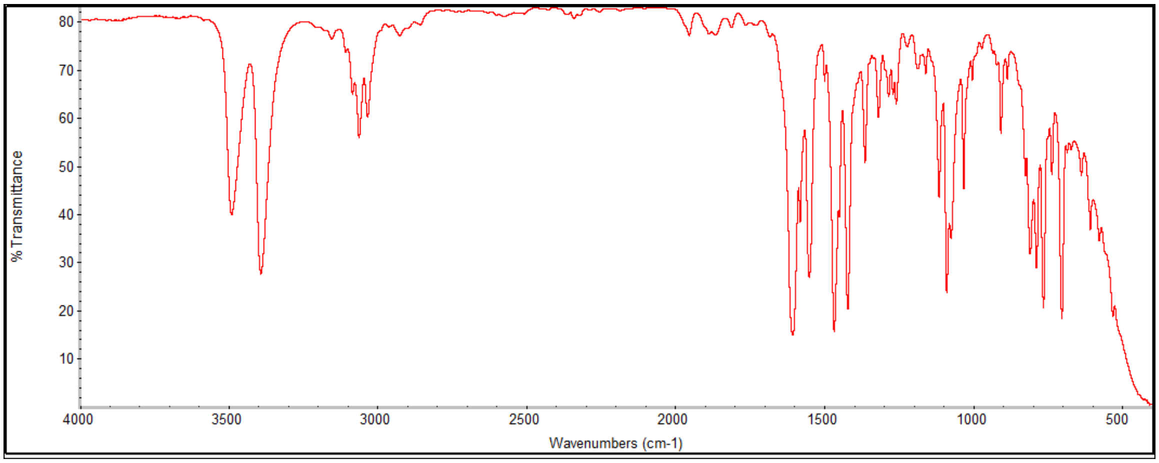
% Transmittance
80 70 60 50 40 30 20 10
4000 3500 3000 2500 2000 1500 1000 500
Wavenumbers (cm-1)

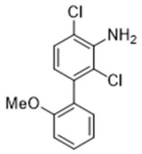

Cl
NH2
Cl
MeO

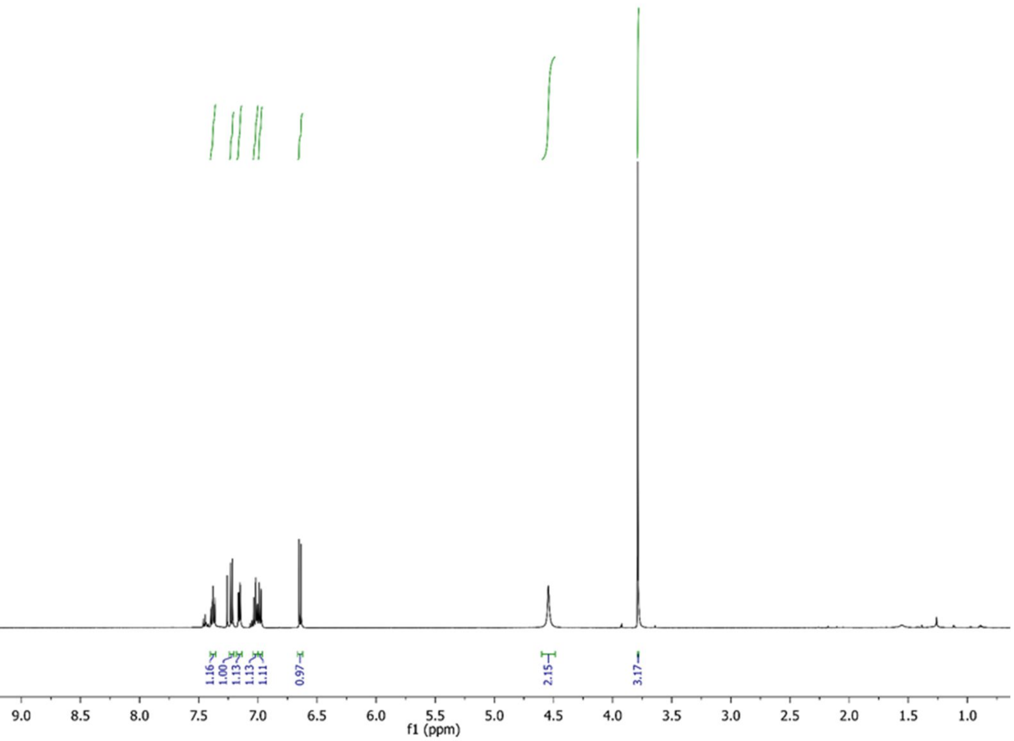

1.16
1.00
1.13
1.13
1.11
0.97
2.15
3.17
10.0
9.5
9.0
8.5
8.0
7.5
7.0
6.5
6.0
5.5
5.0
4.5
4.0
3.5
3.0
2.5
2.0
1.5
1.0
f1 (ppm)

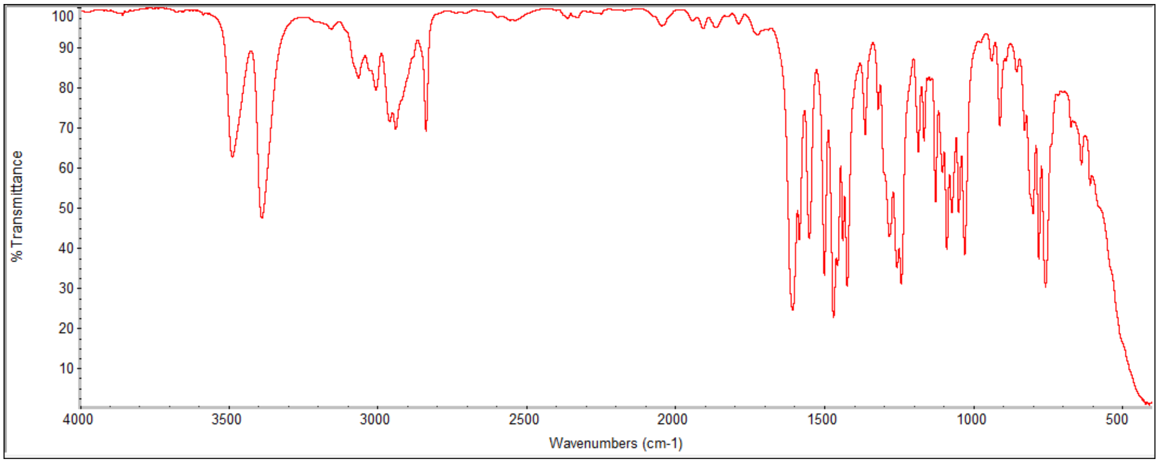

% Transmittance
100
90
80
70
60
50
40
30
20
10
4000
3500
3000
2500
2000
1500
1000
500
Wavenumbers (cm-1)

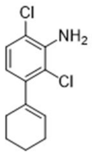
Cl
NH2
Cl

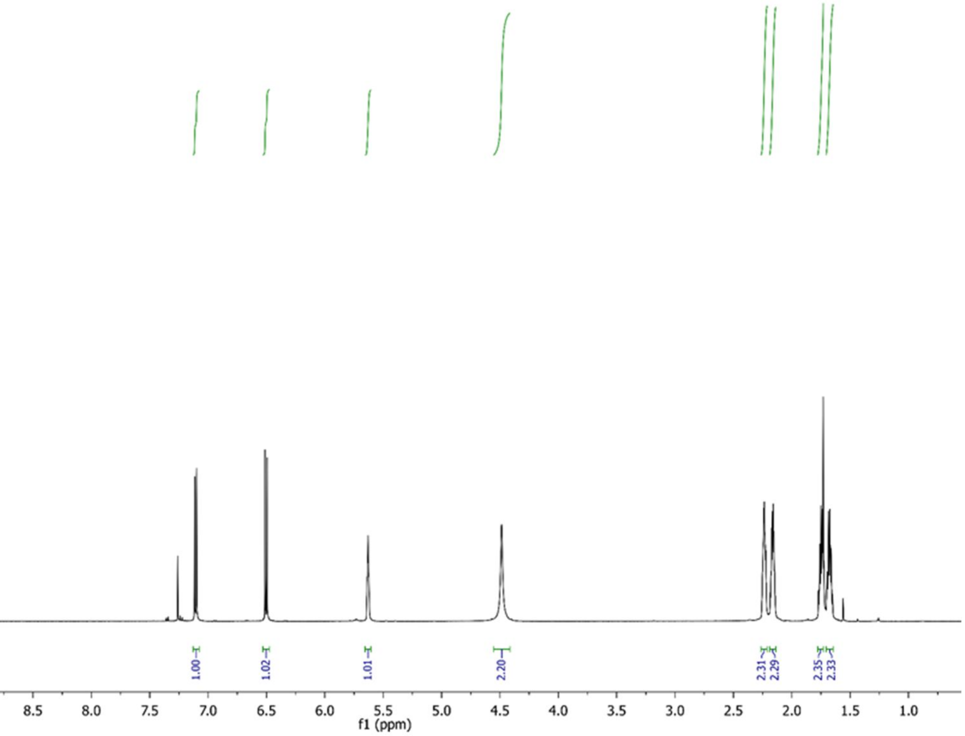
1.00
1.02
1.01
2.20
2.31
2.29
2.35
2.33
10.0
9.5
9.0
8.5
8.0
7.5
7.0
6.5
6.0
5.5
5.0
4.5
4.0
3.5
3.0
2.5
2.0
1.5
1.0
f1 (ppm)

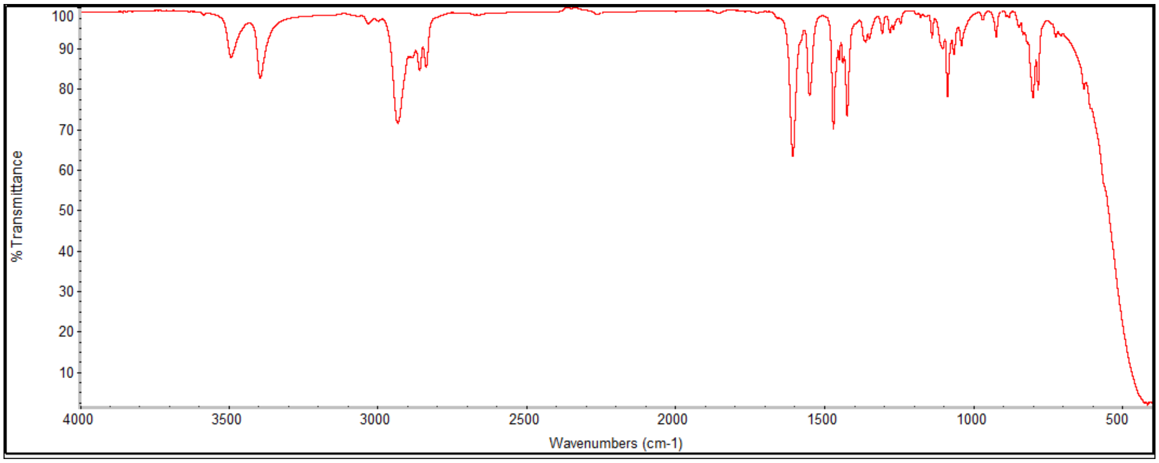
% Transmittance
100
90
80
70
60
50
40
30
20
10
4000
3500
3000
2500
2000
1500
1000
500
Wavenumbers (cm-1)

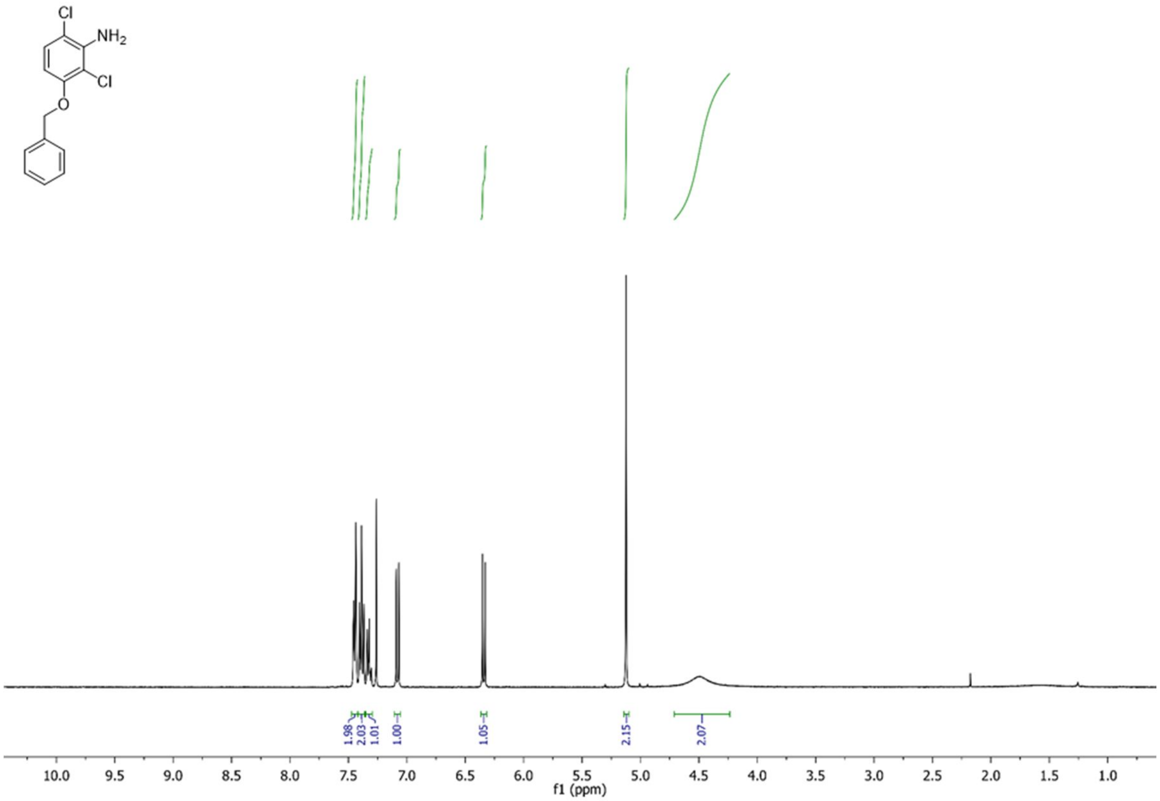
Cl
NH2
Cl
O
1.98
2.03
1.01
1.00
1.05
2.15
2.07
10.0 9.5 9.0 8.5 8.0 7.5 7.0 6.5 6.0 5.5 5.0 4.5 4.0 3.5 3.0 2.5 2.0 1.5 1.0
f1 (ppm)

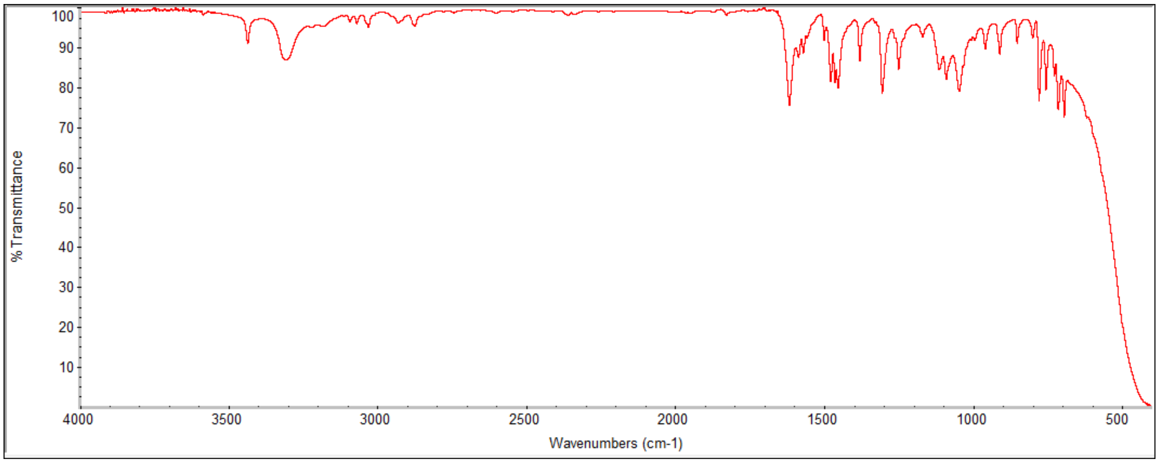
% Transmittance
100 90 80 70 60 50 40 30 20 10
4000 3500 3000 2500 2000 1500 1000 500
Wavenumbers (cm-1)

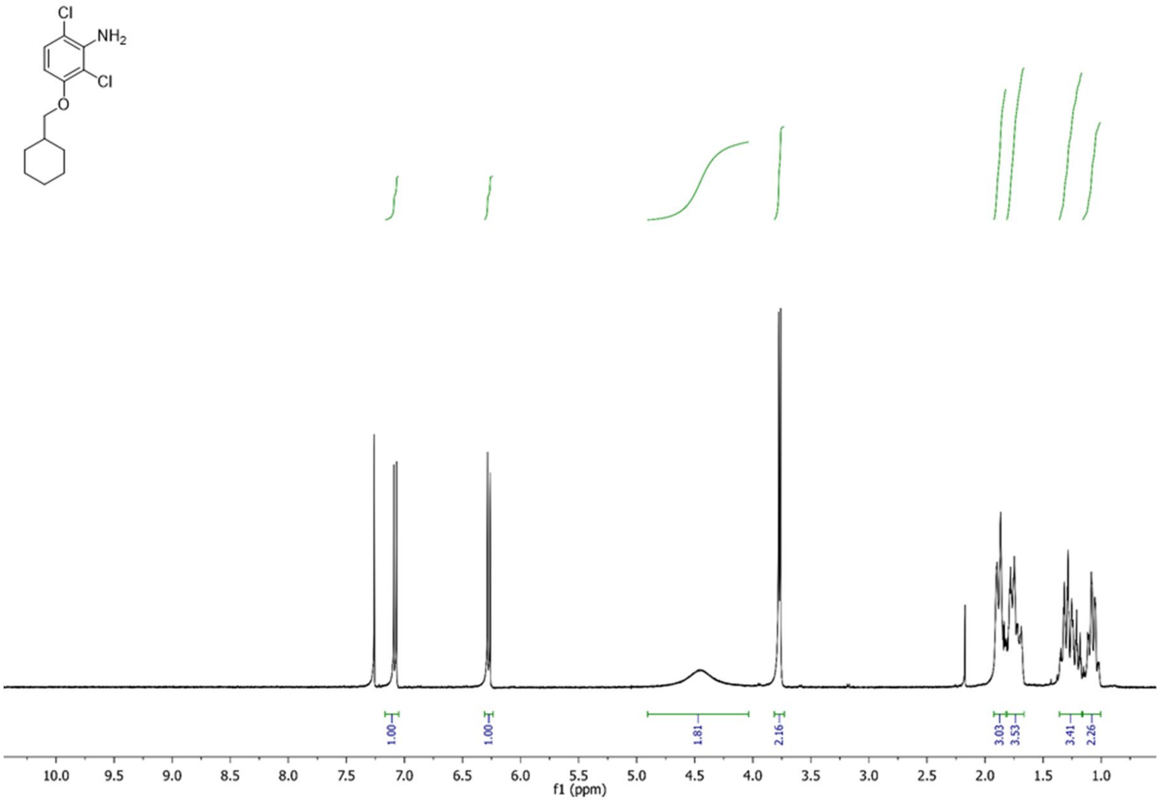
Cl
NH2
Cl
O
1.00
1.00
1.81
2.16
3.03
3.53
3.41
2.26
10.0 9.5 9.0 8.5 8.0 7.5 7.0 6.5 6.0 5.5 5.0 4.5 4.0 3.5 3.0 2.5 2.0 1.5 1.0
f1 (ppm)

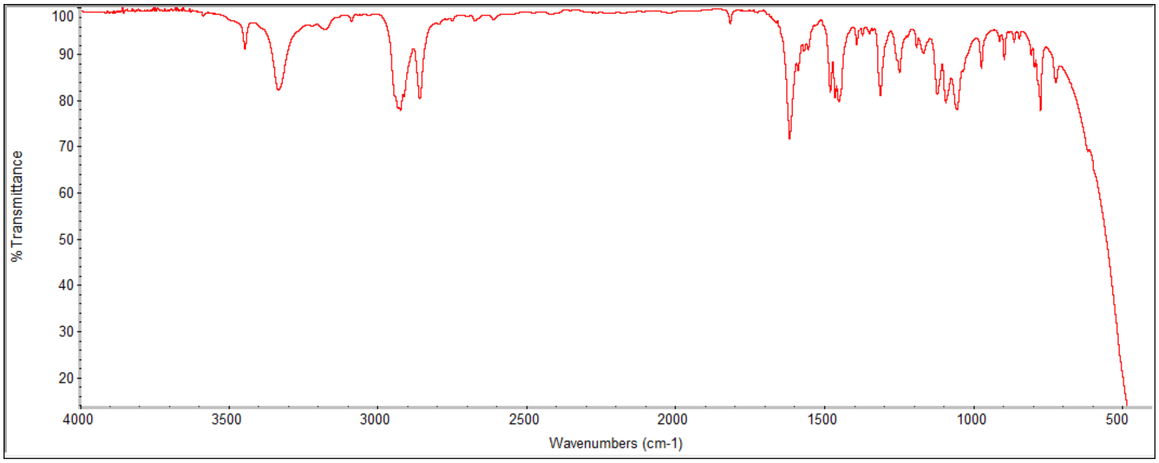
% Transmittance
100
90
80
70
60
50
40
30
20
4000 3500 3000 2500 2000 1500 1000 500
Wavenumbers (cm-1)

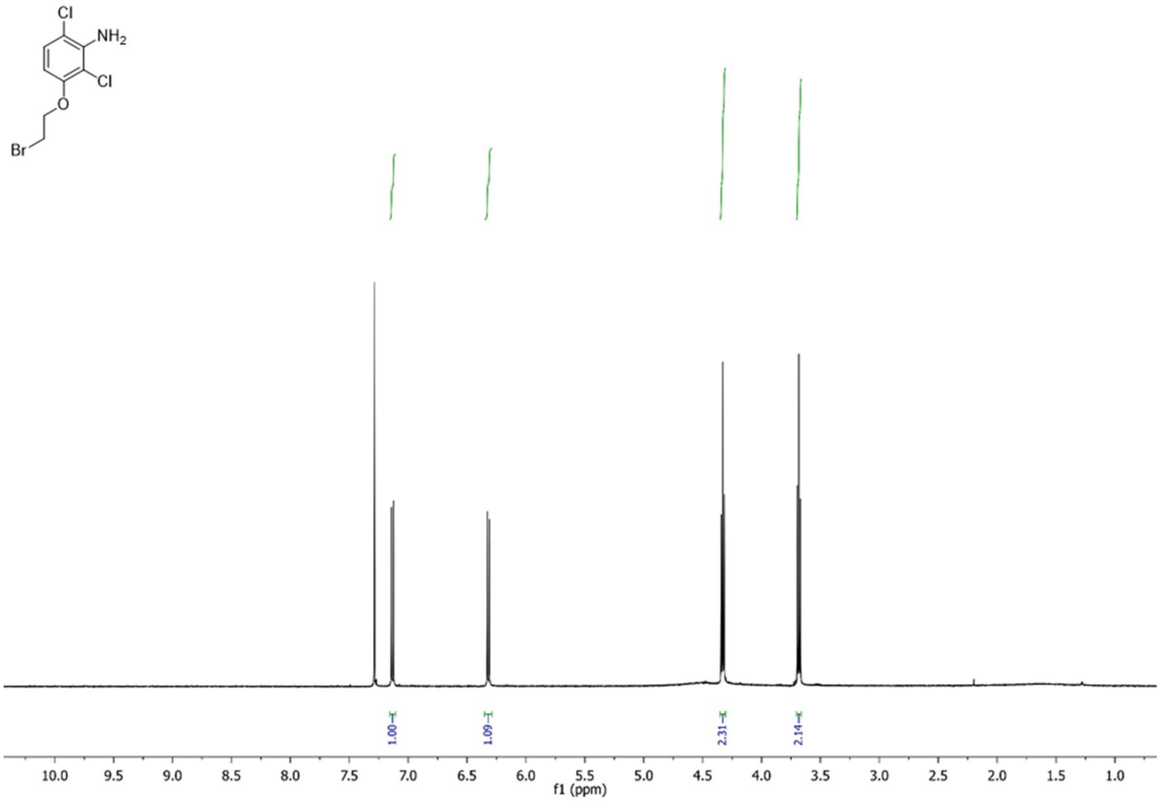
Cl
NH2
Cl
O
Br
1.00
1.09
2.31
2.14
10.0 9.5 9.0 8.5 8.0 7.5 7.0 6.5 6.0 5.5 5.0 4.5 4.0 3.5 3.0 2.5 2.0 1.5 1.0
f1 (ppm)

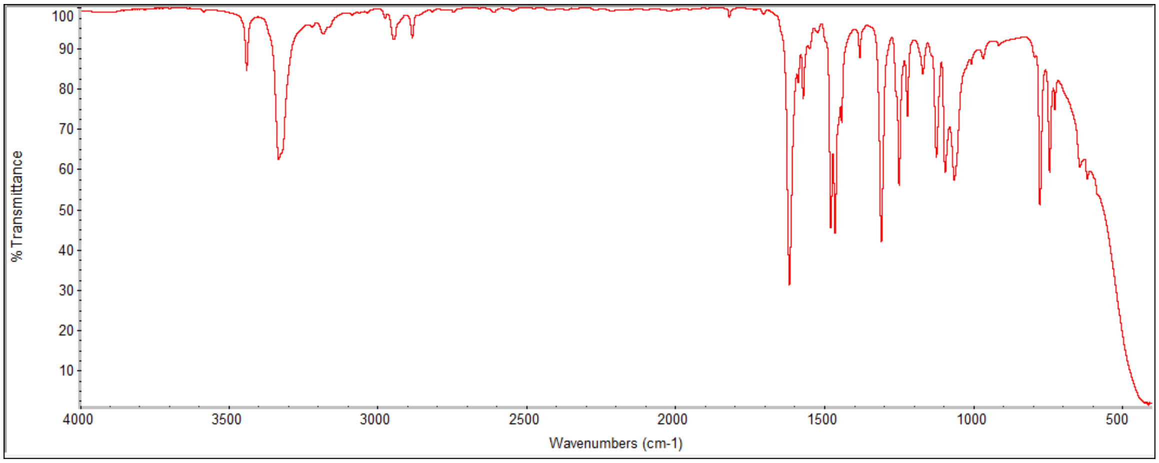
% Transmittance
100 90 80 70 60 50 40 30 20 10
4000 3500 3000 2500 2000 1500 1000 500
Wavenumbers (cm-1)

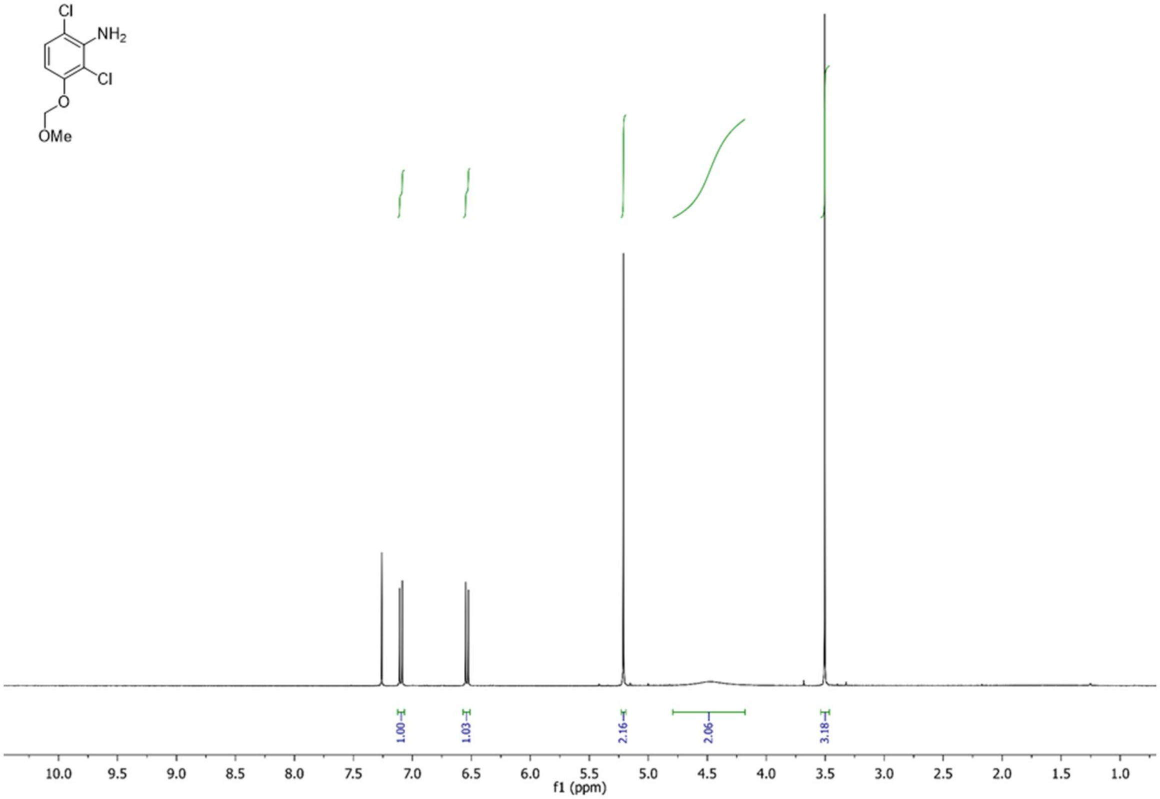
Cl
NH2
Cl
O
OMe
1.00
1.03
2.16
2.06
3.18
10.0 9.5 9.0 8.5 8.0 7.5 7.0 6.5 6.0 5.5 5.0 4.5 4.0 3.5 3.0 2.5 2.0 1.5 1.0
f1 (ppm)

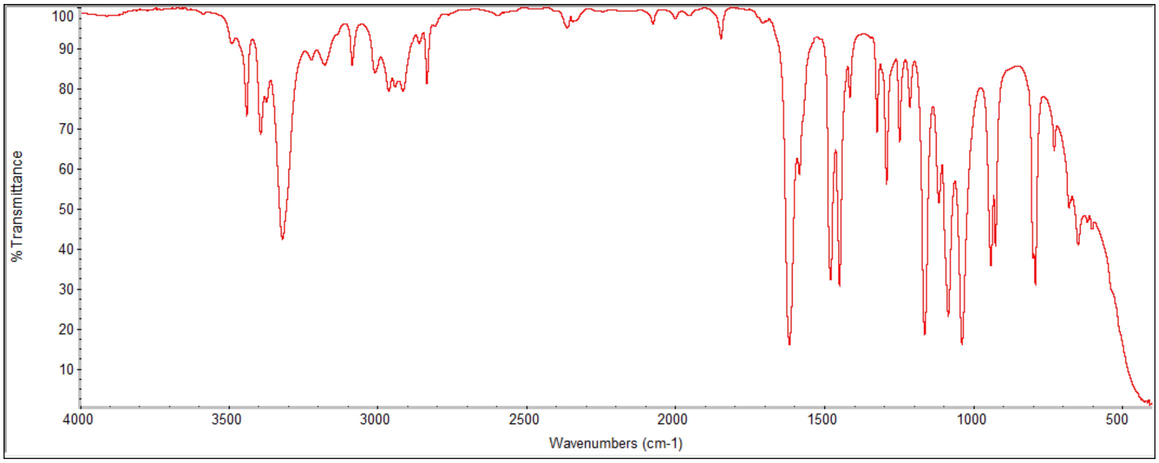
% Transmittance
100 90 80 70 60 50 40 30 20 10
4000 3500 3000 2500 2000 1500 1000 500
Wavenumbers (cm-1)

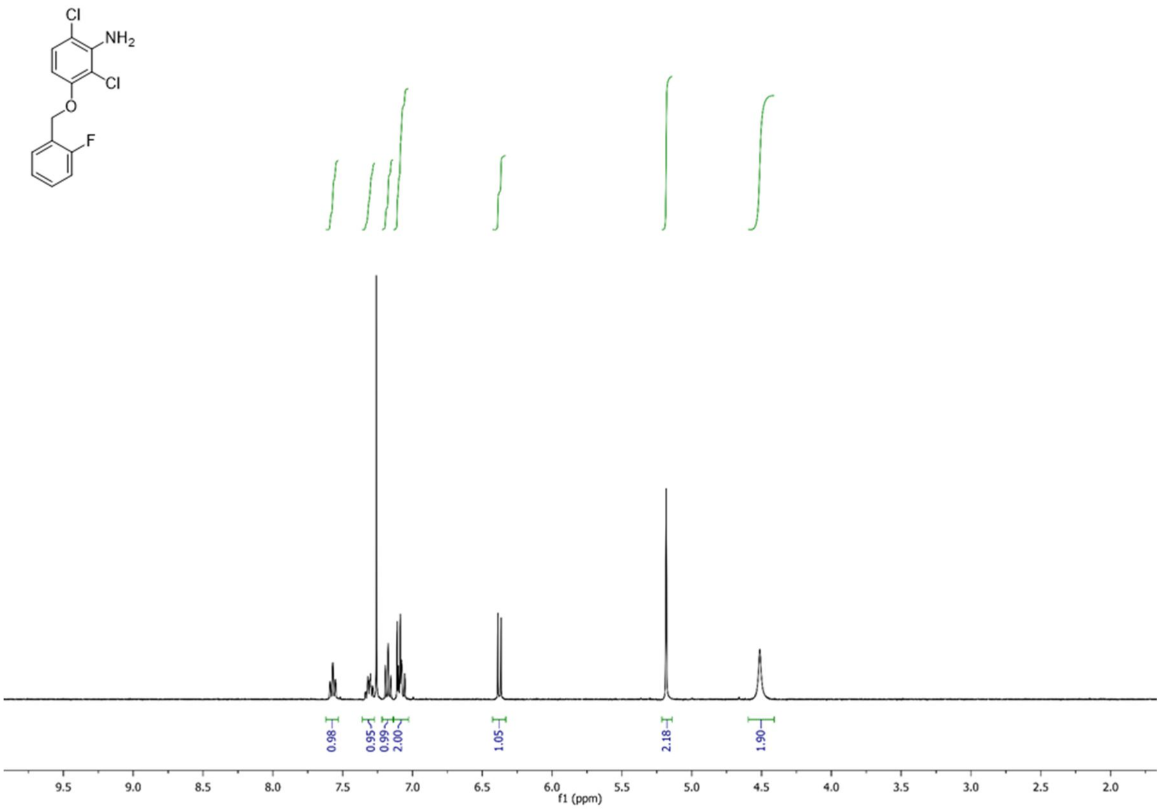
Cl
NH2
Cl
O
F
0.98
0.95
0.99
2.00
1.05
2.18
1.90
9.5
9.0
8.5
8.0
7.5
7.0
6.5
6.0
5.5
5.0
4.5
4.0
3.5
3.0
2.5
2.0
f1 (ppm)

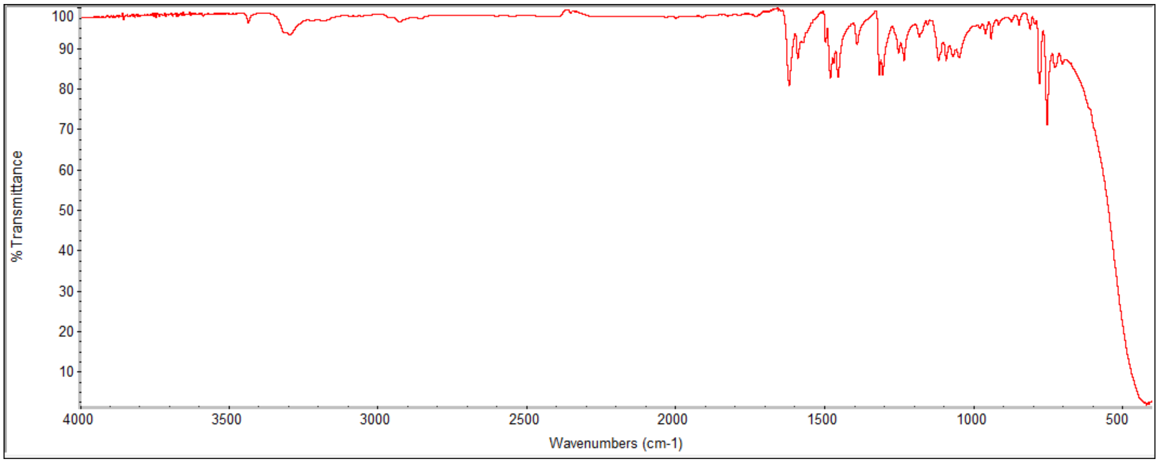
% Transmittance
100
90
80
70
60
50
40
30
20
10
4000
3500
3000
2500
2000
1500
1000
500
Wavenumbers (cm-1)

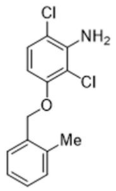
Cl
NH2
Cl
O
Me

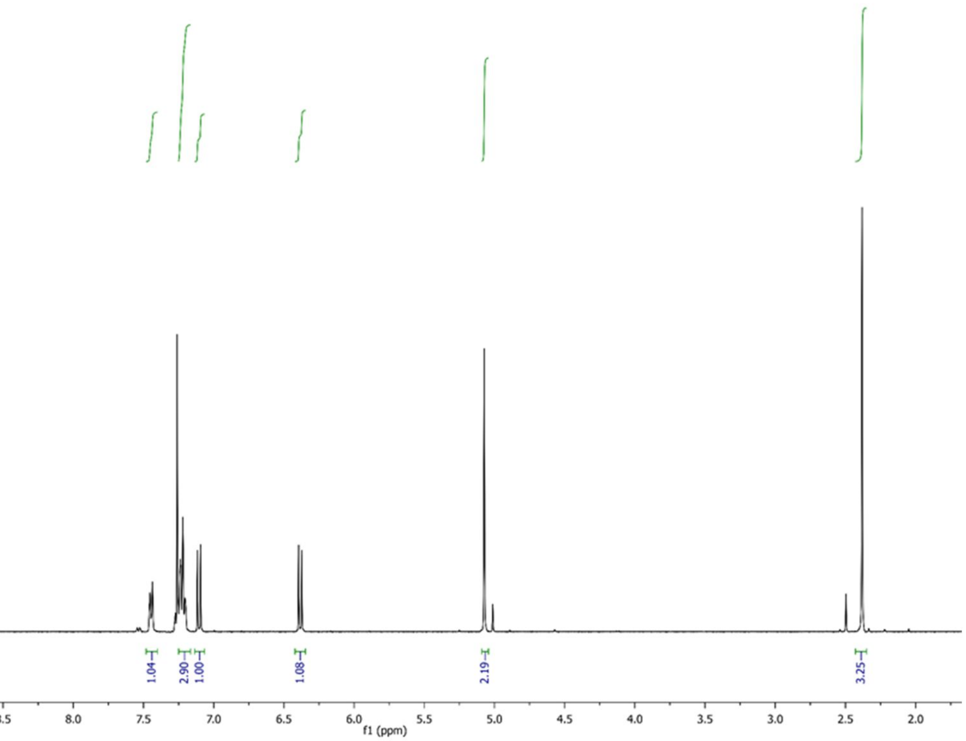
1.04
2.90
1.00
1.08
2.19
3.25
9.5
9.0
8.5
8.0
7.5
7.0
6.5
6.0
5.5
5.0
4.5
4.0
3.5
3.0
2.5
2.0
f1 (ppm)

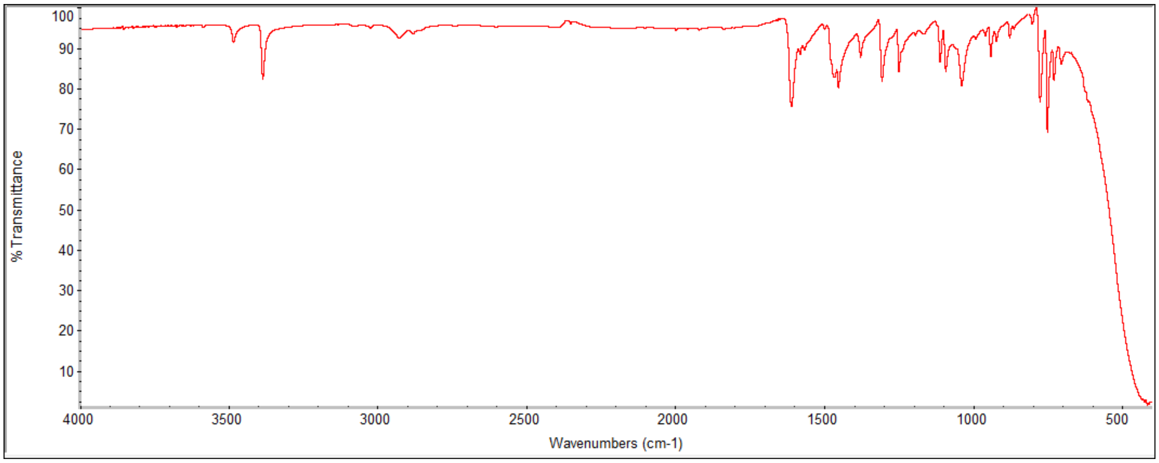
% Transmittance
100
90
80
70
60
50
40
30
20
10
4000
3500
3000
2500
2000
1500
1000
500
Wavenumbers (cm-1)

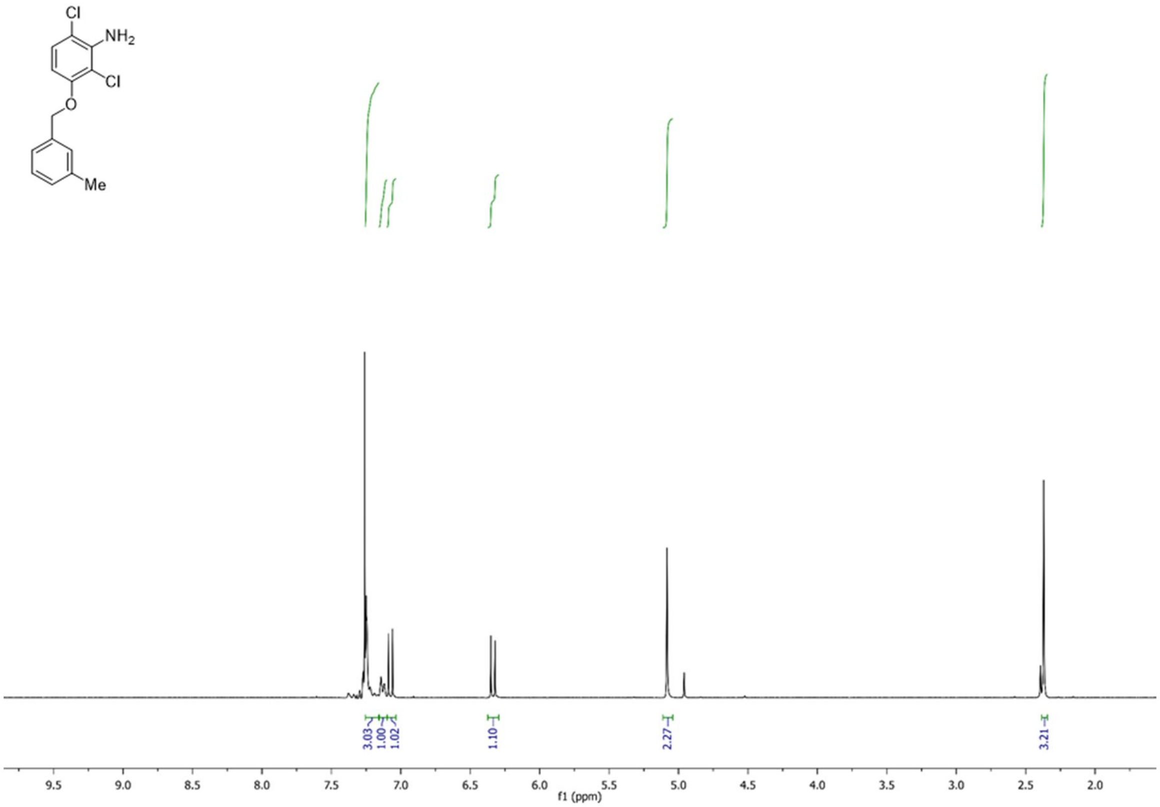
Cl
NH2
Cl
O
Me
3.03
1.00
1.02
1.10
2.27
3.21
9.5 9.0 8.5 8.0 7.5 7.0 6.5 6.0 5.5 5.0 4.5 4.0 3.5 3.0 2.5 2.0
f1 (ppm)

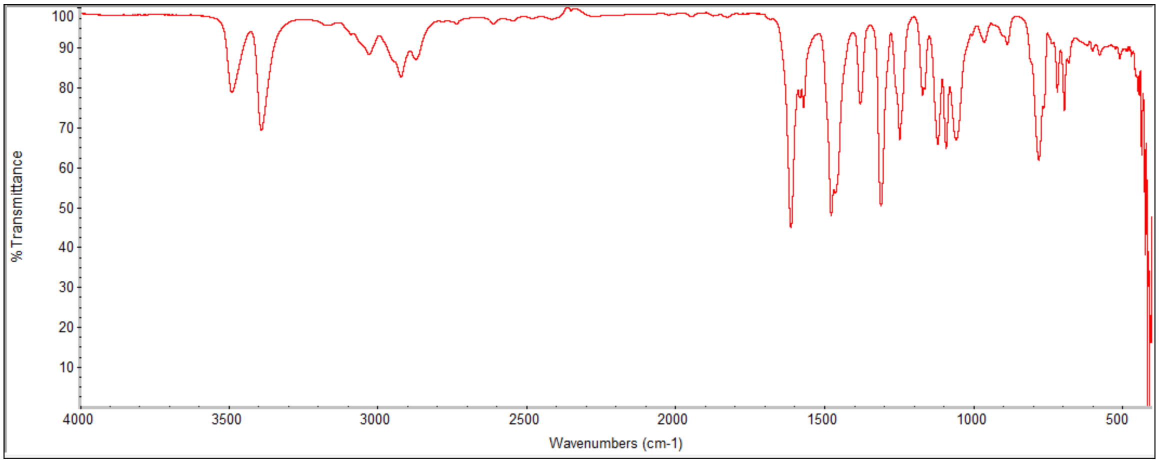
% Transmittance
100 90 80 70 60 50 40 30 20 10
4000 3500 3000 2500 2000 1500 1000 500
Wavenumbers (cm-1)

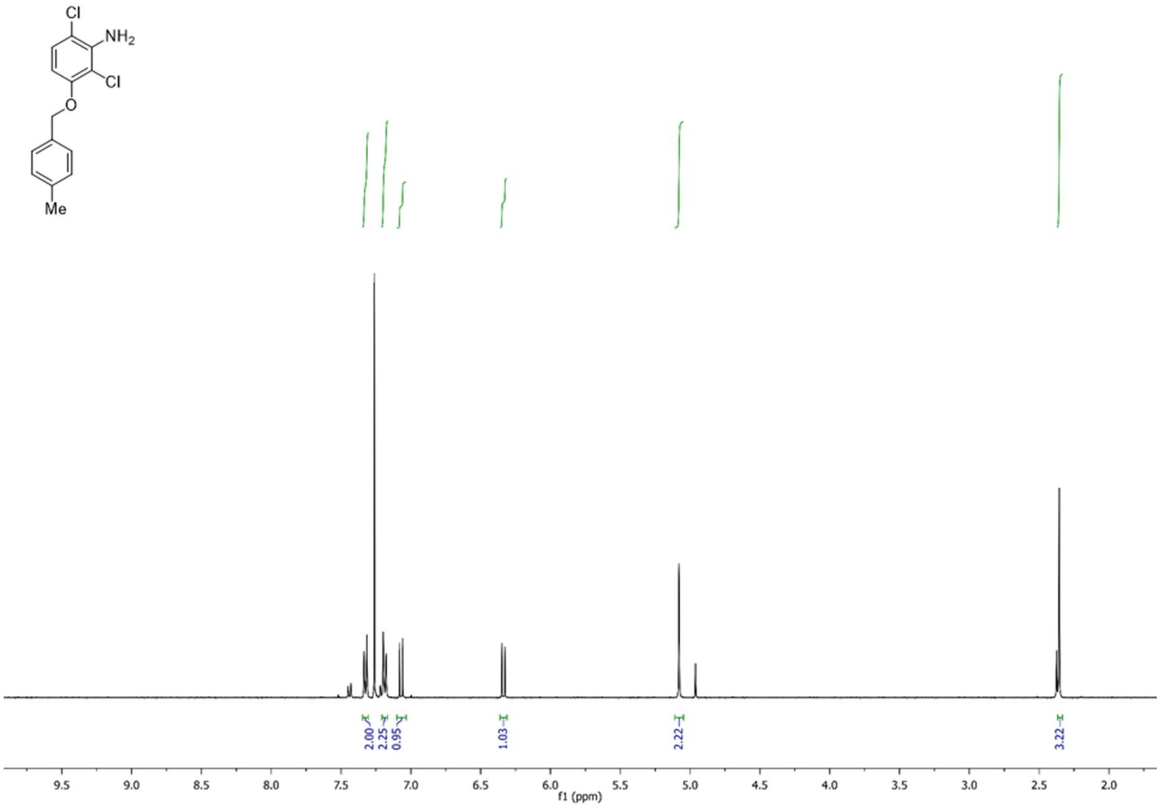
Cl
NH2
Cl
O
Me
2.00
2.25
0.95
1.03
2.22
3.22
f1 (ppm)

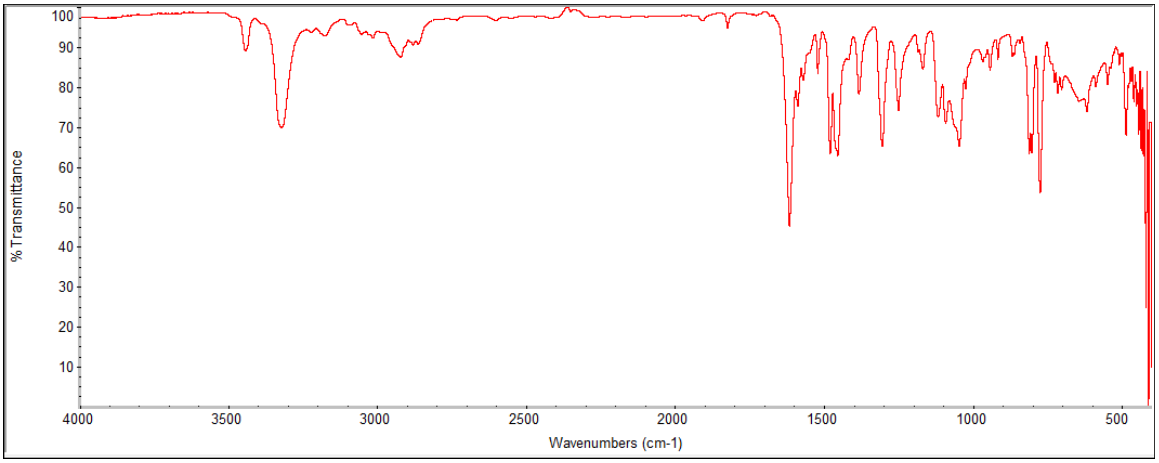
% Transmittance
Wavenumbers (cm-1)

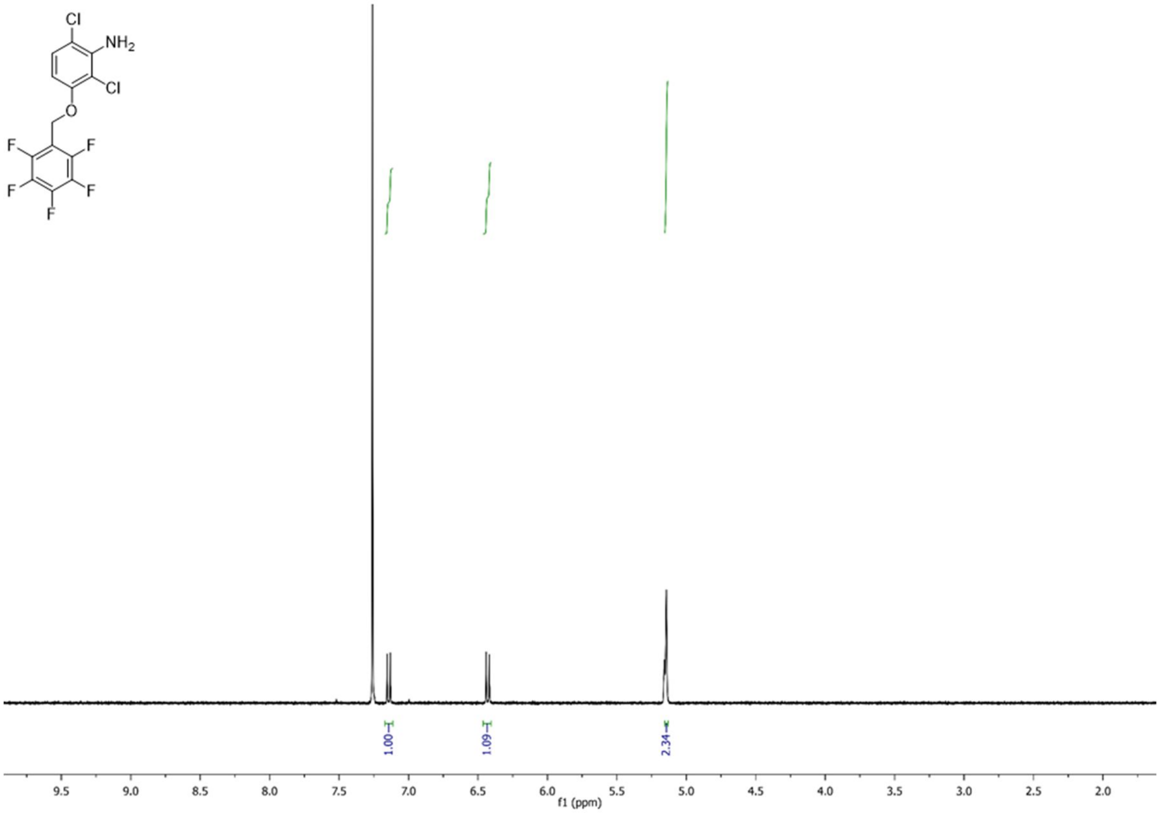
Cl
NH2
Cl
O
F
F
F
F
F
1.00
1.09
2.34
9.5
9.0
8.5
8.0
7.5
7.0
6.5
6.0
5.5
5.0
4.5
4.0
3.5
3.0
2.5
2.0
f1 (ppm)

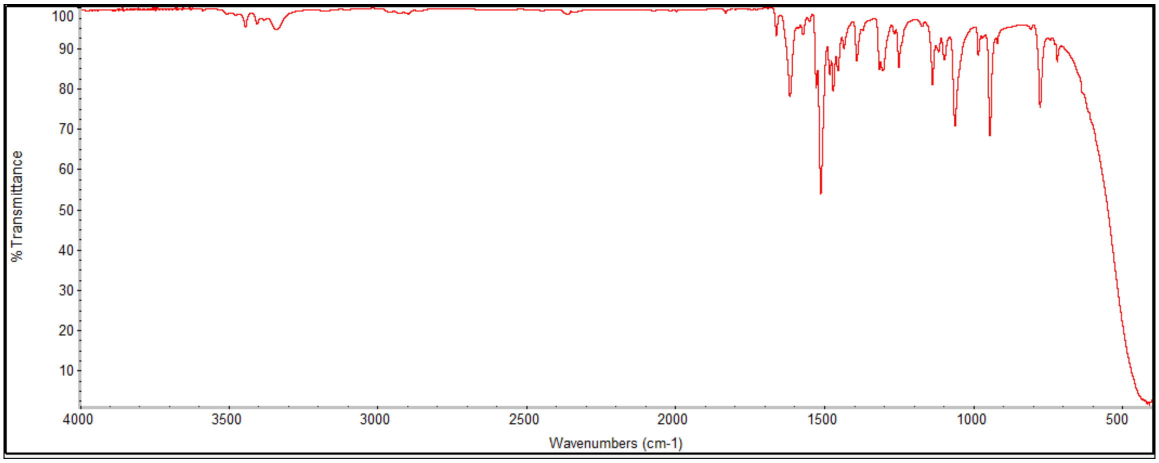
% Transmittance
100
90
80
70
60
50
40
30
20
10
4000
3500
3000
2500
2000
1500
1000
500
Wavenumbers (cm-1)

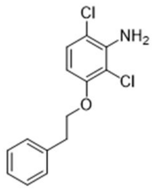
Cl
NH2
Cl
O

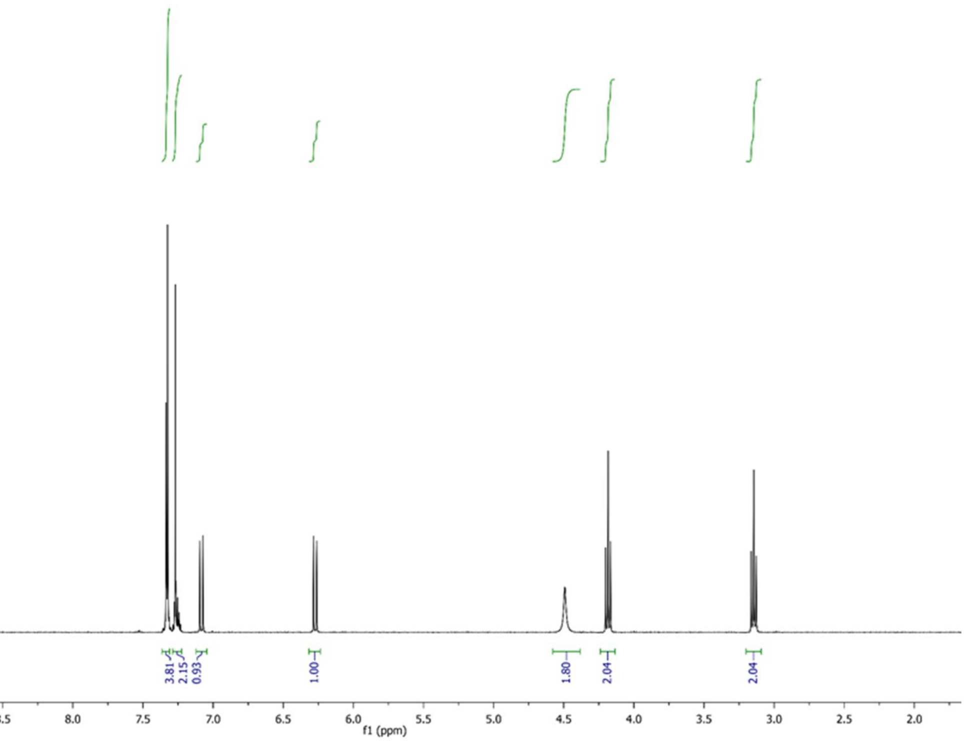
3.81
2.15
0.93
1.00
1.80
2.04
2.04
9.5
9.0
8.5
8.0
7.5
7.0
6.5
6.0
5.5
5.0
4.5
4.0
3.5
3.0
2.5
2.0
f1 (ppm)

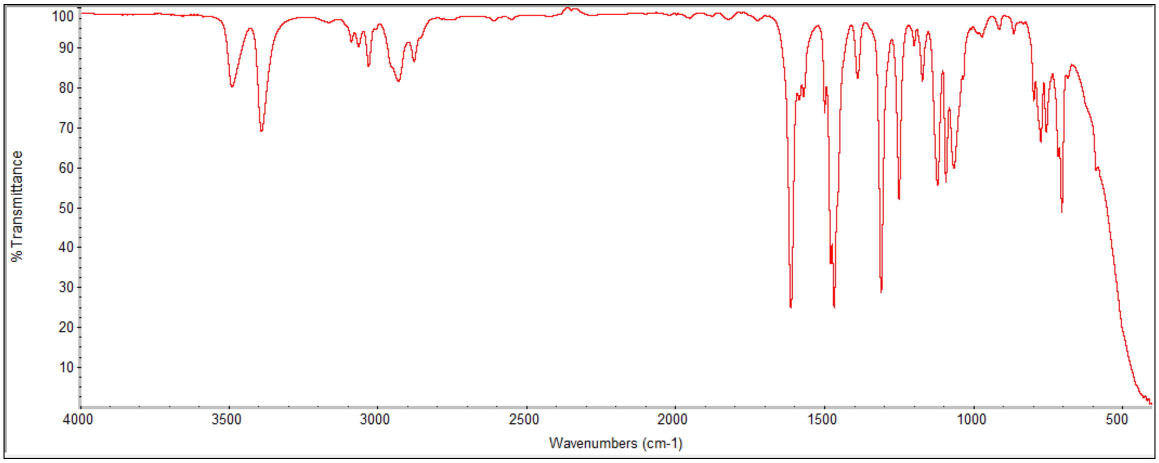
% Transmittance
100
90
80
70
60
50
40
30
20
10
4000
3500
3000
2500
2000
1500
1000
500
Wavenumbers (cm-1)

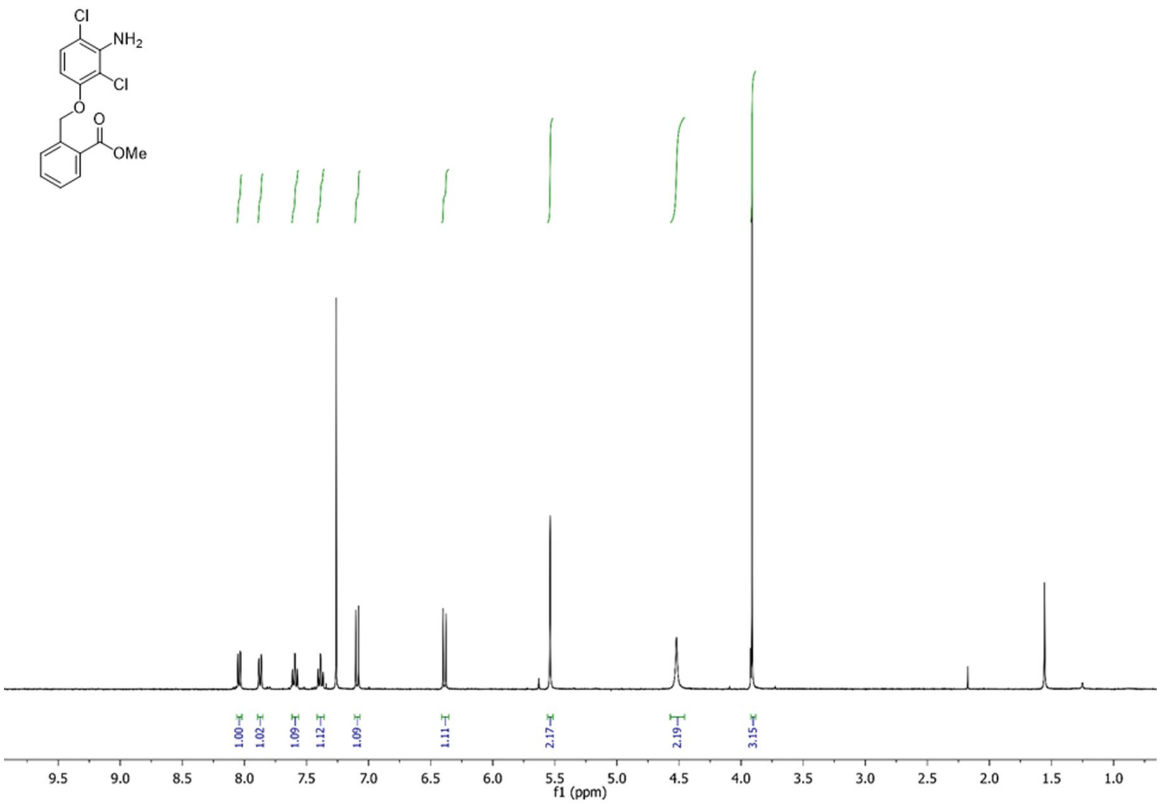
Cl
NH2
Cl
O
O
OMe
1.00
1.02
1.09
1.12
1.09
1.11
2.17
2.19
3.15
9.5 9.0 8.5 8.0 7.5 7.0 6.5 6.0 5.5 5.0 4.5 4.0 3.5 3.0 2.5 2.0 1.5 1.0
f1 (ppm)

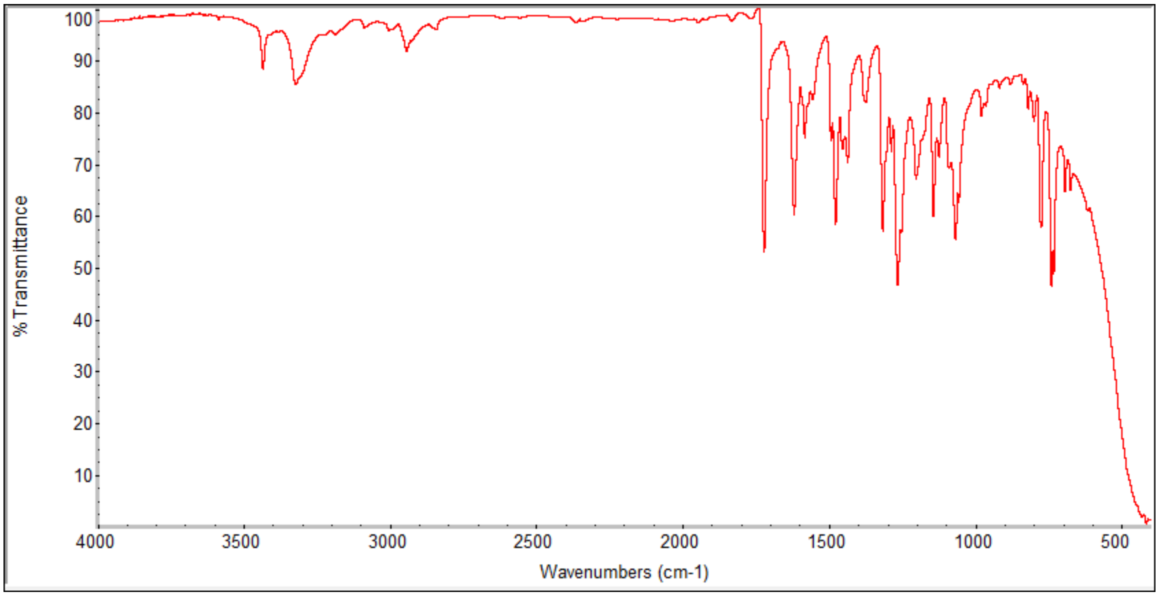
% Transmittance
100
90
80
70
60
50
40
30
20
10
4000 3500 3000 2500 2000 1500 1000 500
Wavenumbers (cm-1)

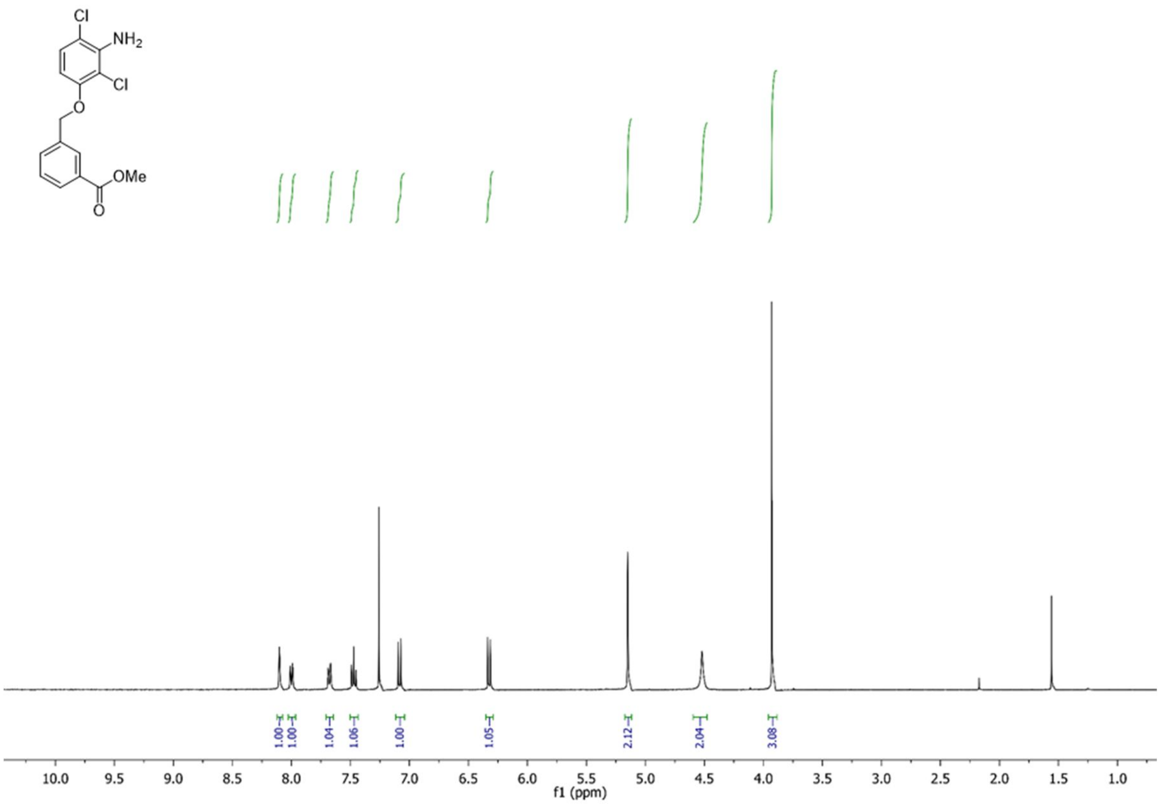

Cl
NH2
Cl
O
OMe
O
1.00
1.00
1.04
1.06
1.00
1.05
2.12
2.04
3.08
10.0
9.5
9.0
8.5
8.0
7.5
7.0
6.5
6.0
5.5
5.0
4.5
4.0
3.5
3.0
2.5
2.0
1.5
1.0
f1 (ppm)

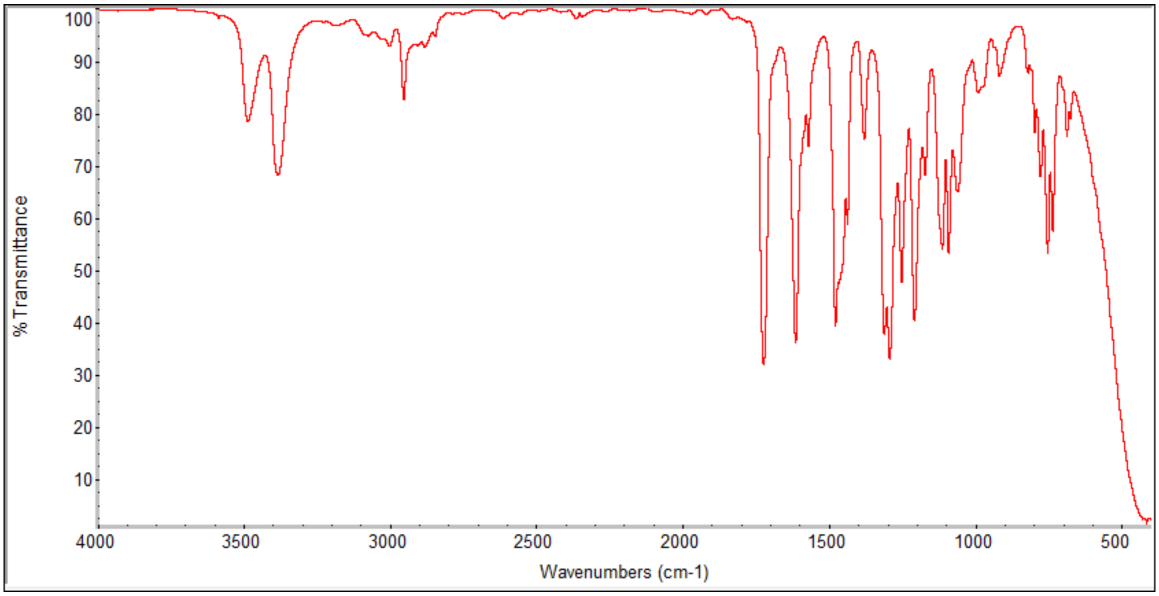

% Transmittance
Wavenumbers (cm-1)
100
90
80
70
60
50
40
30
20
10
4000
3500
3000
2500
2000
1500
1000
500

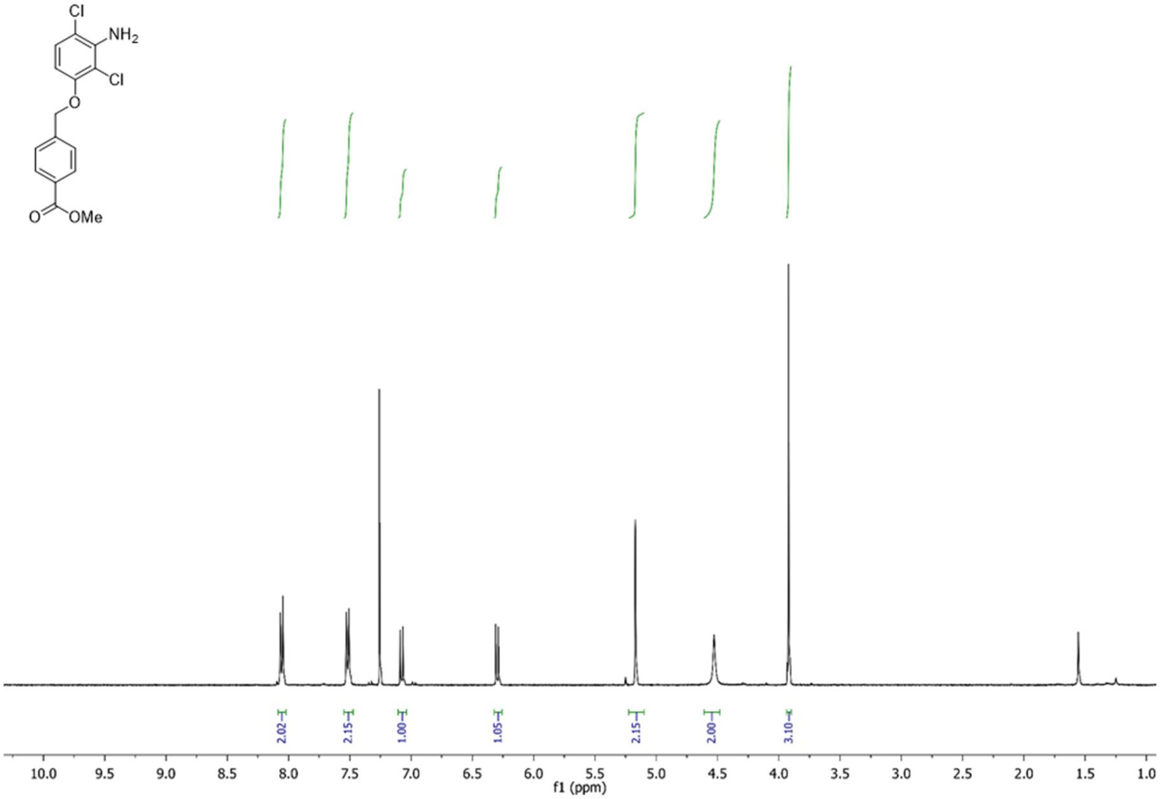
Cl
NH2
Cl
O
O
OMe
2.02
2.15
1.00
1.05
2.15
2.00
3.10
10.0 9.5 9.0 8.5 8.0 7.5 7.0 6.5 6.0 5.5 5.0 4.5 4.0 3.5 3.0 2.5 2.0 1.5 1.0
f1 (ppm)

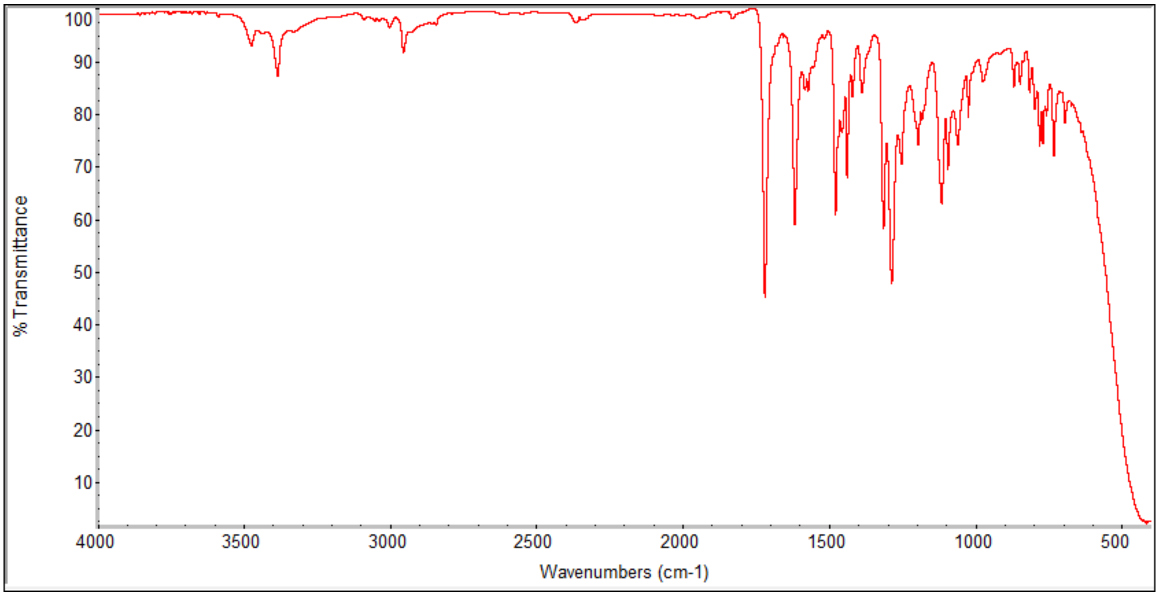
% Transmittance
100
90
80
70
60
50
40
30
20
10
4000 3500 3000 2500 2000 1500 1000 500
Wavenumbers (cm-1)

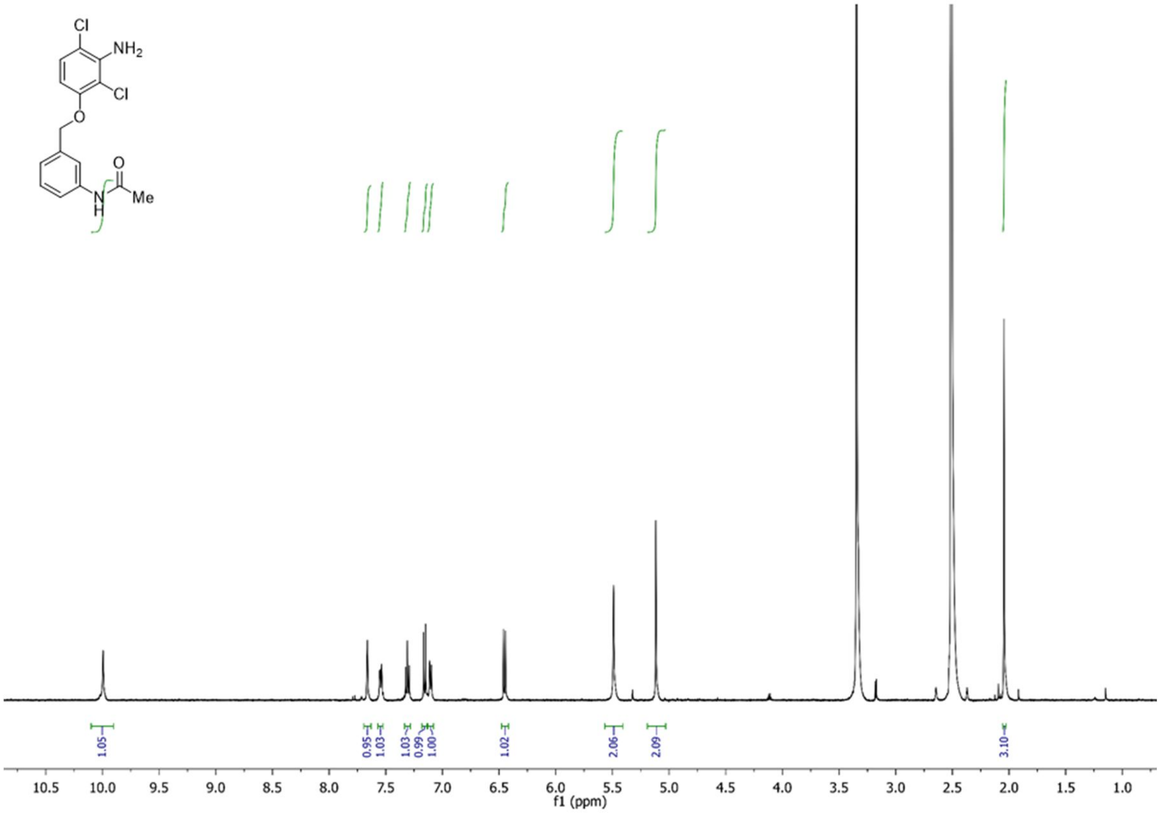

Cl
NH2
Cl
O
O
N
H
Me
1.05
0.95
1.03
1.03
0.99
1.00
1.02
2.06
2.09
3.10
10.5
10.0
9.5
9.0
8.5
8.0
7.5
7.0
6.5
6.0
5.5
5.0
4.5
4.0
3.5
3.0
2.5
2.0
1.5
1.0
f1 (ppm)

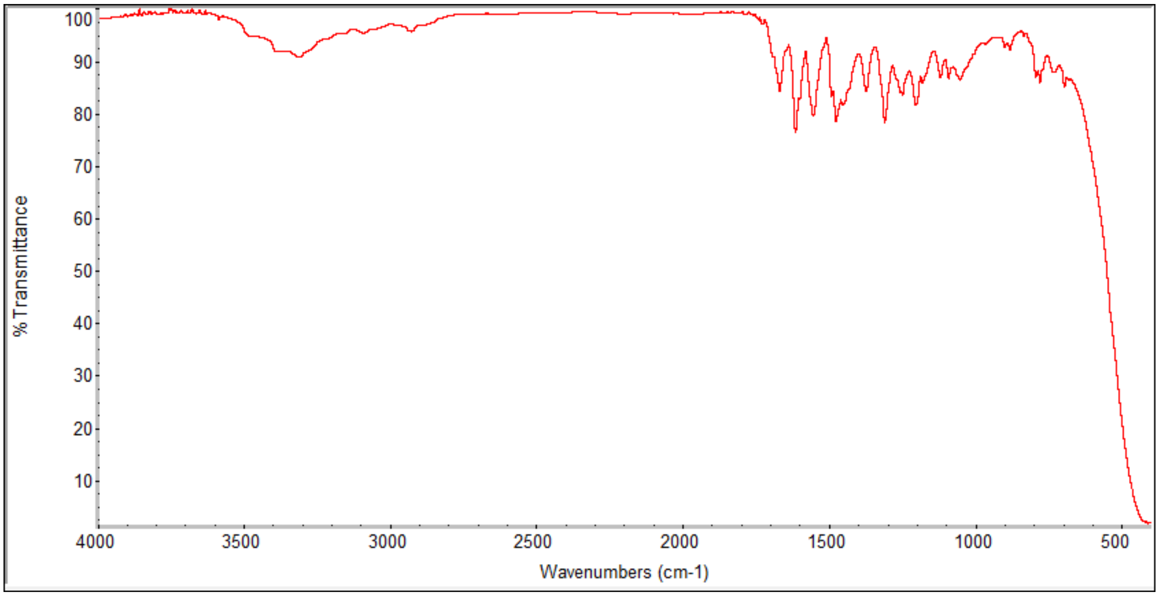

% Transmittance
100
90
80
70
60
50
40
30
20
10
4000
3500
3000
2500
2000
1500
1000
500
Wavenumbers (cm-1)

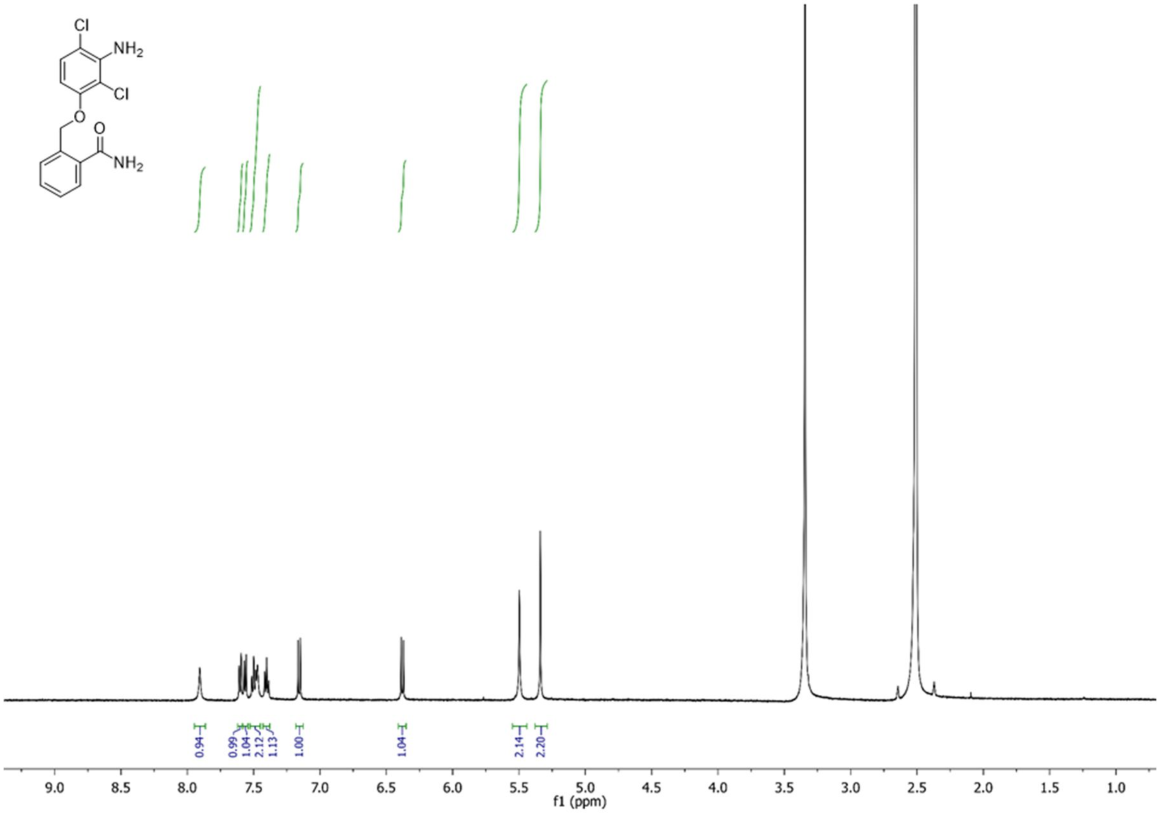

Cl
NH2
Cl
O
O
NH2
0.94
0.99
1.04
2.12
1.13
1.00
1.04
2.14
2.20
9.0
8.5
8.0
7.5
7.0
6.5
6.0
5.5
5.0
4.5
4.0
3.5
3.0
2.5
2.0
1.5
1.0
f1 (ppm)

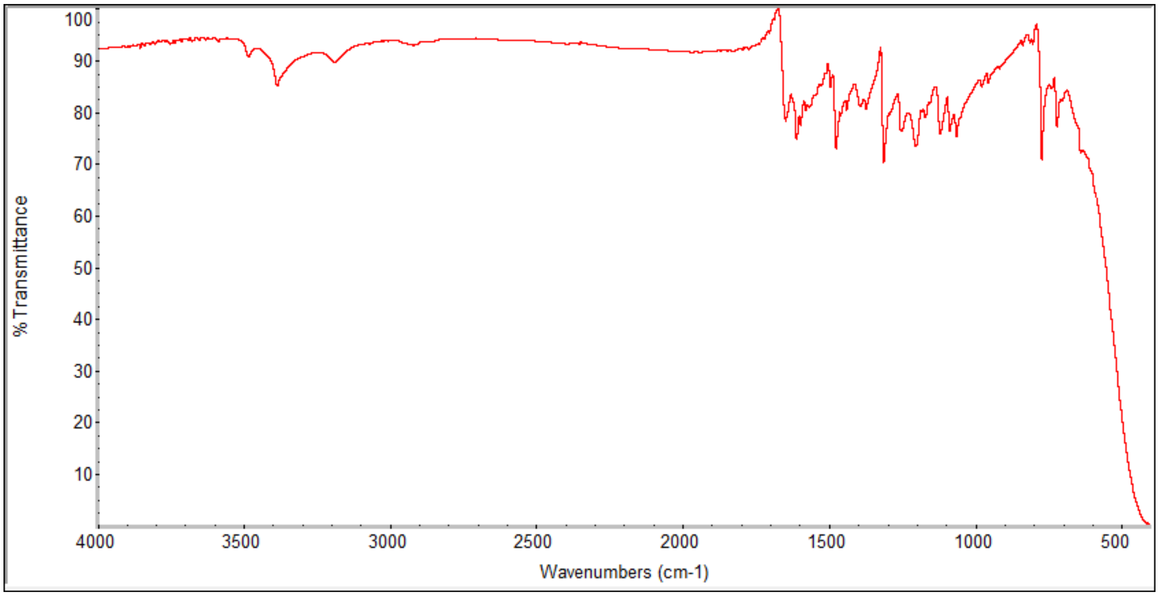

% Transmittance
Wavenumbers (cm-1)
100
90
80
70
60
50
40
30
20
10
4000
3500
3000
2500
2000
1500
1000
500

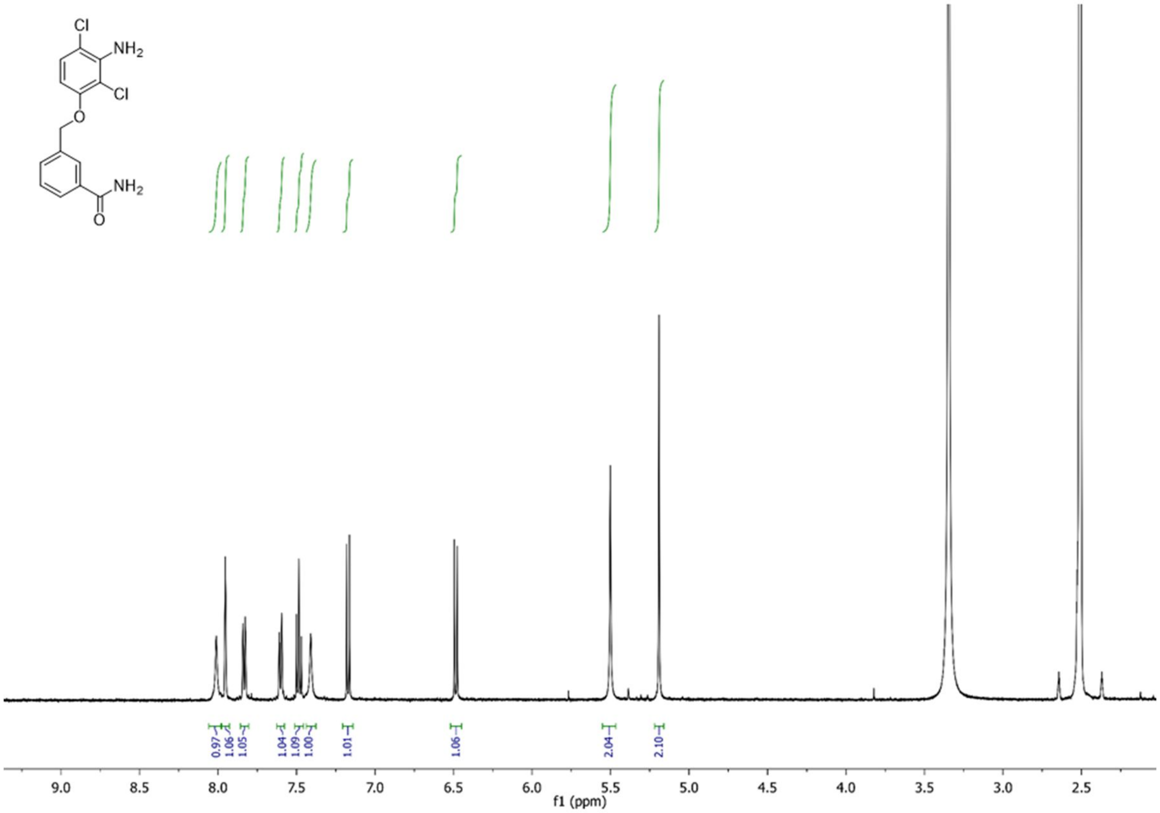
Cl
NH2
Cl
O
NH2
O
0.97
1.06
1.05
1.04
1.09
1.00
1.01
1.06
2.04
2.10
9.0
8.5
8.0
7.5
7.0
6.5
6.0
5.5
5.0
4.5
4.0
3.5
3.0
2.5
f1 (ppm)

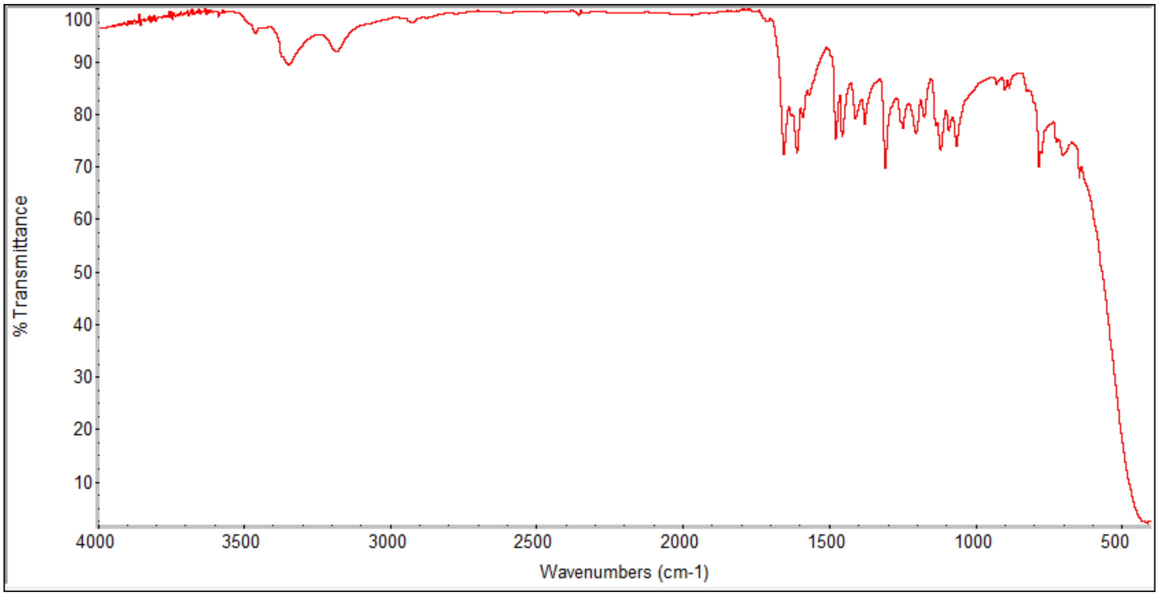
% Transmittance
100
90
80
70
60
50
40
30
20
10
4000
3500
3000
2500
2000
1500
1000
500
Wavenumbers (cm-1)

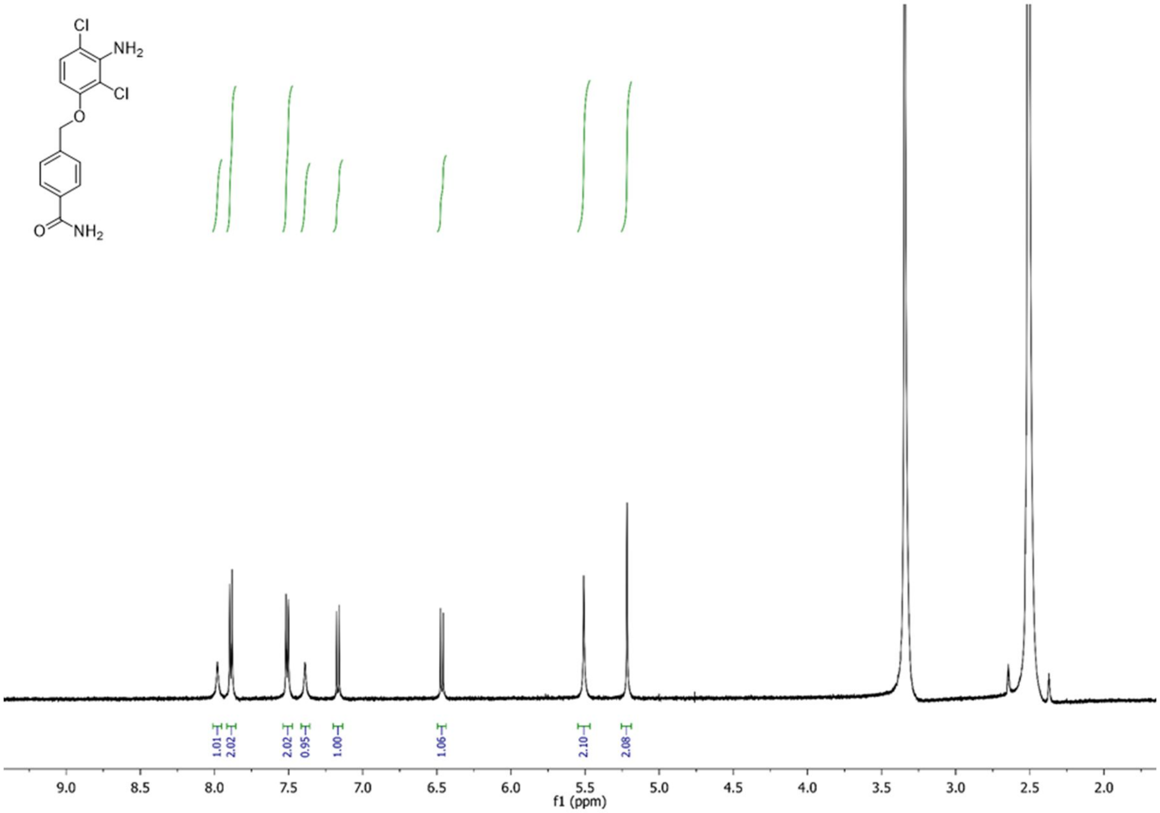
Cl
NH2
Cl
O
O
NH2
1.01
2.02
2.02
0.95
1.00
1.06
2.10
2.08
9.0
8.5
8.0
7.5
7.0
6.5
6.0
5.5
5.0
4.5
4.0
3.5
3.0
2.5
2.0
f1 (ppm)

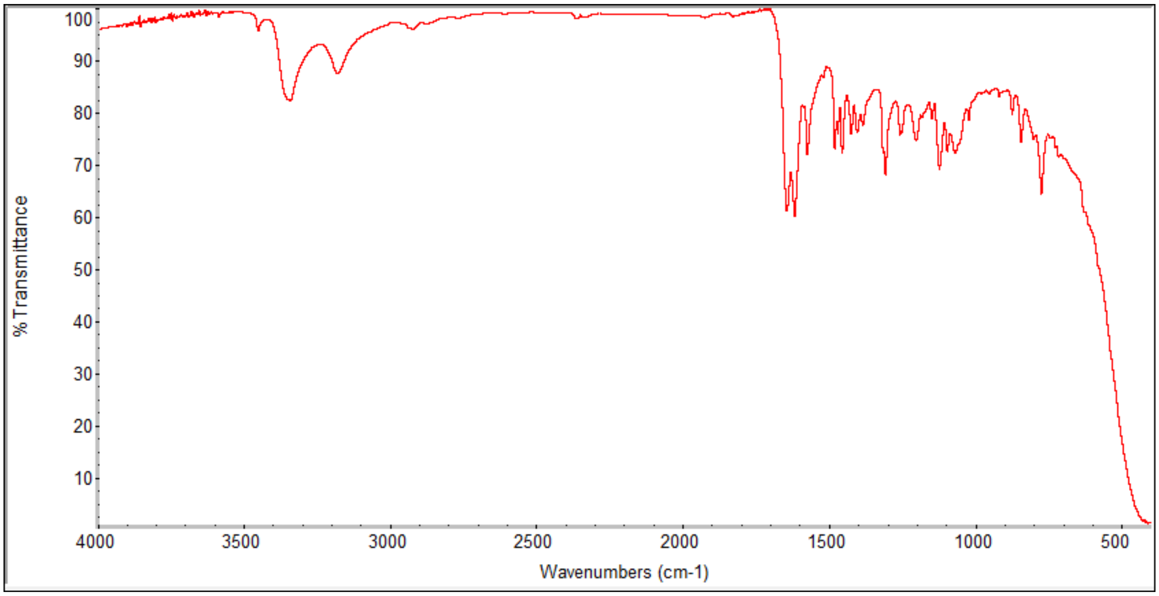
% Transmittance
100
90
80
70
60
50
40
30
20
10
4000
3500
3000
2500
2000
1500
1000
500
Wavenumbers (cm-1)

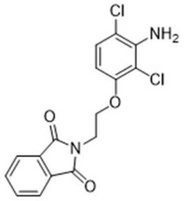

Cl
NH2
Cl
O
O
N
O

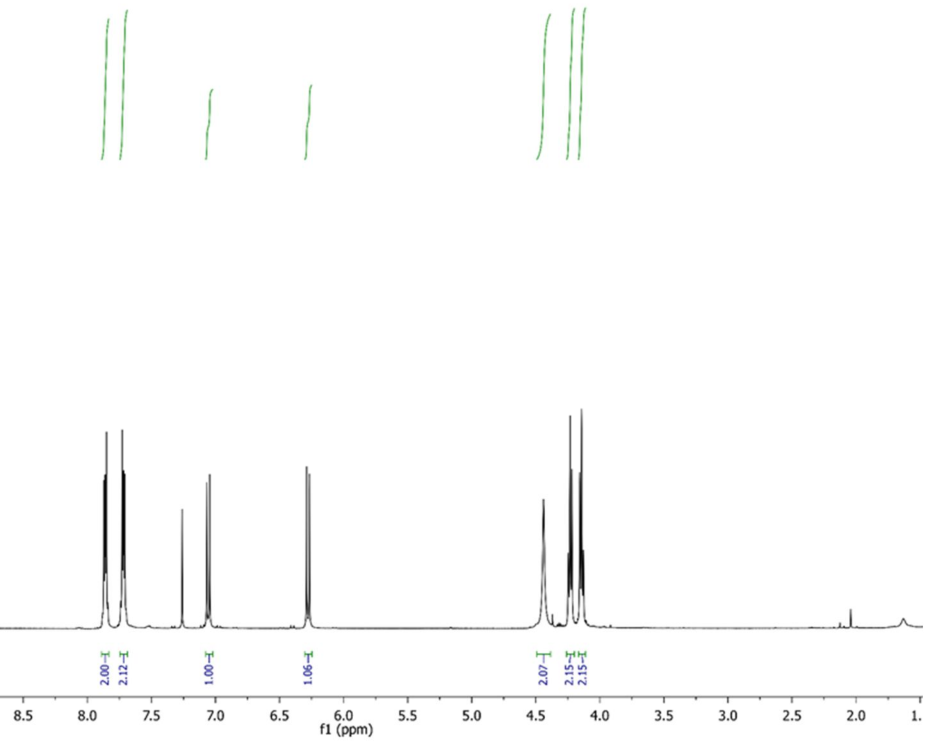

2.00
2.12
1.00
1.06
2.07
2.15
2.15
10.0
9.5
9.0
8.5
8.0
7.5
7.0
6.5
6.0
5.5
5.0
4.5
4.0
3.5
3.0
2.5
2.0
f1 (ppm)

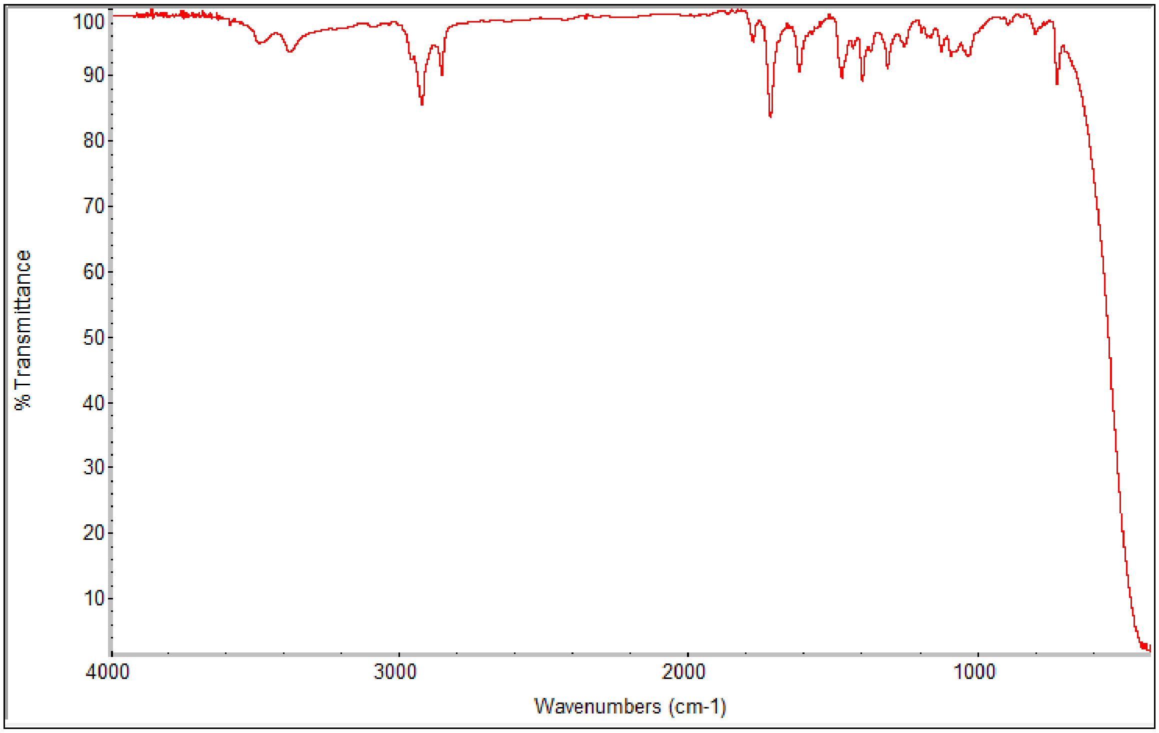

% Transmittance
100
90
80
70
60
50
40
30
20
10
4000
3000
2000
1000
Wavenumbers (cm-1)

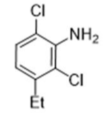
Cl
NH2
Cl
Et

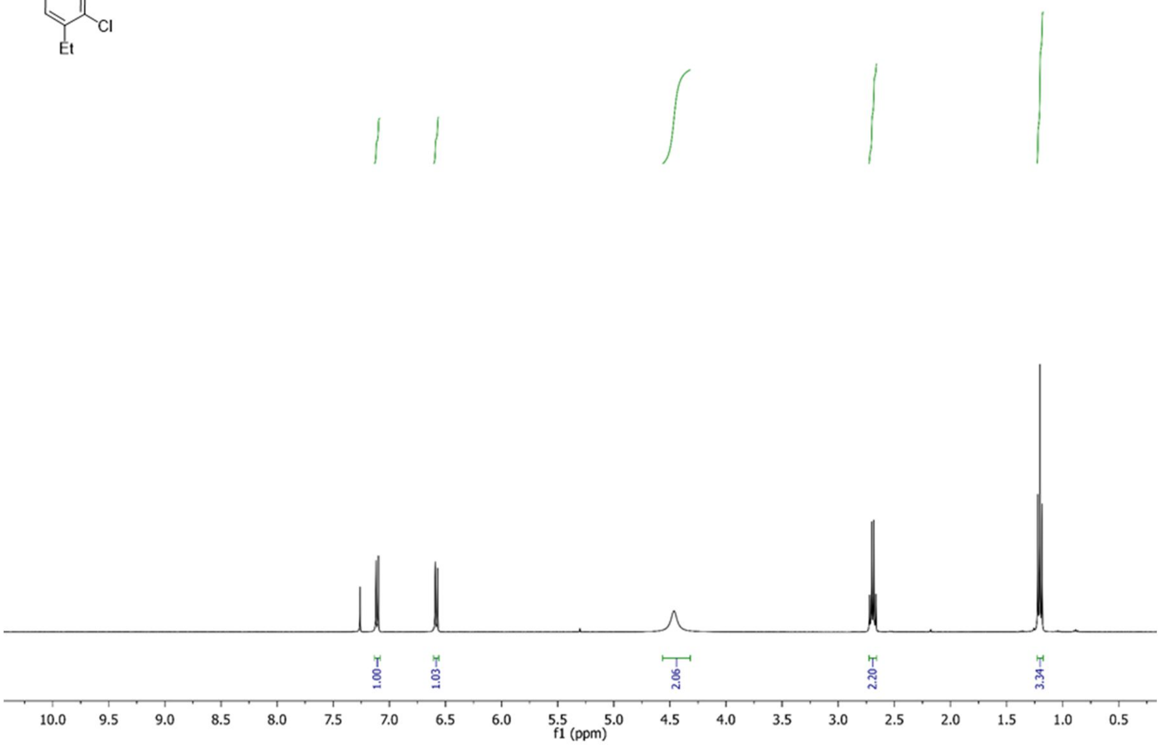
1.00
1.03
2.06
2.20
3.34
10.0 9.5 9.0 8.5 8.0 7.5 7.0 6.5 6.0 5.5 5.0 4.5 4.0 3.5 3.0 2.5 2.0 1.5 1.0 0.5
f1 (ppm)

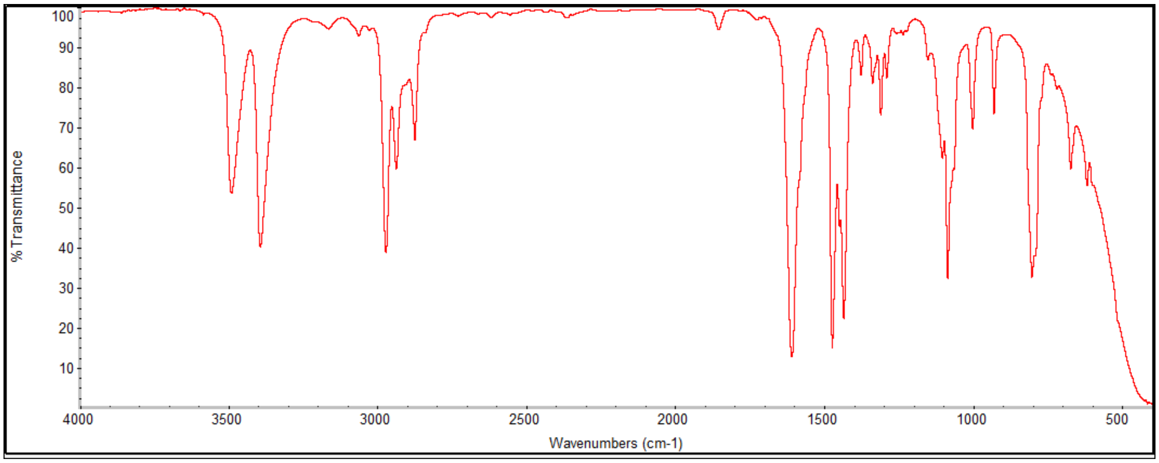
100
90
80
70
60
50
40
30
20
10
% Transmittance
4000 3500 3000 2500 2000 1500 1000 500
Wavenumbers (cm-1)

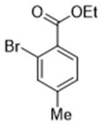
O
OEt
Br
Me

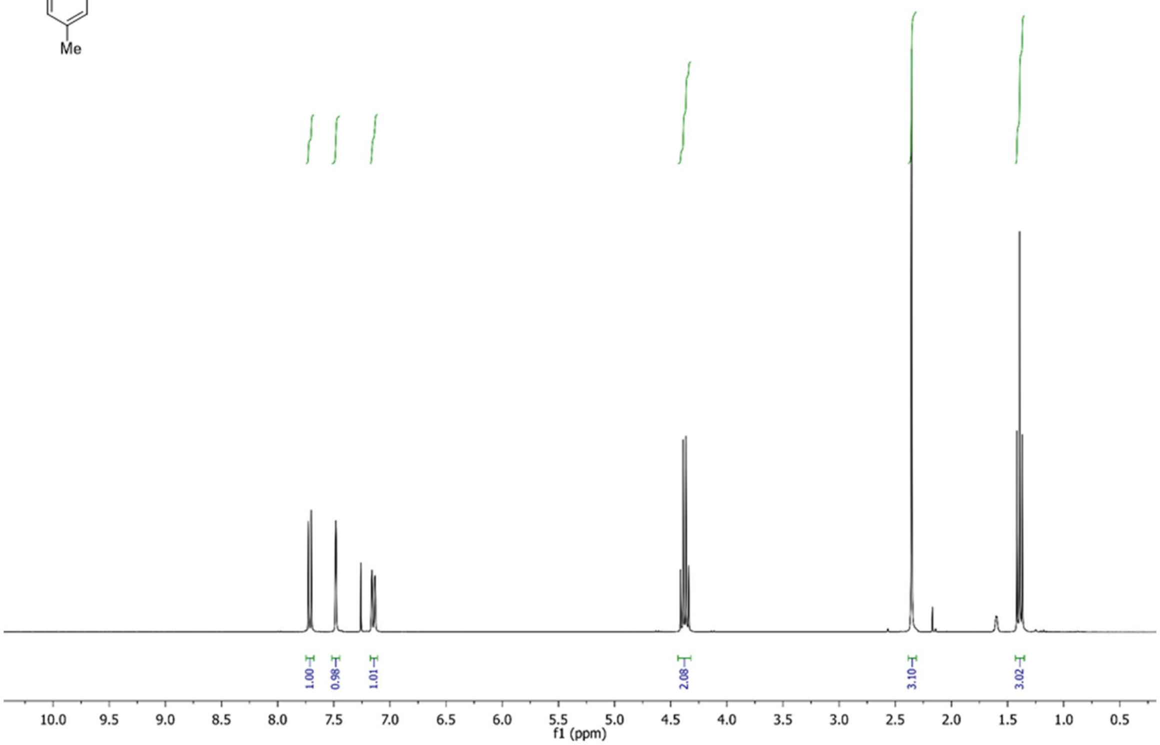
1.00
0.98
1.01
2.08
3.10
3.02
10.0 9.5 9.0 8.5 8.0 7.5 7.0 6.5 6.0 5.5 5.0 4.5 4.0 3.5 3.0 2.5 2.0 1.5 1.0 0.5
f1 (ppm)

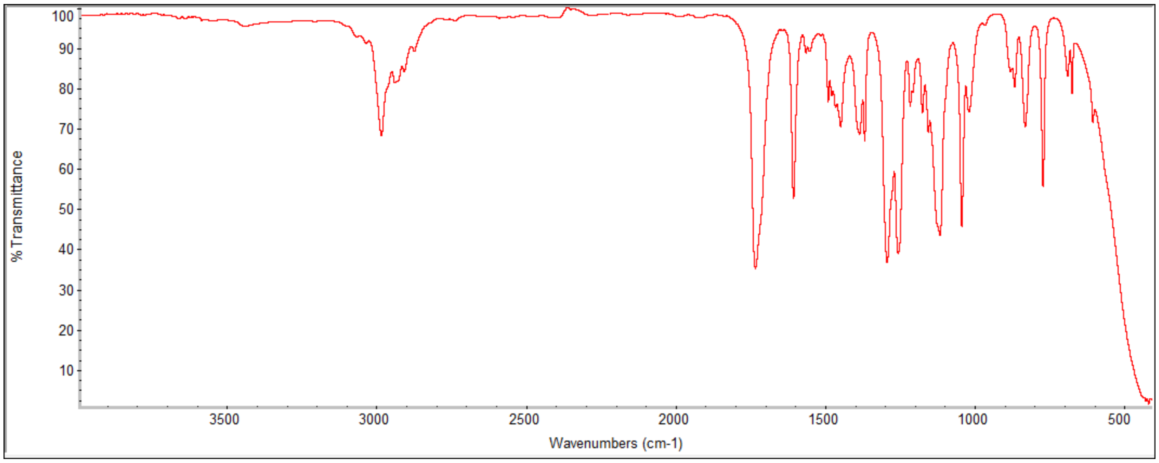
% Transmittance
100 90 80 70 60 50 40 30 20 10
3500 3000 2500 2000 1500 1000 500
Wavenumbers (cm-1)

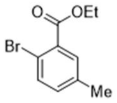
O
OEt
Br
Me

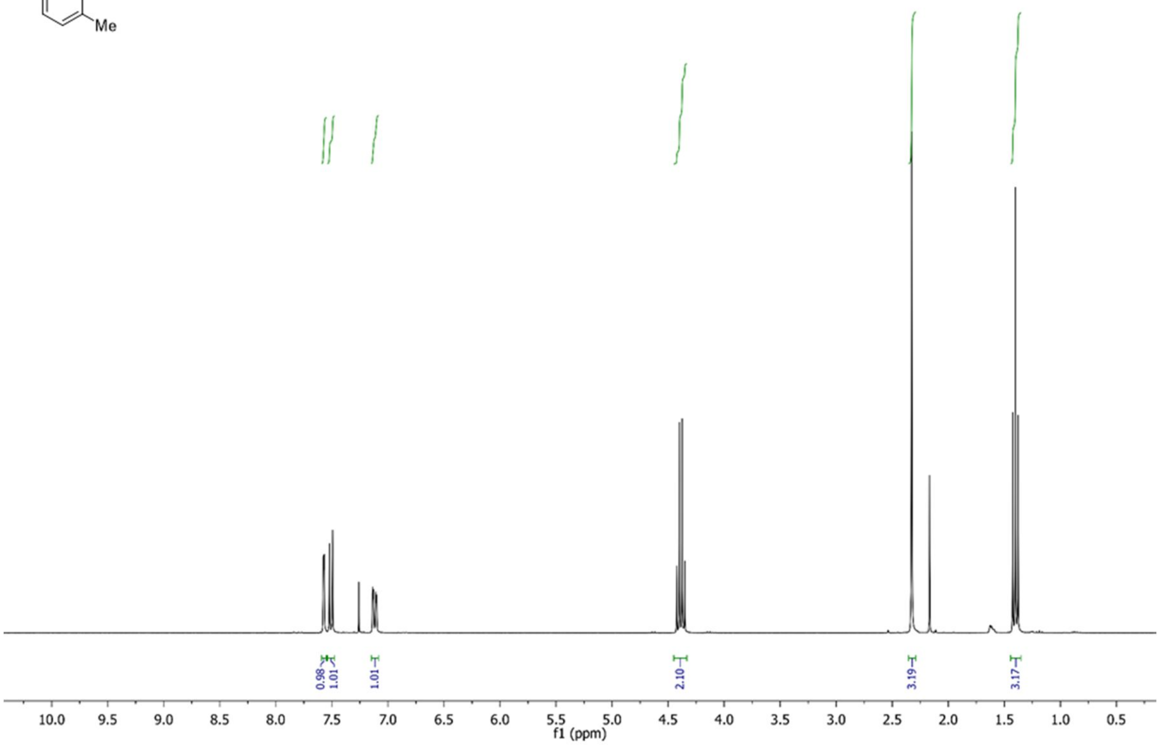
0.98
1.01
1.01
2.10
3.19
3.17
10.0 9.5 9.0 8.5 8.0 7.5 7.0 6.5 6.0 5.5 5.0 4.5 4.0 3.5 3.0 2.5 2.0 1.5 1.0 0.5
f1 (ppm)

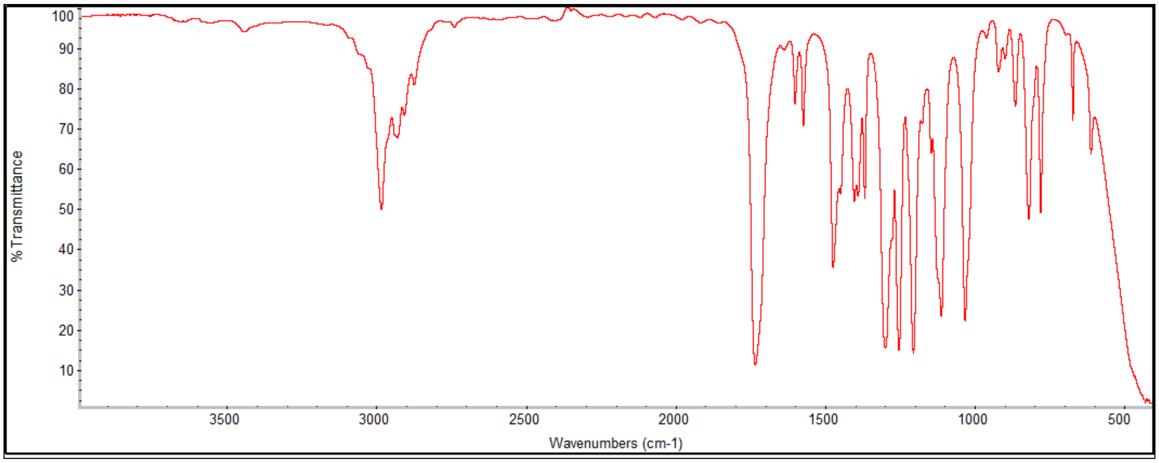
100
90
80
70
60
50
40
30
20
10
% Transmittance
3500 3000 2500 2000 1500 1000 500
Wavenumbers (cm-1)

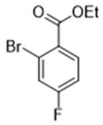
O
OEt
Br
F

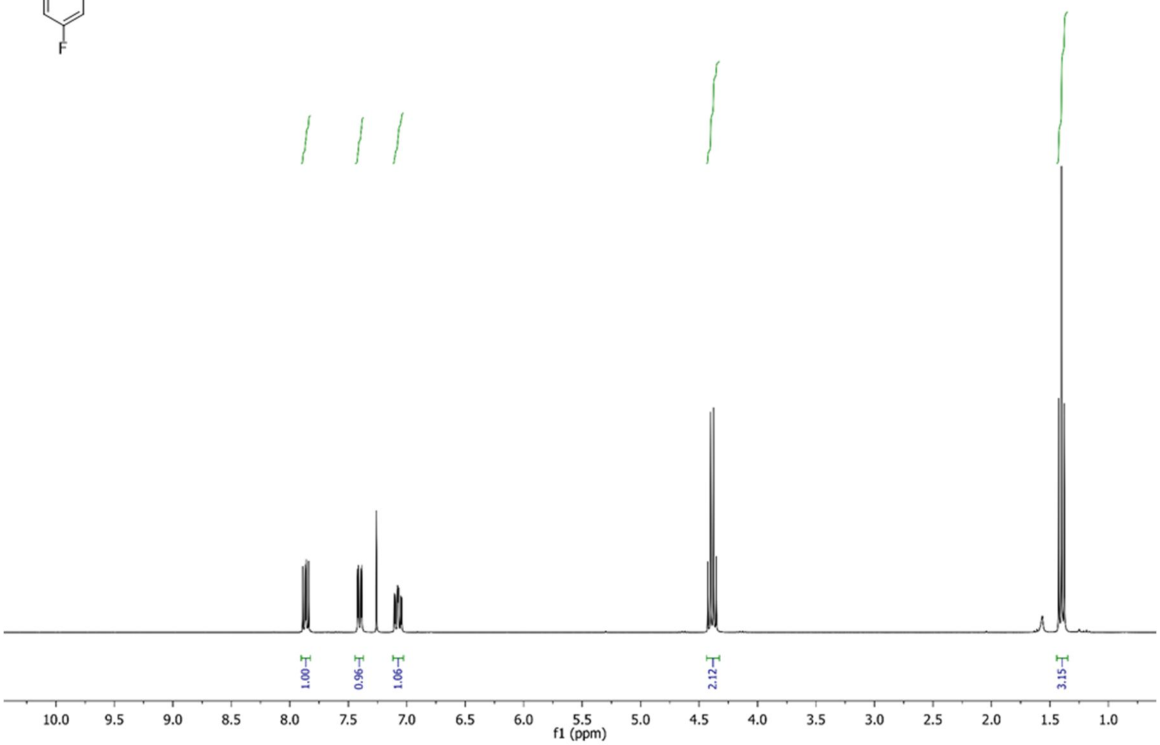
1.00
0.96
1.06
2.12
3.15
10.0 9.5 9.0 8.5 8.0 7.5 7.0 6.5 6.0 5.5 5.0 4.5 4.0 3.5 3.0 2.5 2.0 1.5 1.0
f1 (ppm)

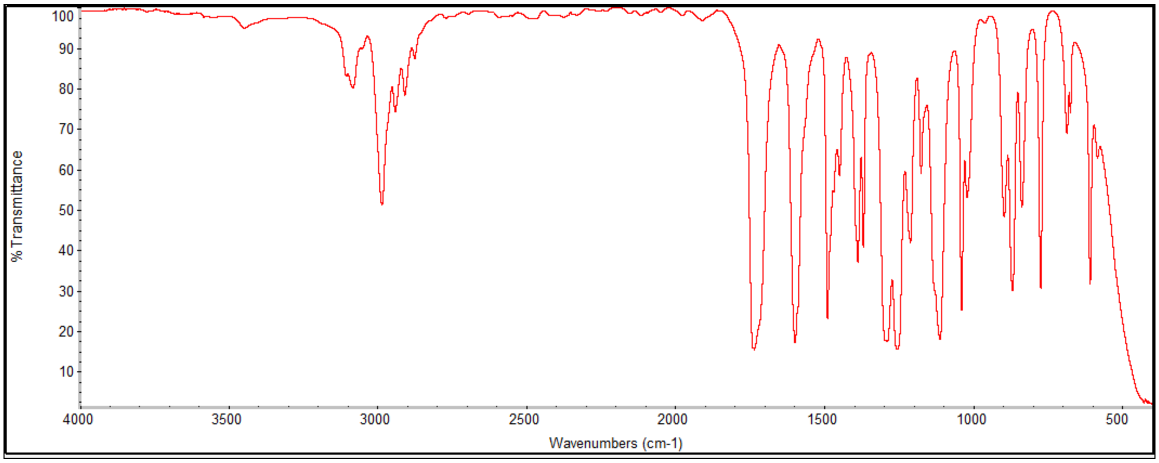
%Transmittance
100 90 80 70 60 50 40 30 20 10
4000 3500 3000 2500 2000 1500 1000 500
Wavenumbers (cm-1)

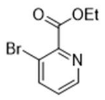
O
OEt
Br
N

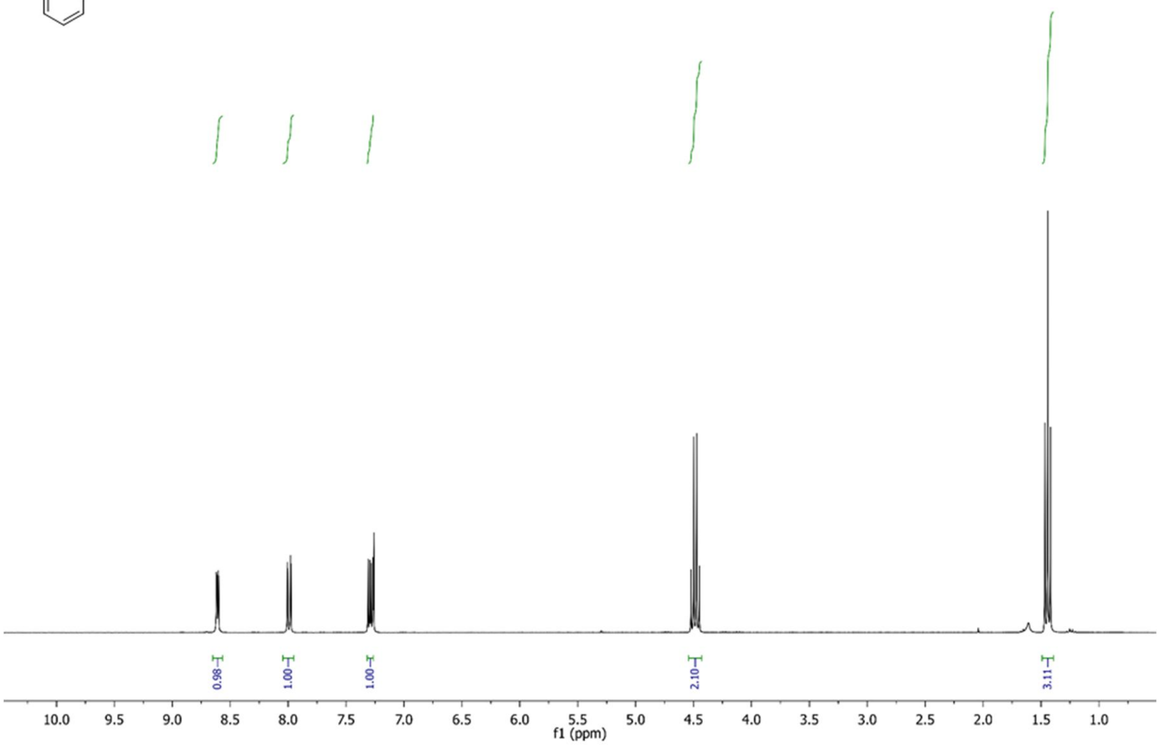
0.98
1.00
1.00
2.10
3.11
10.0 9.5 9.0 8.5 8.0 7.5 7.0 6.5 6.0 5.5 5.0 4.5 4.0 3.5 3.0 2.5 2.0 1.5 1.0
f1 (ppm)

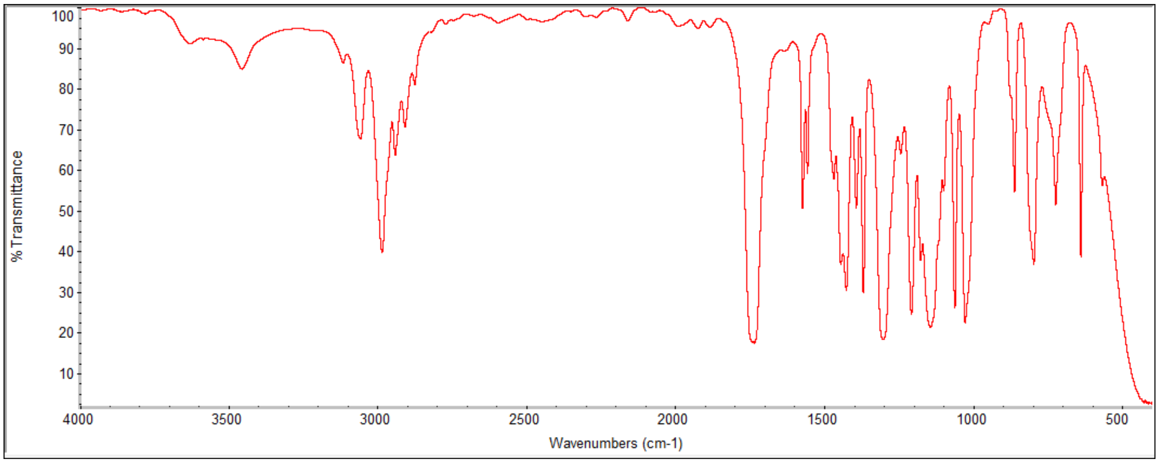
% Transmittance
100 90 80 70 60 50 40 30 20 10
4000 3500 3000 2500 2000 1500 1000 500
Wavenumbers (cm-1)

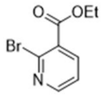
O
OEt
Br
N

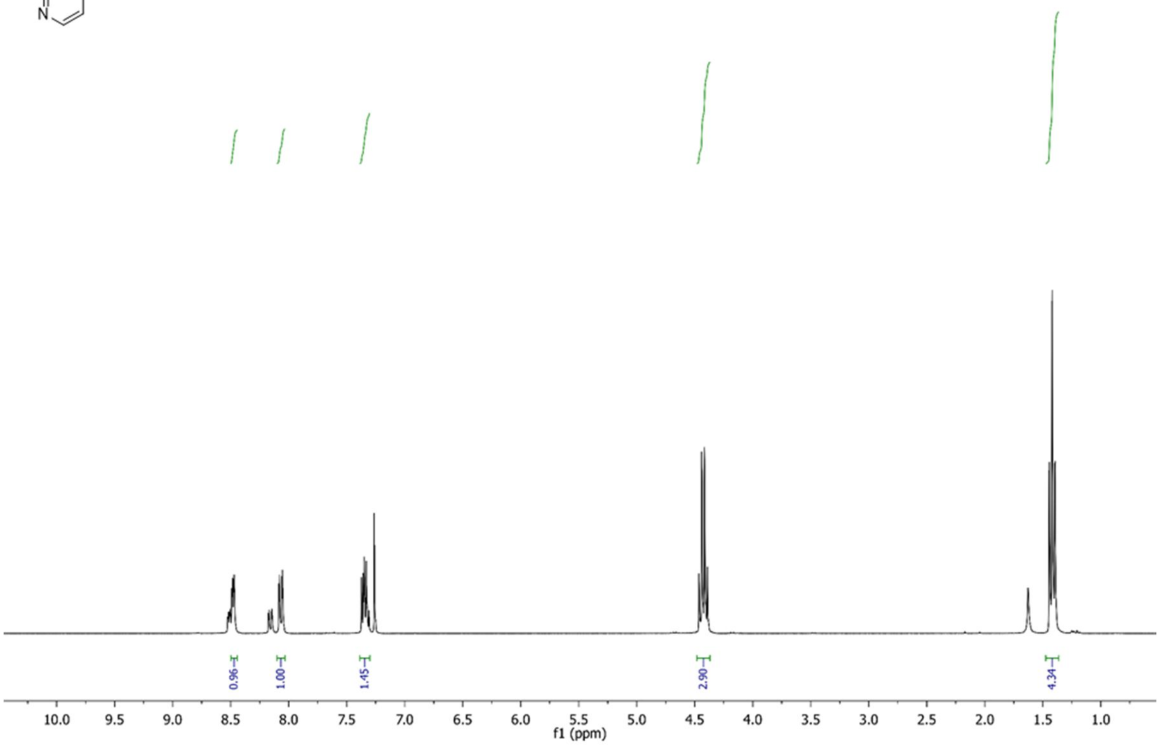
0.96
1.00
1.45
2.90
4.34
10.0
9.5
9.0
8.5
8.0
7.5
7.0
6.5
6.0
5.5
5.0
4.5
4.0
3.5
3.0
2.5
2.0
1.5
1.0
f1 (ppm)

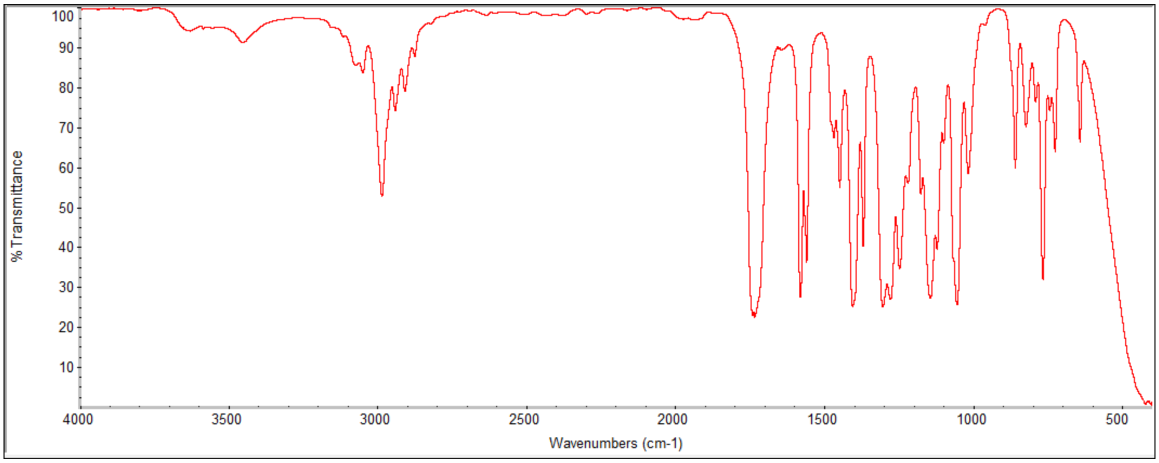
% Transmittance
100
90
80
70
60
50
40
30
20
10
4000
3500
3000
2500
2000
1500
1000
500
Wavenumbers (cm-1)

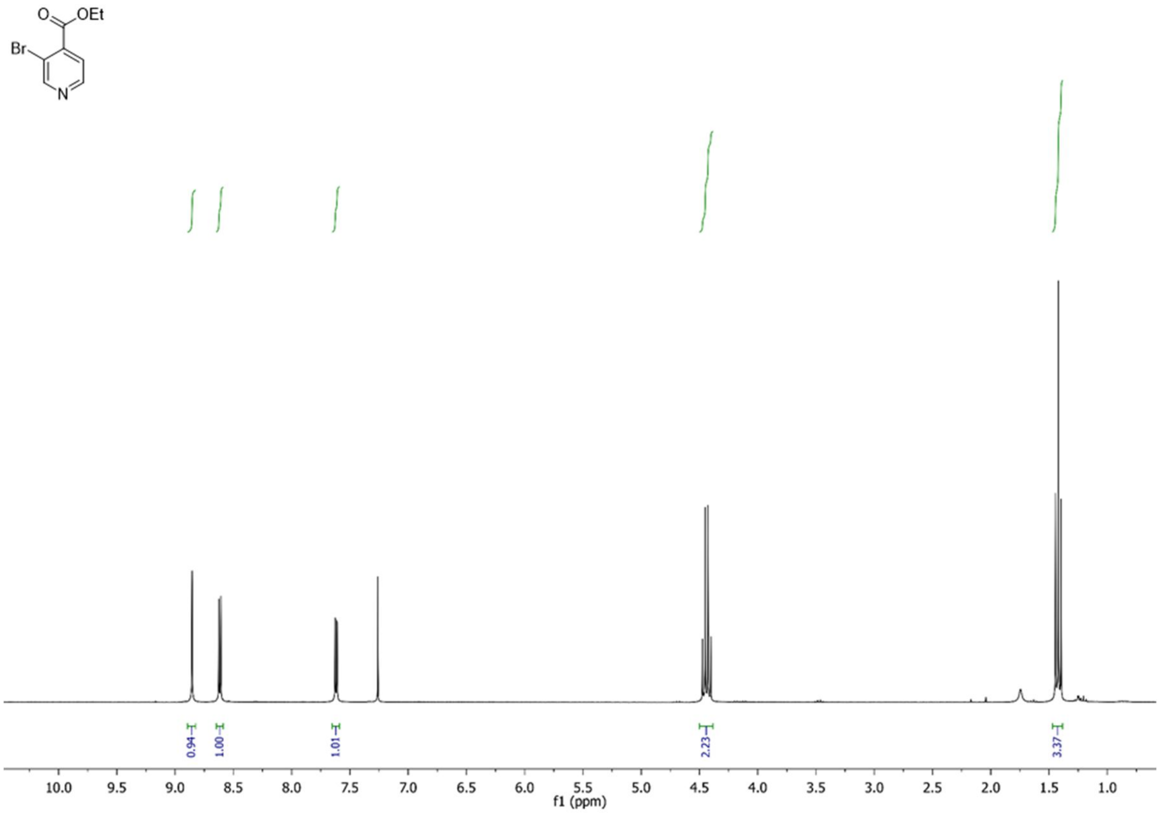
O
OEt
Br
N
0.94
1.00
1.01
2.23
3.37
10.0 9.5 9.0 8.5 8.0 7.5 7.0 6.5 6.0 5.5 5.0 4.5 4.0 3.5 3.0 2.5 2.0 1.5 1.0
f1 (ppm)

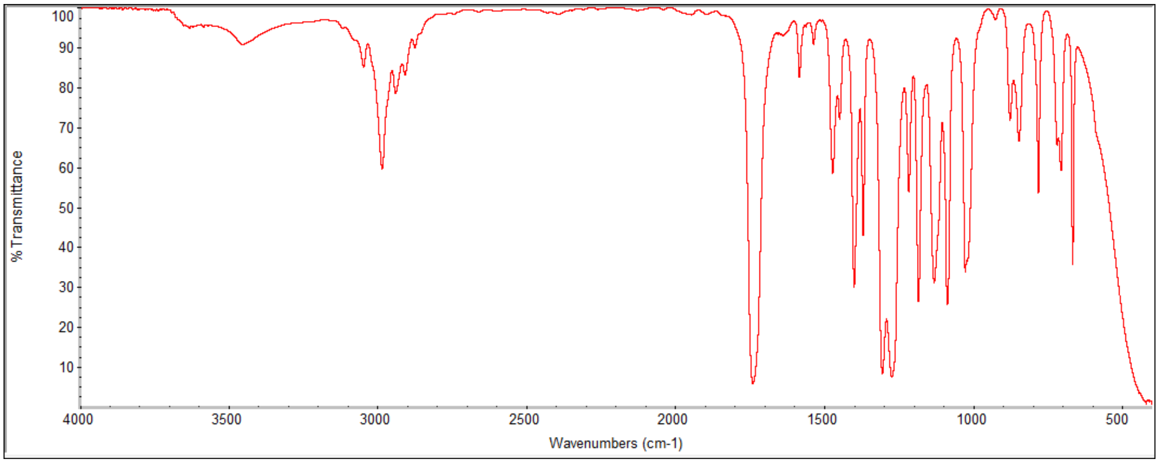
% Transmittance
100 90 80 70 60 50 40 30 20 10
4000 3500 3000 2500 2000 1500 1000 500
Wavenumbers (cm-1)

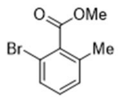
O
OMe
Br
Me

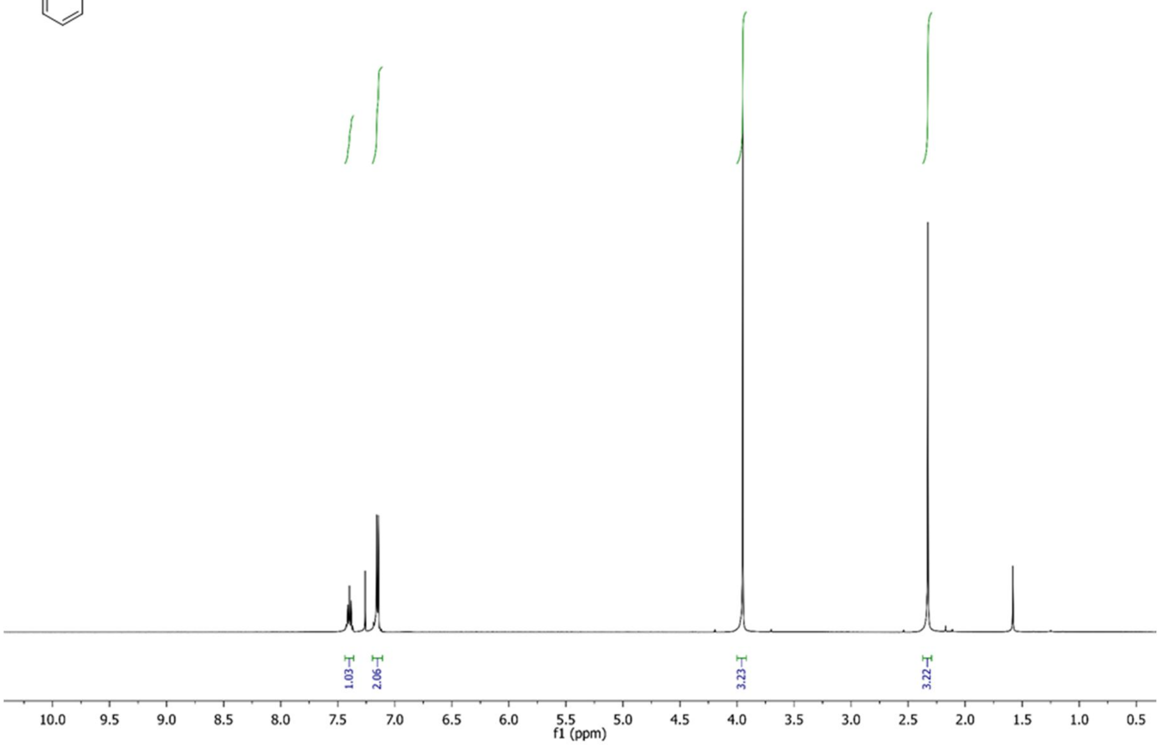
1.03
2.06
3.23
3.22
10.0
9.5
9.0
8.5
8.0
7.5
7.0
6.5
6.0
5.5
5.0
4.5
4.0
3.5
3.0
2.5
2.0
1.5
1.0
0.5
f1 (ppm)

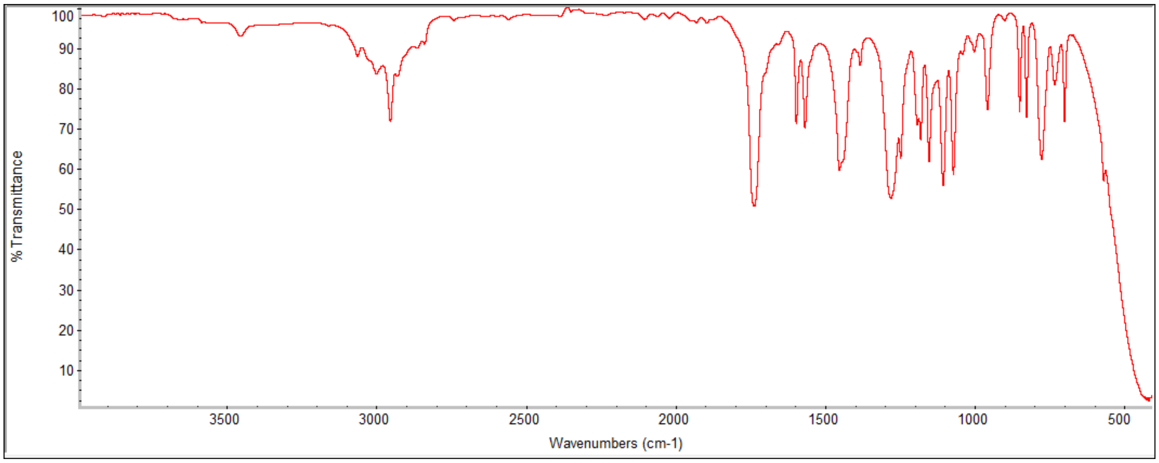
% Transmittance
100
90
80
70
60
50
40
30
20
10
3500
3000
2500
2000
1500
1000
500
Wavenumbers (cm-1)

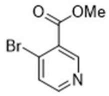
O
OMe
Br
N

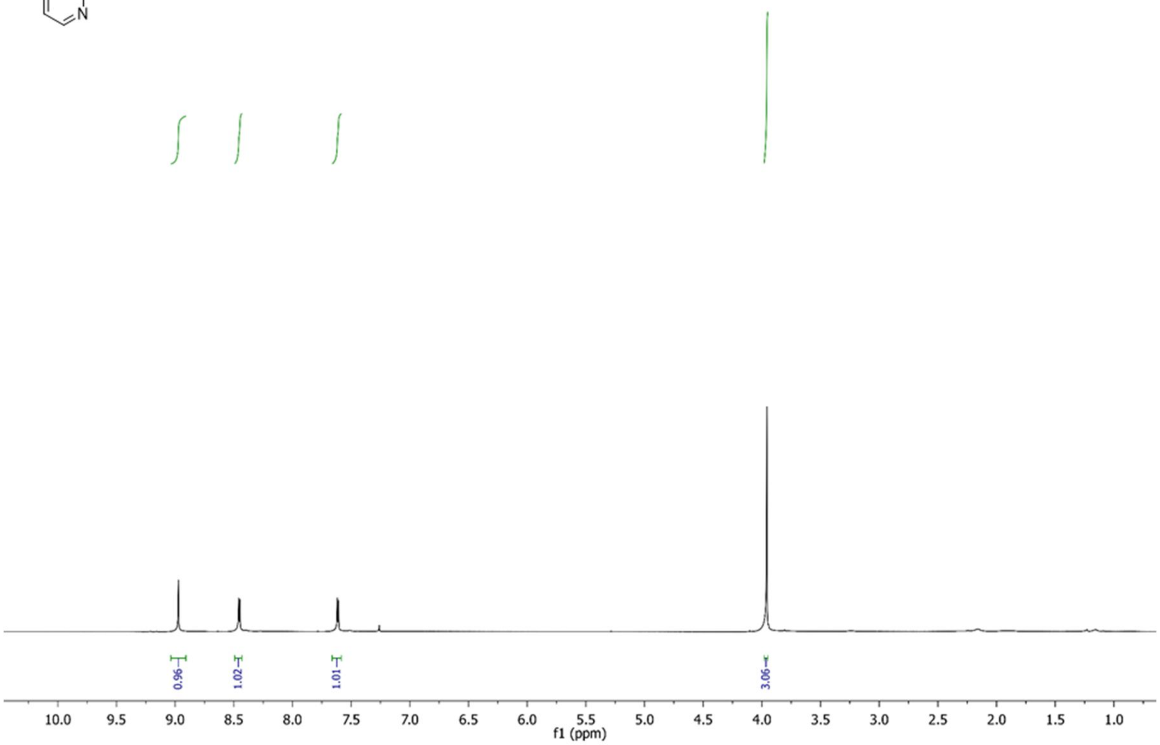
0.96
1.02
1.01
3.06
10.0 9.5 9.0 8.5 8.0 7.5 7.0 6.5 6.0 5.5 5.0 4.5 4.0 3.5 3.0 2.5 2.0 1.5 1.0
f1 (ppm)

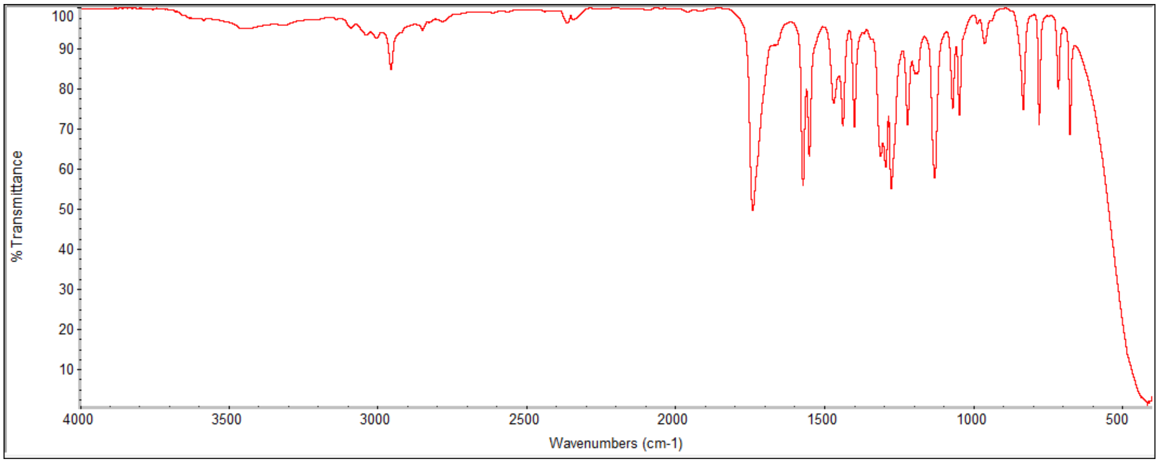
% Transmittance
100 90 80 70 60 50 40 30 20 10
4000 3500 3000 2500 2000 1500 1000 500
Wavenumbers (cm-1)

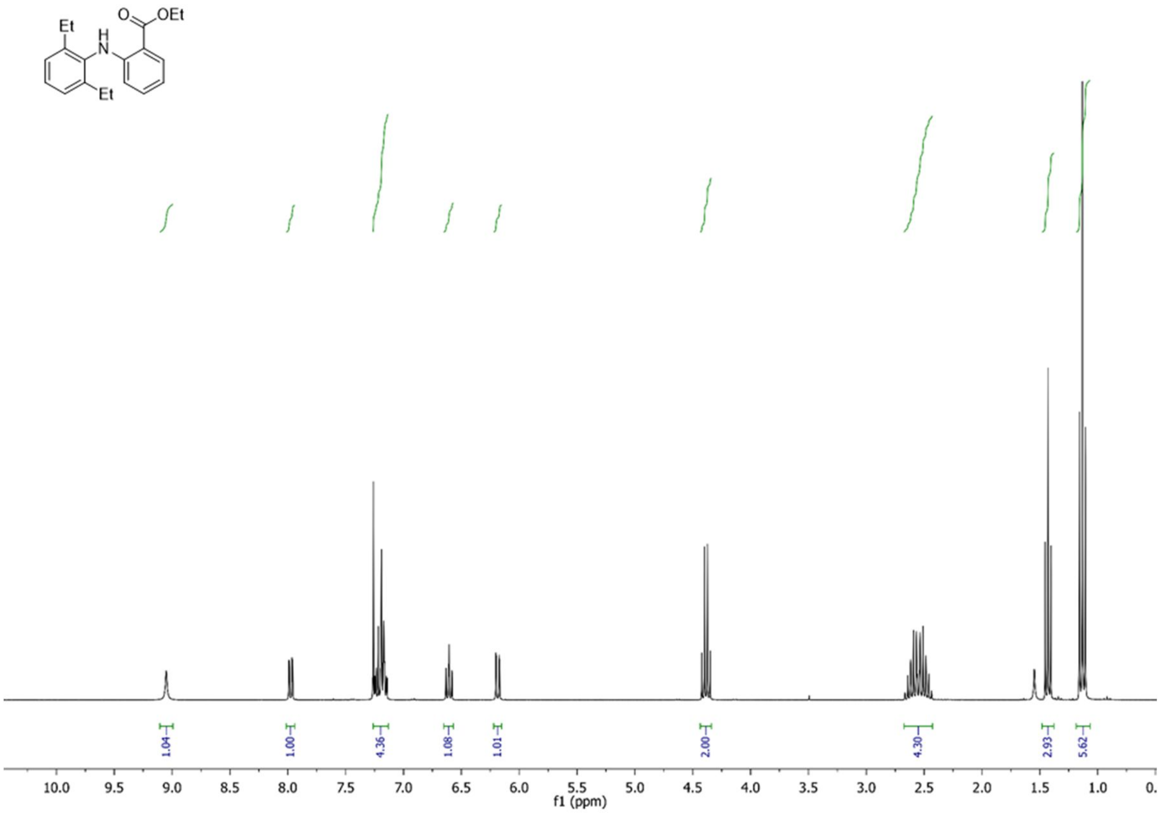
Et
H
N
O
OEt
Et
1.04
1.00
4.36
1.08
1.01
2.00
4.30
2.93
5.62
10.0
9.5
9.0
8.5
8.0
7.5
7.0
6.5
6.0
5.5
5.0
4.5
4.0
3.5
3.0
2.5
2.0
1.5
1.0
f1 (ppm)

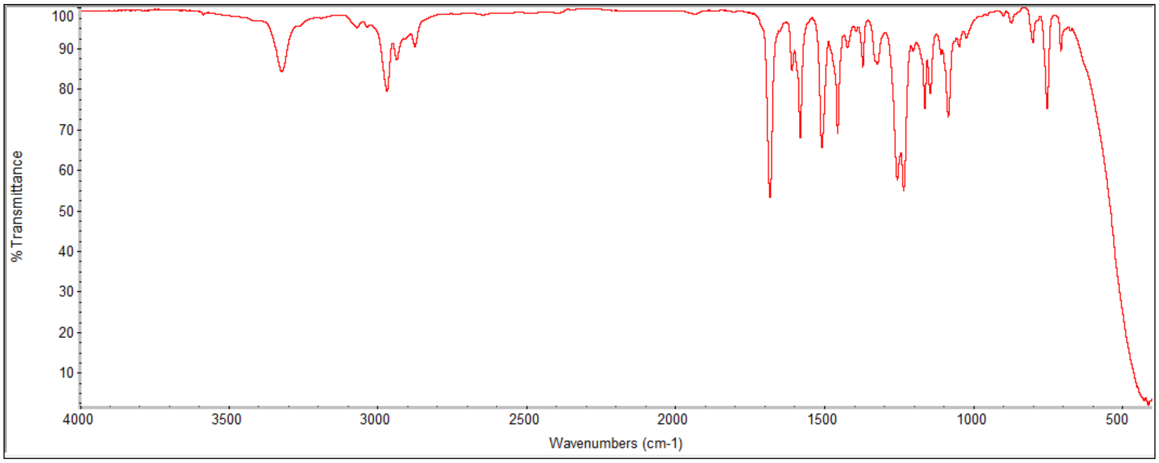
% Transmittance
100
90
80
70
60
50
40
30
20
10
4000
3500
3000
2500
2000
1500
1000
500
Wavenumbers (cm-1)

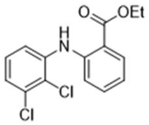

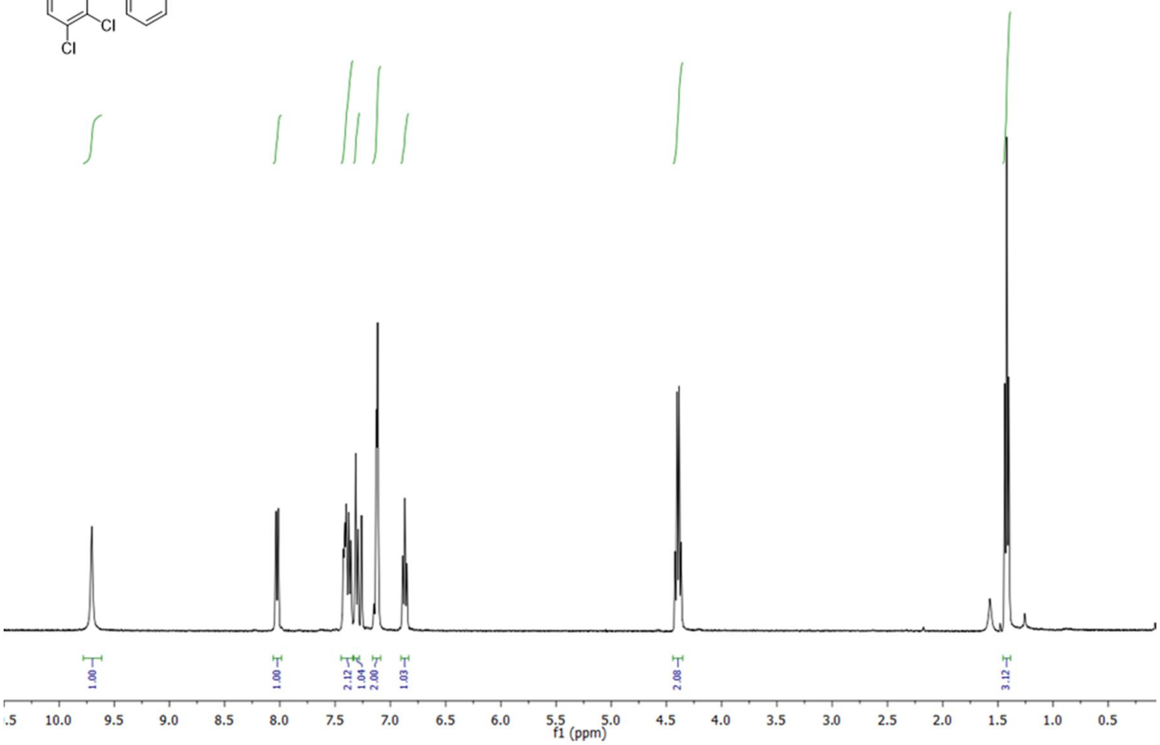

1.00
1.00
2.12
1.04
2.00
1.03
2.08
3.12
f1 (ppm)

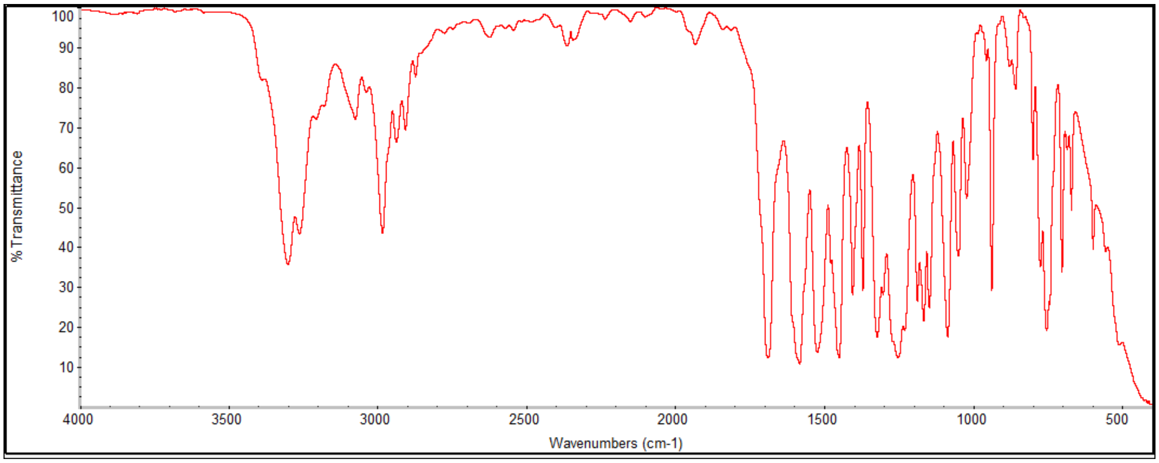

% Transmittance
Wavenumbers (cm-1)

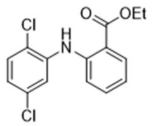
O
OEt
Cl
H
N
Cl

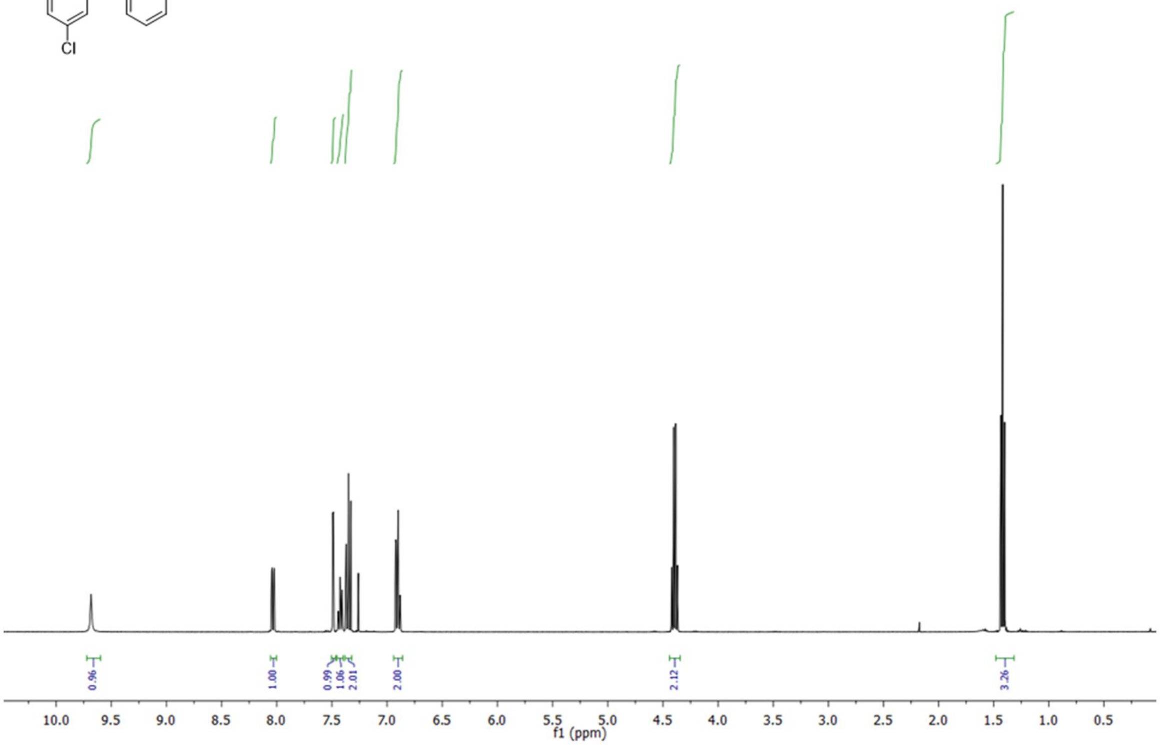
0.96
1.00
0.99
1.06
2.01
2.00
2.12
3.26
10.0 9.5 9.0 8.5 8.0 7.5 7.0 6.5 6.0 5.5 5.0 4.5 4.0 3.5 3.0 2.5 2.0 1.5 1.0 0.5
f1 (ppm)

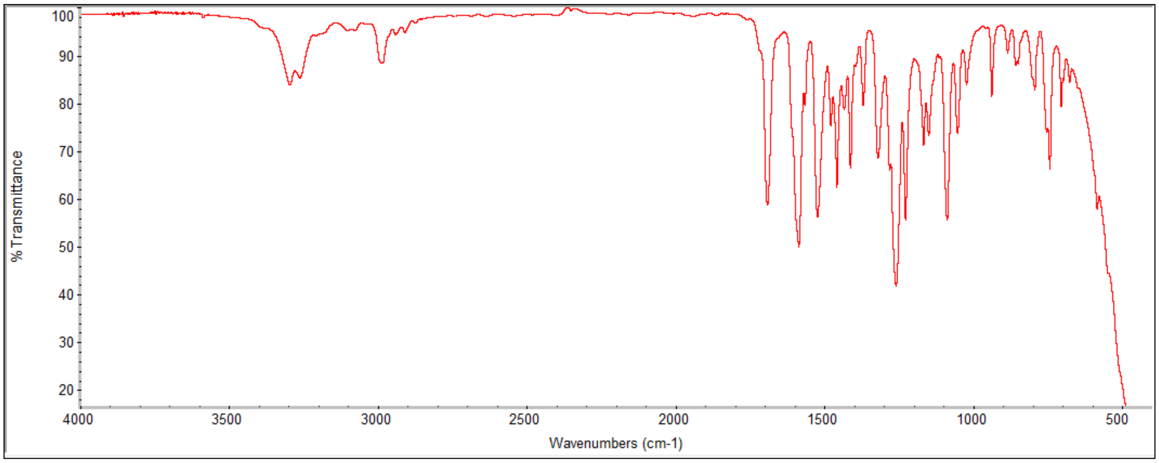
% Transmittance
100
90
80
70
60
50
40
30
20
4000 3500 3000 2500 2000 1500 1000 500
Wavenumbers (cm-1)

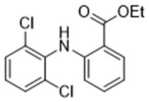
Cl
H
N
O
OEt
Cl

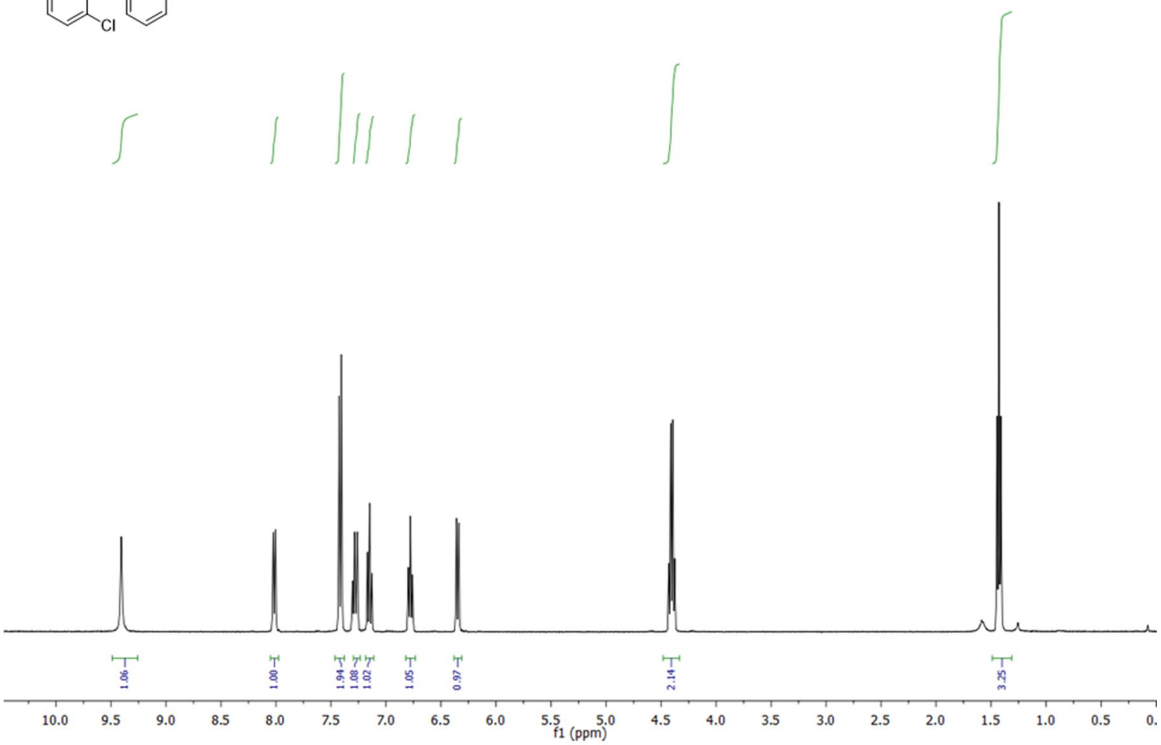
1.06
1.00
1.94
1.08
1.02
1.05
0.97
2.14
3.25
10.0 9.5 9.0 8.5 8.0 7.5 7.0 6.5 6.0 5.5 5.0 4.5 4.0 3.5 3.0 2.5 2.0 1.5 1.0 0.5 0.
f1 (ppm)

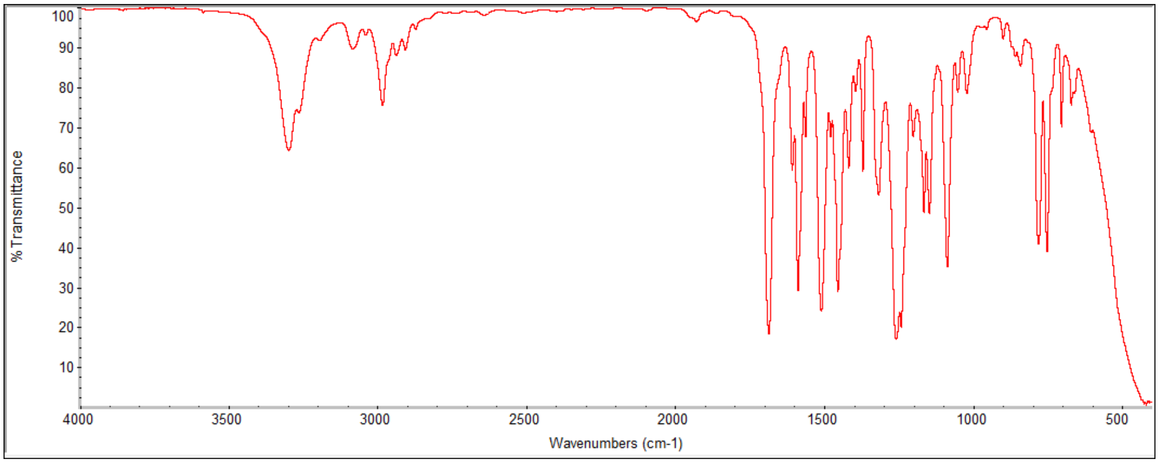
% Transmittance
100
90
80
70
60
50
40
30
20
10
4000 3500 3000 2500 2000 1500 1000 500
Wavenumbers (cm-1)

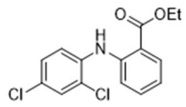

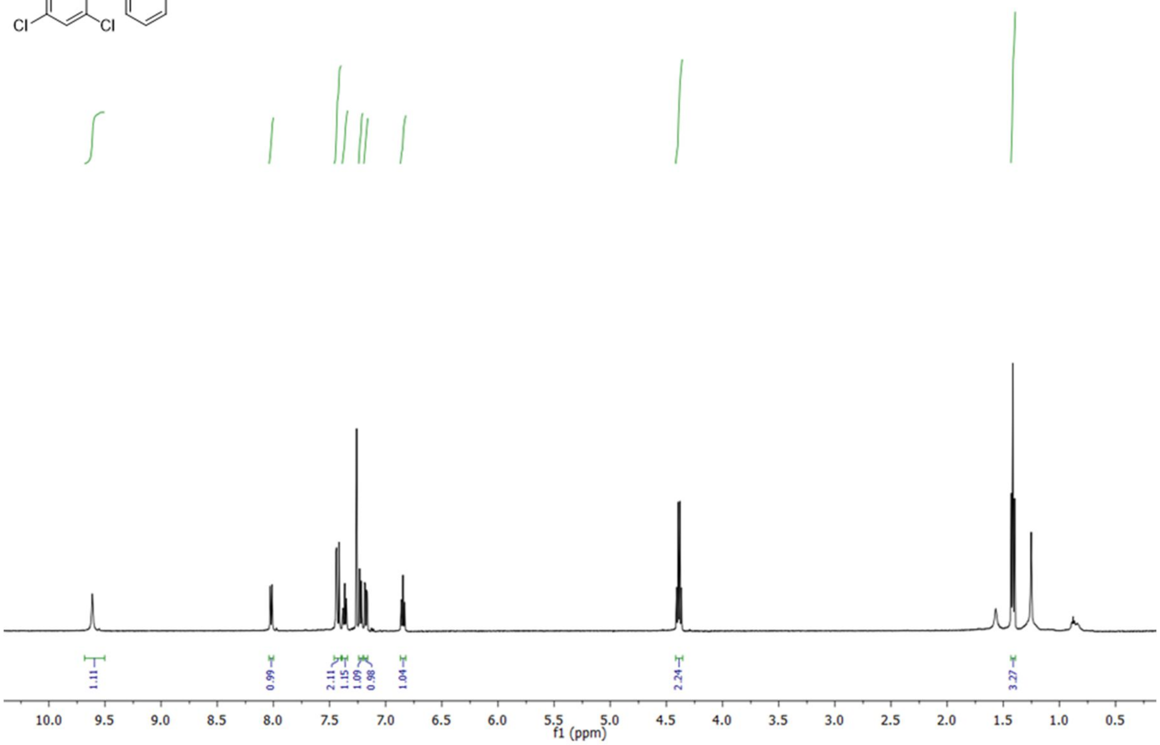

1.11
0.99
2.11
1.15
1.09
0.98
1.04
2.24
3.27
10.0
9.5
9.0
8.5
8.0
7.5
7.0
6.5
6.0
5.5
5.0
4.5
4.0
3.5
3.0
2.5
2.0
1.5
1.0
0.5
f1 (ppm)

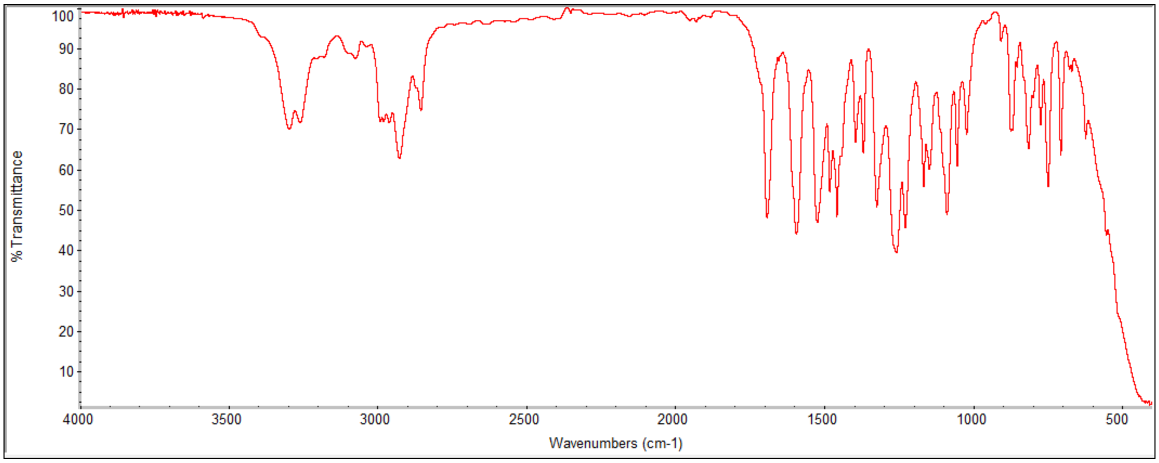

% Transmittance
100
90
80
70
60
50
40
30
20
10
4000
3500
3000
2500
2000
1500
1000
500
Wavenumbers (cm-1)

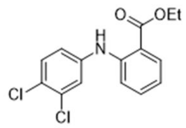
OEt
O
H
N
Cl
Cl

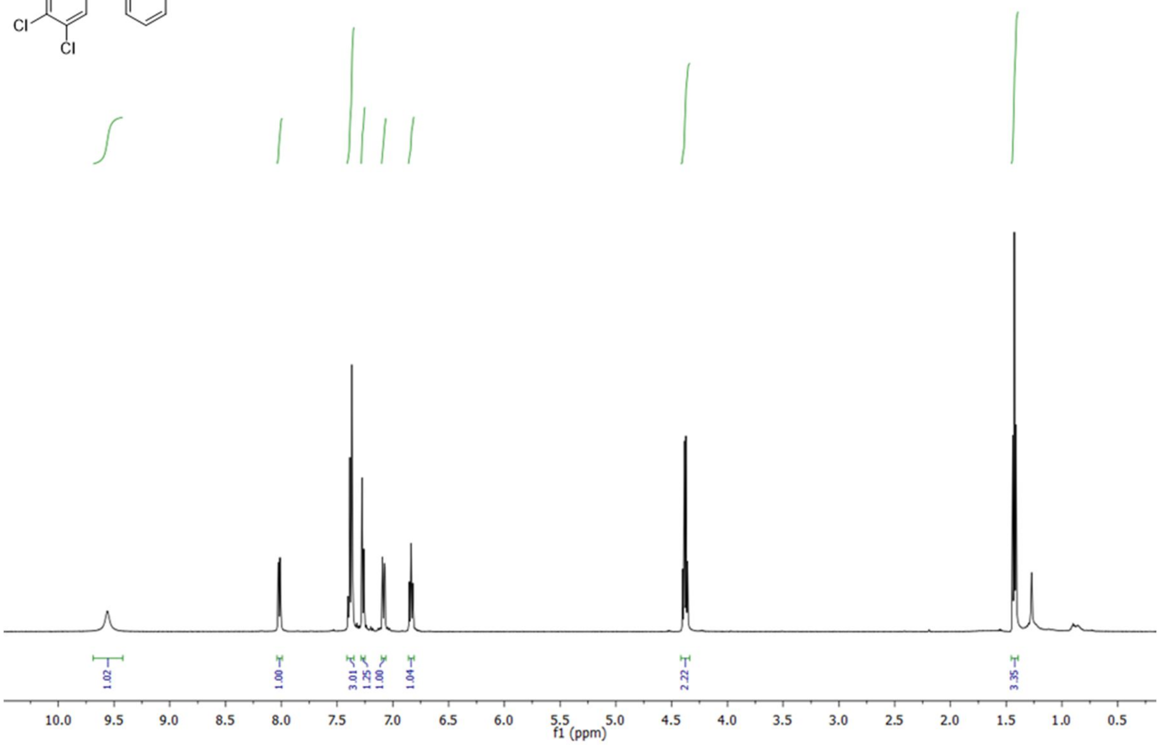
1.02
1.00
3.01
1.25
1.00
1.04
2.22
3.35
10.0
9.5
9.0
8.5
8.0
7.5
7.0
6.5
6.0
5.5
5.0
4.5
4.0
3.5
3.0
2.5
2.0
1.5
1.0
0.5
f1 (ppm)

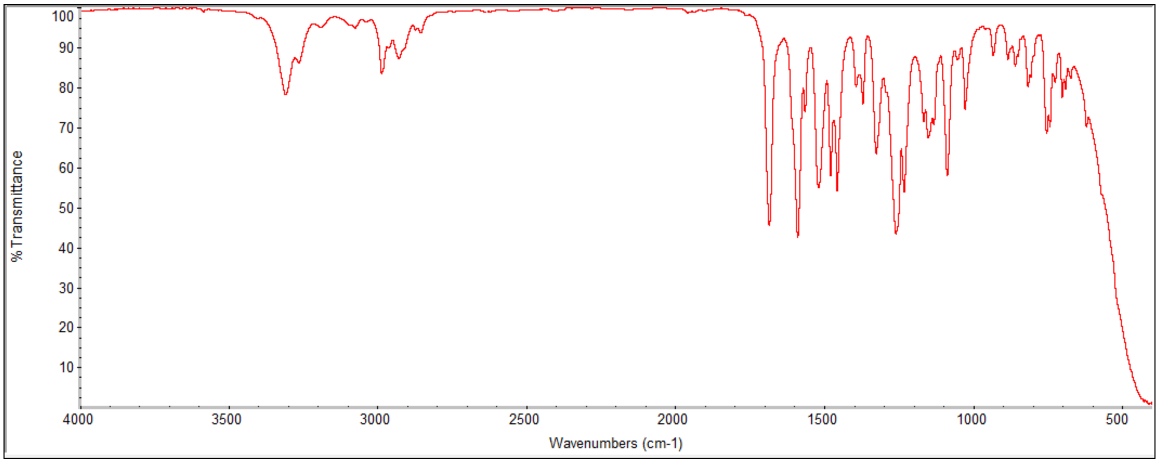
% Transmittance
100
90
80
70
60
50
40
30
20
10
4000
3500
3000
2500
2000
1500
1000
500
Wavenumbers (cm-1)

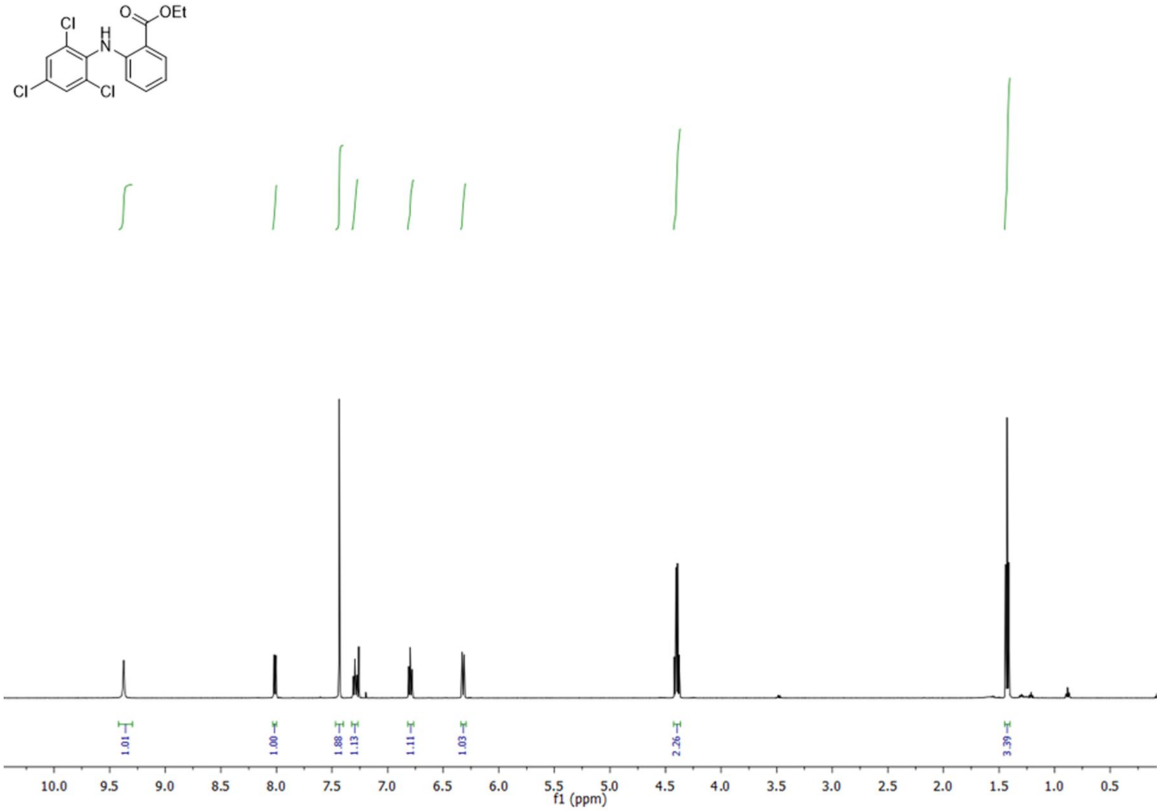

OEt
O
H
N
Cl
Cl
Cl
1.01
1.00
1.88
1.13
1.11
1.03
2.26
3.39
10.0
9.5
9.0
8.5
8.0
7.5
7.0
6.5
6.0
5.5
5.0
4.5
4.0
3.5
3.0
2.5
2.0
1.5
1.0
0.5
f1 (ppm)

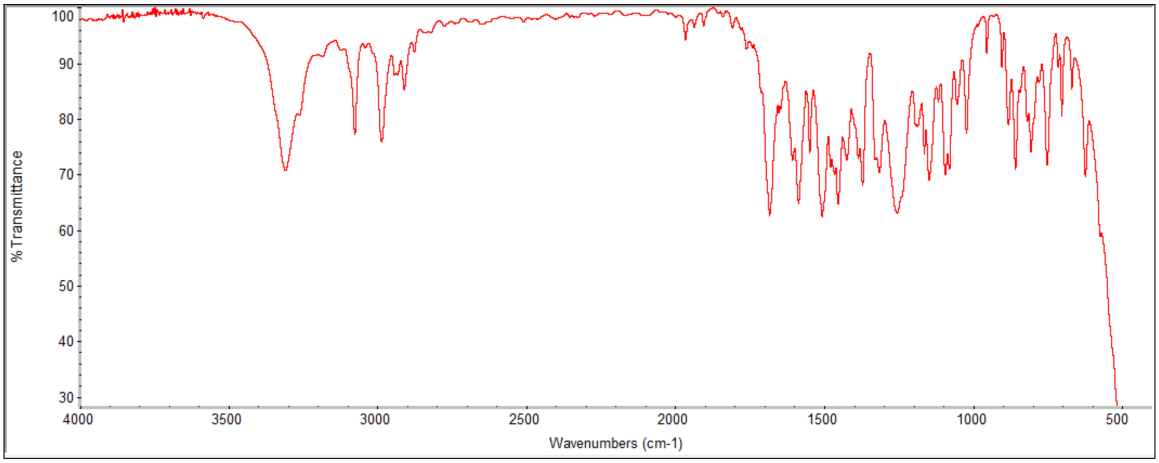

% Transmittance
100
90
80
70
60
50
40
30
4000
3500
3000
2500
2000
1500
1000
500
Wavenumbers (cm-1)

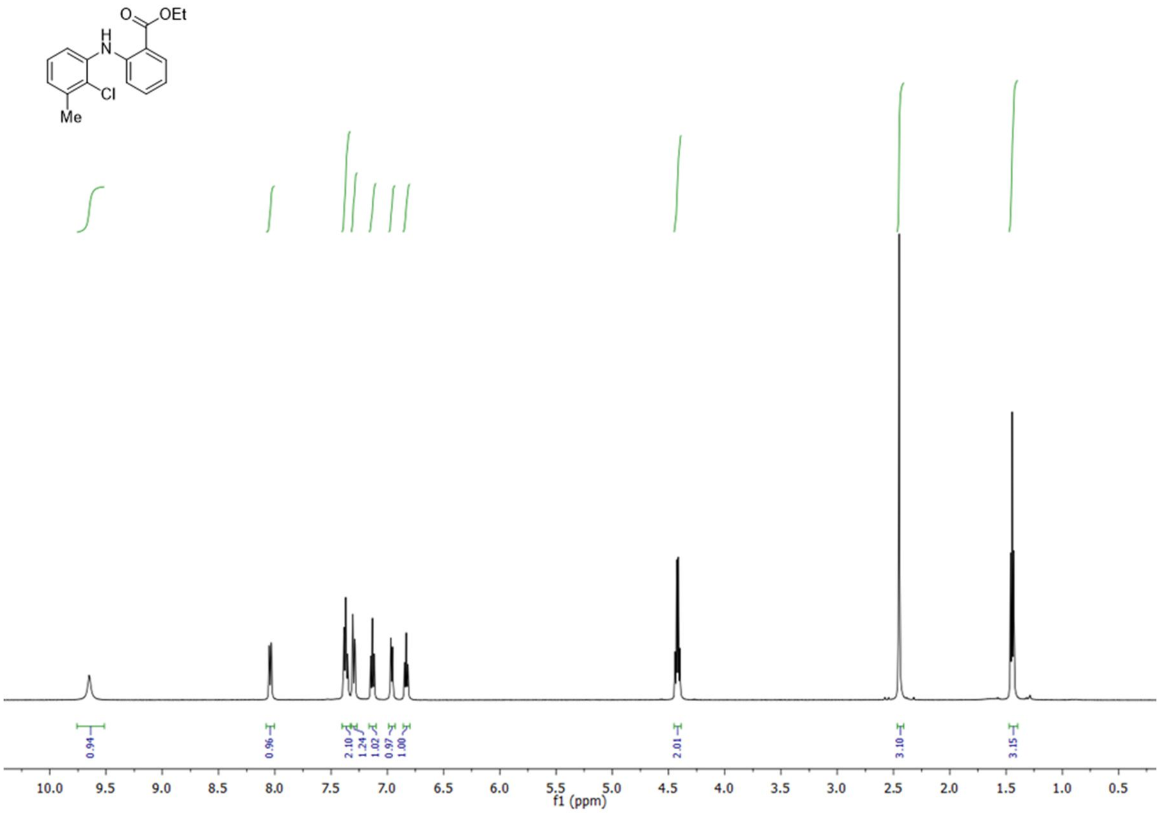
O
OEt
H
N
Cl
Me
0.94
0.96
2.10
1.24
1.02
0.97
1.00
2.01
3.10
3.15
10.0 9.5 9.0 8.5 8.0 7.5 7.0 6.5 6.0 5.5 5.0 4.5 4.0 3.5 3.0 2.5 2.0 1.5 1.0 0.5
f1 (ppm)

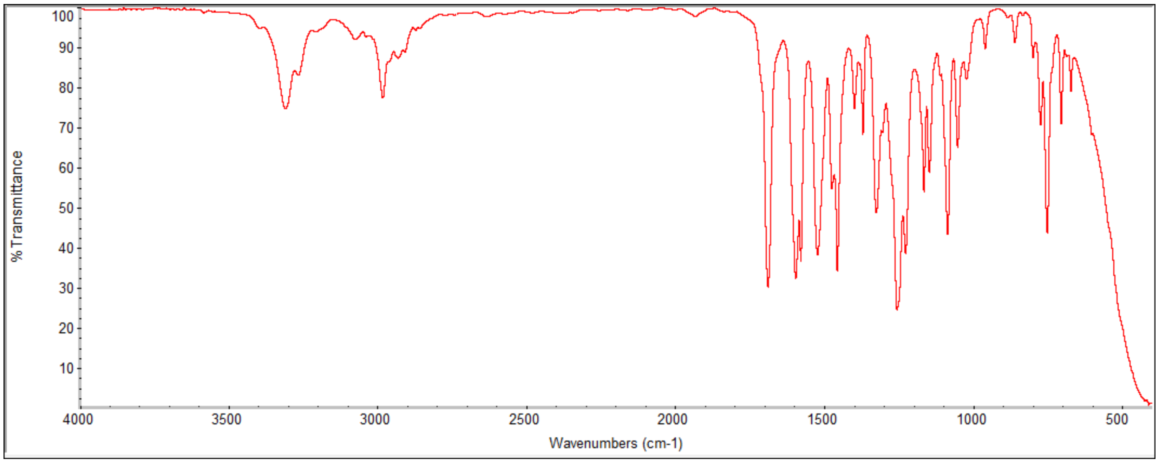
% Transmittance
100 90 80 70 60 50 40 30 20 10
4000 3500 3000 2500 2000 1500 1000 500
Wavenumbers (cm-1)

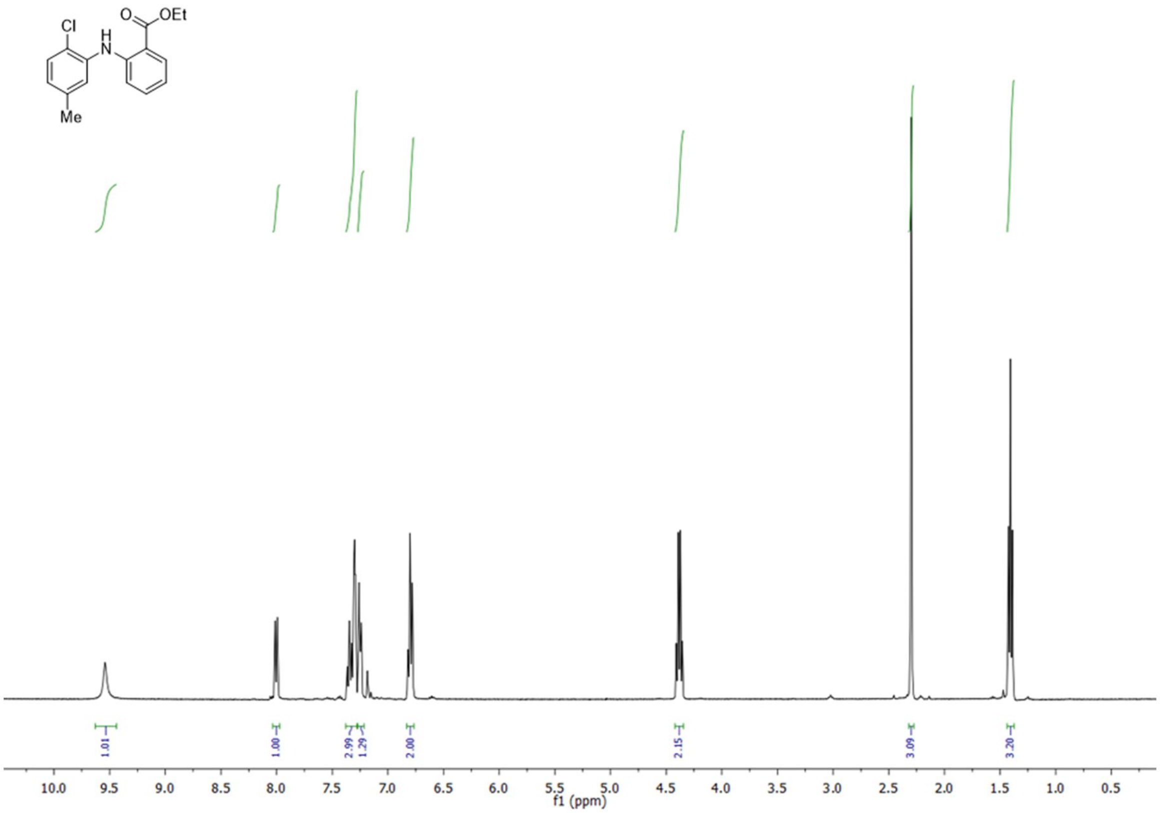
Cl
O
OEt
H
N
Me
1.01
1.00
2.99
1.29
2.00
2.15
3.09
3.20
10.0 9.5 9.0 8.5 8.0 7.5 7.0 6.5 6.0 5.5 5.0 4.5 4.0 3.5 3.0 2.5 2.0 1.5 1.0 0.5
f1 (ppm)

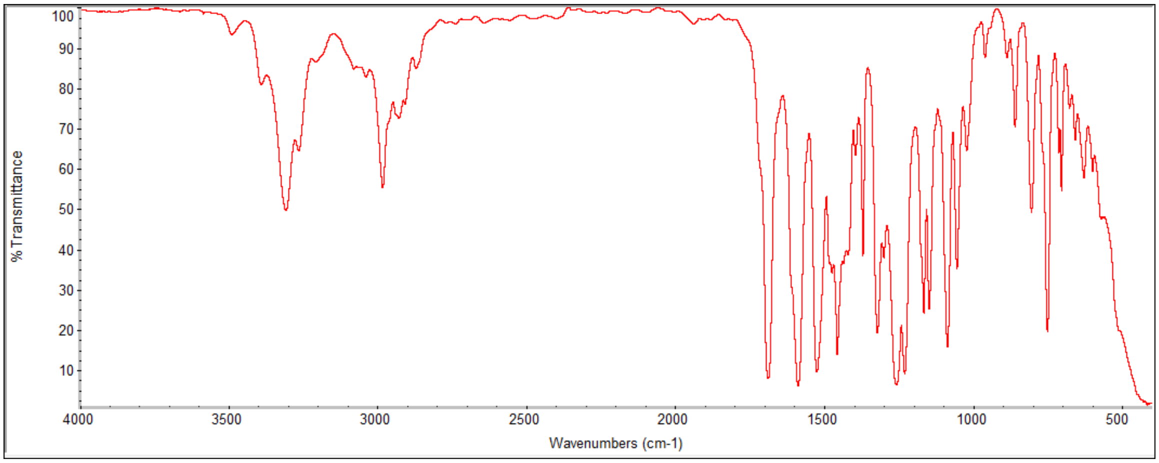
% Transmittance
100 90 80 70 60 50 40 30 20 10
4000 3500 3000 2500 2000 1500 1000 500
Wavenumbers (cm-1)

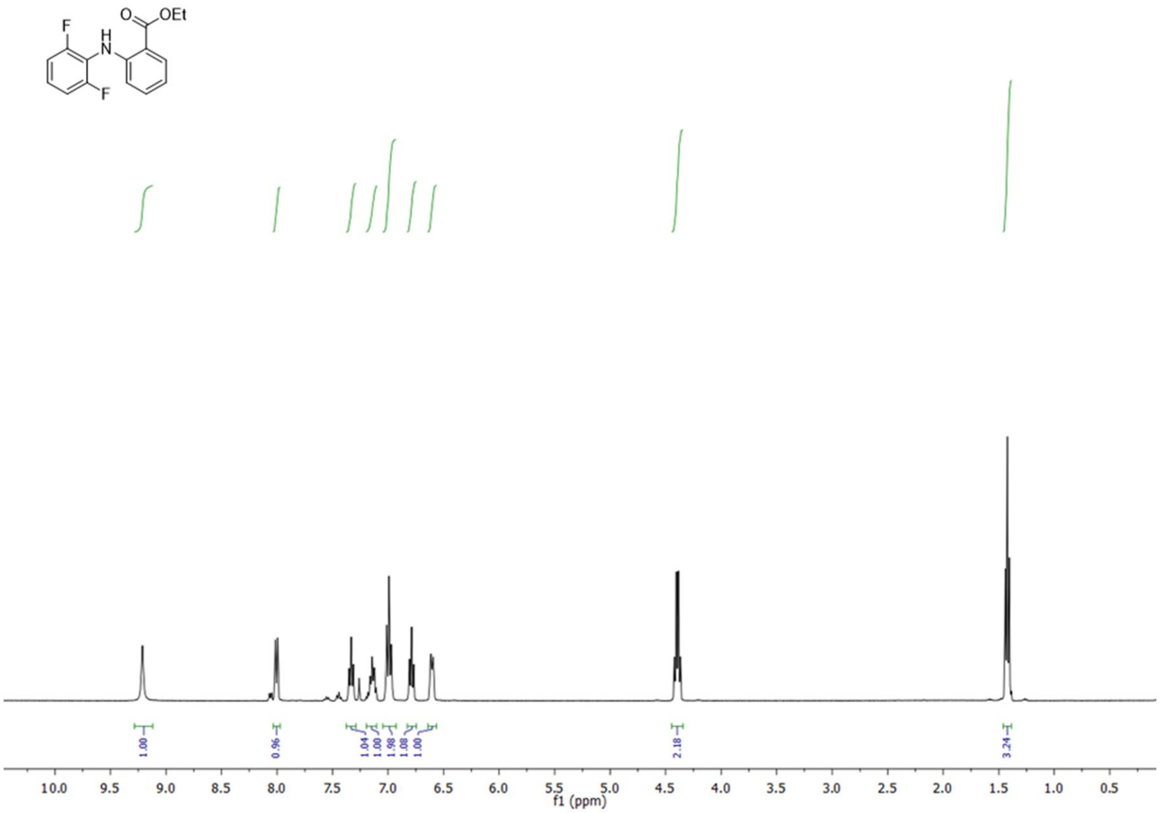
O
OEt
F
H
N
F
1.00
0.96
1.04
1.00
1.98
1.08
1.00
2.18
3.24
10.0 9.5 9.0 8.5 8.0 7.5 7.0 6.5 6.0 5.5 5.0 4.5 4.0 3.5 3.0 2.5 2.0 1.5 1.0 0.5
f1 (ppm)

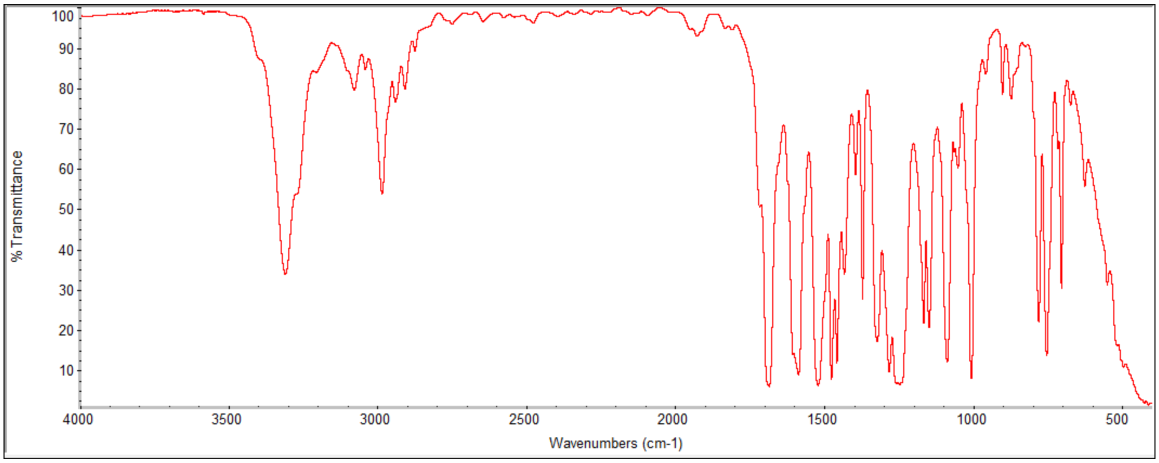
% Transmittance
100
90
80
70
60
50
40
30
20
10
4000 3500 3000 2500 2000 1500 1000 500
Wavenumbers (cm-1)

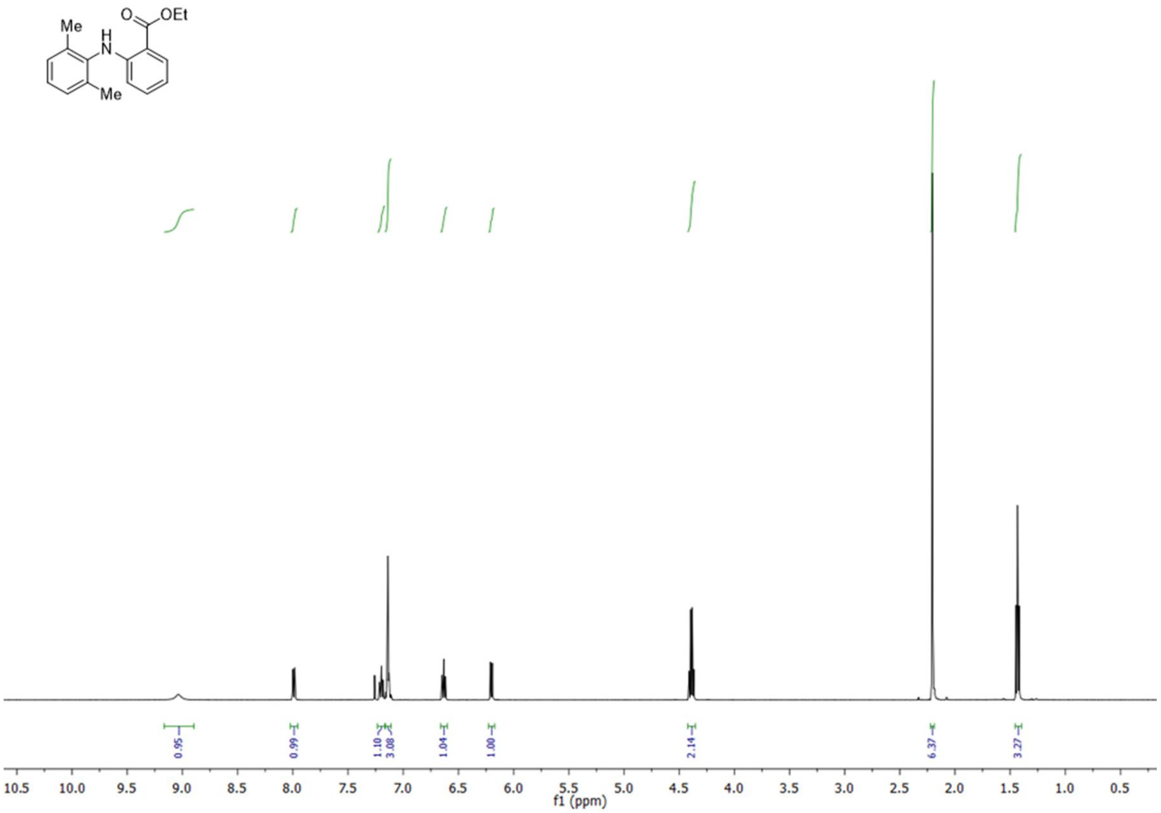
Me
H
N
O
OEt
Me
0.95
0.99
1.10
3.08
1.04
1.00
2.14
6.37
3.27
10.5 10.0 9.5 9.0 8.5 8.0 7.5 7.0 6.5 6.0 5.5 5.0 4.5 4.0 3.5 3.0 2.5 2.0 1.5 1.0 0.5
f1 (ppm)

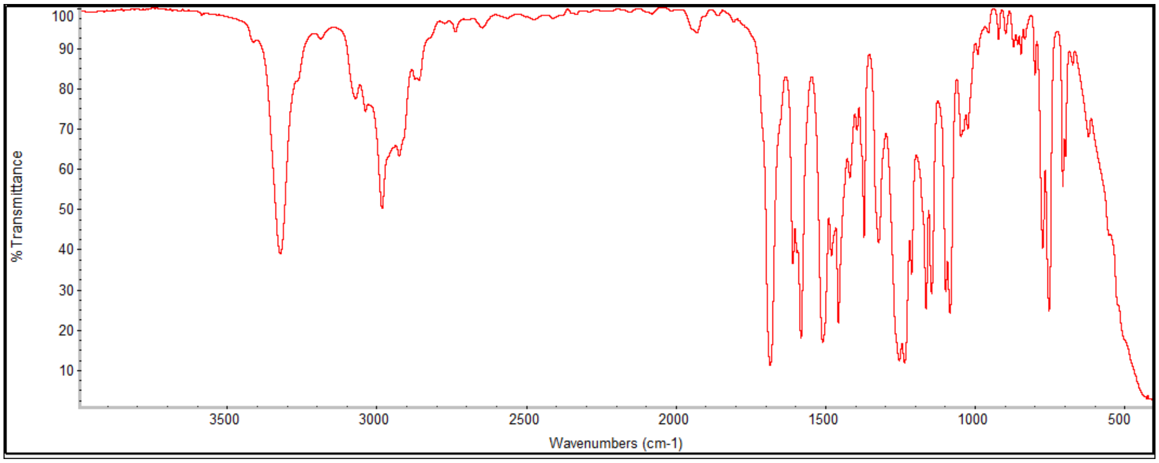
% Transmittance
100 90 80 70 60 50 40 30 20 10
3500 3000 2500 2000 1500 1000 500
Wavenumbers (cm-1)

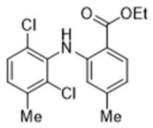
Cl
H
N
O
OEt
Cl
Me
Me

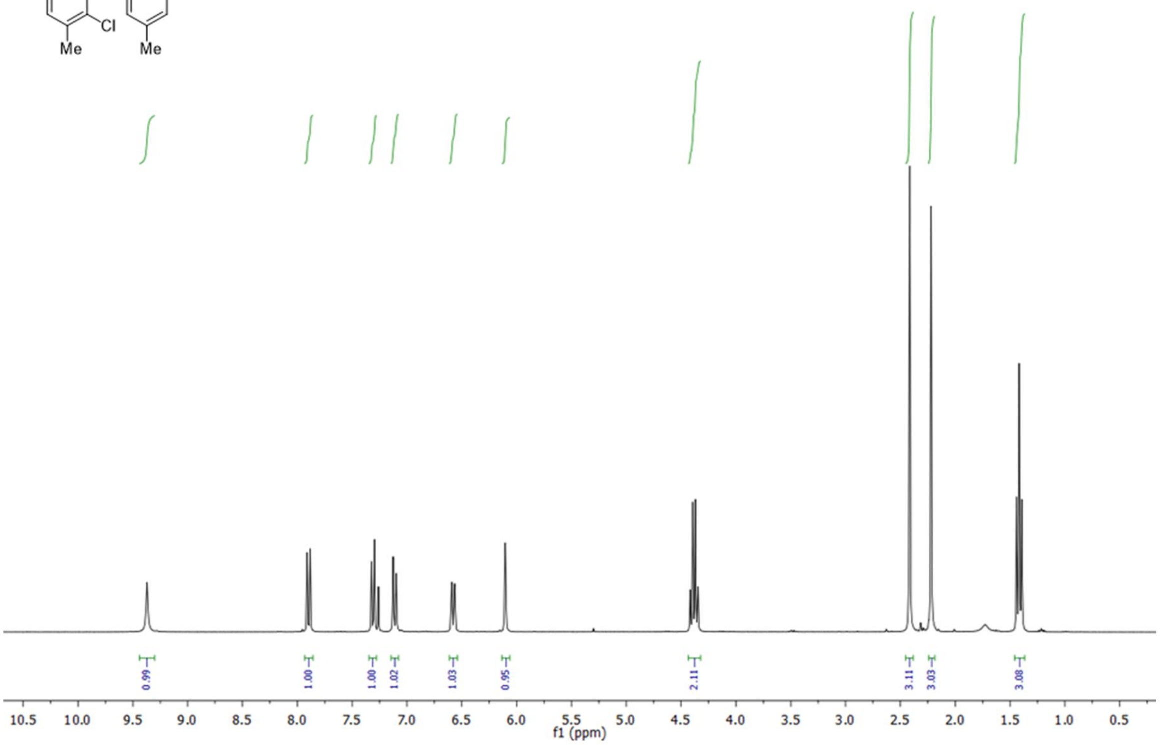
0.99
1.00
1.00
1.02
1.03
0.95
2.11
3.11
3.03
3.08
10.5
10.0
9.5
9.0
8.5
8.0
7.5
7.0
6.5
6.0
5.5
5.0
4.5
4.0
3.5
3.0
2.5
2.0
1.5
1.0
0.5
f1 (ppm)

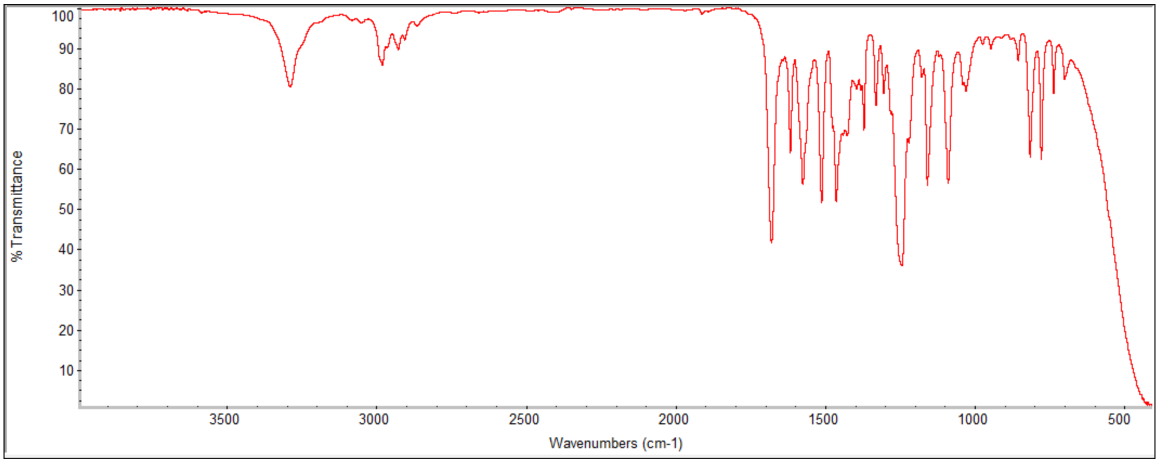
% Transmittance
100
90
80
70
60
50
40
30
20
10
3500
3000
2500
2000
1500
1000
500
Wavenumbers (cm-1)

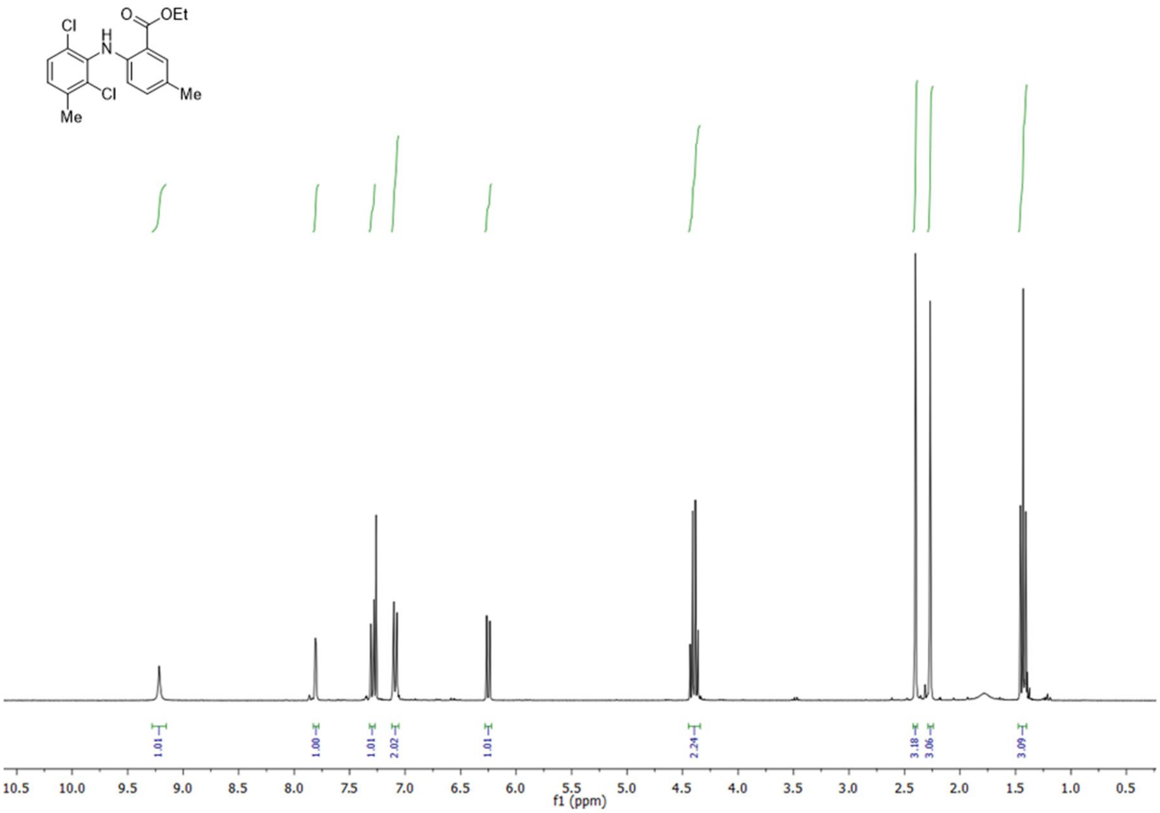

Cl
H
N
O
OEt
Cl
Me
Me
1.01
1.00
1.01
2.02
1.01
2.24
3.18
3.06
3.09
10.5 10.0 9.5 9.0 8.5 8.0 7.5 7.0 6.5 6.0 5.5 5.0 4.5 4.0 3.5 3.0 2.5 2.0 1.5 1.0 0.5
f1 (ppm)

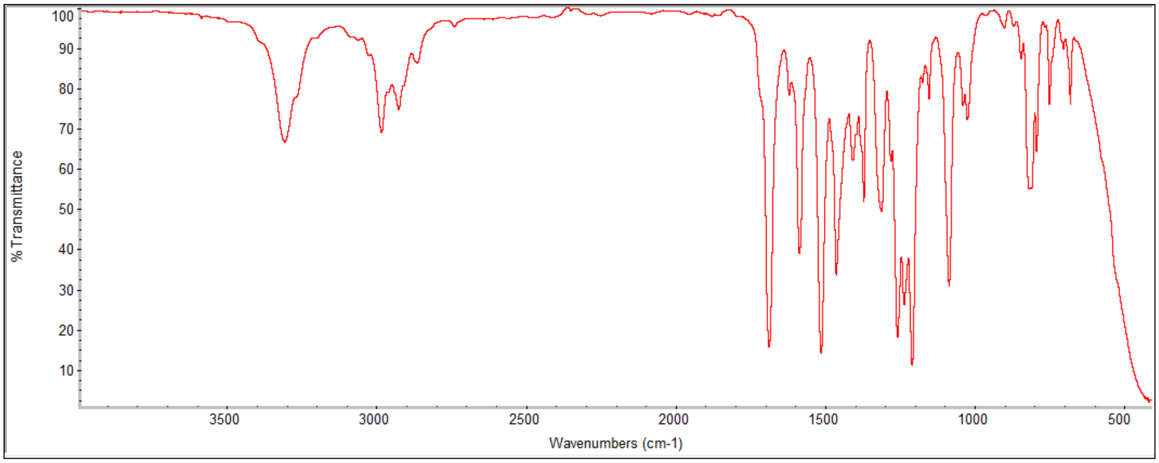

% Transmittance
100
90
80
70
60
50
40
30
20
10
3500 3000 2500 2000 1500 1000 500
Wavenumbers (cm-1)

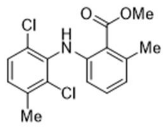
Cl
O
OMe
H
N
Me
Cl
Me

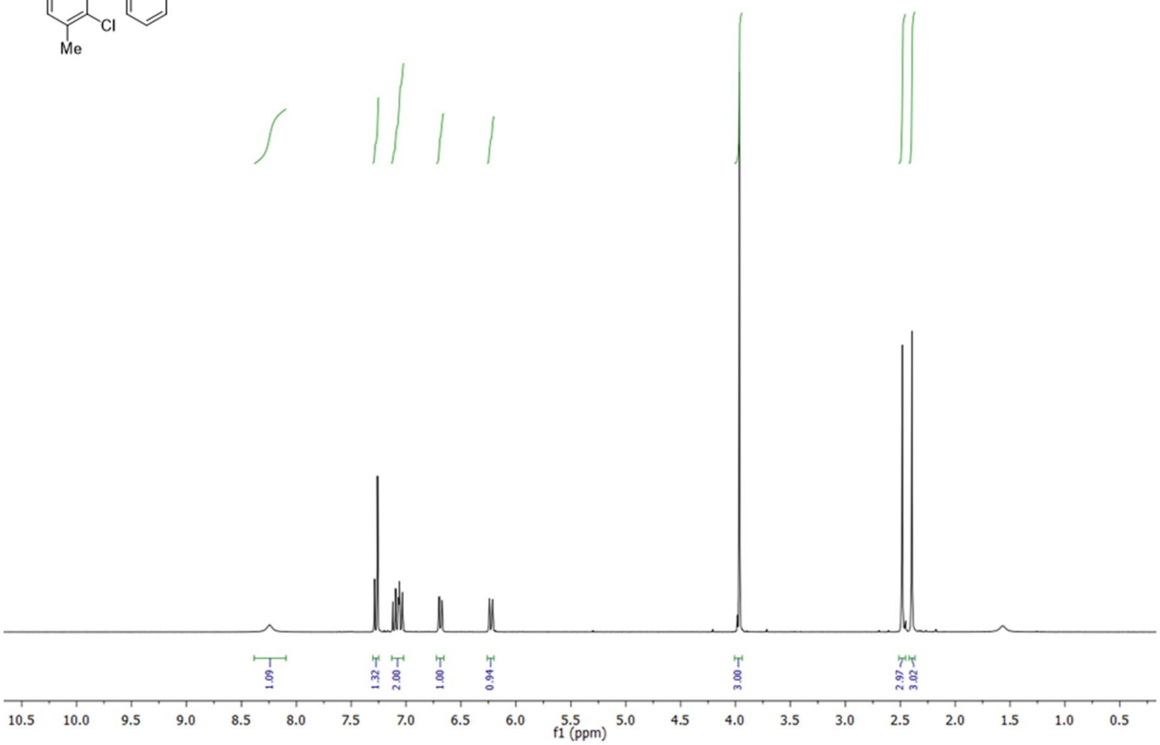
1.09
1.32
2.00
1.00
0.94
3.00
2.97
3.02
10.5 10.0 9.5 9.0 8.5 8.0 7.5 7.0 6.5 6.0 5.5 5.0 4.5 4.0 3.5 3.0 2.5 2.0 1.5 1.0 0.5
f1 (ppm)

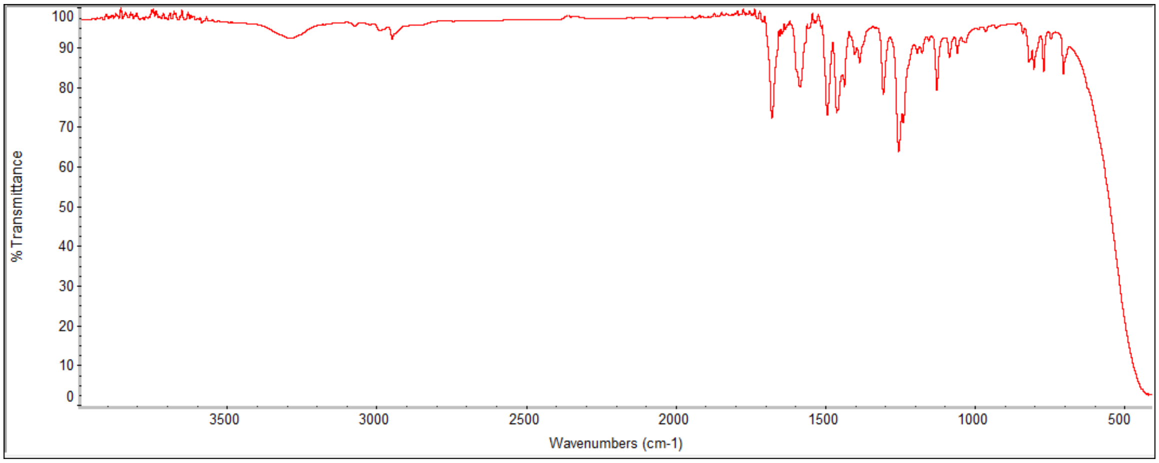
% Transmittance
100 90 80 70 60 50 40 30 20 10 0
3500 3000 2500 2000 1500 1000 500
Wavenumbers (cm-1)

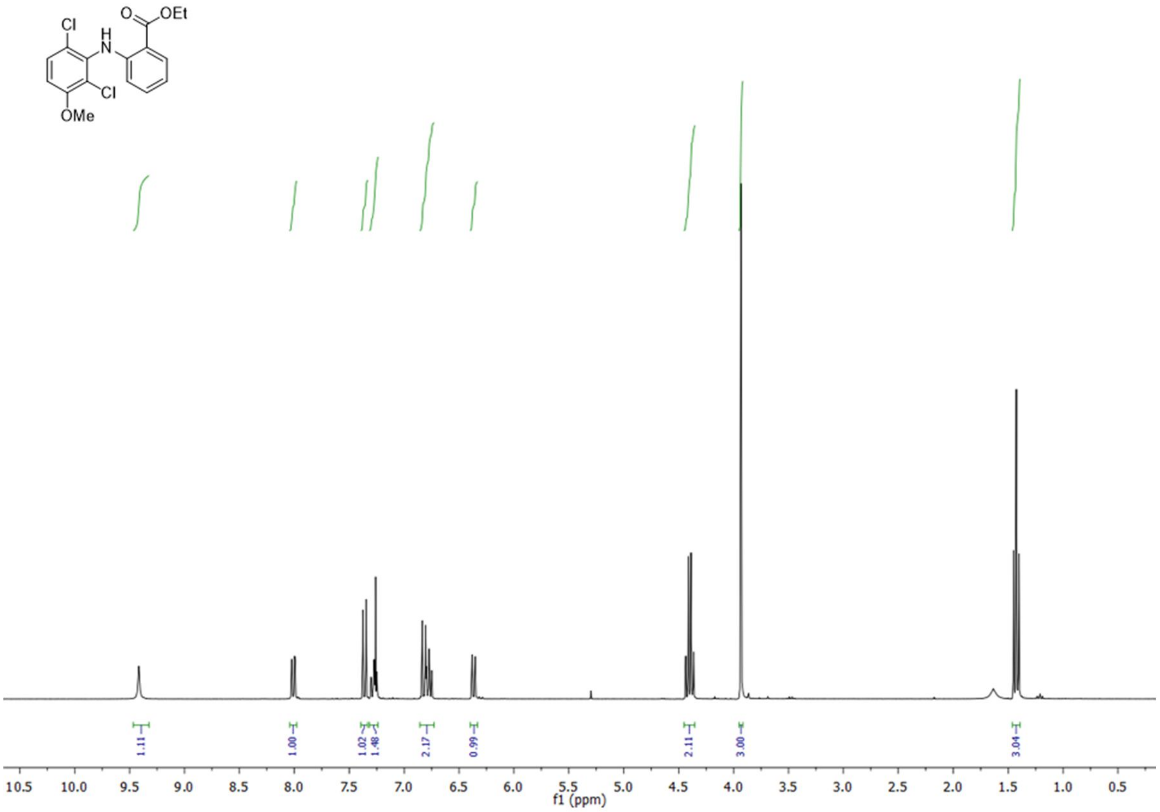
Cl
O
OEt
H
N
Cl
OMe
1.11
1.00
1.02
1.48
2.17
0.99
2.11
3.00
3.04
10.5
10.0
9.5
9.0
8.5
8.0
7.5
7.0
6.5
6.0
5.5
5.0
4.5
4.0
3.5
3.0
2.5
2.0
1.5
1.0
0.5
f1 (ppm)

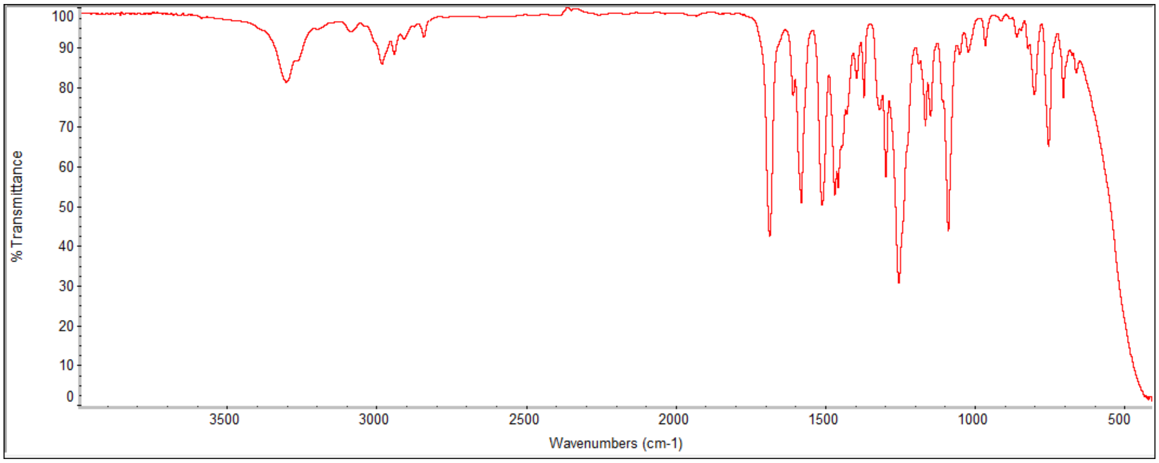
100
90
80
70
60
50
40
30
20
10
0
% Transmittance
3500
3000
2500
2000
1500
1000
500
Wavenumbers (cm-1)

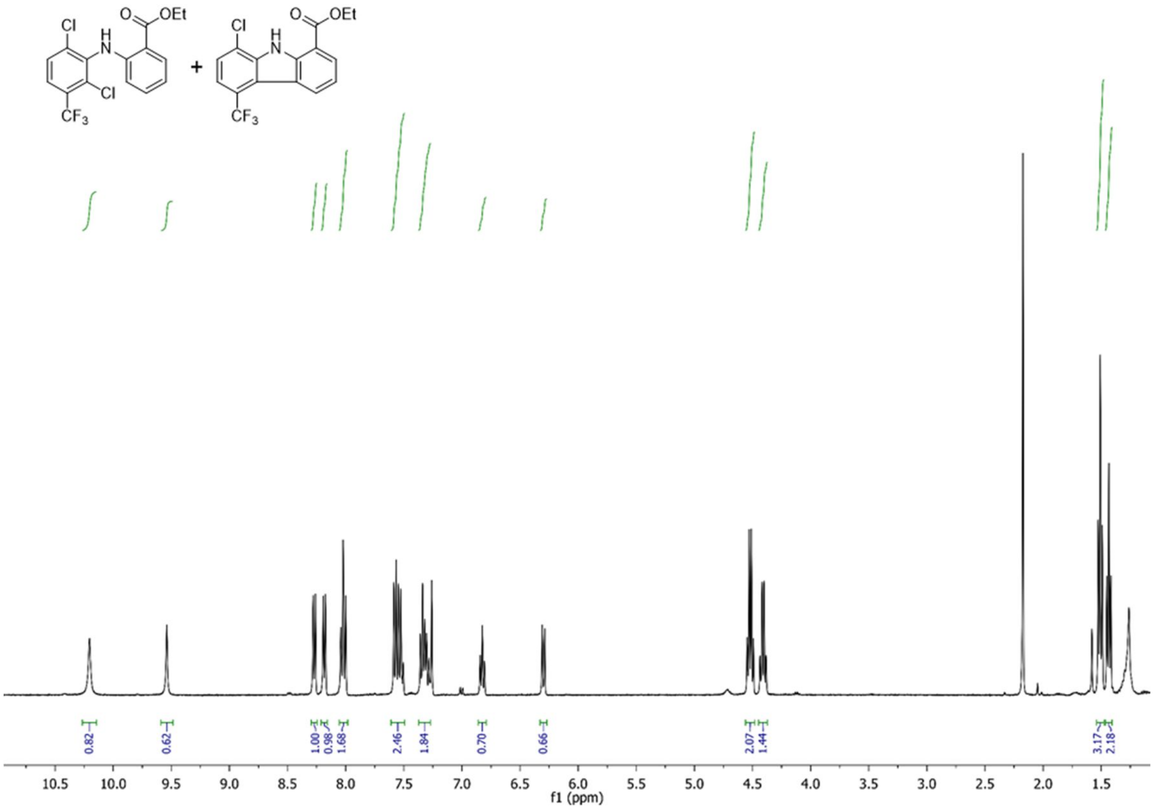
Cl
O
OEt
H
N
Cl
CF3
+
Cl
H
N
O
OEt
CF3
0.82
0.62
1.00
0.98
1.68
2.46
1.84
0.70
0.66
2.07
1.44
3.17
2.18
10.5 10.0 9.5 9.0 8.5 8.0 7.5 7.0 6.5 6.0 5.5 5.0 4.5 4.0 3.5 3.0 2.5 2.0 1.5
f1 (ppm)

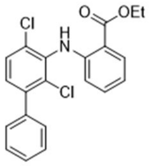

Cl
H
N
O
OEt
Cl

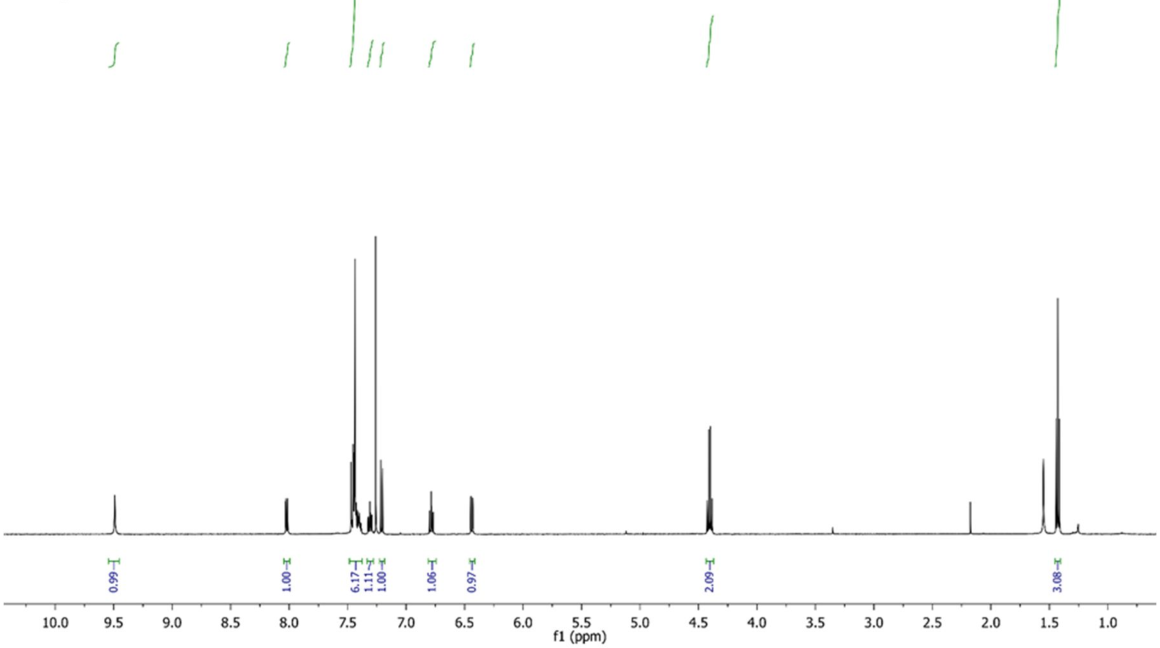

0.99
1.00
6.17
1.11
1.00
1.06
0.97
2.09
3.08
10.0
9.5
9.0
8.5
8.0
7.5
7.0
6.5
6.0
5.5
5.0
4.5
4.0
3.5
3.0
2.5
2.0
1.5
1.0
f1 (ppm)

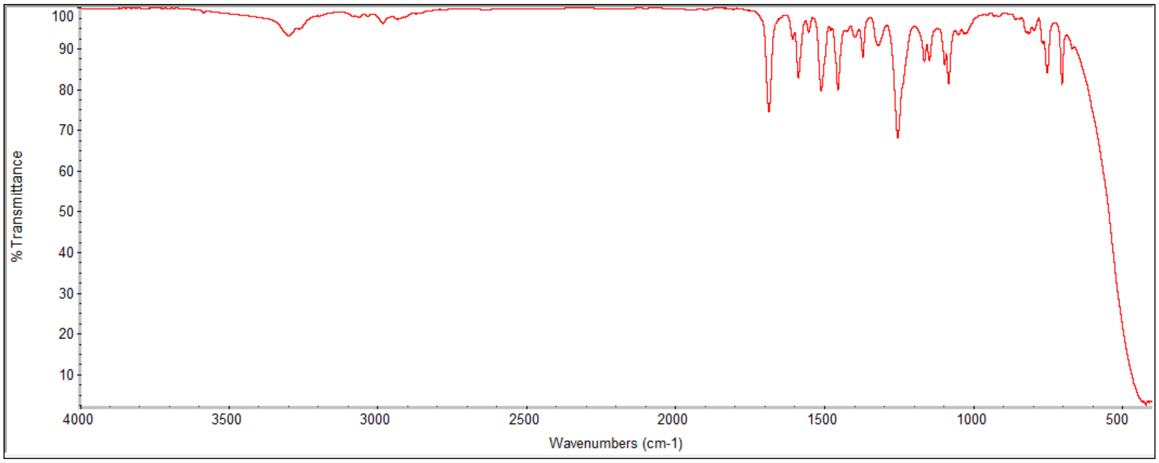

% Transmittance
100
90
80
70
60
50
40
30
20
10
4000
3500
3000
2500
2000
1500
1000
500
Wavenumbers (cm-1)

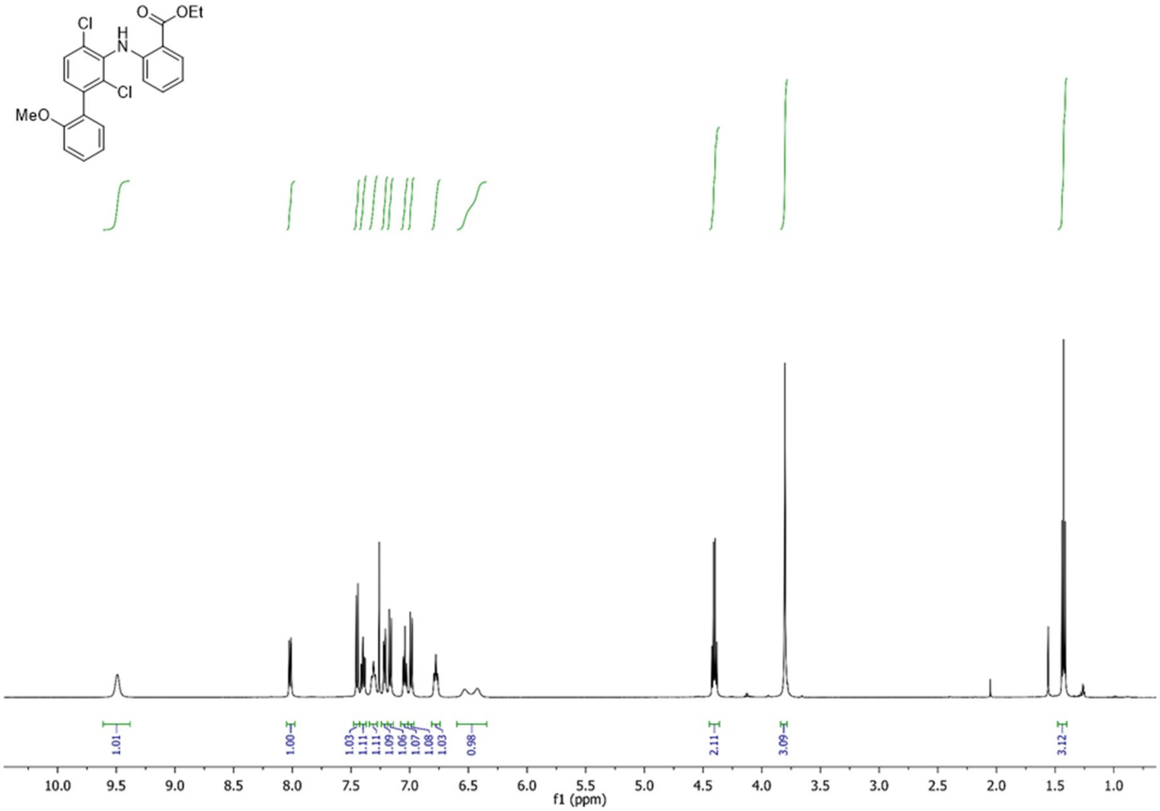
Cl
H
N
O
OEt
Cl
MeO
1.01
1.00
1.03
1.11
1.11
1.09
1.06
1.07
1.08
1.03
0.98
2.11
3.09
3.12
10.0
9.5
9.0
8.5
8.0
7.5
7.0
6.5
6.0
5.5
5.0
4.5
4.0
3.5
3.0
2.5
2.0
1.5
1.0
f1 (ppm)

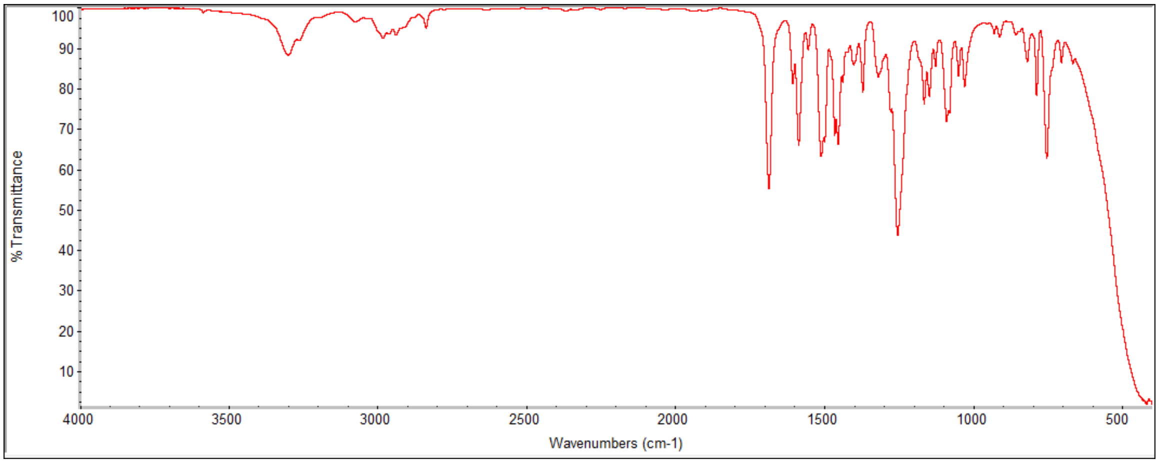
% Transmittance
100
90
80
70
60
50
40
30
20
10
4000
3500
3000
2500
2000
1500
1000
500
Wavenumbers (cm-1)

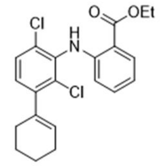
Cl
H
N
O
OEt
Cl

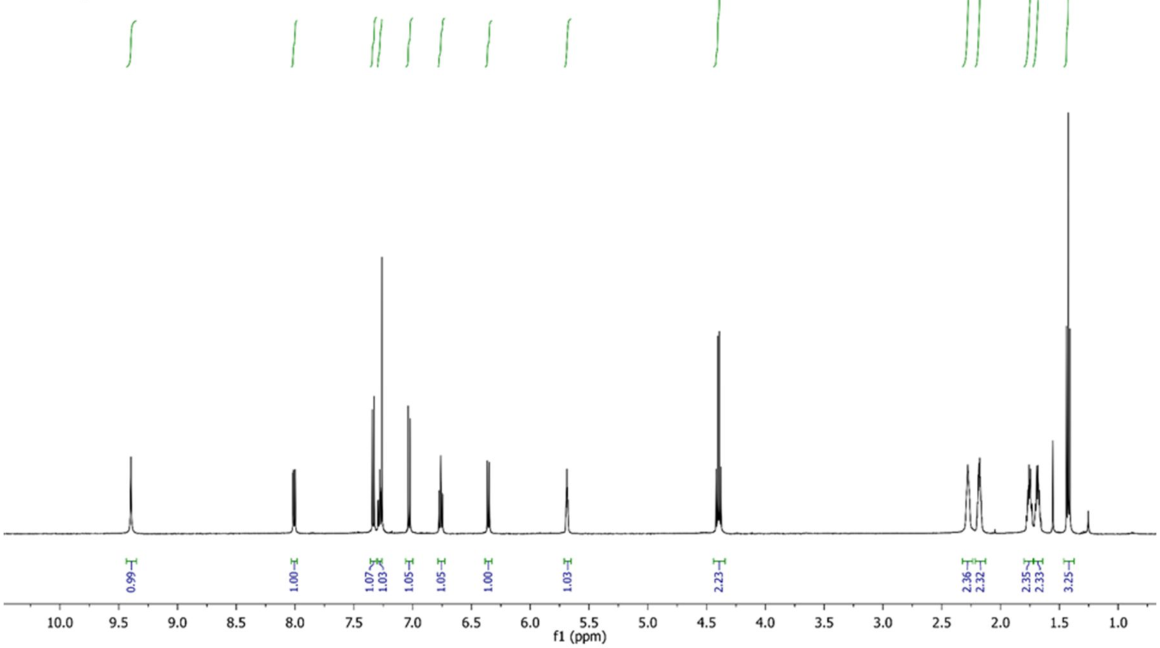
0.99
1.00
1.07
1.03
1.05
1.05
1.00
1.03
2.23
2.36
2.32
2.35
2.33
3.25
10.0
9.5
9.0
8.5
8.0
7.5
7.0
6.5
6.0
5.5
5.0
4.5
4.0
3.5
3.0
2.5
2.0
1.5
1.0
f1 (ppm)

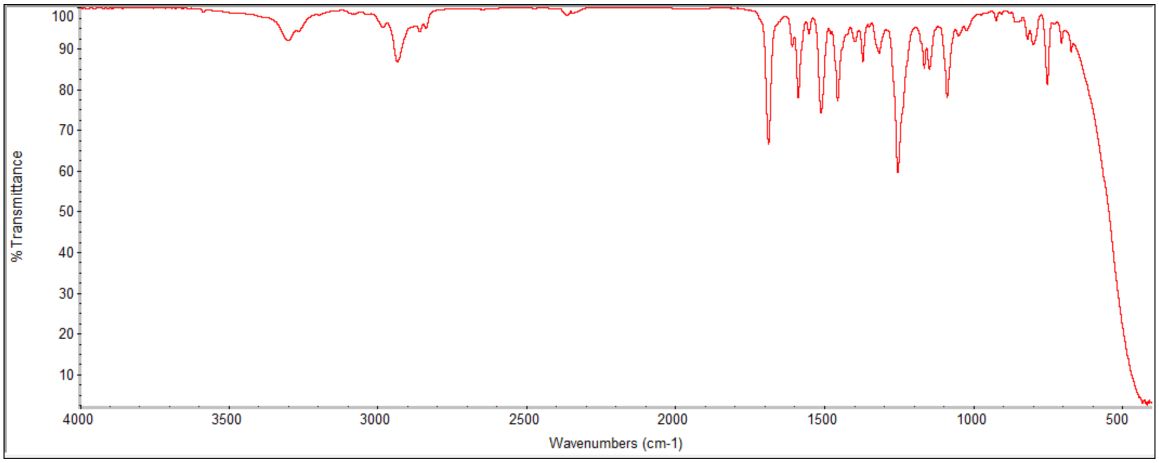
100
90
80
70
60
50
40
30
20
10
% Transmittance
4000
3500
3000
2500
2000
1500
1000
500
Wavenumbers (cm-1)

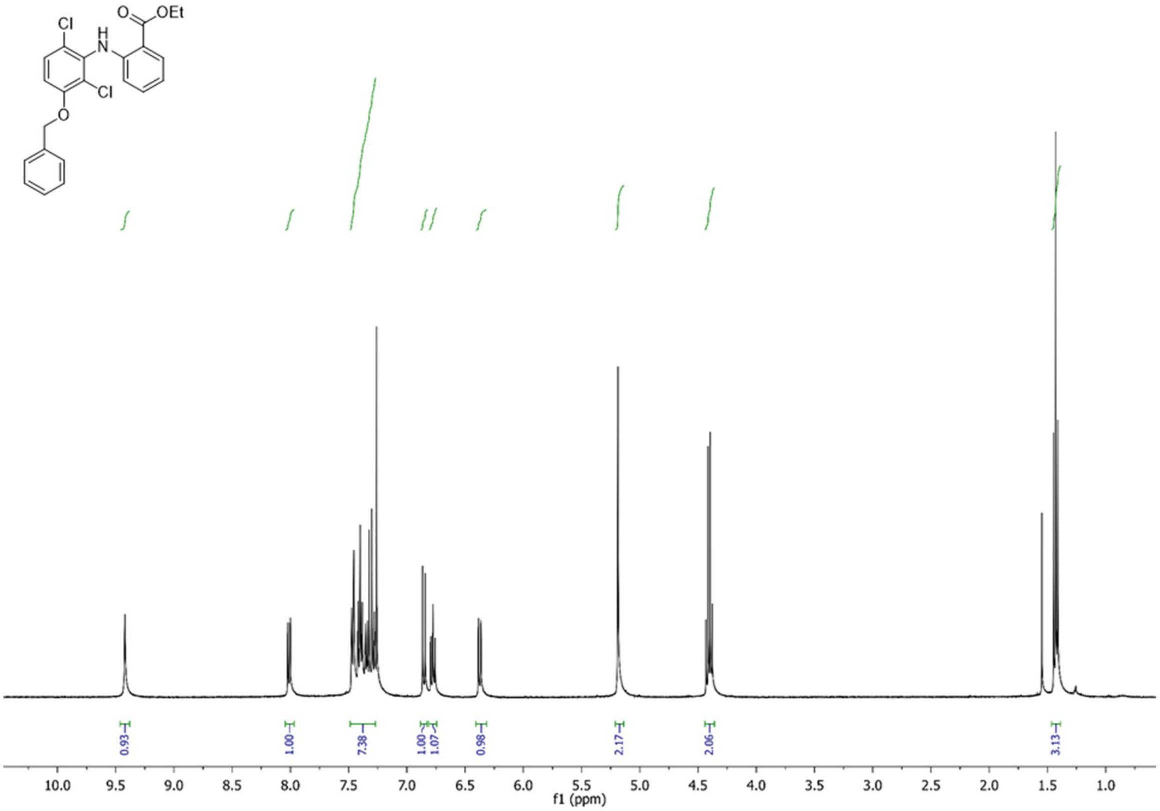
OEt
Cl
H
N
O
Cl
O
0.93
1.00
7.38
1.00
1.07
0.98
2.17
2.06
3.13
10.0
9.5
9.0
8.5
8.0
7.5
7.0
6.5
6.0
5.5
5.0
4.5
4.0
3.5
3.0
2.5
2.0
1.5
1.0
f1 (ppm)

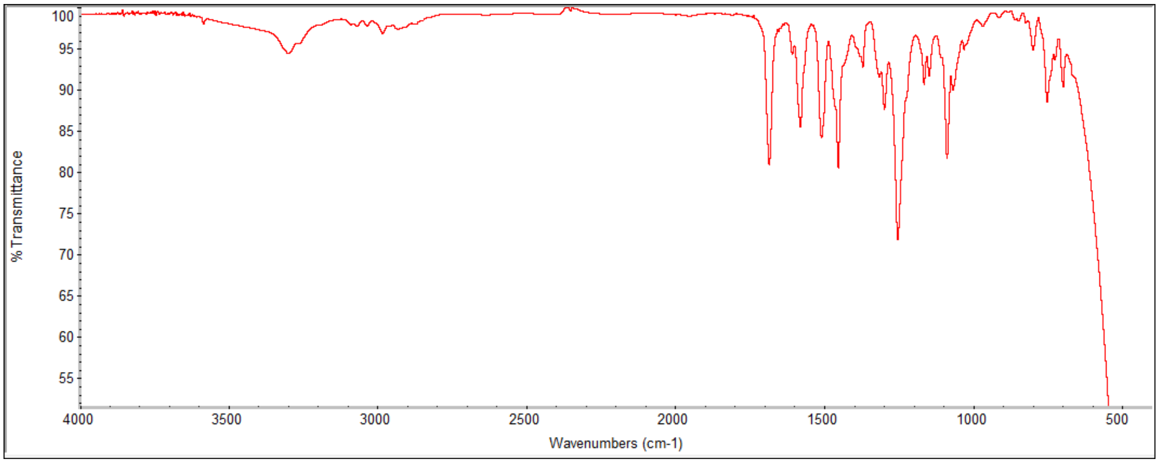
% Transmittance
100
95
90
85
80
75
70
65
60
55
4000
3500
3000
2500
2000
1500
1000
500
Wavenumbers (cm-1)

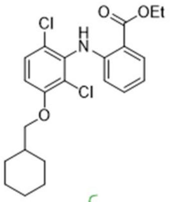

O
OEt
Cl
H
N
Cl
O

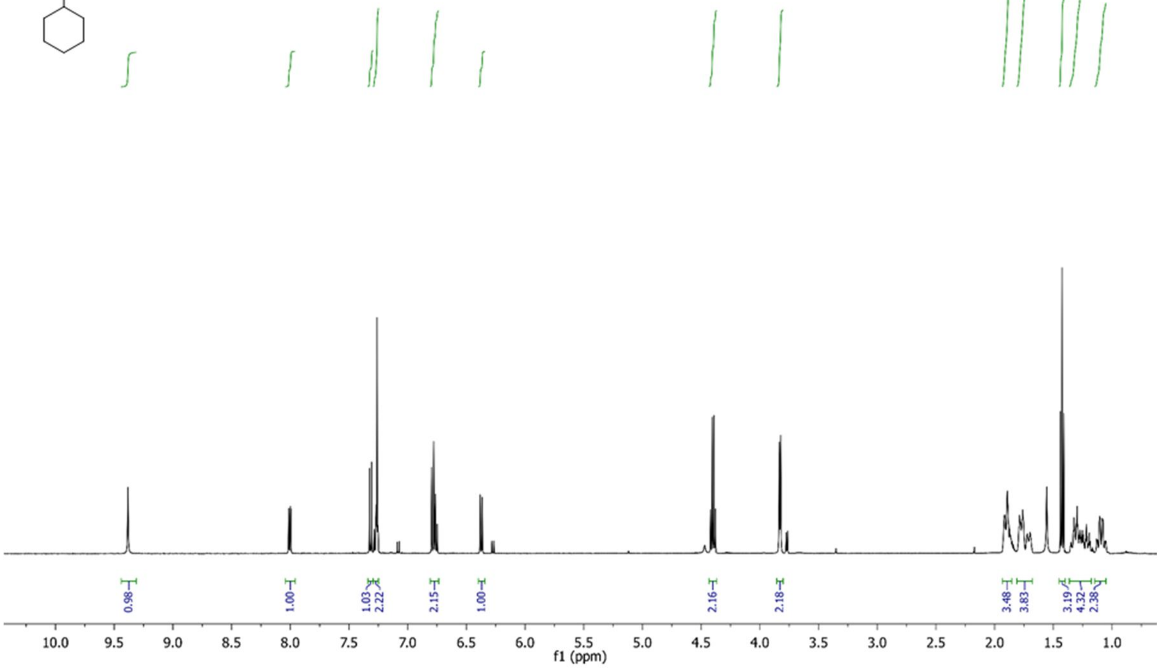

0.98
1.00
1.03
2.22
2.15
1.00
2.16
2.18
3.48
3.83
3.19
4.32
2.38
10.0 9.5 9.0 8.5 8.0 7.5 7.0 6.5 6.0 5.5 5.0 4.5 4.0 3.5 3.0 2.5 2.0 1.5 1.0
f1 (ppm)

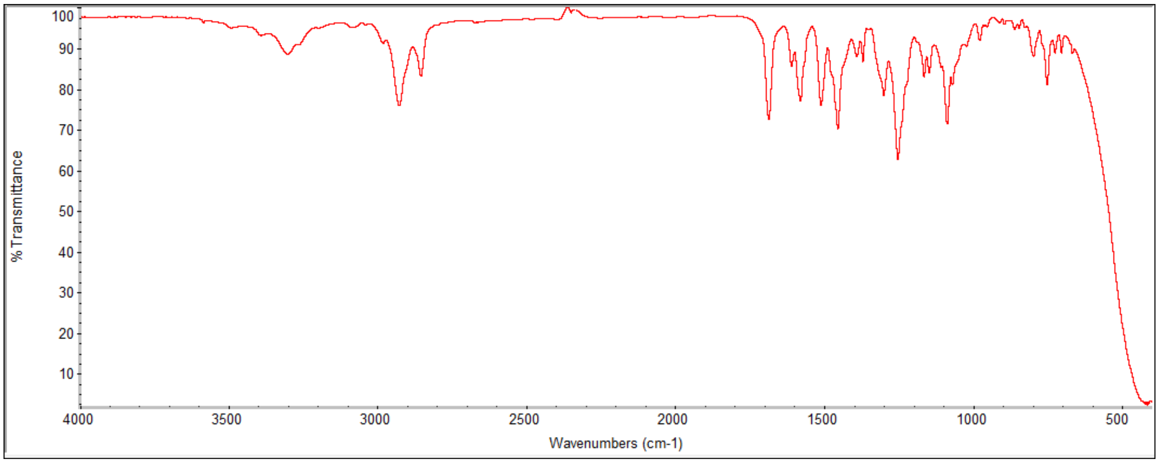

% Transmittance
100 90 80 70 60 50 40 30 20 10
4000 3500 3000 2500 2000 1500 1000 500
Wavenumbers (cm-1)

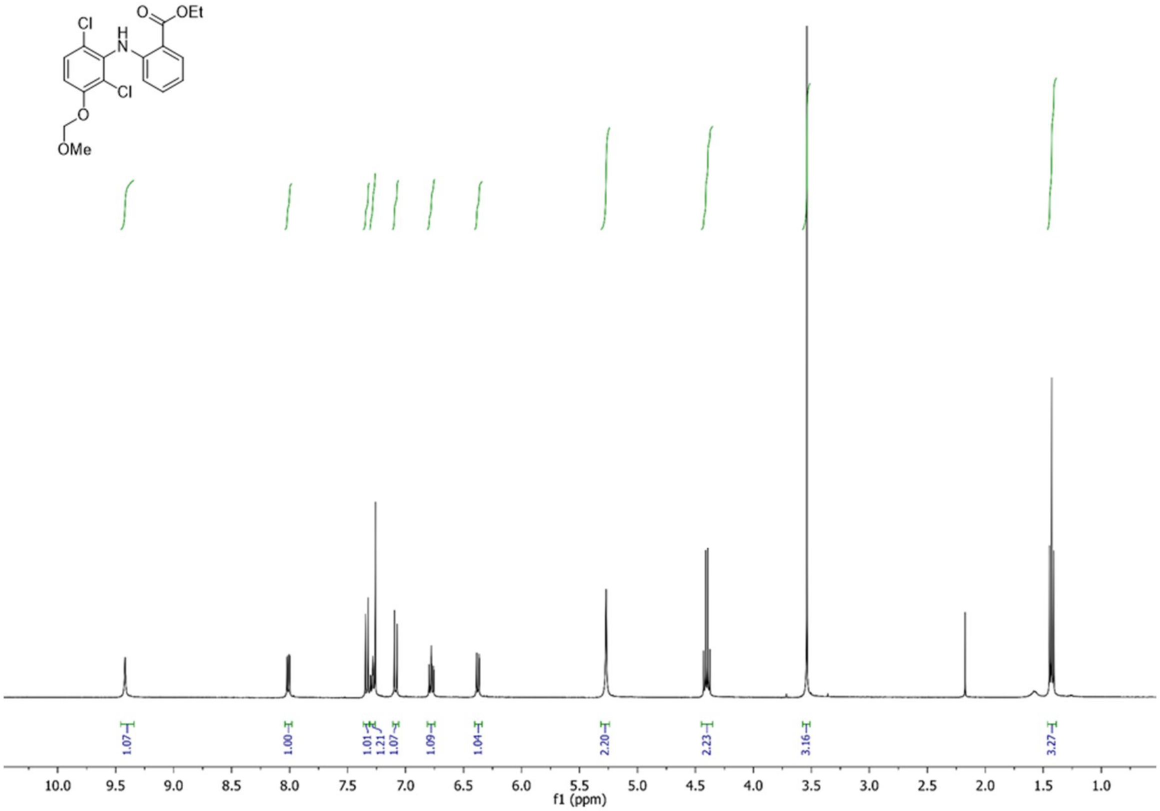

Cl
H
N
O
OEt
Cl
O
OMe
1.07
1.00
1.01
1.21
1.07
1.09
1.04
2.20
2.23
3.16
3.27
10.0 9.5 9.0 8.5 8.0 7.5 7.0 6.5 6.0 5.5 5.0 4.5 4.0 3.5 3.0 2.5 2.0 1.5 1.0
f1 (ppm)

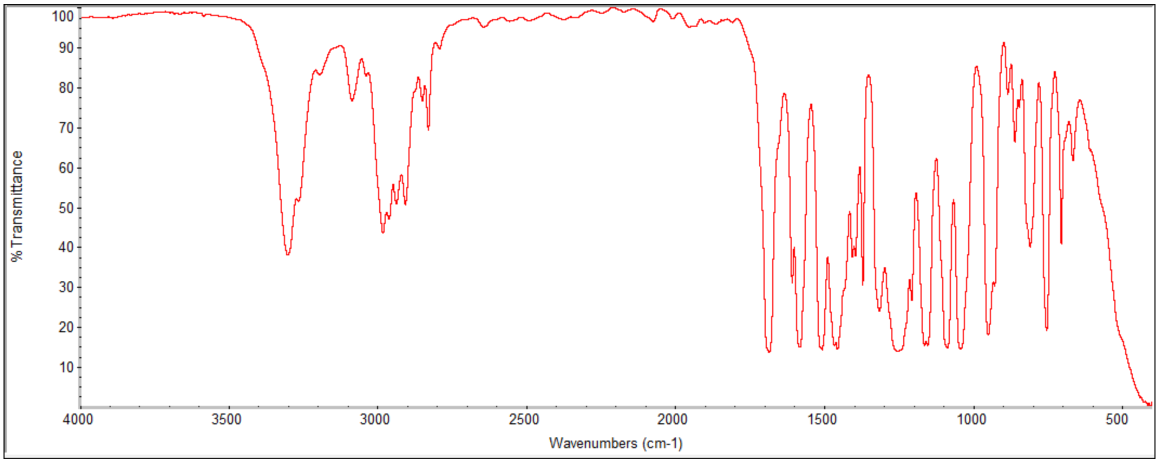

% Transmittance
100 90 80 70 60 50 40 30 20 10
4000 3500 3000 2500 2000 1500 1000 500
Wavenumbers (cm-1)

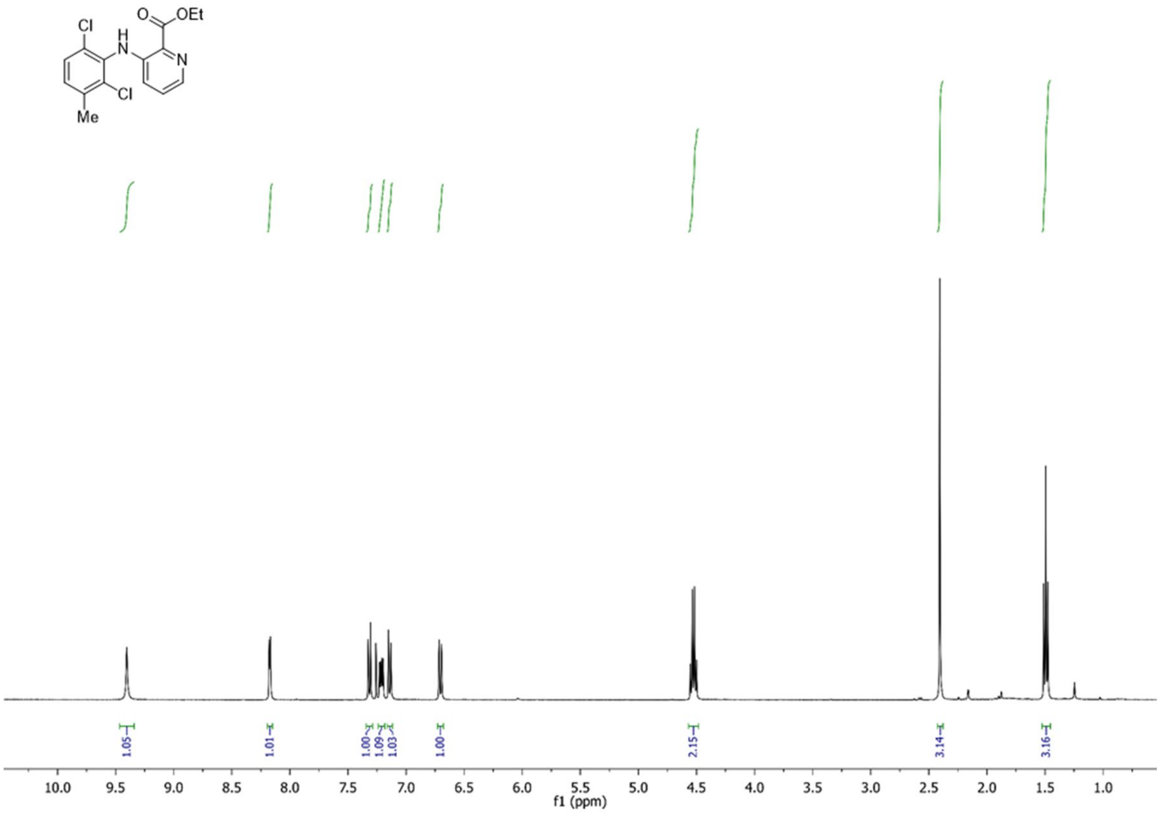
Cl
H
N
O
OEt
N
Cl
Me
1.05
1.01
1.00
1.09
1.03
1.00
2.15
3.14
3.16
10.0 9.5 9.0 8.5 8.0 7.5 7.0 6.5 6.0 5.5 5.0 4.5 4.0 3.5 3.0 2.5 2.0 1.5 1.0
f1 (ppm)

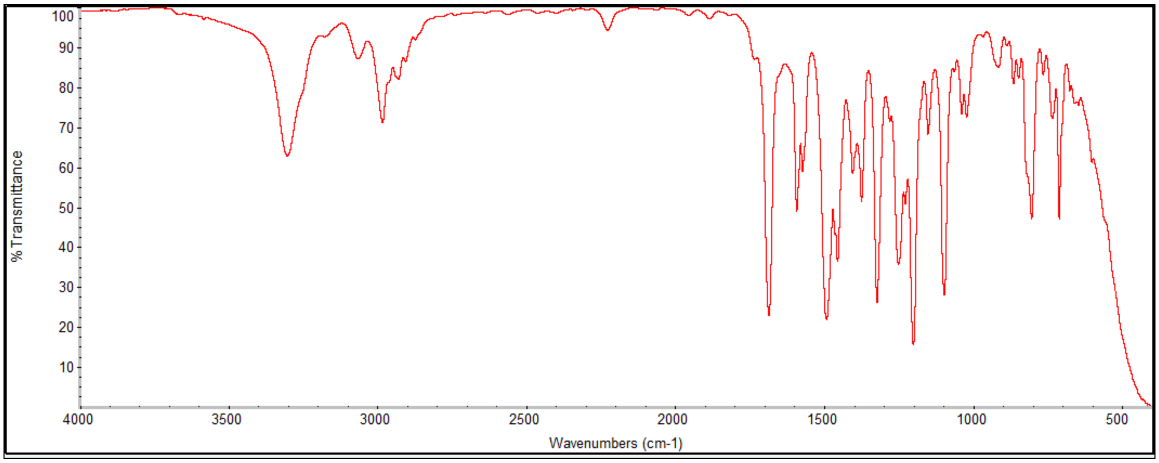
% Transmittance
100 90 80 70 60 50 40 30 20 10
4000 3500 3000 2500 2000 1500 1000 500
Wavenumbers (cm-1)

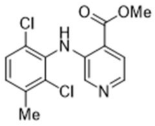

Cl
H
N
O
OMe
Cl
N
Me

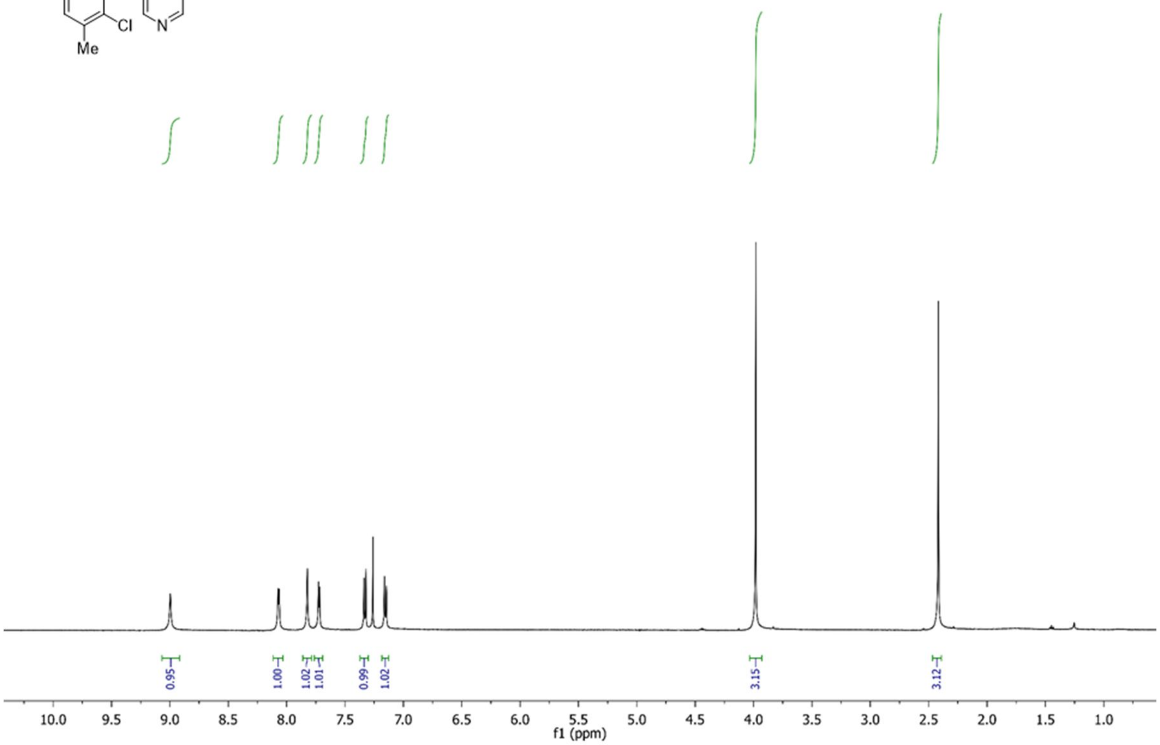

0.95
1.00
1.02
1.01
0.99
1.02
3.15
3.12
10.0
9.5
9.0
8.5
8.0
7.5
7.0
6.5
6.0
5.5
5.0
4.5
4.0
3.5
3.0
2.5
2.0
1.5
1.0
f1 (ppm)

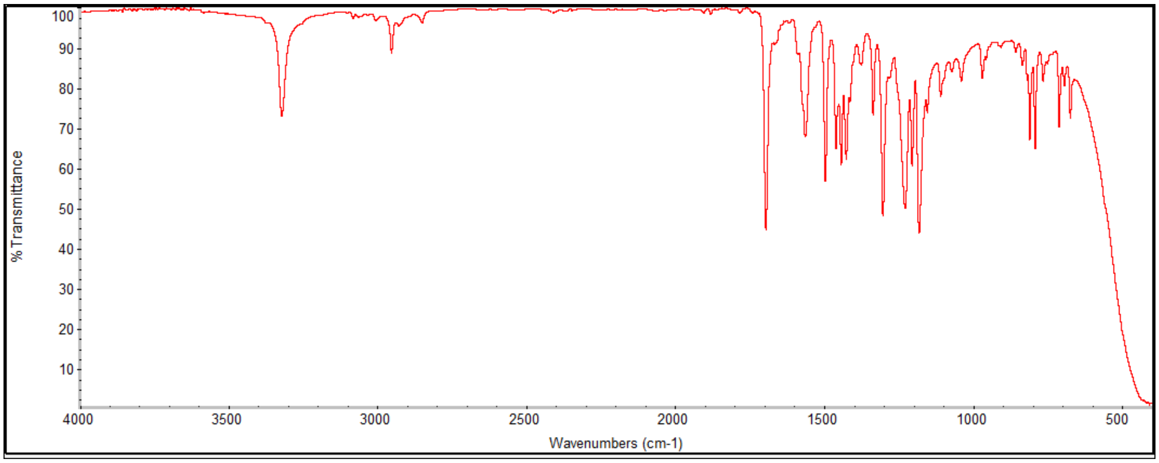

100
90
80
70
60
50
40
30
20
10
% Transmittance
4000
3500
3000
2500
2000
1500
1000
500
Wavenumbers (cm-1)

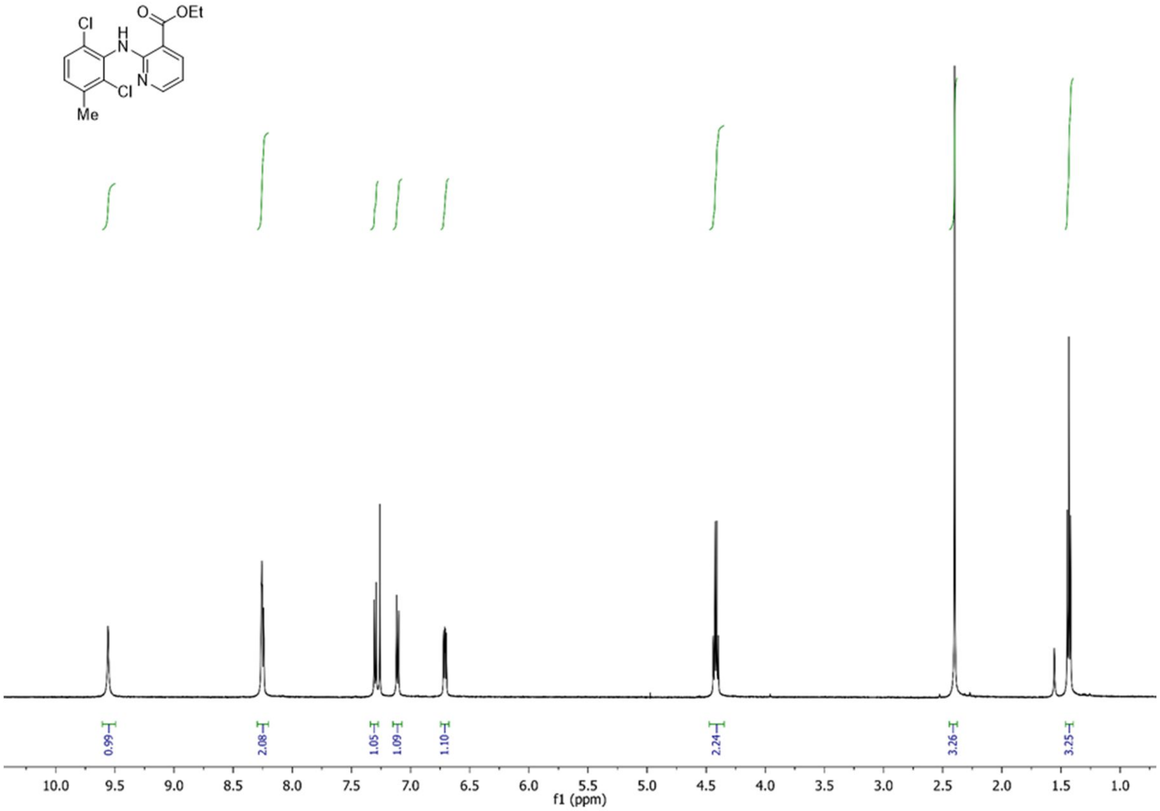

O
OEt
Cl
H
N
N
Cl
Me
0.99
2.08
1.05
1.09
1.10
2.24
3.26
3.25
10.0 9.5 9.0 8.5 8.0 7.5 7.0 6.5 6.0 5.5 5.0 4.5 4.0 3.5 3.0 2.5 2.0 1.5 1.0
f1 (ppm)

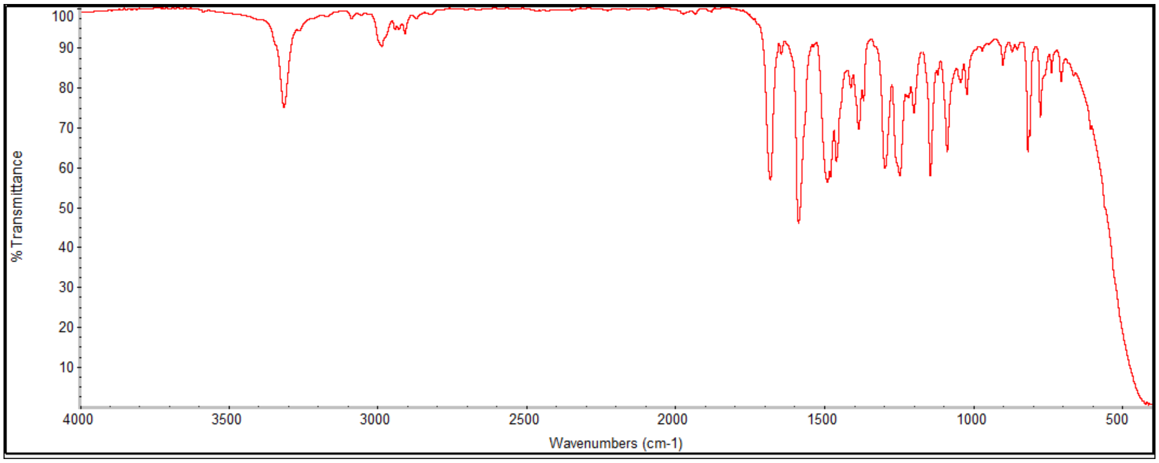

% Transmittance
100 90 80 70 60 50 40 30 20 10
4000 3500 3000 2500 2000 1500 1000 500
Wavenumbers (cm-1)

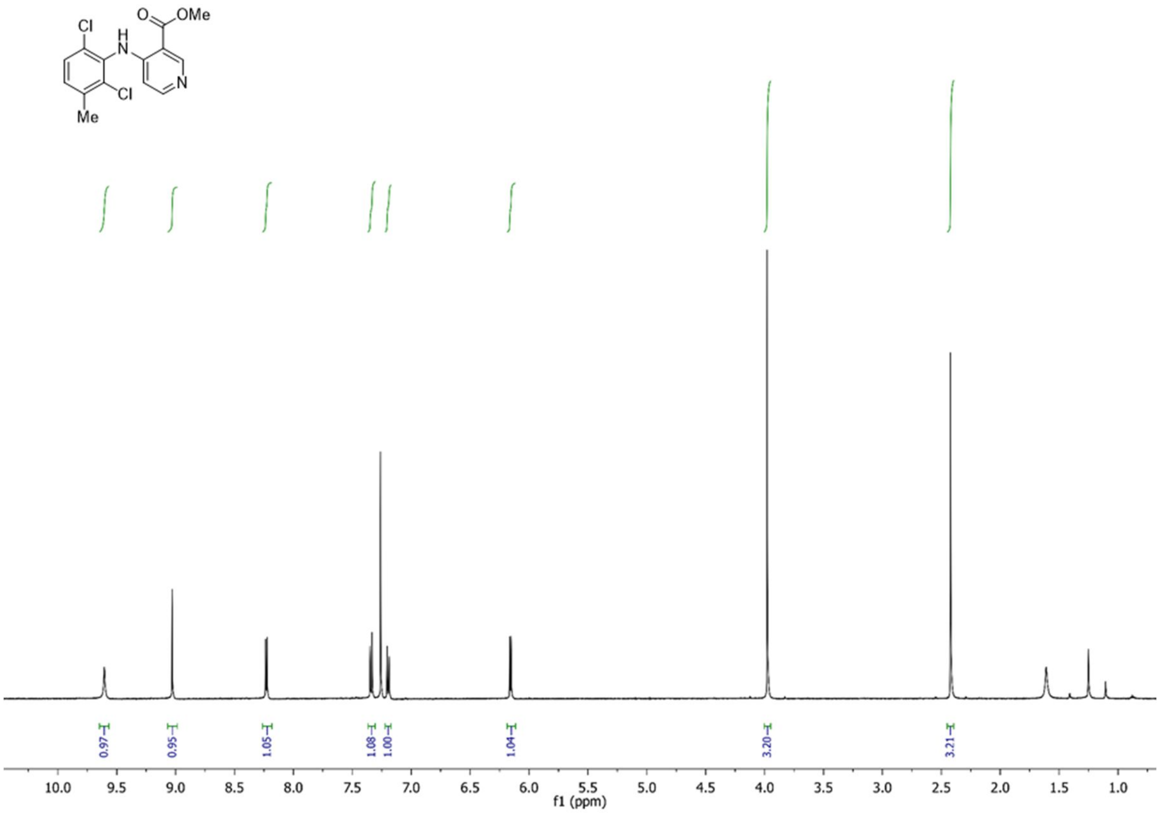

OMe
Cl
Cl
Me
N
0.97
0.95
1.05
1.08
1.00
1.04
3.20
3.21
10.0
9.5
9.0
8.5
8.0
7.5
7.0
6.5
6.0
5.5
5.0
4.5
4.0
3.5
3.0
2.5
2.0
1.5
1.0
f1 (ppm)

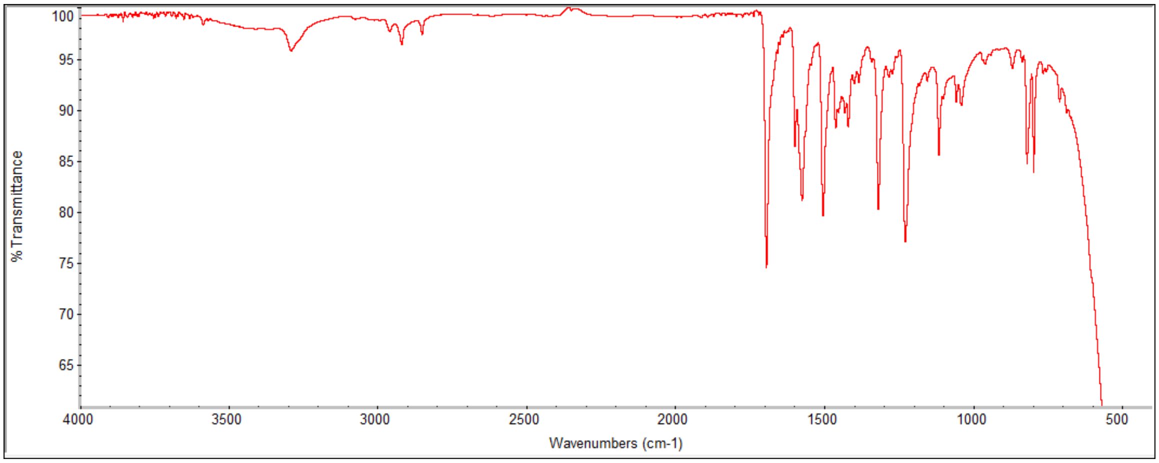

% Transmittance
100
95
90
85
80
75
70
65
4000
3500
3000
2500
2000
1500
1000
500
Wavenumbers (cm-1)

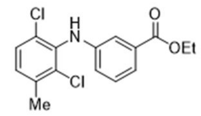
Cl
H
N
O
OEt
Cl
Me

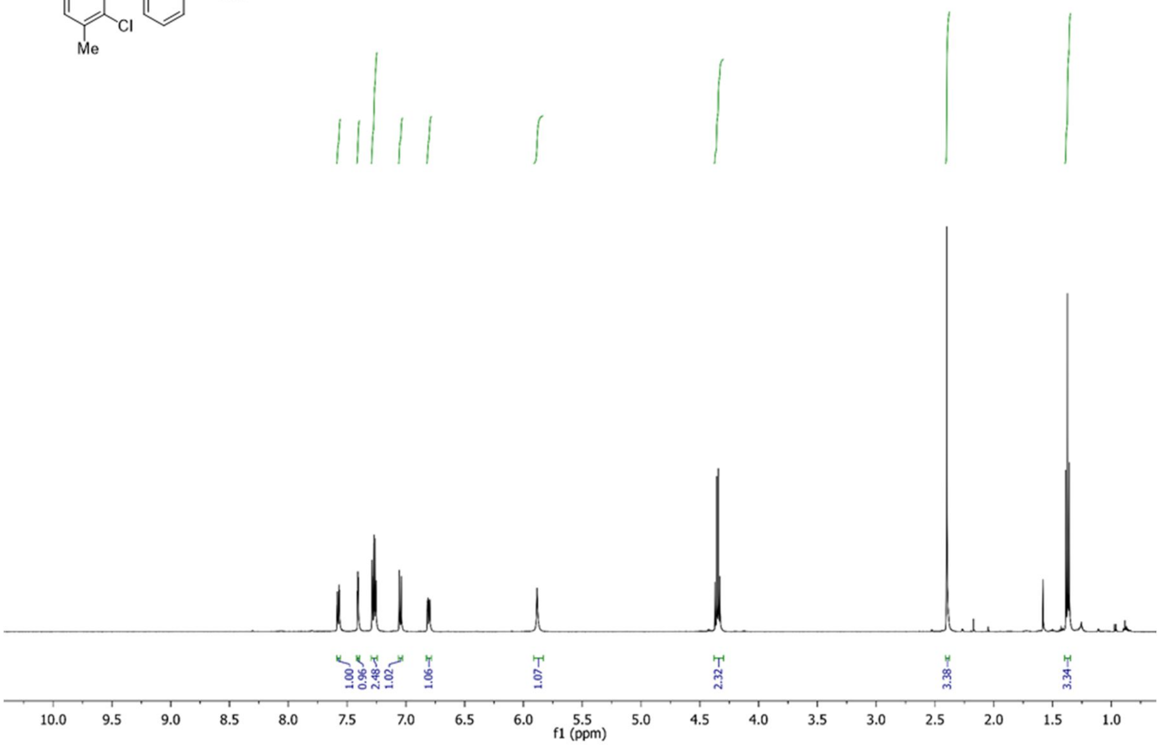
1.00
0.96
2.48
1.02
1.06
1.07
2.32
3.38
3.34
10.0 9.5 9.0 8.5 8.0 7.5 7.0 6.5 6.0 5.5 5.0 4.5 4.0 3.5 3.0 2.5 2.0 1.5 1.0
f1 (ppm)

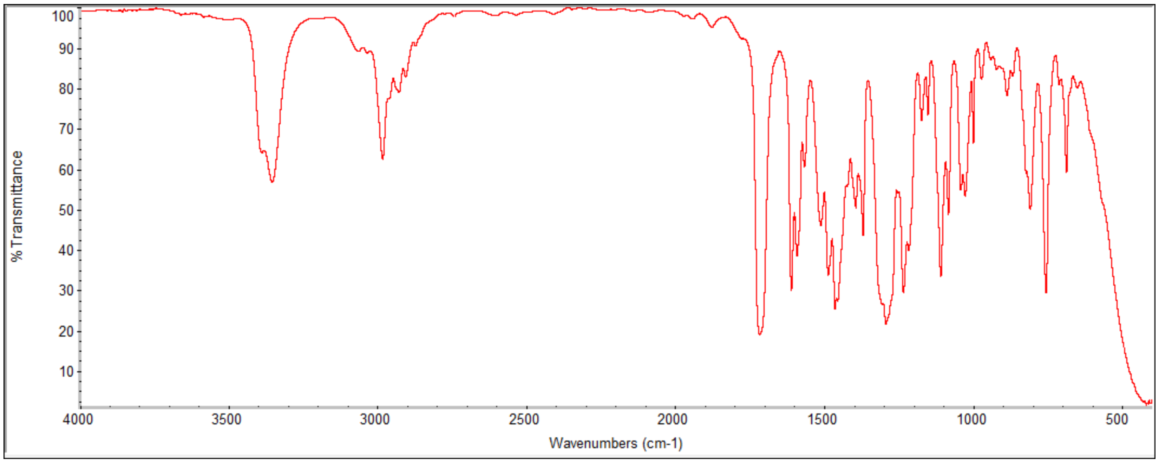
% Transmittance
100 90 80 70 60 50 40 30 20 10
4000 3500 3000 2500 2000 1500 1000 500
Wavenumbers (cm-1)

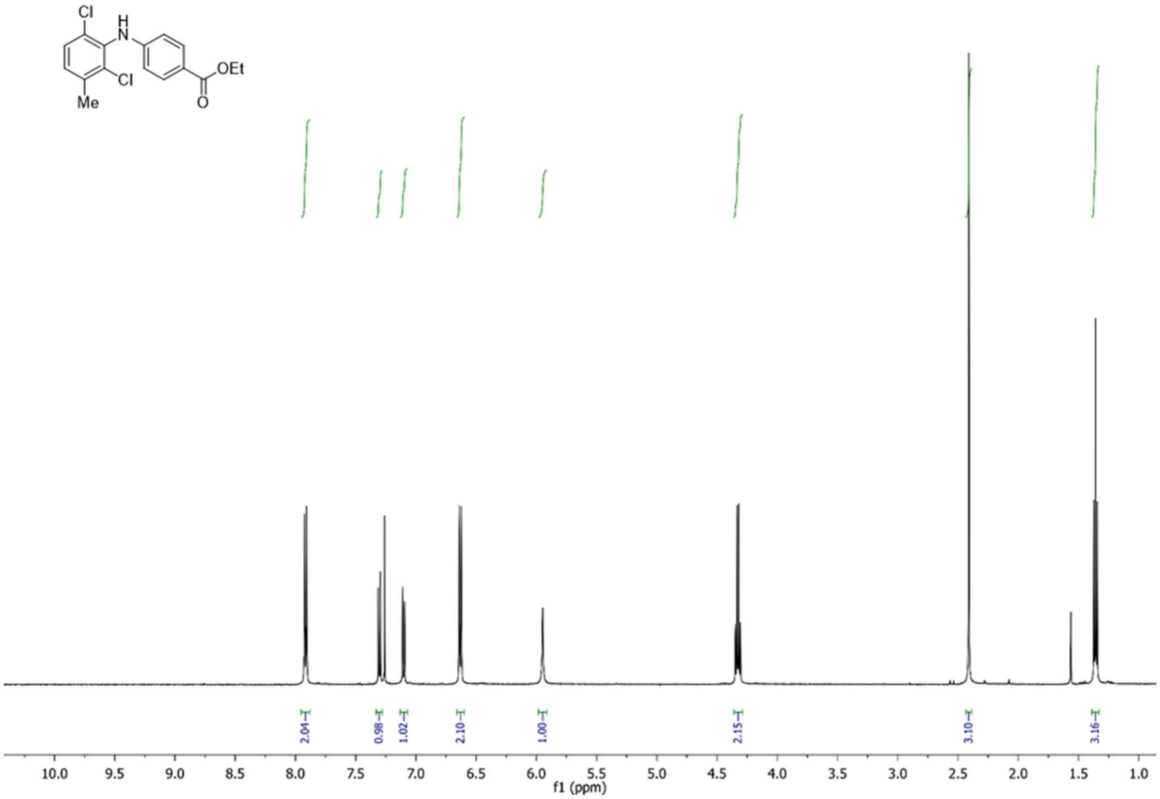
Cl
H
N
Cl
OEt
O
Me
2.04
0.98
1.02
2.10
1.00
2.15
3.10
3.16
10.0 9.5 9.0 8.5 8.0 7.5 7.0 6.5 6.0 5.5 5.0 4.5 4.0 3.5 3.0 2.5 2.0 1.5 1.0
f1 (ppm)

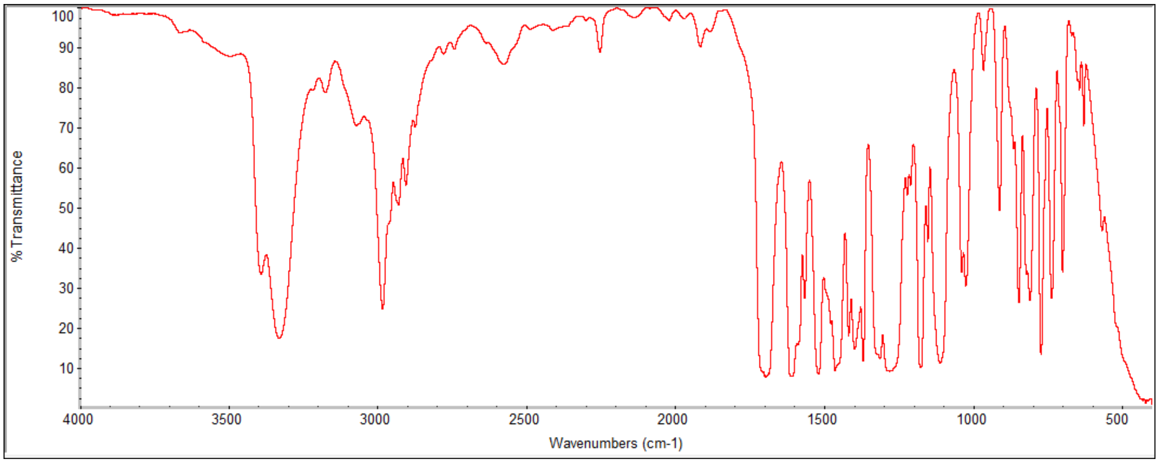
% Transmittance
100 90 80 70 60 50 40 30 20 10
4000 3500 3000 2500 2000 1500 1000 500
Wavenumbers (cm-1)

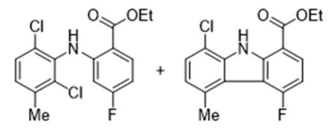

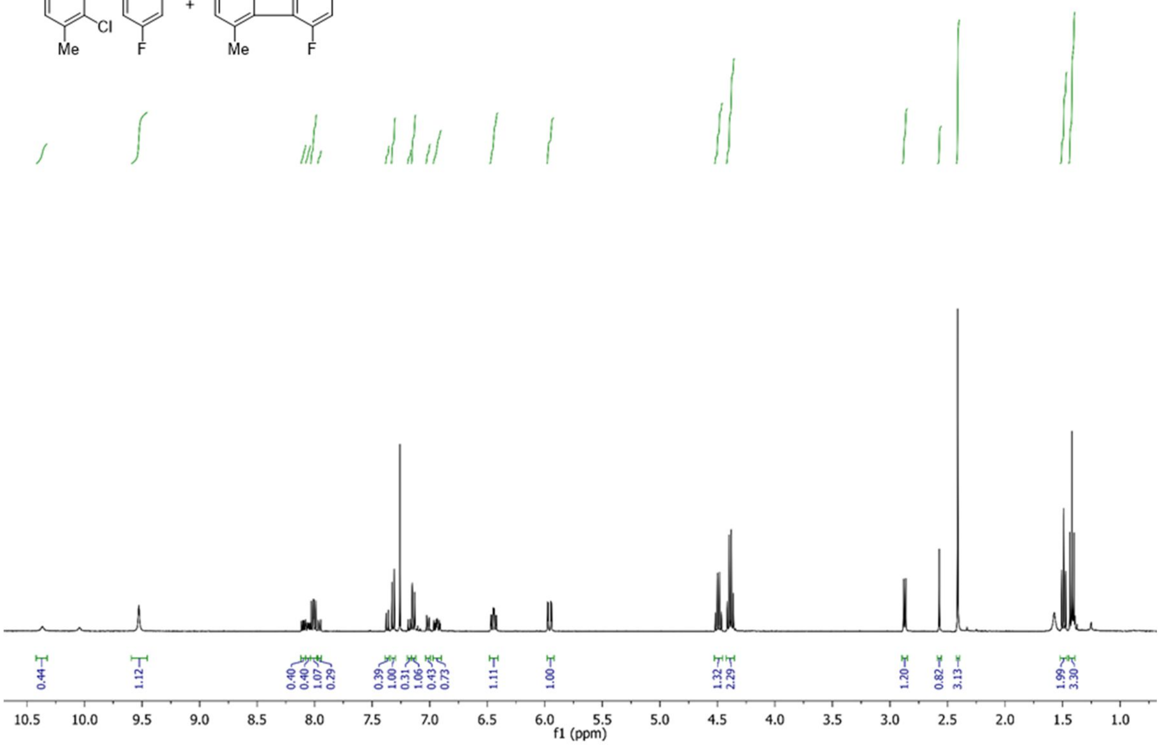

f1 (ppm)

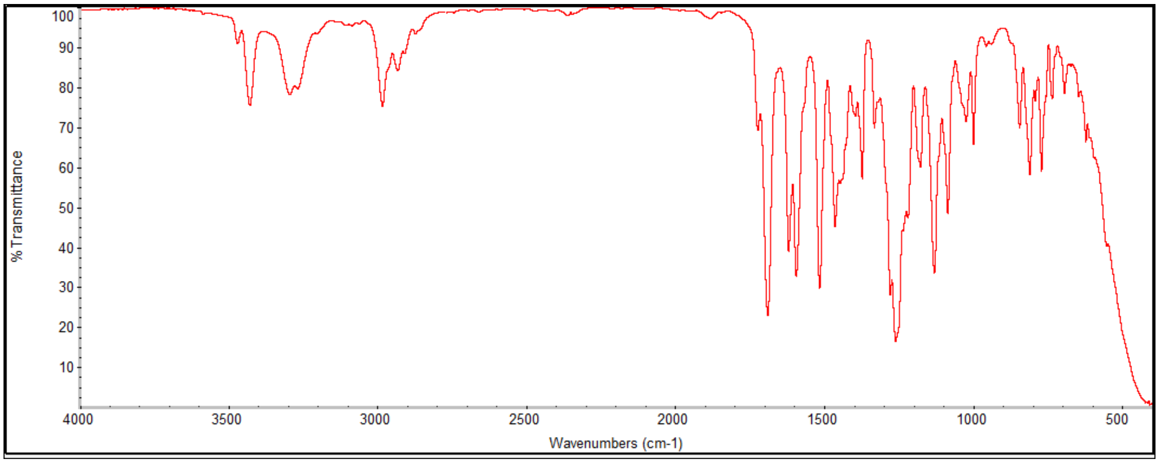

% Transmittance
Wavenumbers (cm-1)

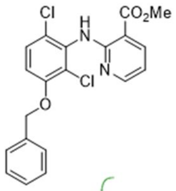
Cl
H
N
CO2Me
Cl
N
O

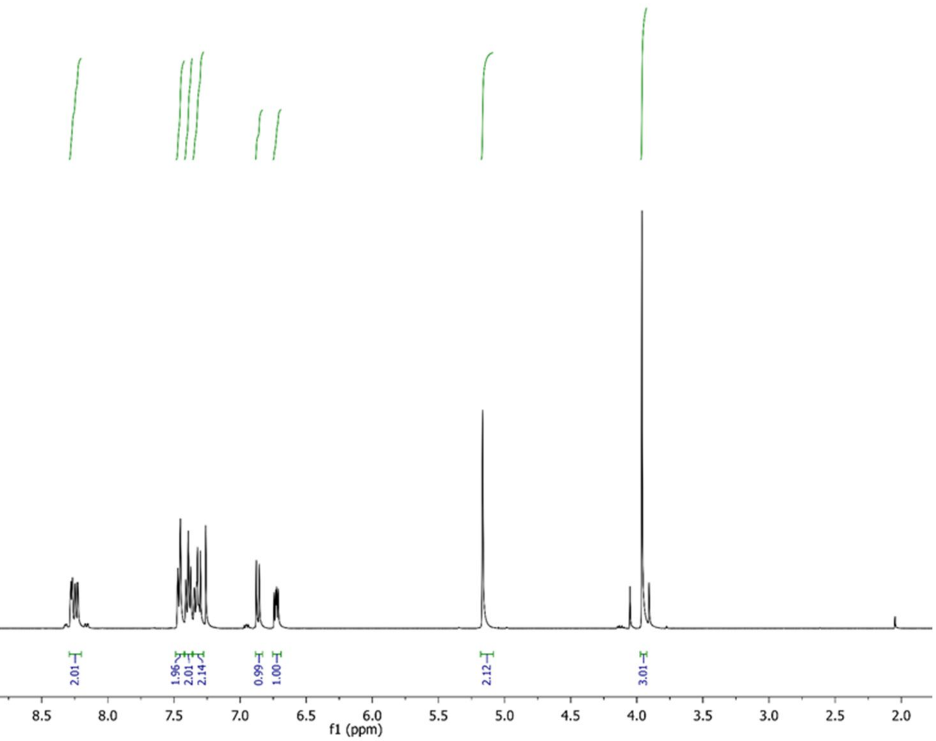
0.95
2.01
1.96
2.01
2.14
0.99
1.00
2.12
3.01
10.0
9.5
9.0
8.5
8.0
7.5
7.0
6.5
6.0
5.5
5.0
4.5
4.0
3.5
3.0
2.5
2.0
f1 (ppm)

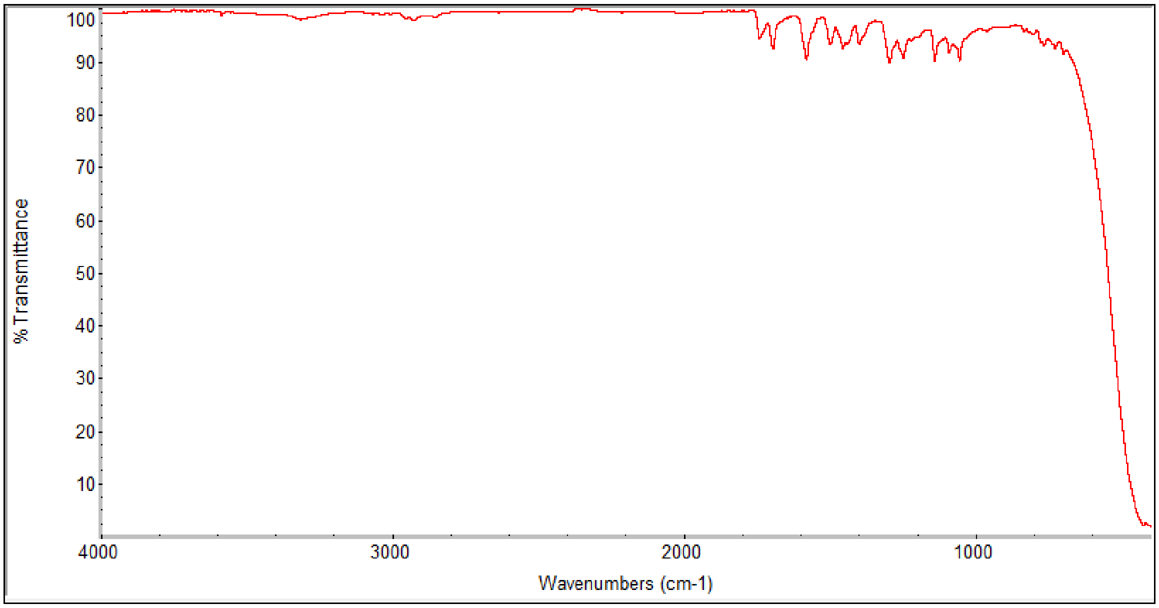
% Transmittance
100
90
80
70
60
50
40
30
20
10
4000
3000
2000
1000
Wavenumbers (cm-1)

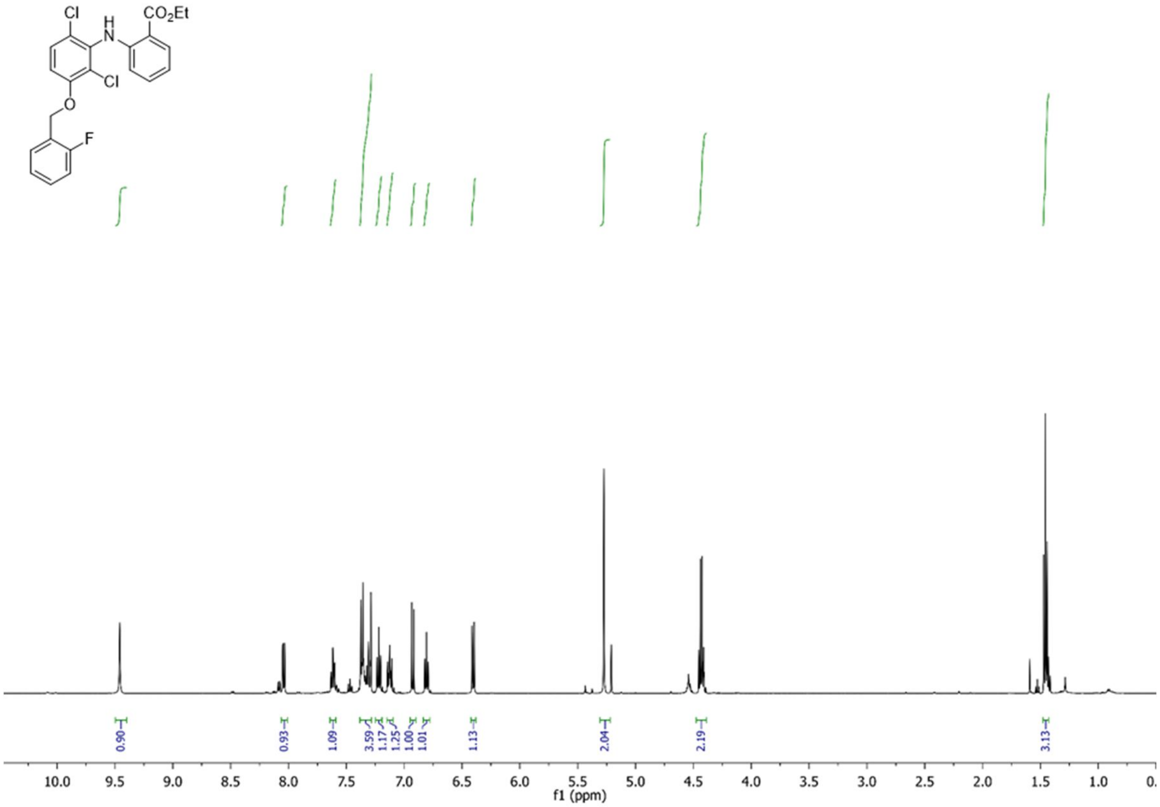
Cl
H
N
CO2Et
Cl
O
F
0.90
0.93
1.09
3.59
1.17
1.25
1.00
1.01
1.13
2.04
2.19
3.13
10.0 9.5 9.0 8.5 8.0 7.5 7.0 6.5 6.0 5.5 5.0 4.5 4.0 3.5 3.0 2.5 2.0 1.5 1.0
f1 (ppm)

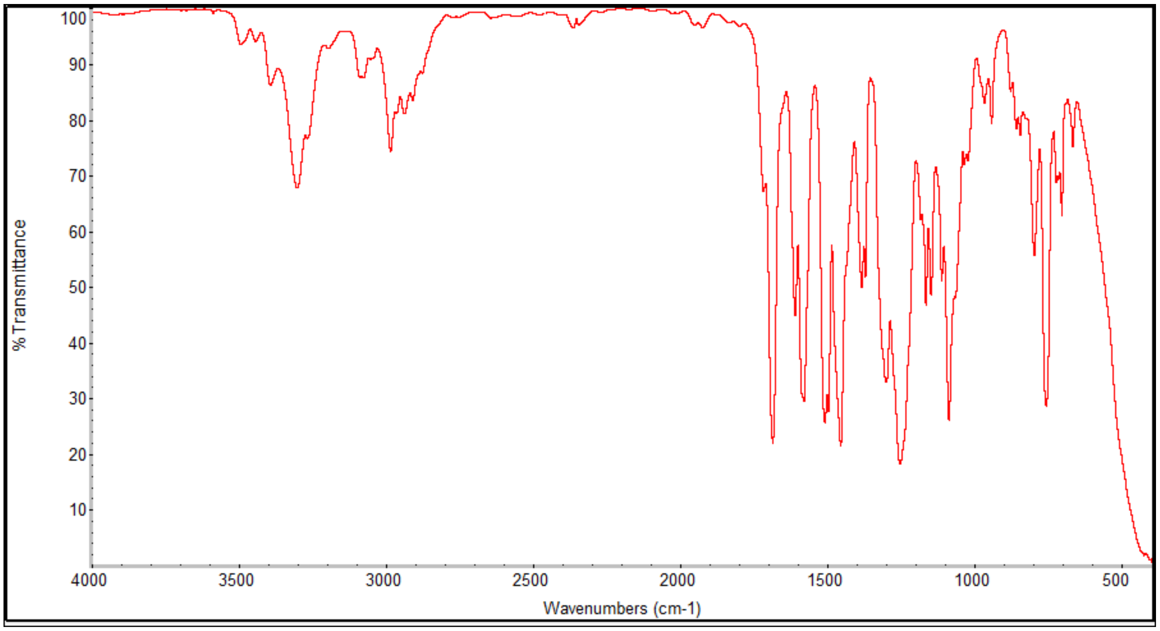
%Transmittance
100 90 80 70 60 50 40 30 20 10
4000 3500 3000 2500 2000 1500 1000 500
Wavenumbers (cm-1)

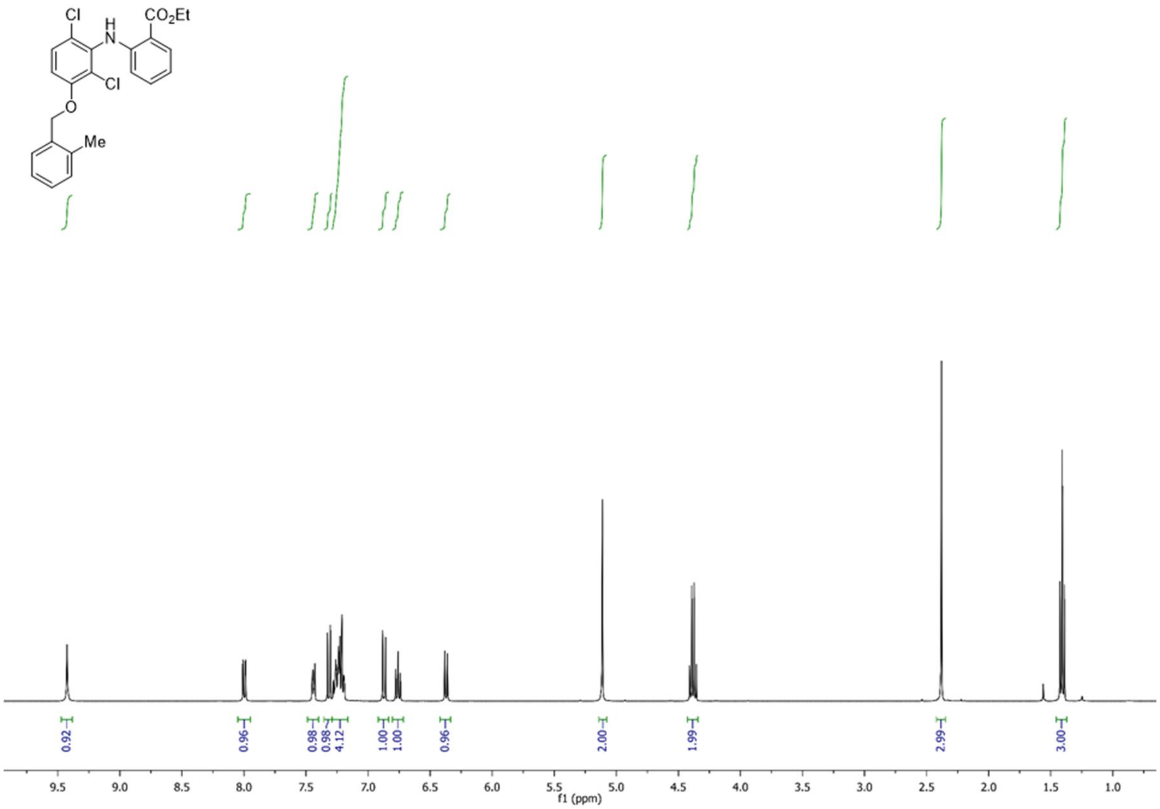
Cl
H
N
CO2Et
Cl
O
Me
0.92
0.96
0.98
0.98
4.12
1.00
1.00
0.96
2.00
1.99
2.99
3.00
9.5 9.0 8.5 8.0 7.5 7.0 6.5 6.0 5.5 5.0 4.5 4.0 3.5 3.0 2.5 2.0 1.5 1.0
f1 (ppm)

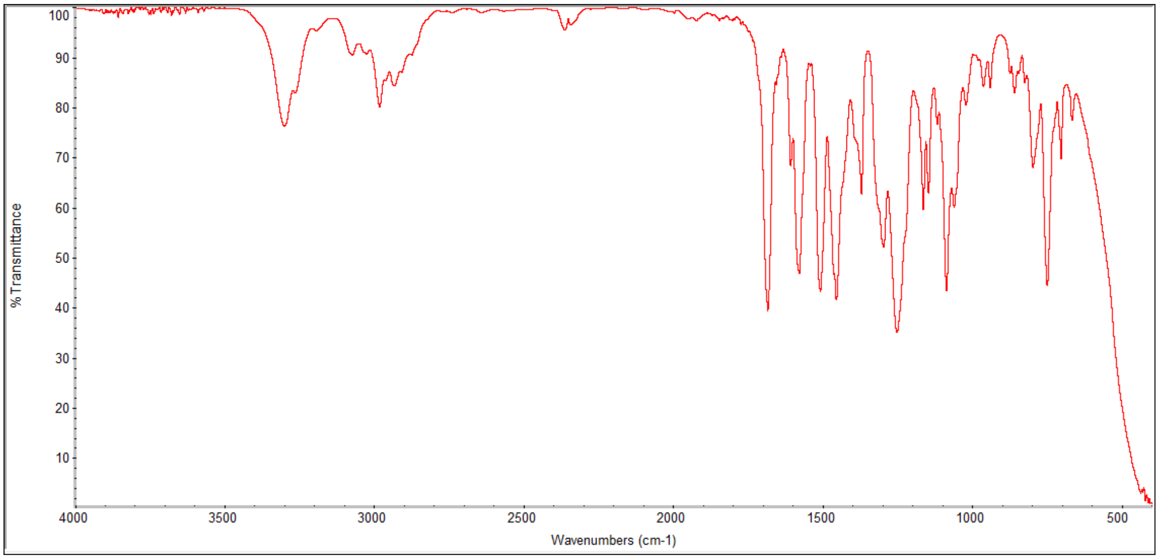
% Transmittance
100
90
80
70
60
50
40
30
20
10
4000 3500 3000 2500 2000 1500 1000 500
Wavenumbers (cm-1)

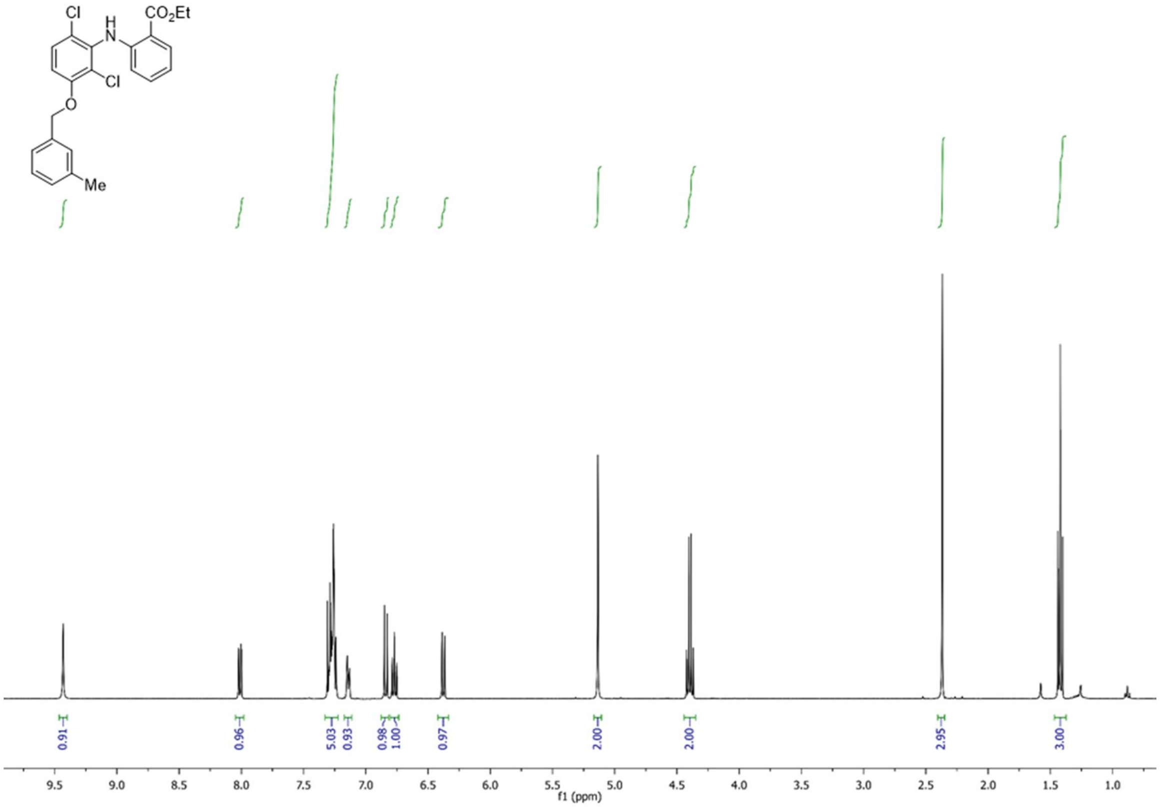

Cl
H
N
CO2Et
Cl
O
Me
0.91
0.96
5.03
0.93
0.98
1.00
0.97
2.00
2.00
2.95
3.00
9.5
9.0
8.5
8.0
7.5
7.0
6.5
6.0
5.5
5.0
4.5
4.0
3.5
3.0
2.5
2.0
1.5
1.0
f1 (ppm)

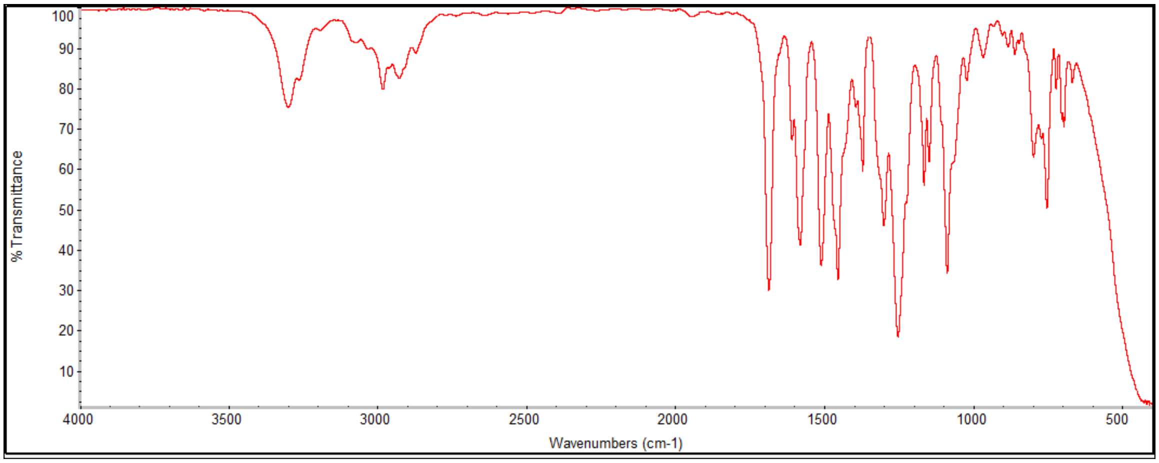

% Transmittance
100
90
80
70
60
50
40
30
20
10
4000
3500
3000
2500
2000
1500
1000
500
Wavenumbers (cm-1)

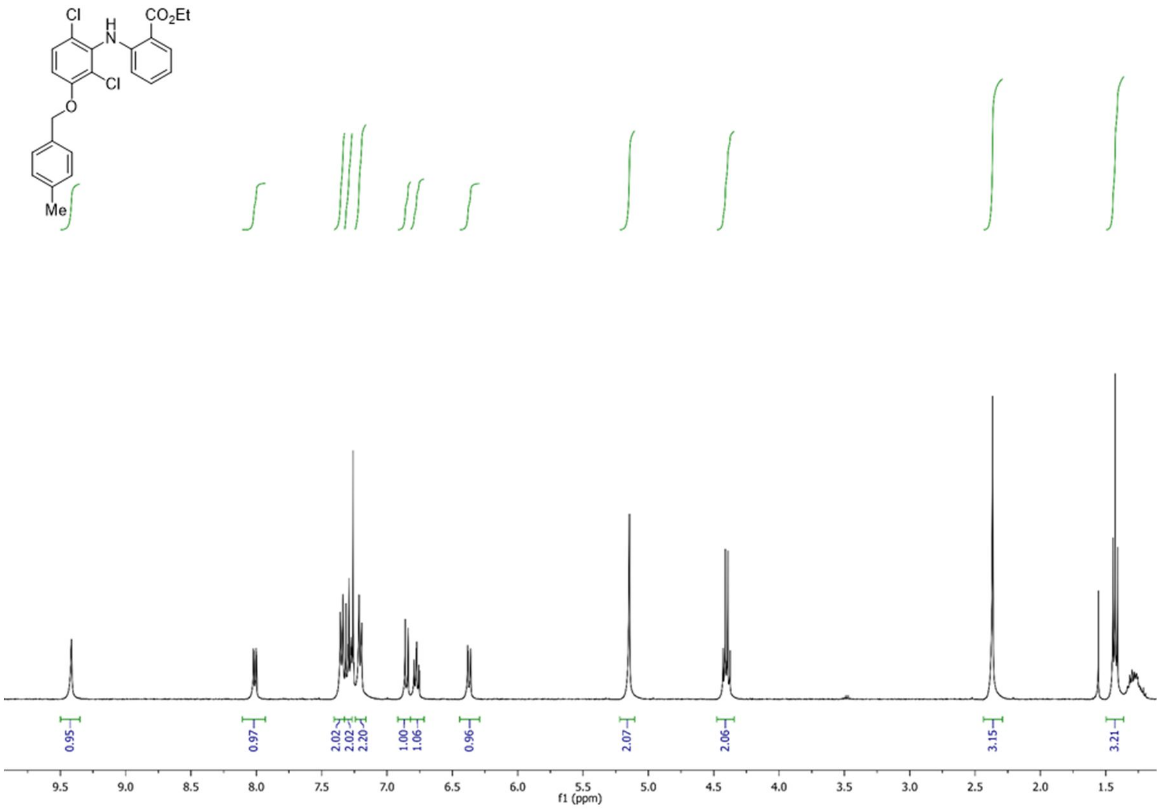
Cl
H
N
CO2Et
Cl
O
Me
0.95
0.97
2.02
2.02
2.20
1.00
1.06
0.96
2.07
2.06
3.15
3.21
9.5 9.0 8.5 8.0 7.5 7.0 6.5 6.0 5.5 5.0 4.5 4.0 3.5 3.0 2.5 2.0 1.5
f1 (ppm)

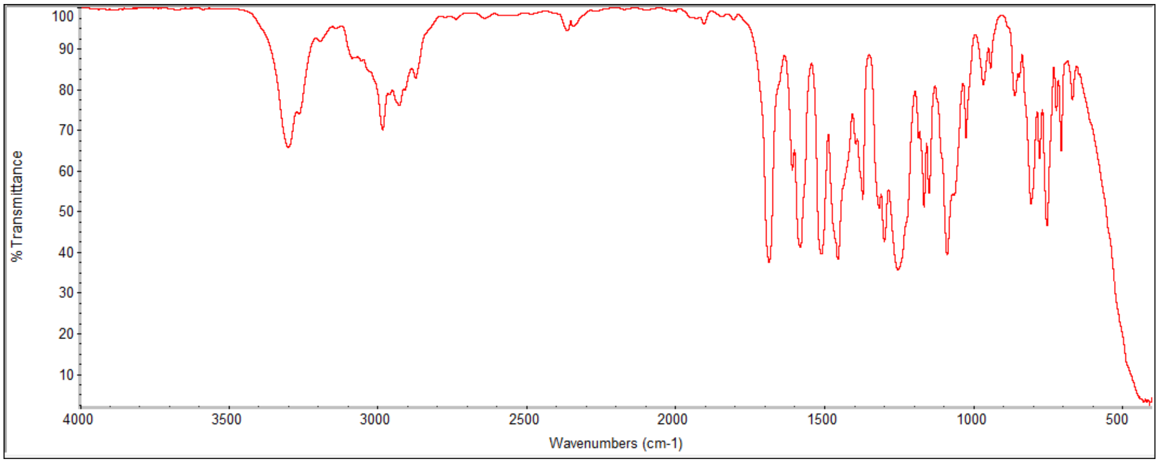
% Transmittance
100 90 80 70 60 50 40 30 20 10
4000 3500 3000 2500 2000 1500 1000 500
Wavenumbers (cm-1)

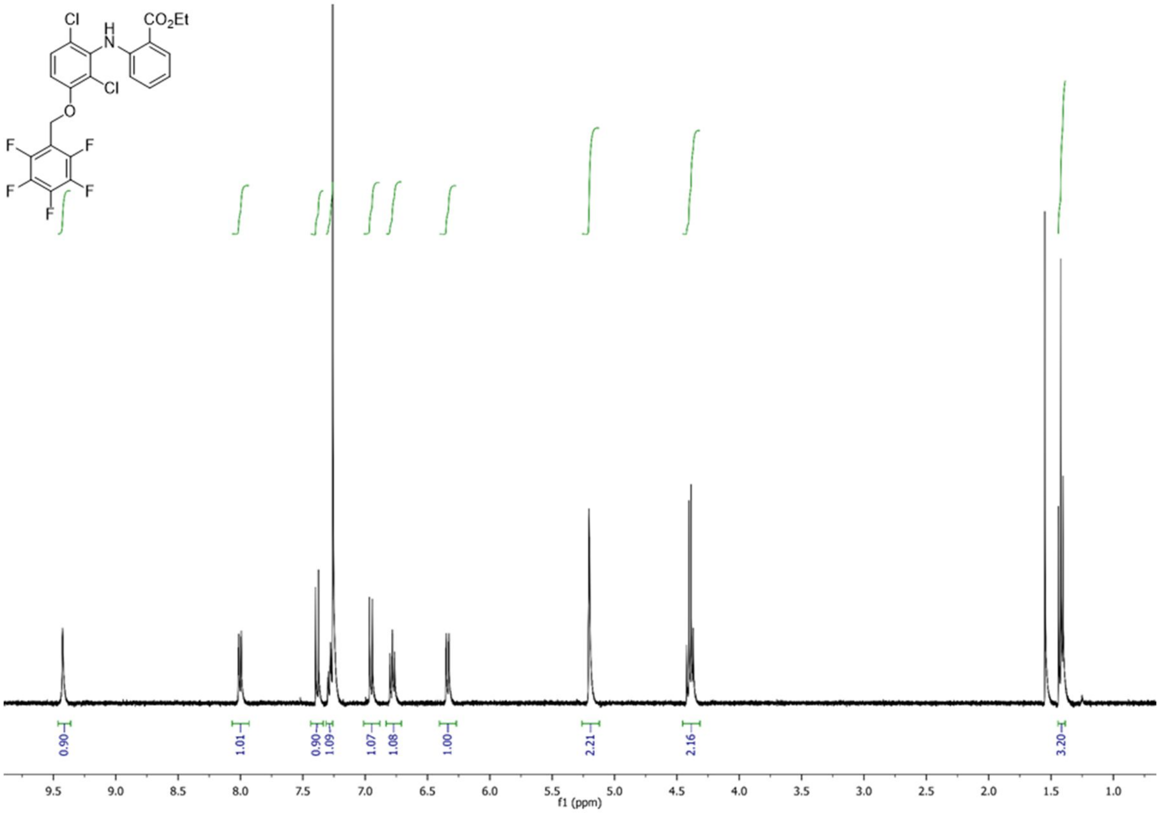
Cl
H
N
CO2Et
Cl
O
F
F
F
F
F
0.90
1.01
0.90
1.09
1.07
1.08
1.00
2.21
2.16
3.20
9.5
9.0
8.5
8.0
7.5
7.0
6.5
6.0
5.5
5.0
4.5
4.0
3.5
3.0
2.5
2.0
1.5
1.0
f1 (ppm)

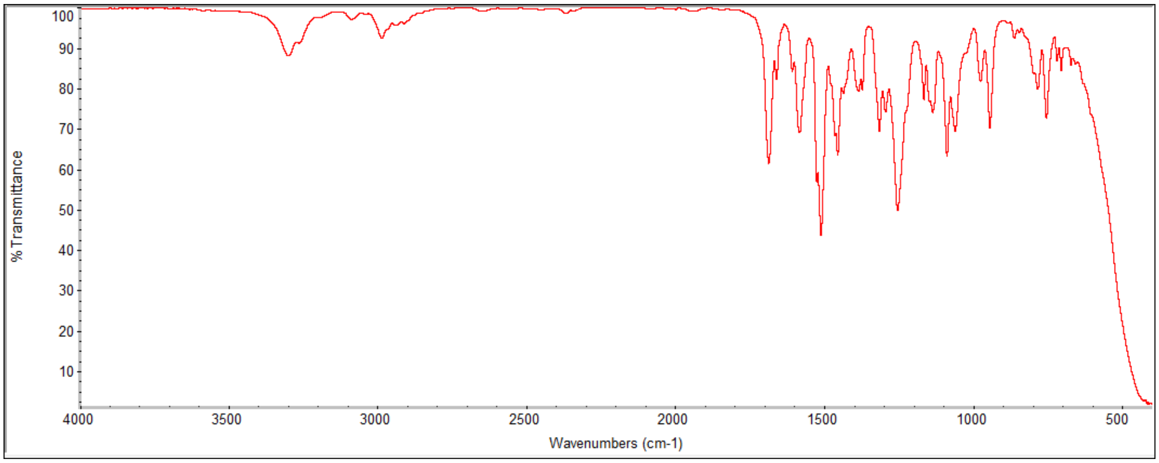
% Transmittance
100
90
80
70
60
50
40
30
20
10
4000
3500
3000
2500
2000
1500
1000
500
Wavenumbers (cm-1)

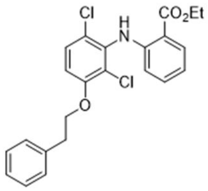
Cl
H
N
CO2Et
Cl
O

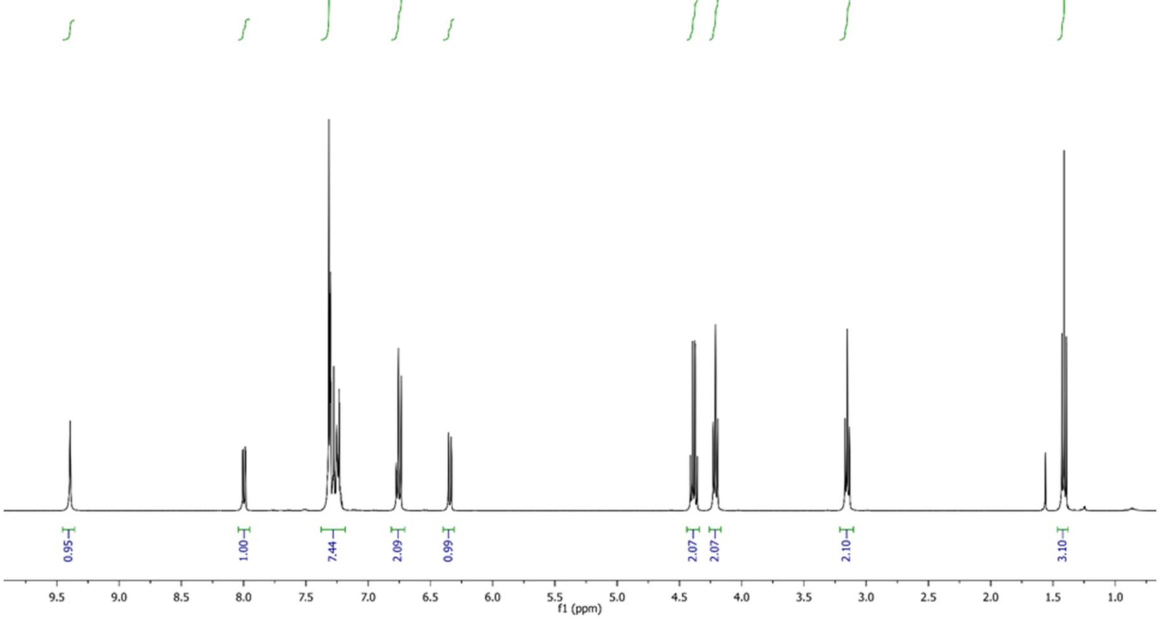
0.95
1.00
7.44
2.09
0.99
2.07
2.07
2.10
3.10
9.5
9.0
8.5
8.0
7.5
7.0
6.5
6.0
5.5
5.0
4.5
4.0
3.5
3.0
2.5
2.0
1.5
1.0
f1 (ppm)

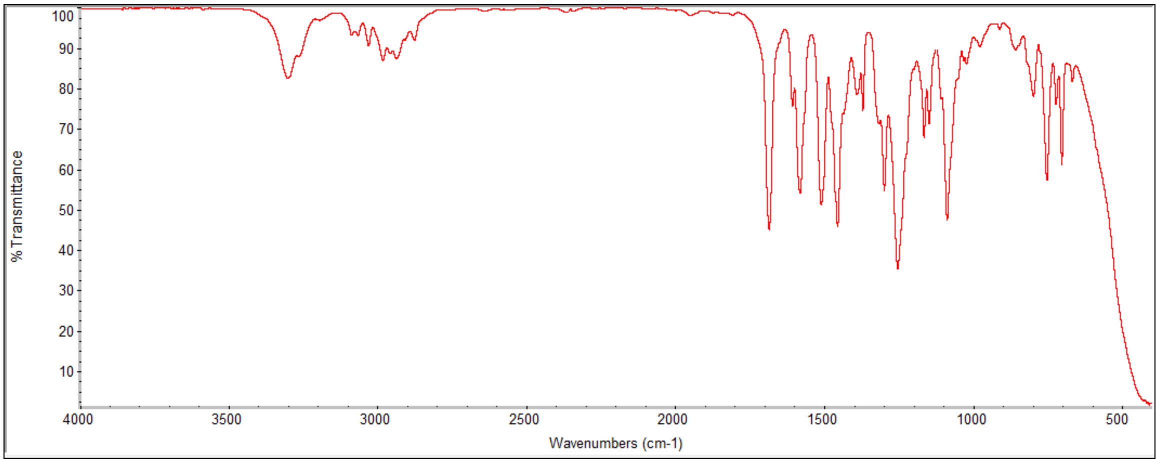
% Transmittance
100
90
80
70
60
50
40
30
20
10
4000
3500
3000
2500
2000
1500
1000
500
Wavenumbers (cm-1)

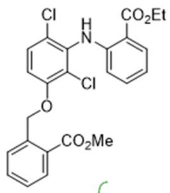

Cl
H
N
CO2Et
Cl
O
CO2Me

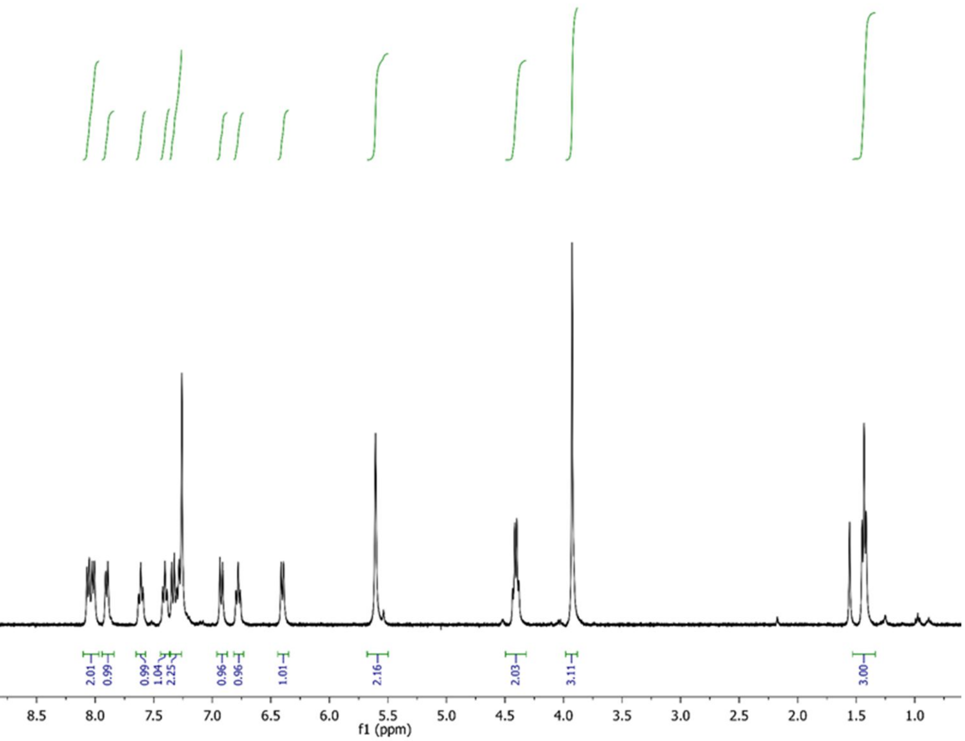

0.93
2.01
0.99
0.99
1.04
2.25
0.96
0.96
1.01
2.16
2.03
3.11
3.00
10.0
9.5
9.0
8.5
8.0
7.5
7.0
6.5
6.0
5.5
5.0
4.5
4.0
3.5
3.0
2.5
2.0
1.5
1.0
f1 (ppm)

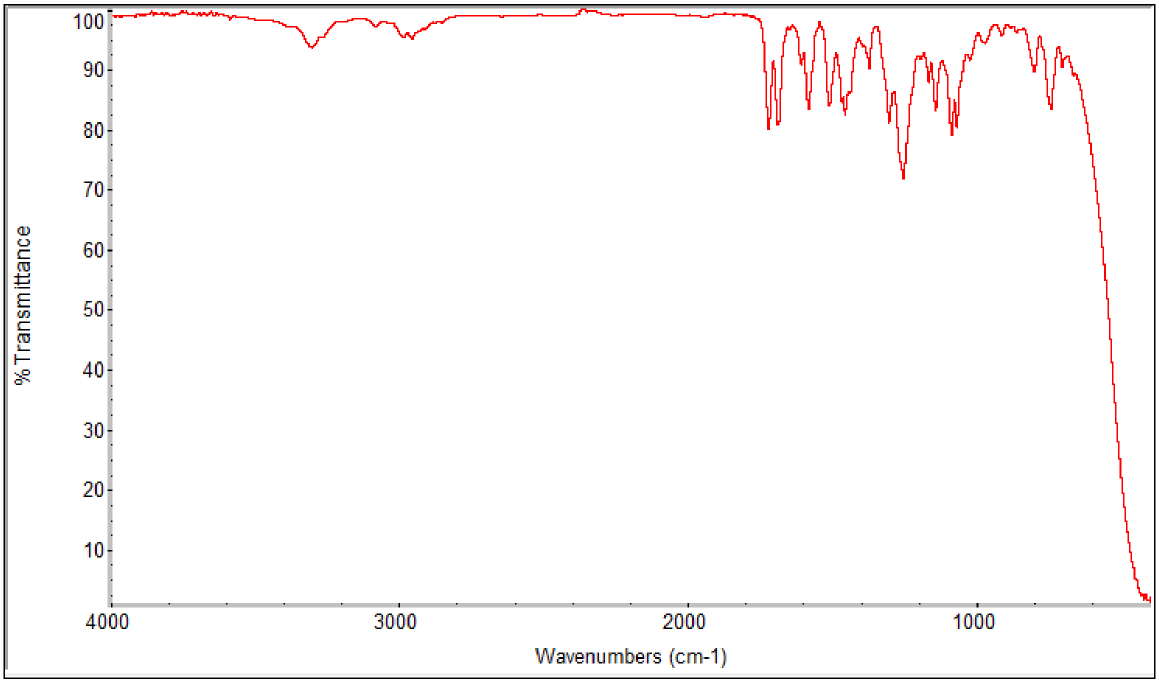

%Transmittance
100
90
80
70
60
50
40
30
20
10
4000
3000
2000
1000
Wavenumbers (cm-1)

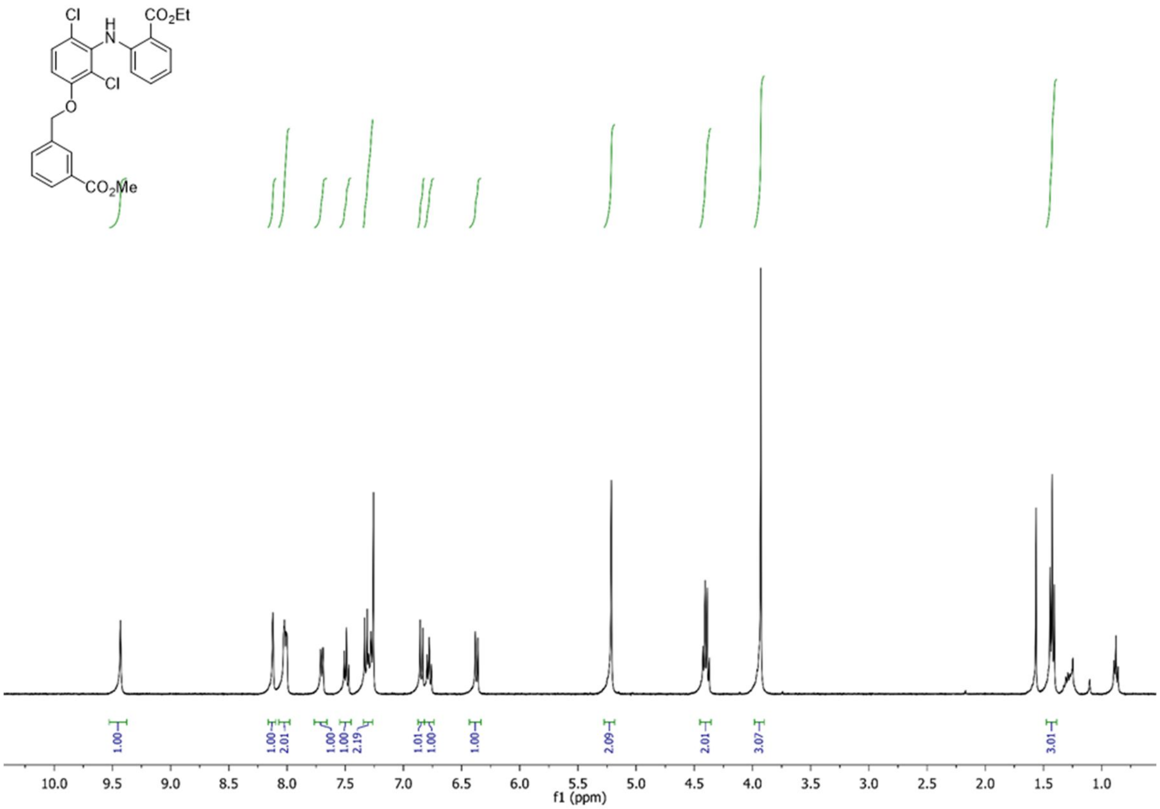
Cl
H
N
CO2Et
Cl
O
CO2Me
1.00
1.00
2.01
1.00
1.00
2.19
1.01
1.00
1.00
2.09
2.01
3.07
3.01
10.0
9.5
9.0
8.5
8.0
7.5
7.0
6.5
6.0
5.5
5.0
4.5
4.0
3.5
3.0
2.5
2.0
1.5
1.0
f1 (ppm)

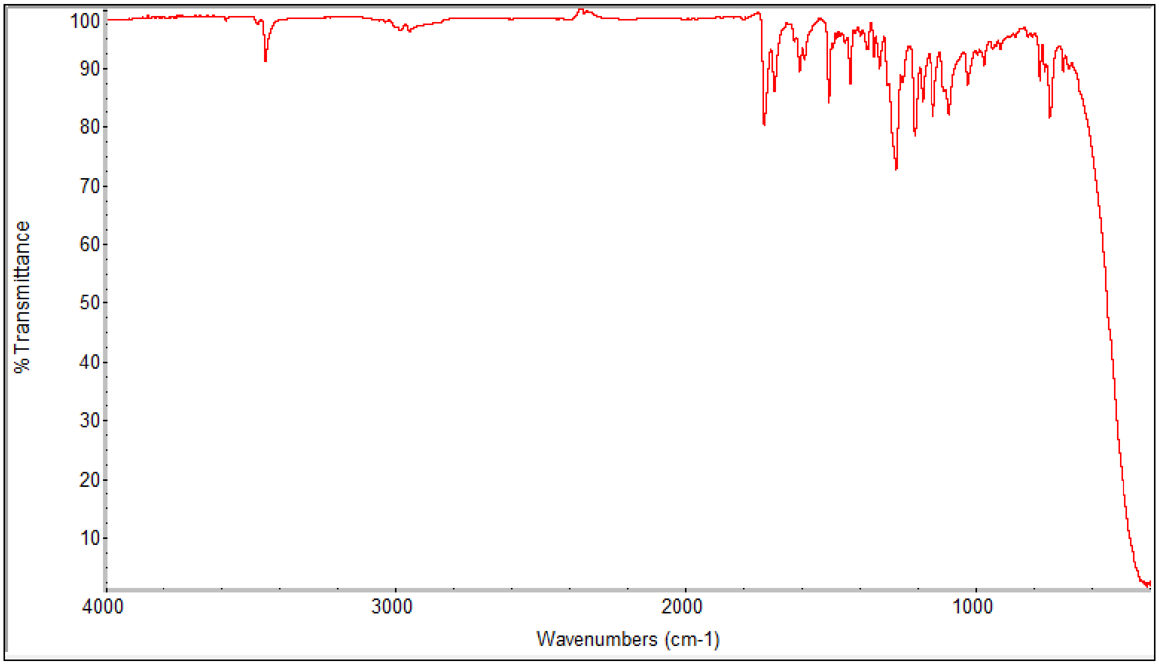
% Transmittance
100
90
80
70
60
50
40
30
20
10
4000
3000
2000
1000
Wavenumbers (cm-1)

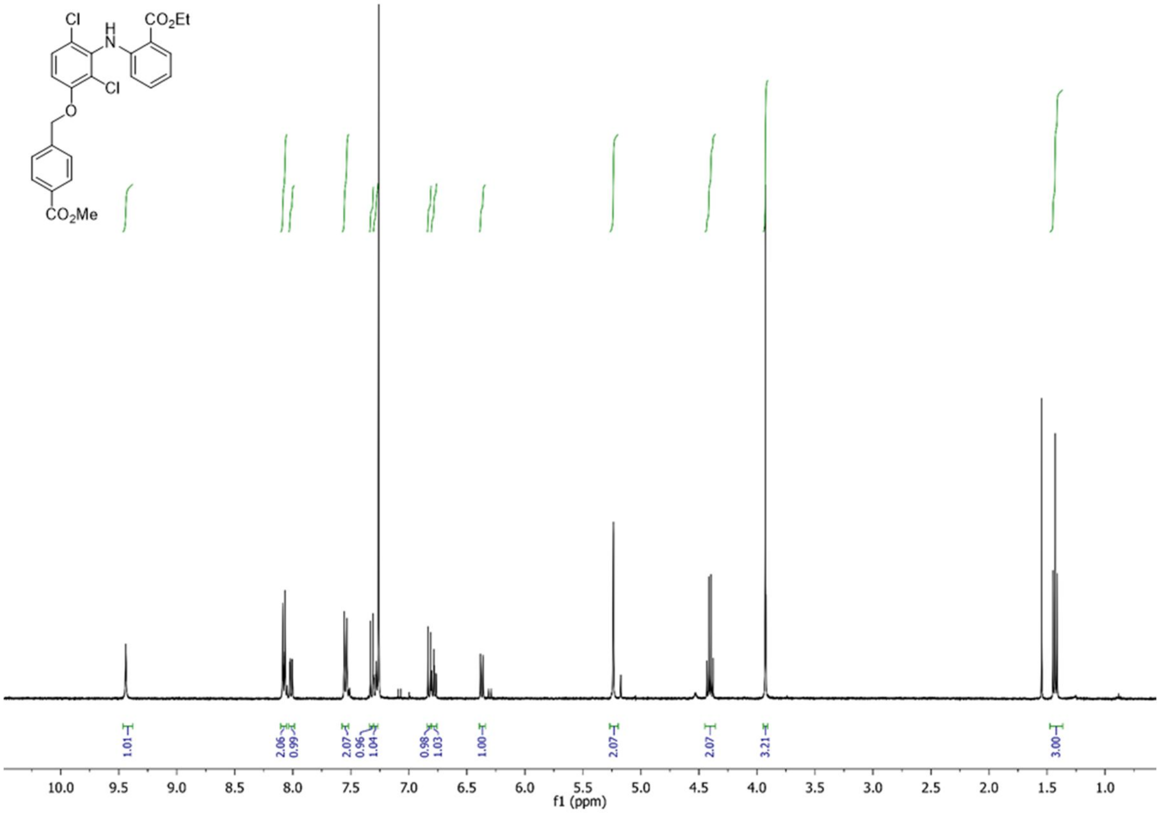
Cl
H
N
CO2Et
Cl
O
CO2Me
1.01
2.06
0.99
2.07
0.96
1.04
0.98
1.03
1.00
2.07
2.07
3.21
3.00
10.0 9.5 9.0 8.5 8.0 7.5 7.0 6.5 6.0 5.5 5.0 4.5 4.0 3.5 3.0 2.5 2.0 1.5 1.0
f1 (ppm)

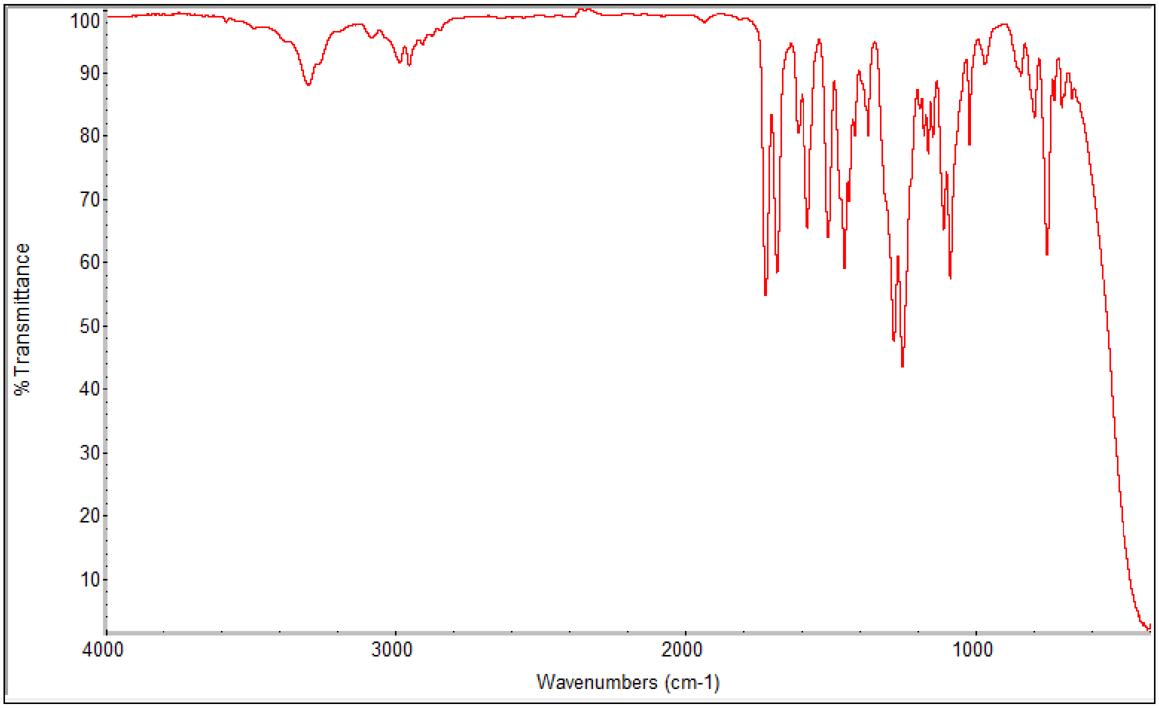
%Transmittance
100
90
80
70
60
50
40
30
20
10
4000
3000
2000
1000
Wavenumbers (cm-1)

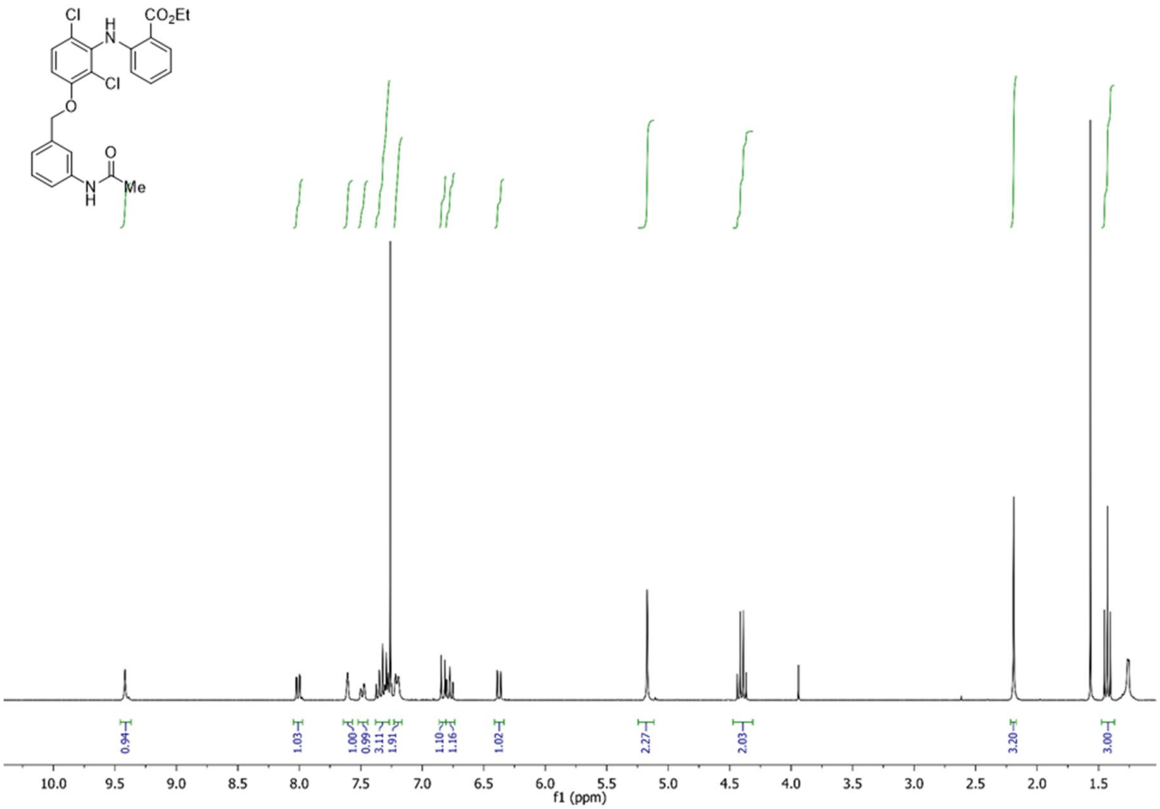
Cl
H
N
CO2Et
Cl
O
O
N
H
Me
0.94
1.03
1.00
0.99
3.11
1.91
1.10
1.16
1.02
2.27
2.03
3.20
3.00
10.0 9.5 9.0 8.5 8.0 7.5 7.0 6.5 6.0 5.5 5.0 4.5 4.0 3.5 3.0 2.5 2.0 1.5
f1 (ppm)

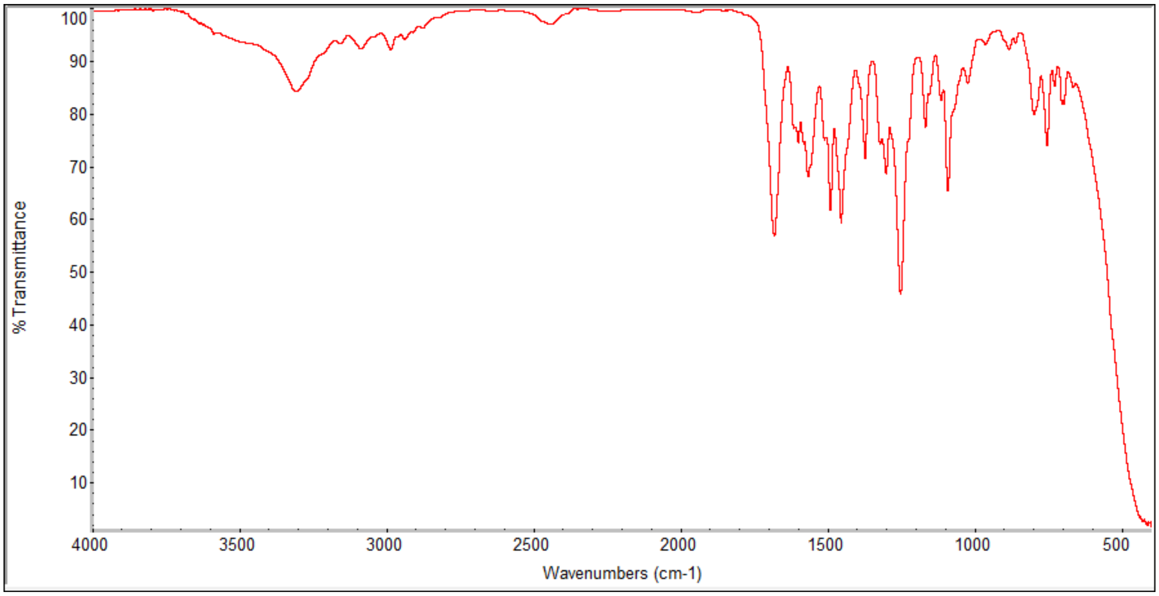
% Transmittance
100 90 80 70 60 50 40 30 20 10
4000 3500 3000 2500 2000 1500 1000 500
Wavenumbers (cm-1)

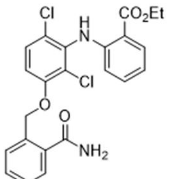
Cl
H
N
CO2Et
Cl
O
O
NH2

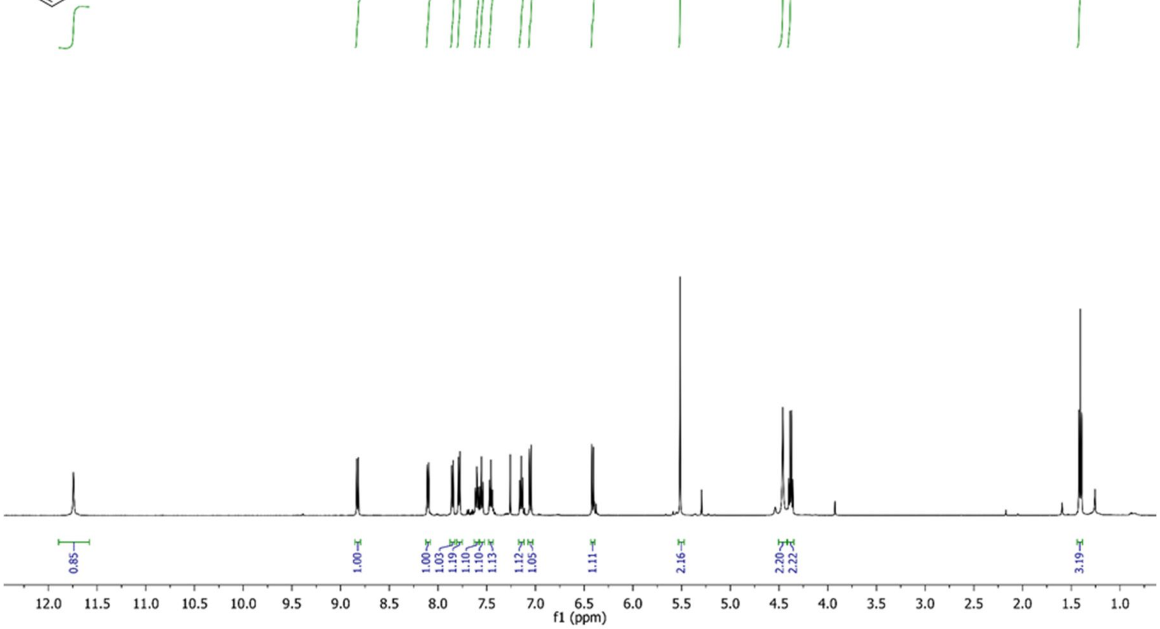
0.85
1.00
1.00
1.03
1.19
1.10
1.10
1.13
1.12
1.05
1.11
2.16
2.20
2.22
3.19
12.0 11.5 11.0 10.5 10.0 9.5 9.0 8.5 8.0 7.5 7.0 6.5 6.0 5.5 5.0 4.5 4.0 3.5 3.0 2.5 2.0 1.5 1.0
f1 (ppm)

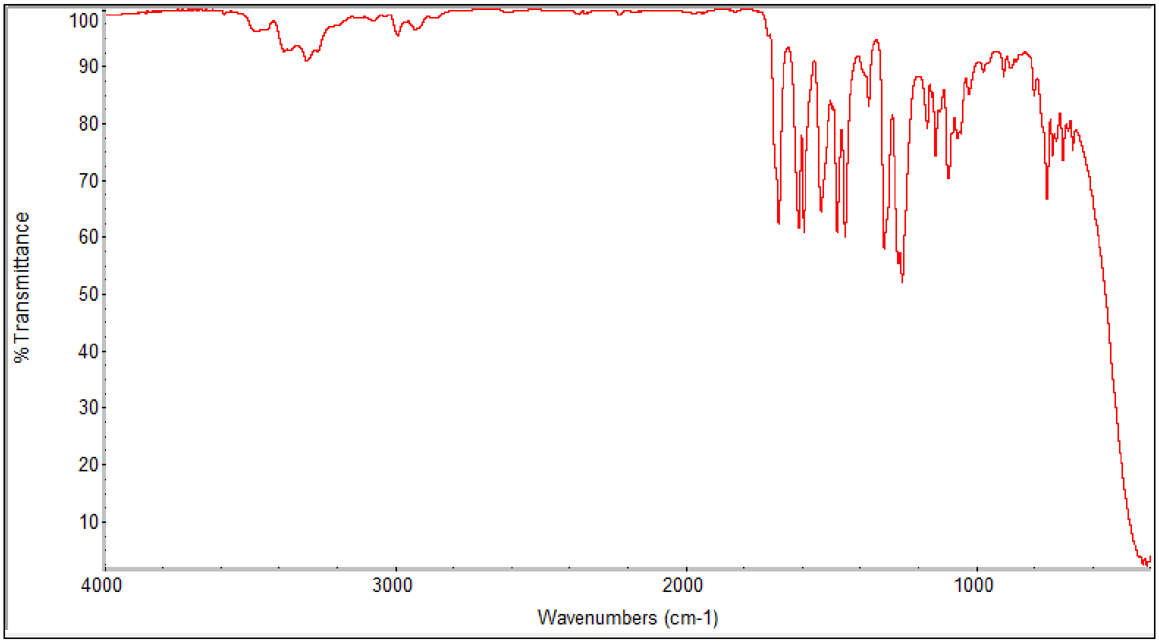
%Transmittance
100
90
80
70
60
50
40
30
20
10
4000
3000
2000
1000
Wavenumbers (cm-1)

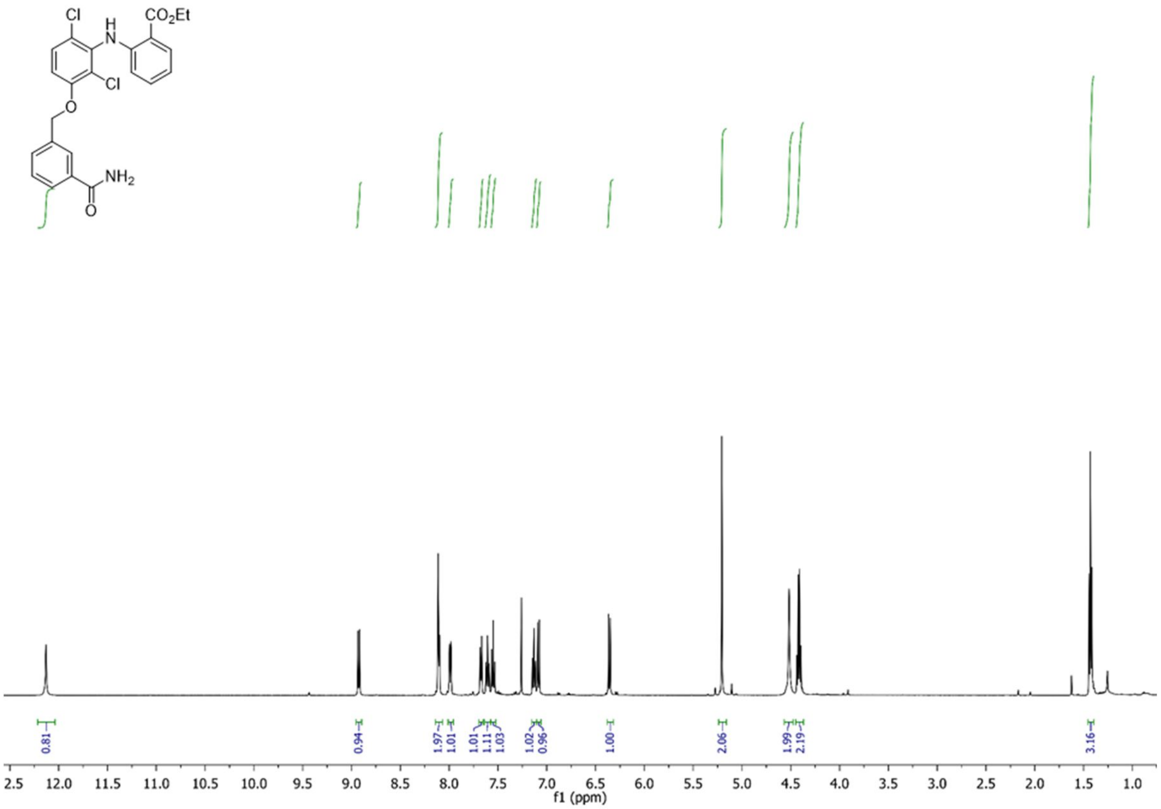
Cl
H
N
CO2Et
Cl
O
NH2
O
f1 (ppm)

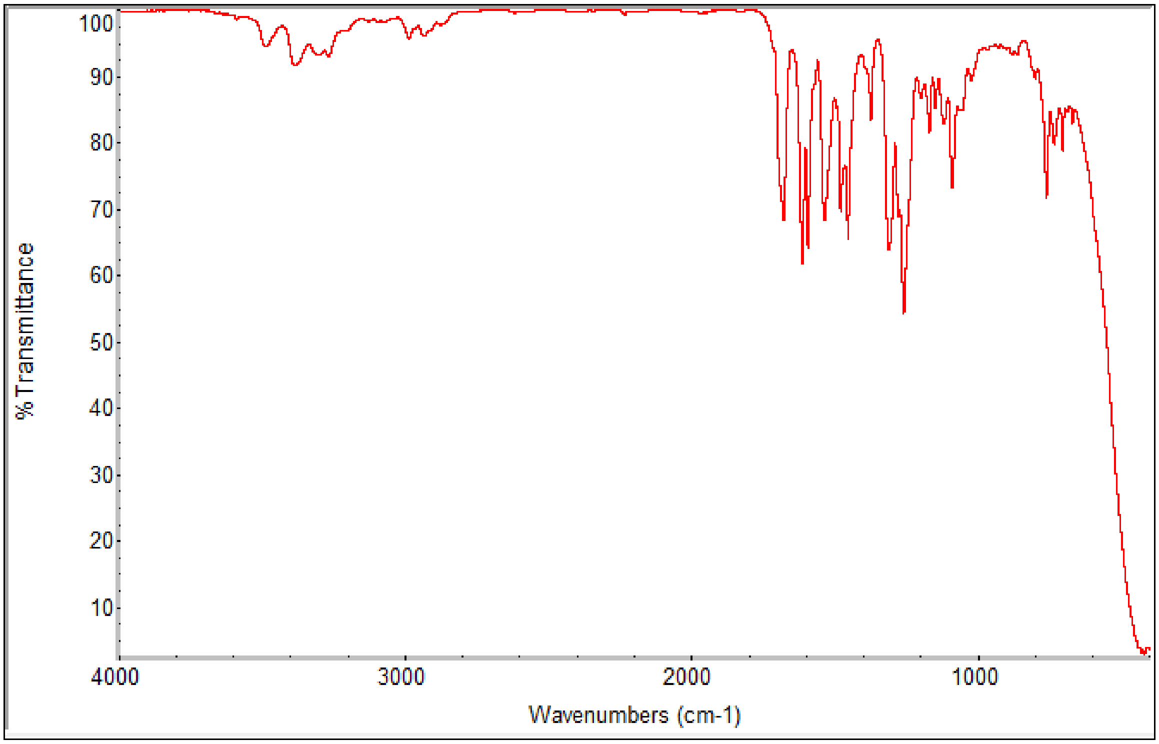
% Transmittance
Wavenumbers (cm-1)

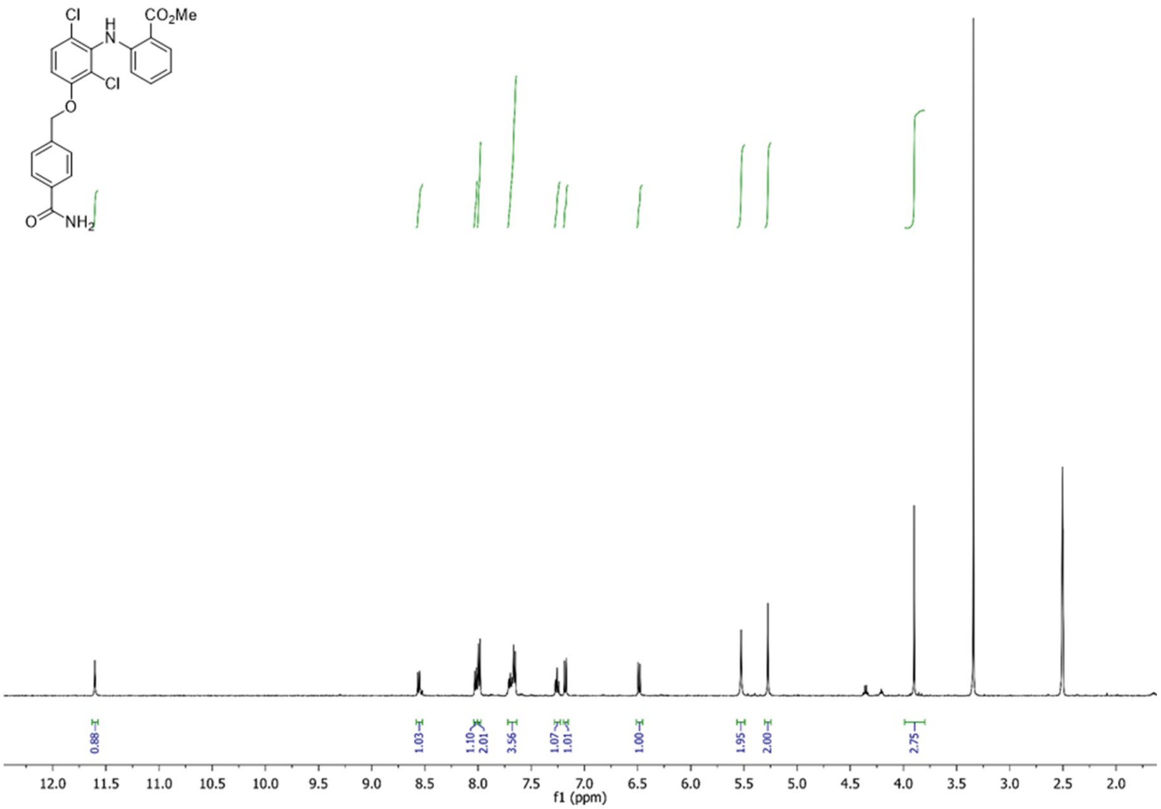

Cl
H
N
CO2Me
Cl
O
O
NH2
0.88
1.03
1.10
2.01
3.56
1.07
1.01
1.00
1.95
2.00
2.75
12.0
11.5
11.0
10.5
10.0
9.5
9.0
8.5
8.0
7.5
7.0
6.5
6.0
5.5
5.0
4.5
4.0
3.5
3.0
2.5
2.0
f1 (ppm)

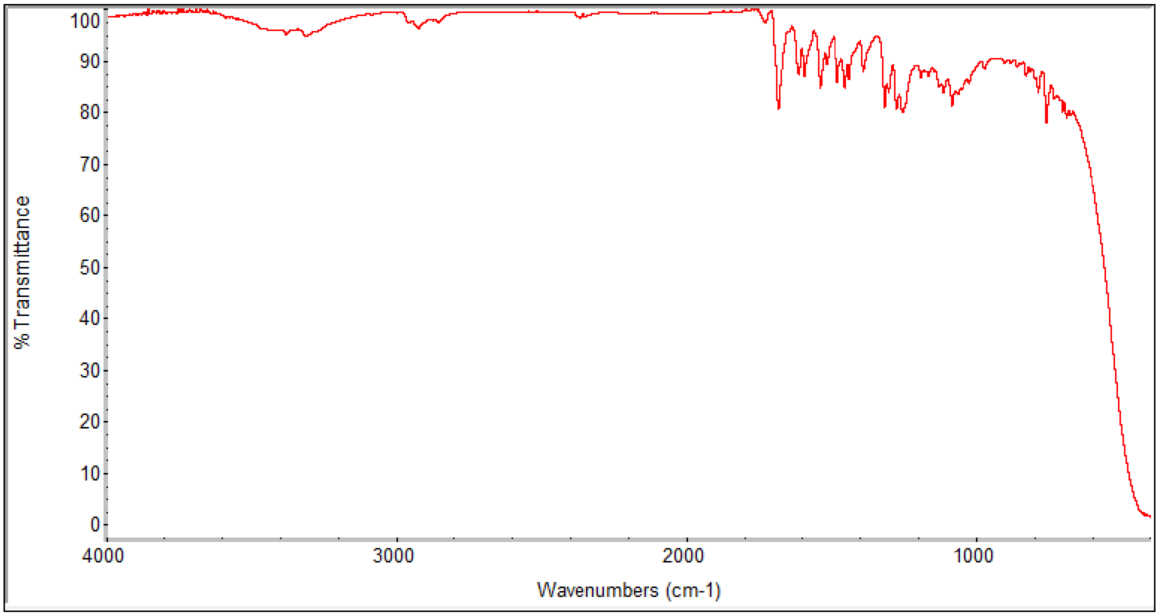

% Transmittance
100
90
80
70
60
50
40
30
20
10
0
4000
3000
2000
1000
Wavenumbers (cm-1)

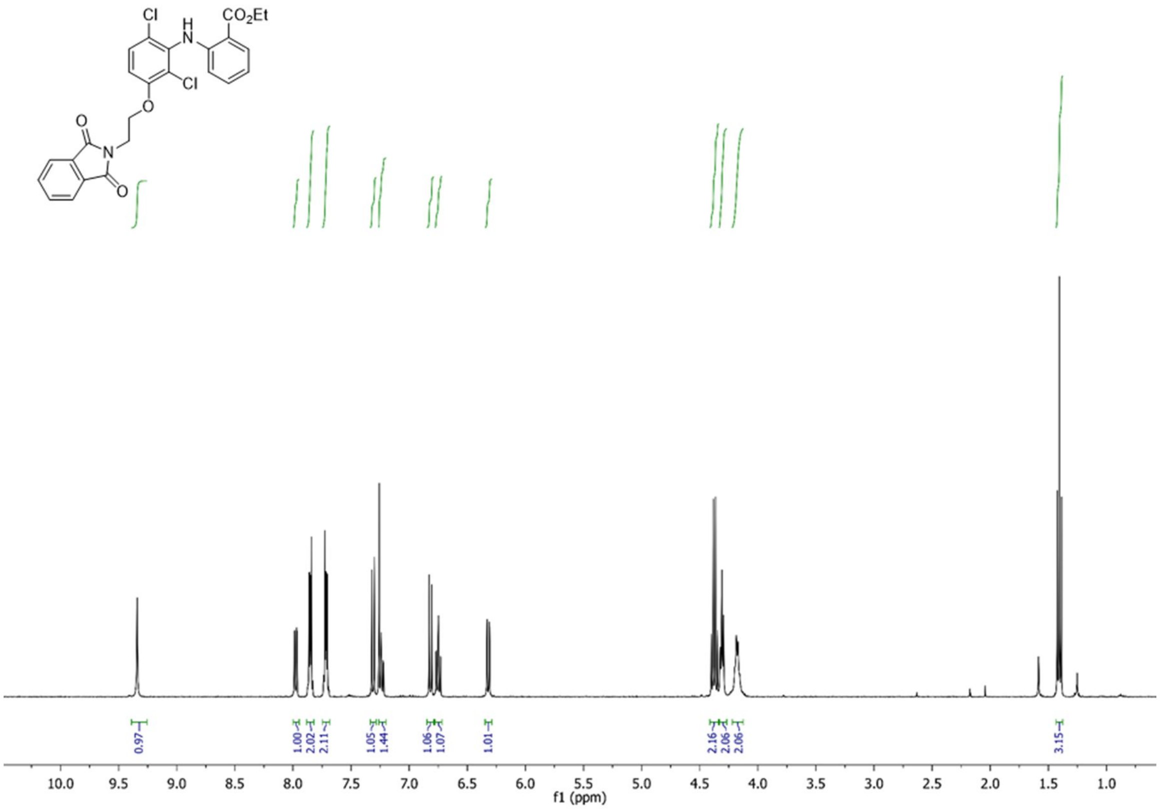

Cl
H
N
CO2Et
Cl
O
O
N
O
0.97
1.00
2.02
2.11
1.05
1.44
1.06
1.07
1.01
2.16
2.06
2.06
3.15
10.0 9.5 9.0 8.5 8.0 7.5 7.0 6.5 6.0 5.5 5.0 4.5 4.0 3.5 3.0 2.5 2.0 1.5 1.0
f1 (ppm)

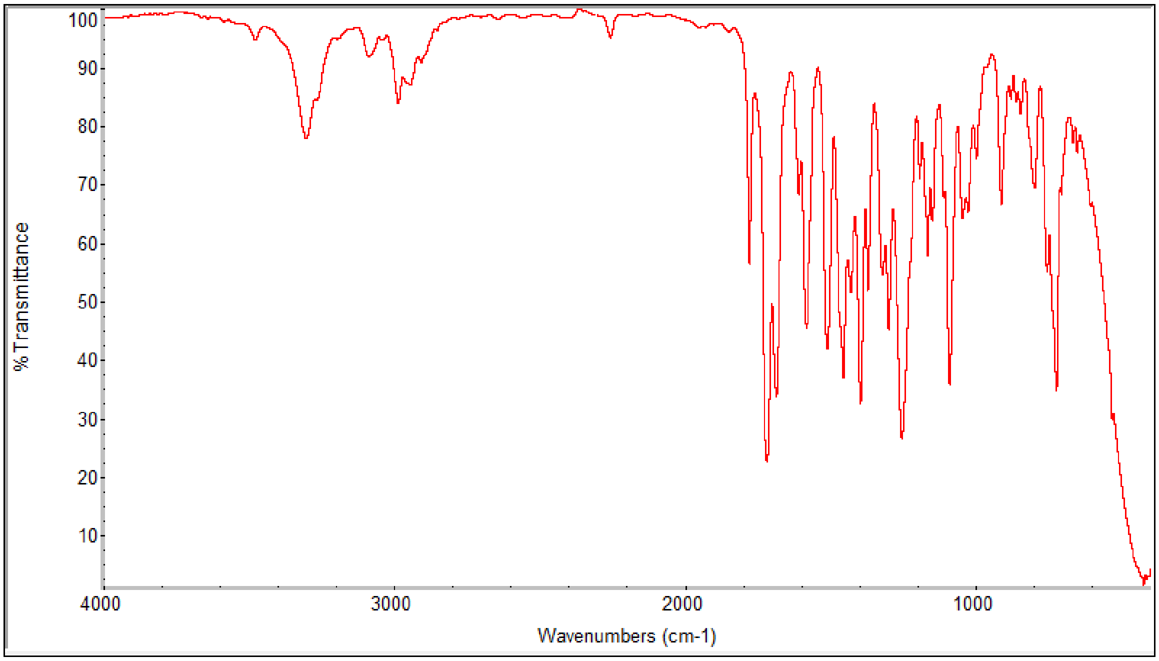

%Transmittance
100 90 80 70 60 50 40 30 20 10
4000 3000 2000 1000
Wavenumbers (cm-1)

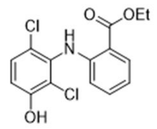

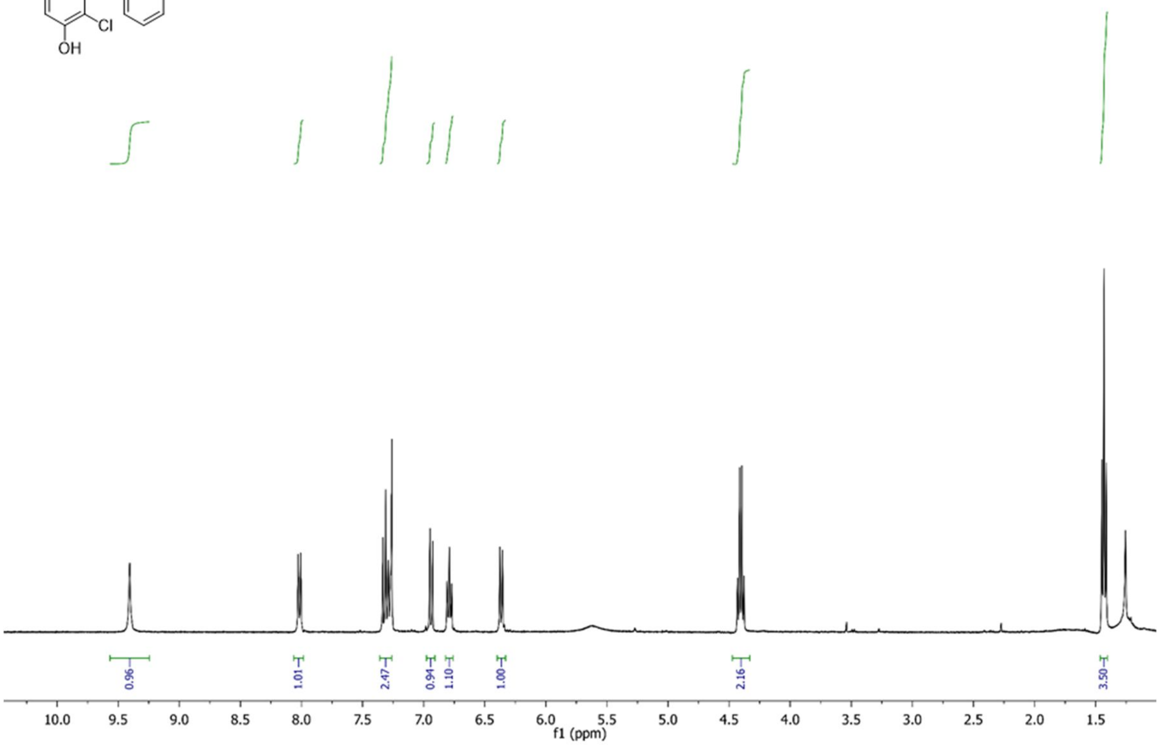
0.96
1.01
2.47
0.94
1.10
1.00
2.16
3.50
10.0
9.5
9.0
8.5
8.0
7.5
7.0
6.5
6.0
5.5
5.0
4.5
4.0
3.5
3.0
2.5
2.0
1.5
f1 (ppm)

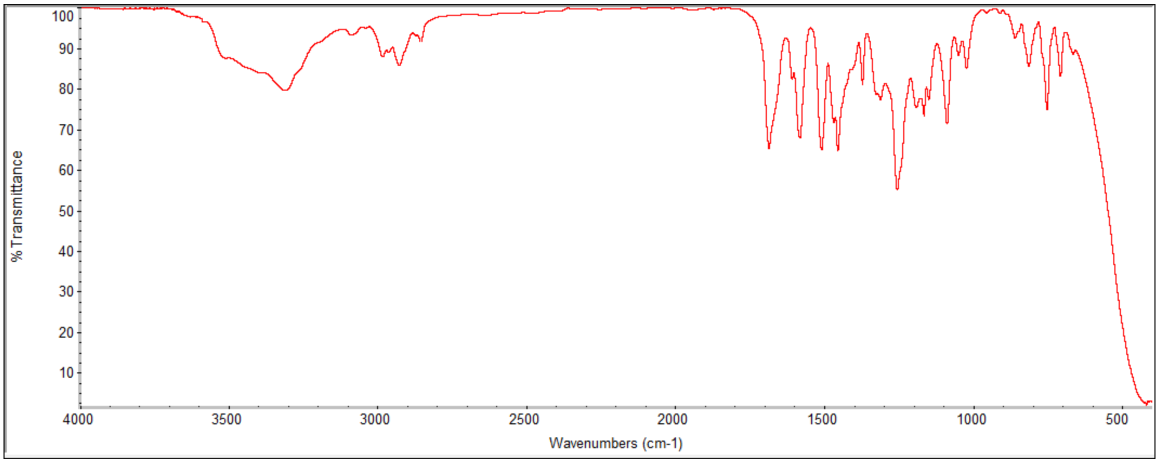
% Transmittance
Wavenumbers (cm-1)
4000
3500
3000
2500
2000
1500
1000
500

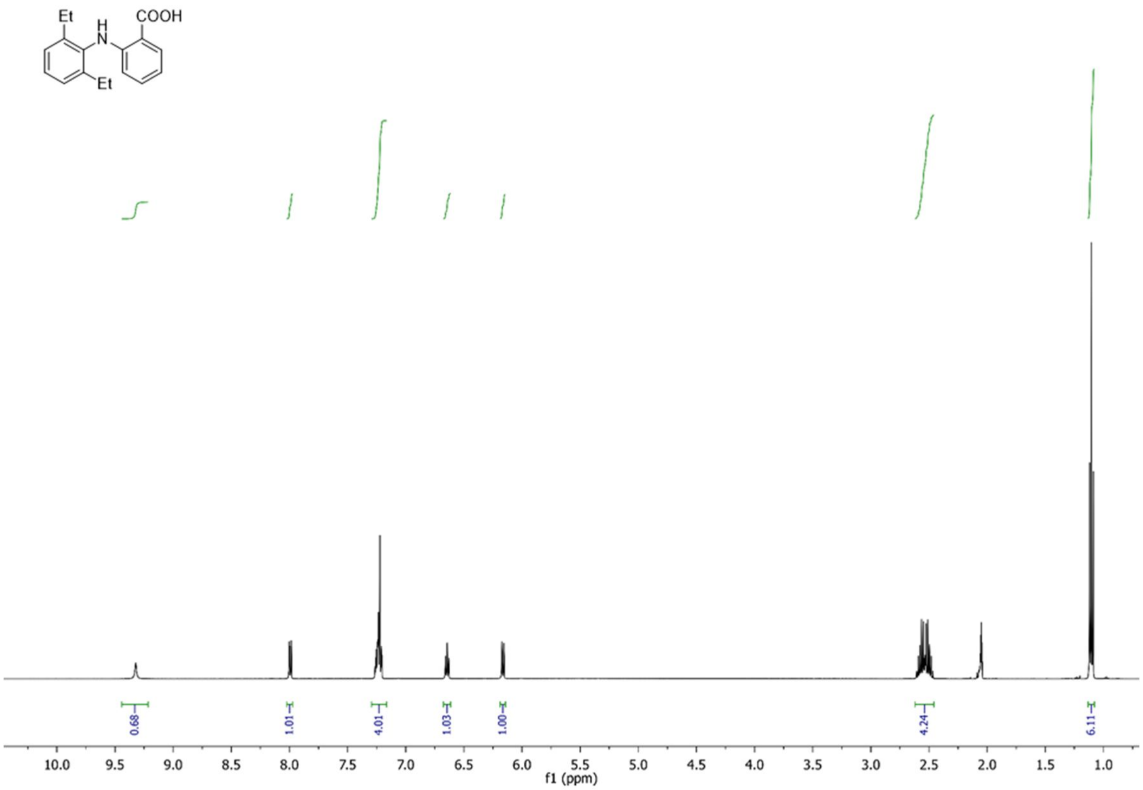
Et
H
N
COOH
Et
0.68
1.01
4.01
1.03
1.00
4.24
6.11
10.0 9.5 9.0 8.5 8.0 7.5 7.0 6.5 6.0 5.5 5.0 4.5 4.0 3.5 3.0 2.5 2.0 1.5 1.0
f1 (ppm)

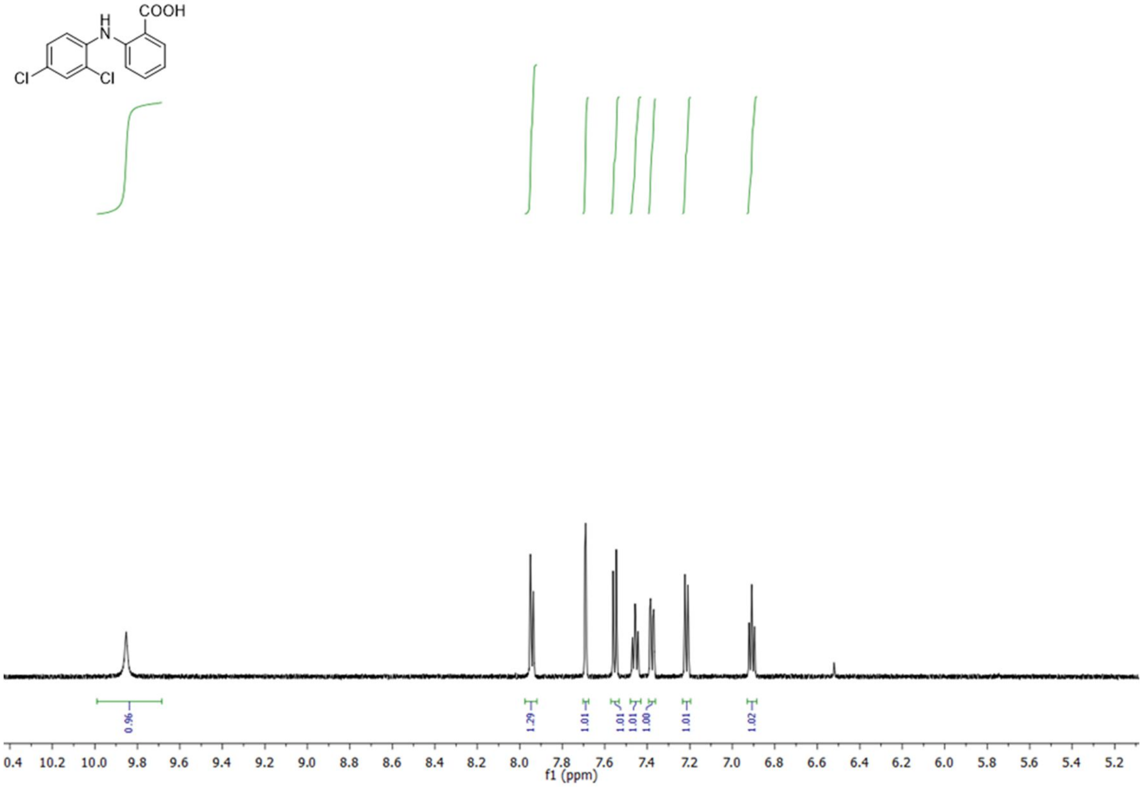
H
N
COOH
Cl
Cl
0.96
1.29
1.01
1.01
1.01
1.00
1.01
1.02
0.4 10.2 10.0 9.8 9.6 9.4 9.2 9.0 8.8 8.6 8.4 8.2 8.0 7.8 7.6 7.4 7.2 7.0 6.8 6.6 6.4 6.2 6.0 5.8 5.6 5.4 5.2
f1 (ppm)

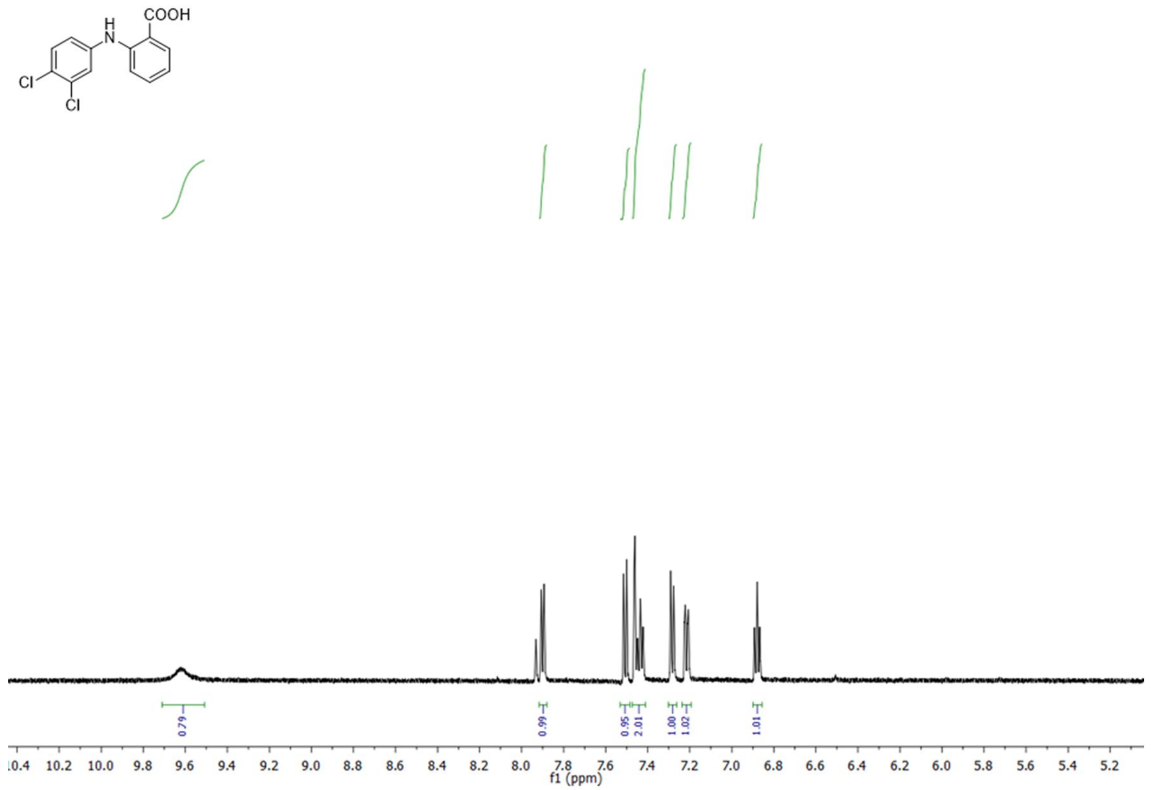
COOH
H
N
Cl
Cl
0.79
0.99
0.95
2.01
1.00
1.02
1.01
10.4 10.2 10.0 9.8 9.6 9.4 9.2 9.0 8.8 8.6 8.4 8.2 8.0 7.8 7.6 7.4 7.2 7.0 6.8 6.6 6.4 6.2 6.0 5.8 5.6 5.4 5.2
f1 (ppm)

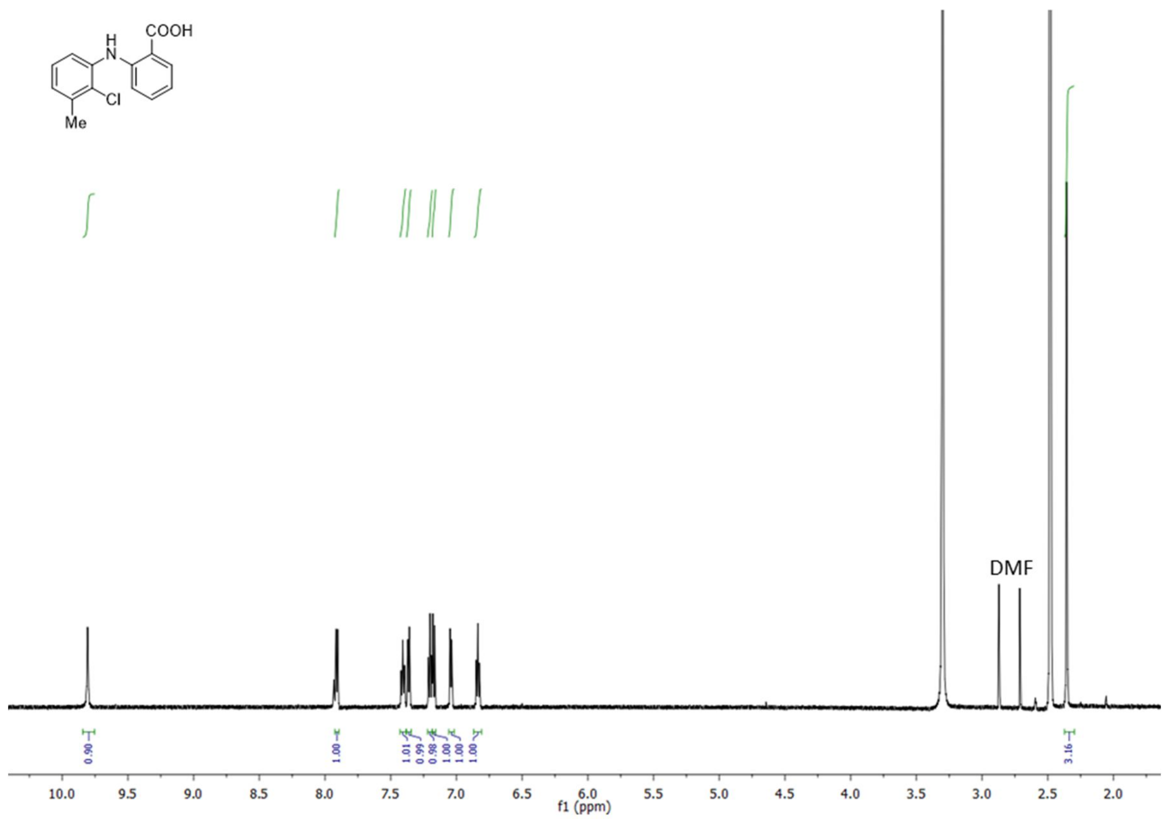
COOH
H
N
Cl
Me
DMF
0.90
1.00
1.01
0.99
0.98
1.00
1.00
3.16
10.0 9.5 9.0 8.5 8.0 7.5 7.0 6.5 6.0 5.5 5.0 4.5 4.0 3.5 3.0 2.5 2.0
f1 (ppm)

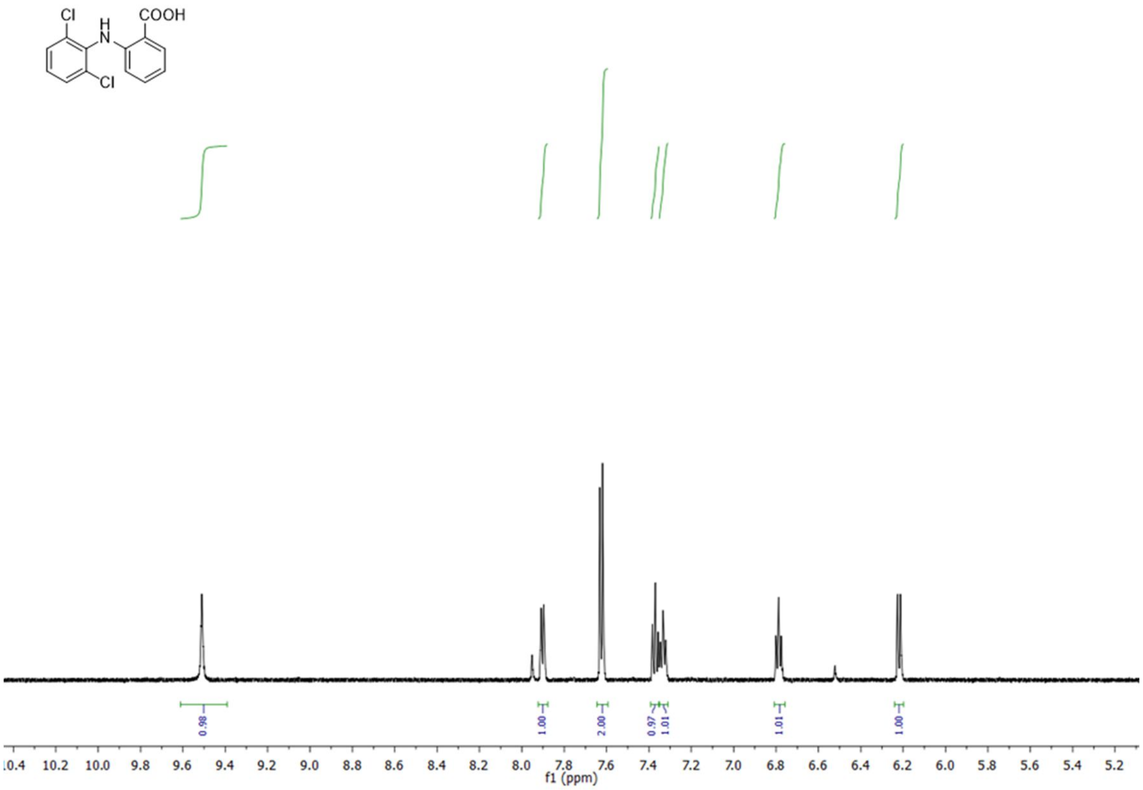

Cl
H
N
COOH
Cl
0.98
1.00
2.00
0.97
1.01
1.01
1.00
10.4 10.2 10.0 9.8 9.6 9.4 9.2 9.0 8.8 8.6 8.4 8.2 8.0 7.8 7.6 7.4 7.2 7.0 6.8 6.6 6.4 6.2 6.0 5.8 5.6 5.4 5.2
f1 (ppm)

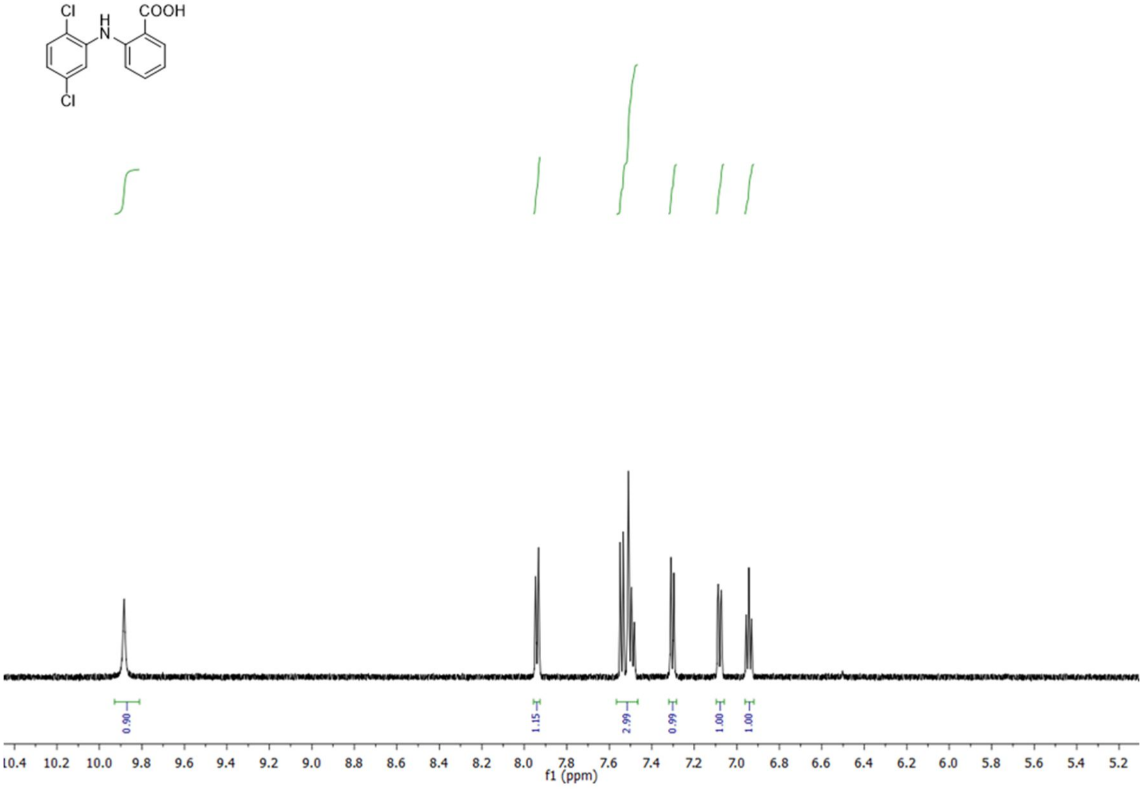

Cl
H
N
COOH
Cl
0.90
1.15
2.99
0.99
1.00
1.00
10.4 10.2 10.0 9.8 9.6 9.4 9.2 9.0 8.8 8.6 8.4 8.2 8.0 7.8 7.6 7.4 7.2 7.0 6.8 6.6 6.4 6.2 6.0 5.8 5.6 5.4 5.2
f1 (ppm)

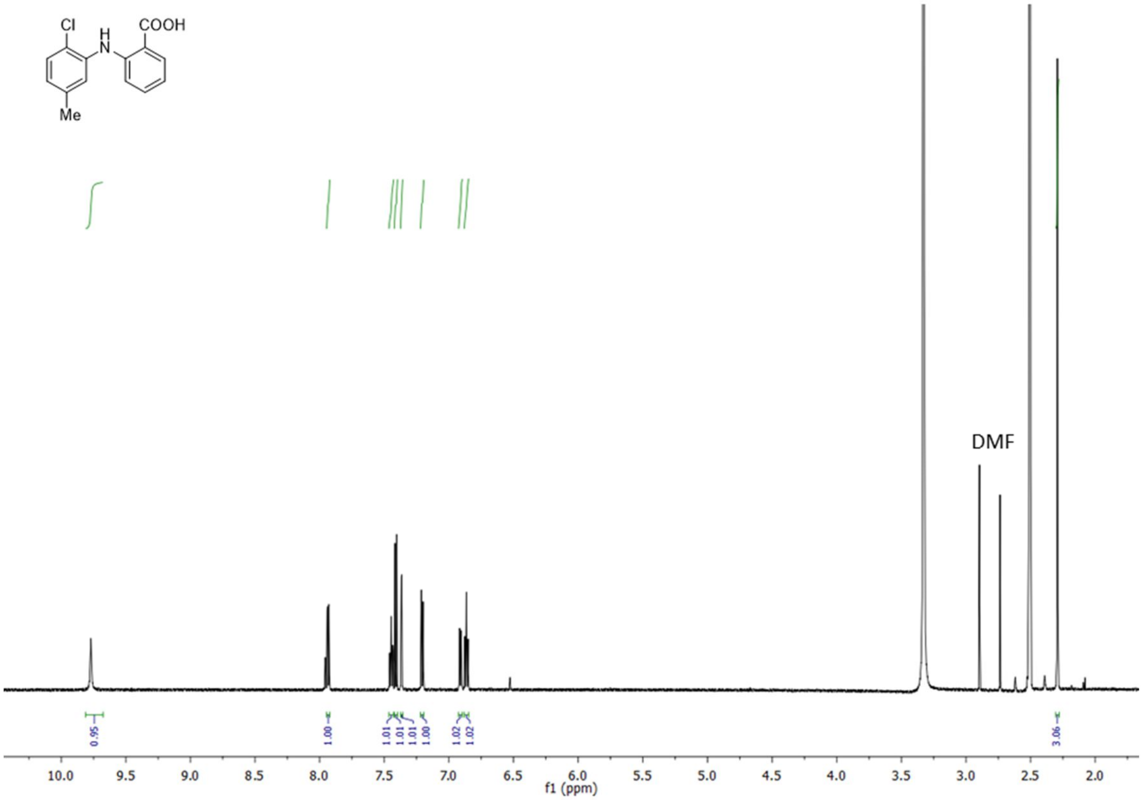
Cl
COOH
H
N
Me
DMF
0.95
1.00
1.01
1.01
1.01
1.00
1.02
1.02
3.06
f1 (ppm)

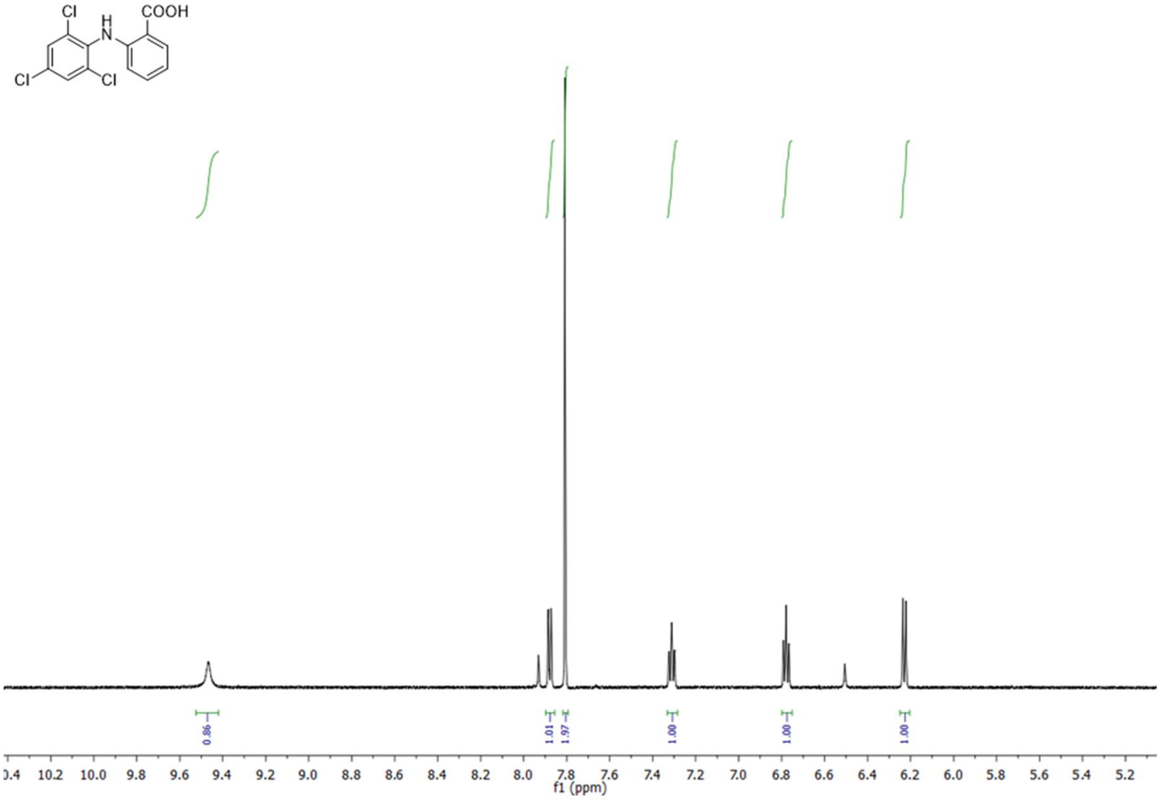
Cl
H
N
COOH
Cl
Cl
0.86
1.01
1.97
1.00
1.00
1.00
f1 (ppm)

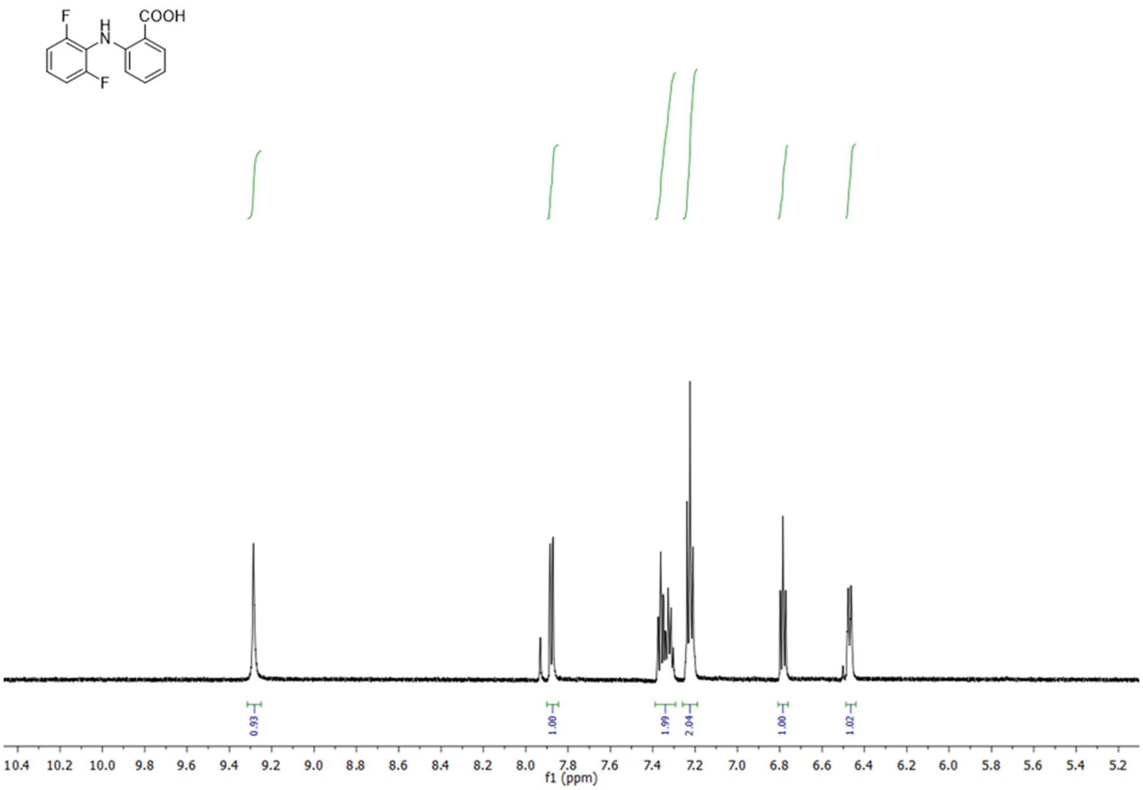
COOH
F
H
N
F
0.93
1.00
1.99
2.04
1.00
1.02
10.4
10.2
10.0
9.8
9.6
9.4
9.2
9.0
8.8
8.6
8.4
8.2
8.0
7.8
7.6
7.4
7.2
7.0
6.8
6.6
6.4
6.2
6.0
5.8
5.6
5.4
5.2
f1 (ppm)

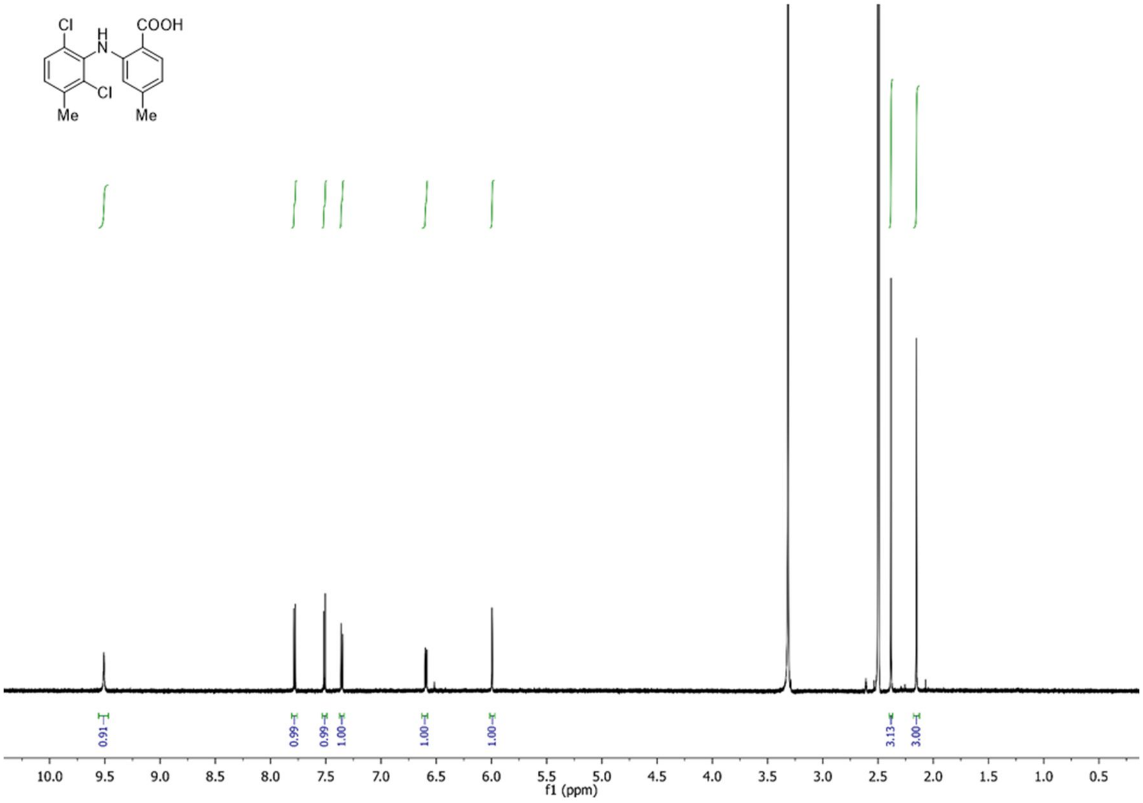
Cl
H
N
COOH
Cl
Me
Me
0.91
0.99
0.99
1.00
1.00
1.00
3.13
3.00
10.0
9.5
9.0
8.5
8.0
7.5
7.0
6.5
6.0
5.5
5.0
4.5
4.0
3.5
3.0
2.5
2.0
1.5
1.0
0.5
f1 (ppm)

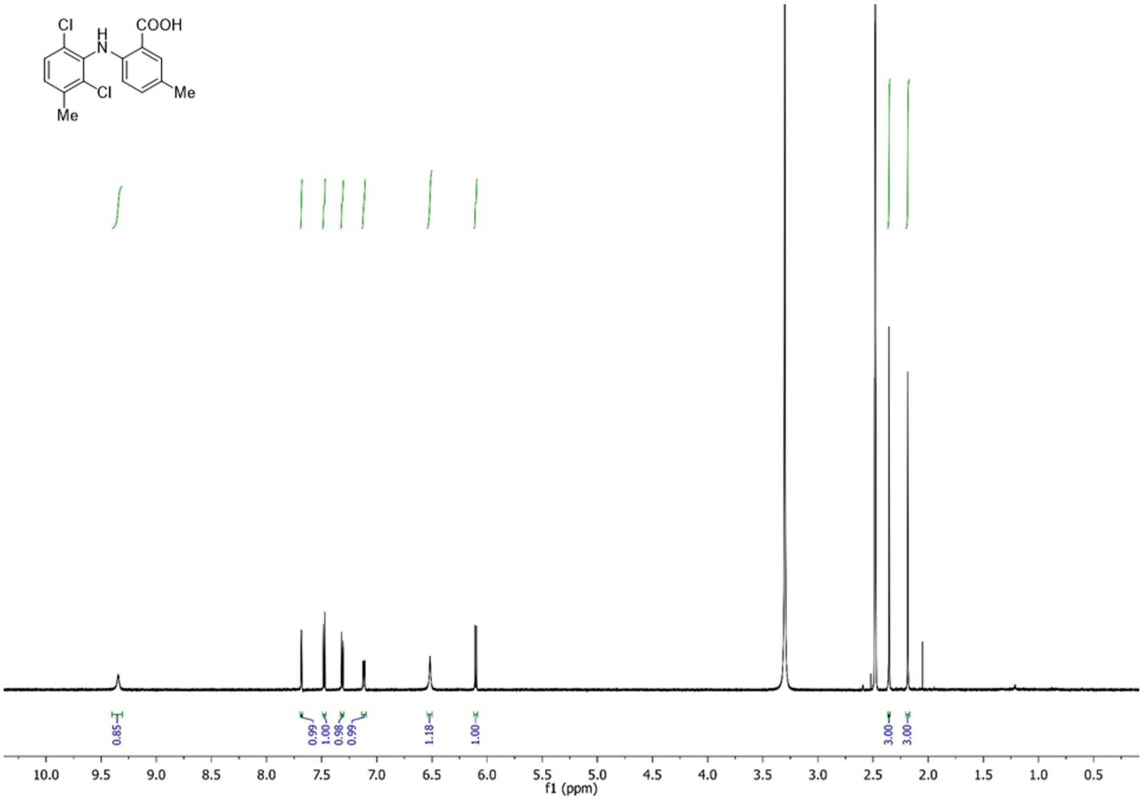
Cl
H
N
COOH
Cl
Me
Me
0.85
0.99
1.00
0.98
0.99
1.18
1.00
3.00
3.00
10.0 9.5 9.0 8.5 8.0 7.5 7.0 6.5 6.0 5.5 5.0 4.5 4.0 3.5 3.0 2.5 2.0 1.5 1.0 0.5
f1 (ppm)

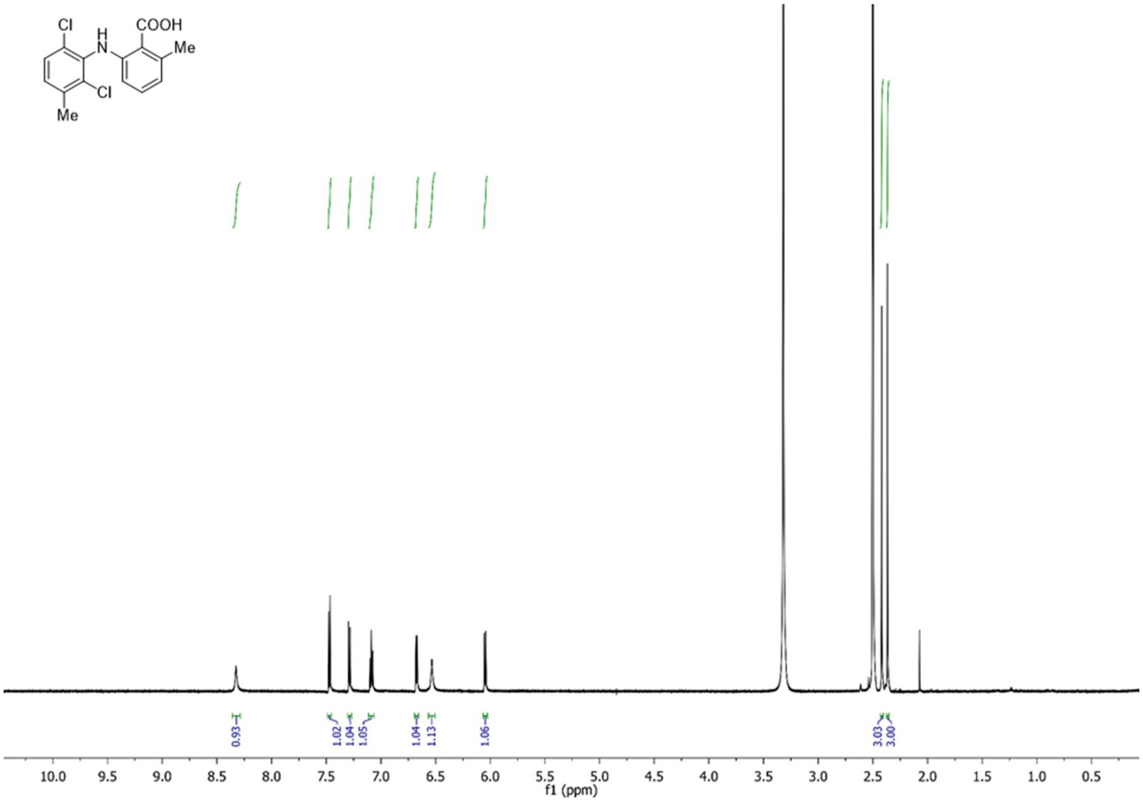
Cl
H
N
COOH
Me
Cl
Me
0.93
1.02
1.04
1.05
1.04
1.13
1.06
3.03
3.00
10.0 9.5 9.0 8.5 8.0 7.5 7.0 6.5 6.0 5.5 5.0 4.5 4.0 3.5 3.0 2.5 2.0 1.5 1.0 0.5
f1 (ppm)

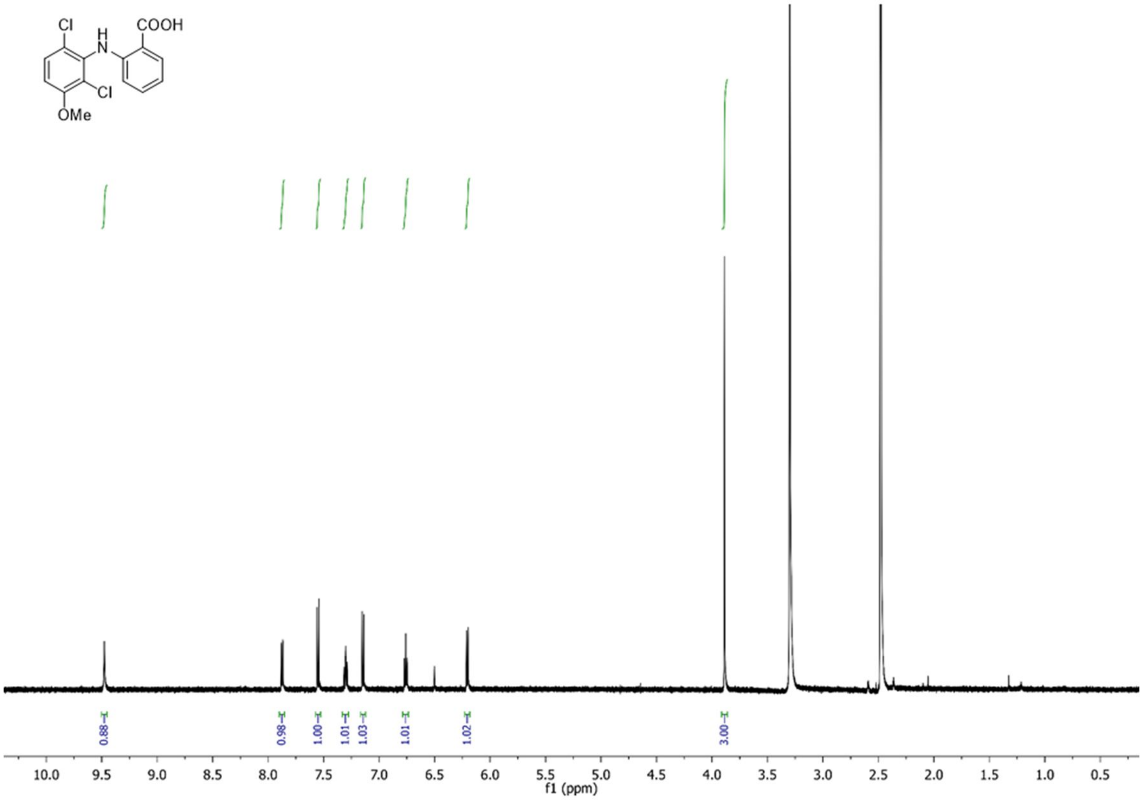

COOH
Cl
H
N
Cl
OMe
0.88
0.98
1.00
1.01
1.03
1.01
1.02
3.00
10.0 9.5 9.0 8.5 8.0 7.5 7.0 6.5 6.0 5.5 5.0 4.5 4.0 3.5 3.0 2.5 2.0 1.5 1.0 0.5
f1 (ppm)

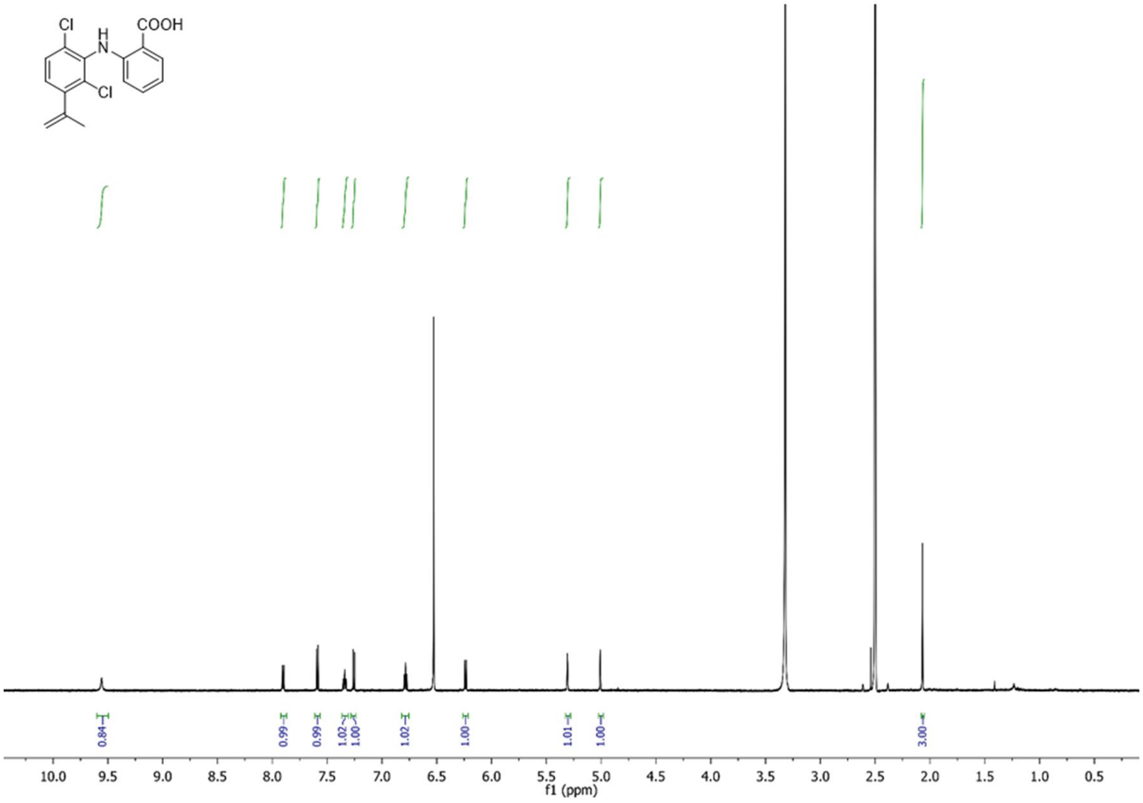

COOH
Cl
H
N
Cl
0.84
0.99
0.99
1.02
1.00
1.02
1.00
1.01
1.00
3.00
10.0 9.5 9.0 8.5 8.0 7.5 7.0 6.5 6.0 5.5 5.0 4.5 4.0 3.5 3.0 2.5 2.0 1.5 1.0 0.5
f1 (ppm)

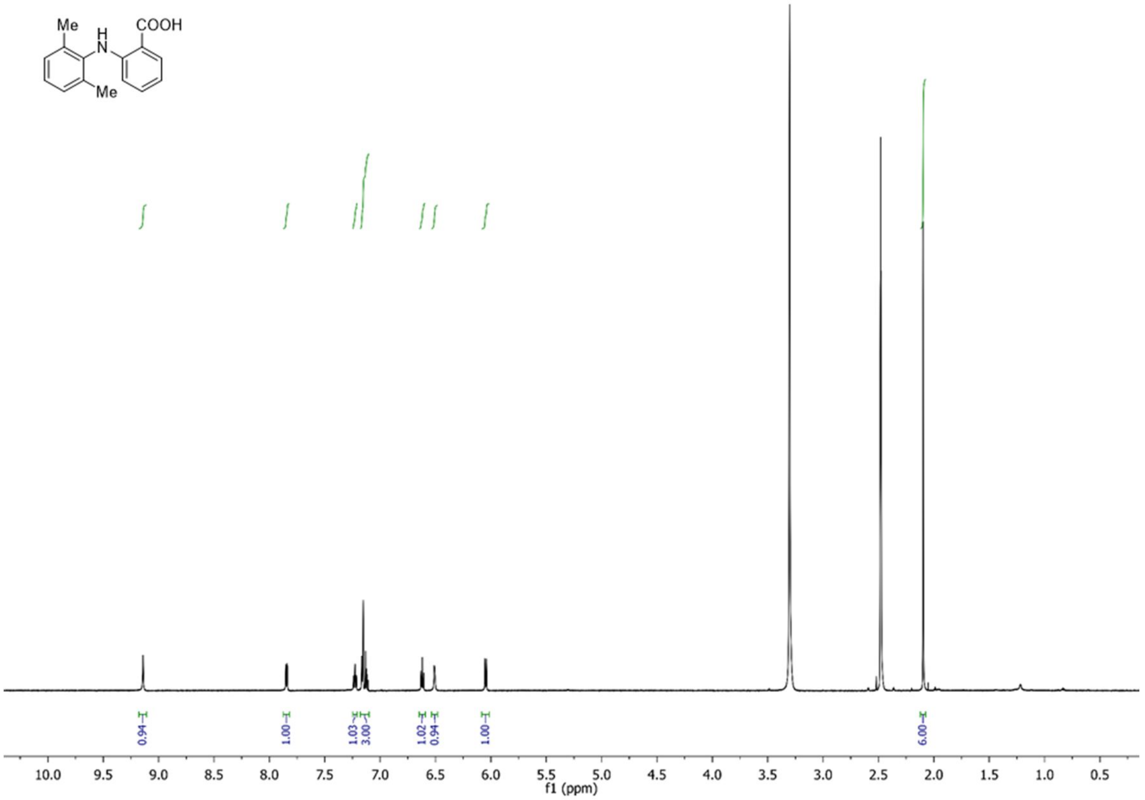
Me
H
N
COOH
Me
0.94
1.00
1.03
3.00
1.02
0.94
1.00
6.00
10.0 9.5 9.0 8.5 8.0 7.5 7.0 6.5 6.0 5.5 5.0 4.5 4.0 3.5 3.0 2.5 2.0 1.5 1.0 0.5
f1 (ppm)

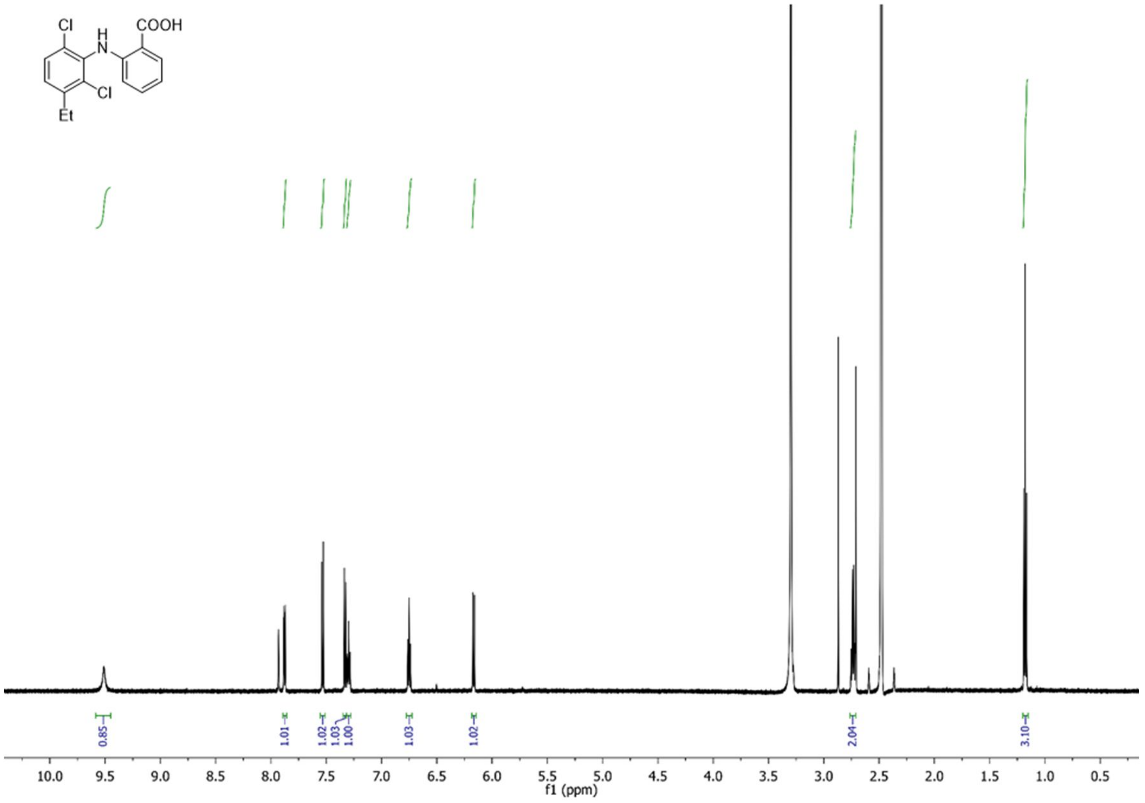
Cl
H
N
COOH
Cl
Et
0.85
1.01
1.02
1.03
1.00
1.03
1.02
2.04
3.10
10.0 9.5 9.0 8.5 8.0 7.5 7.0 6.5 6.0 5.5 5.0 4.5 4.0 3.5 3.0 2.5 2.0 1.5 1.0 0.5
f1 (ppm)

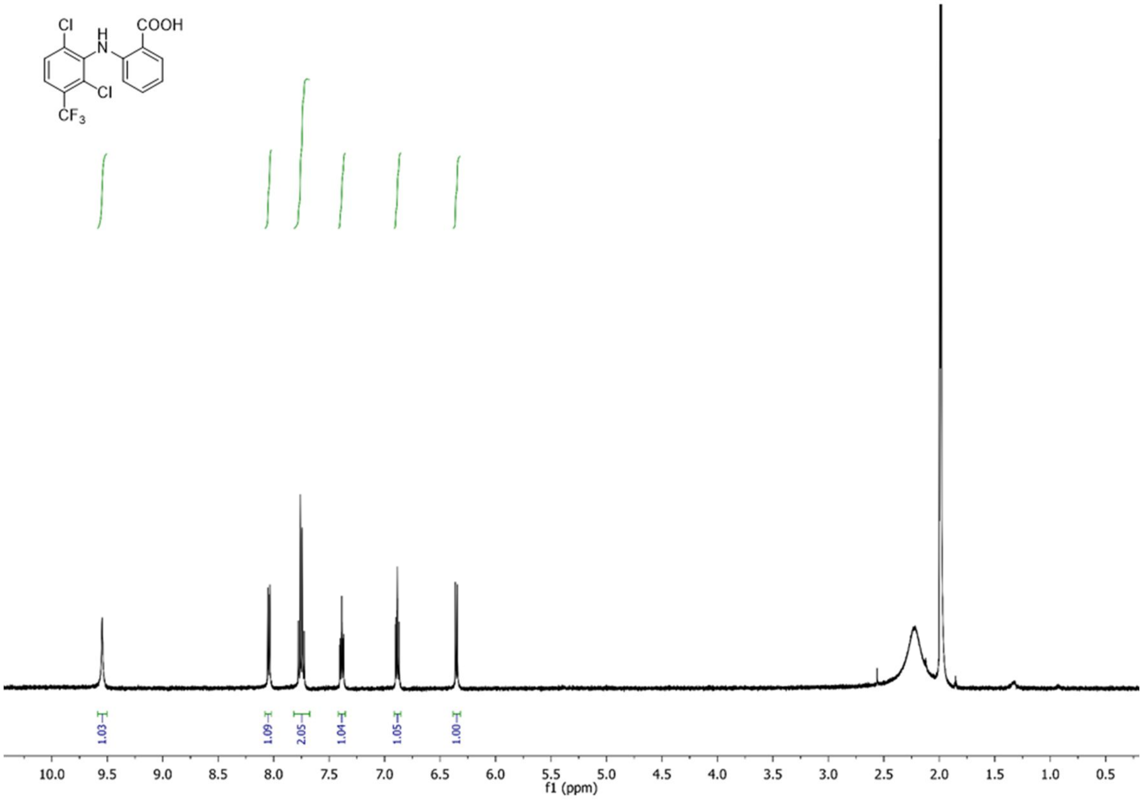
Cl
H
N
COOH
Cl
CF3
1.03
1.09
2.05
1.04
1.05
1.00
10.0 9.5 9.0 8.5 8.0 7.5 7.0 6.5 6.0 5.5 5.0 4.5 4.0 3.5 3.0 2.5 2.0 1.5 1.0 0.5
f1 (ppm)

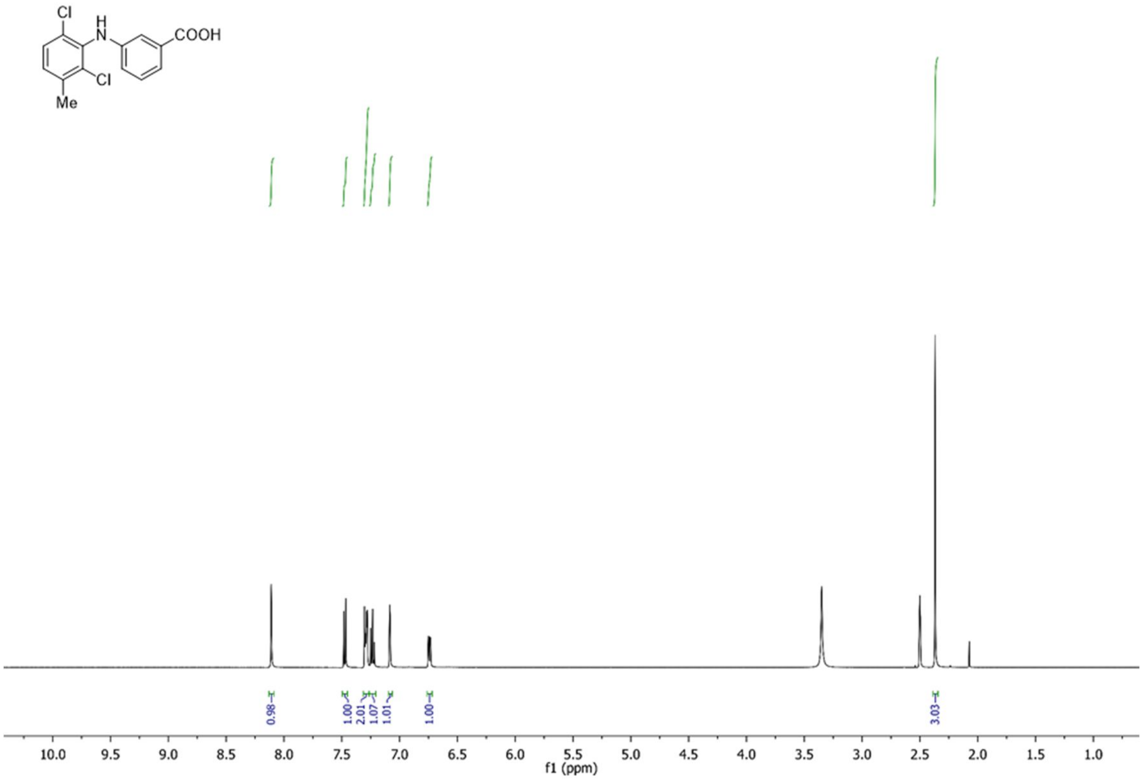
Cl
H
N
COOH
Cl
Me
0.98
1.00
2.01
1.07
1.01
1.00
3.03
10.0 9.5 9.0 8.5 8.0 7.5 7.0 6.5 6.0 5.5 5.0 4.5 4.0 3.5 3.0 2.5 2.0 1.5 1.0
f1 (ppm)

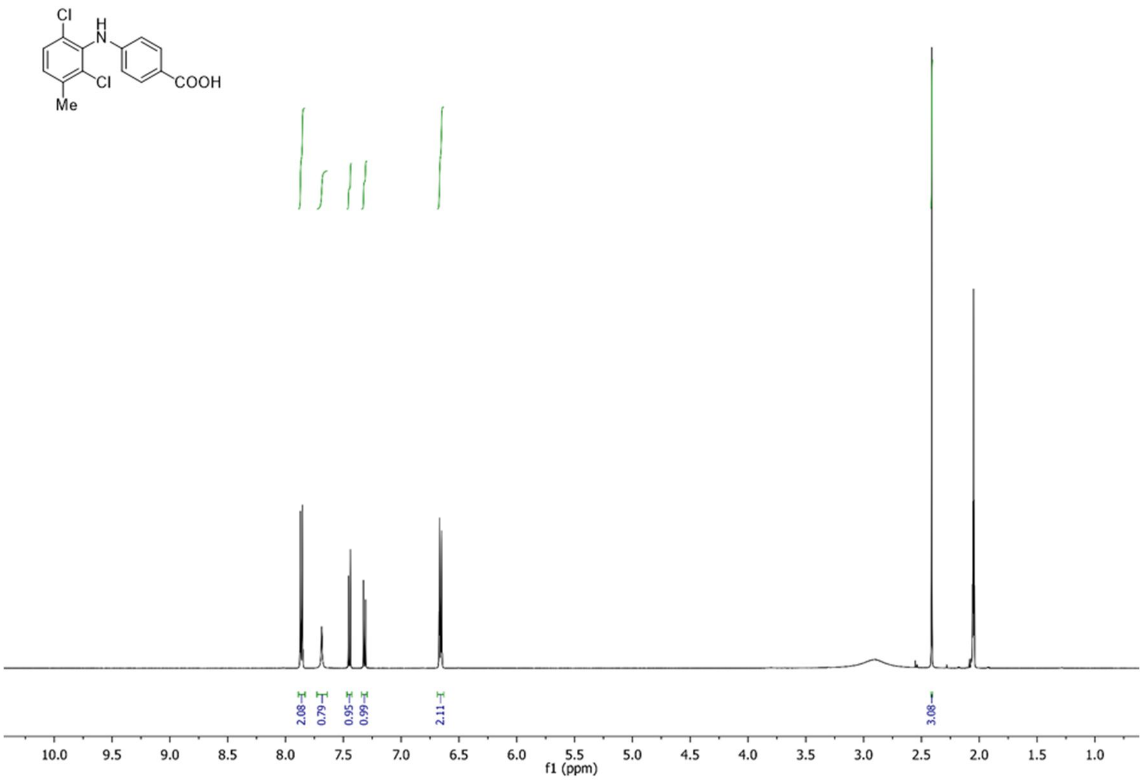

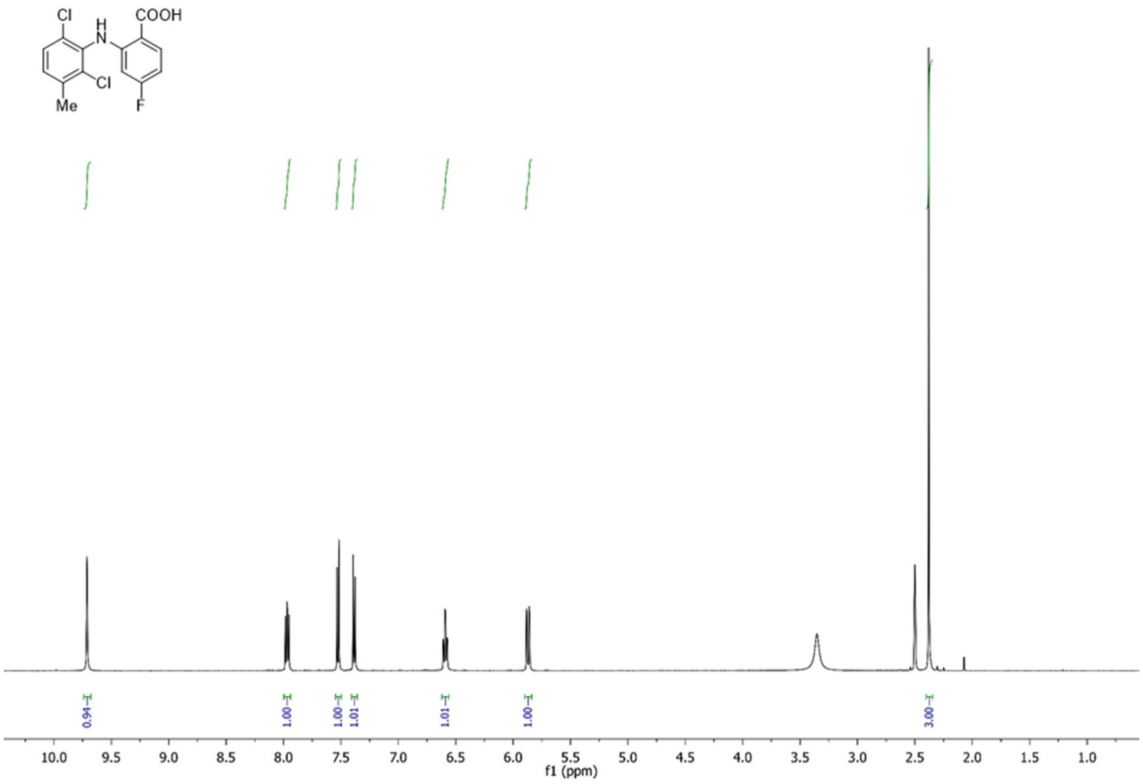

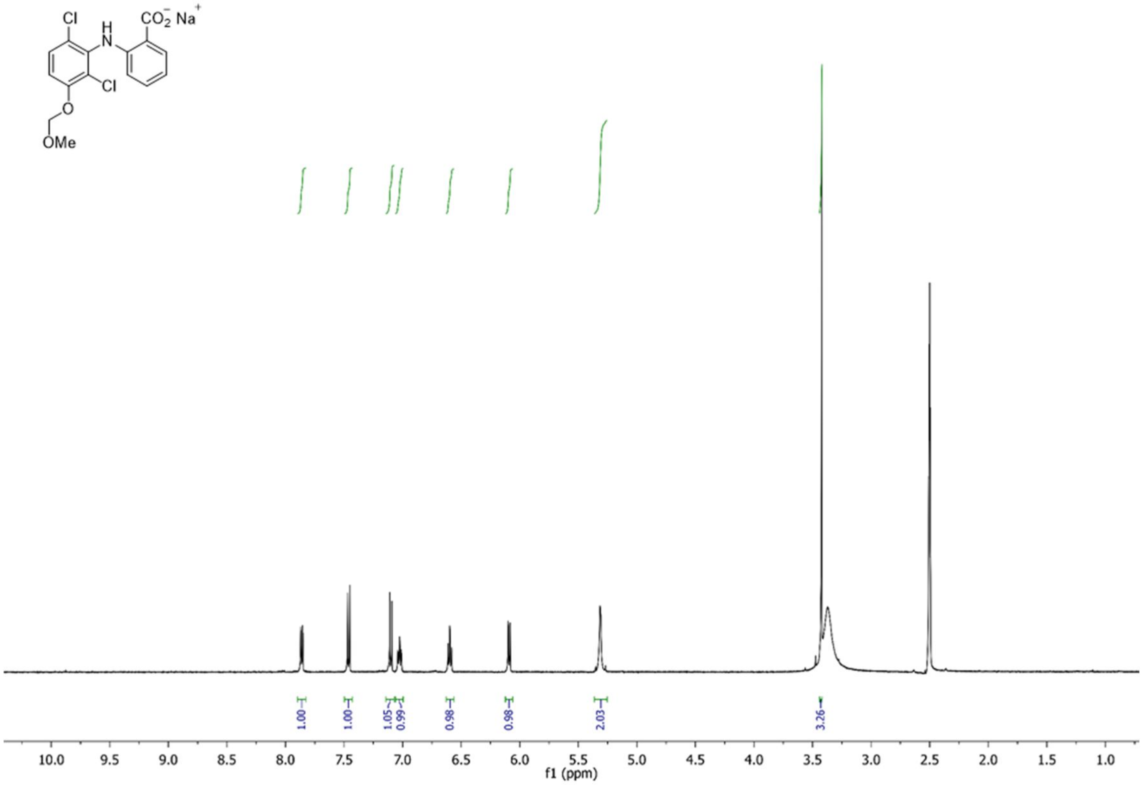
Cl
H
N
$CO_2^-$ $Na^+$
Cl
O
OMe
1.00
1.00
1.05
0.99
0.98
0.98
2.03
3.26
10.0 9.5 9.0 8.5 8.0 7.5 7.0 6.5 6.0 5.5 5.0 4.5 4.0 3.5 3.0 2.5 2.0 1.5 1.0
f1 (ppm)

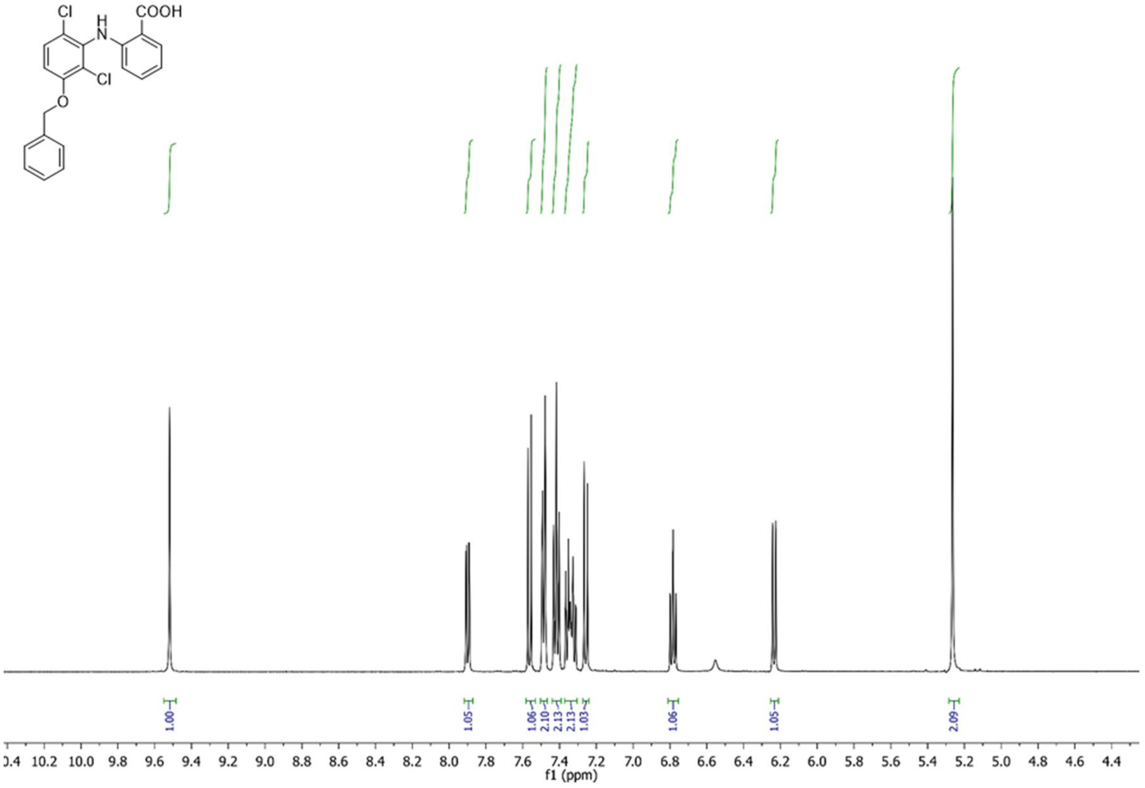
Cl
H
N
COOH
Cl
O
1.00
1.05
1.06
2.10
2.13
2.13
1.03
1.06
1.05
2.09
10.2 10.0 9.8 9.6 9.4 9.2 9.0 8.8 8.6 8.4 8.2 8.0 7.8 7.6 7.4 7.2 7.0 6.8 6.6 6.4 6.2 6.0 5.8 5.6 5.4 5.2 5.0 4.8 4.6 4.4
f1 (ppm)

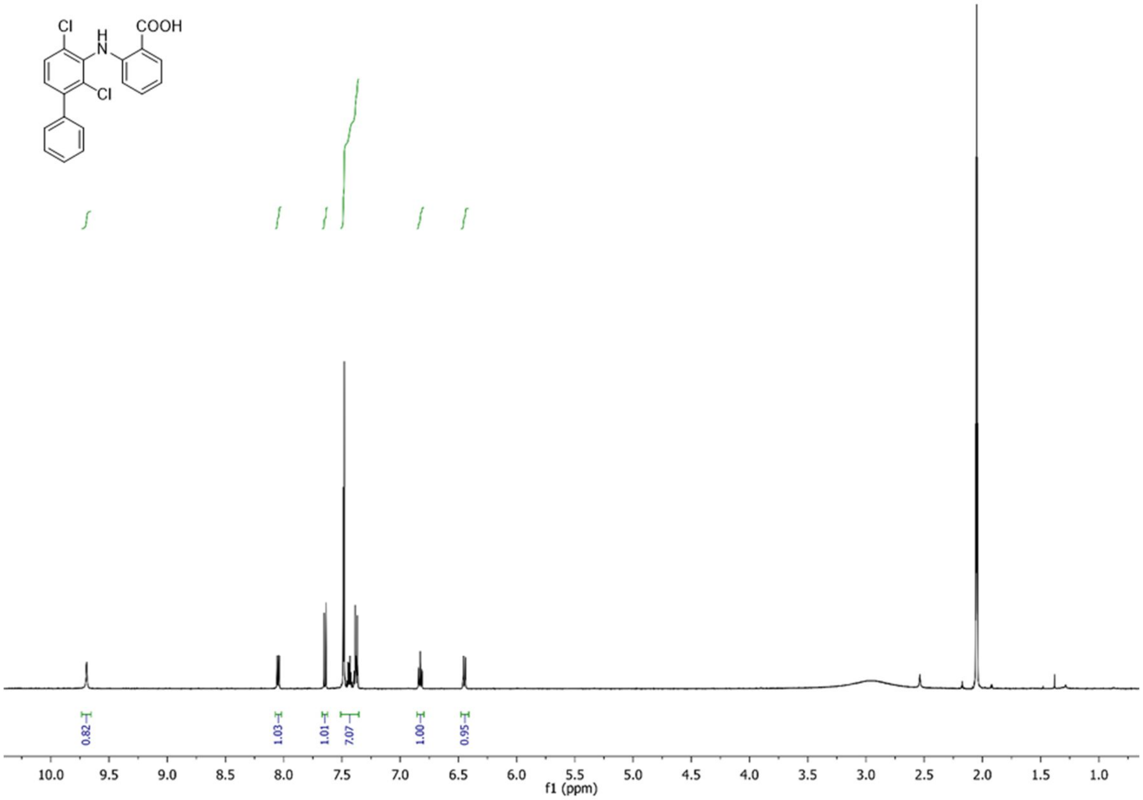
Cl
H
N
COOH
Cl
0.82
1.03
1.01
7.07
1.00
0.95
10.0 9.5 9.0 8.5 8.0 7.5 7.0 6.5 6.0 5.5 5.0 4.5 4.0 3.5 3.0 2.5 2.0 1.5 1.0
f1 (ppm)

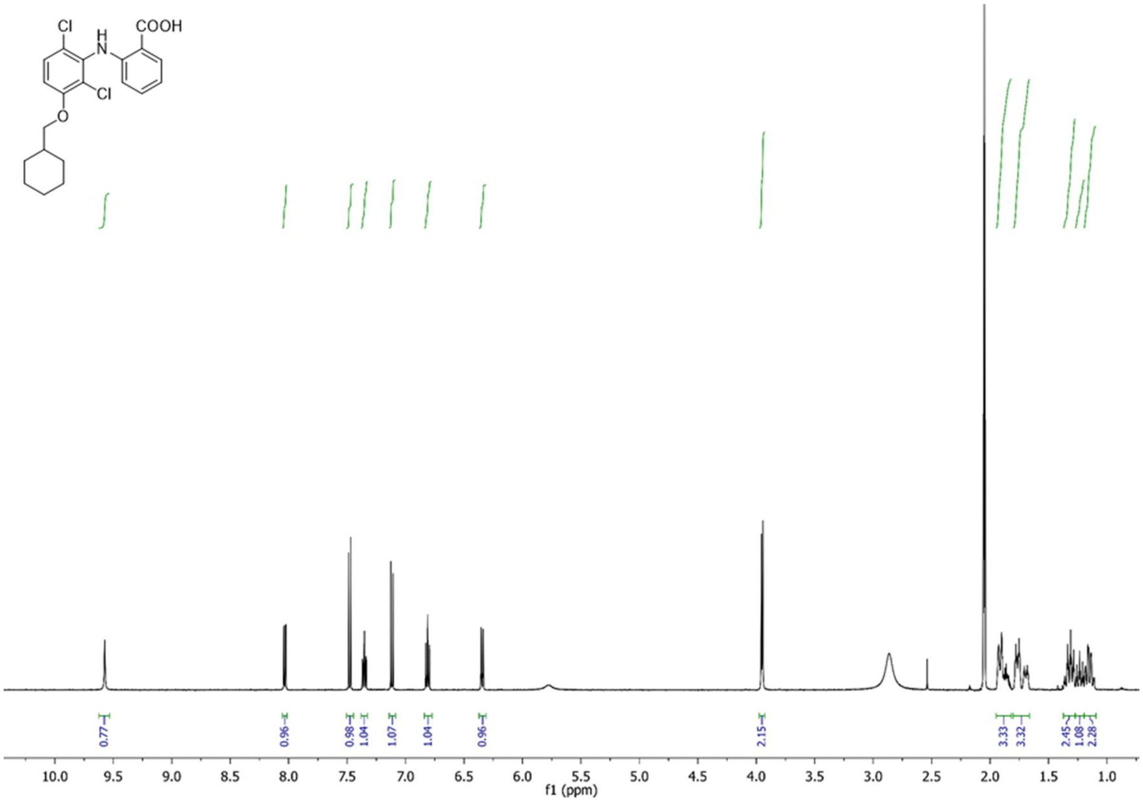
Cl
H
N
COOH
Cl
O
0.77
0.96
0.98
1.04
1.07
1.04
0.96
2.15
3.33
3.32
2.45
1.08
2.28
10.0 9.5 9.0 8.5 8.0 7.5 7.0 6.5 6.0 5.5 5.0 4.5 4.0 3.5 3.0 2.5 2.0 1.5 1.0
f1 (ppm)

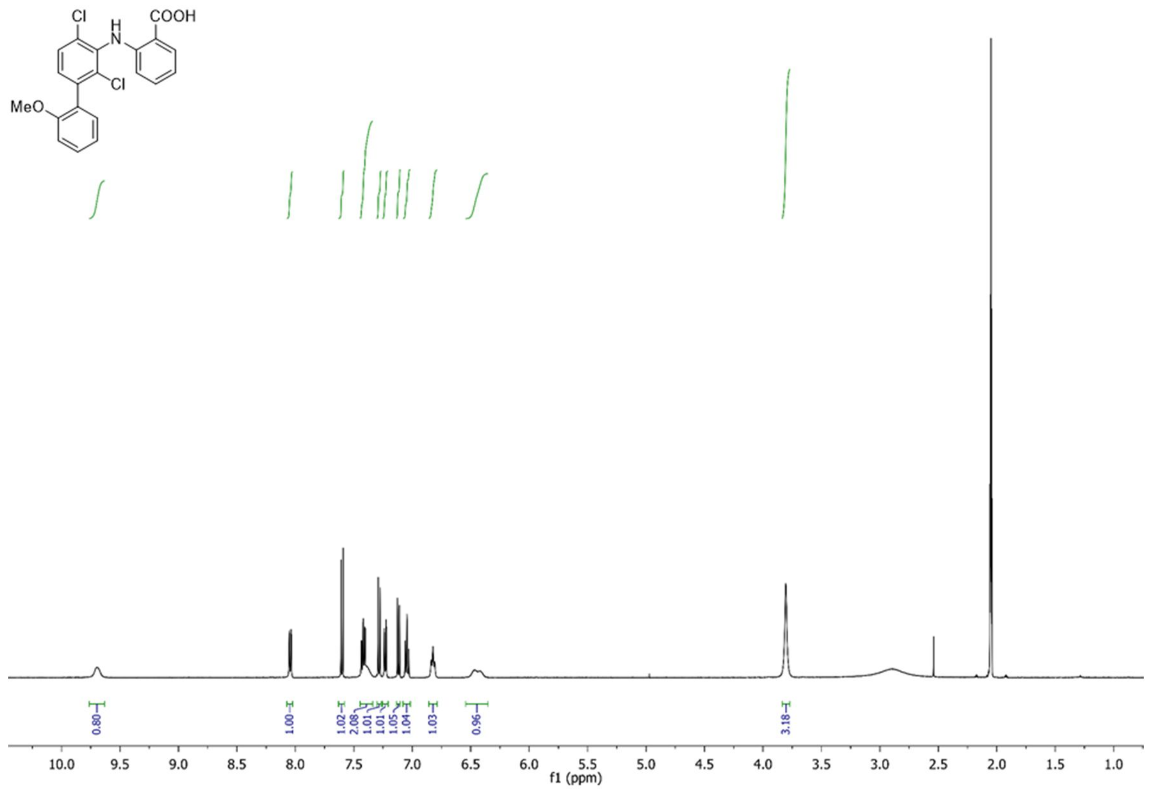

Cl
H
N
COOH
Cl
MeO
0.80
1.00
1.02
2.08
1.01
1.01
1.05
1.04
1.03
0.96
3.18
10.0 9.5 9.0 8.5 8.0 7.5 7.0 6.5 6.0 5.5 5.0 4.5 4.0 3.5 3.0 2.5 2.0 1.5 1.0
f1 (ppm)

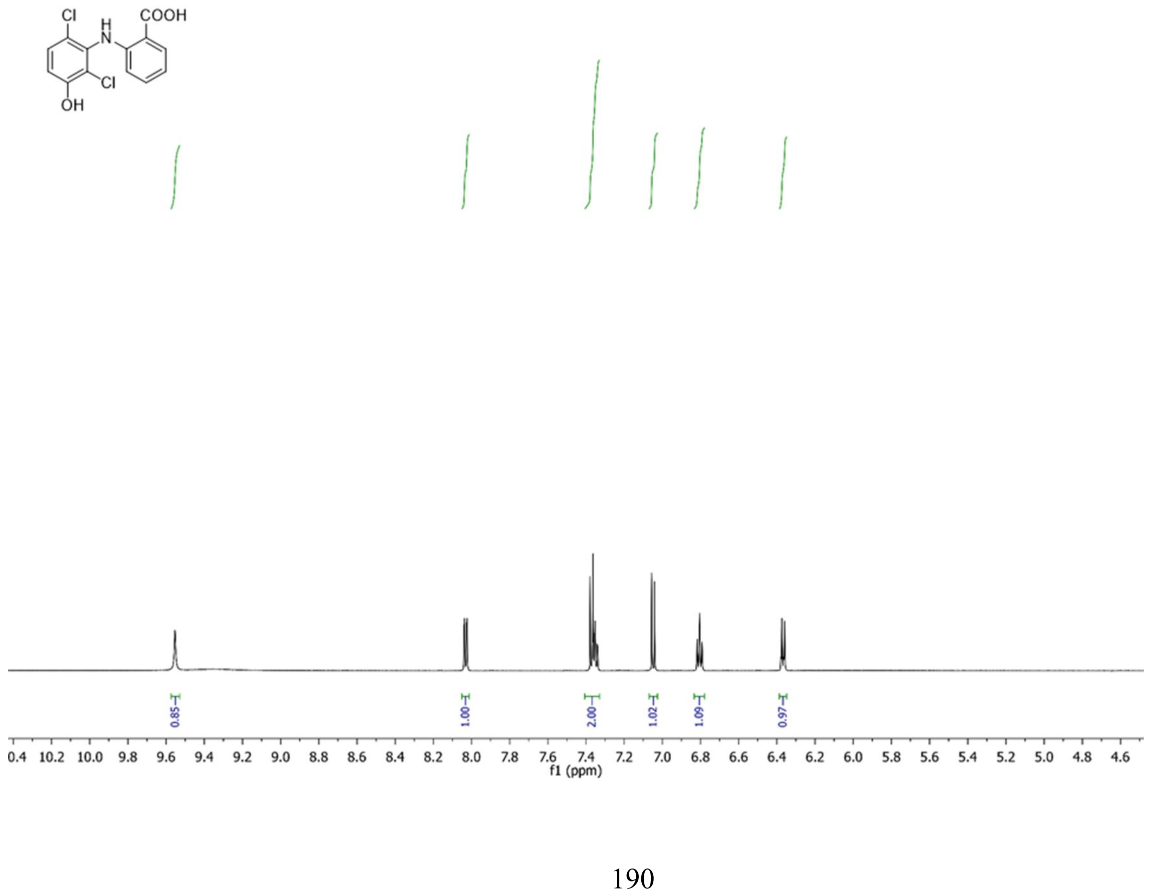

Cl
H
N
COOH
Cl
OH
0.85
1.00
2.00
1.02
1.09
0.97
0.4 10.2 10.0 9.8 9.6 9.4 9.2 9.0 8.8 8.6 8.4 8.2 8.0 7.8 7.6 7.4 7.2 7.0 6.8 6.6 6.4 6.2 6.0 5.8 5.6 5.4 5.2 5.0 4.8 4.6
f1 (ppm)

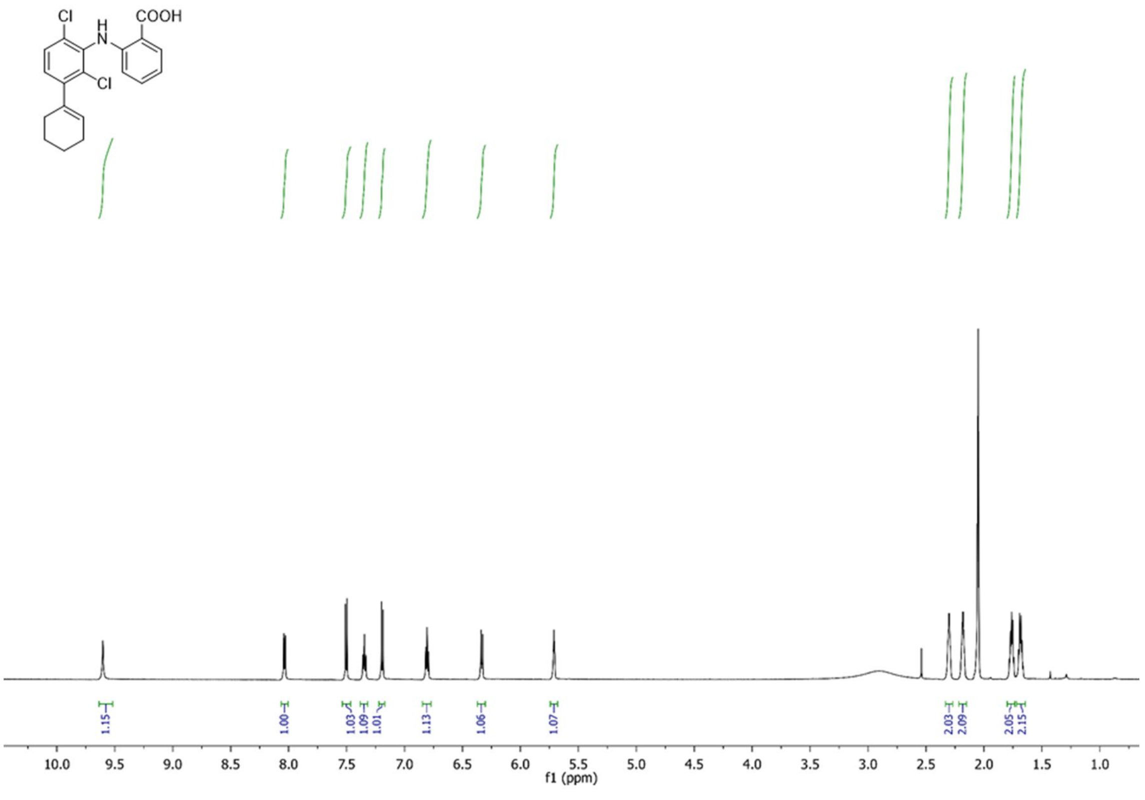
Cl
H
N
COOH
Cl
1.15
1.00
1.03
1.09
1.01
1.13
1.06
1.07
2.03
2.09
2.05
2.15
10.0 9.5 9.0 8.5 8.0 7.5 7.0 6.5 6.0 5.5 5.0 4.5 4.0 3.5 3.0 2.5 2.0 1.5 1.0
f1 (ppm)

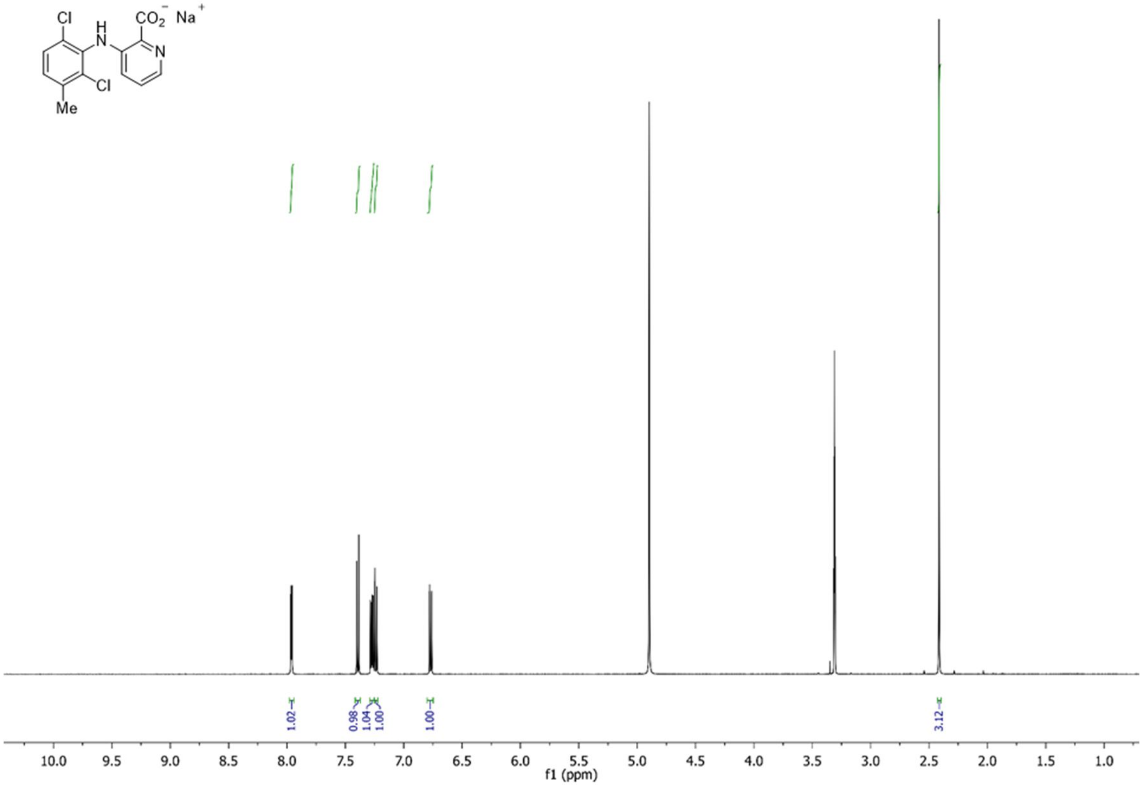
Cl
H
N
$CO_2^-$ $Na^+$
N
Cl
Me
1.02
0.98
1.04
1.00
1.00
3.12
10.0 9.5 9.0 8.5 8.0 7.5 7.0 6.5 6.0 5.5 5.0 4.5 4.0 3.5 3.0 2.5 2.0 1.5 1.0
f1 (ppm)

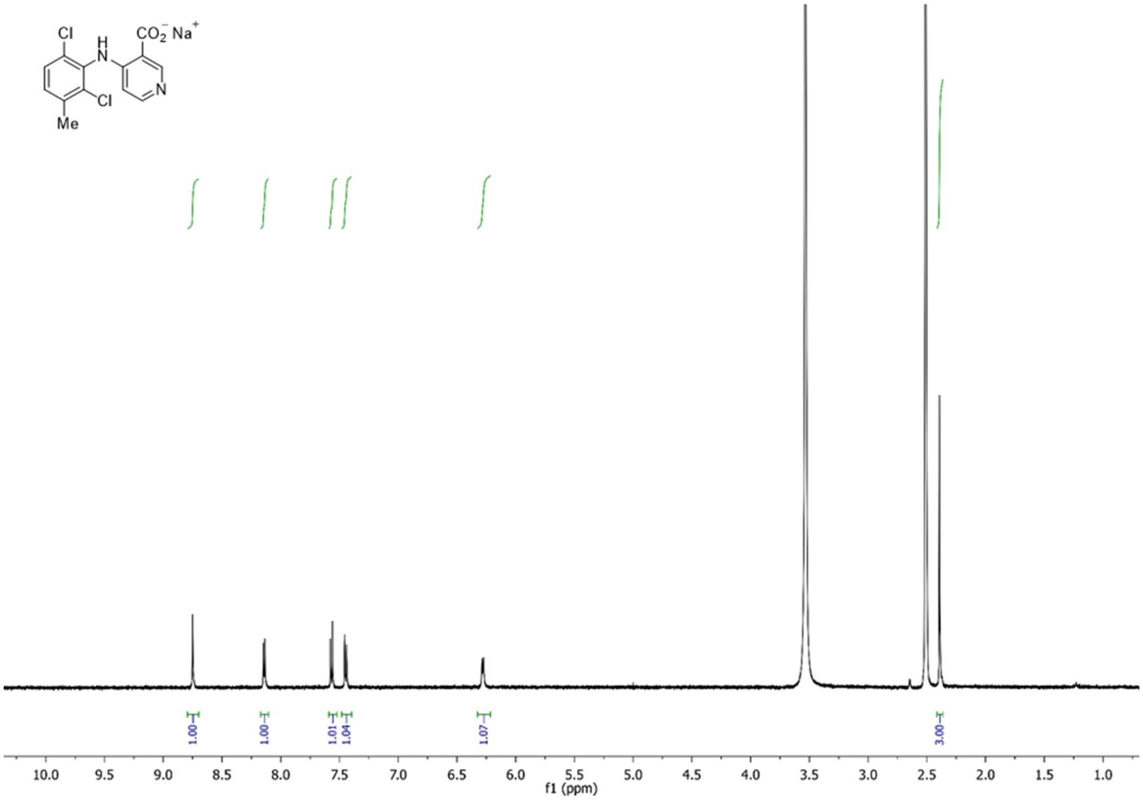

Cl
H
N
CO2⁻ Na⁺
N
Cl
Me
1.00
1.00
1.01
1.04
1.07
3.00
10.0 9.5 9.0 8.5 8.0 7.5 7.0 6.5 6.0 5.5 5.0 4.5 4.0 3.5 3.0 2.5 2.0 1.5 1.0
f1 (ppm)

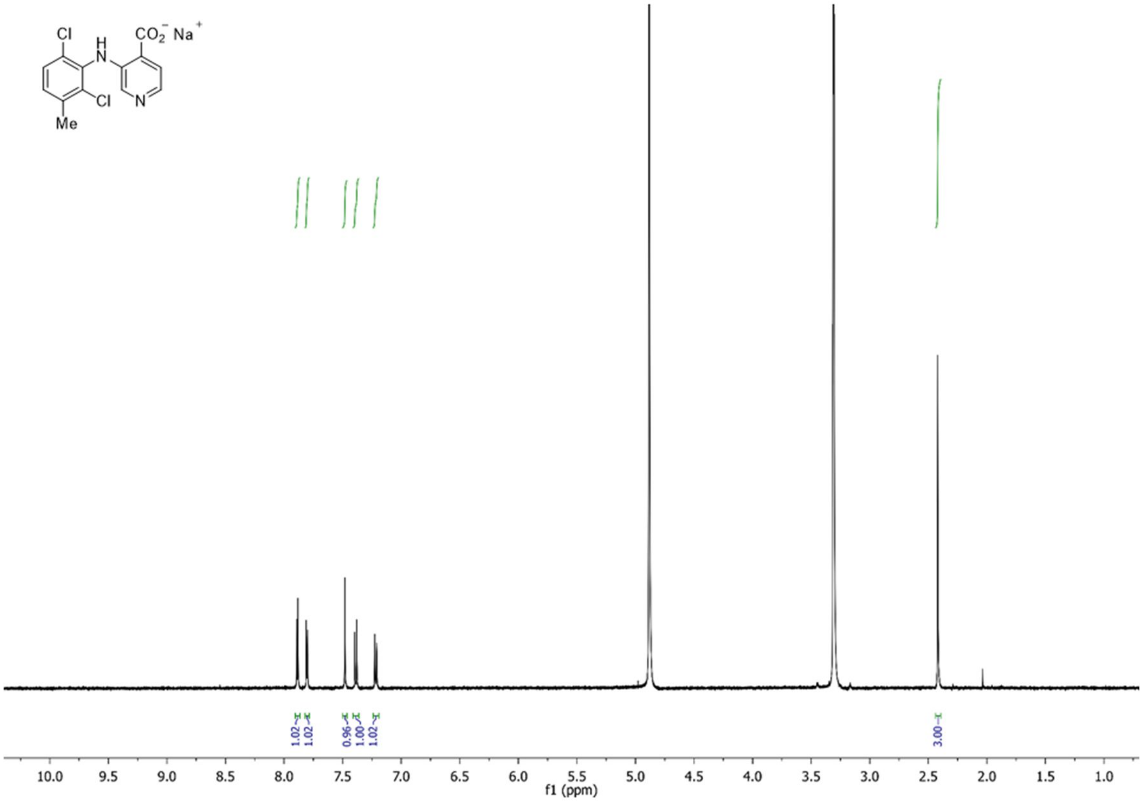

Cl
H
N
CO2⁻ Na⁺
N
Cl
Me
1.02
1.02
0.96
1.00
1.02
3.00
10.0 9.5 9.0 8.5 8.0 7.5 7.0 6.5 6.0 5.5 5.0 4.5 4.0 3.5 3.0 2.5 2.0 1.5 1.0
f1 (ppm)

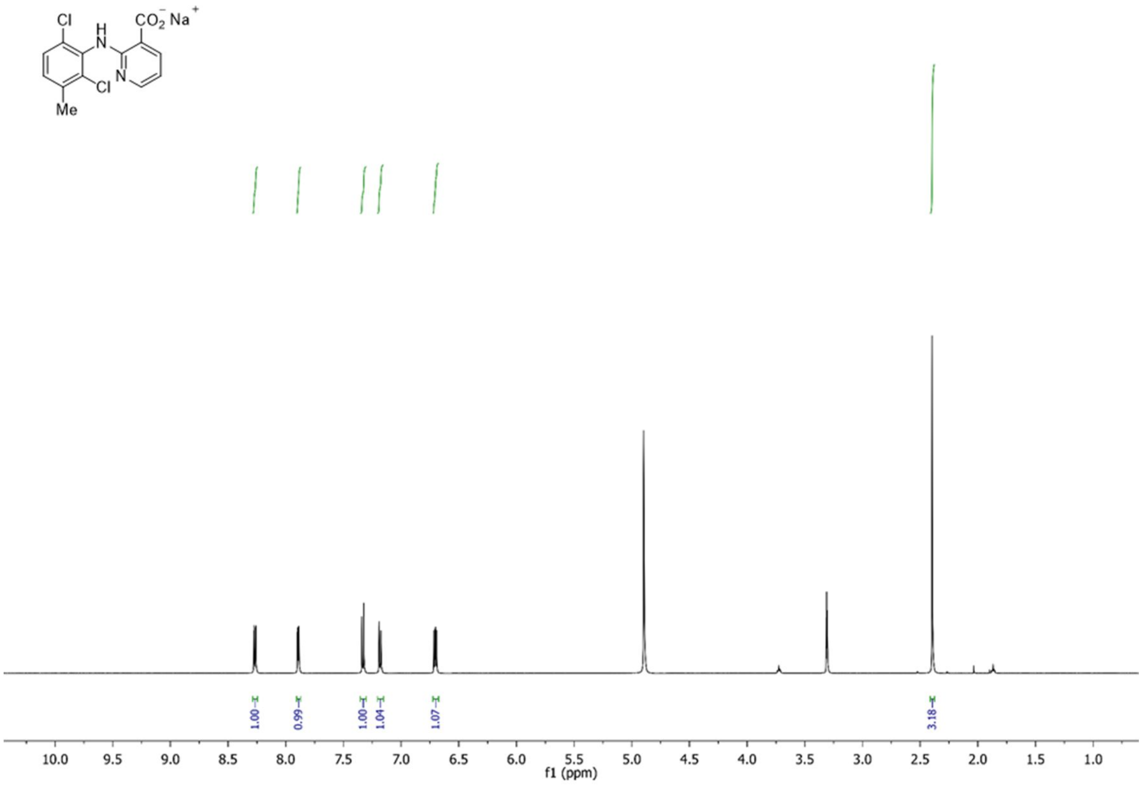

Cl
H
N
CO2- Na+
N
Cl
Me
1.00
0.99
1.00
1.04
1.07
3.18
10.0 9.5 9.0 8.5 8.0 7.5 7.0 6.5 6.0 5.5 5.0 4.5 4.0 3.5 3.0 2.5 2.0 1.5 1.0
f1 (ppm)

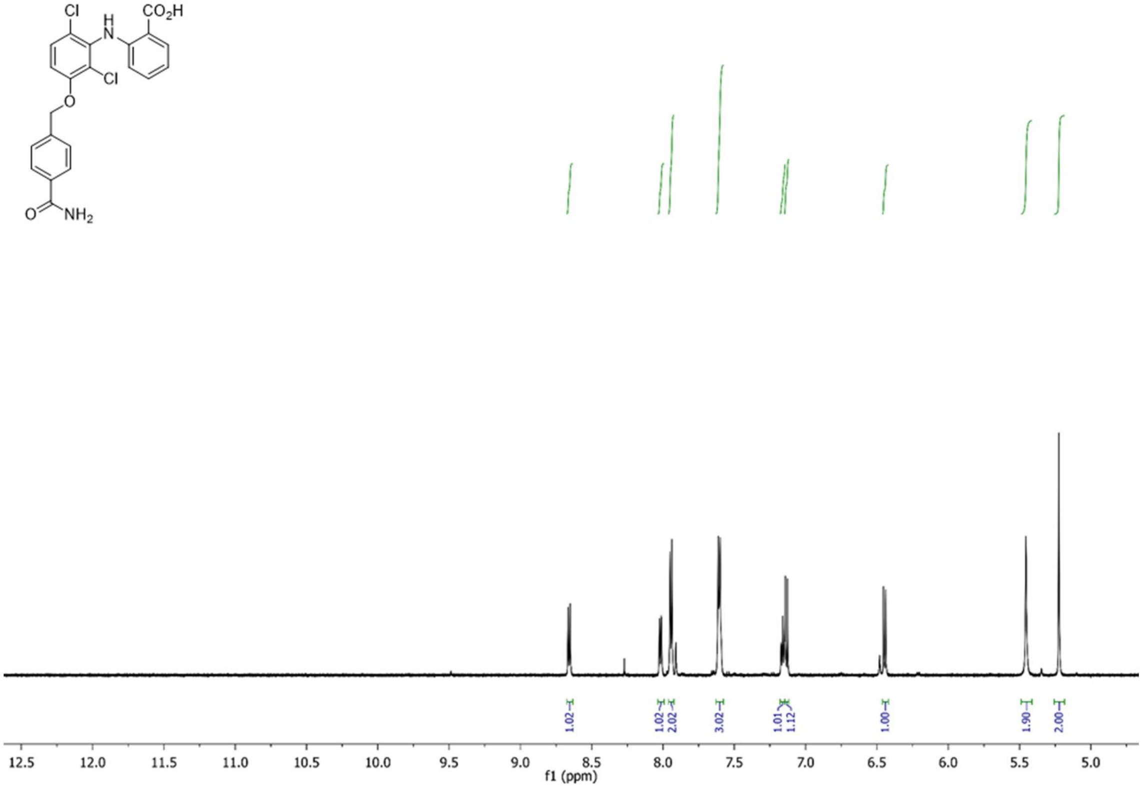

Cl
H
N
CO2H
Cl
O
O
NH2
1.02
1.02
2.02
3.02
1.01
1.12
1.00
1.90
2.00
12.5 12.0 11.5 11.0 10.5 10.0 9.5 9.0 8.5 8.0 7.5 7.0 6.5 6.0 5.5 5.0
f1 (ppm)

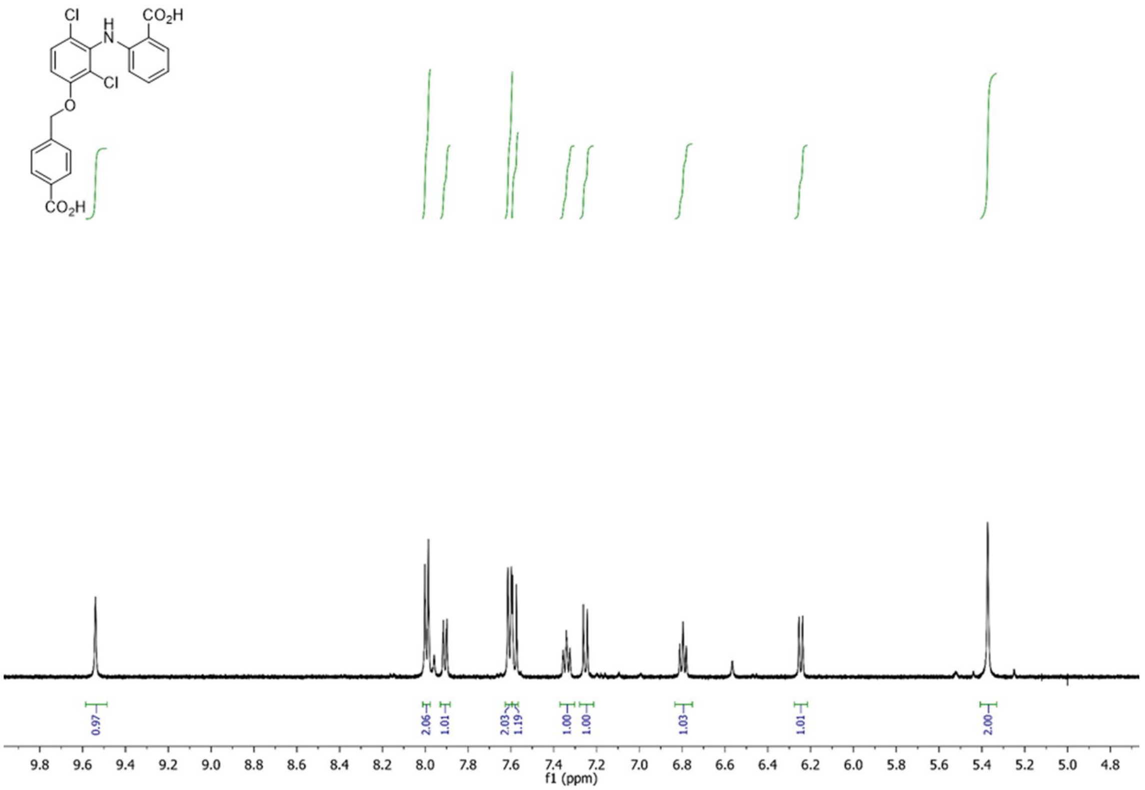
Cl
H
N
CO2H
Cl
O
CO2H
0.97
2.06
1.01
2.03
1.19
1.00
1.00
1.03
1.01
2.00
9.8 9.6 9.4 9.2 9.0 8.8 8.6 8.4 8.2 8.0 7.8 7.6 7.4 7.2 7.0 6.8 6.6 6.4 6.2 6.0 5.8 5.6 5.4 5.2 5.0 4.8
f1 (ppm)

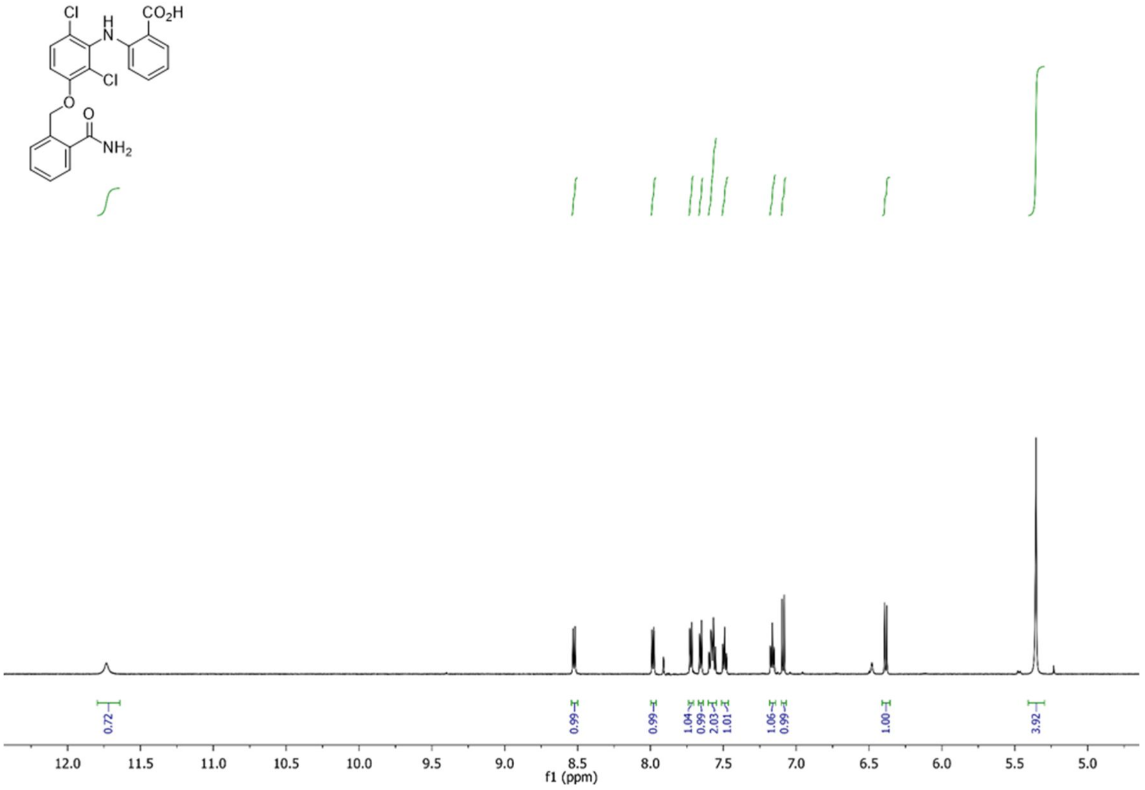
Cl
H
N
CO2H
Cl
O
O
NH2
0.72
0.99
0.99
1.04
0.99
2.03
1.01
1.06
0.99
1.00
3.92
12.0 11.5 11.0 10.5 10.0 9.5 9.0 8.5 8.0 7.5 7.0 6.5 6.0 5.5 5.0
f1 (ppm)

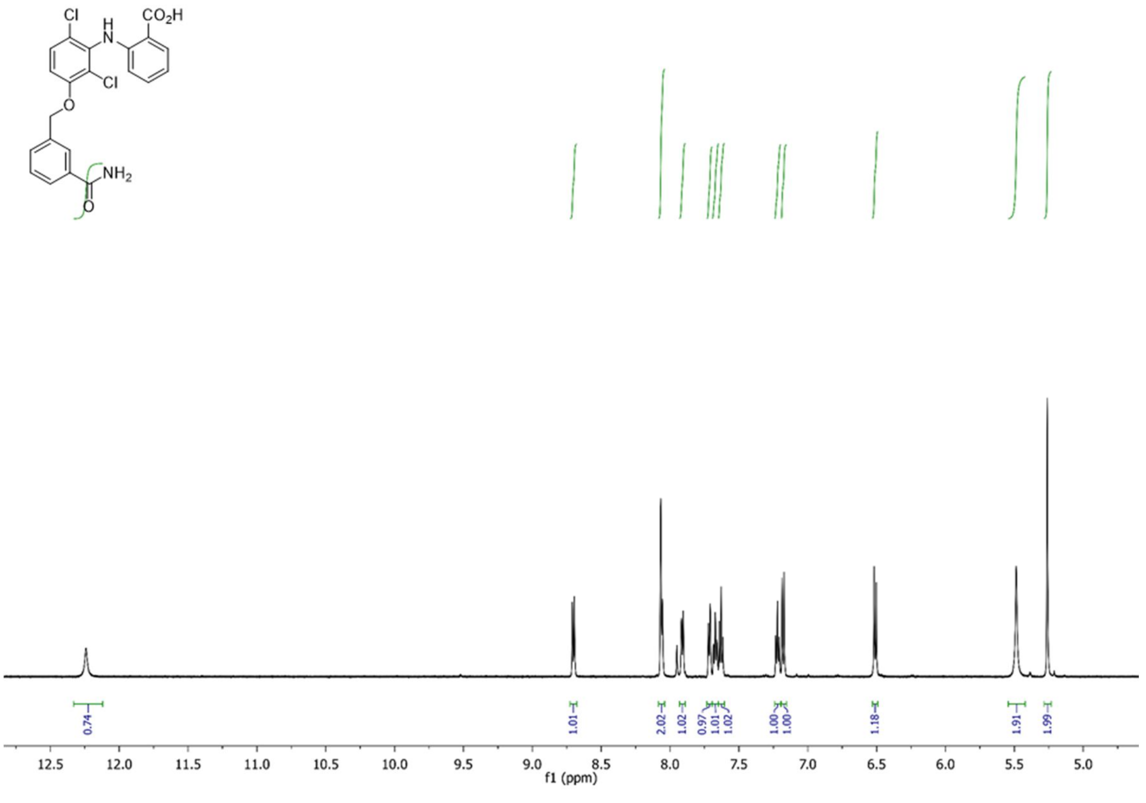
Cl
H
N
CO2H
Cl
O
NH2
O
0.74
1.01
2.02
1.02
0.97
1.01
1.02
1.00
1.00
1.18
1.91
1.99
12.5
12.0
11.5
11.0
10.5
10.0
9.5
9.0
8.5
8.0
7.5
7.0
6.5
6.0
5.5
5.0
f1 (ppm)

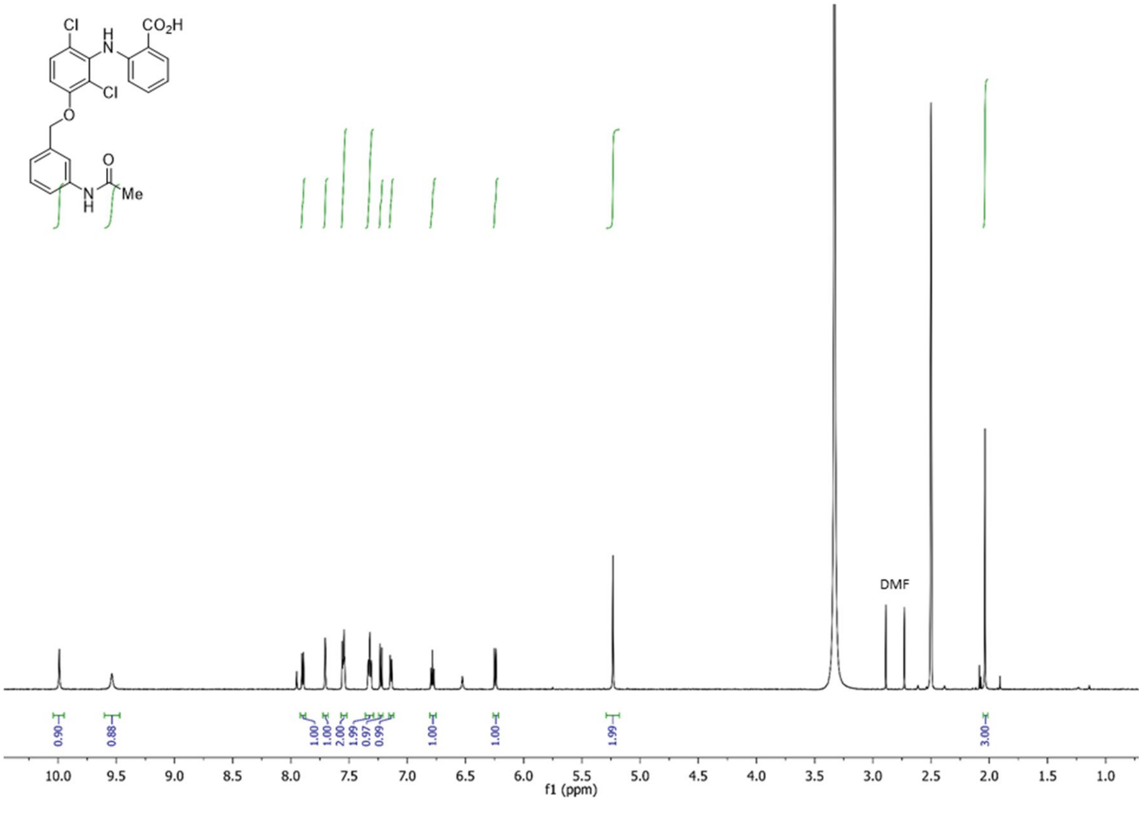
Cl
H
N
CO2H
Cl
O
N
H
O
Me
DMF
0.90
0.88
1.00
1.00
2.00
1.99
0.97
0.99
1.00
1.00
1.99
3.00
10.0
9.5
9.0
8.5
8.0
7.5
7.0
6.5
6.0
5.5
5.0
4.5
4.0
3.5
3.0
2.5
2.0
1.5
1.0
f1 (ppm)

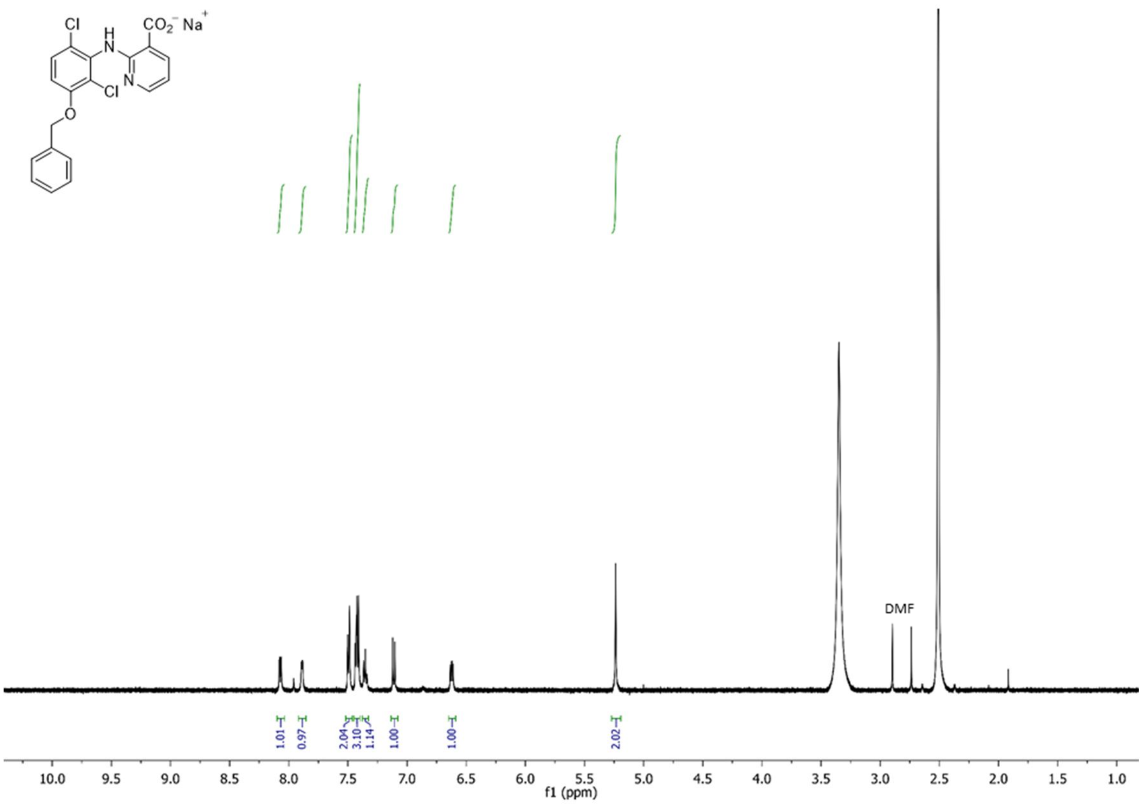

DMF
1.01
0.97
2.04
3.10
1.14
1.00
1.00
2.02
10.0 9.5 9.0 8.5 8.0 7.5 7.0 6.5 6.0 5.5 5.0 4.5 4.0 3.5 3.0 2.5 2.0 1.5 1.0
f1 (ppm)

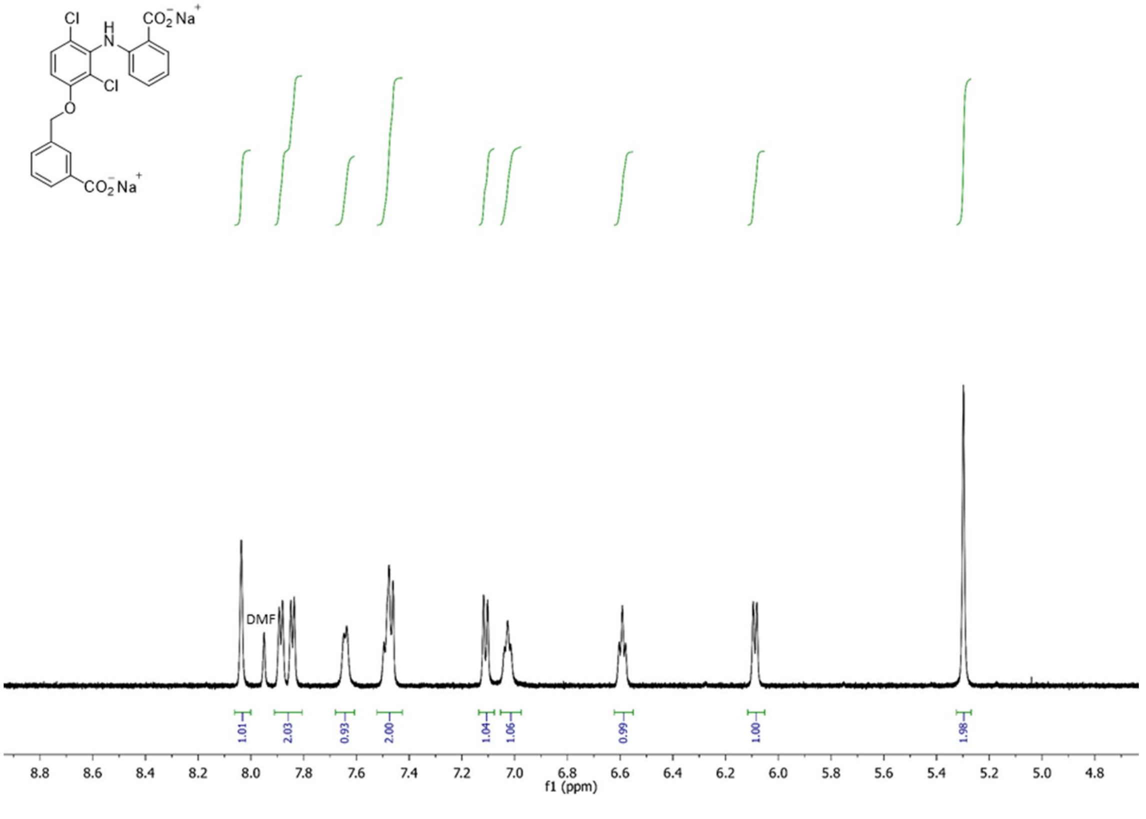

DMF
1.01
2.03
0.93
2.00
1.04
1.06
0.99
1.00
1.98
8.8 8.6 8.4 8.2 8.0 7.8 7.6 7.4 7.2 7.0 6.8 6.6 6.4 6.2 6.0 5.8 5.6 5.4 5.2 5.0 4.8
f1 (ppm)

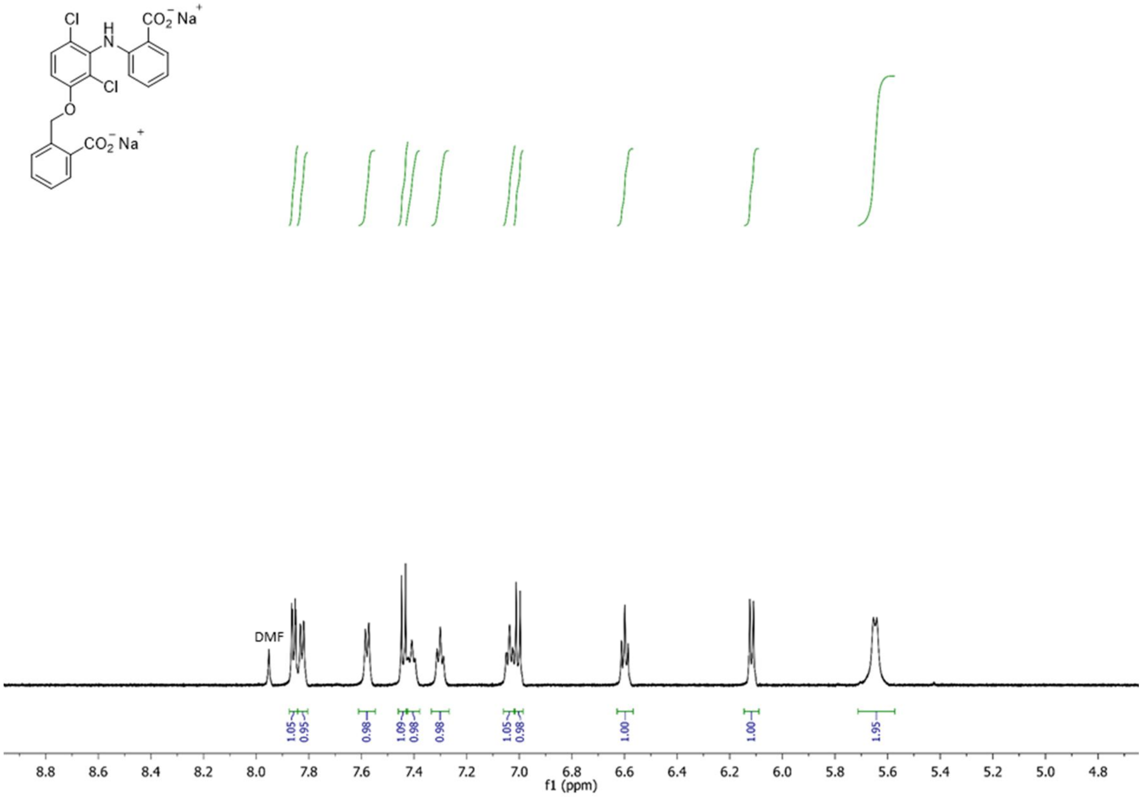
Cl
H
N
CO2− Na+
Cl
O
CO2− Na+
DMF
1.05
0.95
0.98
1.09
0.98
0.98
1.05
0.98
1.00
1.00
1.95
8.8 8.6 8.4 8.2 8.0 7.8 7.6 7.4 7.2 7.0 6.8 6.6 6.4 6.2 6.0 5.8 5.6 5.4 5.2 5.0 4.8
f1 (ppm)

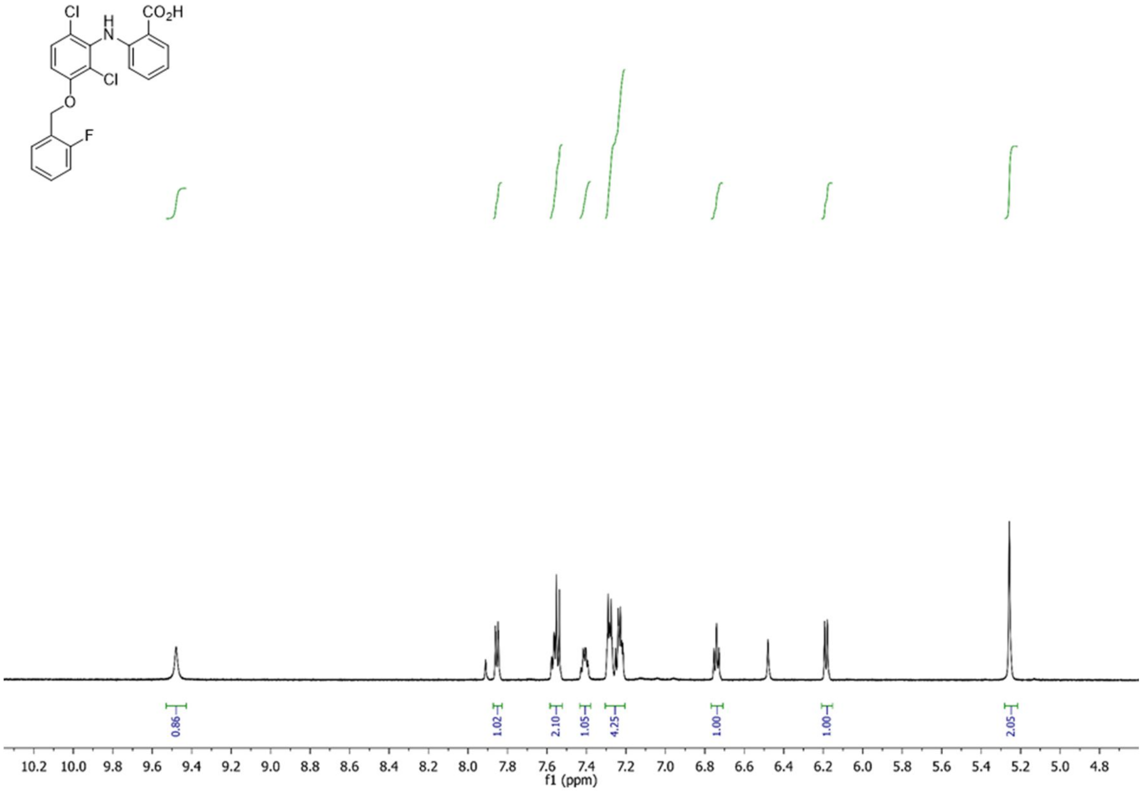
Cl
H
N
CO2H
Cl
O
F
0.86
1.02
2.10
1.05
4.25
1.00
1.00
2.05
10.2 10.0 9.8 9.6 9.4 9.2 9.0 8.8 8.6 8.4 8.2 8.0 7.8 7.6 7.4 7.2 7.0 6.8 6.6 6.4 6.2 6.0 5.8 5.6 5.4 5.2 5.0 4.8
f1 (ppm)

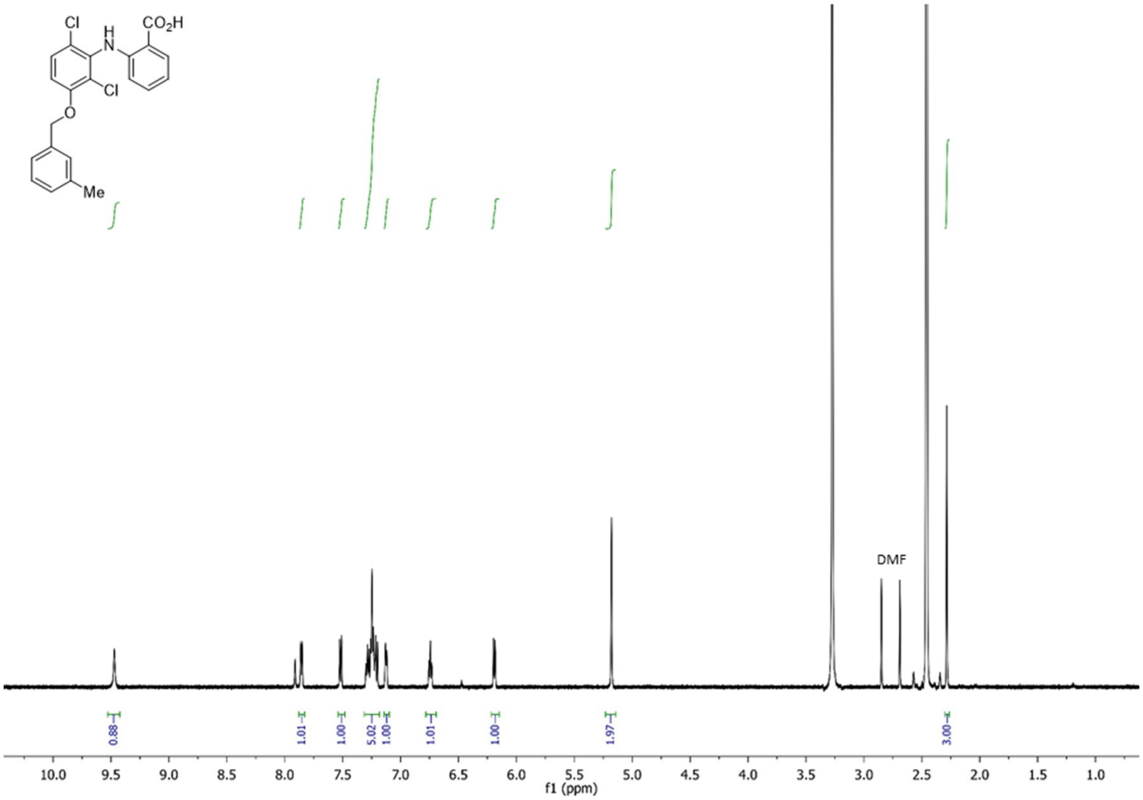
Cl
H
N
$CO_2H$
Cl
O
Me
DMF
0.88
1.01
1.00
5.02
1.00
1.01
1.00
1.97
3.00
f1 (ppm)

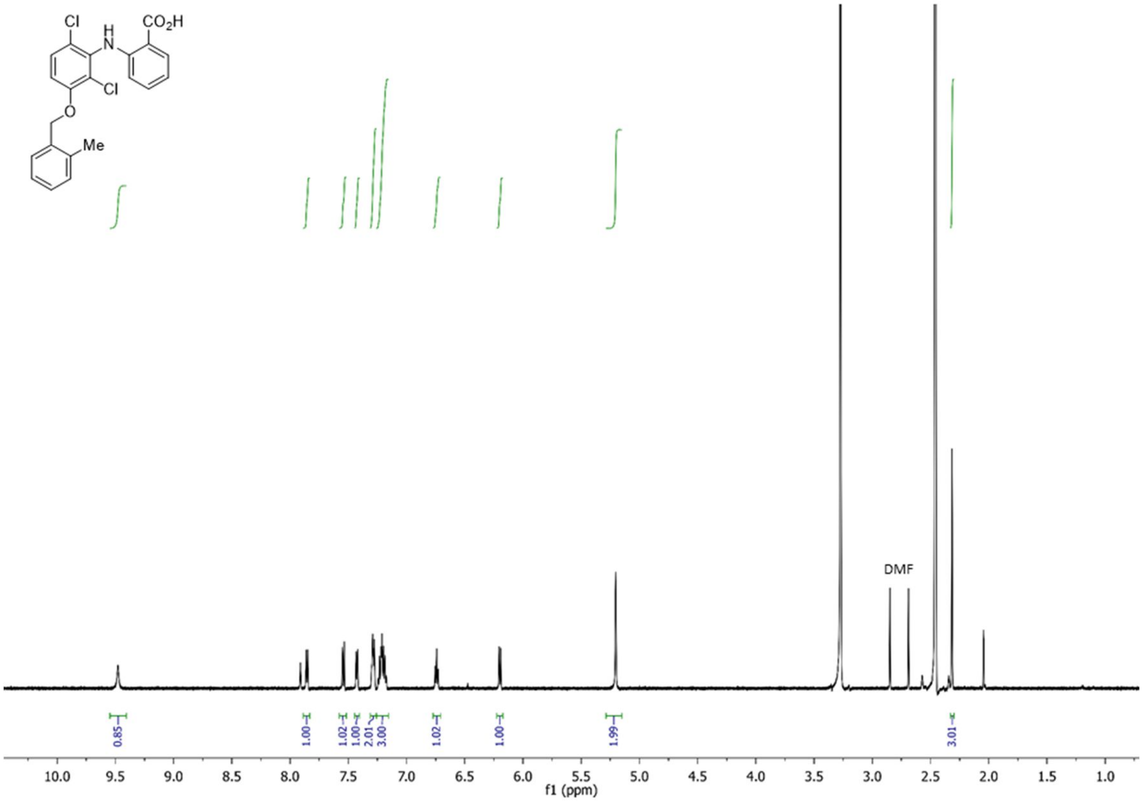
Cl
H
N
$CO_2H$
Cl
O
Me
DMF
0.85
1.00
1.02
1.00
2.01
3.00
1.02
1.00
1.99
3.01
f1 (ppm)

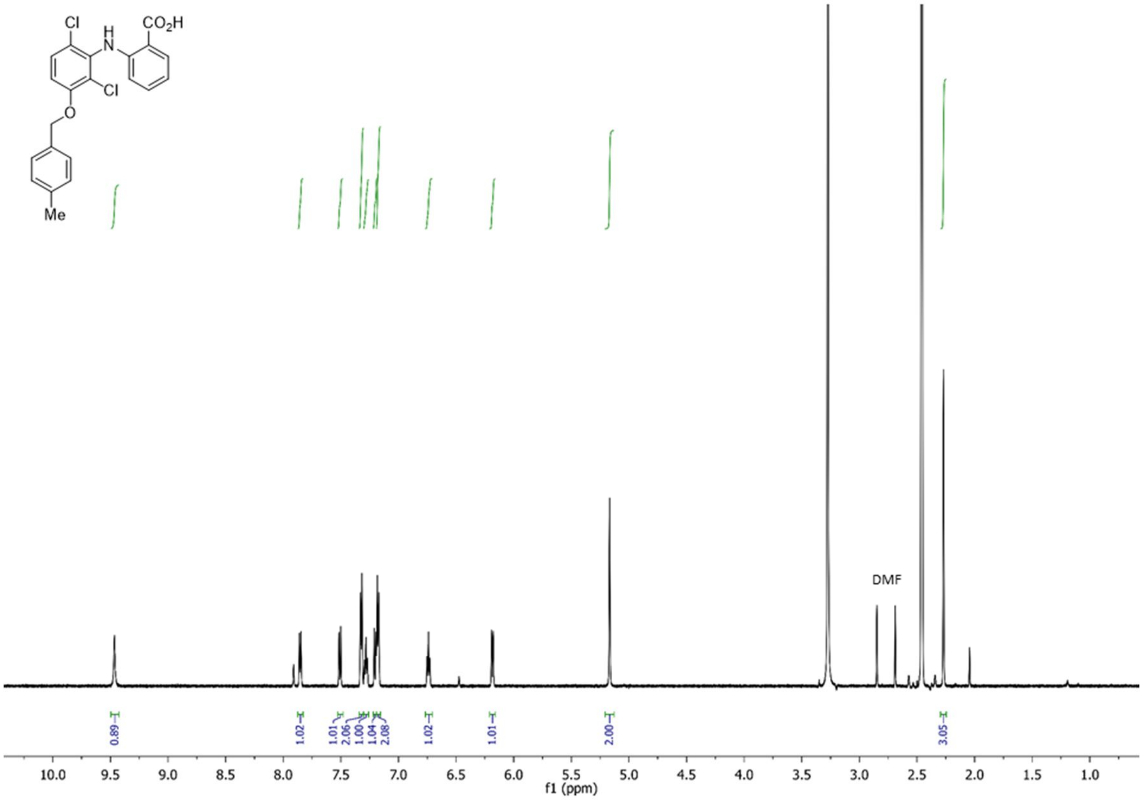
Cl
H
N
CO2H
Cl
O
Me
DMF
0.89
1.02
1.01
2.06
1.00
1.04
2.08
1.02
1.01
2.00
3.05
10.0 9.5 9.0 8.5 8.0 7.5 7.0 6.5 6.0 5.5 5.0 4.5 4.0 3.5 3.0 2.5 2.0 1.5 1.0
f1 (ppm)

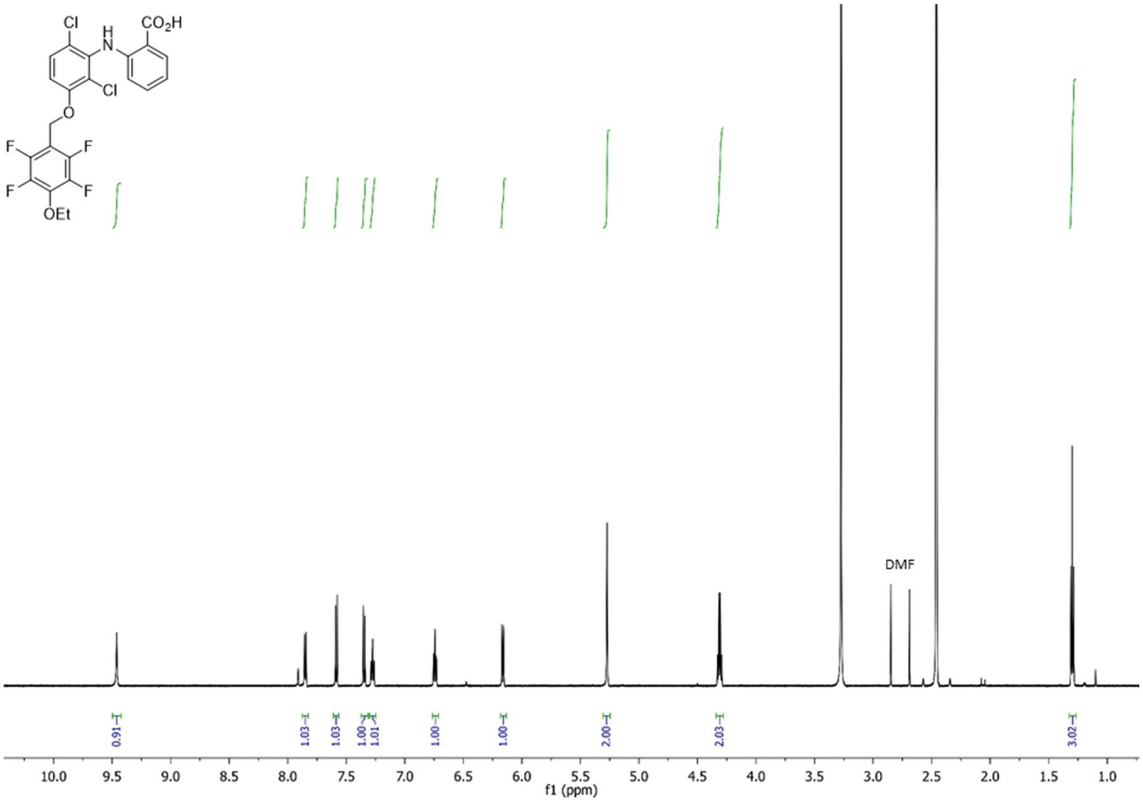
Cl
H
N
CO2H
Cl
O
F
F
F
F
OEt
DMF
0.91
1.03
1.03
1.00
1.01
1.00
1.00
2.00
2.03
3.02
10.0 9.5 9.0 8.5 8.0 7.5 7.0 6.5 6.0 5.5 5.0 4.5 4.0 3.5 3.0 2.5 2.0 1.5 1.0
f1 (ppm)

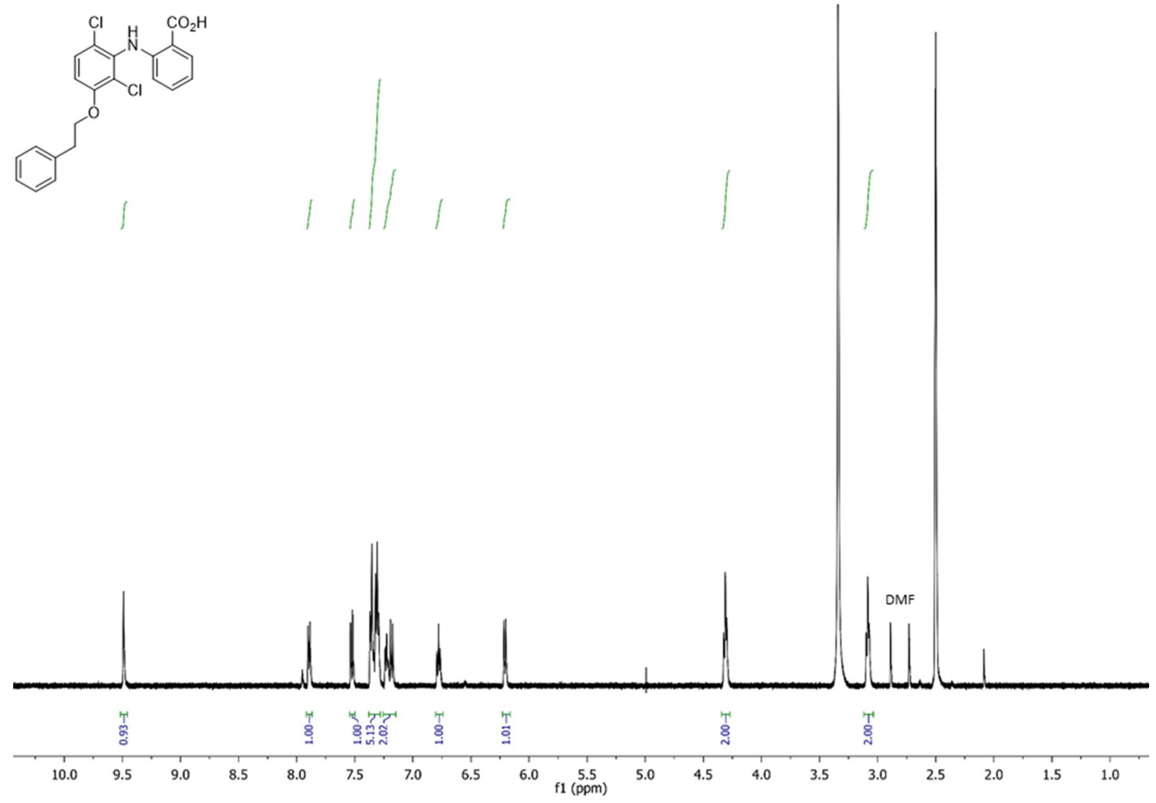
CO2H
Cl
H
N
Cl
O
DMF
0.93
1.00
1.00
5.13
2.02
1.00
1.01
2.00
2.00
f1 (ppm)

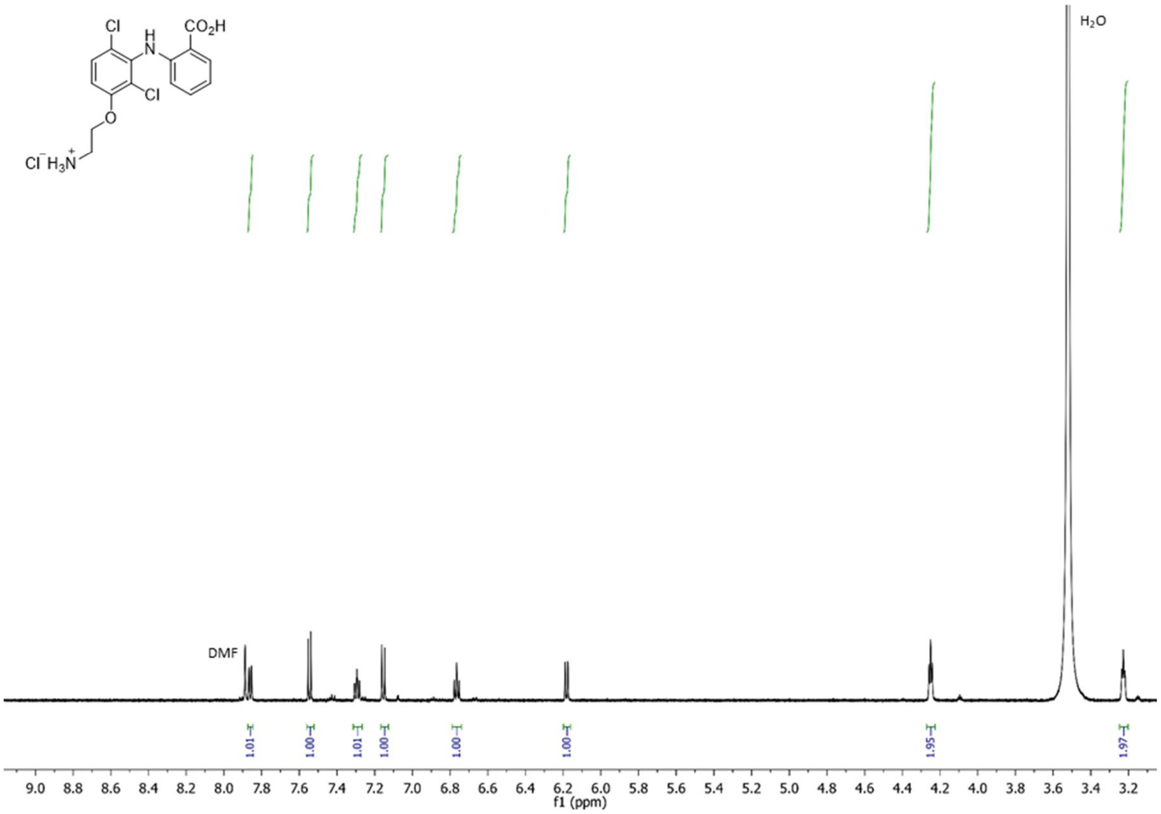
CO2H
Cl
H
N
Cl
O
Cl⁻ H3N⁺
H2O
DMF
1.01
1.00
1.01
1.00
1.00
1.00
1.95
1.97
f1 (ppm)

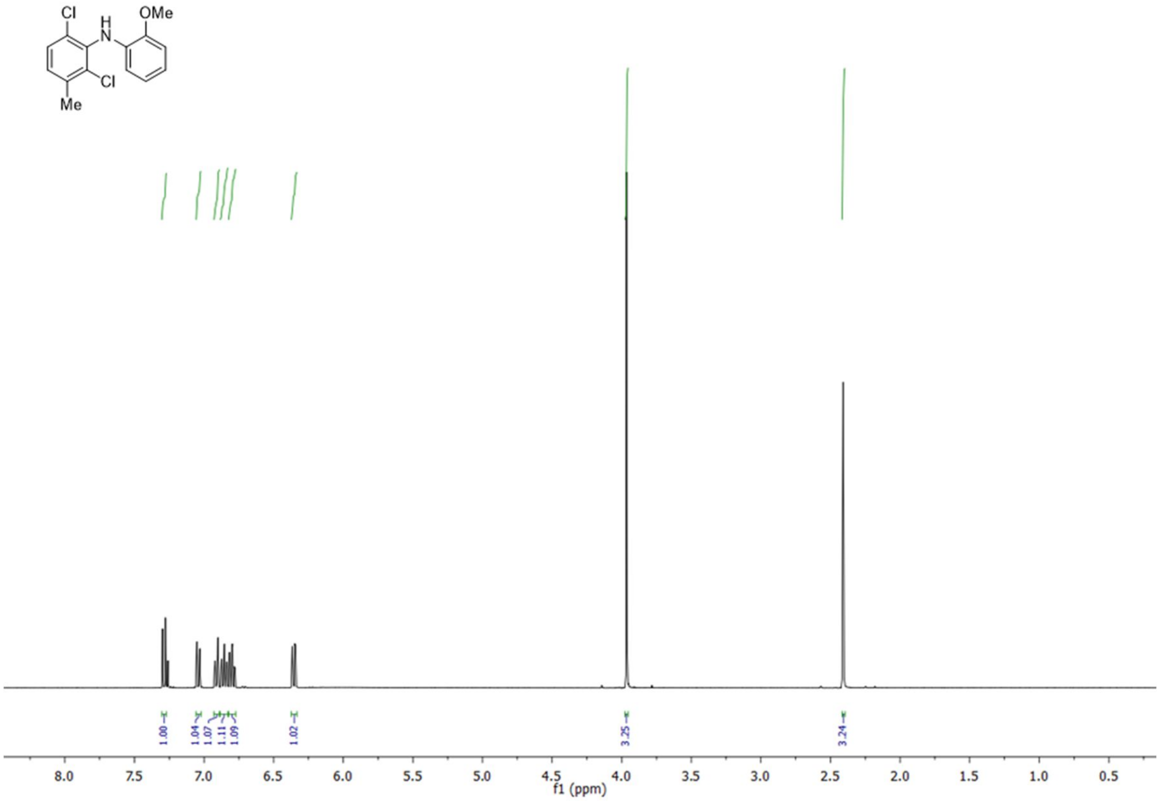
Cl
H
N
OMe
Cl
Me
1.00
1.04
1.07
1.11
1.09
1.02
3.25
3.24
8.0 7.5 7.0 6.5 6.0 5.5 5.0 4.5 4.0 3.5 3.0 2.5 2.0 1.5 1.0 0.5
f1 (ppm)

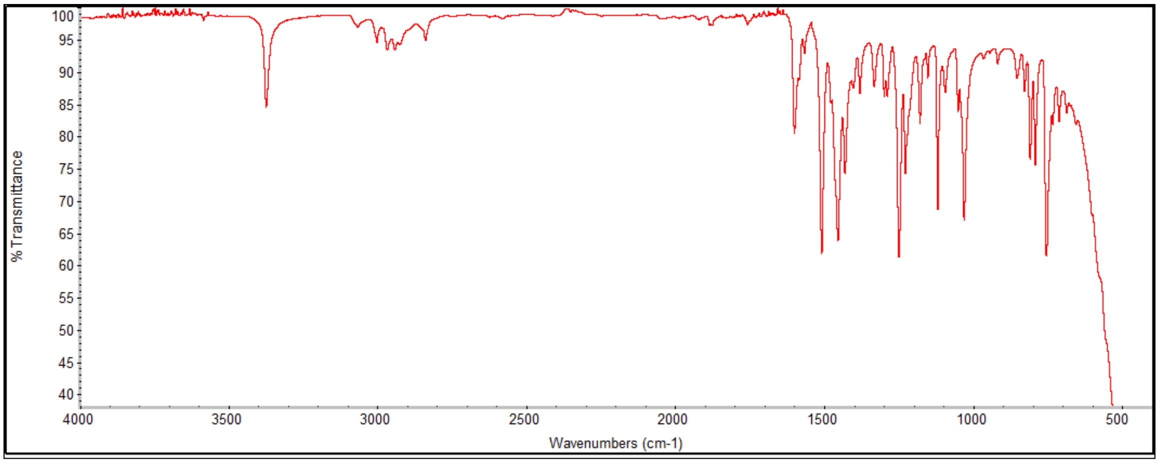
% Transmittance
100 95 90 85 80 75 70 65 60 55 50 45 40
4000 3500 3000 2500 2000 1500 1000 500
Wavenumbers (cm-1)

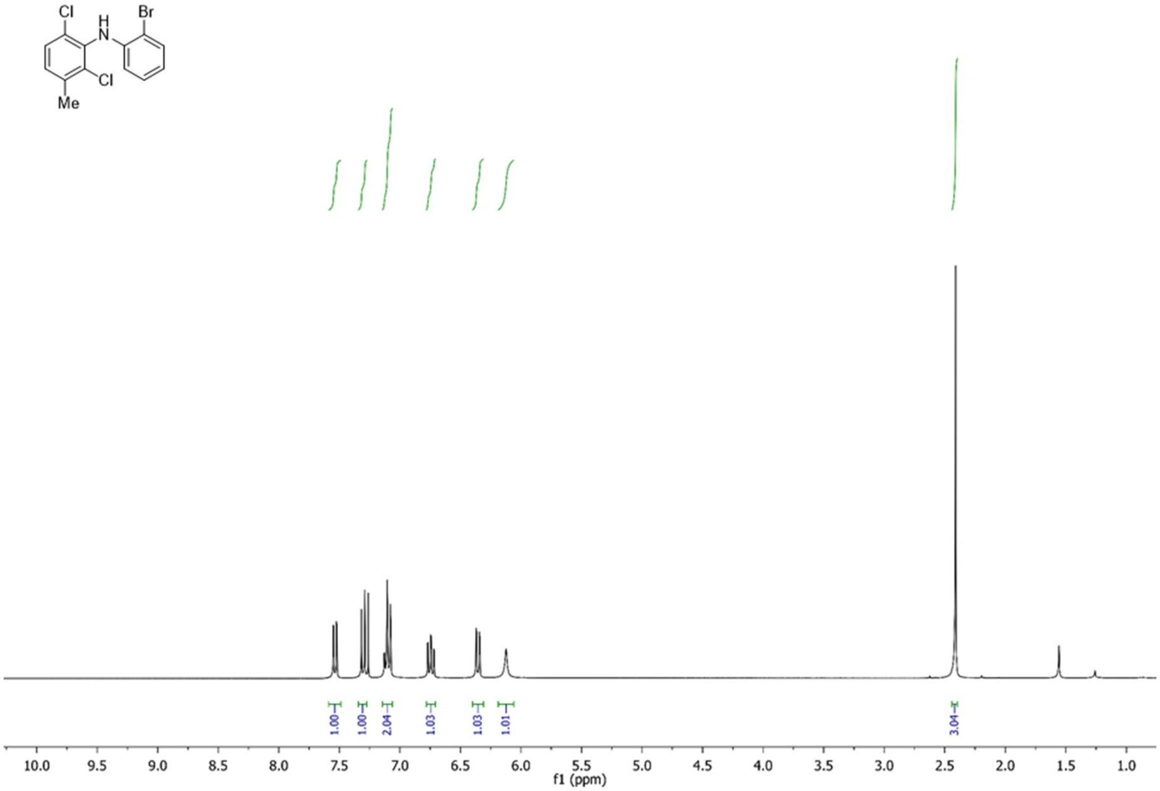
Cl
H
N
Br
Cl
Me
1.00
1.00
2.04
1.03
1.03
1.01
3.04
10.0 9.5 9.0 8.5 8.0 7.5 7.0 6.5 6.0 5.5 5.0 4.5 4.0 3.5 3.0 2.5 2.0 1.5 1.0
f1 (ppm)

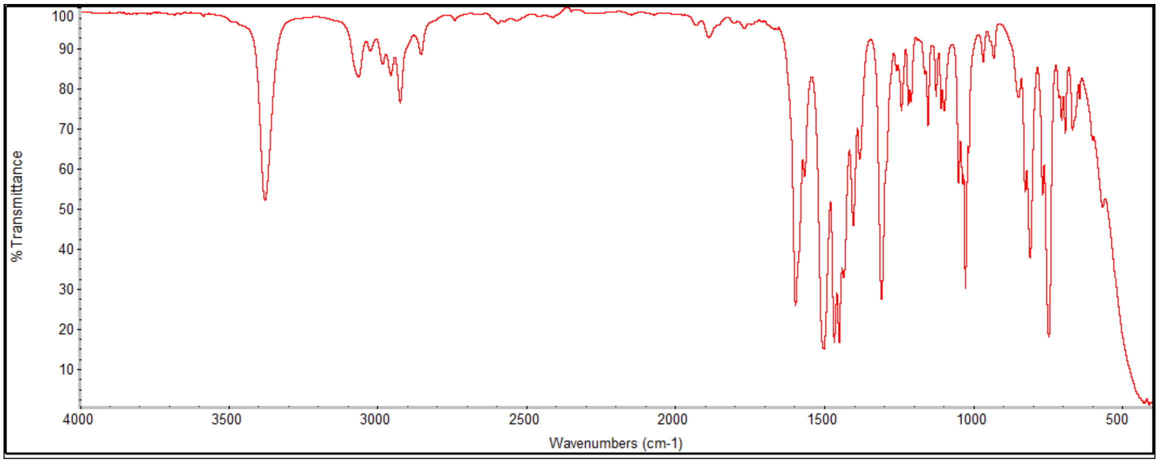
% Transmittance
100 90 80 70 60 50 40 30 20 10
4000 3500 3000 2500 2000 1500 1000 500
Wavenumbers (cm-1)

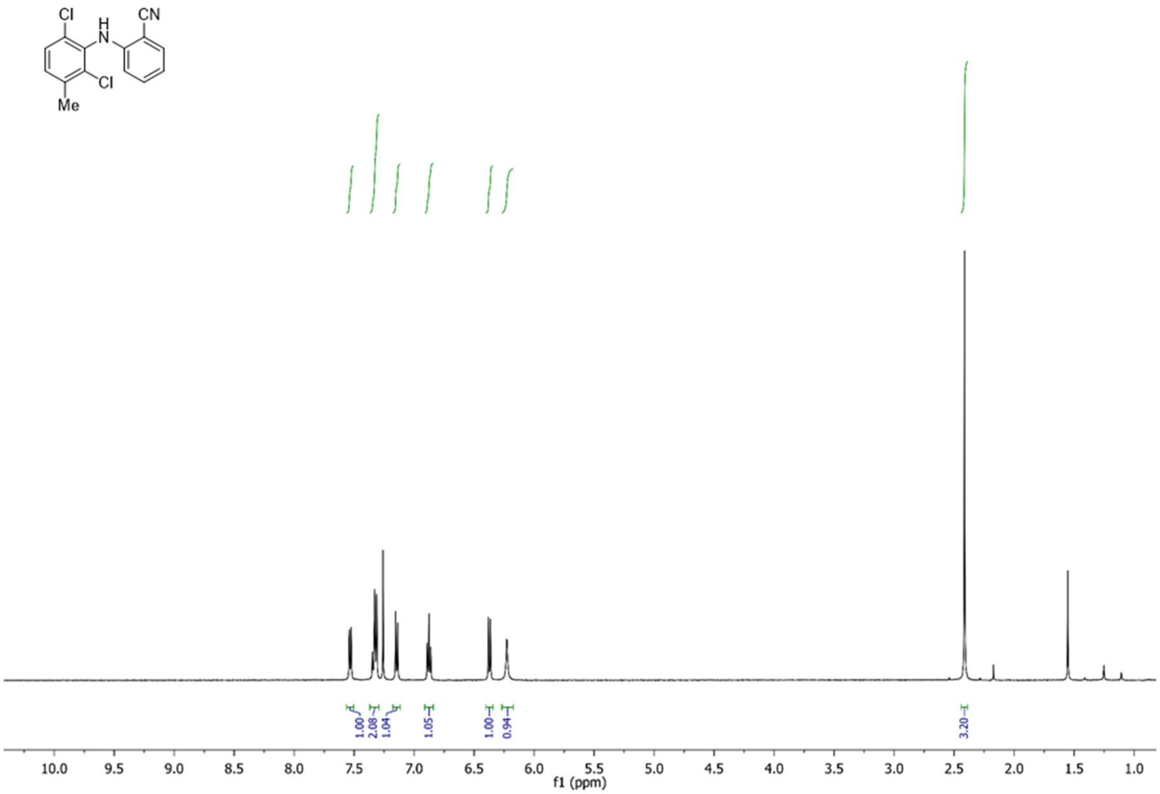
Cl
H
N
CN
Cl
Me
1.00
2.08
1.04
1.05
1.00
0.94
3.20
10.0 9.5 9.0 8.5 8.0 7.5 7.0 6.5 6.0 5.5 5.0 4.5 4.0 3.5 3.0 2.5 2.0 1.5 1.0
f1 (ppm)

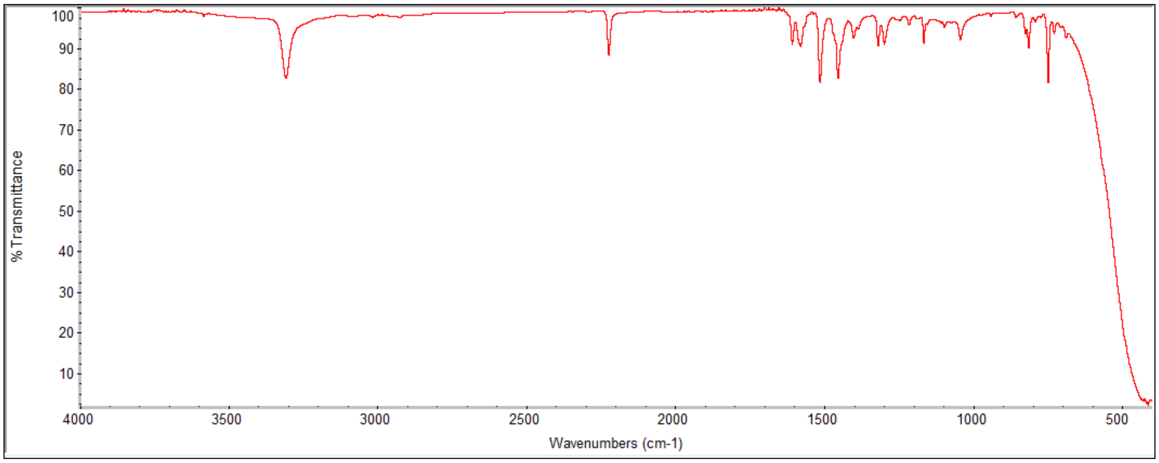
% Transmittance
100 90 80 70 60 50 40 30 20 10
4000 3500 3000 2500 2000 1500 1000 500
Wavenumbers (cm-1)

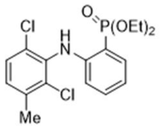
Cl
H
N
O
P(OEt)2
Cl
Me

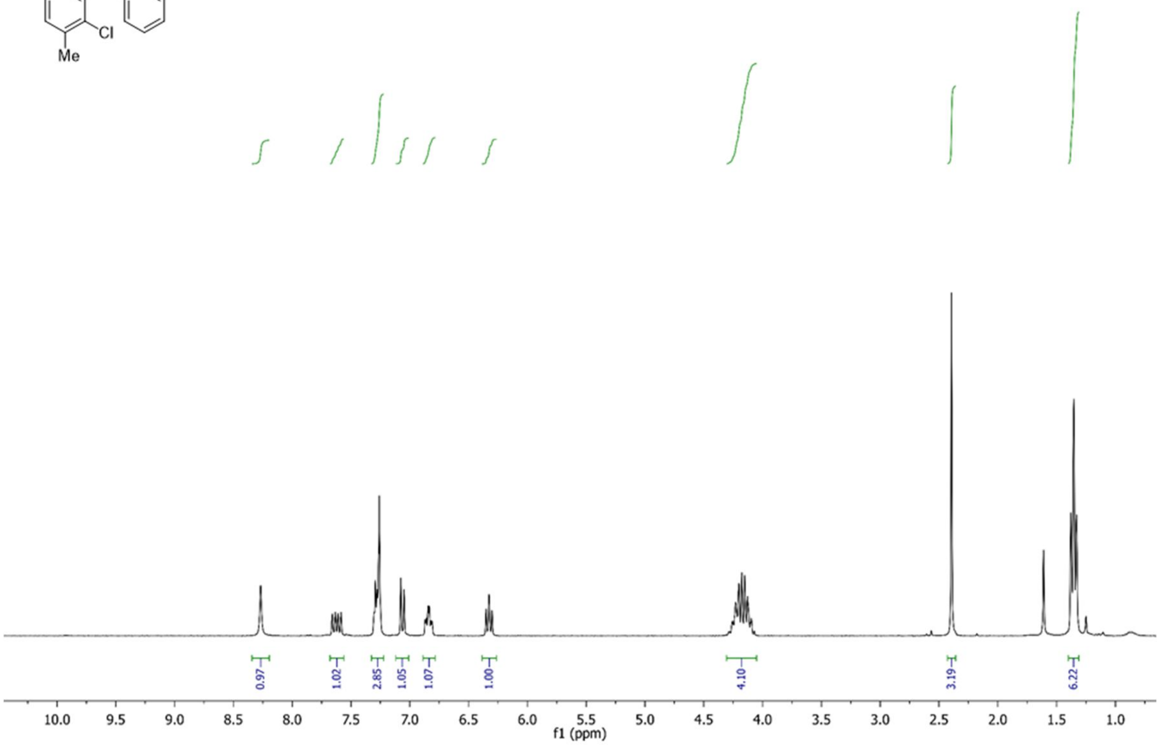
0.97
1.02
2.85
1.05
1.07
1.00
4.10
3.19
6.22
10.0 9.5 9.0 8.5 8.0 7.5 7.0 6.5 6.0 5.5 5.0 4.5 4.0 3.5 3.0 2.5 2.0 1.5 1.0
f1 (ppm)

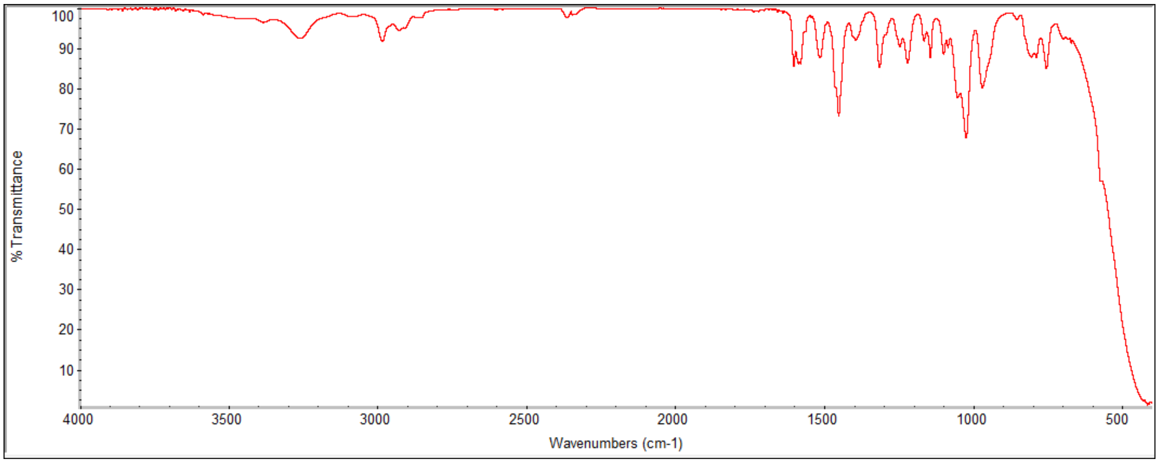
% Transmittance
100 90 80 70 60 50 40 30 20 10
4000 3500 3000 2500 2000 1500 1000 500
Wavenumbers (cm-1)

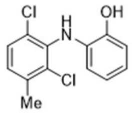

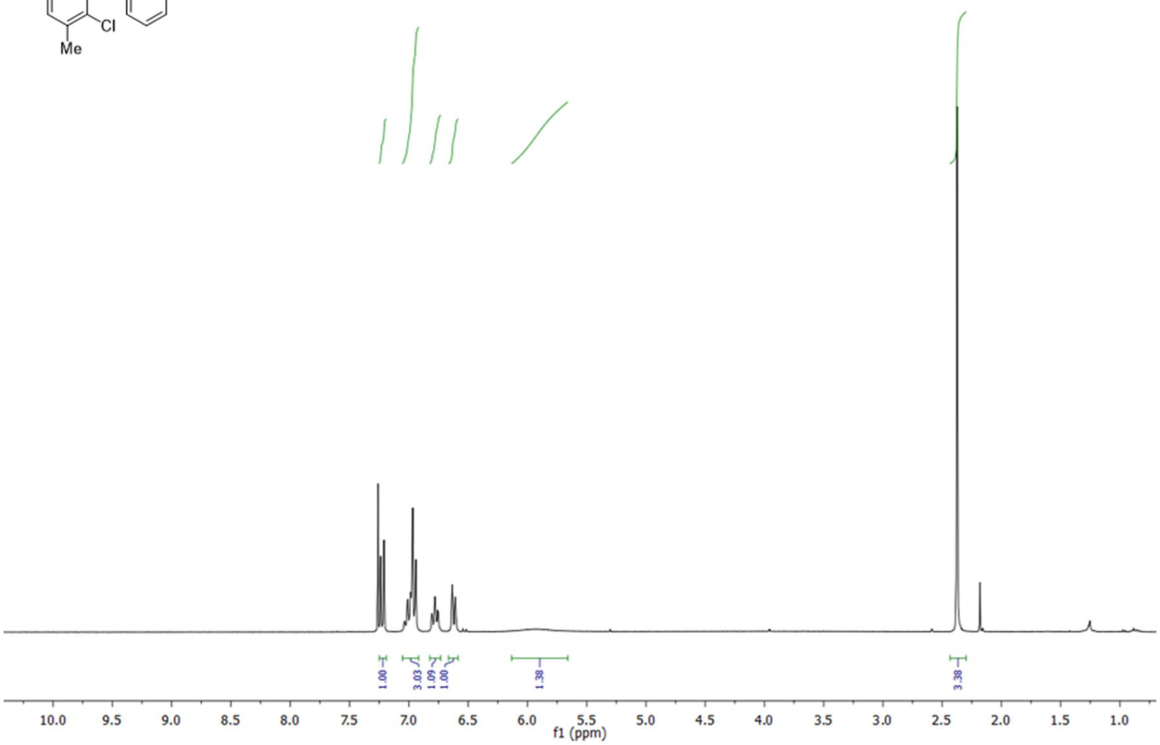
1.00
3.03
1.09
1.00
1.38
3.38
10.0
9.5
9.0
8.5
8.0
7.5
7.0
6.5
6.0
5.5
5.0
4.5
4.0
3.5
3.0
2.5
2.0
1.5
1.0
f1 (ppm)

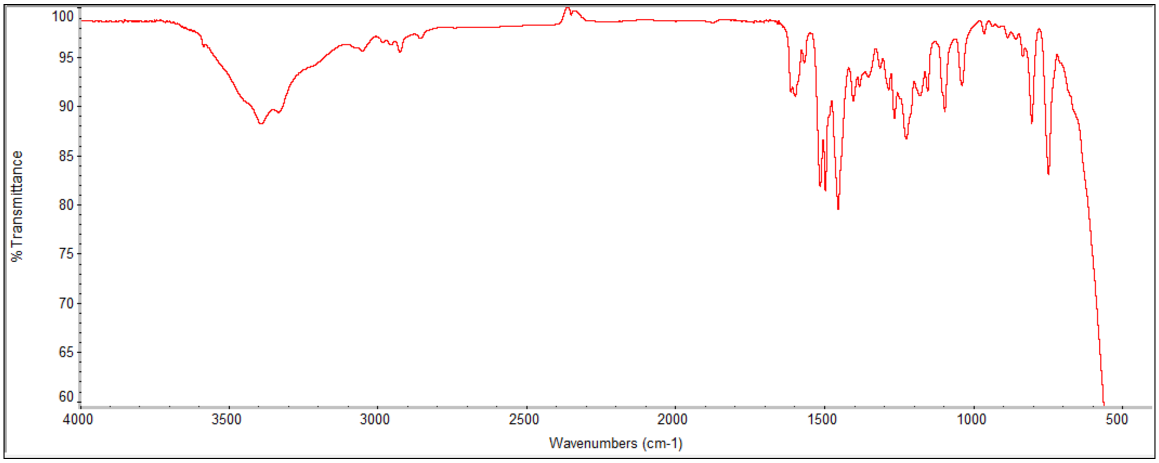
% Transmittance
Wavenumbers (cm-1)

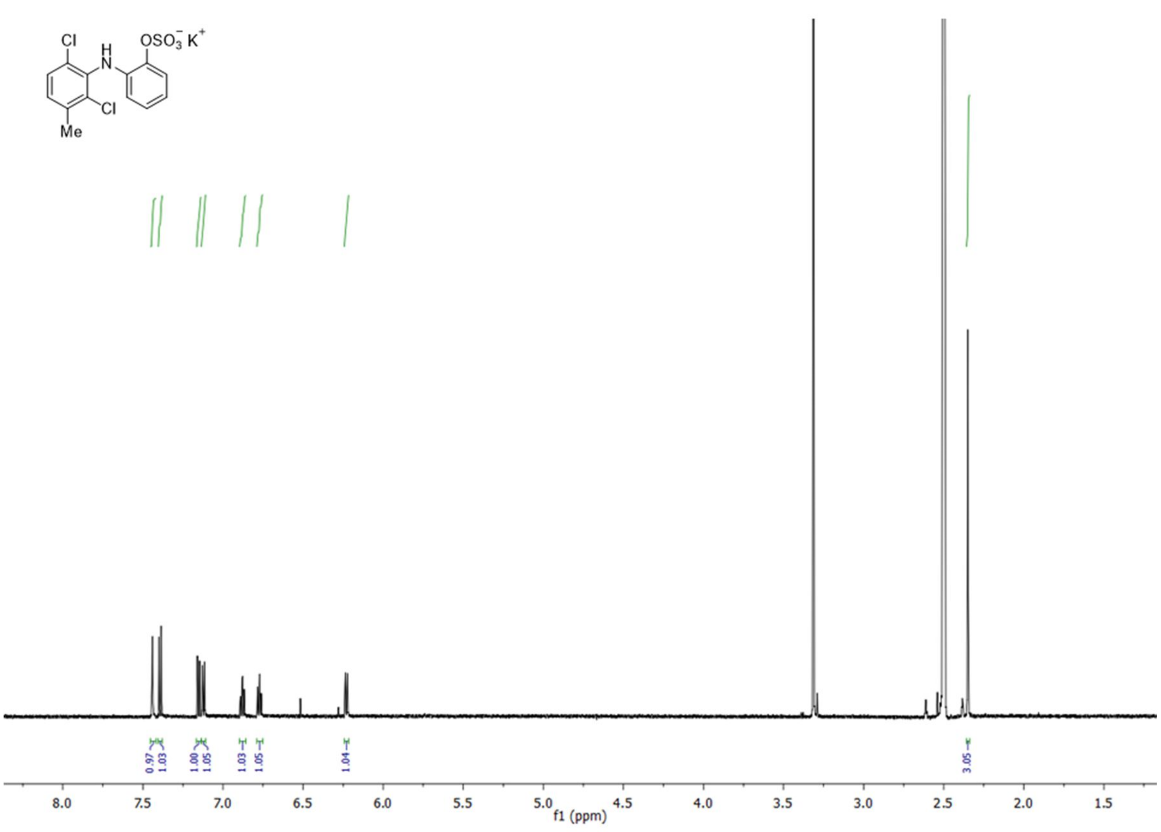

OSO3- K+
Cl
H
N
Cl
Me
0.97
1.03
1.00
1.05
1.03
1.05
1.04
3.05
8.0
7.5
7.0
6.5
6.0
5.5
5.0
4.5
4.0
3.5
3.0
2.5
2.0
1.5
f1 (ppm)

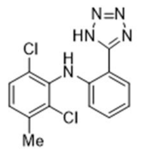
N=N
HN
N
Cl
H
N
Cl
Me

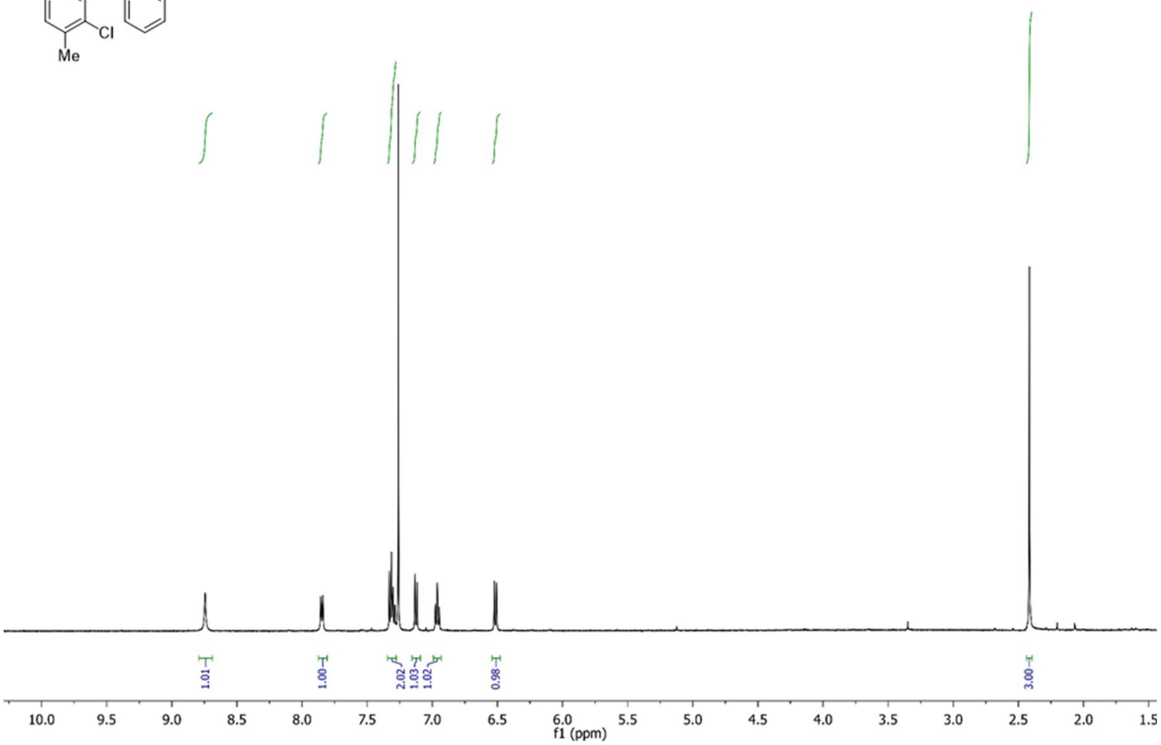
1.01
1.00
2.02
1.03
1.02
0.98
3.00
f1 (ppm)

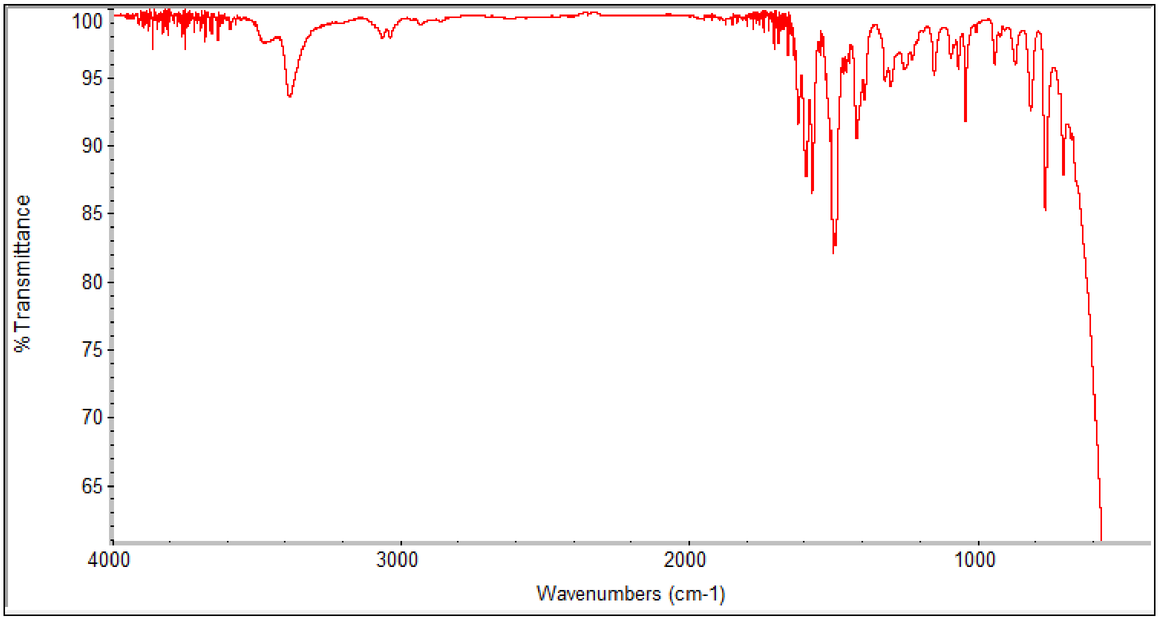
% Transmittance
Wavenumbers (cm-1)

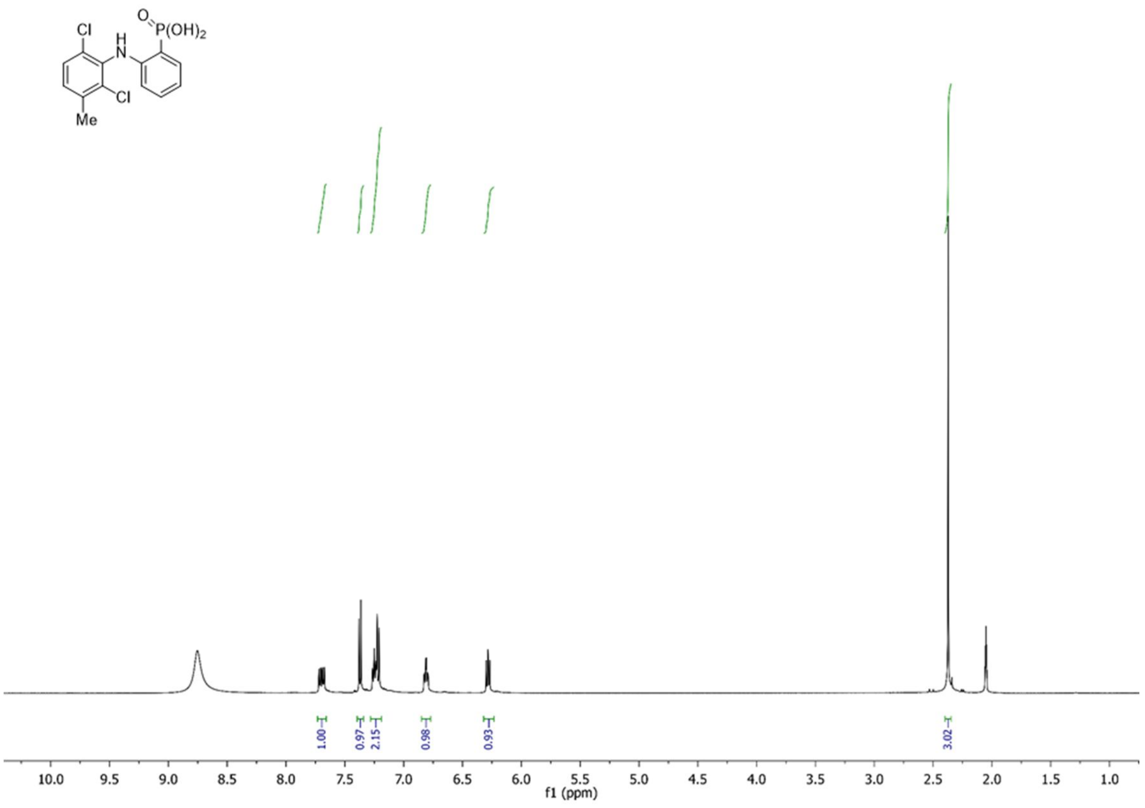
Cl
H
N
O
P(OH)2
Cl
Me
1.00
0.97
2.15
0.98
0.93
3.02
10.0 9.5 9.0 8.5 8.0 7.5 7.0 6.5 6.0 5.5 5.0 4.5 4.0 3.5 3.0 2.5 2.0 1.5 1.0
f1 (ppm)

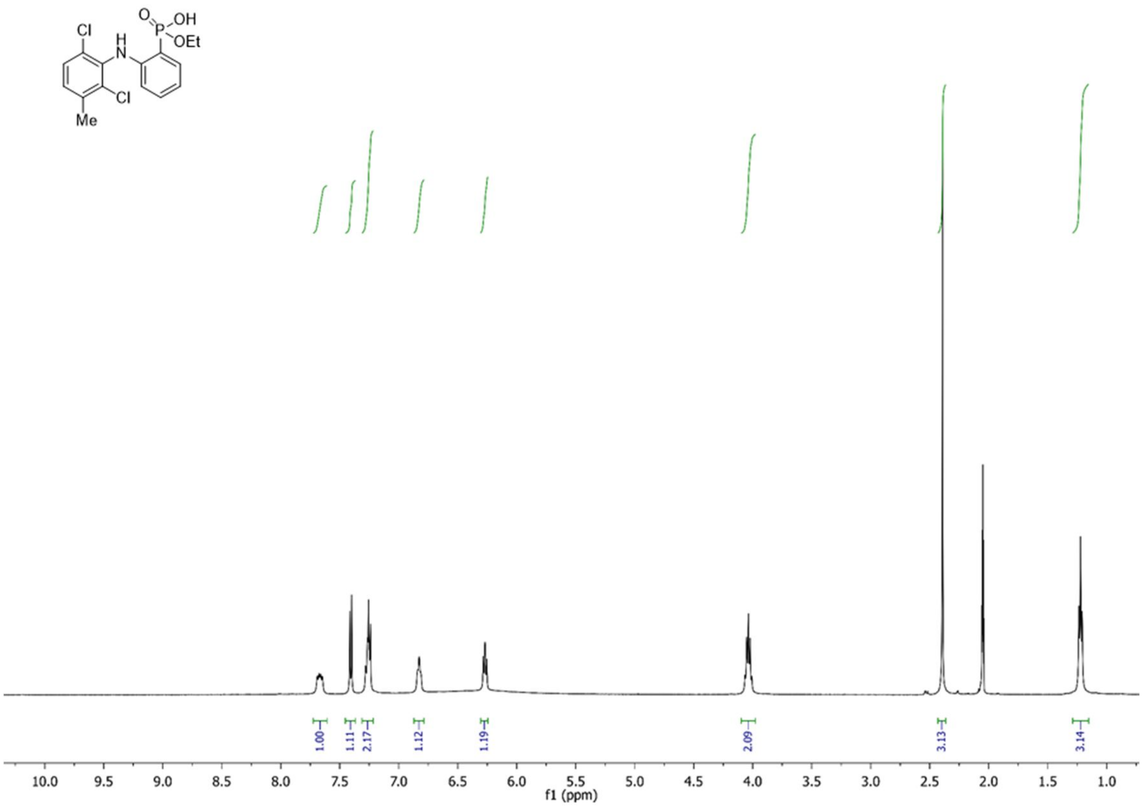
Cl
H
N
O
P
OH
OEt
Cl
Me
1.00
1.11
2.17
1.12
1.19
2.09
3.13
3.14
10.0 9.5 9.0 8.5 8.0 7.5 7.0 6.5 6.0 5.5 5.0 4.5 4.0 3.5 3.0 2.5 2.0 1.5 1.0
f1 (ppm)
